# RNPS1 stabilizes NAT10 protein to facilitate translation in cancer via tRNA ac^4^C modification

**DOI:** 10.1101/2023.02.11.528122

**Authors:** Xiaochen Wang, Kang Li, Maosheng Cheng, Hao Xu, Jie Chen, Xin Peng, Rongsong Ling, Jianwen Chen, Yuehan Wan, Lixin Ke, Caihua Zhang, Qitong Zhang, Yutong Zou, Fangfang Chen, Zhi Chen, Shuang Chen, Jingting Li, Liang Peng, Qianming Chen, Cheng Wang, Qi Liu, Demeng Chen

**Affiliations:** Center For Translational Medicine, the First Affiliated Hospital, Sun Yat-Sen University, Guangzhou, China.; State Key Laboratory of Oral Diseases, National Clinical Research Center for Oral Diseases, Research Unit of Oral Carcinogenesis and Management, Chinese Academy of Medical Sciences, West China Hospital of Stomatology, Sichuan University, Chengdu, Sichuan 610041, P. R. China.; Rice Research Institute & Guangdong Key Laboratory of New Technology in Rice Breeding, Guangdong Academy of Agricultural Sciences, Guangzhou, China.; College of Agriculture, South China Agricultural University, Guangzhou 510642, China; Institute for Advanced Study, Shenzhen University, Shenzhen, China.; Department of Oral and Maxillofacial Surgery, Hospital of Stomatology, Guanghua School of Stomatology, Sun Yat-Sen University, Guangzhou 510055, China; Guangdong Provincial Key Laboratory of Stomatology, Sun Yat-Sen University, Guangzhou 510080, China.; Senior Department of Oncology, the Fifth Medical Center of PLA General Hospital, NO.8 the east street, Fengtai District, Beijing, 100071, P.R.China; Stomatology Hospital, School of Stomatology, Cancer Center, Zhejiang University School of Medicine, and Key Laboratory of Oral Biomedical Research of Zhejiang Province, Clinical Research Center of Oral Diseases of Zhejiang Province, Hangzhou, 310006, Zhejiang, China.

**Keywords:** NAT10, RNPS1, tRNA, ac^4^C, TRMC-seq

## Abstract

NAT10 is dysregulated and plays an essential role in various types of cancers. However, the exact machenism of how NAT10 regulates cancer progression remains debatable. In this report, we show that NAT10 affects tumorigeneis mainly based on its acetylation function on tRNA. In addition, we found NAT10 regulate the ac^4^C of tRNA in cancer via interaction with RNPS1, which in turn protect NAT10 from degradation by E3 ubiquitin ligase ZSWIM6. We developed TRMC-seq method to compreshensively profile tRNA ac^4^C sites and uncovered the presence of ac^4^C in a broader range of tRNA isoacceptors than previous studies. Multi-omics analysis identified AP-1 signaling pathway as a major downstream mediator of NAT10. Mechanistically, we found NAT10 is responsible for the translation efficiency genes which contain higher ac^4^C-tRNA codon. Importantly, our genetic mouse model validated our in vitro findings of NAT10 in cancer. Our study highlights a role of NAT10 in mediating tRNA ac^4^C to regulate the translation and tumorigenesis of cancer.

## Introduction

In recent years, increasing evidence suggests that RNA epitranscriptomics and related enzymes are misregulated in tumorigenic processes and serve as ideal targets of cancer therapy (*1, 2*). To date, more than 170 different modifications have been identified in RNA molecules and the majority of them are located in the anticodon loop region of tRNA (*3, 4*). Among these modifications, N4-acetylcytidine (ac^4^C) on cytidine is a conserved acetylation on RNA across various species and has important regulatory functions in diverse biological processes (*5*). Ac^4^C can be catalyzed by N- acetyltransferase 10 (NAT10) in mammals or by its homolog Kre33 in yeast. In mammals, NAT10 has been found to catalyze the ac^4^C of tRNA-Ser, tRNA-Leu and 18S rRNA (*3, 6*). However, whether ac^4^C exist in the mammalian mRNA remains controversial. By using an ac^4^C antibody-based high-throughput sequencing approach (acRIP-seq), some studies claimed that there are abundant ac^4^C sites across the transcriptome and ac^4^C occurs in the cytidine enriched within the wobble sites of mRNA, which is crucial for mRNA stability and translation (*7-9*). However, another group argued the existence of ac^4^C on eukaryotic mRNA by using ac^4^C-seq, a chemical-based sequencing method which is able to identify ac^4^C at single-nucleotide resolution (*10*). Therefore, the distribution and biological functions of ac^4^C on mammalian RNA need in-depth investigation.

The poor outcomes of current treatment for head and neck squamous cell carcinoma (HNSCC) patients are underscored by the high mortality and morbidity rates (*11*). Therefore, more effective therapies for HNSCC are needed. Recently, we and others have shown that targeting dysregulated RNA modification enzymes, including METTL1 and METTL3, which are responsible for mRNA m^6^A and tRNA m^7^G modification, respectively, represent promising strategies for treatment of HNSCC (*12-14*). This also prompts us to pursue a more comprehensive understanding of how RNA modification can drive the tumorigenesis and progression of HNSCC.

In this study, we found that upregulated NAT10 is significantly correlated with poor prognosis of HNSCC patients using different cohorts. Knockdown (KD) of NAT10 or treatment of NAT10 small molecule inhibitor Remodelin can suppress the tumorigenic properties of HNSCC cells. In addition, we identified RNPS1 is associated with NAT10 in HNSCC. Using both antibody-and chemical-based sequencing approaches, we found NAT10 and RNPS1 are required for ac^4^C modification of tRNA, but not mRNA. In addition, we developed a tRNA reduction and misincorporation sequencing method to profile the ac^4^C on tRNA and found several novel ac^4^C sites, which play a crucial role in tRNA-based translation and tRNA turnover. Multi-omics analysis revealed that depletion of NAT10 or RNPS1 resulted in inhibition of JUN oncogene in HNSCC cells. Mechanistically, RNPS1 can stabilize NAT10 protein by preventing the ubiquitination of NAT10. Overall, this study presents a novel mechanism of how oncogenic program is tightly controlled by NAT10-mediated tRNA ac^4^C acetylation in HNSCC.

## Results

### NAT10 is elevated in HNSCC and correlates with HNSCC progression

To evaluate the NAT10 expression and investigate its correlation with HNSCC progression, we initially characterized the mRNA expression level of NAT10 in local HNSCC samples. We found that NAT10 mRNA was significantly upregulated in HNSCC compared with normal tissue (Fig. 1A). In addition, Northwestern & Northern blot assays revealed that ac^4^C levels in total RNA was elevated in HNSCC compared with normal tissue (Fig. 1B). Compared with human primary HNSCC with T1/2 stage, NAT10 mRNA expression was increased significantly in those with T3/4 stage (Fig. 1C). A dramatic upregulation of NAT10 was observed in HNSCC with lymph node metastasis as compared to those without metastasis (Fig. 1D). Of note, increased expression of NAT10 was also observed during HNSCC progression and correlated with more advanced HNSCC (Fig. 1E). Moreover, high expression of NAT10 indicates a poor overall survival and disease-free survival (Fig. 1F, G). We then examined the expression of NAT10 protein using immunohistochemistry staining (IHC). As expected, NAT10 protein expression is significantly increased in HNSCC as compared to the non-cancerous adjacent tissues (Fig. 1H, I). To further validate these findings, The Cancer Genome Atlas Head-Neck Squamous Cell Carcinoma (TCGA-HNSC) cohort was used to examine NAT10 expression and determine its prognostic value. Consistently, NAT10 expression was significantly upregulated in HNSCC samples and associated with cancer progression (Fig. S1A-D). Importantly, high expression of NAT10 mRNA indicates a poor overall survival and progression-free survival (Fig. S1E-H). Collectively, our data suggest that NAT10 is markedly upregulated and is associated with malignant progression of HNSCC, suggesting NAT10 might enhance aggressiveness of HNSCC.

**Figure 1.**
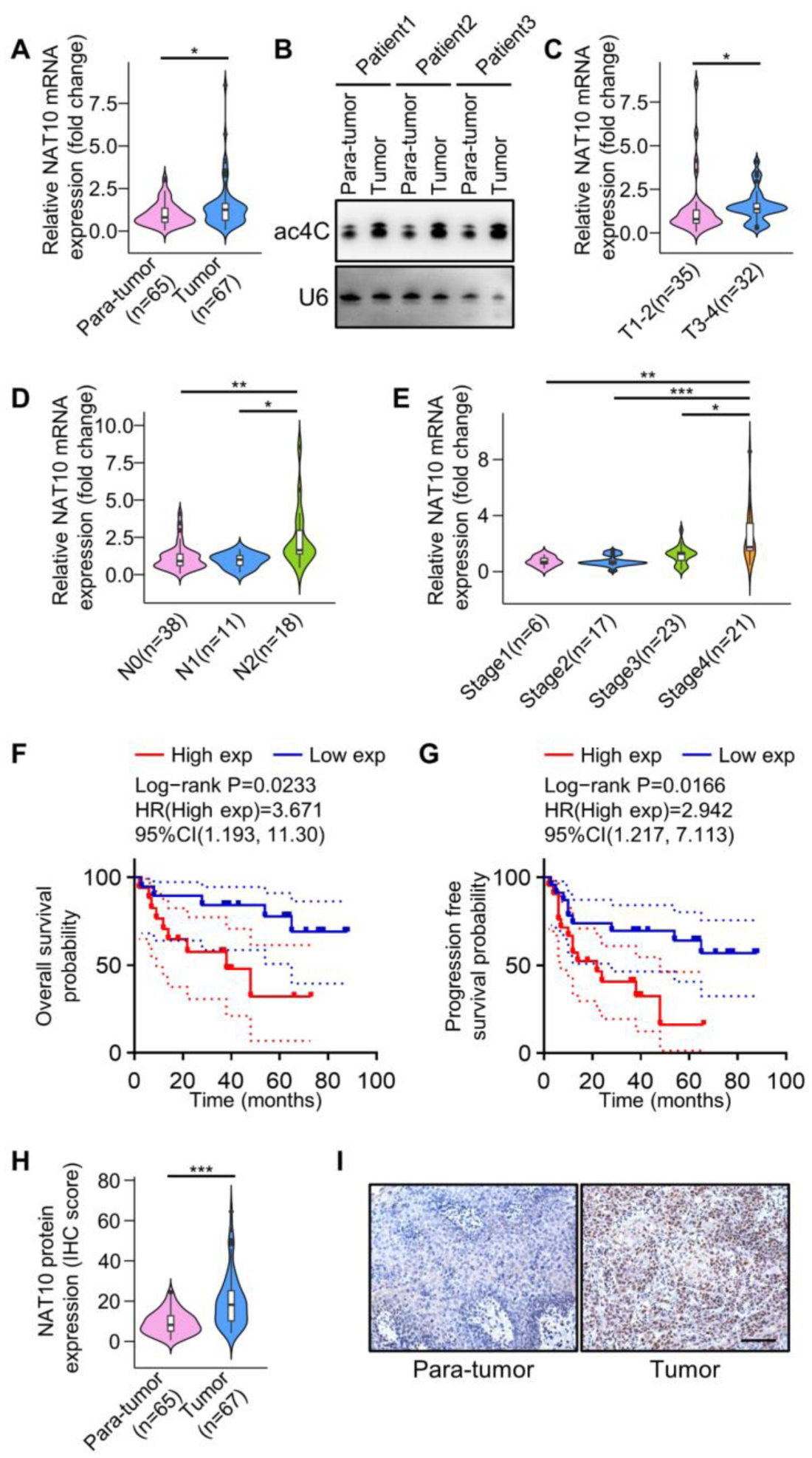
NAT10 is elevated in HNSCC and correlates with HNSCC progression. (A) Expression of NAT10 mRNA by qPCR in HNSCC samples from the Hospital of Stomatology, Sun Yat-sen University. The violin plot represents a hybrid of a box plot and a kernel density plot. In the box plot, the upper side of the box indicates the third quartile (Q3), the lower side indicates the first quartile (Q1) and the length of this box represents interquartile range (IQR). The second quartile(Q2), also called median (Md), is depicting as a horizontal line inside the box. The endpoints of the two vertical lines outside the box represent the minimum and maximum values respectively and the points outside the box represent outliers. On each side of the box plot is a kernel density estimation to show the distribution shape of the data. Wider sections of the violin plot represent a higher probability that members of the population will take on the given value; the skinnier sections represent a lower probability. Para-tumor, n = 65; Tumor, n = 67. *p < 0.05 by unpaired Student’s t test.(A) (B) The ac^4^C levels of para-tumor and tumor RNA samples in HNSCC patients by Northwestern blot (NWB) and Northern blot (NB). U6 snRNA was used a loading control. (B) Violin plot showing the correlation between NAT10 expression and HNSCC tumor size and invasion depth of patients. T1: tumor size ≤ 2 cm, infiltration depth ≤ 0.5 cm; T2: tumor size ≤ 2 cm, 0.5 < infiltration depth ≤ 1 cm, or 2 cm < tumor size ≤ 4 cm, infiltration depth ≤ 1 cm; T3: tumor size > 4 cm, or infiltration depth > 1 cm; T4: the tumor infiltrated the bone cortex, or inferior alveolar nerve, or orofacial, or masticatory muscles, etc. The violin plot parameters are as in Figure 1A. T1-2, n = 35, T3-4, n = 32. *p < 0.05 by unpaired Student’s t test. (A) (D) The correlation between NAT10 expression and HNSCC nodal metastasis status of patients. N0: no regional lymph node metastasis; N1: metastases in 1 to 3 axillary lymph nodes; N2: metastases in 4 to 9 axillary lymph nodes. N0, n = 38; N1, n = 11; N2, n = 18. *p < 0.05 and **p < 0.01 by One-way ANOVA. (B) (E) The correlation between NAT10 expression and HNSCC individual cancer stages of patients. Stage1, n = 6; Stage2, n = 17; Stage3, n = 23; Stage4, n = 21. *p < 0.05, **p < 0.01 and ***p < 0.001 by One-way ANOVA. (F-G) Kaplan-Meier survival curves revealing NAT10 expression and overall survival probability or progression free survival probability in HNSCC patients. Among them, different groups were tested by log rank. HR (high exp) represents the risk coefficient of the high expression group relative to the low expression group. If HR > 1 represents the risk factor of the gene, and if HR < 1 represents the protective factor of the gene; 95% Cl represents HR confidence interval. p = 0.0233 (F) and p = 0.0166 (G) by Log-rank test. (H-I) The immunocytochemistry (IHC) score and IHC protein expression of NAT10 in para-tumor (n = 65) and tumor (n = 67) of HNSCC samples from the Hospital of Stomatology, Sun Yat-sen University. Scale bars of (I) are 100 μm. ***p < 0.001 (H) by unpaired Student’s t test.

### NAT10 plays a vital role in HNSCC progression and metastasis

To elucidate the role of NAT10 in HNSCC, we first compared the expression of the NAT10 in a panel of HNSCC cell lines and human normal oral epithelial keratinocytes (HOK). Western blot results showed all HNSCC cell lines expressed higher level of NAT10 compared to HOK cells (Fig. 2A). Among these cell lines, SCC-9 and SCC-15 cells expressed the highest levels of NAT10 (Fig. 2A). We then constructed SCC-9 and SCC-15 NAT10 KD stable lines using two different lentiviruses (Fig. 2B-C). We used mass spectrometry to quantify the levels of ac^4^C RNAs and found NAT10 KD led to dramatic decrease of ac^4^C levels (Fig. 2D). In addition, dot blot assays revealed that KD of NAT10 significantly reduced ac^4^C levels in total RNA relative to control cells (Fig. 2E). Functionally, depletion of NAT10 resulted in inhibition of cell proliferation and migratory abilities (Fig. 2F; Fig. S2A). We also detected an increase in the number of apoptotic cells in NAT10 KD cells compared with the control cells (Fig. 2G). To examine whether NAT10 is important for the stem cell-like properties of HNSCC cells, we performed sphere-forming assay and aldehyde dehydrogenase (ALDH) activity assay. Our resulted demonstrated that depletion of NAT10 significantly decreased the tumor sphere formation frequency and the percentage of ALDH^high^ population of SCC-9 and SCC-15 cells (Fig. S2B and C). Furthermore, we performed orthotopic implantation experiments in nude mice to examine the tumorigenic and metastatic ability of NAT10 stable KD SCC-15 cells in vivo. The tumor volume in the KD group were significantly smaller than those in the control group (Fig. 2H-J). Depletion of NAT10 also resulted in the cell proliferation and metastatic capability to cervical lymph nodes (Fig. 2K).

**Figure 2.**
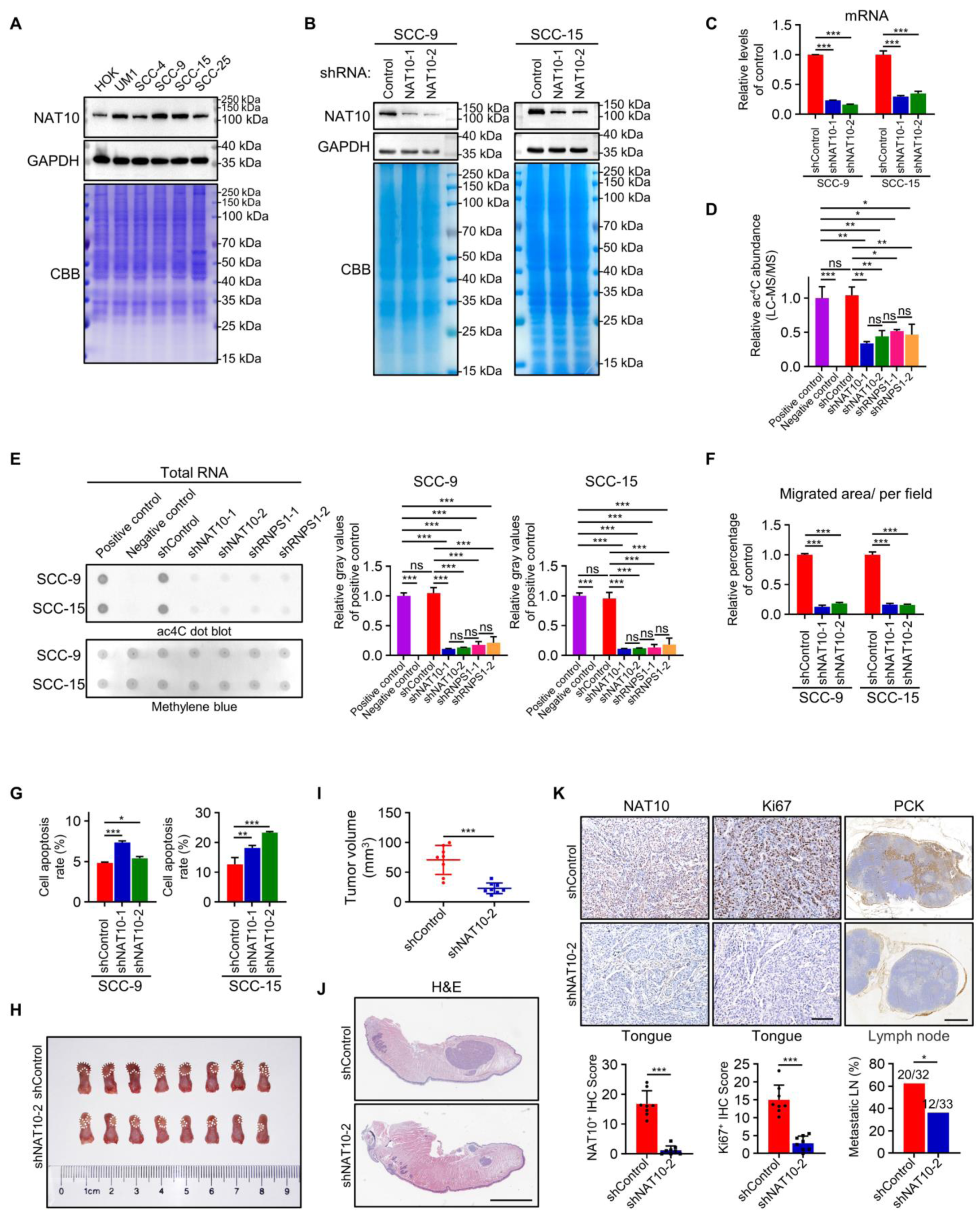
NAT10 plays a vital role in HNSCC progression and metastasis. (A) Expression of NAT10 by Western blot (WB) in different cell lines, including HOK, UM1, SCC-4, SCC-9, SCC-15 and SCC-25. GAPDH and Coomassie brilliant blue (CBB) staining were used as loading control. (B and C) WB and qPCR showed that NAT10 was knocked down by shRNA in SCC-9 and SCC-15. GAPDH and CBB staining were used as loading control. Data are represented as mean ± standard deviation (SD). ***p < 0.001 (C) by One-way ANOVA. (I) (D) The bar chart of relative ac^4^C abundance of RNA by LC-MS/MS from SCC-15 cell line. Positive control: ribosomal RNA; Negative control: poly(A)-enriched RNA; The remaining samples are total RNA from control, NAT10-KD and RNPS1-KD groups. N = 2 independent biological replicates. Data are represented as mean ± SD. *p < 0.05, **p < 0.01 and ***p < 0.001 by One-way ANOVA. (I) (E) Dot blot (left) and relative gray values (right) showed the ac^4^C level of RNA in Positive control, Negative control, Control, NAT10-KD and RNPS1-KD groups with methylene blue staining as loading control. n = 2 independent biological replicates. Data are represented as mean ± SD. ***p < 0.001 by One-way ANOVA. (F and G) Migration (F) and cell apoptosis (G) between the shControl, shNAT10-1 and shNAT10-2 in HNSCC cell lines. In the migration assay (F), Mitomycin C (MMC) was added to inhibit the effect of cell proliferation. Data are represented as mean ± SD. *p < 0.05, **p < 0.01 and ***p < 0.001 in (F) and (G) by One-way ANOVA. (A) (H) Image of xenografted SCC-15 cell line including shControl and shNAT10-2 in tongue of nude mice. The white dotted line indicates tumor boundary. n = 8 mice in each group. (I) Tumor volume(mm^3^) of orthotopic transplantation tumor between shControl and shNAT10-2. n = 8 mice in each group. Data are represented as mean ± SD. ***p < 0.001 by unpaired Student’s t test. (A) (J) H&E staining of tongue with orthotopic transplantation tumor. The scale bar is 2 mm. n = 8 mice in each group. (B) (K) The immunohistochemical (IHC) staining and IHC Score of NAT10 (left), Ki67 (middle) in orthotopic transplantation tumor and PCK (right) in lymph node. The scale bar of lymph node is 500 μm and the rest are 100 μm. n = 8 mice in each group. Data are represented as mean ± SD. ***p < 0.001 in IHC Score of NAT10 and Ki67 by unpaired Student’s t test. *p < 0.05 in Metastatic LN (%) by Chi-square test.

We also inhibited the activity of NAT10 in HNSCC cells using small molecular inhibitor, Remodelin (*15*). First, we treated SCC-9 cells with gradient concentrations of Remodelin to define its pharmacokinetic properties. We found that Remodelin inhibited proliferation of SCC-9 and SCC-15 HNSCC cell lines with mean ± SD IC_50_ of 7.166 ± 1.104 μM or 14.38 ± 1.21 μM, respectively (Fig. S2D). Treatment of Remodelin evidently decreased the levels of NAT10 protein and ac^4^C modification in total RNA (Fig. S2E-G). Furthermore, Remodelin impaired the cell proliferation, migration ability of SCC-9 and SCC-15 cells (Fig. S2H and I). Remodelin also induced apoptosis of HNSCC cells (Fig. S2J). Sphere-forming assay and ALDH assay showed that Remodelin significantly decreased the tumor sphere formation frequency and the percentage of ALDH^high^ population compared with the control groups (Fig. S2K and L). Results from orthotopic model demonstrated that Remodelin inhibited the tumor formation of SCC-15 cells in vivo (Fig. S3A and B). IHC results showed Remodelin diminished the expression of NAT10 in SCC-15 in vivo (Fig. S3C). In addition, Remodelin significantly decreased the proliferative and metastatic ability of SCC-15 (Fig. S3D and E).

Complementarily, we transfected SCC4 cells, which exhibited lowest level of NAT10 (Fig. 2A), with wild type NAT10 (NAT10-WT) or enzymatic mutant NAT10 (NAT10-MUT) (*15*). Western blot results confirmed the expression of NAT10 was drastically elevated after transfection (Fig. S3F). In addition, the ac^4^C level of total RNA in the NAT10-WT transfection group by dot blot assay was increased compared with that of the control group and NAT10-MUT group (Fig. S3G and H). Functional assays revealed that cell proliferation, migration ability and stem cell-like properties were enhanced, and apoptosis rate was inhibited in NAT10-WT group compared with the control or NAT10-MUT group (Fig. S3I-M). Collectively, our results indicated that NAT10 is essential for HNSCC tumorigenic properties.

### NAT10 interacts with RNPS1 to regulate tRNA ac^4^C modifications

To discover potential NAT10 associated proteins, we performed coimmunoprecipitation (co-IP) to pulldown endogenous NAT10 in SCC-15 cells. The product were run in a gel for a short distance then cut out and digested for a selective and sensitive high-performance liquid chromatography tandem mass spectrometry (LC-MS/MS) analysis. NAT10 was identified in the eluates by LC-MS/MS from SCC-15 cells pulldown by anti-NAT10 antibody but not IgG control, indicating the successful pull down of NAT10 from cell extracts. To identify the unique NAT10 associated proteins in HNSCC cells, proteins that appeared in the IgG and NAT10 KD control groups were then subtracted. In total, LC-MS/MS of the co-IP proteins identified 18 different proteins (Table S1). In particular, we found RNA-binding protein with serine-rich domain 1 (RNPS1) were enriched among these 18 candidates (Table S1). To test that, we pulled them down from SCC-15 cell extracts using anti-NAT10 or anti-RNPS1 antibodies. Indeed, the pulldown in each IP experiment contained the other protein (Fig. 3A). We also perform Coomassie brilliant blue to confirmed the presence of RNPS1 band in the gel (Fig. S4A). Unexpectedly, we did not detect THUMPD1 by LC-MS/MS analysis (Table S1). We then performed the co-IP assay of THUMPD1 and NAT10 using different cells. Although we can clearly detected interaction between THUMPD1 and NAT10 in previous published cell line HCT116 (Fig. S4B) (*16*), we did not detect positive co-IP event of THUMPD1 and NAT10 in SCC-9 and SCC-15 cells (Fig. S4C). To map the domains of RNPS1 that are responsible for its interaction with NAT10. We constructed multiple plasmids to express different domains of RNPS1 with HA (hemagglutinin) tag, including full-length RNPS1 (WT), RNPS1 N-terminal region (NT), a serine-rich domain (S domain) in the N-terminal region (NT+S), RNPS1 without an arginine/serine/proline-rich domain (RS/P domain) in the C-terminal region (ΔRS/P) and RNPS1 without S domain (ΔS) in SCC4 cells (Fig. 3B). We found RNPS1-NT+S and RNPS1-ΔRS/P, interacted with NAT10 (Fig. 3C and D), suggesting that S domain of RNPS1 was responsible for the interactions with NAT10. Furthermore, double immunofluorescence staining for NAT10 and RNPS1 clearly demonstrated that NAT10 and RNPS1 co-localized in the nucleoli (Fig. 3E) of SCC-15 cells. While NAT10 is predominantly located in the nucleoli, we noticed a broader expression pattern of RNPS1 (Fig. 3E). In addition, a significant positive correlation between NAT10 and RNPS1 protein expression was identified by IHC staining in HNSCC patients (Fig. 3F). Using TCGA database, we found mRNA expression of NAT10 and RNPS1 positively correlated in HNSCC (Fig. S4D). we also found expression of RNPS1 was significantly higher in HNSCC compared with normal tissues and correlated with advanced tumor grade, advanced tumor stages and nodal metastasis status (Fig. S4E-H). Intriguingly, NAT10 knock down reduced the RNPS1 protein levels and *vice versa* in both SCC-9 and SCC-15 cell lines (Fig. 3G). Notably, depletion of RNPS1 reduced levels of ac^4^C in total RNA (Fig. 2D and E).

**Figure 3.**
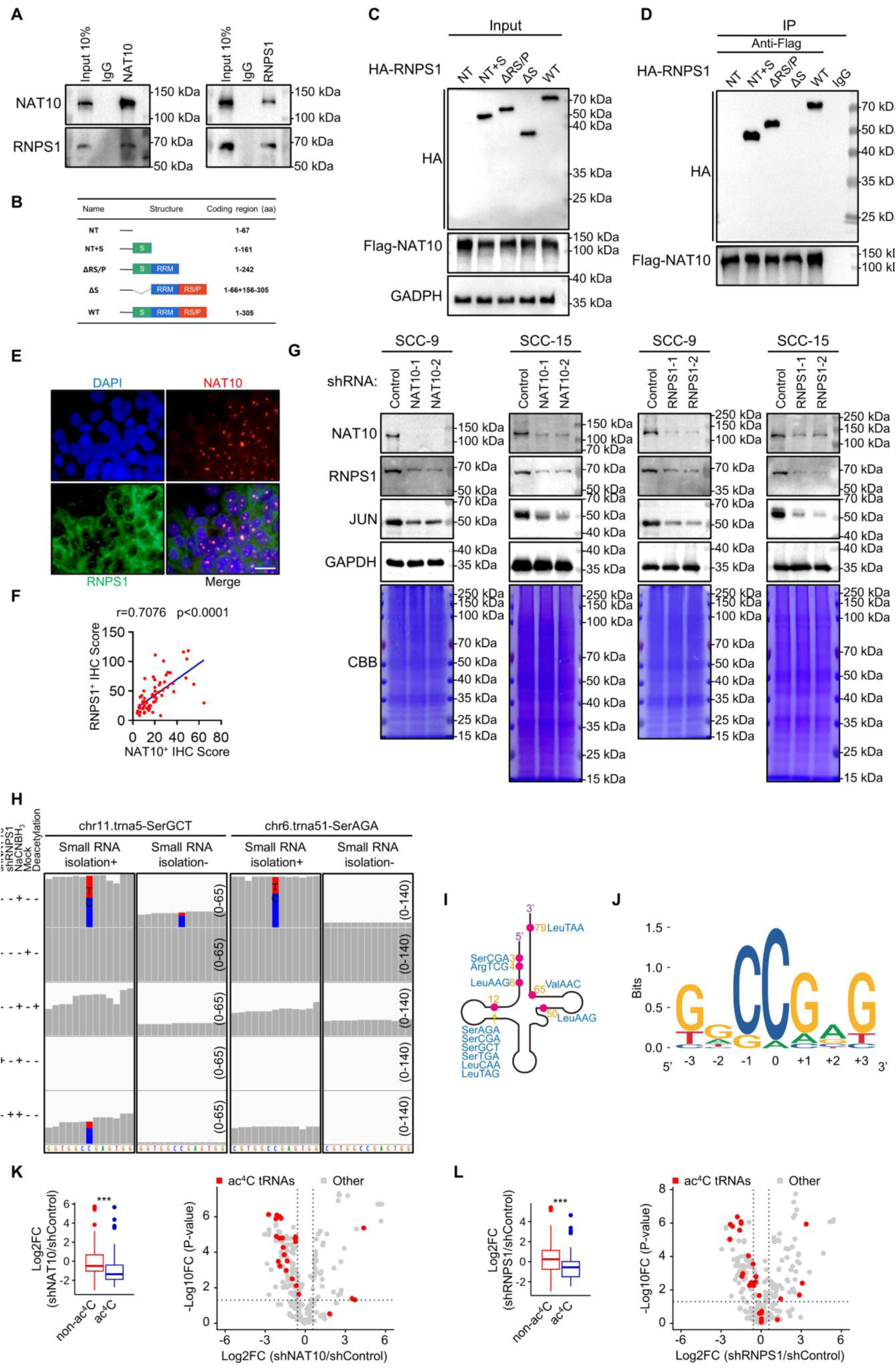
NAT10 interacts with RNPS1 to regulate tRNA ac^4^C modifications. (A) Association of endogenous NAT10 with RNPS1 in SCC-15 by co-IP with anti-NAT10 antibody or anti-RNPS1 antibody. Anti-IgG antibody was used as a negative control. (B) Strategy of RNPS1 variant proteins for mapping interaction domains with NAT10. (C) The expression of RNPS1 variant proteins after pcDNA3.1-HA-RNPS1 variant proteins and pICE-FLAG-NAT10-siR-WT were co-transfected into SCC-4 cells. (D) SCC-4 cells were co-transfected with NAT10-Flag and HA-RNPS1 variant proteins as indicated. The NAT10-RNPS1 complex was immunoprecipitated with anti-Flag antibody and detected with anti-HA antibody. Anti-IgG antibody was used as a negative control. (E) Immunofluorescence colocalization staining of NAT10 (red) and RNPS1 (green) expression in SCC-15 cells. The nuclear is counterstained with DAPI (blue). (F) The protein expression of NAT10 and RNPS1 was correlated in HNSCC patient tissues from the Hospital of Stomatology, Sun Yat-sen University. r = 0.7076, p < 0.0001 by Spearman correlation analysis. (G) The expression of NAT10, RNPS1, JUN and GAPDH in control group and KD groups of HNSCC cell lines. GAPDH and CBB staining were used as loading control. (H) Misincorporation sites of tRNA^Ser(GCT)^ and tRNA^Ser(AGA)^ in small RNA and total RNA from SCC-15 cells were detected by TRMC-seq or total RNA reduction and misincorporation sequencing. blue letters (C) and bars, cytidine; red letters (T) and bars, thymidine; green letters(A), adenosine; orange letters (G), guanosine. Small RNA isolation+, detected by TRMC-seq; Small RNA isolation-, detected by total RNA reduction and misincorporation sequencing. The number on the right indicates the number of nucleotides in the ordinate. (I) A total of 7 different ac^4^C sites were found in 10 tRNA isoacceptors by TRMC-seq. (J) Sequence motif surrounding the ac^4^C sites identified by TRMC-Seq in HNSCC cells. (K and L) The total (left) and individual (right) expression levels of non-ac^4^C tRNAs and ac^4^C tRNAs in the NAT10 KD/Control group (K) and RNPS1 KD/Control group (L). In the box diagram (left), the upper side of the box is the third quartile (Q3), the lower side of the box is the first quartile (Q1) and the length of this box represents interquartile range (IQR). The second quartile(Q2), also called median (Md), is depicting as a horizontal line inside the box. The endpoints of the two vertical lines outside the box represent the minimum and maximum values respectively and the points outside the box represent outliers. For Scatter plot (right), x-axis indicates the changes of tRNA expression levels, y-axis indicates p-value. ***p < 0.001 by Mann-Whitney test.

NAT10 has known to participate in acetylation of critical oncogene proteins(*17, 18*). Hence, we perform acetylated-lysine (Ac-K-100) assay to investigate whether NAT10 is involved in protein acetylation of HNSCC cells. However, our Ac-K-100 results showed no discernible difference between control and KD of NAT10 or RNPS1 HNSCC cells (Fig. S4I and J).

Conservatively, NAT10 can regulate ac^4^C of different type of RNAs. However, it is still debatable whether ac^4^C modification is present in the mRNA of eukaryotic cells (*10, 19*). To tackle that, we first applied antibody-based acRIPseq approach to the total RNAs of SCC-15 cells (*7*). However, only 1 consensus peak was identified after filtering for replication, indicating ac^4^C does not occur on mRNA. We then modified chemical-based ac^4^C-seq method to examine mRNA ac^4^C in control, mock, deacetylation, shNAT10 and shRNPS1 HNSCC groups (*10*) (Fig. S4K). Our sequencing data identified ac^4^C modification in 18S rRNA helix 34 and helix 45, two well-known sites that have been reported previously (*10*) (Fig. S4L; Fig. 2D and E), supporting the legitimacy of our approach. However, analysis of the sequencing results revealed only 2 C>T misincorporations in the mRNA, but they were not eliminated after the KD of NAT10, suggesting these were not *bone fide* ac^4^C modification. To further validate these findings, we isolated poly(A)-enriched RNA from HNSCC cells and performed mass spectrometry and ac^4^C dot-blot assays. We found no detectable ac^4^C signaling (Fig. 2D and E). Together, our results indicated ac^4^C modification is not present in the mRNA of HNSCC cells.

Since NAT10 is well-known for its catalyzing the formation of ac^4^C on tRNA^Ser^ and tRNA^Leu^ (*10, 20, 21*), we then analyzed ac^4^C on tRNA in HNSCC cells. Surprisingly, we found 13 ac^4^C sites in 9 tRNA isoacceptors, including ac^4^C_12_ on tRNA^Ser(GCT)^; ac^4^C79 on tRNA^Leu(TAA)^; ac^4^C12 on tRNA^Leu(TAG)^; ac^4^C12 on tRNA^Ser(TGA)^; ac^4^C12 on tRNA^Ser(AGA)^; ac^4^C12 on tRNA^Ser(CGA)^; ac^4^C_4_ on tRNA^Arg(TCG)^; ac^4^C6 on tRNA^Leu(AAG)^ and ac^4^C12 on tRNA^Leu(CAA)^ in our ac^4^C-seq data (Fig. 3H; Fig. S5A), suggesting a broader effect of NAT10 on tRNA ac^4^C. Interestingly, KD of RNPS1 also led to decreased of misincorporation proportion in the same tRNA ac^4^C sites (Fig. 3H; Fig. S5A). Bolstered by these finding, we then isolated small RNAs (< 200 nt) and performed mass spectrometry and ac^4^C dot blot assays. We found levels of ac^4^C on small RNAs were reduced after NAT10 or RNPS1 KD (Fig. S5B-E). In order to find the specific sites of acetylation modification on tRNA, we developed a method more suitable for tRNA reduction and misincorporation sequencing (TRMC-seq) (Fig. S4K). Based on this approach, we found ac^4^C on 10 tRNA isoacceptors, including 16 tRNA isodecoders (17 sites) using TRMC-Seq (Fig. 3H and I; Fig. S5A; Fig. S6A). Besides the sites described above, TRMC-seq results revealed several novel sites, including ac^4^C_3_ on tRNA^Ser(CGA)^; ac^4^C_50_ on tRNA^Leu(AAG)^; ac^4^C_65_ on tRNA^Val(AAC)^ (Fig. 3H; Fig. S5A), supporting higher sensitivity of TRMC-seq in detecting ac^4^C on tRNAs. Motif analysis revealed that ac^4^C sites were strongly enriched for CCG trimers (Fig. 3J). To probe the function of ac^4^C modification in tRNA regulation, we compared the expression levels of non-ac^4^C-modified tRNA and ac^4^C-modified tRNA. We found decrease of ac^4^C-modified tRNA levels after KD of NAT10 or RNPS1 compared to non-ac^4^C-modified tRNA (Fig. 3K and L). Northern blot results confirmed that KD of NAT10 or RNPS1 led to suppression of ac^4^C-modified tRNA detected by TRMC-seq, including tRNA^Arg(TCG)^, tRNA^Ser(TGA)^, tRNA^Leu(CAA)^ and tRNA^Val(AAC)^ (Fig. S6B and C). It’s worth noting that KD of NAT10 or RNPS1 led to a greater inhibition of ac^4^C in tRNA compared with rRNA (Fig. S6D and E).

To explore the global translation dynamics by NAT10 or RNPS1, we performed polysome profiling assay to isolate the polysome-associated translated mRNAs from untranslated ones based on sucrose-gradient separation. Both SCC-9 and SCC-15 cells after NAT10 or RNPS1 KD showed clear reduction in polysome peaks compared to the control cells (Fig. 4A). To test whether decreased of polysome peaks reflected a less active translating mRNA, we measured protein synthesis using SUnSET, a puromycin/antibody-based method. Our results revealed inhibition of translation at the protein level after NAT10 or RNPS1 KD in SCC-9 and SCC-15 cells (Fig. 4B). In addition, results from ribosome profiling assays showed that KD of NAT10 or RNPS1 increased the codon-dependent ribosome pause of ac^4^C tRNA, supporting that NAT10 or RNPS1-mediated ac^4^C tRNA modification was critical for efficient codon recognition during the ribosomal transition of ac^4^C tRNA decoding codons (Fig. 4C and D). In addition, codon frequency analysis demonstrated that the mRNAs with increased translation efficiency (TE) possess significantly lower frequencies of ac^4^C-modified tRNA decoding codons (Fig. 4E and F). To study whether ac^4^C tRNAs play a crucial role in regulating the translation of HNSCC cells, we overexpressed SerGCT, SerAGA, LeuCAA in NAT10 KD cells (Fig. S6F and G). Although the ac^4^C levels of tRNA were not altered (Fig. S6H and I), overexpression of SerGCT, SerAGA, LeuCAA led to partially rescue of the translation level after NAT10 KD (Fig. 4G; Fig. S6F and G).

**Figure 4.**
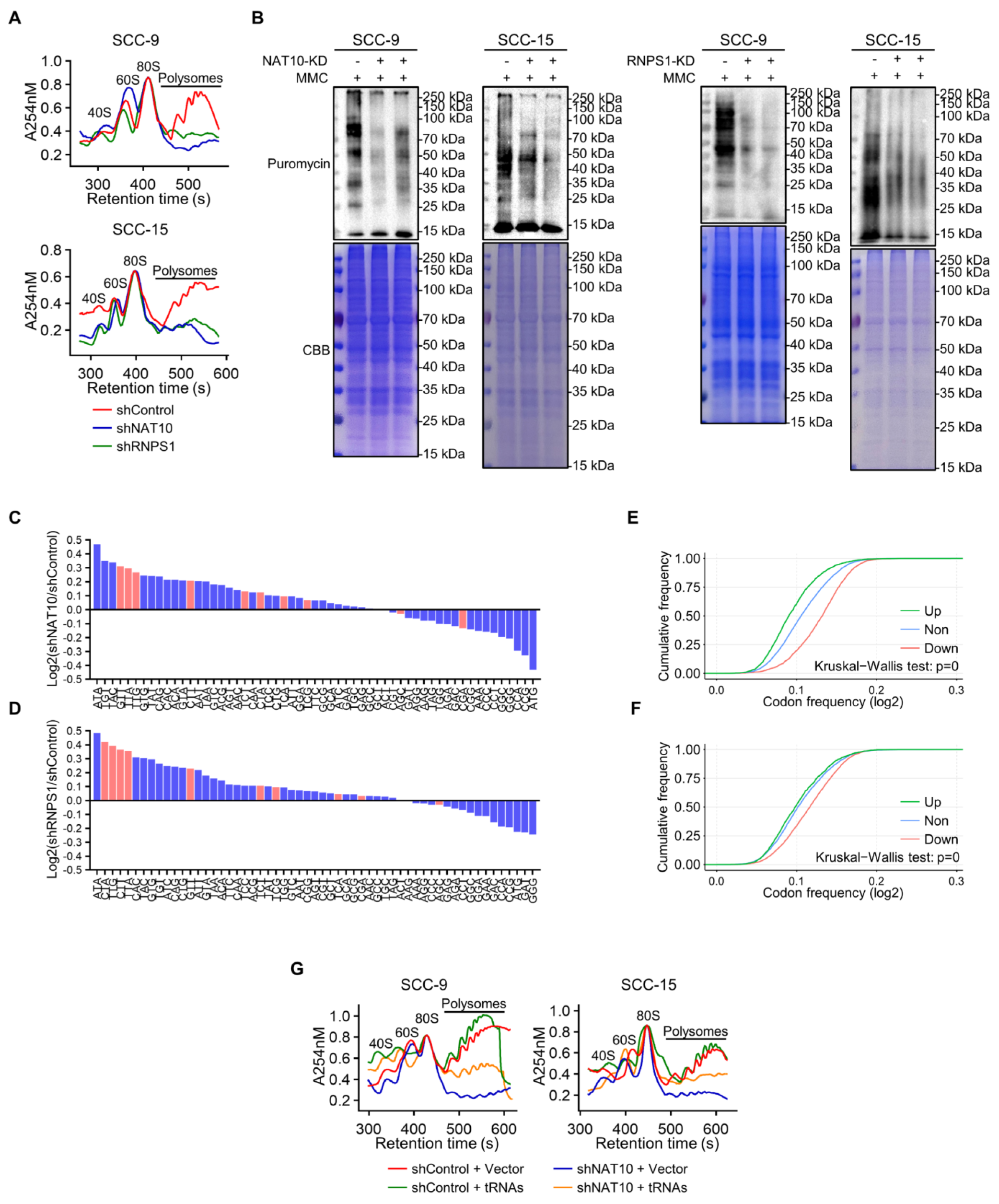
NAT10 or RNPS1 affects translation efficiency. (A) Polysome profiling of the Control, NAT10 KD and RNPS1 KD groups in SCC-9 (up) and SCC-15 (down) cell lines. MMC was added to inhibit the effect of cell proliferation. (B) Puromycin assay revealed reduced protein production in NAT10 KD and RNPS1 KD cells. MMC was added to inhibit the effect of cell proliferation and CBB staining was used as loading control. (C-D) Codon occupancy bias of NAT10 KD (C) and RNPS1 KD (D) groups in SCC-15. The ac^4^C tRNA corresponding to the codons were depicted by the red bars. (E and F) Codon frequency in the CDS region of the genes with increased TE (up, green), decreased TE (down, red) and other genes (non, blue) of NAT10 KD cells (E) and RNPS1 KD cells (F). p = 0 by Kruskal-Wallis test. (I) (G) Polysome profiling of the Control + Vector, NAT10-KD + Vector, Control + tRNAs- OE and NAT10-KD + tRNAs-OE groups in SCC-9 (left) and SCC-15 (right) cell lines. The overexpressed tRNAs include SerGCT, SerAGA and LeuCAA. MMC was added to inhibit the effect of cell proliferation.

### Integrated analysis of multiple ‘omics’ reveal NAT10 and RNPS1 regulate JUN

To fully understand how multifaceted aspects of translation are controlled by NAT10 in HNSCC, we determined the translatomic and proteomic profiles of the NAT10 KD and control SCC-15 cells. We first used ribosome nascent-chain complex-bound mRNA sequencing (RNC-seq) to compared the translating mRNA profile between control and NAT10 or RNPS1 KD samples. We found 4271 differentially expressed translational active mRNA after NAT10 KD, including 3206 down-regulated translating mRNA and 1065 up-regulated translating mRNA (Fig. 5A; Table S2). Canonical pathway analysis on the down-regulated translating genes using ingenuity pathway analysis (IPA) revealed significant enrichment in pathways involved in molecular mechanisms of cancer, Rac signaling, PTEN signaling, actin cytoskeleton signaling and HGF signaling (Table S4). Meanwhile, we found 1878 significantly different translating mRNA after RNPS1 depletion (Fig. 5B; Table S3). Among these, 1481 translating mRNA were down regulated and 971 of them (65.6%) overlapped with down-regulated translating mRNAs with NAT10 KD ones (Table S2 and S3). IPA analysis results demonstrated that RNPS1 decreased the translation of genes participating in a myriad of signaling pathways (Table S4). Interestingly, 141 out of 155 (90.9%) pathways overlapped with those after NAT10 KD (Fig. 5C). To determine the proteomic profiles of HNSCC cells after NAT10 or RNPS1 KD, we conducted isobaric tags for relative and absolute quantitation (iTRAQ) assay. In 6190 total proteins identified in our iTRAQ datasets, we detected 692 or 621 down-expressed proteins after NAT10 KD or RNPS1 KD, respectively (Table S5 and S6). IPA analysis demonstrated that the ablation of NAT10 in HNSCC cells down-expressed the proteins involved in cholesterol biosynthesis, acute phase response signaling and IL-8 signaling (Table S7). Importantly, 81 out of 91 (89%) pathways that down-regulating genes involved in after NAT10 ablation overlapped with pathways enriched in RNPS1 KD group (Fig. 5C and Table S7). We then calculated the codon usage score of ac^4^C tRNA in genes and found genes that were downregulated in both NAT10 and RNPS1 KD iTRAQ sample showed biased towards tRNA codon that were ac^4^C modified (Fig. 5E and Fig. S7A).

**Figure 5.**
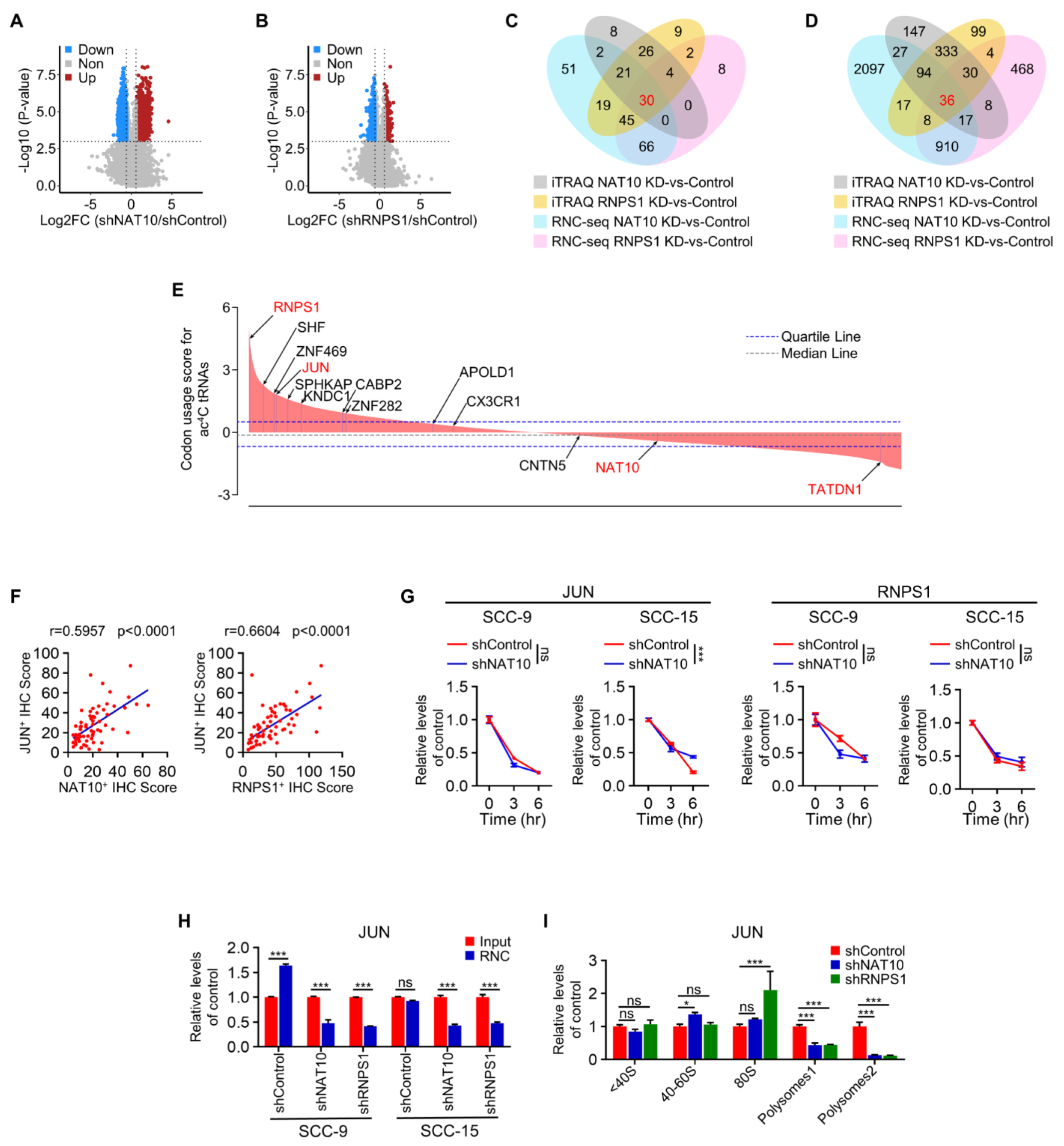
Integrated analysis of multiple ‘omics’ reveal NAT10 and RNPS1 regulate JUN. (A-B) Volcano plots of translation ratio (TR) in NAT10 KD cells (A) and RNPS1 KD cells (B). Red dots represent the genes with up-regulated TE. Blue dots represent the genes with down-regulated TE. p values by unpaired Student’s t test. (C-D) Venn diagram of IPA pathway analysis (C) and down regulated genes (D) for RNC-seq in NAT10 KD and RNPS1 KD cells, and iTRAQ in in NAT10 KD and RNPS1 KD cells. grey, RNC-seq (NAT10 KD-vs-Control); yellow, RNC-seq (RNPS1 KD-vs-Control); blue, iTRAQ (NAT10 KD-vs-Control); red, iTRAQ (RNPS1 KD-vs-Control). (A) (E) The bar graph showed the usage score of codons corresponding to 10 ac^4^C tRNAs in each gene. The genes in red indicate that they had been verified by RNC-qPCR assay or Polysome-qPCR assay. Blue dotted line represents the quartile; Grey dotted line represents the median. (B) (F) The protein expression of NAT10 vs JUN (left) and RNPS1 vs JUN (right) were correlated in HNSCC patient tissues from the Hospital of Stomatology, Sun Yat-sen University. NAT10 vs JUN, r = 0.5957, p < 0.0001 by Spearman correlation analysis. RNPS1 vs JUN, r = 0.6604, p < 0.0001 by Spearman correlation analysis. (C) (G) Actinomycin D assay showed that the half-life of JUN (left) and RNPS1 (right) mRNA did not change after KD of NAT10 in SCC-9 and SCC-15 cells. Data are represented as mean ± SD. ***p < 0.001 by Two-way ANOVA. (D) (H) The level of JUN mRNA at the ribosome combined stages and input stages after KD of NAT10 or RNPS1 in SCC-9 and SCC-15 cells. Data are represented as mean ± SD. ***p < 0.001 by Two-way ANOVA. (I) The level of JUN mRNA at different stages of ribosome or polysome binding and stages of ribosome unbinding after KD of NAT10 or RNPS1 in SCC-15 cells. Data are represented as mean ± SD. *p < 0.05 and ***p < 0.001 by Two-way ANOVA.

To further expand our finding on the involvement of NAT10 or RNPS1 in HNSCC, we integrated these multiple ‘omics’ analysis. By overlapping the ingenuity canonical pathways associated with depleted NAT10 and RNPS1 in the mRNA and protein levels, we showed 30 pathways consistently implicated in HNSCC, including IL-2, IL-3, IL- 4, IL-6, IL-8 signaling, PTEN signaling, regulation of the epithelial mesenchymal transition by growth factors pathway, *et al* (Fig. 5C and Table S4, S7). Overall, we identified 36 genes that were downregulated in RNCseq and iTRAQ datasets after NAT10 or RNPS1 KD (Fig. 5D; Table S2, S3, S5 and S6). Interestingly, JUN, a well-known oncogene that is frequently upregulated in HNSCC, was found among these 36 genes. We then wondered whether JUN can serve as a downstream target of NAT10 or RNPS1 to promote tumorigenesis of HNSCC cells. First of all, we found the protein level of JUN correlated positively with NAT10 or RNPS1 in HNSCC patient samples (Fig. 5F). Secondly, our Western blot results showed that JUN expression was inhibited after KD of NAT10 or RNPS1 in SCC-9 and SCC-15 cells (Fig. 3G). However, KD of NAT10 or RNPS1 affected neither the mRNA expression level nor the mRNA stability of RNPS1 and JUN (Fig. 5G and Fig. S7B). Instead, both RNC-qPCR and polysome-qPCR results demonstrated that KD of NAT10 or RNPS1 suppressed the translating activities of JUN mRNA (Fig. 5H and I). To test whether the translation of JUN is regulated via ac^4^C-sensitive codons, we cloned a wild type JUN transcript and two mutated JUN transcripts, where part of the ac^4^C sensitive codons were replaced with non-ac^4^C modified synonymous mutation, into vectors carrying luciferase reporter (Fig.S7C). We then transfected SCC cells using these clones and performed luciferase assay. We found that translation ratio of wild type JUN was drastically repressed after NAT10 knockdown. Compared to the wild type, both mutated transcripts reduced JUN translation ratios to a much lesser extent (Fig. S7D). These evidence indicated that the translation of JUN heavily relied on the ac^4^C sensitive codons. Furthermore, we over-expressed JUN in NAT10-KD SCC-9 and SCC-15. Although this did not reverse the effects of loss of tRNA ac^4^C and inhibition of translation (Fig. S7E-J), it could partially rescue the effects of cell proliferation, migration, apoptosis, and stem-cell like properties of HNSCC cells (Fig. S8A-E). Finally, overexpression of JUN was able to rescue the effect caused by Remodelin in HNSCC cells (Fig. S8F-I and Fig. S9A-E).

Since RNPS1 possess a high codon usage score of ac^4^C tRNA (Fig. 5E), we also tested whether loss of NAT10 can affect its translation. Our results showed that KD of NAT10 led to repressed translation of RNPS1, but not TATDN1, a negative control (Fig. S9F). However, depletion of RNPS1 showed no effect on the translation of NAT10 or TATDN1 (Fig. S9F).

### RNPS1 ensures the stability of NAT10 by inhibiting the ubiquitination of NAT10

RNPS1 is known for its role in modulating mRNA metabolism (*22*). To test whether RNPS1 is directly involved in ac^4^C modification, we overexpressed RNPS1 in the SCC-4 cells. We found that the levels of ac^4^C on RNA remained unaltered after overexpression of RNPS1 alone or with NAT10-MUT (Fig. S3G and H). In addition, overexpression of RNPS1 did not affect the global translation level of SCC-4 cells (Fig. S9 G and H). Hence, RNPS1 may regulate RNA acetylation in a NAT10-dependent manner. We noticed that RNPS1 KD led to decreased level of NAT10. We then examined our RNC-seq data and found the mRNA level and translation ratio of NAT10 was not affected after RNPS1 KD (Fig. S9F), indicating destabilization of NAT10 protein after RNPS1 KD. To explore how NAT10 is degraded after RNPS1 KD, we first treated cells with RNPS1 siRNA. We then determined the half-life of NAT10 protein (Fig. S10A). Subsequently, we used either the lysosomal inhibitor chloroquine (CQ) or proteasome inhibitor MG132 to dissect the pathway that is responsible for NAT10 degradation. Our data demonstrated that NAT10 degradation is mediated by proteasome (Fig. S10B). Since the ubiquitin–proteasome system (UPS) is responsible for degrading most proteins, we wondered which E3 ubiquitin ligase was recruited to NAT10 protein. At this point, we looked into our LC-MS/MS data to search for the E3 ubiquitin ligase candidates. We found ZSWIM6, an E3 ubiquitin ligase (*23, 24*), was associated with NAT10. We confirmed this interaction by co-IP experiments between NAT10 and ZSWIM6 both endogenously and exogenously (Fig. S10C-F) after RNPS1 KD. Furthermore, our data showed RNPS1 KD significantly increase level of ubiquitination of NAT10 (Fig. S10G and H), which can be blocked by knockdown of ZSWIM6 (Fig.7G; Fig. S10I). Overall, our data suggested that RNPS1 can stabilize NAT10 by protecting its associated with E3 ubiquitin ligase ZSWIM6.

### Nat10 drives chemically-induced HNSCC tumorigenesis

To examine the role of Nat10 in spontaneous HNSCC, we first generated conditional knockout *Nat10^flox/flox^* and knockin *Nat10^knockin^* mutant mice (Fig. 6A and Fig. 7A). *Nat10^flox/flox^* were crossed with *K14CreER* mice to generate *K14CreER*; *Nat10^flox/flox^* (Nat10^cKO^) and *K14CreER*; *Nat10^wt/wt^* (Nat10^cKO-Ctrl^) mice. 6-8 weeks old Nat10^cKO-Ctrl^ and Nat10^cKO^ mice were first injected with tamoxifen for five consecutive days, then treated with 4NQO for 16 weeks and normal drinking water for additional 10 weeks to induce HNSCC formation. Mice were then euthanized for collection of tongues and cervical lymph nodes. We observed much less morphological changes in the Nat10^cKO^ tongue tissues and found the number and area of oral lesions in the Nat10^cKO^ mice were significantly lower compared with the NAT10^cKO-Ctrl^ group (Fig. 6B and C). Histological analysis of all lesions obtained from NAT10^cKO^ and NAT10^cKO-Ctrl^ groups revealed that loss of Nat10 nearly abolished the formation of grade 3 HNSCC (Fig. 6D). Furthermore, ablation of Nat10 led to decreased proliferation of hyperplastic regions in the tongue and fewer metastatic lymph nodes (Fig. 6E and F). IHC staining results showed that NAT10 was not present in the mucosa of Nat10^cKO^ tongues (Fig. 6G). Moreover, levels of Rnps1 and Jun in tumor cells greatly declined after Nat10 knockout (Fig. 6H and I), supporting our finding using HNSCC cell lines. To test whether the translational activities of *Rnps1* and *Jun* genes were affected by NAT10 in vivo, we took advantage of the RiboTag mice, which can express hemagglutinin (HA) epitope tagging ribosomal protein L22 (RPL22) in a Cre-mediated manner(*25*). We crossed *Ribotag* (RiboTag) mice with Nat10^cKO^ mice to obtain *K14CreER*; *Ribotag*; *Nat10^flox/flox^* mice, which allowed us to capture translated mRNA transcripts after Nat10 knockout. We then pulled down the RPL22-HA-bounded mRNA using HA antibody and performed qRT-PCR using primers designed for Rnps1 and Jun transcripts. As shown in Fig. 6J and 6K, we detected decreased levels of translating Rnps1 and Jun after Nat10 ablation. In addition, we also found that these ten ac^4^C tRNAs were also degraded in HNSCC of *K14CreER*; Nat*10^flox/flox^* mice (Fig. 6L). In the meantime, we induced HNSCC in wild type mice and tested the effects of Remodelin on spontaneous HNSCC. In line with the results from Nat10 conditional knockout mice, treatment of Remodelin inhibited the HNSCC tumorigenesis and grades of malignancy (Fig. 6M-O). The proliferative and metastatic ability of tumor cells were also repressed by Remodelin (Fig. S11A and B). IHC results showed that levels of Nat10, Rnps1 and Jun were diminished in HNSCC after Remodelin treatment (Fig. S11C-E).

**Figure 6.**
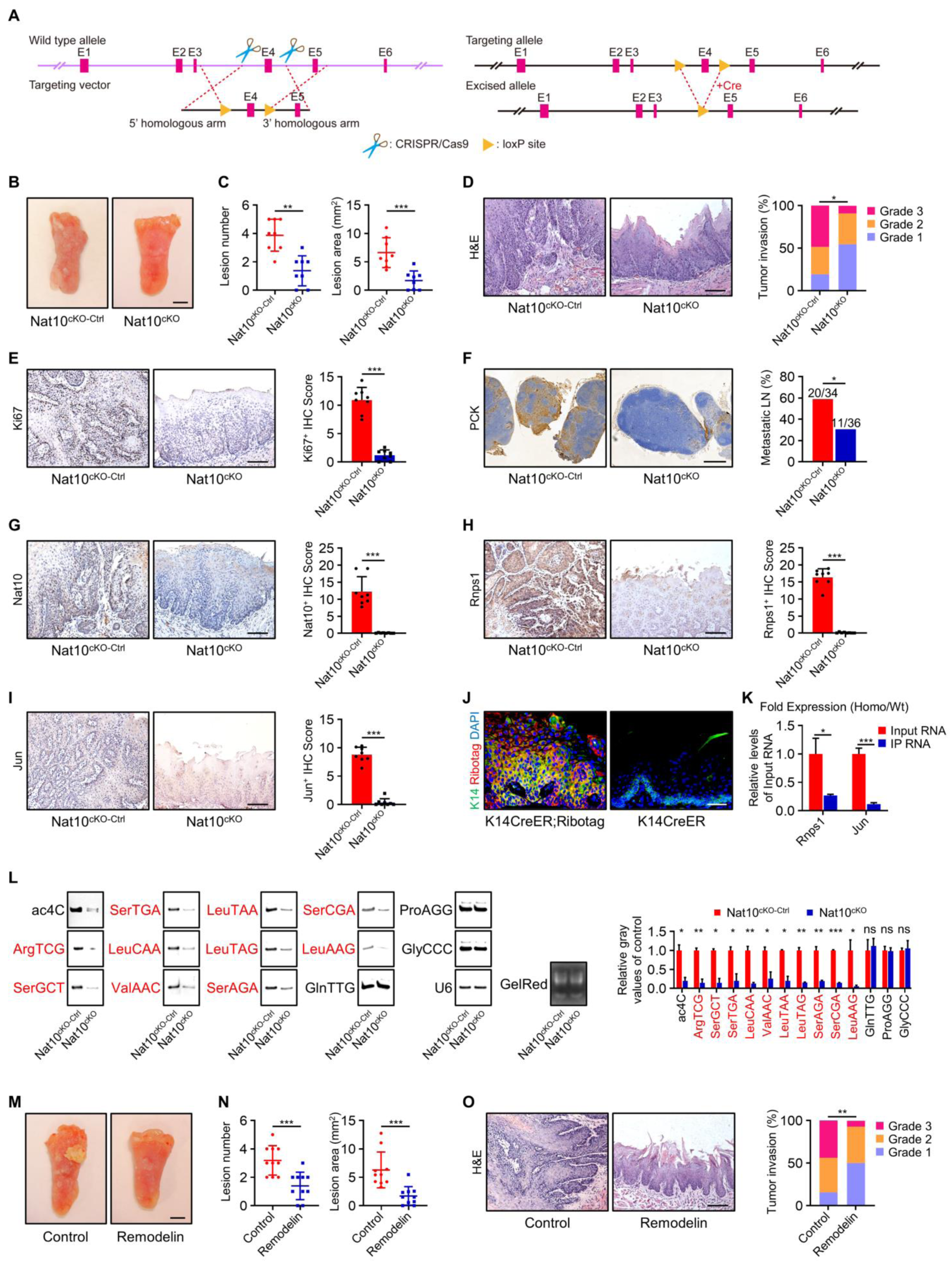
Nat10 drives chemically-induced HNSCC tumorigenesis. (A) Targeting strategy of Nat10^cKO^ mouse. This design is to conditionally knockout the exon 4 by Cre-loxP system. (B) Representative image of tongue visible lesions. n = 8 mice in each group. Scale bar, 2 mm. (C) Quantification of HNSCC lesion number and lesion area (mm^3^). n = 8 mice in each group. Data are represented as mean ± SD. **p < 0.01 and ***p < 0.001 by unpaired Student’s t test. (D) Representative H&E staining of HNSCC and Quantification of HNSCC invasion grades. Scale bar, 100 μm. n = 8 mice in each group. *p < 0.05 by Mantel-Haenszel chi-square test (E-I) Representative IHC staining and IHC Score of Ki67 (E), Nat10 (G), Rnps1 (H) and Jun (I) in HNSCC and PCK (F) in lymph node. The scale bar of (F) is 500 μm and the rest are 100 μm. n = 8 mice in each group. Data are represented as mean ± SD. *p < 0.05 and ***p < 0.001 in (E), (G), (H), and (I) by unpaired Student’s t test. *p < 0.05 in (F) by Chi-square test. (A) (J) Immunofluorescence staining of *K14CreER; RiboTag* mice. Epithelial tissues were stained with anti-K14 antibody (green) and anti-HA antibody (red). Nuclei are stained with DAPI (blue). Scale bar, 50 μm. (B) (K) The levels of Rnps1 mRNA and Jun mRNA at the ribosome combined stages and input stages detected by Ribo-tag assay between WT and *K14CreER; RiboTag; Nat10^flox/flox^* mice. Data are represented as mean ± SD. *p < 0.05 and ***p < 0.001 by unpaired Student’s t test. (C) (L) NWB & NB assay (left) and relative gray values (right) showed ac^4^C modification levels of RNA and expression of non-ac^4^C tRNAs and ac^4^C tRNAs (red font) between NAT10^cKO-Ctrl^ and Nat10^cKO^ mice. NB of U6 snRNA and GelRed staining were used as loading control. n = 2 independent biological replicates. Data are represented as mean ± SD. *p < 0.05, **p < 0.01 and ***p < 0.001 in relative gray values by unpaired Student’s t test. (A) (M) Representative image of tongue visible lesions between control and treatment group. Scale bar, 2 mm. n = 10 mice in each group. (B) (N) Quantification of HNSCC lesion number and lesion area (mm^3^). n = 10 mice in each group. Data are represented as mean ± SD. ***p < 0.001 by unpaired Student’s t test. (C) (O) Representative H&E staining of HNSCC and Quantification of HNSCC invasion grades between control group and Remodelin-treated group. Scale bar, 100 μm. n = 10 mice in each group. **p < 0.01 by Mantel-Haenszel chi-square test.

**Figure 7.**
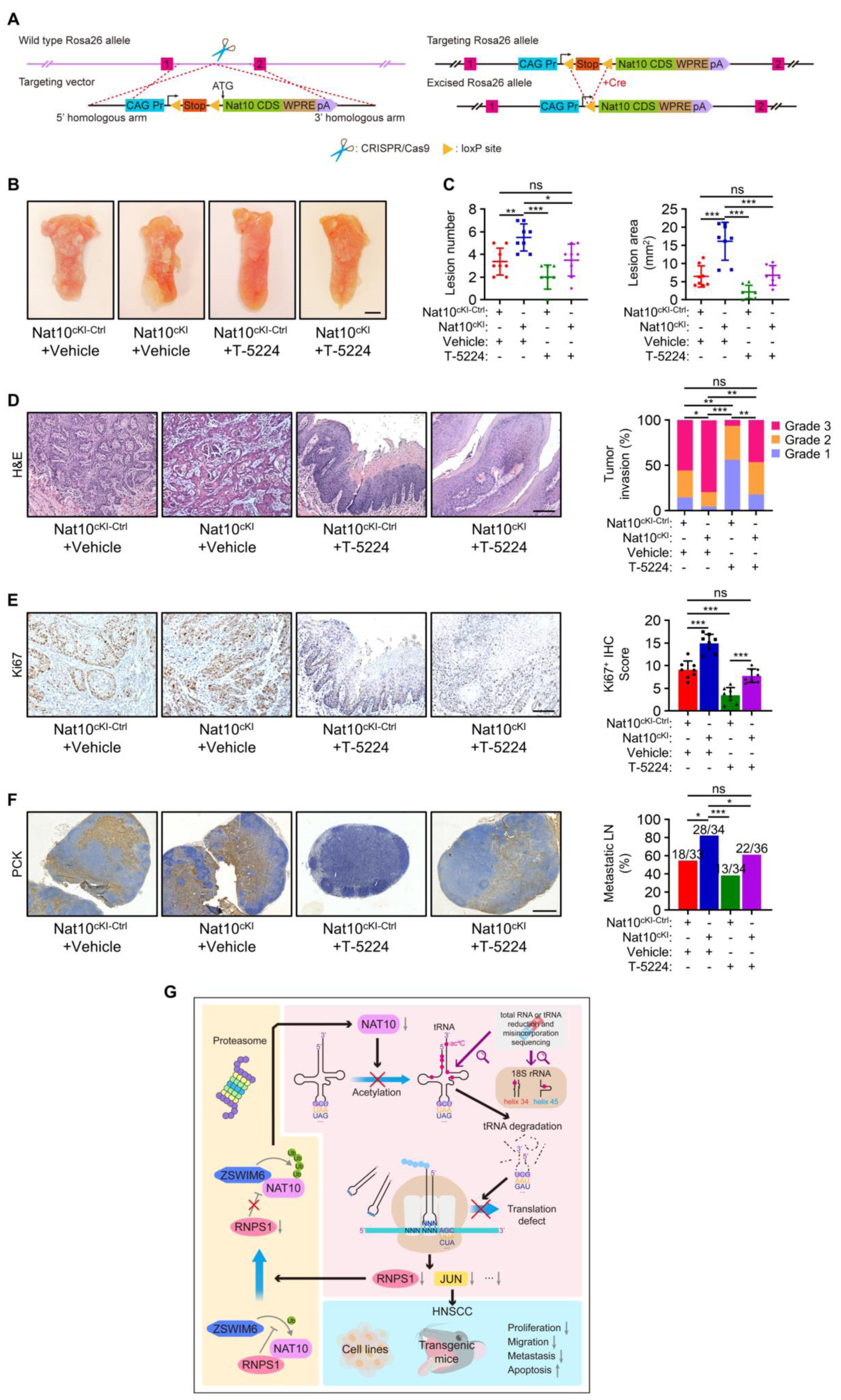
Nat10 drives chemically-induced HNSCC tumorigenesis. (A) Targeting strategy of Nat10^cKI^ mouse. Insertion of the targeting vector Nat10 mROSA-KI-12p-A into the Rosa26 locus by homologous recombination in mouse oosperm cells. (B) Representative image of tongue visible lesions in different treatment groups. n = 8 mice in each group. Scale bar, 2 mm. (C) Quantification of HNSCC lesion number and lesion area (mm^3^). n = 8 mice in each group. Data are represented as mean ± SD. *p < 0.05, **p < 0.01 and ***p < 0.001 by One-way ANOVA. (D) Representative H&E staining of HNSCC and quantification of HNSCC invasion grades in different treatment groups. n = 8 mice in each group. Scale bar, 100 μm. *p < 0.05, **p < 0.01 and ***p < 0.001 by Mantel-Haenszel chi-square test. (E and F) Representative IHC staining and IHC Score of Ki67 (E) in HNSCC and PCK (A) (F) in lymph node. n = 8 mice in each group. The scale bar of (E) is 100 μm and (F) is 500 μm. Data are represented as mean ± SD in (E). ***p < 0.001 in (E) by One-way ANOVA. *p < 0.05 and ***p < 0.001 in (F) by Chi-square test. (B) (G) Schematic model of the underlying mechanism of ac^4^C tRNA modification regulated by NAT10 in the regulation of HNSCC tumorigenesis and development.

To further explore the role of Nat10 and functional relevance of Jun regulated by Nat10 in mouse HNSCC formation, we applied 4NQO to *K14Cre; Nat10^knockin/knockin^* (Nat10^cKI^) and *K14Cre; Nat10^wt/wt^* (Nat10^cKI-Ctrl^) mice. One month before harvesting, Nat10^cKI^ and Nat10^cKI-Ctrl^ were intraperitoneally injected daily with control vehicle or T-5224, a chemical inhibitor of FOS/JUN activator protein-1 (AP1) pathway (Fig. 7B). 26 weeks after initial 4NQO treatment, we found more lesions and larger areas of lesions in the Nat10^cKI^ mice (Fig. 7C). In addition, we observed more poorly-differentiated and muscle-invasive tumors formed after overexpression of Nat10 (Fig. 7D). Notably, more proliferation and metastases of tumor cells were detected in Nat10^cKI^ mice compared to control mice (Fig. 7E and F). The rescue experiments indicated that treatment of T-5224 abrogated the oncogenic effects of Nat10 overexpression (Fig. 7B-F and Fig. S11F-H). It is worth noting that we observed no discernible defect in the homeostasis of oral epithelium after knockout or overexpression of Nat10 (Fig. S11I and J), suggesting Nat10 is more relevant to tumorigenesis instead of tissue homeostasis.

## Discussion

RNA modifications hold a promising function not only because of their prognostic value but also because of their role as potential therapeutic targets in cancer (*26*). For example, mRNA N6-methyladenosine (m^6^A), 5-methylcytosine (m^5^C) and pseudouridine (Ψ) regulators have been reported to promote proliferation, invasion, metastasis, stem cell-like properties or immune system evasion in various malignant tumors, including HNSCC (*12, 27, 28*). Recently, advance in mRNA modification research have led to successful development of novel potential cancer drugs targeting epitranscriptomic modifiers (*2, 29*). On the other hand, dysregulation of tRNA modifications or tRNA modifier enzymes contribute greatly to the development of various cancers (*26, 30*). However, the roles of the tRNA modification changes in HNSCC remain poorly characterized. Here, we observed upregulation of RNA ac^4^C and its catalyzing enzyme NAT10 in HNSCC. We also found that expression of NAT10 is linked to poor overall survival in HNSCC patients, suggesting a role of RNA ac^4^C in HNSCC.

In the early days, ac^4^C were identified in tRNA or rRNA (*3*). Recently, contradictory functions of NAT10 have been reported in terms of whether NAT10 was able to catalyze the ac^4^C on mRNAs in mammalian cells (*7, 8, 10, 31*). To parse the role of NAT10 in HNSCC, we modified previous published approaches and examined the distribution of ac^4^C signal across RNA in HNSCC. In our study, both acRIP-seq and ac^4^C-seq do not recognize reliable ac^4^C signal on mRNA, indicating that ac^4^C does not occur on mRNA, at least in HNSCC cells. Instead, we consistently identified tRNA ac^4^C in our sequencing data. In general, chemical-based approaches are considered as more reliable ways to detect RNA modification in densely modified tRNA than antibody-based methods using next-generation sequencing (*5, 10*). However, the nature of tRNAs, such as highly-structured, intensively modified and relatively small percentage in total RNA, makes it difficult to fully identify the ac^4^C modification sites. To overcome this, we first isolated tRNA and used the purified bacterial demethylase AlkB to remove tRNA methylation to achieve more efficient and accurate tRNA sequencing. By doing so, we were able to better profile ac^4^C tRNA and identify several novel tRNA ac^4^C sites in HNSCC. One common mechanism for tRNA epitranscriptome in cancer progression is to increase high-fidelity translation of oncogenic transcripts that possess skewed usage patterns of related codons (*32*). In fact, codon usage analysis of the most downregulated genes in our data showed that these gene contained higher percentage of tRNA codon that were ac^4^C modified.

Intriguingly, we found RNPS1 can interact with NAT10 in HNSCC. Genes regulated by NAT10 overlapped greatly with the downstream targets of RNPS1. Depletion of NAT10 or RNPS1 resulted in inhibition of HNSCC tumor growth and metastasis via downregulation of IL-6 and IL-8 signaling pathways, which are tightly governed by AP-1 (activator protein 1) transcription factors, including JUN, FOS and ATF (*33-36*). In our study, we found the translation of JUN is consistently down-regulated in NAT10 or RNPS1. We also found that JUN transcript contains high proportion of ac^4^C codons, which are functionally important to the translation of JUN. In addition, we showed that exogenous over-expression of JUN partially rescued the effects of depletion of NAT10 or RNPS1 in HNSCC, supporting JUN as a downstream mediator of NAT10 or RNPS1.We found an intriguing phenomenon that depletion of RNPS1 led to decrease level of NAT10 protein. Although RNPS1 possesses RNA binding activity, our result showed it was not able to directly catalyze ac^4^C on tRNA. Rather, RNPS1 was able to stabilize NAT10 by inhibiting the binding of E3 ubiquitin ligase ZSWIM6 (Fig. 7G), uncovering a novel function of RNPS1 in RNA modification.

Recent studies have highlighted the importance of RNA modifications in controlling gene expression in multiple cellular and biological processes, including tumorigenesis (*26, 37*). To interrogate the role of Nat10 in cancer development in vivo, we generated Nat10 conditional knockout and knockin mouse model. We confirmed an essential role of Nat10 in mouse HNSCC model. More interestingly, we found perturbation of Nat10 did not cause major defect in mouse oral epithelium tissue, suggesting Nat10 targeting might not lead to severe adverse effects. Excitingly, recently studies have shown that Remodelin is able to block the growth of tumor cells in a variety of cancers, including HNSCC (*38, 39*).

Together, our study, revealing NAT10 as a robust pro-tumorigenic factor in HNSCC, highlights an important role for tRNA ac^4^C in cancer pathogenesis. Furthermore, our results support tRNA ac^4^C as a main function of NAT10 to regulate the translation activities in HNSCC (Fig. 7G). Our findings might promote a novel insight for HNSCC prognosis and treatment.

---

**Figure S1.**
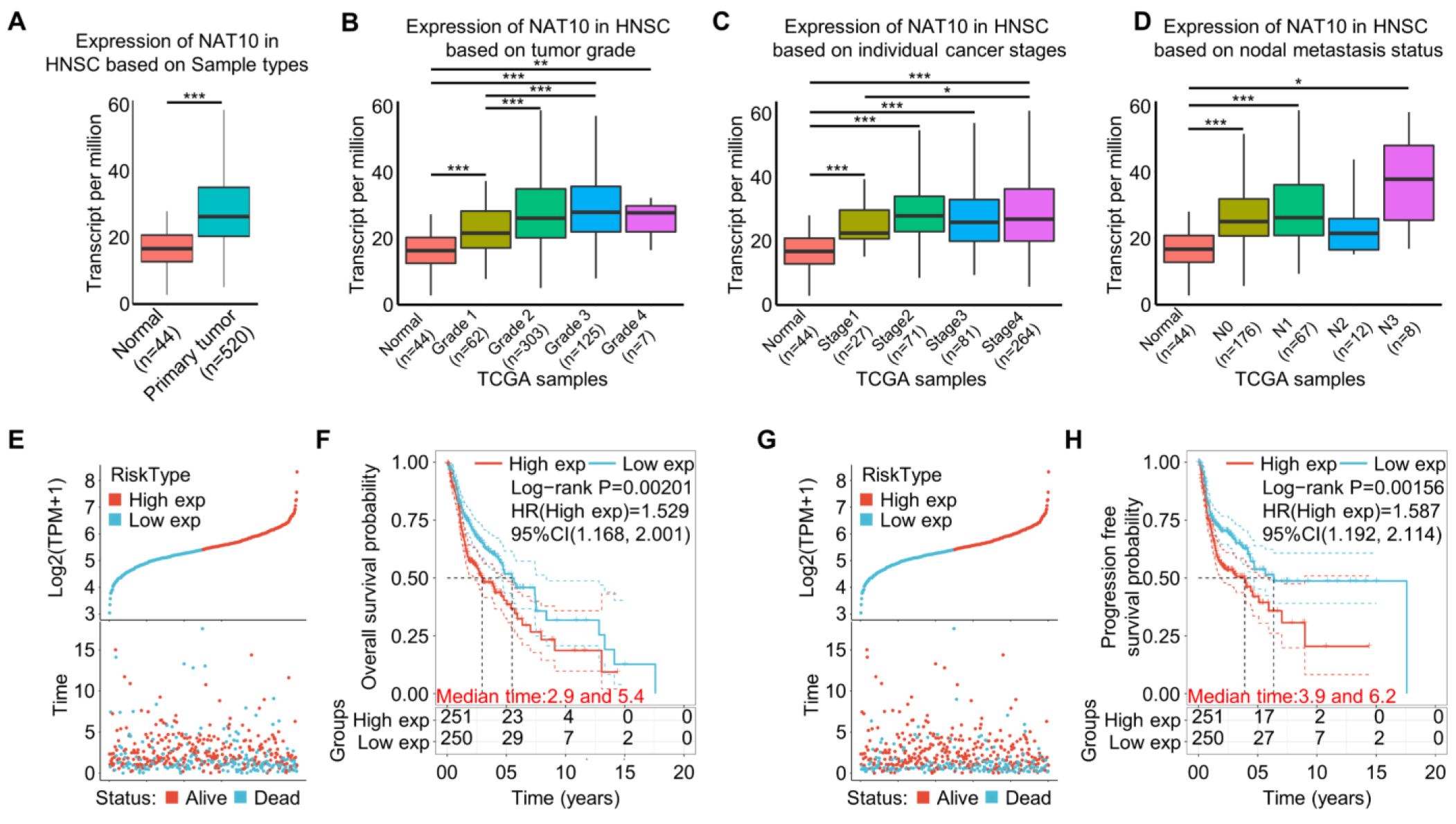
NAT10 is elevated in HNSCC and correlates with HNSCC progression using TCGA datasets, related to Figure 1. (A) Expression of NAT10 between normal (n = 44) and primary tumor (n = 520) of TCGA data. ***p < 0.001 by unpaired Student’s t test. (B) The expression of NAT10 increased with the degree of HNSCC tumor grade in TCGA dataset. Grade 1: well differentiated (low grade); Grade 2: moderately differentiated (intermediate grade); Grade 3: poorly differentiated (high grade); Grade 4: Undifferentiated (high grade). Normal, n = 44; Grade 1, n = 62; Grade 2, n = 303; Grade 3, n = 125; Grade 4, n = 7. **p < 0.01 and ***p < 0.001 by One-way ANOVA. (C) Box plot showing the correlation between NAT10 expression and HNSCC individual cancer stages of TCGA dataset. Normal, n = 44; Stage1, n = 27; Stage2, n = 71; Stage3, n = 81; Stage4, n = 264. *p < 0.05 and ***p < 0.001 by One-way ANOVA. (D) The correlation between NAT10 expression and HNSCC nodal metastasis status of TCGA dataset. N0: no regional lymph node metastasis; N1: metastases in 1 to 3 axillary lymph nodes; N2: metastases in 4 to 9 axillary lymph nodes; N3: metastases in 10 or more axillary lymph nodes. Normal, n = 44; N0, n = 176; N1, n = 67; N2, n = 12; N3, n = 8 mice in each group. *p < 0.05 and ***p < 0.001 by One-way ANOVA. (A) (E) The correlation between NAT10 expression and overall survival time or overall survival status in TCGA dataset. The top represents the scatter plot of the gene expression from low to high, and different colors represent different expression groups; the bottom represents the distribution of survival time and survival status corresponding to gene expression of different samples. (B) (F) The Kaplan-Meier survival curve distribution of NAT10 expression and overall survival probability in TCGA dataset was tested by log rank. HR (high exp) represents the risk coefficient of the high expression group relative to the low expression group. If HR > 1 represents the risk factor of the gene, and if HR < 1 represents the protective factor of the gene; 95% Cl represents HR confidence interval; Median time represents the time when the survival rate of high expression group or low expression group is 50%. p = 0.00201 by Log-rank test. (C) (G) NAT10 expression, progression free survival time and progression free survival status in TCGA dataset. (D) (H) The Kaplan-Meier survival curve distribution of NAT10 expression and progression free survival probability in TCGA dataset. p = 0.00156 by Log-rank test.

**Figure S2.**
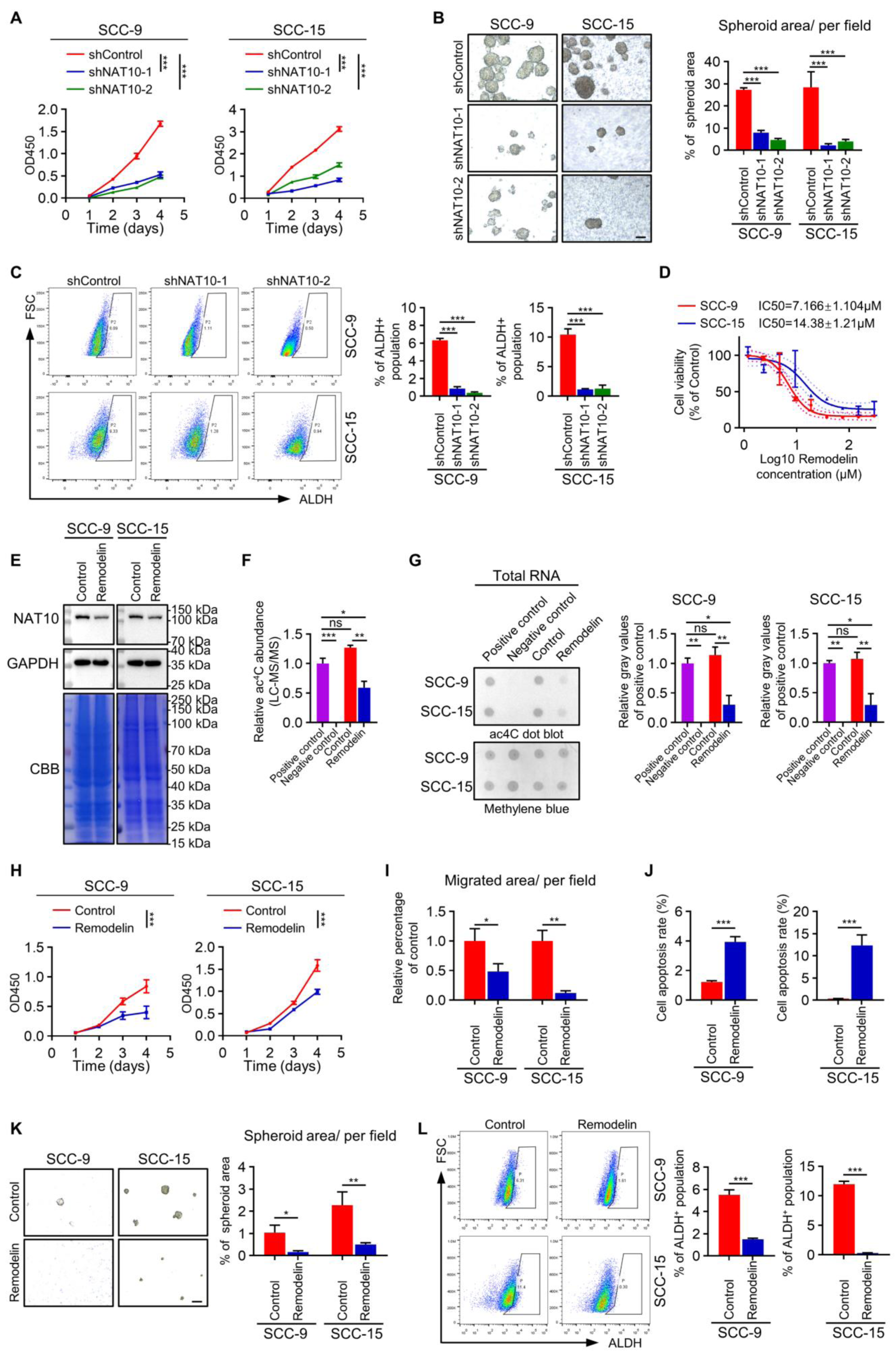
NAT10 plays a vital role in HNSCC progression and metastasis, related to Figure 2. (A-C) Proliferation (A), sphere formation (B) and aldehyde dehydrogenase activity (C) between the shControl, shNAT10-1 and shNAT10-2 in HNSCC cell lines. The scale bar of (B) is 100 μm. Data are represented as mean ± SD. ***p < 0.001 in (A) by Two-way ANOVA. ***p < 0.001 in (B) and (C) by One-way ANOVA. (A) (D) IC_50_ values for Remodelin in cytotoxicity assays with HNSCC cell lines. SCC-9, IC_50_ ± SE = 7.166 ± 1.104 μM; SCC-15, IC_50_ ± SE = 14.38 ± 1.21 μM. Data are represented as mean ± SD. (B) (E) The expression of NAT10 in SCC-9 and SCC-15 after with/without Remodelin treatment. Cells were treated with Remodelin at a concentration of 8 μM (SCC-9) or 15 μM (SCC-15) for 24-48 h. GAPDH and CBB staining were used as loading control. (C) (F) The bar chart of relative ac^4^C abundance of RNA by LC-MS/MS from SCC-15 cells. Positive control: ribosomal RNA; Negative control: poly(A)-enriched RNA; The remaining samples are total RNA from Control (DMSO) and Remodelin groups. Cells were treated with Remodelin at a concentration of 8 μM (SCC-9) or 15 μM (SCC-15) for 24-48 h. n = 2 independent biological replicates. Data are represented as mean ± SD. *p < 0.05, **p < 0.01 and ***p < 0.001 by One-way ANOVA. (D) (G) Dot blot (left) and relative gray values (right) showed the ac^4^C level of RNA in positive control, negative control, Control, Remodelin groups with methylene blue staining as loading control. Cells were treated with Remodelin at a concentration of 8 μM (SCC-9) or 15 μM (SCC-15) for 24-48 h. n = 2 independent biological replicates. Data are represented as mean ± SD. *p < 0.05 and **p < 0.01 by One-way ANOVA. (H-L) Remodelin reduced the proliferation ability (H), migration (I), sphere formation ability (K), aldehyde dehydrogenase activity (L) and increased rate of cell apoptosis (J) of HNSCC cell lines. Cells were treated with Remodelin at a concentration of 8 μM (SCC-9) or 15 μM (SCC-15) for 4 d (H), 48 h (I, J and L) or 3-4 d (K), respectively. The scale bar of (K) is 100 μm. Data are represented as mean ± SD. ***p < 0.001 in (E) (H) by Two-way ANOVA. *p < 0.05, **p < 0.01 and ***p < 0.001 in (I-L) by unpaired Student’s t test.

**Figure S3.**
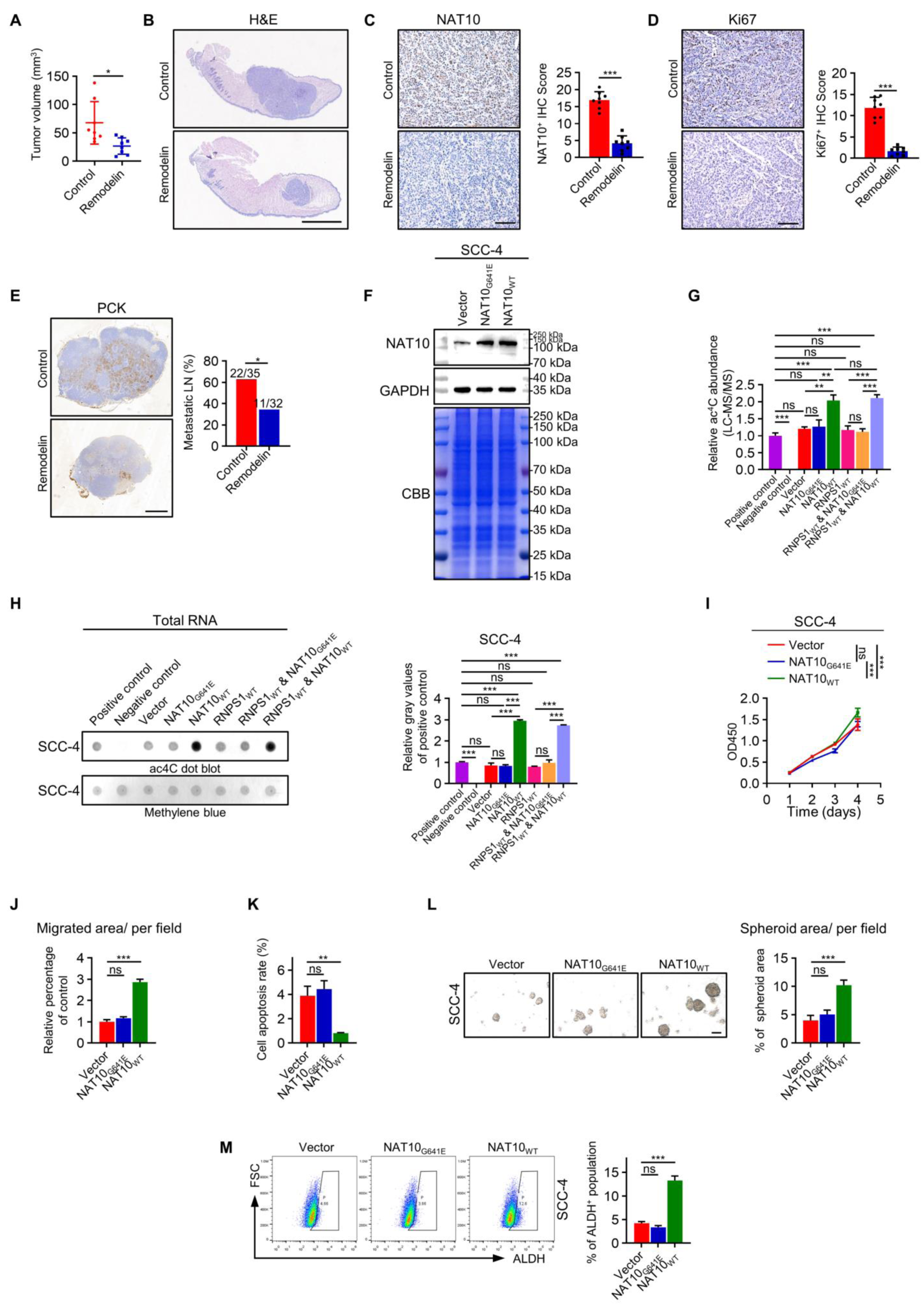
NAT10 plays a vital role in HNSCC progression and metastasis, related to Figure 2. (A) Tumor volume (mm^3^) of orthotopic transplantation tumor between controlled group and treated group. n = 8 mice in each group. Data are represented as mean ± SD. *p < 0.05 by unpaired Student’s t test (B-E) H&E staining of tongue (B) with orthotopic transplantation tumor and IHC staining with score of NAT10 (C), Ki67 (D) in orthotopic transplantation tumor and PCK (E) in lymph node between Control and Remodelin groups. The scale bar represents 2mm in panel (B), 500 μm in panel (E) and 100 μm for the rest. n = 8 mice in each group. Data are represented as mean ± SD in (C) and (D). ***p < 0.001 in (C) and (D) by unpaired Student’s t test. *p < 0.05 in (E) by Chi-square test. (A) (F) WB showed the protein expression of NAT10 when NAT10_WT_ or NAT10_G641E_ was exogenously overexpressed in SCC-4. GAPDH and CBB staining were used as loading control. (B) (G) The relative ac^4^C level of RNA detected by LC-MS/MS in Positive control, Negative control, Vector, NAT10-MUT, NAT10-WT, RNPS1-WT, RNPS1-WT & NAT10-MUT and RNPS1-WT & NAT10-WT groups of SCC-4 cells. Positive control: ribosomal RNA; Negative control: poly(A)-enriched RNA; The remaining samples are total RNA from Vector, NAT10-MUT, NAT10-WT, RNPS1-WT, RNPS1-WT & NAT10-MUT and RNPS1-WT & NAT10-WT groups. n = 2 independent biological replicates. Data are represented as mean ± SD. **p < 0.01 and ***p < 0.001 by One-way ANOVA. (C) (H) Dot blot (left) and relative gray values (right) showed the ac^4^C modification level of RNA in positive control, negative control, Vector, NAT10-MUT, NAT10-WT, RNPS1-WT, RNPS1-WT & NAT10-MUT and RNPS1-WT & NAT10-WT groups with methylene blue staining as loading control. n = 2 independent biological replicates. Data are represented as mean ± SD. ***p < 0.001 by One-way ANOVA. (I-M) The cell phenotype assay including proliferation ability (I), migration (J), cell apoptosis (K), sphere formation (L) and aldehyde dehydrogenase activity (M) of SCC- 4 cell line with overexpression of Vector, NAT10_WT_ or NAT10_G641E_. The scale bar of (L) is 100 μm. Data are represented as mean ± SD. ***p < 0.001 in (I) by Two-way ANOVA. **p < 0.01 and ***p < 0.001 in (J-M) by One-way ANOVA.

**Figure S4.**
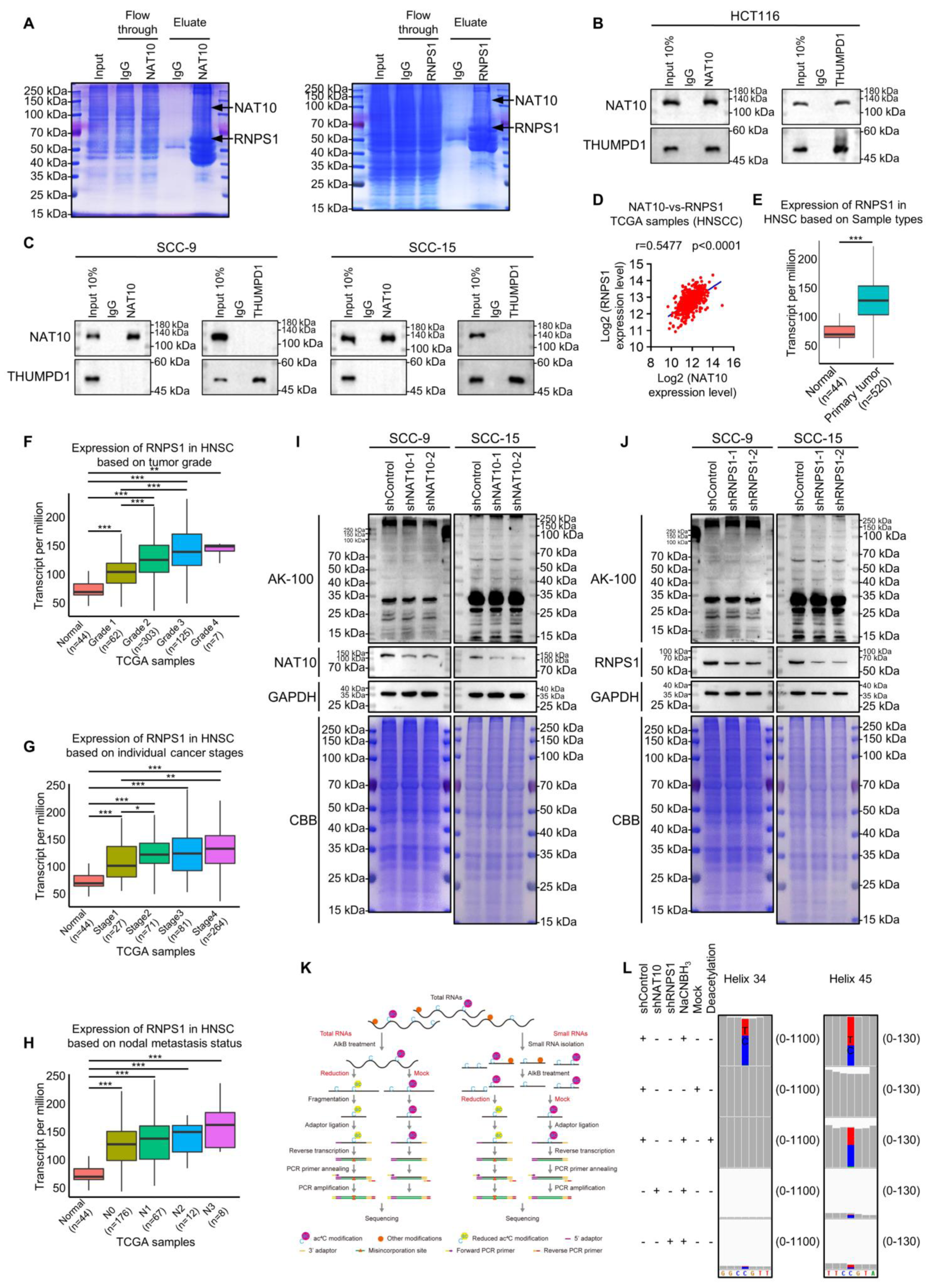
NAT10 interacts with RNPS1 to regulate tRNA ac^4^C modifications, related to Figure 3. (A) Protein samples from different steps of protein immunoprecipitation by NAT10 antibody(left), RNPS1 antibody (right) and IgG antibody from SCC-15 were separated by SDS-PAGE and stained with CBB. (B and C) The co-IP assay of NAT10 and THUMPD1 in HCT116 (B), SCC-9 and SCC- 15 (C). Anti-IgG antibody was used as a negative control. (A) (D) The correlation plot of NAT10 and RNPS1 expression level in HNSCC using TCGA dataset. n = 502, r = 0.5477, p < 0.0001 by Spearman correlation analysis. (B) (E) Expression of RNPS1 between normal (n = 44) and primary tumor (n = 520) of TCGA data. ***p < 0.001 by unpaired Student’s t test. (F-H) The correlation between RNPS1 expression and HNSCC tumor grade (F), individual cancer stages (G) and nodal metastasis status (H) of TCGA dataset. *p < 0.05, **p < 0.01 and ***p < 0.001 by One-way ANOVA. (I and J) The lysine acetylation levels of protein detected by WB with anti-acetylated-Lysine (Ac-K2-100) antibody in NAT10 KD (I) or RNPS1 KD (J) cells. GAPDH and CBB staining were used as loading control. (A) (K) Schematic diagram of total RNA reduction and misincorporation sequencing (left) and tRNA reduction and misincorporation sequencing (TRMC-seq) (right). (B) (L) Misincorporation rates in total RNA from SCC-15 cells are shown for known sites in 18S rRNA. blue letters C and bars, cytidine; red letters T and bars, thymidine; green letters A, adenosine; orange letters G, guanosine. The number on the right indicates the number of nucleotides in the ordinate.

**Figure S5.**
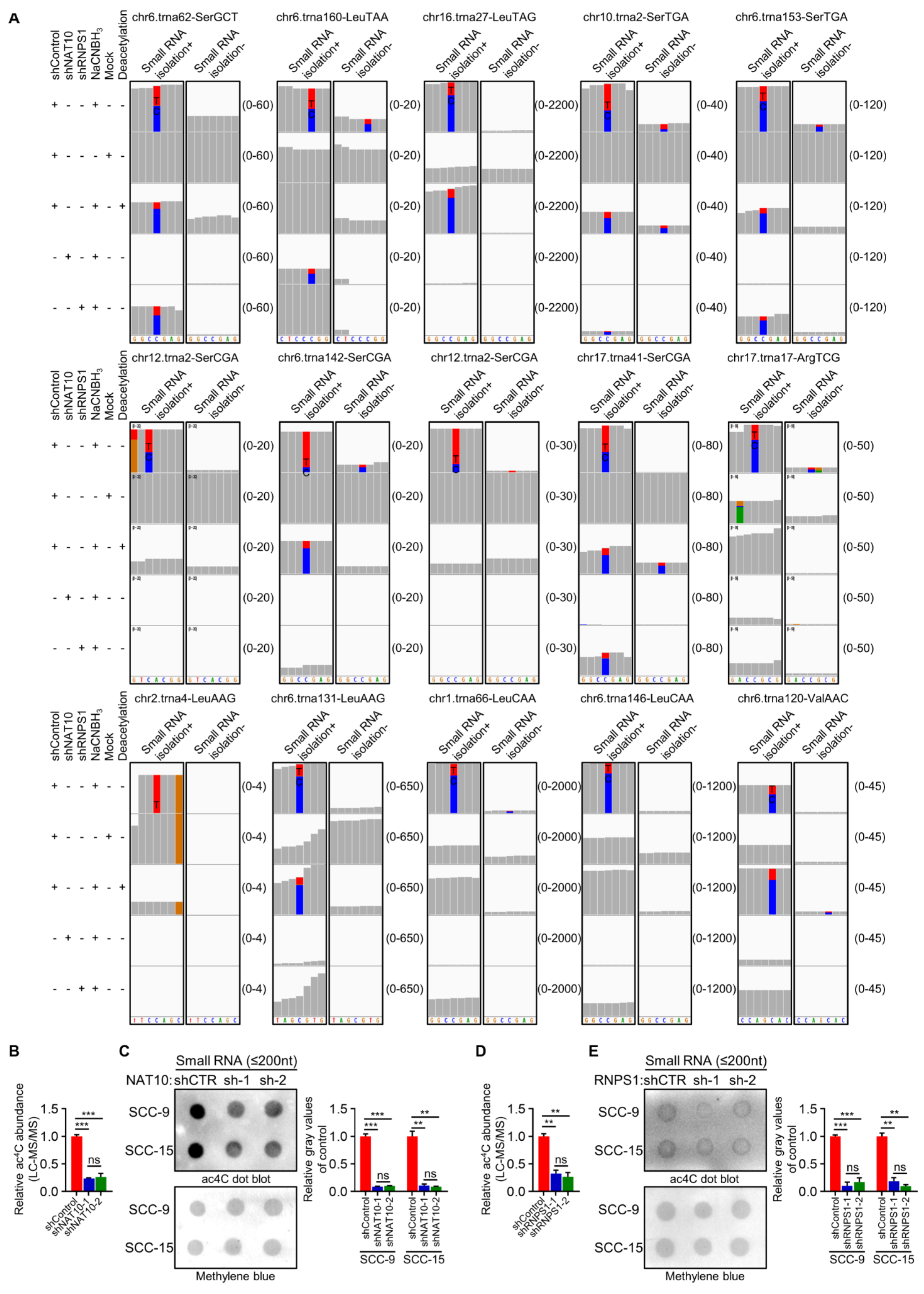
NAT10 interacts with RNPS1 to regulate tRNA ac^4^C modifications, related to Figure 3. (A) From left to right, from top to bottom, Misincorporation rates of ac^4^C_12_ in tRNA^Ser(GCT)^, ac^4^C_79_ in tRNA^Leu(TAA)^, ac^4^C_12_ in tRNA^Leu(TAG)^, ac^4^C_12_ in tRNA^Ser(TGA)^, ac^4^C_12_ in tRNA ^Ser(TGA)^, ac^4^C_3_ in tRNA^Ser(CGA)^, ac^4^C_12_ in tRNA^Ser(CGA)^, ac^4^C_12_ in tRNA^Ser(CGA)^, ac^4^C_12_ in tRNA^Ser(CGA)^, ac^4^C_4_ in tRNA^Arg(TCG)^, ac^4^C_50_ in tRNA^Leu(AAG)^, ac^4^C_6_ in tRNA^Leu(AAG)^, ac^4^C_12_ in tRNA^Leu(CAA)^, ac^4^C_12_ in tRNA^Leu(CAA)^ and ac^4^C_65_ in tRNA^Val(AAC)^ in small RNA and total RNA from SCC-15 cells were detected by TRMC- seq or total RNA reduction and misincorporation sequencing. blue letters C and bars, cytidine; red letters T and bars, thymidine; green letters A and bars, adenosine; orange letters G and bars, guanosine. Small RNA isolation+, detected by TRMC-seq; Small RNA isolation-, detected by total RNA reduction and misincorporation sequencing. The number on the right indicates the number of nucleotides in the ordinate. (B) The relative ac^4^C level of small RNA (≤200nt) detected by LC-MS/MS in Control and NAT10-KD groups of SCC-15 cells. n = 2 independent biological replicates. Data are represented as mean ± SD. ***p < 0.001 by One-way ANOVA. (A) (C) Dot blot (left) and relative gray values (right) showed the ac^4^C content of small RNA in Control and NAT10-KD groups of SCC-9 or SCC-15 cells with methylene blue staining as loading control. n = 2 independent biological replicates. Data are represented as mean ± SD. **p < 0.01 and ***p < 0.001 by One-way ANOVA. (B) (D) The bar chart showed relative ac^4^C abundance of small RNA by LC-MS/MS between Control and RNPS1-KD groups of SCC-15 cells. n = 2 independent biological replicates. Data are represented as mean ± SD. **p < 0.01 by One-way ANOVA. (C) (E) Dot blot assay (left) and relative gray statistical values (right) showed that compared with the Control, RNPS1-KD group displayed the downward trend in ac^4^C levels of small RNA of SCC-9 or SCC-15 cells with methylene blue staining as loading control. n = 2 independent biological replicates. Data are represented as mean ± SD. **p < 0.01 and ***p < 0.001 by One-way ANOVA.

**Figure S6.**
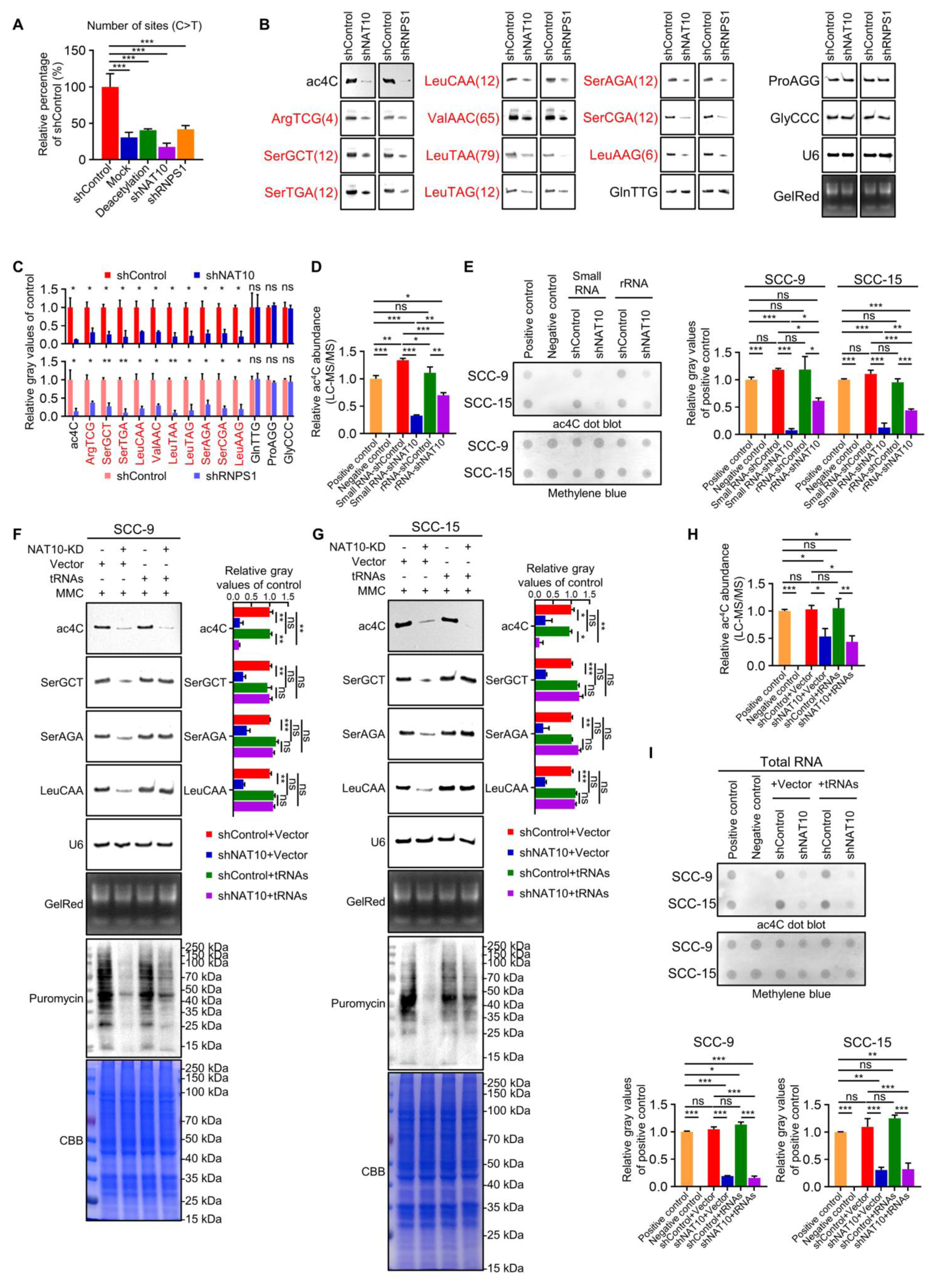
NAT10 interacts with RNPS1 to regulate tRNA ac^4^C modifications, related to Figure 3 and Figure 4. (A) The number of misincorporation (C>T) sites detected in shControl, Mock, Deacetylation, shNAT10, shRNPS1 groups. Data are represented as mean ± SD. ***p < 0.001 by One-way ANOVA. (B and C) NWB & NB (B) and relative gray values (C) showed ac^4^C modification levels of RNA and expression of non-ac^4^C tRNAs and ac^4^C tRNAs (red font) between Control, NAT10 KD and RNPS1 KD groups. NB of U6 snRNA and GelRed staining were used as loading control. The numbers in brackets represent the ac^4^C sites. n = 2 independent biological replicates. Data are represented as mean ± SD. *p < 0.05 and **p < 0.01 in (C) by unpaired Student’s t test. (I) (D) The bar chart of relative ac^4^C abundance of RNA by LC-MS/MS in Positive control, Negative control, Small RNA-Control, Small RNA-NAT10-KD, rRNA-Control and rRNA-NAT10-KD of SCC-15 cells. Positive control: ribosomal RNA; Negative control: poly(A)-enriched RNA. n = 2 independent biological replicates. Data are represented as mean ± SD. *p < 0.05, **p < 0.01 and ***p < 0.001 by One-way ANOVA. (II) (E) Dot blot (left) and relative gray values (right) showed the ac^4^C level of RNA in Positive control, Negative control, Small RNA-Control, Small RNA-NAT10-KD, rRNA-Control and rRNA-NAT10-KD groups with methylene blue staining as loading control. n = 2 independent biological replicates. Data are represented as mean ± SD. *p < 0.05, **p < 0.01 and ***p < 0.001 by One-way ANOVA. (F and G) NWB & NB showed ac^4^C modification levels of RNA and expression of SerGCT, SerAGA and LeuCAA tRNAs between Control + Vector, NAT10-KD + Vector, Control + tRNAs-OE (SerGCT, SerAGA and LeuCAA) and NAT10-KD + tRNAs-OE groups in SCC-9 (F) and SCC-15 (G). The corresponding relative gray values is on the right. NB of U6 snRNA and GelRed staining were used as loading control. The following is the corresponding puromycin assay and CBB staining was used as loading control. MMC was added to inhibit the effect of cell proliferation. Data are represented as mean ± SD. *p < 0.05, **p < 0.01 and ***p < 0.001 by One-way ANOVA. (A) (H) The relative ac^4^C level of RNA by LC-MS/MS in SCC-15 cells. Positive control: ribosomal RNA; Negative control: poly(A)-enriched RNA; The remaining samples are total RNA from Control + Vector, NAT10-KD + Vector, Control + tRNAs-OE (SerGCT, SerAGA and LeuCAA) and NAT10-KD + tRNAs-OE groups. n = 2 independent biological replicates. Data are represented as mean ± SD. **p < 0.01 and ***p < 0.001 by One-way ANOVA. (I) Dot blot (up) and relative gray values (down) showed the ac^4^C content of RNA in Positive control, Negative control, Control + Vector, NAT10-KD + Vector, Control + tRNAs-OE (SerGCT, SerAGA and LeuCAA) and NAT10-KD + tRNAs-OE groups with methylene blue staining as loading control. n = 2 independent biological replicates. Data are represented as mean ± SD. ***p < 0.001 in (H) by One-way ANOVA.

**Figure S7.**
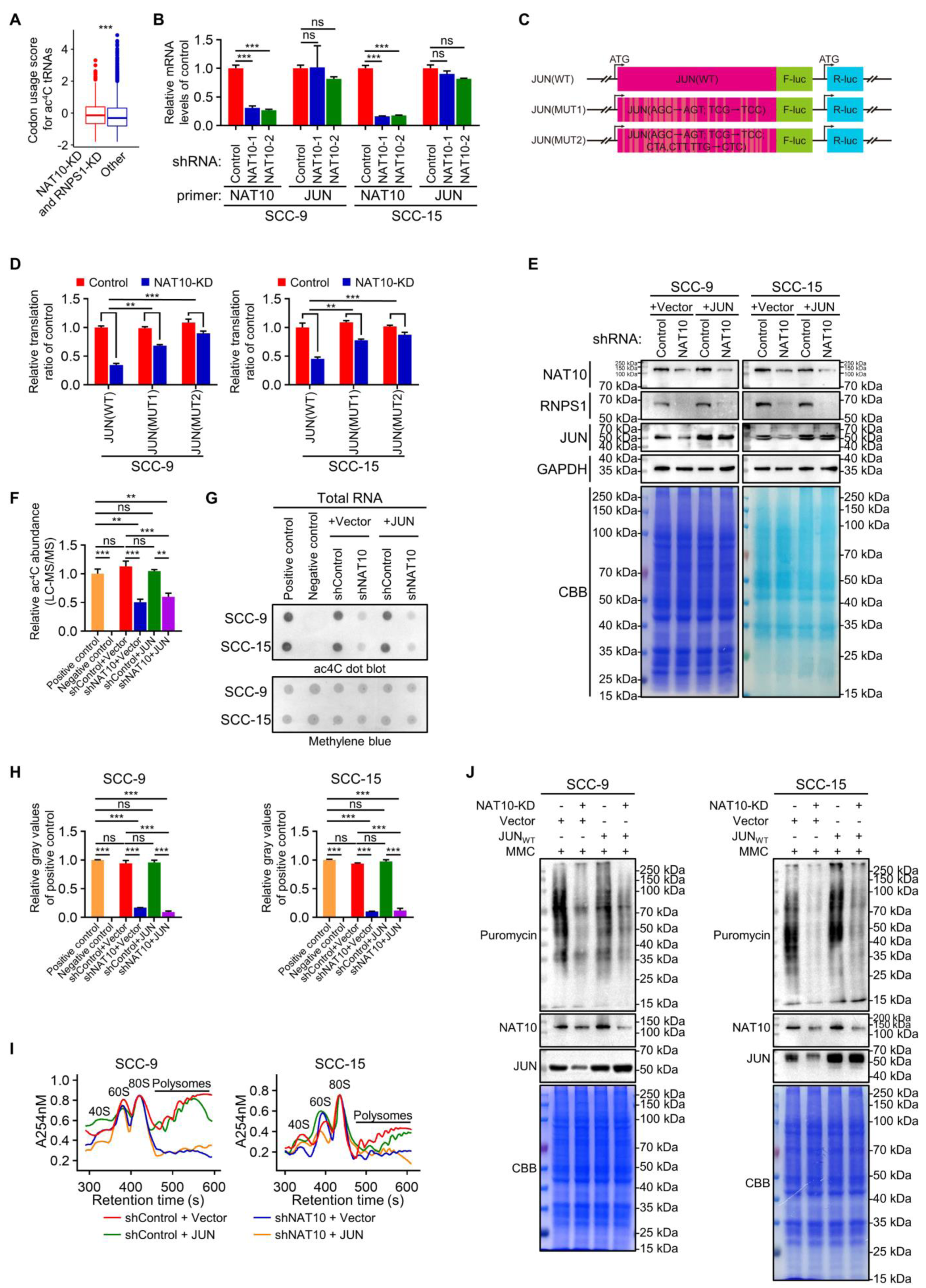
Integrated analysis of multiple ‘omics’ reveal NAT10 and RNPS1 regulate JUN, related to Figure 5. (A) The codon usage scores of common down regulated genes detected by iTRAQ in NAT10-KD and RNPS1-KD were higher than scores of other genes. The box plot parameters are as in Figure 3J. ***p < 0.001 by unpaired Student’s t test. (B) The expression of JUN mRNA remained unchanged after KD of NAT10 in SCC-9 and SCC-15 cells. Data are represented as mean ± SD. ***p < 0.001 by One-way ANOVA. (C and D) Schematic diagram (C) and relative translation ratio column chart (D) of luciferase reporter system. The coding sequence of JUN was inserted into a vector carrying a luciferase reporter system. The mutated JUN transcripts were constructed by replacing the codon AGC to ACT; TCG to TCC (MUT1) or AGC to AGT; TCG to TCC; CTA, CTT, TTG to CTC (MUT2). The relative translation ratios were calculated as the ratio of F-luc translation efficiency (TE) to R-luc translation efficiency and the TE of F-luc and R-luc were calculated by normalizing the luciferase protein levels to mRNA levels. Data are represented as mean ± SD. **p < 0.01 and ***p < 0.001 in (D) by One-way ANOVA. (A) (E) The WB showed that JUN was overexpressed after KD of NAT10 in SCC-9 and SCC-15 cells. GAPDH and CBB staining were used as loading control. (B) (F) The bar chart of relative ac^4^C abundance of RNA by LC-MS/MS from SCC-15 cells. Positive control: ribosomal RNA; Negative control: poly(A)-enriched RNA; The remaining samples are total RNA from Control + Vector, NAT10-KD + Vector, Control + JUN-OE and NAT10-KD + JUN-OE groups. n = 2 independent biological replicates. Data are represented as mean ± SD. **p < 0.01 and ***p < 0.001 by One-way ANOVA. (G and H) Dot blot (G) and relative gray values (H) showed the ac^4^C level of RNA in Positive control, Negative control, Control + Vector, NAT10-KD + Vector, Control + JUN-OE and NAT10-KD + JUN-OE groups with methylene blue staining as loading control. n = 2 independent biological replicates. Data are represented as mean ± SD. ***p < 0.001 in (H) by One-way ANOVA. (I and J) Polysome profiling (I) and Puromycin assay (J) of the Control + Vector, NAT10-KD + Vector, Control + JUN-OE and NAT10-KD + JUN-OE groups in SCC-9 (left) and SCC-15 (right) cells. MMC was added to inhibit the effect of cell proliferation and CBB staining was used as loading control.

**Figure S8.**
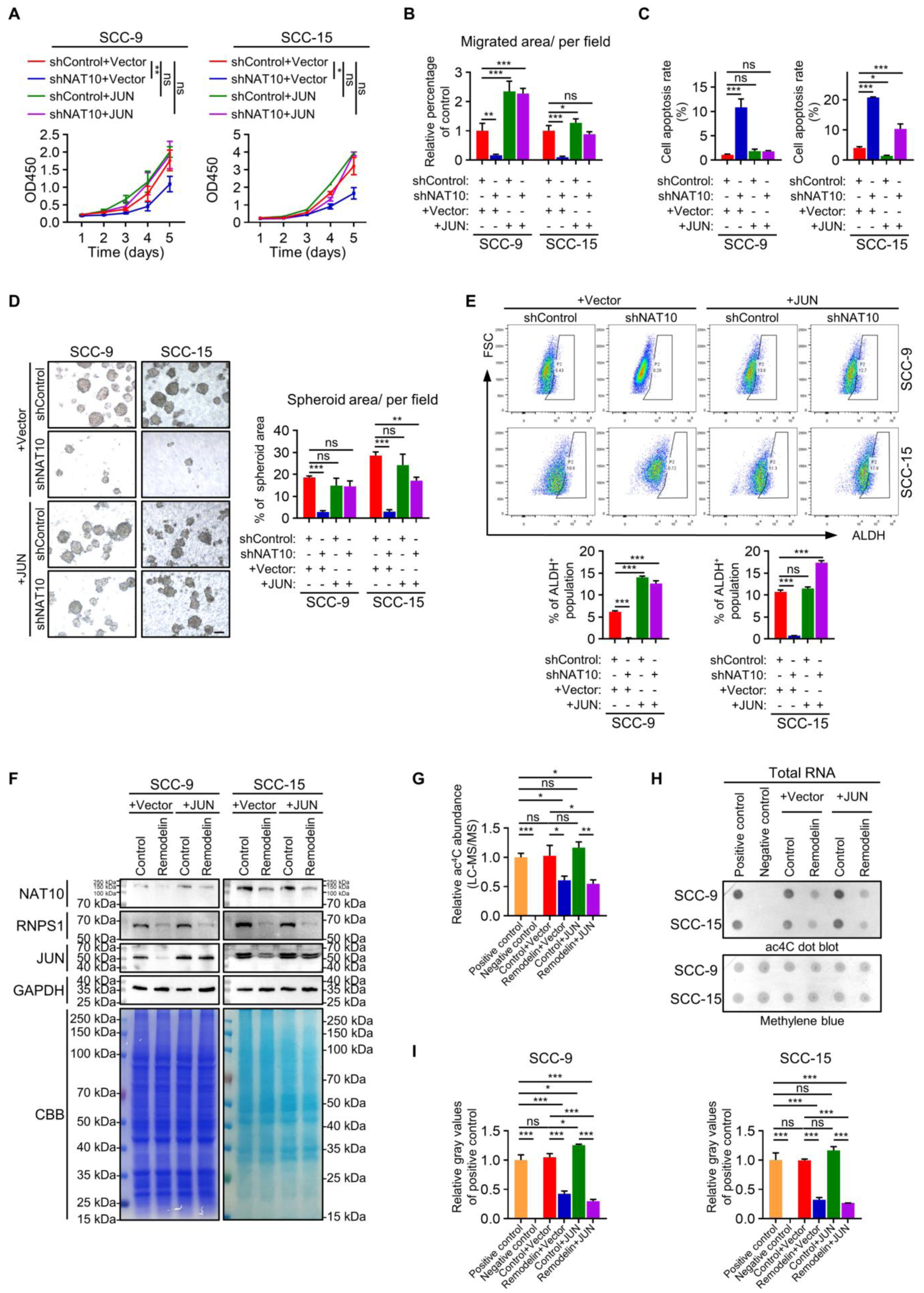
Integrated analysis of multiple ‘omics’ reveal NAT10 and RNPS1 regulate JUN, related to Figure 5. (A-E) In the rescue assay of JUN overexpression after KD of NAT10, the phenotypic differences including proliferation ability (A), migration (B), cell apoptosis (C), sphere formation (D) and aldehyde dehydrogenase activity (E) between the shControl + Vector, shNAT10 + Vector, shControl + JUN and shNAT10 + JUN in HNSCC cell lines were observed. The scale bar of (D) is 100 μm. Data are represented as mean ± SD. *p < 0.05, **p < 0.01 in (A) by Two-way ANOVA. *p < 0.05, **p < 0.01 and ***p < 0.001 in (B-E) by One-way ANOVA. (A) (F) The WB assay showed that JUN was overexpressed after treatment of Remodelin in SCC-9 and SCC-15 cells. Cells were treated with Remodelin at a concentration of 8 μM (SCC-9) or 15 μM (SCC-15) for 24-48 h. GAPDH and CBB staining were used as loading control. (B) (G) The bar chart of relative ac^4^C abundance of RNA by LC-MS/MS from SCC-15 cells. Positive control: ribosomal RNA; Negative control: poly(A)-enriched RNA; The remaining samples are total RNA from DMSO treatment + Vector, Remodelin treatment + Vector, DMSO treatment + JUN-OE and Remodelin treatment + JUN-OE groups. Cells were treated with Remodelin at a concentration of 8 μM (SCC-9) or 15 μM (SCC- (A) 15) for 24-48 h. n = 2 independent biological replicates. Data are represented as mean ± SD. *p < 0.05, **p < 0.01 and ***p < 0.001 by One-way ANOVA. (H and I) Dot blot (H) and relative gray values (I) showed the ac^4^C level of RNA in Positive control, Negative control, DMSO treatment + Vector, Remodelin treatment + Vector, DMSO treatment + JUN-OE and Remodelin treatment + JUN-OE groups with methylene blue staining as loading control. Cells were treated with Remodelin at a concentration of 8 μM (SCC-9) or 15 μM (SCC-15) for 24-48 h. n = 2 independent biological replicates. Data are represented as mean ± SD. *p < 0.05 and ***p < 0.001 in (I) by One-way ANOVA.

**Figure S9.**
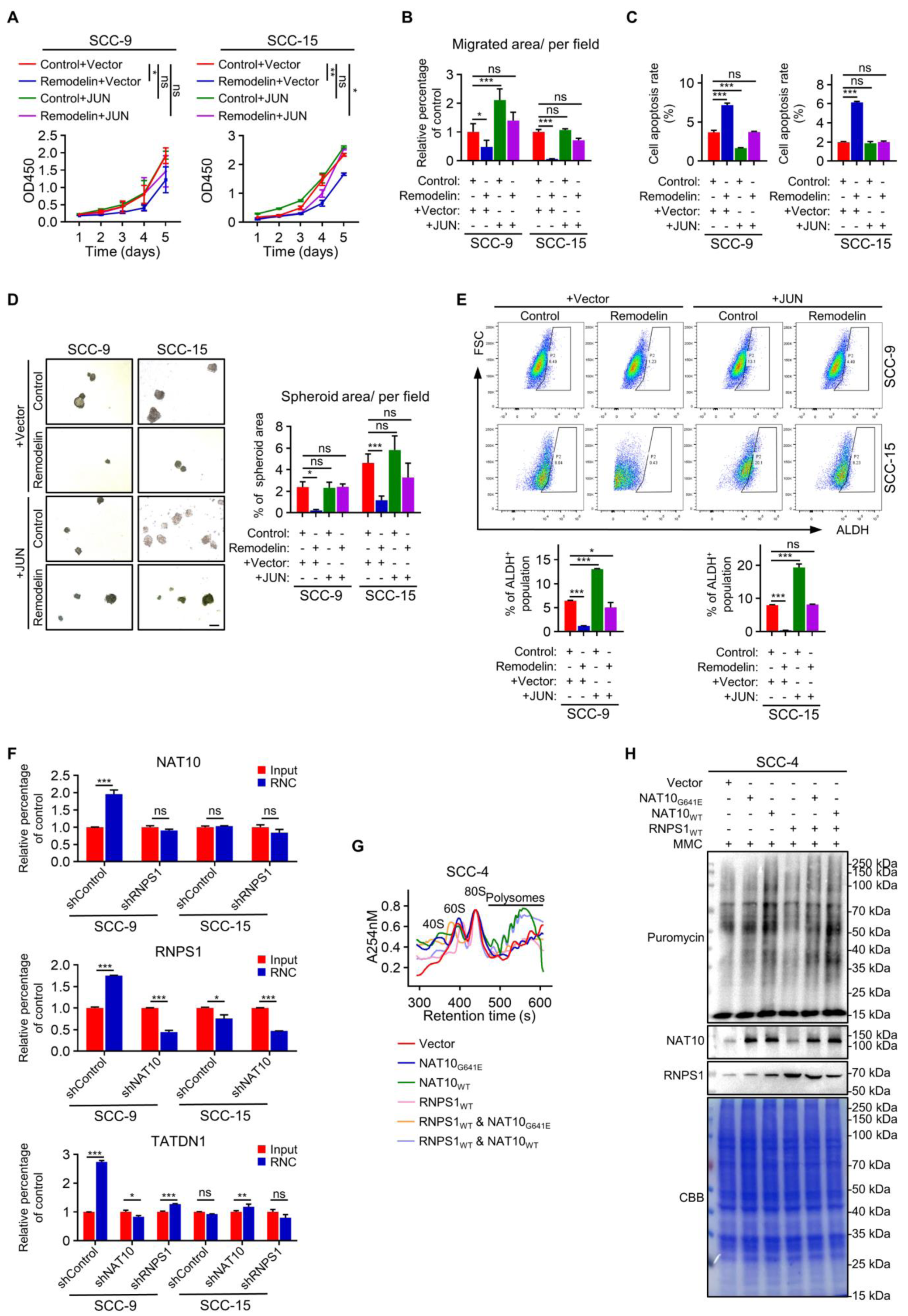
Integrated analysis of multiple ‘omics’ reveal NAT10 and RNPS1 regulate JUN, related to Figure 5. (A-E) In the rescue assay of JUN overexpression after treatment of Remodelin, the phenotypic differences including proliferation ability (A), migration (B), cell apoptosis (C), sphere formation (D) and aldehyde dehydrogenase activity (E) between the Control + Vector, Remodelin + Vector, Control + JUN and Remodelin + JUN in HNSCC cell lines were observed. Cells were treated with Remodelin at a concentration of 8 μM (SCC-9) or 15 μM (SCC-15) for 5 d (A), 48 h (B, C and E) or 3-4 d (D), respectively. The scale bar of (D) is 100 μm. Data are represented as mean ± SD. *p < 0.05, **p < 0.01 in (A) by Two-way ANOVA. *p < 0.05 and ***p < 0.001 in (B-E) by One-way ANOVA. (A) (F) The content of NAT10 mRNA, RNPS1 mRNA and TATDN1 mRNA at the ribosome combined stages and input stages after KD of NAT10 or RNPS1. Data are represented as mean ± SD. *p < 0.05, **p < 0.01 and ***p < 0.001 by Two-way ANOVA. (G and H) Polysome profiling (G) and Puromycin assay (H) of the Vector, NAT10- MUT, NAT10-WT, RNPS1-WT, RNPS1-WT & NAT10-MUT and RNPS1-WT & NAT10-WT groups in SCC-4 cells. MMC was added to inhibit the effect of cell proliferation and CBB staining was used as loading control.

**Figure S10.**
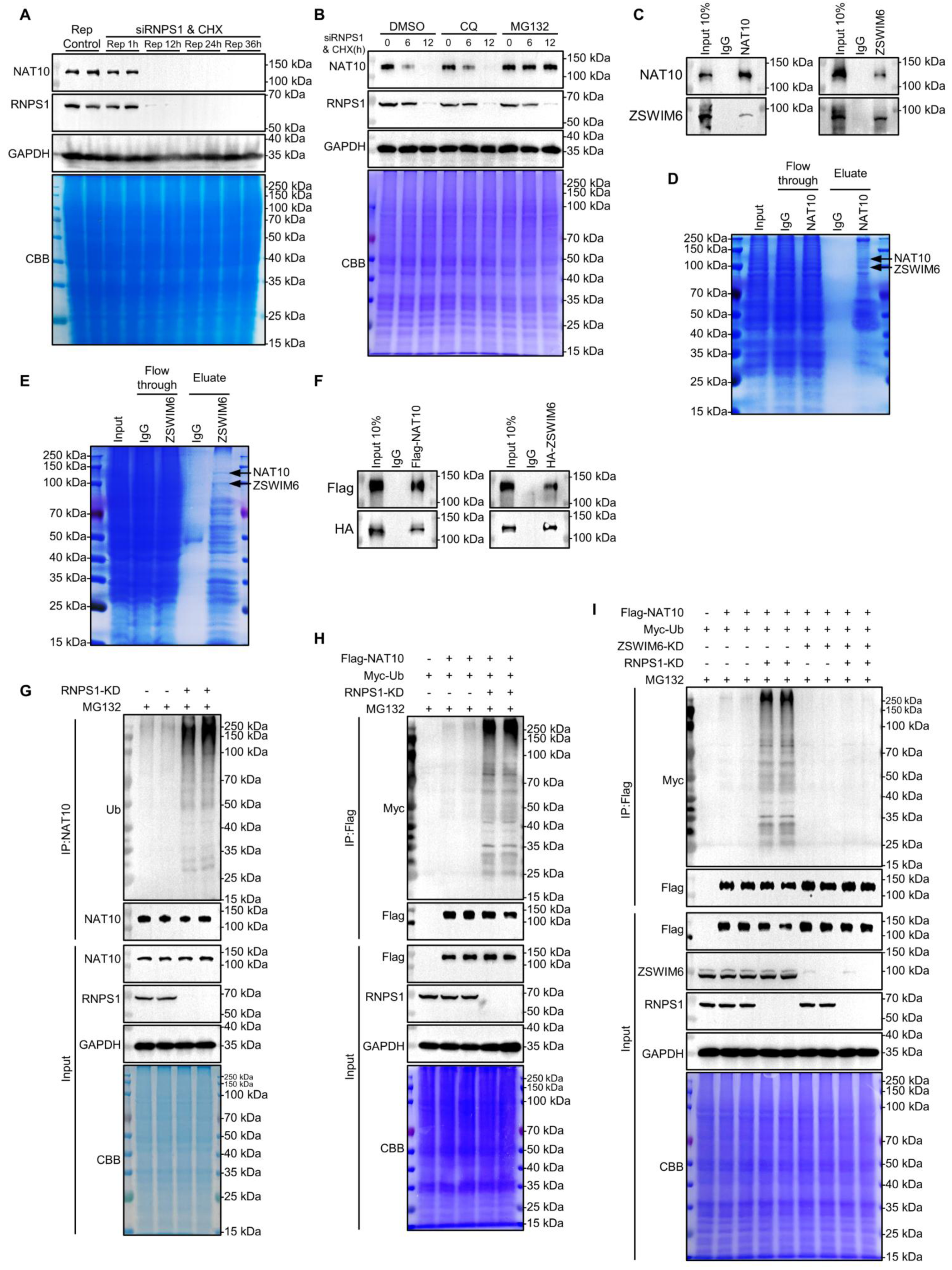
RNPS1 ensures the stability of NAT10 by inhibiting the ubiquitination of NAT10. (A) WB image and analysis showing NAT10 protein expression level in SCC-15 cells for serial RNPS1 knock-down and cycloheximide (CHX, 100 μg/mL) treatment durations. WB of GAPDH and Coomassie brilliant blue staining were used as loading control. (B) WB showing expression level in cells treated with 100 μg/mL cycloheximide (CHX) for the indicated durations in the presence of RNPS1 knock-down with 20 μM chloroquine (CQ) or 10 μM MG132. WB of GAPDH and Coomassie brilliant blue staining were used as loading control. (C) Association of endogenous NAT10 with ZSWIM6 in SCC-15 by co-IP with anti-NAT10 antibody or anti-ZSWIM6 antibody after RNPS1 knock-down and MG132 treatment. Anti-IgG antibody was used as a negative control. (D and E) Protein samples from different steps of protein immunoprecipitation by NAT10 antibody(D), ZSWIM6 antibody (E) and IgG antibody from SCC-15 after RNPS1 knock-down and MG132 treatment were separated by SDS-PAGE and stained with Coomassie brilliant blue. (I) (F) Co-IP assay of Flag-NAT10 and HA-ZSWIM6 in 293T cells, which were transiently co-transfected with Flag-NAT10 and HA-ZSWIM6 plasmids and treated with RNPS1 knock-down and MG132. (II) (G) The WB images of NAT10 ubiquitination in SCC-15 cells after RNPS1 knock-down and MG132 (10 μM) treatment. GAPDH and CBB staining were used as loading control. (III) (H) The ubiquitination level of Flag-NAT10 analyzed by precipitating Flag-NAT10 with an anti-Flag antibody followed by WB using anti-Myc antibody in 293T cells transfected with Flag-NAT10 and Myc-Ub, after RNPS1 knock-down and MG132 (10 μM) treatment as indicated. GAPDH and CBB staining were used as loading control. (I) The WB images shows the Myc-Ubiquitin enrichment level of Flag-NAT10 under different conditions including MG132 (10 μM) treatment, RNPS1 knock-down and/or ZSWIM6 knock-down. GAPDH and CBB staining were used as loading control.

**Figure S11.**
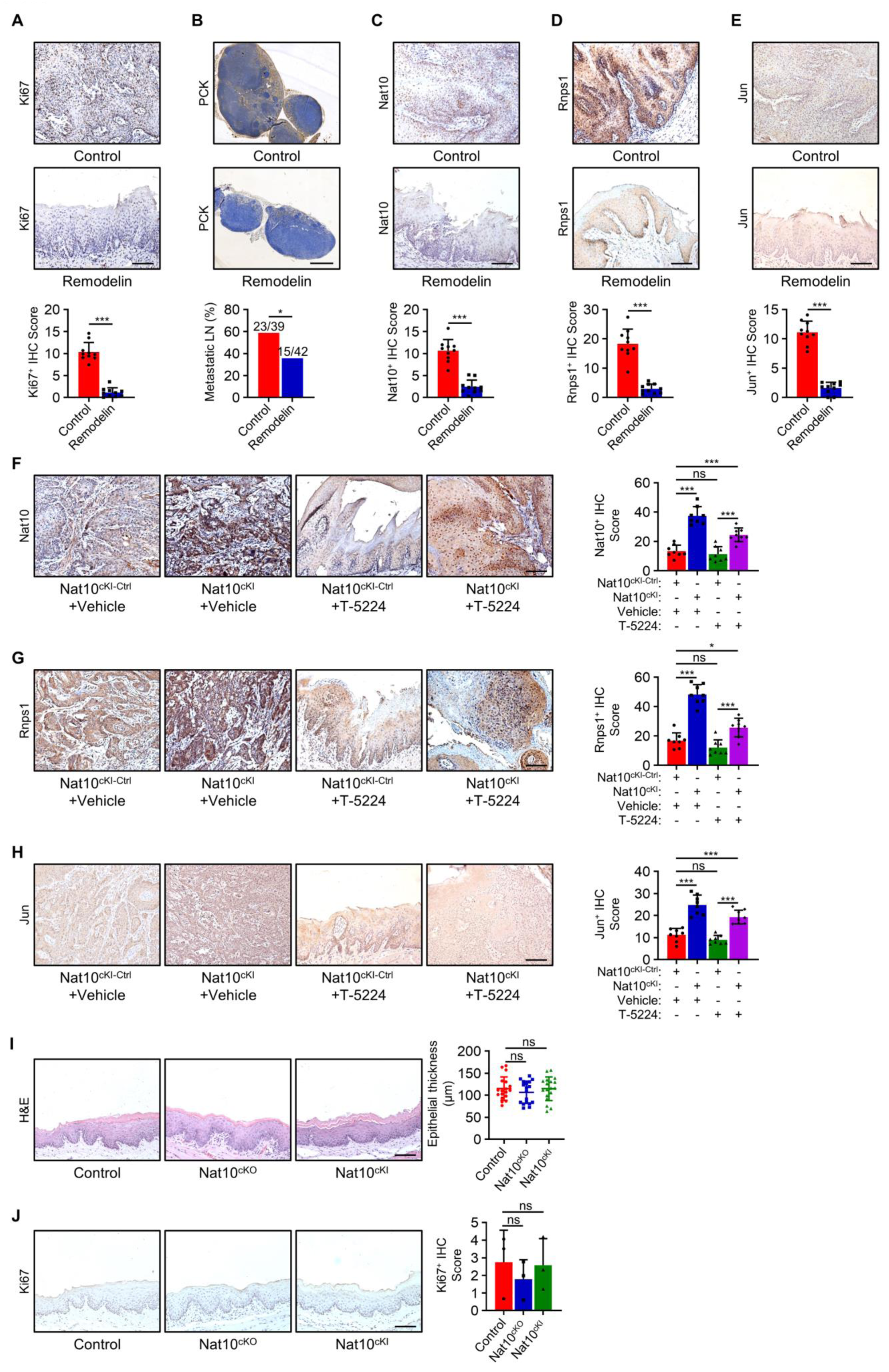
Nat10 drives chemically-induced HNSCC tumorigenesis, related to Figure 6 and Figure 7. (A-E) Representative IHC staining and IHC Score of Ki67 (A), Nat10 (C), Rnps1(D) and Jun (E) in HNSCC and PCK (B) in lymph node. The scale bar of (B) is 500 μm and the rest are 100 μm. n = 10 mice in each group. Data are represented as mean ± SD. *p < 0.05 and ***p < 0.001 in (A) and (C-E) by unpaired Student’s t test. *p < 0.05 in (B) by Chi-square test. (F-H) Representative IHC staining and IHC Score of Nat10 (F), Rnps1(G) and Jun (H) in HNSCC between different treatment groups. Scale bars are 100 μm. Data are represented as mean ± SD. *p < 0.05 and ***p < 0.001 by One-way ANOVA. (I) Representative H&E staining of oral epithelium and quantification of oral epithelial thickness. Mice were not fed with 4NQO drinking water and six random locations of each mouse were selected for epithelial thickness measurement. n = 3 mice for each group. Scale bar, 100 μm. Data are represented as mean ± SD. p values by One-way ANOVA. 1. (J) Representative IHC staining and IHC Score of Ki67 in normal oral epithelium between Control, NAT10-KO and NAT10-KI mice groups. n = 3 mice for each group. Scale bar, 100 μm. Data are represented as mean ± SD. Data are represented as mean ± SD. p values by One-way ANOVA.

## NB replicates

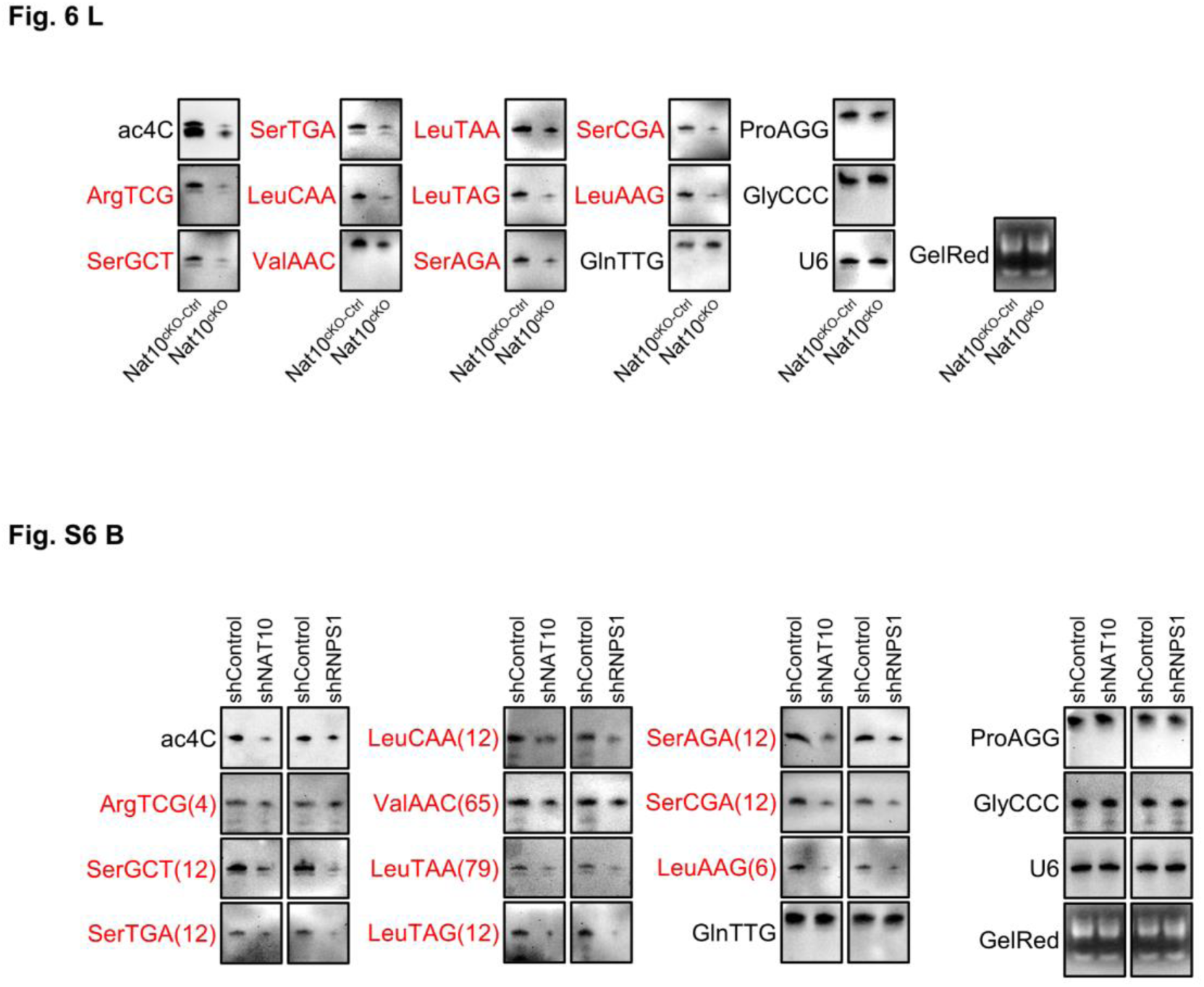

## WB replicates

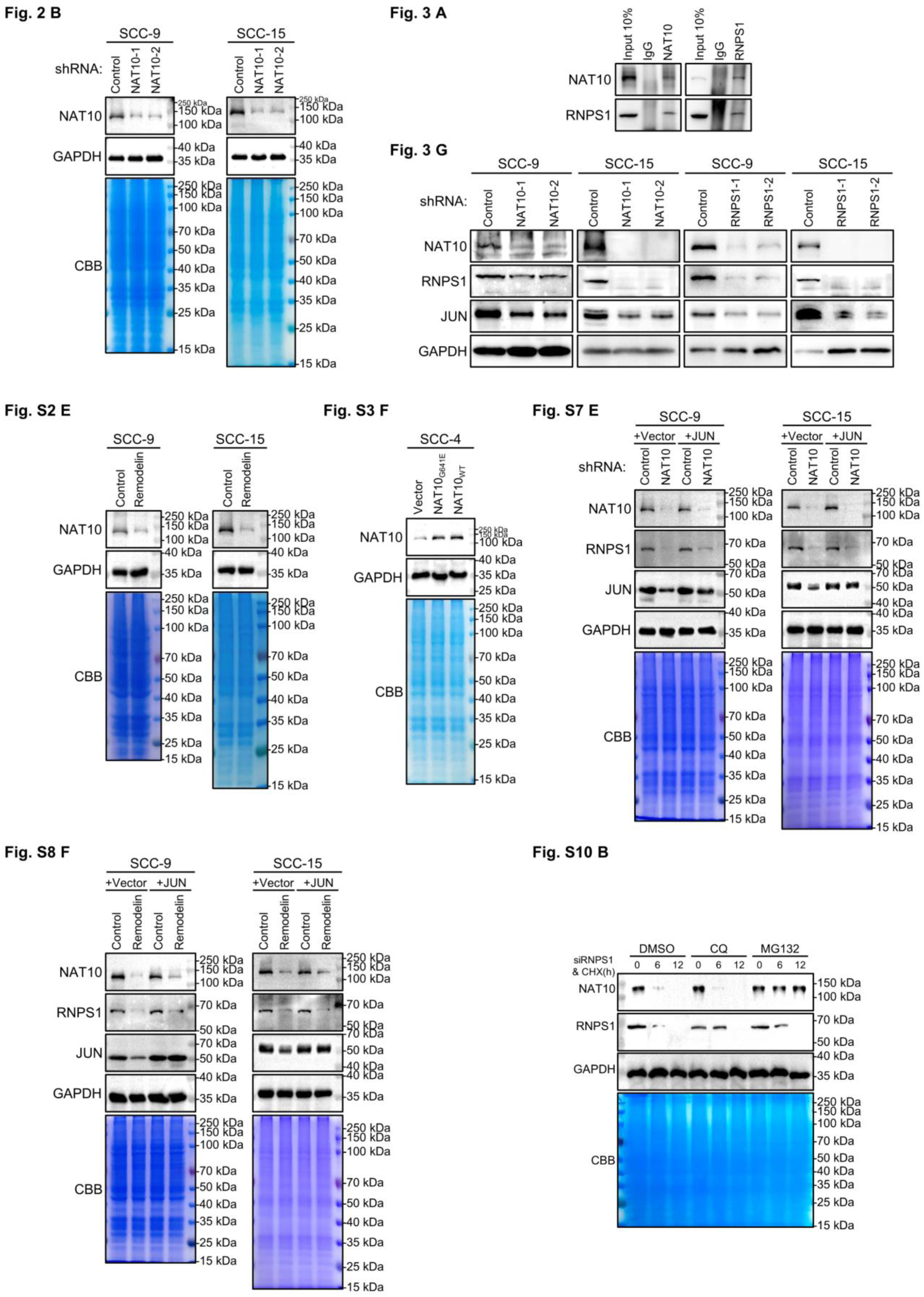

## Acknowledgments

We thank the members of the Lin, Guo, Cai, Zhou, and Zuo labs for reagents, cells, helpful discussions and data-analysis supports. We thank the Hospital of Stomatology, Sun Yat-sen University for providing clinical samples. This work is supported by National Natural Science Foundation of China (82173362 and 81872409), Natural Science Foundation of Guangdong Province (2018A030313610), the Open Funding of the State Key Laboratory of Oral Diseases (SKLOD2021OF02), Guangdong Basic and Applied Basic Research Foundation (2019A1515110110).

## Author Contributions

Conceptualization, X.W., Q.C., C.W., Q.L., and D.C.; Methodology, X.W., K.L., M.C., and J.C.; Data Analysis and Curation, H.X., X.P., K.L., R.L., L.K., Q.L., J.L., L.P., and D.C.; Investigation and Validation, J.W.C., Y.W., R.L., L.K., C.Z., Q.Z., Y.Z., Z.C., F.G., and S.C.; Resources, M.C., J.C., Y.W., L.P., and C.W.; Writing-Original Draft, X.W., K.L., and D.C.; Writing-Review & Editing, J.L., L.P., Q.C., C.W., Q.L., and D.C.; Supervision and funding acquisition, M.C., J.W.C., and D.C.

## Materials and Methods

### Data availability

The raw data including RNC-seq, Ribosome-seq, total RNA reduction and misincorporation sequencing and tRNA reduction and misincorporation sequencing data have been deposited at GSA (accession number HRA002348).

### Lead contact

Further information and requests for resources and reagents should be directed to and will be fulfilled by the Lead Contact, Demeng Chen.

### Materials availability

This study did not generate any new reagents.

### Experimental Model and Subject Details

### Cell culture and generation of mutant cell lines

HOK cells were purchased from ScienCell. SCC-4, SCC-9, SCC-15, SCC-25 and 293T were purchased from the American Type Culture Collection (ATCC) and UM1 was provided by Xiaofeng Zhou. UM1, SCC-4, SCC-9, SCC-15 and SCC-25 were maintained in 1:1 mixture of Dulbecco’s modified Eagle’s medium and Ham’s F12 medium (DMEM/F12, Gibco, C11330500BT). HOK (ScienCell, CP2610) was maintained in Dulbecco’s modified Eagle’s medium (DMEM, Gibco, C11995500BT-1) at a 37°C incubator containing 5% CO_2_. All cell line media were supplemented with 10% fetal bovine serum (FBS, Gibco, 10270-106) and 1% penicillin/streptomycin (Gibco, 15140-122).

For stable KD of NAT10 and RNPS1 in HNSCC cells, we used lentivirus to integrate shRNA into the host cell lines. Briefly, the shRNAs targeting NAT10 or RNPS1 were cloned into pLKO.1 plasmid, and then co-transfected into 293T cells with packaging vector psPAX2 and enveloped vector pMD2.G using Lipofectamine 2000 reagent (Invitrogen, 11668019). Subsequently, the medium supernatant containing lentivirus was collected and added into the medium of SCC-9 or SCC-15 cells with 10 μg/mL Polybrene (YEASEN, 40804ES76). After 48 hours, the positive clones were screened with 2.5 μg/mL puromycin (Beyotime, ST551-250 mg). Similarly, we used retrovirus system to construct the stable overexpression strain of JUN as described previously (*40*). For NAT10_WT_ or NAT10_G641E_ overexpression, Lipofectamine 2000 reagent was used for transfection of SCC-4 with pICE-FLAG-NAT10-siR-WT or pICE-FLAG-NAT10- siR-G641E plasmids. After 24-48 hours, series of experiments were carried out for phenotypic analysis. For mapping of RNPS1 domains that interact with NAT10, we constructed various truncated RNPS1 with HA tag into pcDNA3.1 vector and cotransfected with pICE-FLAG-NAT10-siR-WT into SCC-4 through Lipofectamine 2000 reagent. 24-48 hours later, cell lysate was immunoprecipitated with anti-Flag antibody (Proteintech, 80010-1-RR) and Protein A/G magnetic beads (Thermo Fisher Scientific, 88803). The NAT10-RNPS1 complex was detected with anti-HA antibody after immunoblotting. In the rescue assay of tRNAs overexpression with/without knock down of NAT10, similarly as mentioned above, the plasmids pUC19-tRNA-SerGCT, pUC19-tRNA-SerAGA and pUC19-tRNA-LeuCAA were co-transfected into SCC-9 or SCC-15 cells with Lipofectamine 2000 reagent (Invitrogen, 11668019). For ubiquitination assay, RNPS1-KD and/or ZSWIM6-KD cells transfected with the indicated plasmids pICE-FLAG-NAT10-siR-WT and pCDH-MYC-Ubiquitin were cultured with 10 μM MG-132 (Selleck, S2619) and lysed. The supernatants were subjected to immunoprecipitation and immunoblot analysis with the indicated antibodies as described below. For luciferase reporter assay, pZX001-JUN-WT, pZX001-JUN-MUT1 or pZX001-JUN-MUT2 was transfected into control or NAT10 KD cell lines of SCC-9 or SCC-15 respectively. After incubation at 37 ℃ for 48 h, the luciferase activity was measured and analyzed using Renilla-Firefly Luciferase Dual Assay Kit (MCE, HY-K1013) according to the manufacturer’s instructions.

### Human subjects

HNSCC tissues and adjacent tissues were collected from patients with HNSCC who underwent surgery at the Hospital of Stomatology, Sun Yat-sen University. This study received the informed written consent from all patients before the experiment.

### Animal studies

All animal studies were approved by the Institutional Animal Care and Use Committee, Sun Yat-Sen University (IACUC, SYSU). All animals were housed under specific pathogen-free conditions and handled in the Laboratory Animal Center, Sun Yat-Sen University. The approval number is SYSU-IACUC-2020-000437 and SYSU-IACUC- 2021-000122. *K14CreER* (Stock No: 005107) and *RiboTag* (Stock No: 011029) mice were purchased from Jackson Laboratory. BALB/c-nu/nu (Application No: 20191230- 00047) mice were purchased from the Laboratory Animal Center, Sun Yat-Sen University.

### Generation of Nat10^flox/flox^ mice and Nat10^knockin^ mice

To generate *Nat10* conditional knockout mice, we used CRISPR-Cas9 system. After scanning the gene structure and the size of exons, we found that exon 4 can be conditionally removed and the deletion of exon 4 will result in a 71 aa (66 native aa plus 5 frameshift aa) truncated protein which may be subject to non-sense mediated decay (Fig. 6A). The intron 3 and intron 4 are large in size and insertion of *loxP* element did not interfere mRNA splicing. To minimize the possibility of disruption of Nat10 expression, both *loxP* sites were inserted into non-conserved regions. In addition, Cre-dependent *Nat10* knockin mice were generated by using a CRISPR-Cas9 strategy. Briefly, the synthetic fragment Nat10-CDS (3103bp, digested with AscI and EcoRV) were ligated into mROSA-KI-12p (digested with AscI and EcoRV) to form Nat10 mROSA-KI-12p-A targeting vector (Fig. 7A). Correctly targeted colonies, Cas9-expression mRNAs and sgRNAs were microinjected into C57BL/6N mice oosperm and implanting embryos into the oviducts of pseudopregnant female mice for generating chimeric mice.

### Mouse models for HNSCC

After obtaining the desired mouse, we proceeded to validate the functionality of the floxed allele by crossing with the well-established Cre-expressing mouse strain: the *K14CreER*. For analyzing gene expression and mRNA translation, *K14CreER*, *Nat10^flox/flox^* and *RiboTag* mouse strains were interbred to obtain *K14CreER; RiboTag; Nat10^flox/flox^* mice. To introduce Nat10 conditional overexpression, *K14CreER* and *Nat10^knockin^* mouse strains were cross mated to generate *K14CreER; Nat10^knockin/knockin^* mice. For induction of HNSCC in mice, 6 weeks aged mice were fed with drinking water containing 50 μg/mL 4NQO (Sigma-Aldrich, N8141) for 16 weeks and then given normal drinking water for additional 10 weeks. As for conditional *Nat10* knockout and knockin studies, mice were intraperitoneally injected with tamoxifen (0.08mg/g body weight per day for 3 days; Sigma-Aldrich, T5648-1G). For chemical-treated animals’ assay, Remodelin (MCE or TargetMol, 1622921-15-6) was administered once every two days by oral gavage at 80 mg/kg body weight for 4 weeks. T-5224 (120 mg/kg body weight, TargetMol, 530141-72-1) was injected intraperitoneally every three days for 4 weeks.

### Establishment of orthotopic transplanted tumor model in nude mice

For orthotopic transplant assay, nude mice (BALB/c-nu/nu) maintained under specific pathogen-free conditions were used. Five weeks after birth, the nude mice were anesthetized with 87.5 mg/kg Ketamine/12.5 mg/kg Xylazine intraperitoneally, routinely disinfected, and injected with a total 50 μL mixture of cell suspension and phosphate buffer saline (1:1, approximately 5×10^5^ cancer cells) into the tongue of the mice. The whole operation was completed within 1 hour after the preparation of single cell suspension. After inoculation, the mice were kept in the sterile laminar flow room, and the vitality of the mice were observed for 24 hours to determine whether they were affected with anesthetics.

## Method Details

### RNA extraction and real-time PCR

Cell pellets were collected and then subjected to total RNA extraction using AG RNAex Pro Reagent (Accurate Biotechnology, AG21102) according to the manufacturer’s instructions. Next, in accordance with the instructions, real-time PCR was performed using PerfectStart® Green qPCR SuperMix (TransGen Biotech, 1), by a Bio-Rad CXF96 real-time system (Bio-Rad, USA). Applying an internal control, the relative quantity was calculated.

For the extraction of rRNA, after the cells were washed with pre-chilled PBS, 2 ml of cell lysis buffer (1% Triton™ X-100 (Sigma-Aldrich, X-100) in ribosome buffer) was added. The ribosome buffer consists of 20 mM HEPES-KOH (pH 7.4; BestBio, BB- 1932-100mL), 15 mM MgCl_2_ (Sigma-Aldrich, M8266-1KG), 200mM KCl (Sigma-Aldrich, 746436-500G) and 2 mM dithiothreitol (DTT, BBI Life Sciences, A620058- 0025). After 30 min on ice, the cells were scraped off and transferred to a pre-chilled 1.5 ml tube. Cell debris was removed by centrifuging at 15000 rpm for 10 min at 4℃.

The supernatant was transferred on the surface of 11.5 mL of sucrose buffer (30% sucrose in ribosome buffer) and centrifuged at the speed of 36000 rpm for 4 h at 4℃.

Then discard the supernatant and add AG RNAex Pro Reagent for extraction as described above.

### Total protein extraction and Western blot analysis

Protein lysates were isolated in lysis buffer and followed by the addition of sample loading buffer (4Abio, 4APA008-15), separated by 10% SDS polyacrylamide gel electrophoresis (EpiZyme, PG11X) and transferred onto PVDF membranes (Merck Millipore, IPVH00010). The membranes incubated with primary antibodies against: NAT10 (Santa Cruz, [sc-271770] or Proteintech, [13365-1-AP], 1:2000), RNPS1 (Proteintech, 10555-1-AP, 1:2000), JUN (Proteintech, 24909-1-AP, 1:2000), HA (Abcam, ab9110, 1:4000), GAPDH (Cell Signaling Technology, 2118S, 1:2000). Enhanced chemiluminescence (ECL) method with appropriate species-specific horseradish peroxidase-conjugated secondary antibodies (Proteintech, anti-rabbit [SA00001-2] 1:4000, anti-mouse [SA00001-1] 1:4000) were used to visualize the blots.

### Cell proliferation assay and IC50 measurements

CCK-8 assay (DOJINDO, CK04-500T) was conducted to determine the cell proliferation. According to the manufacturer’s instructions, cell suspension was added into 96-well plate (0.01-0.02 × 10^6^ cells/well). The absorbance at 490 nm was measured after addition of CCK-8 reagent to cells. For IC_50_ measurements, cell lines were treated with vehicle or a range of Remodelin concentrations (1.17, 2.34, 4.69, 9.38, 18.75, 37.5, 75, 150 and 300 μM) for 72 hours. Similarly, CCK-8 reagent was added and measured by microplate reader (TECAN, infinite m200 pro).

### Cell migration assay

Cell migration assay was carried out using Cell Culture Insert (Merck millipore, MCHT06H48). Cells were put into the chamber containing 600mL medium for 4 to 6 hours of culture in cell incubator. And DMEM containing 20% FBS was served as a stimulating factor. Then cells were wiped out carefully and stained with crystal purple (Beyotime, C0121). 1 hour later, the chamber was moved to the empty 24-well plate, followed by an observation under the microscope.

### Dot blot

RNAs including total RNA, rRNA, mRNA and small RNA were extracted and enriched (mRNA was enriched through Poly(A)Purist™ MAG Kit (Invitrogen, AM1922) and small RNA was enriched through MirVana miRNA Isolation Kit (Invitrogen, AM1561)) before dot blots were performed using rabbit monoclonal anti-ac^4^C antibodies (Abcam, ab252215). Briefly, equal amounts of diluted RNAs were denatured at 90℃ for 5 min and 4℃ for 1 min. After that, RNAs were added into an Amersham Hybond N+ membrane (GE Healthcare, RPN203B) and crosslinked twice with 1200 mJ for 25-50s in the Stratalinker 2400 UV Crosslinker (Stratalinker, USA). Membrane was blocked with 5% non-fat milk in 1 × PBST for 1 h at room temperature and then was inducted overnight with anti-ac^4^C antibody in 1% non-fat milk at 4℃. After several washing steps with 0.1% PBST, horseradish peroxidase coupled secondary antibody was applied at 4℃ overnight. Membrane was developed with Chemiluminescent Imaging System (Tanon 5200 SF, China).

### Tumor sphere formation assays

To establish tumor spheres, cells were seeded onto ultra-low attachment 6-well plate at 10,000 cells/mL and cultured seven days in the tumor sphere medium as previously described (*41*). The cells were cultured for seven days, during which serum-free media were changed every other day until the spheres formed. Three dishes were used for each group and all experiments were repeated three times.

### Flow cytometric analysis of apoptosis

Cells were collected, washed, spun down and labelled with fluorescein isothiocyanate (FITC)-Annexin V and propidium iodide using the FITC Annexin V Apoptosis Detection Kit (DOJINDO, AD10). Apoptotic cells were analyzed by using FACS LSRFortessa flow cytometry (BD Biosciences, USA). Graphs show percentage of viable cells, dead cells, early apoptotic cells, and late apoptotic cells.

### Aldehyde dehydrogenase activity assay and flow cytometry

The ALDEFLUOR™ fluorescent reagent system (STEMCELL Technologies, 01700) was used to identify the cells expressing ALDH activity by using FACS LSRFortessa flow cytometer (BD Biosciences, USA). Briefly, cells were digested, suspended and labelled with ALDEFLUOR Stem Cell Identification Kit (STEMCELL Technologies, 01700), following the manufacturer’s instructions. DEAB-treated cells served as control to set the ALDH^high^ regions. Data analysis was performed using the Guava InCyte software. The analysis was performed in triplicate and repeated at least two times.

### Acetylated RNA immunoprecipitation and sequencing

To investigate acetylated RNA regions across the whole transcriptome, acRIP-seq analysis was performed with the 150 μg total RNA from control or the NAT10 depletion groups. Briefly, DNase-treated total RNA was fragmented by the EpiTM RNA Fragmentation buffer for 6 min at 70 ℃, after which EDTA was added immediately to terminate the reaction. Then RNA was purified and collected by the Zymo RNA clean and concentrator-25 kit. For further immunoprecipitation, Anti-ac^4^C or rabbit monoclonal IgG Isotype control were pre-incubated with Protein G Dynabeads (Thermo Fisher Scientific) in PBS for 4-6 hours at 4 ℃. Then RNA fragments were added for immunoprecipitation at 4 ℃ overnight. After that, beads were washed two times in acRIP low salt precipitate buffer and another two times in acRIP high salt precipitate buffer. Then, recycling of RNA was carried out by HiPure cell miRNA Kit. The subsequent inputs and acRIPs libraries construction were completed by EpiTM mini longRNA-seq kit. Bioptic Qsep100 Analyzer was used for the quality control of the libraries. Libraries were multiplexed on the NovaSeq high-throughput sequencing platform and sequenced in PE150-sequencing mode.

### tRNA reduction and misincorporation sequencing (TRMC-seq)

The isolated small RNAs’ other domain modifications were removed by AlkB and AlkB-mut, which quenched with 0.5M EDTA to a final concentration of 5 mM EDTA. Next the AlkB-treated RNAs were recovered by Oligo Clean & Concentrator kit (ZYMO RESEARCH, D4060) and divided into three groups: NaCNBH_3_ (treated with NaCNBH_3_ and without pre-treatment of mild alkali), Deacetylation (treated with NaCNBH_3_ and pre-treatment of mild alkali), Mock (without any treatment). Then the RNAs were precipitated with 2.5 volume cold ethanol at −20 °C for at least four hours and desalted with 75% cold ethanol. After precipitation, the RNAs were ligated with 3’ adaptors, hybridized with reverse transcription primers and ligated with 5’ adaptors. Ligated RNAs were reverse transcribed using TGIRT-III (InGex, TGIRT50) and performed PCR amplification. Subsequently, the products were purified by QIAquick PCR Purification kit (QIAGEN, 28104) and circularized by the splint oligo sequence forming the single strand circle DNA, followed by sequencing with MGISEQ-2000 (Fig. S4K). For total RNA reduction and misincorporation sequencing, we fragmented RNA and skipped the step of small RNA isolation. We then carried out the remaining steps as described in the TRMC-seq (Fig. S4K).

### RNC-seq

The total ribosome bound and unbound mRNAs were separated by gradient centrifugation. The content and quality of RNA were detected by gel electrophoresis and Nano-300 Micro Spectrophotometer. The library preparations were subsequently sequenced on a platform of BGISEQ-500. TR was calculated by dividing the ribosome combined fragments signals by the input RNA-seq signals.

### Polysome assay

For polysome assay, cells were incubated with PBS containing 100 μg/mL cycloheximide (CHX, Sigma-Aldrich, C7698). 15 min later, cells were harvested with polysome cell extraction buffer. We then sperated the supernatant, measured the OD value and added the supernatant on the top of 11mL 10-50% sucrose gradient tube. After centrifugation (36K rpm for 2hrs-2.5hrs at 4℃ with max break), the samples were analyzed and plotted by BR-188 Density Gradient Fractionation System (Brandel, USA).

### Puromycin assay

For puromycin assay, also known as SUnSET assay (*42*), cells were first incubated with puromycin (1 µM final concentration) for 30 min, then harvested for protein extraction. 15 µg protein were loaded onto a SDS polyacrylamide gel for electropherosis. The following steps are carried out like regular Western blot analysis described above. The concentration of the first antibody-anti-Puromycin is 1:2000 (Merck millipore, MABE342).

### Actinomycin D assay

Actinomycin D (MCE, HY-17559) was added into culture medium and incubated with cells for different periods of time (0 hour, 3 hours and 6 hours). The follow-up operation steps are same as above RNA extraction and real-time PCR.

### Northwestern blot (NWB) and Northern blot (NB)

Northern blot was performed as previous described (*43*). Briefly, 2 or 3 µg total RNA samples were mixed with 2 × RNA loading buffer and denatured at 95℃ for 5 min and 4℃ for 1 min. After 15% UREA-PAGE electrophoresis, the RNAs were transferred onto a positive charged nylon membrane mentioned above. The RNAs on the membrane were crosslinked with UV and then blotted with digoxigenin-labeled probes against tRNAs or U6 snRNA. For Northwestern blot, the RNA containing membranes were blotted with anti-ac^4^C antibody overnight. Finally, the digoxigenin or anti-ac^4^C antibody signals were detected following the Western blot protocol described above.

### Liquid chromatography tandem mass spectrometry (LC-MS/MS)

Proteins were first separated using SDS polyacrylamide gels. Gels were then stained with Coomassie blue to visulize proteins bands. We then cut the gel bands with proteins and put them into a clean 1.5 mL centrifuge tube. Gel bands were rinsed with ddH_2_O and decolorized with decolorizing solution (50% acetonitrile (Thermo Fisher Scientific, A998-4L), 25 mM ammonium bicarbonate (Thermo Fisher Scientific, 1066-33-7)) to completely white with acetonitrile and vacuum dried. Next, we added dithiothreitol (DTT, Amresco, 0281-BEJ-100G) to the tube and performed reduction reaction at 56 ℃ for 1 hour. After removing DTT, the reduced protein was alkylated by iodoacetamide (IAM, Sigma-Aldrich, I6125-10G) and incubated at room temperature in dark for 45 minutes. Then ammonium bicarbonate and acetonitrile were successively used for cleaning and decolorization. The dried gel bands were digested with trypsin at 37 ℃ overnight. Subsequently, formic acid (Thermo Fisher Scientific, 64-18-6) was added to stop the digestion reaction and detected by Q Exactive Mass Spectrometer (Thermo scientific).

### Protein digestion and iTRAQ labeling

The proteins were reduced and alkylated according to the method (LC-MS/MS) mentioned above and we purified samples through acetone precipitation. Then Bradford protein assay was used to determine the protein concentration and 50 μg of protein was diluted with 8 M urea in 100 mM TEAB. Trypsin was used for protein digestion. The samples were digested overnight at 37 °C. After that, peptides were desalted and vacuum-dried, according to the manufacturer’s protocol. Then the peptides were labeled according to the instructions provided by iTRAQ Reagents-8plex kit (AB SCIEX, 4390812).

### Mass spectrometry of ac^4^C RNAs

RNA samples were extracted as previously shown. Add 1 μg of RNA samples into the buffer solution, completely enzymolysis the samples into nucleosides at 37 ℃ under the action of phosphodiesterase (0.002 U/μL; Sigma-Aldrich, P3243), S1 nuclease (180 U/μL; Takara, 2410A) and alkaline phosphatase (30 U/μL; Takara, 2250A), and re extract the enzymatically hydrolyzed sample with chloroform-method. Put the obtained upper aqueous solution into the injection bottle and analyze it via LC-MS/MS. The chromatographic column mainly adopts Waters ACQUITY UPLC HSS T3 C18 column (1.8 μm, 100 mm × 2.1 mm i.d.). At 40 °C, the column flow rate was set to 0.3 mL/min. For mass spectrum, the temperature of the electrospray ion source was set to 550 °C, and the mass spectrum voltage is set to 5500 V under positive electrospray ionization mode.

### Histologic evaluation and immunohistochemical staining

For hematoxylin and eosin (H&E) staining, tissues were fixed in formalin, paraffin-embedded, sectioned (5 μm), deparaffinized and stained with H&E staining kit (Solarbio, G1120-3). For IHC staining, sections were deparaffinized and treated with 3% H_2_O_2_ in water for 10 min. Antigen retrieval procedure was conducted to sections with 10 mM citrate buffer (pH 6.0) for 10 min. Next, tissue sections were incubated with the Biocare blocking reagent for 10 min, followed by an overnight incubation at 4℃ with anti-NAT10 (Santa Cruz, sc-271770 1:200), anti-RNPS1 (Proteintech, 10555- 1-AP, 1:200), anti-JUN (Proteintech, 24909-1-AP, 1:200), anti-Ki67 (Novus, NB500-170, 1:200), anti-PCK (pan-Cytokeratin, Santa Cruz, sc-8018, 1:200). Slides were then incubated with goat anti-rabbit horseradish peroxidase-conjugated secondary antibodies for 30 min at room temperature, treated with 3,3′-diami-nobenzidine and counter stained with hematoxylin. For analysis, the staining intensity was scored and the percentage of positive stained areas of tumor cells per the whole tumor area were calculated.

### Immunofluorescence staining

Mouse HNSCC tumors were dissected and fixed with 4% paraformaldehyde in PBS overnight. Samples were then rinsed with cold PBS, equilibrated in 30% sucrose in PBS overnight and embedded in OCT (Tissue Tek, 25608-930). Tissue sections or cell climbing sheets were stained with the following primary antibody: anti-Cytokeratin 14 (Abcam, ab7800; 1:100) & anti-HA tag (Abcam, ab9110; 1:100) or anti-NAT10 (Santa Cruz, sc-271770 1:100) & anti-RNPS1 (Proteintech, 10555-1-AP, 1:100) after permeated with 1% Triton™ X-100 (Sigma-Aldrich, X-100). Then, the antigens were visualized by the corresponding secondary antibody conjugated with DyLight 488 and Fluor 594. The nuclei were counterstained with DAPI (Solarbio, C0065) at 1:1000 for 1 minute. Images were captured with an upright fluorescence microscope (ZEISS, Germany).

## Quantification and Statistical Analysis

### Quantification of TRMC-seq

SRNAtools (*44*) was used to infer tRNA expressions. In brief, after trimming the adapter and filtering low-quality sequences, the clean sequencing reads were mapped to a reference genome and different small RNA libraries using Bowtie (*45*) with a maximum of two mismatch. The mapped reads are used to identify and profile tRNAs. DEseq (*46*) was then used to infer the statistical significance of differential expression of tRNAs. As the default, a tRNA is considered to be significantly differentially expressed when the P value is ≤0.05 and the fold change is at least 1.5-fold.

### Analysis of Ribo-seq

RiboToolkit (RiboToolkit: an integrated platform for analysis and annotation of ribosome profiling data to decode mRNA translation at codon resolution) was used to analyze the codon occupancy based on Ribo-seq. In the analyzing process, the clean Ribo-seq sequences were first aligned to rRNAs, tRNA and snRNA to exclude the RPFs coming from rRNA, tRNA and snRNA using Bowtie (*45*) with a maximum of two mismatches. Cleaned RPF sequences were then mapped to the reference genome using STAR (*47*). The unique genome-mapped RPFs are then mapped against protein coding transcripts using Bowtie. For codon-based analyses, 5’ mapped sites of RPFs (26-32 nt) translated in 0-frame were used to infer the P-sites and the occupancy on each codon was calculated. The codon occupancy was further normalized by the basal occupancy which was calculated as the average occupancy of +1, +2 and +3 position downstream of A-sites. TE was calculated by dividing the ribosome protected fragments (RPF) signals by the input RNA-seq signals.

### Quantification of codon usage for ac^4^C tRNAs

For the calculation of the score of codon usage of acetylated tRNAs-UUU in each gene, the range method ((observed value - minimum value) / range) was used to normalize the codon usage obtained from the Codon Usage Database (http://www.kazusa.or.jp/codon/) of ten ac^4^C tRNAs-UUU to get the δ (value range 0∼1). According to the data scaling method (https://sebastianraschka.com/Articles/2014_about_feature_scaling.html), the exponent 10δ of the natural constant e is set for each codon. Then multiply by the usage rate γ of the codon in each gene, as shown in the following formula to get the first level score m of each gene. Finally, the z-score method was used to standardize all m, and the score of codon usage for each gene containing ac^4^C tRNAs-UUU was obtained.

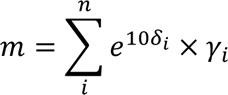

### Other statistical analyses

Statistical analyses were performed using GraphPad Prism 8 software. All data were presented as mean ± SD unless otherwise specified. Statistical significance was determined by p < 0.05. For statistical comparisons, unpaired Student’s T test (and nonparametric test for two groups), one-way ANOVA analyses (three groups or more groups), two-way ANOVA analyses (three groups or more groups for two factors), Pearson chi-square test (metastatic lymph node for transgenic mice) and Mantel-Haenszel chi-square test (tumor invasion for transgenic mice) were performed. Significance was defined *p < 0.05, **p < 0.01, ***p < 0.001.

## Refrences

1. R. Su et al., Targeting FTO Suppresses Cancer Stem Cell Maintenance and Immune Evasion. Cancer Cell 38, 79–96 e11 (2020).

2. E. Yankova et al., Small-molecule inhibition of METTL3 as a strategy against myeloid leukaemia. Nature 593, 597–601 (2021).

3. P. Boccaletto et al., MODOMICS: a database of RNA modification pathways. 2017 update. Nucleic Acids Res 46, D303–D307 (2018).

4. S. Chimnaronk et al., RNA helicase module in an acetyltransferase that modifies a specific tRNA anticodon. EMBO J 28, 1362–1373 (2009).

5. J. M. Thomas et al., A Chemical Signature for Cytidine Acetylation in RNA. J Am Chem Soc 140, 12667–12670 (2018).

6. S. Ito et al., Human NAT10 is an ATP-dependent RNA acetyltransferase responsible for N4-acetylcytidine formation in 18 S ribosomal RNA (rRNA). J Biol Chem 289, 35724–35730 (2014).

7. D. Arango et al., Acetylation of Cytidine in mRNA Promotes Translation Efficiency. Cell 175, 1872–1886 e1824 (2018).

8. Y. Zhang et al., NAT10 promotes gastric cancer metastasis via N4-acetylated COL5A1. Signal Transduct Target Ther 6, 173 (2021).

9. G. Guo et al., Epitranscriptomic N4-Acetylcytidine Profiling in CD4(+) T Cells of Systemic Lupus Erythematosus. Front Cell Dev Biol 8, 842 (2020).

10. A. Sas-Chen et al., Dynamic RNA acetylation revealed by quantitative cross-evolutionary mapping. Nature 583, 638–643 (2020).

11. R. L. Siegel, K. D. Miller, H. E. Fuchs, A. Jemal, Cancer Statistics, 2021. CA Cancer J Clin 71, 7–33 (2021).

12. L. Liu et al., METTL3 Promotes Tumorigenesis and Metastasis through BMI1 m(6)A Methylation in Oral Squamous Cell Carcinoma. Mol Ther 28, 2177–2190 (2020).

13. S. Jin et al., The m6A demethylase ALKBH5 promotes tumor progression by inhibiting RIG-I expression and interferon alpha production through the IKKepsilon/TBK1/IRF3 pathway in head and neck squamous cell carcinoma. Mol Cancer 21, 97 (2022).

14. J. Chen et al., Aberrant translation regulated by METTL1/WDR4-mediated tRNA N7- methylguanosine modification drives head and neck squamous cell carcinoma progression. Cancer Commun (Lond) 42, 223–244 (2022).

15. D. Larrieu, S. Britton, M. Demir, R. Rodriguez, S. P. Jackson, Chemical inhibition of NAT10 corrects defects of laminopathic cells. Science 344, 527–532 (2014).

16. S. Sharma et al., Yeast Kre33 and human NAT10 are conserved 18S rRNA cytosine acetyltransferases that modify tRNAs assisted by the adaptor Tan1/THUMPD1. Nucleic Acids Res 43, 2242–2258 (2015).

17. X. Liu et al., NAT10 regulates p53 activation through acetylating p53 at K120 and ubiquitinating Mdm2. EMBO Rep 17, 349–366 (2016).

18. Q. Li et al., NAT10 is upregulated in hepatocellular carcinoma and enhances mutant p53 activity. BMC Cancer 17, 605 (2017).

19. D. Dominissini, G. Rechavi, N(4)-acetylation of Cytidine in mRNA by NAT10 Regulates Stability and Translation. Cell 175, 1725–1727 (2018).

20. L. Kotelawala, E. J. Grayhack, E. M. Phizicky, Identification of yeast tRNA Um(44) 2’-O- methyltransferase (Trm44) and demonstration of a Trm44 role in sustaining levels of specific tRNA(Ser) species. RNA 14, 158–169 (2008).

21. S. Thalalla Gamage, A. Sas-Chen, S. Schwartz, J. L. Meier, Quantitative nucleotide resolution profiling of RNA cytidine acetylation by ac4C-seq. Nat Protoc 16, 2286–2307 (2021).

22. J. W. Mabin et al., The Exon Junction Complex Undergoes a Compositional Switch that Alters mRNP Structure and Nonsense-Mediated mRNA Decay Activity. Cell Rep 25, 2431–2446 e2437 (2018).

23. E. E. Palmer et al., A Recurrent De Novo Nonsense Variant in ZSWIM6 Results in Severe Intellectual Disability without Frontonasal or Limb Malformations. Am J Hum Genet 101, 995–1005 (2017).

24. D. J. Tischfield et al., Loss of the neurodevelopmental gene Zswim6 alters striatal morphology and motor regulation. Neurobiol Dis 103, 174–183 (2017).

25. E. Sanz et al., Cell-type-specific isolation of ribosome-associated mRNA from complex tissues. Proc Natl Acad Sci U S A 106, 13939–13944 (2009).

26. I. Barbieri, T. Kouzarides, Role of RNA modifications in cancer. Nat Rev Cancer 20, 303–322 (2020).

27. P. Nombela, B. Miguel-Lopez, S. Blanco, The role of m(6)A, m(5)C and Psi RNA modifications in cancer: Novel therapeutic opportunities. Mol Cancer 20, 18 (2021).

28. L. Yi, G. Wu, L. Guo, X. Zou, P. Huang, Comprehensive Analysis of the PD-L1 and Immune Infiltrates of m(6)A RNA Methylation Regulators in Head and Neck Squamous Cell Carcinoma. Mol Ther Nucleic Acids 21, 299–314 (2020).

29. Y. Huang et al., Small-Molecule Targeting of Oncogenic FTO Demethylase in Acute Myeloid Leukemia. Cancer Cell 35, 677–691 e610 (2019).

30. T. Suzuki, The expanding world of tRNA modifications and their disease relevance. Nat Rev Mol Cell Biol 22, 375–392 (2021).

31. J. M. Thomas, K. M. Bryson, J. L. Meier, Nucleotide resolution sequencing of N4- acetylcytidine in RNA. Methods Enzymol 621, 31–51 (2019).

32. G. Kawai, T. Hashizume, T. Miyazawa, J. A. McCloskey, S. Yokoyama, Conformational characteristics of 4-acetylcytidine found in tRNA. Nucleic Acids Symp Ser, 61–62 (1989).

33. Q. Xu et al., The interaction of interleukin-8 and PTEN inactivation promotes the malignant progression of head and neck squamous cell carcinoma via the STAT3 pathway. Cell Death Dis 11, 405 (2020).

34. T. A. Karakasheva et al., IL-6 Mediates Cross-Talk between Tumor Cells and Activated Fibroblasts in the Tumor Microenvironment. Cancer Res 78, 4957–4970 (2018).

35. C. Georganas et al., Regulation of IL-6 and IL-8 expression in rheumatoid arthritis synovial fibroblasts: the dominant role for NF-kappa B but not C/EBP beta or c-Jun. J Immunol 165, 7199–7206 (2000).

36. X. Ding et al., Epigenetic activation of AP1 promotes squamous cell carcinoma metastasis. Sci Signal 6, ra28 21-13, S20-15 (2013).

37. I. A. Roundtree, M. E. Evans, T. Pan, C. He, Dynamic RNA Modifications in Gene Expression Regulation. Cell 169, 1187–1200 (2017).

38. W. Tao et al., NAT10 as a potential prognostic biomarker and therapeutic target for HNSCC. Cancer Cell Int 21, 413 (2021).

39. J. Wu, H. Zhu, J. Wu, W. Chen, X. Guan, Inhibition of N-acetyltransferase 10 using remodelin attenuates doxorubicin resistance by reversing the epithelial-mesenchymal transition in breast cancer. Am J Transl Res 10, 256–264 (2018).

40. B. K. Park et al., NF-kappaB in breast cancer cells promotes osteolytic bone metastasis by inducing osteoclastogenesis via GM-CSF. Nat Med 13, 62–69 (2007).

41. D. Chen et al., Targeting BMI1(+) Cancer Stem Cells Overcomes Chemoresistance and Inhibits Metastases in Squamous Cell Carcinoma. Cell Stem Cell 20, 621–634 e626 (2017).

42. E. K. Schmidt, G. Clavarino, M. Ceppi, P. Pierre, SUnSET, a nonradioactive method to monitor protein synthesis. Nat Methods 6, 275–277 (2009).

43. S. Lin et al., Mettl1/Wdr4-Mediated m(7)G tRNA Methylome Is Required for Normal mRNA Translation and Embryonic Stem Cell Self-Renewal and Differentiation. Mol Cell 71, 244–255 e245 (2018).

44. Q. Liu et al., Small noncoding RNA discovery and profiling with sRNAtools based on high-throughput sequencing. Brief Bioinform 22, 463–473 (2021).

45. B. Langmead, C. Trapnell, M. Pop, S. L. Salzberg, Ultrafast and memory-efficient alignment of short DNA sequences to the human genome. Genome Biol 10, R25 (2009).

46. S. Anders, W. Huber, Differential expression analysis for sequence count data. Genome Biol 11, R106 (2010).

47. A. Dobin et al., STAR: ultrafast universal RNA-seq aligner. Bioinformatics 29, 15–21 (2013).

