## Supplemental Figure 1-11 and Supplemental Table 1-7 for "RNPS1 stabilizes NAT10 protein to facilitate translation in cancer via tRNA ac^4^C modification"

**Figure S1**

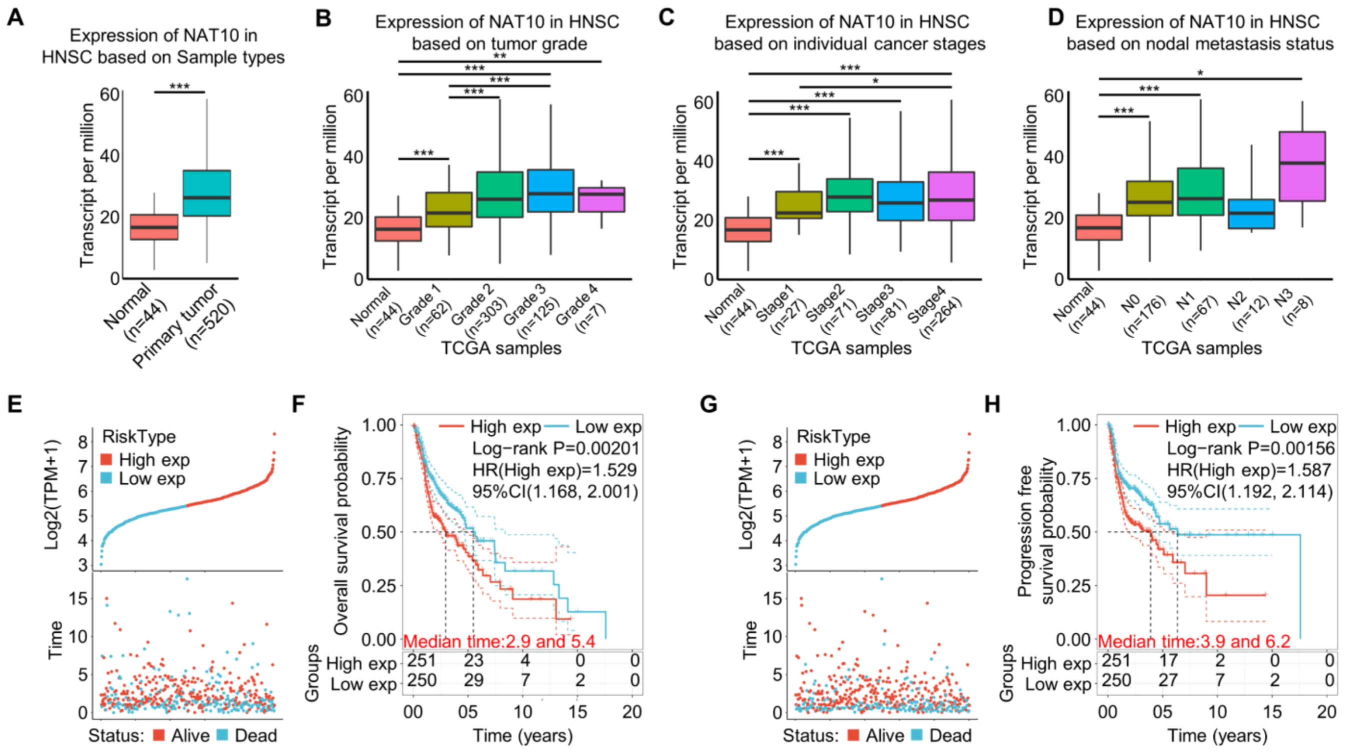


**Figure S1. NAT10 is elevated in HNSCC and correlates with HNSCC progression using TCGA datasets, related to Figure 1**

(A) Expression of NAT10 between normal (n = 44) and primary tumor (n = 520) of TCGA data. ***p < 0.001 by unpaired Student's t test.

(B) The expression of NAT10 increased with the degree of HNSCC tumor grade in TCGA dataset. Grade 1: well differentiated (low grade); Grade 2: moderately differentiated (intermediate grade); Grade 3: poorly differentiated (high grade); Grade 4: Undifferentiated (high grade). Normal, n = 44; Grade 1, n = 62; Grade 2, n = 303; Grade 3, n = 125; Grade 4, n = 7. **p < 0.01 and ***p < 0.001 by One-way ANOVA.

(C) Box plot showing the correlation between NAT10 expression and HNSCC individual cancer stages of TCGA dataset. Normal, n = 44; Stage1, n = 27; Stage2, n = 71; Stage3, n = 81; Stage4, n = 264. *p < 0.05 and ***p < 0.001 by One-way ANOVA.

(D) The correlation between NAT10 expression and HNSCC nodal metastasis status of TCGA dataset. N0: no regional lymph node metastasis; N1: metastases in 1 to 3 axillary lymph nodes; N2: metastases in 4 to 9 axillary lymph nodes; N3: metastases in 10 or more axillary lymph nodes. Normal, n = 44; N0, n = 176; N1, n = 67; N2, n = 12; N3, n = 8 mice in each group. *$p < 0.05$ and ***$p < 0.001$ by One-way ANOVA.

(E) The correlation between NAT10 expression and overall survival time or overall survival status in TCGA dataset. The top represents the scatter plot of the gene expression from low to high, and different colors represent different expression groups; the bottom represents the distribution of survival time and survival status corresponding to gene expression of different samples.

(F) The Kaplan-Meier survival curve distribution of NAT10 expression and overall survival probability in TCGA dataset was tested by log rank. HR (high exp) represents the risk coefficient of the high expression group relative to the low expression group. If HR > 1 represents the risk factor of the gene, and if HR < 1 represents the protective factor of the gene; 95% Cl represents HR confidence interval; Median time represents the time when the survival rate of high expression group or low expression group is 50%. $p = 0.00201$ by Log-rank test.

(G) NAT10 expression, progression free survival time and progression free survival status in TCGA dataset.

(H) The Kaplan-Meier survival curve distribution of NAT10 expression and progression free survival probability in TCGA dataset. $p = 0.00156$ by Log-rank test.

**Figure S2**

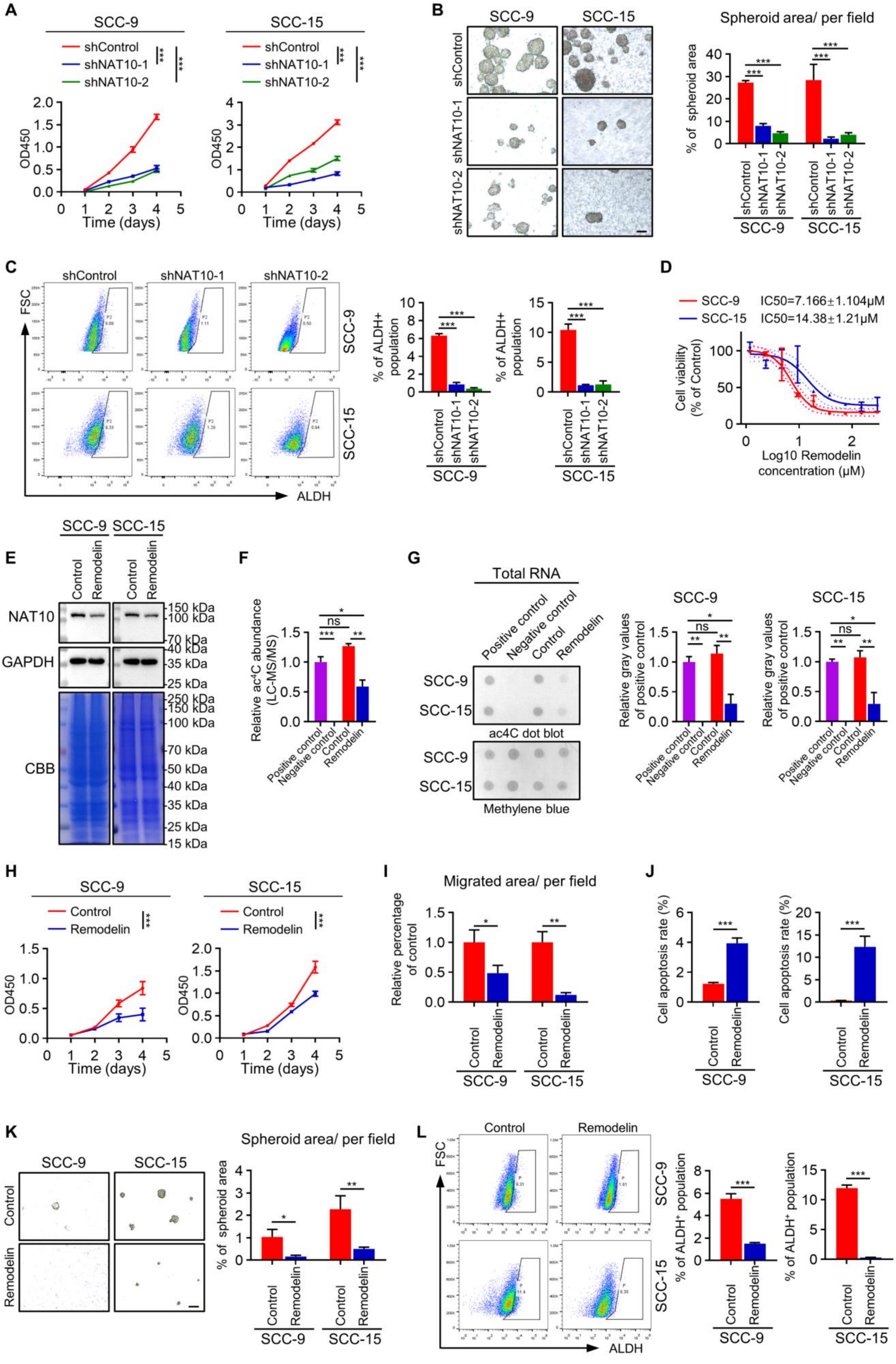
A
SCC-9
SCC-15
shControl
shNAT10-1
shNAT10-2
OD450
Time (days)
B
SCC-9
SCC-15
shControl
shNAT10-1
shNAT10-2
Spheroid area/ per field
% of spheroid area
C
shControl
shNAT10-1
shNAT10-2
FSC
ALDH
SCC-9
SCC-15
% of ALDH+ population
D
SCC-9 IC50=7.166±1.104μM
SCC-15 IC50=14.38±1.21μM
Cell viability (% of Control)
Log10 Remodelin concentration (μM)
E
SCC-9 SCC-15
Control
Remodelin
NAT10
GAPDH
CBB
150 kDa
100 kDa
70 kDa
40 kDa
35 kDa
25 kDa
250 kDa
150 kDa
100 kDa
70 kDa
50 kDa
40 kDa
35 kDa
25 kDa
15 kDa
F
Relative ac4C abundance (LC-MS/MS)
Positive control
Negative control
Control
Remodelin
G
Total RNA
Positive control
Negative control
Control
Remodelin
SCC-9
SCC-15
ac4C dot blot
Methylene blue
Relative gray values of positive control
H
SCC-9
SCC-15
Control
Remodelin
OD450
Time (days)
I
Migrated area/ per field
Relative percentage of control
Control
Remodelin
SCC-9
SCC-15
J
Cell apoptosis rate (%)
Control
Remodelin
SCC-9
SCC-15
K
SCC-9
SCC-15
Control
Remodelin
Spheroid area/ per field
% of spheroid area
L
Control
Remodelin
FSC
ALDH
SCC-9
SCC-15
% of ALDH+ population

**Figure S2. NAT10 plays a vital role in HNSCC progression and metastasis, related to Figure 2**

(A-C) Proliferation (A), sphere formation (B) and aldehyde dehydrogenase activity (C) between the shControl, shNAT10-1 and shNAT10-2 in HNSCC cell lines. The scale bar of (B) is 100 μm. Data are represented as mean ± SD. ***p < 0.001 in (A) by Two-way ANOVA. ***p < 0.001 in (B) and (C) by One-way ANOVA.

(D) $IC_{50}$ values for Remodelin in cytotoxicity assays with HNSCC cell lines. SCC-9, $IC_{50}$ ± SE = 7.166 ± 1.104 μM; SCC-15, $IC_{50}$ ± SE = 14.38 ± 1.21 μM. Data are represented as mean ± SD.

(E) The expression of NAT10 in SCC-9 and SCC-15 after with/without Remodelin treatment. Cells were treated with Remodelin at a concentration of 8 μM (SCC-9) or 15 μM (SCC-15) for 24-48 h. GAPDH and CBB staining were used as loading control.

(F) The bar chart of relative $ac^4C$ abundance of RNA by LC-MS/MS from SCC-15 cells. Positive control: ribosomal RNA; Negative control: poly(A)-enriched RNA; The remaining samples are total RNA from Control (DMSO) and Remodelin groups. Cells were treated with Remodelin at a concentration of 8 μM (SCC-9) or 15 μM (SCC-15) for 24-48 h. n = 2 independent biological replicates. Data are represented as mean ± SD. *p < 0.05, **p < 0.01 and ***p < 0.001 by One-way ANOVA.

(G) Dot blot (left) and relative gray values (right) showed the $ac^4C$ level of RNA in positive control, negative control, Control, Remodelin groups with methylene blue staining as loading control. Cells were treated with Remodelin at a concentration of 8 μM (SCC-9) or 15 μM (SCC-15) for 24-48 h. n = 2 independent biological replicates. Data are represented as mean ± SD. *p < 0.05 and **p < 0.01 by One-way ANOVA.

(H-L) Remodelin reduced the proliferation ability (H), migration (I), sphere formation ability (K), aldehyde dehydrogenase activity (L) and increased rate of cell apoptosis (J) of HNSCC cell lines. Cells were treated with Remodelin at a concentration of 8 μM (SCC-9) or 15 μM (SCC-15) for 4 d (H), 48 h (I, J and L) or 3-4 d (K), respectively. The scale bar of (K) is 100 μm. Data are represented as mean ± SD. ***p < 0.001 in (H) by Two-way ANOVA. *p < 0.05, **p < 0.01 and ***p < 0.001 in (I-L) by unpaired Student's t test.

**Figure S3**

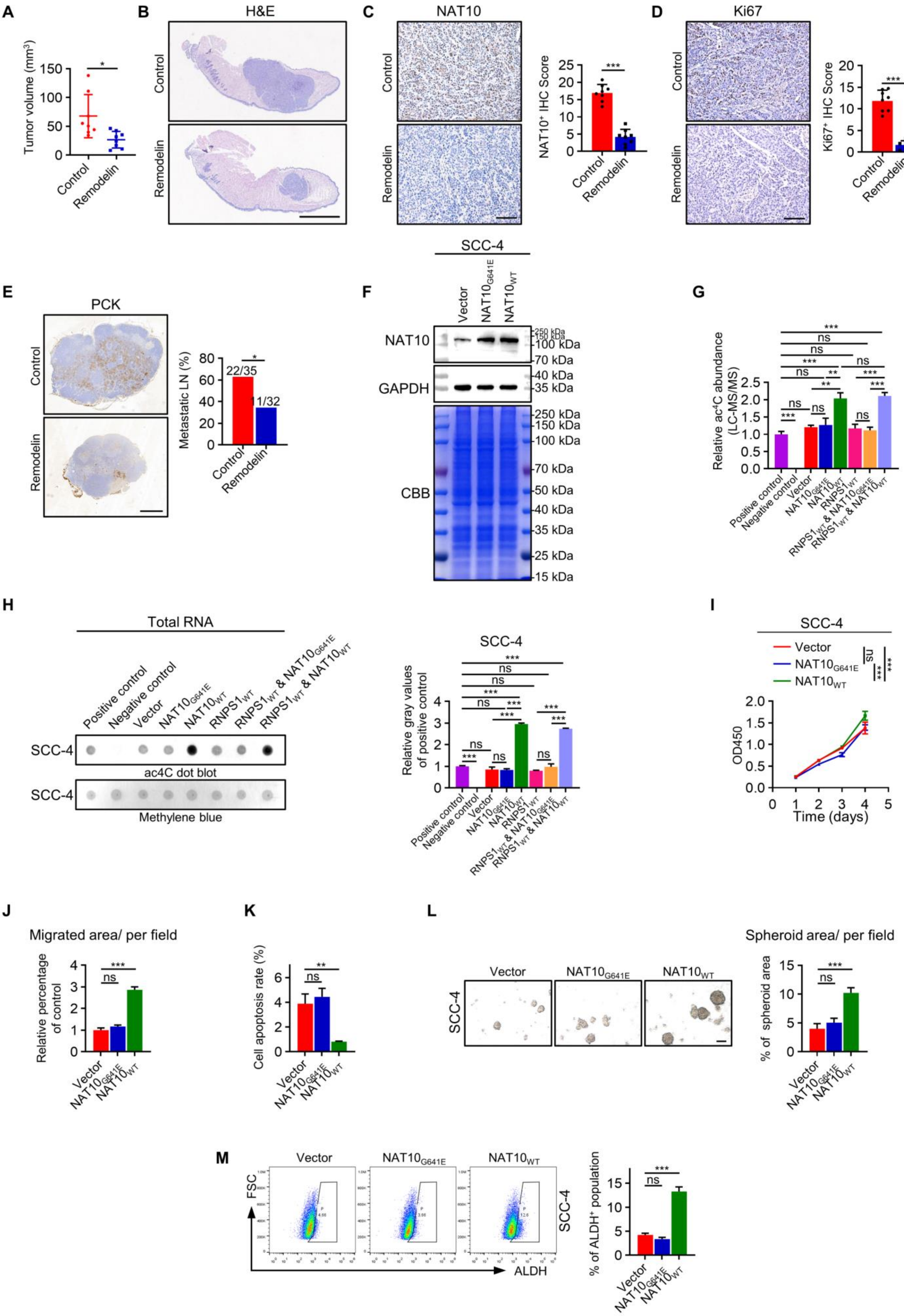
A
Tumor volume (mm3)
Control
Remodelin
B
H&E
C
NAT10
NAT10+ IHC Score
D
Ki67
Ki67+ IHC Score
E
PCK
Metastatic LN (%)
22/35
11/32
F
SCC-4
Vector
NAT10G641E
NAT10WT
NAT10
GAPDH
CBB
G
Relative ac4C abundance (LC-MS/MS)
H
Total RNA
Positive control
Negative control
RNPS1WT
ac4C dot blot
Methylene blue
Relative gray values of positive control
I
OD450
Time (days)
J
Migrated area/ per field
Relative percentage of control
K
Cell apoptosis rate (%)
L
Spheroid area/ per field
% of spheroid area
M
FSC
ALDH
% of ALDH+ population

**Figure S3. NAT10 plays a vital role in HNSCC progression and metastasis, related to Figure 2**

(A) Tumor volume ($mm^3$) of orthotopic transplantation tumor between controlled group and treated group. n = 8 mice in each group. Data are represented as mean ± SD. *p < 0.05 by unpaired Student's t test

(B-E) H&E staining of tongue (B) with orthotopic transplantation tumor and IHC staining with score of NAT10 (C), Ki67 (D) in orthotopic transplantation tumor and PCK (E) in lymph node between Control and Remodelin groups. The scale bar represents 2mm in panel (B), 500 µm in panel (E) and 100 µm for the rest. n = 8 mice in each group. Data are represented as mean ± SD in (C) and (D). ***p < 0.001 in (C) and (D) by unpaired Student's t test. *p < 0.05 in (E) by Chi-square test.

(F) WB showed the protein expression of NAT10 when $NAT10_{WT}$ or $NAT10_{G641E}$ was exogenously overexpressed in SCC-4. GAPDH and CBB staining were used as loading control.

(G) The relative $ac^4C$ level of RNA detected by LC-MS/MS in Positive control, Negative control, Vector, NAT10-MUT, NAT10-WT, RNPS1-WT, RNPS1-WT & NAT10-MUT and RNPS1-WT & NAT10-WT groups of SCC-4 cells. Positive control: ribosomal RNA; Negative control: poly(A)-enriched RNA; The remaining samples are total RNA from Vector, NAT10-MUT, NAT10-WT, RNPS1-WT, RNPS1-WT & NAT10-MUT and RNPS1-WT & NAT10-WT groups. n = 2 independent biological replicates. Data are represented as mean ± SD. **p < 0.01 and ***p < 0.001 by One-way ANOVA.

(H) Dot blot (left) and relative gray values (right) showed the $ac^4C$ modification level of RNA in positive control, negative control, Vector, NAT10-MUT, NAT10-WT, RNPS1-WT, RNPS1-WT & NAT10-MUT and RNPS1-WT & NAT10-WT groups with methylene blue staining as loading control. n = 2 independent biological replicates. Data are represented as mean ± SD. ***p < 0.001 by One-way ANOVA.

(I-M) The cell phenotype assay including proliferation ability (I), migration (J), cell apoptosis (K), sphere formation (L) and aldehyde dehydrogenase activity (M) of SCC-4 cell line with overexpression of Vector, $NAT10_{WT}$ or $NAT10_{G641E}$. The scale bar of (L) is 100 μm. Data are represented as mean ± SD. ***$p < 0.001$ in (I) by Two-way ANOVA. **$p < 0.01$ and ***$p < 0.001$ in (J-M) by One-way ANOVA.

**Figure S4**

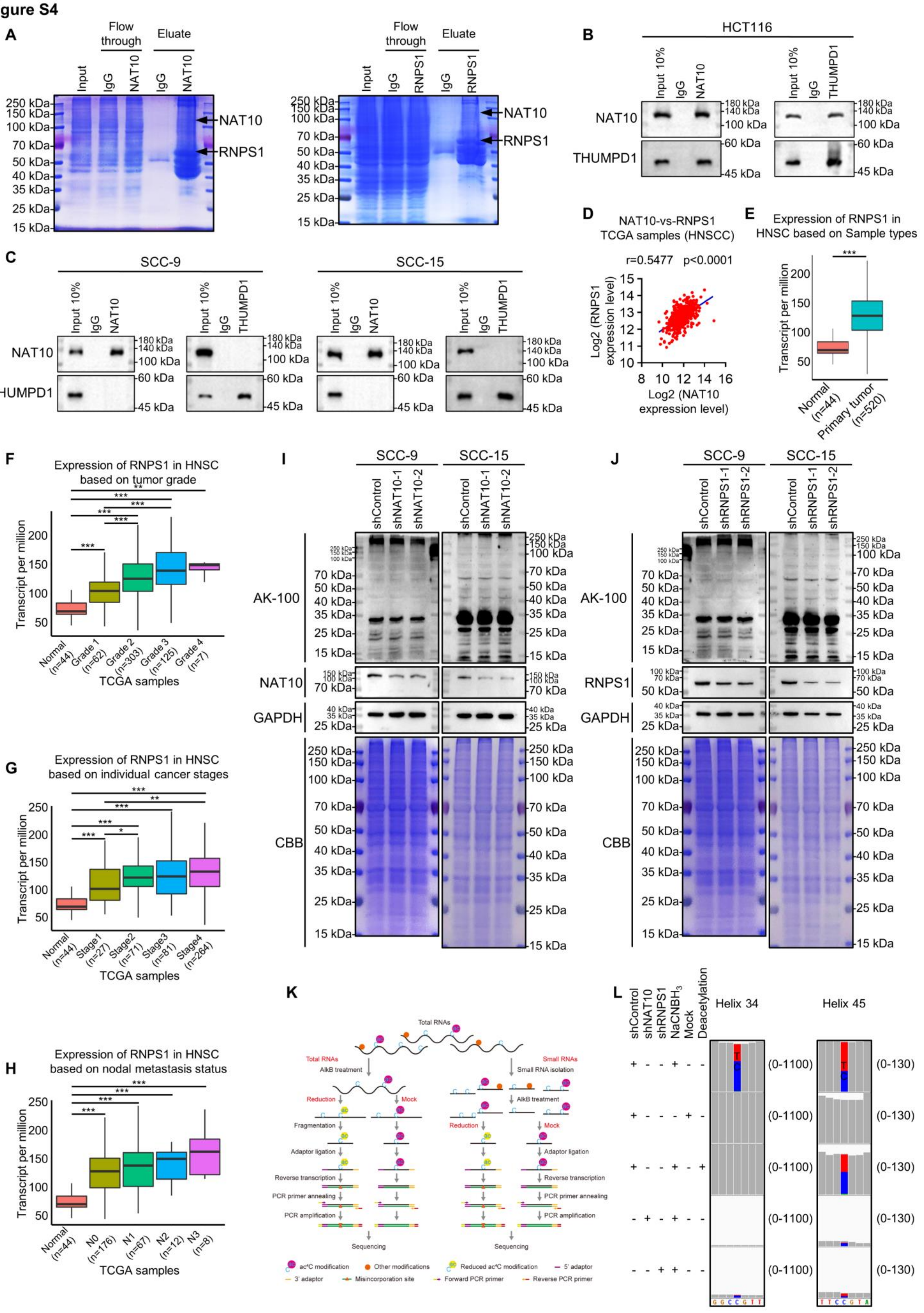
A
B
HCT116
C
SCC-9
SCC-15
D
NAT10-vs-RNPS1
TCGA samples (HNSCC)
r=0.5477 p<0.0001
E
Expression of RNPS1 in HNSC based on Sample types
F
Expression of RNPS1 in HNSC based on tumor grade
G
Expression of RNPS1 in HNSC based on individual cancer stages
H
Expression of RNPS1 in HNSC based on nodal metastasis status
TCGA samples
I
J
AK-100
GAPDH
CBB
K
L
Helix 34
Helix 45

**Figure S4. NAT10 interacts with RNPS1 to regulate tRNA $ac^4C$ modifications, related to Figure 3**

(A) Protein samples from different steps of protein immunoprecipitation by NAT10 antibody(left), RNPS1 antibody (right) and IgG antibody from SCC-15 were separated by SDS-PAGE and stained with CBB.

(B and C) The co-IP assay of NAT10 and THUMPD1 in HCT116 (B), SCC-9 and SCC-15 (C). Anti-IgG antibody was used as a negative control.

(D) The correlation plot of NAT10 and RNPS1 expression level in HNSCC using TCGA dataset. n = 502, r = 0.5477, p < 0.0001 by Spearman correlation analysis.

(E) Expression of RNPS1 between normal (n = 44) and primary tumor (n = 520) of TCGA data. ***p < 0.001 by unpaired Student's t test.

(F-H) The correlation between RNPS1 expression and HNSCC tumor grade (F), individual cancer stages (G) and nodal metastasis status (H) of TCGA dataset. *p < 0.05, **p < 0.01 and ***p < 0.001 by One-way ANOVA.

(I and J) The lysine acetylation levels of protein detected by WB with anti-acetylated-Lysine (Ac-K2-100) antibody in NAT10 KD (I) or RNPS1 KD (J) cells. GAPDH and CBB staining were used as loading control.

(K) Schematic diagram of total RNA reduction and misincorporation sequencing (left) and tRNA reduction and misincorporation sequencing (TRMC-seq) (right).

(L) Misincorporation rates in total RNA from SCC-15 cells are shown for known sites in 18S rRNA. blue letters C and bars, cytidine; red letters T and bars, thymidine; green letters A, adenosine; orange letters G, guanosine. The number on the right indicates the number of nucleotides in the ordinate.

Figure S5

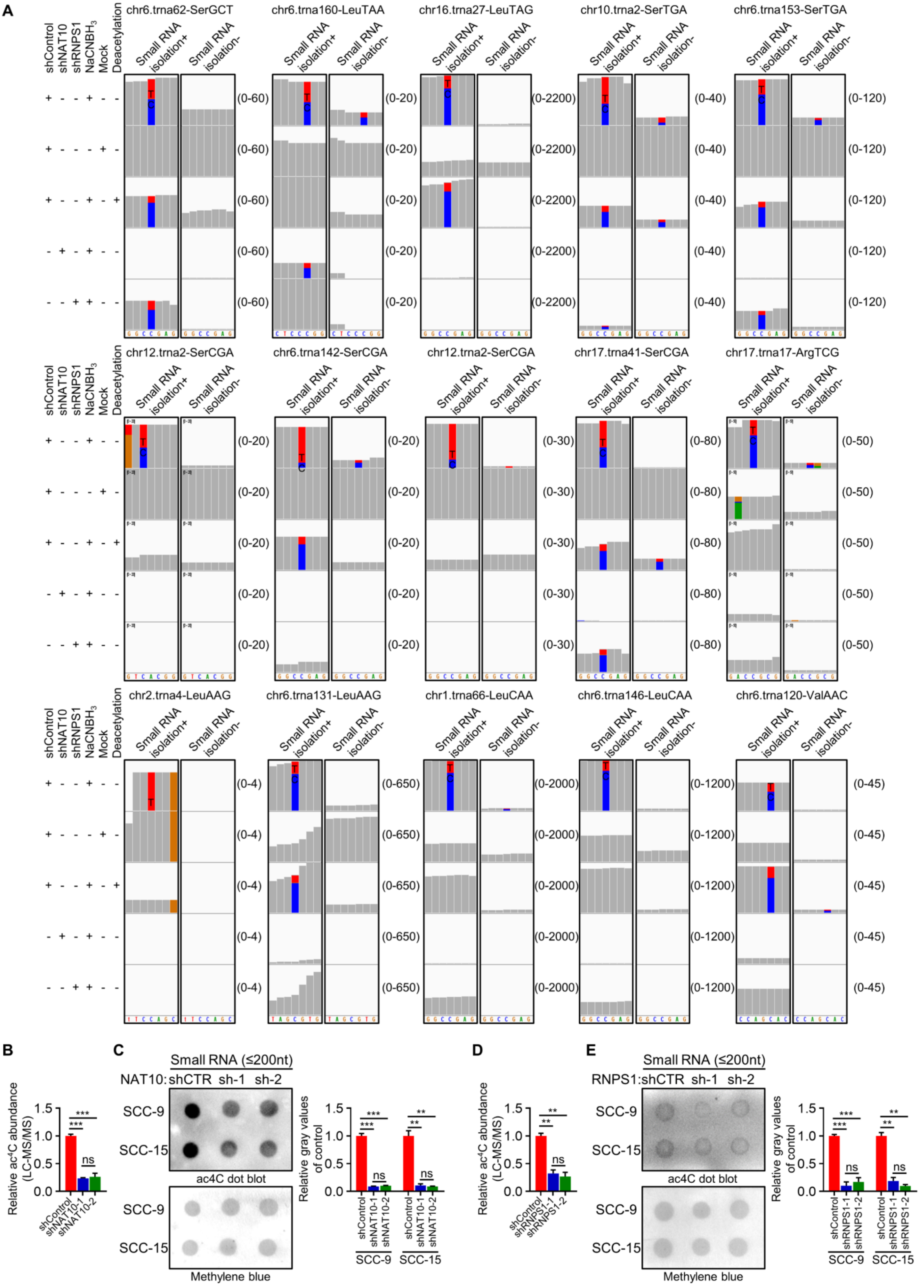
A
chr6.trna62-SerGCT
chr6.trna160-LeuTAA
chr16.trna27-LeuTAG
chr10.trna2-SerTGA
chr6.trna153-SerTGA
chr12.trna2-SerCGA
chr6.trna142-SerCGA
chr12.trna2-SerCGA
chr17.trna41-SerCGA
chr17.trna17-ArgTCG
chr2.trna4-LeuAAG
chr6.trna131-LeuAAG
chr1.trna66-LeuCAA
chr6.trna146-LeuCAA
chr6.trna120-ValAAC
shControl
shNAT10
shRNPS1
NaCNBH$_3$
Mock
Deacetylation
Small RNA isolation+
Small RNA isolation-
B
Relative ac$^4$C abundance (LC-MS/MS)
shControl
shNAT10-1
shNAT10-2
C
Small RNA (≤200nt)
NAT10: shCTR sh-1 sh-2
SCC-9
SCC-15
ac4C dot blot
Methylene blue
Relative gray values of control
D
shRNPS1-1
shRNPS1-2
E
RNPS1: shCTR sh-1 sh-2
ns

**Figure S5. NAT10 interacts with RNPS1 to regulate tRNA $ac^4C$ modifications, related to Figure 3**

(A) From left to right, from top to bottom, Misincorporation rates of $ac^4C_{12}$ in $tRNA^{Ser(GCT)}$, $ac^4C_{79}$ in $tRNA^{Leu(TAA)}$, $ac^4C_{12}$ in $tRNA^{Leu(TAG)}$, $ac^4C_{12}$ in $tRNA^{Ser(TGA)}$, $ac^4C_{12}$ in tRNA $^{Ser(TGA)}$, $ac^4C_3$ in $tRNA^{Ser(CGA)}$, $ac^4C_{12}$ in $tRNA^{Ser(CGA)}$, $ac^4C_{12}$ in $tRNA^{Ser(CGA)}$, $ac^4C_{12}$ in $tRNA^{Ser(CGA)}$, $ac^4C_4$ in $tRNA^{Arg(TCG)}$, $ac^4C_{50}$ in $tRNA^{Leu(AAG)}$, $ac^4C_6$ in $tRNA^{Leu(AAG)}$, $ac^4C_{12}$ in $tRNA^{Leu(CAA)}$, $ac^4C_{12}$ in $tRNA^{Leu(CAA)}$ and $ac^4C_{65}$ in $tRNA^{Val(AAC)}$ in small RNA and total RNA from SCC-15 cells were detected by TRMC-seq or total RNA reduction and misincorporation sequencing. blue letters C and bars, cytidine; red letters T and bars, thymidine; green letters A and bars, adenosine; orange letters G and bars, guanosine. Small RNA isolation+, detected by TRMC-seq; Small RNA isolation-, detected by total RNA reduction and misincorporation sequencing. The number on the right indicates the number of nucleotides in the ordinate.

(B) The relative $ac^4C$ level of small RNA (≤200nt) detected by LC-MS/MS in Control and NAT10-KD groups of SCC-15 cells. n = 2 independent biological replicates. Data are represented as mean ± SD. ***$p < 0.001$ by One-way ANOVA.

(C) Dot blot (left) and relative gray values (right) showed the $ac^4C$ content of small RNA in Control and NAT10-KD groups of SCC-9 or SCC-15 cells with methylene blue staining as loading control. n = 2 independent biological replicates. Data are represented as mean ± SD. **$p < 0.01$ and ***$p < 0.001$ by One-way ANOVA.

(D) The bar chart showed relative $ac^4C$ abundance of small RNA by LC-MS/MS between Control and RNPS1-KD groups of SCC-15 cells. n = 2 independent biological replicates. Data are represented as mean ± SD. **$p < 0.01$ by One-way ANOVA.

(E) Dot blot assay (left) and relative gray statistical values (right) showed that compared with the Control, RNPS1-KD group displayed the downward trend in $ac^4C$ levels of small RNA of SCC-9 or SCC-15 cells with methylene blue staining as loading control. n = 2 independent biological replicates. Data are represented as mean ± SD. **$p < 0.01$ and ***$p < 0.001$ by One-way ANOVA.

**Figure S6**

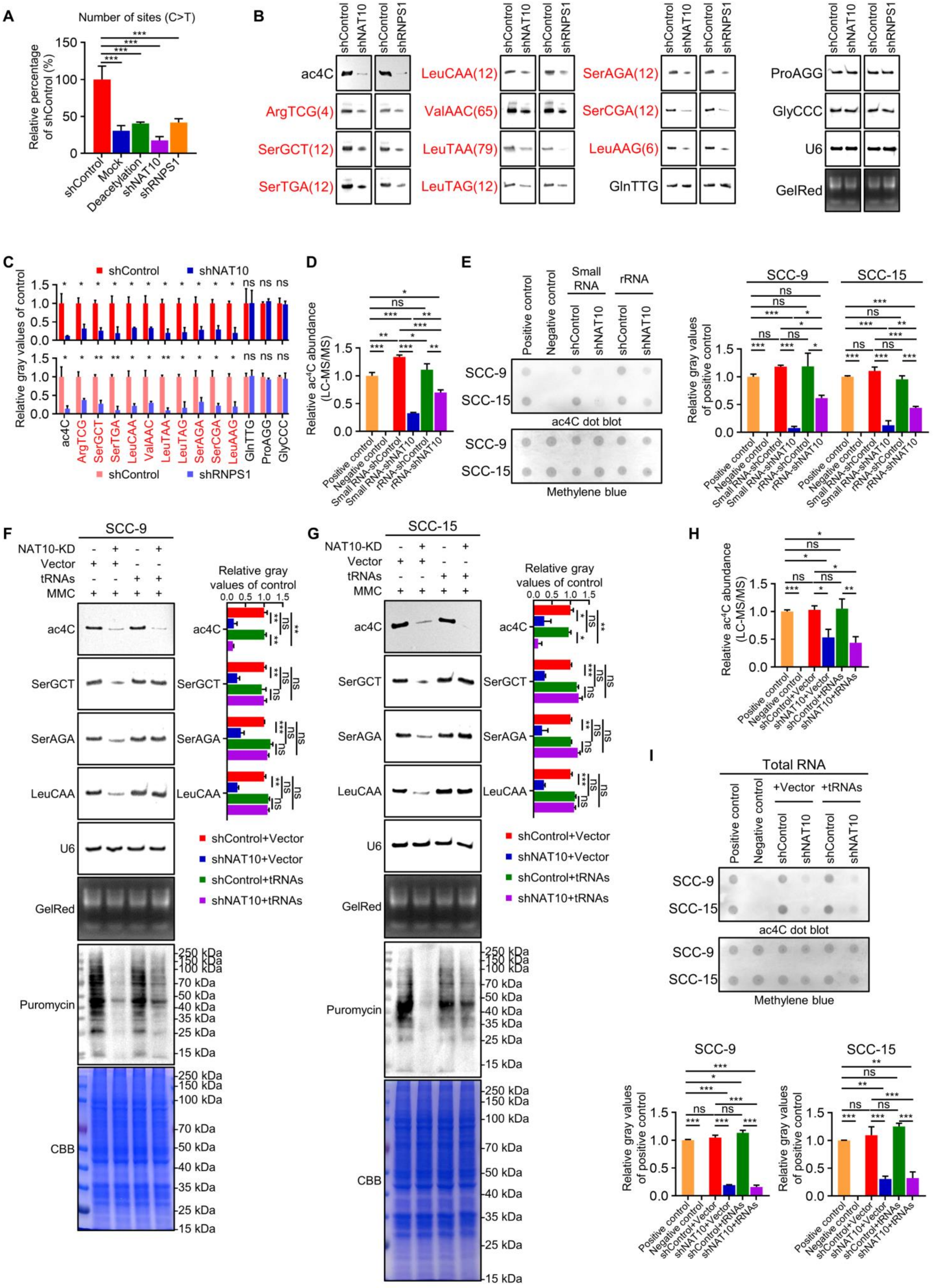
A
Number of sites (C>T)
Relative percentage of shControl (%)
***
shControl
Mock
Deacetylation
shNAT10
shRNPS1
B
shControl
shNAT10
shRNPS1
ac4C
ArgTCG(4)
SerGCT(12)
SerTGA(12)
LeuCAA(12)
ValAAC(65)
LeuTAA(79)
LeuTAG(12)
SerAGA(12)
SerCGA(12)
LeuAAG(6)
GlnTTG
ProAGG
GlyCCC
U6
GelRed
C
Relative gray values of control
ns
ac4C
ArgTCG
SerGCT
SerTGA
LeuCAA
ValAAC
LeuTAA
LeuTAG
SerAGA
SerCGA
LeuAAG
GlnTTG
ProAGG
GlyCCC
D
Relative ac4C abundance (LC-MS/MS)
Positive control
Negative control
Small RNA-shControl
Small RNA-shNAT10
rRNA-shControl
rRNA-shNAT10
E
Small RNA
rRNA
SCC-9
SCC-15
ac4C dot blot
Methylene blue
Relative gray values of positive control
F
NAT10-KD
Vector
tRNAs
MMC
Relative gray values of control
shControl+Vector
shNAT10+Vector
shControl+tRNAs
shNAT10+tRNAs
Puromycin
CBB
250 kDa
150 kDa
100 kDa
70 kDa
50 kDa
40 kDa
35 kDa
25 kDa
15 kDa
G
H
I
Total RNA
+Vector
+tRNAs

**Figure S6. NAT10 interacts with RNPS1 to regulate tRNA $ac^4C$ modifications, related to Figure 3 and Figure 4**

(A) The number of misincorporation (C>T) sites detected in shControl, Mock, Deacetylation, shNAT10, shRNPS1 groups. Data are represented as mean ± SD. ***p < 0.001 by One-way ANOVA.

(B and C) NWB & NB (B) and relative gray values (C) showed $ac^4C$ modification levels of RNA and expression of non-$ac^4C$ tRNAs and $ac^4C$ tRNAs (red font) between Control, NAT10 KD and RNPS1 KD groups. NB of U6 snRNA and GelRed staining were used as loading control. The numbers in brackets represent the $ac^4C$ sites. n = 2 independent biological replicates. Data are represented as mean ± SD. *p < 0.05 and **p < 0.01 in (C) by unpaired Student's t test.

(D) The bar chart of relative $ac^4C$ abundance of RNA by LC-MS/MS in Positive control, Negative control, Small RNA-Control, Small RNA-NAT10-KD, rRNA-Control and rRNA-NAT10-KD of SCC-15 cells. Positive control: ribosomal RNA; Negative control: poly(A)-enriched RNA. n = 2 independent biological replicates. Data are represented as mean ± SD. *p < 0.05, **p < 0.01 and ***p < 0.001 by One-way ANOVA.

(E) Dot blot (left) and relative gray values (right) showed the $ac^4C$ level of RNA in Positive control, Negative control, Small RNA-Control, Small RNA-NAT10-KD, rRNA-Control and rRNA-NAT10-KD groups with methylene blue staining as loading control. n = 2 independent biological replicates. Data are represented as mean ± SD. *p < 0.05, **p < 0.01 and ***p < 0.001 by One-way ANOVA.

(F and G) NWB & NB showed $ac^4C$ modification levels of RNA and expression of SerGCT, SerAGA and LeuCAA tRNAs between Control + Vector, NAT10-KD + Vector, Control + tRNAs-OE (SerGCT, SerAGA and LeuCAA) and NAT10-KD + tRNAs-OE groups in SCC-9 (F) and SCC-15 (G). The corresponding relative gray values is on the right. NB of U6 snRNA and GelRed staining were used as loading control. The following is the corresponding puromycin assay and CBB staining was used as loading control. MMC was added to inhibit the effect of cell proliferation. Data are represented as mean ± SD. *p < 0.05, **p < 0.01 and ***p < 0.001 by One-way ANOVA.

(H) The relative $ac^4C$ level of RNA by LC-MS/MS in SCC-15 cells. Positive control: ribosomal RNA; Negative control: poly(A)-enriched RNA; The remaining samples are total RNA from Control + Vector, NAT10-KD + Vector, Control + tRNAs-OE (SerGCT, SerAGA and LeuCAA) and NAT10-KD + tRNAs-OE groups. n = 2 independent biological replicates. Data are represented as mean ± SD. **p < 0.01 and ***p < 0.001 by One-way ANOVA.

(I) Dot blot (up) and relative gray values (down) showed the $ac^4C$ content of RNA in Positive control, Negative control, Control + Vector, NAT10-KD + Vector, Control + tRNAs-OE (SerGCT, SerAGA and LeuCAA) and NAT10-KD + tRNAs-OE groups with methylene blue staining as loading control. n = 2 independent biological replicates. Data are represented as mean ± SD. ***p < 0.001 in (H) by One-way ANOVA.

### Figure S7

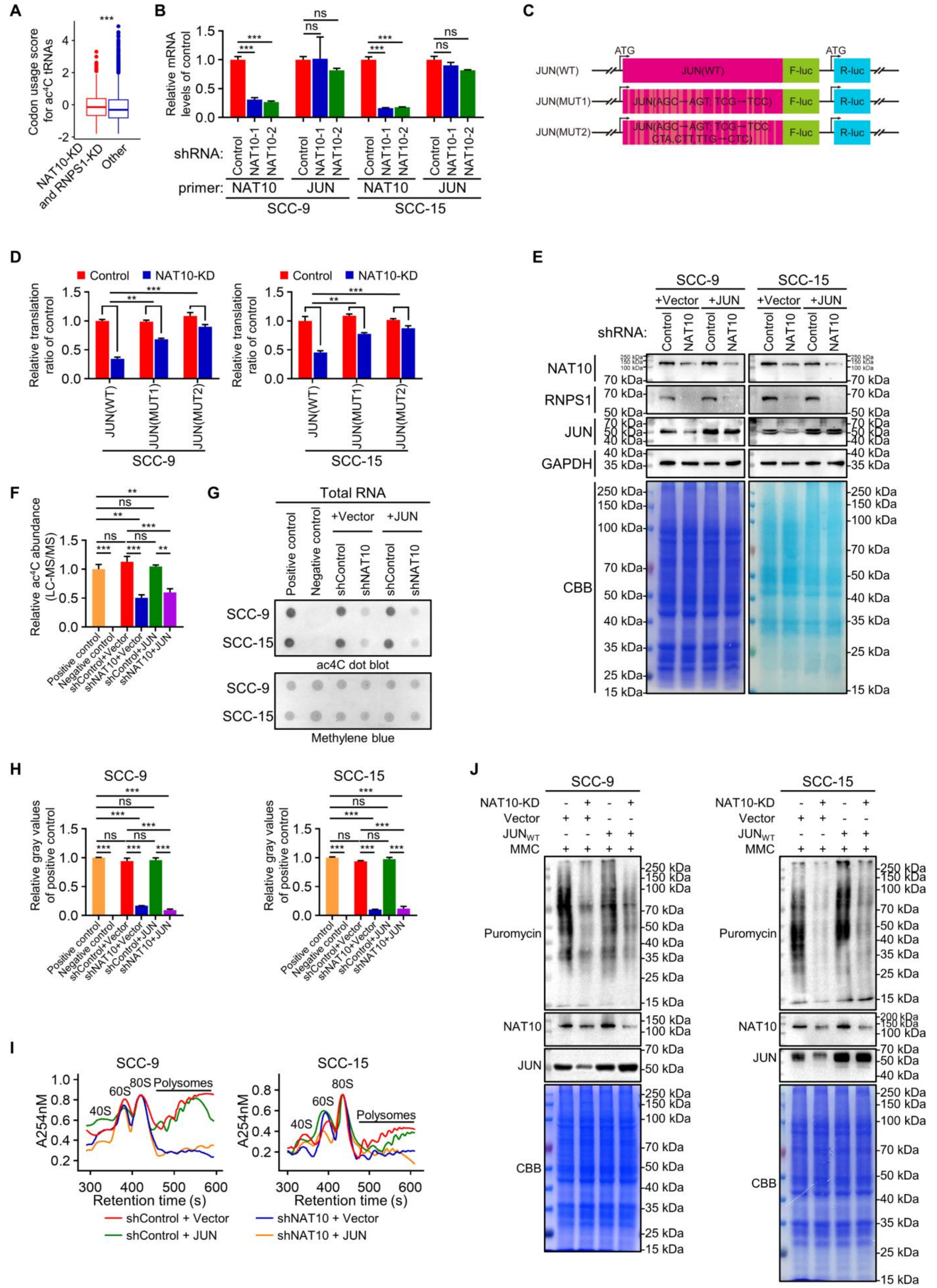

A
Codon usage score for ac4C tRNAs
NAT10-KD and RNPS1-KD
Other
B
Relative mRNA levels of control
shRNA:
Control
NAT10-1
NAT10-2
primer:
NAT10
JUN
SCC-9
SCC-15
C
JUN(WT)
JUN(MUT1)
JUN(MUT2)
ATG
F-luc
R-luc
JUN(AGC→AGT; TCG→TCC)
JUN(AGC→AGT; TCG→TCC CTA,CTT,TTG→CTC)
D
Control
NAT10-KD
Relative translation ratio of control
SCC-9
SCC-15
E
SCC-9
SCC-15
+Vector
+JUN
shRNA:
NAT10
RNPS1
JUN
GAPDH
CBB
F
Relative ac4C abundance (LC-MS/MS)
Positive control
Negative control
shControl+Vector
shNAT10+Vector
shControl+JUN
shNAT10+JUN
G
Total RNA
shControl
shNAT10
ac4C dot blot
Methylene blue
H
Relative gray values of positive control
I
A254nM
40S
60S
80S
Polysomes
Retention time (s)
shControl + Vector
shNAT10 + Vector
shControl + JUN
shNAT10 + JUN
J
NAT10-KD
Vector
JUN_WT
MMC
Puromycin

**Figure S7. Integrated analysis of multiple 'omics' reveal NAT10 and RNPS1 regulate JUN, related to Figure 5**

(A) The codon usage scores of common down regulated genes detected by iTRAQ in NAT10-KD and RNPS1-KD were higher than scores of other genes. The box plot parameters are as in Figure 3J. ***$p < 0.001$ by unpaired Student's t test.

(B) The expression of JUN mRNA remained unchanged after KD of NAT10 in SCC-9 and SCC-15 cells. Data are represented as mean ± SD. ***$p < 0.001$ by One-way ANOVA.

(C and D) Schematic diagram (C) and relative translation ratio column chart (D) of luciferase reporter system. The coding sequence of JUN was inserted into a vector carrying a luciferase reporter system. The mutated JUN transcripts were constructed by replacing the codon AGC to ACT; TCG to TCC (MUT1) or AGC to AGT; TCG to TCC; CTA, CTT, TTG to CTC (MUT2). The relative translation ratios were calculated as the ratio of F-luc translation efficiency (TE) to R-luc translation efficiency and the TE of F-luc and R-luc were calculated by normalizing the luciferase protein levels to mRNA levels. Data are represented as mean ± SD. **$p < 0.01$ and ***$p < 0.001$ in (D) by One-way ANOVA.

(E) The WB showed that JUN was overexpressed after KD of NAT10 in SCC-9 and SCC-15 cells. GAPDH and CBB staining were used as loading control.

(F) The bar chart of relative $ac^4C$ abundance of RNA by LC-MS/MS from SCC-15 cells. Positive control: ribosomal RNA; Negative control: poly(A)-enriched RNA; The remaining samples are total RNA from Control + Vector, NAT10-KD + Vector, Control + JUN-OE and NAT10-KD + JUN-OE groups. n = 2 independent biological replicates. Data are represented as mean ± SD. **$p < 0.01$ and ***$p < 0.001$ by One-way ANOVA.

(G and H) Dot blot (G) and relative gray values (H) showed the $ac^4C$ level of RNA in Positive control, Negative control, Control + Vector, NAT10-KD + Vector, Control + JUN-OE and NAT10-KD + JUN-OE groups with methylene blue staining as loading control. n = 2 independent biological replicates. Data are represented as mean ± SD. ***$p < 0.001$ in (H) by One-way ANOVA.

(I and J) Polysome profiling (I) and Puromycin assay (J) of the Control + Vector, NAT10-KD + Vector, Control + JUN-OE and NAT10-KD + JUN-OE groups in SCC-9 (left) and SCC-15 (right) cells. MMC was added to inhibit the effect of cell proliferation and CBB staining was used as loading control.

**Figure S8**

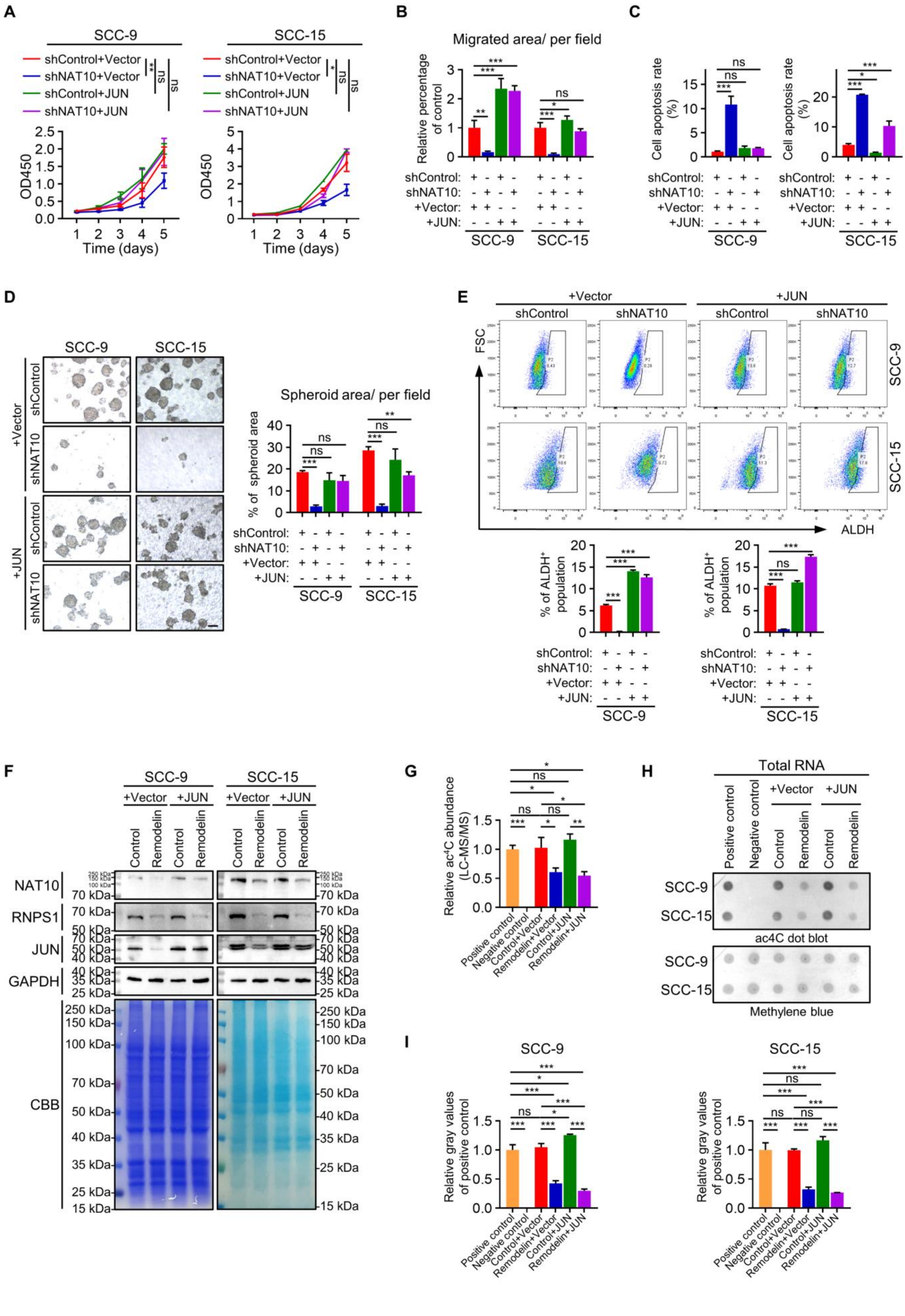

A
SCC-9
SCC-15
shControl+Vector
shNAT10+Vector
shControl+JUN
shNAT10+JUN
OD450
Time (days)
B
Migrated area/ per field
Relative percentage of control
C
Cell apoptosis rate (%)
D
+Vector
+JUN
shControl
shNAT10
Spheroid area/ per field
% of spheroid area
E
FSC
ALDH
% of ALDH+ population
F
NAT10
RNPS1
JUN
GAPDH
CBB
Control
Remodelin
G
Relative ac4C abundance (LC-MS/MS)
Positive control
Negative control
Control+Vector
Remodelin+Vector
Control+JUN
Remodelin+JUN
H
Total RNA
ac4C dot blot
Methylene blue
I
Relative gray values of positive control

**Figure S8. Integrated analysis of multiple 'omics' reveal NAT10 and RNPS1 regulate JUN, related to Figure 5**

(A-E) In the rescue assay of JUN overexpression after KD of NAT10, the phenotypic differences including proliferation ability (A), migration (B), cell apoptosis (C), sphere formation (D) and aldehyde dehydrogenase activity (E) between the shControl + Vector, shNAT10 + Vector, shControl + JUN and shNAT10 + JUN in HNSCC cell lines were observed. The scale bar of (D) is 100 μm. Data are represented as mean ± SD. $*p < 0.05$, $**p < 0.01$ in (A) by Two-way ANOVA. $*p < 0.05$, $**p < 0.01$ and $***p < 0.001$ in (B-E) by One-way ANOVA.

(F) The WB assay showed that JUN was overexpressed after treatment of Remodelin in SCC-9 and SCC-15 cells. Cells were treated with Remodelin at a concentration of 8 μM (SCC-9) or 15 μM (SCC-15) for 24-48 h. GAPDH and CBB staining were used as loading control.

(G) The bar chart of relative $ac^4C$ abundance of RNA by LC-MS/MS from SCC-15 cells. Positive control: ribosomal RNA; Negative control: poly(A)-enriched RNA; The remaining samples are total RNA from DMSO treatment + Vector, Remodelin treatment + Vector, DMSO treatment + JUN-OE and Remodelin treatment + JUN-OE groups. Cells were treated with Remodelin at a concentration of 8 μM (SCC-9) or 15 μM (SCC-15) for 24-48 h. n = 2 independent biological replicates. Data are represented as mean ± SD. $*p < 0.05$, $**p < 0.01$ and $***p < 0.001$ by One-way ANOVA.

(H and I) Dot blot (H) and relative gray values (I) showed the $ac^4C$ level of RNA in Positive control, Negative control, DMSO treatment + Vector, Remodelin treatment + Vector, DMSO treatment + JUN-OE and Remodelin treatment + JUN-OE groups with methylene blue staining as loading control. Cells were treated with Remodelin at a concentration of 8 μM (SCC-9) or 15 μM (SCC-15) for 24-48 h. n = 2 independent biological replicates. Data are represented as mean ± SD. $*p < 0.05$ and $***p < 0.001$ in (I) by One-way ANOVA.

### Figure S9

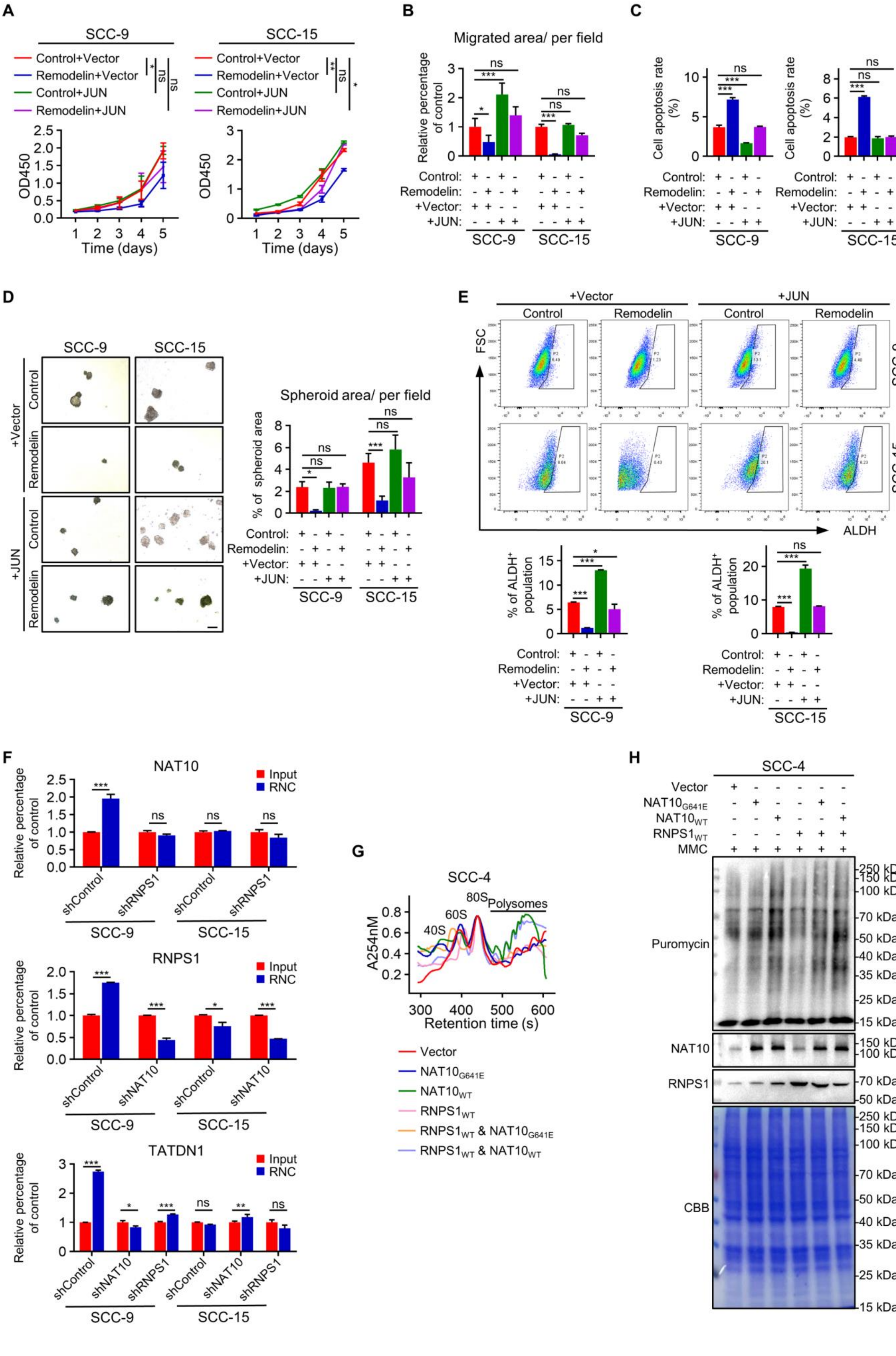
A
SCC-9
SCC-15
Control+Vector
Remodelin+Vector
Control+JUN
Remodelin+JUN
OD450
Time (days)
B
Migrated area/ per field
Relative percentage of control
Control:
Remodelin:
+Vector:
+JUN:
C
Cell apoptosis rate (%)
D
Spheroid area/ per field
% of spheroid area
+Vector
+JUN
Control
Remodelin
E
FSC
ALDH
% of ALDH+ population
F
NAT10
RNPS1
TATDN1
Input
RNC
shControl
shRNPS1
shNAT10
G
SCC-4
80S
60S
40S
Polysomes
A254nM
Retention time (s)
Vector
NAT10G641E
NAT10WT
RNPS1WT
RNPS1WT & NAT10G641E
RNPS1WT & NAT10WT
H
MMC
Puromycin
CBB
250 kDa
150 kDa
100 kDa
70 kDa
50 kDa
40 kDa
35 kDa
25 kDa
15 kDa

**Figure S9. Integrated analysis of multiple 'omics' reveal NAT10 and RNPS1 regulate JUN, related to Figure 5**

(A-E) In the rescue assay of JUN overexpression after treatment of Remodelin, the phenotypic differences including proliferation ability (A), migration (B), cell apoptosis (C), sphere formation (D) and aldehyde dehydrogenase activity (E) between the Control + Vector, Remodelin + Vector, Control + JUN and Remodelin + JUN in HNSCC cell lines were observed. Cells were treated with Remodelin at a concentration of 8 μM (SCC-9) or 15 μM (SCC-15) for 5 d (A), 48 h (B, C and E) or 3-4 d (D), respectively. The scale bar of (D) is 100 μm. Data are represented as mean ± SD. $*p < 0.05$, $**p < 0.01$ in (A) by Two-way ANOVA. $*p < 0.05$ and $***p < 0.001$ in (B-E) by One-way ANOVA.

(F) The content of NAT10 mRNA, RNPS1 mRNA and TATDN1 mRNA at the ribosome combined stages and input stages after KD of NAT10 or RNPS1. Data are represented as mean ± SD. $*p < 0.05$, $**p < 0.01$ and $***p < 0.001$ by Two-way ANOVA.

(G and H) Polysome profiling (G) and Puromycin assay (H) of the Vector, NAT10-MUT, NAT10-WT, RNPS1-WT, RNPS1-WT & NAT10-MUT and RNPS1-WT & NAT10-WT groups in SCC-4 cells. MMC was added to inhibit the effect of cell proliferation and CBB staining was used as loading control.

**Figure S10**

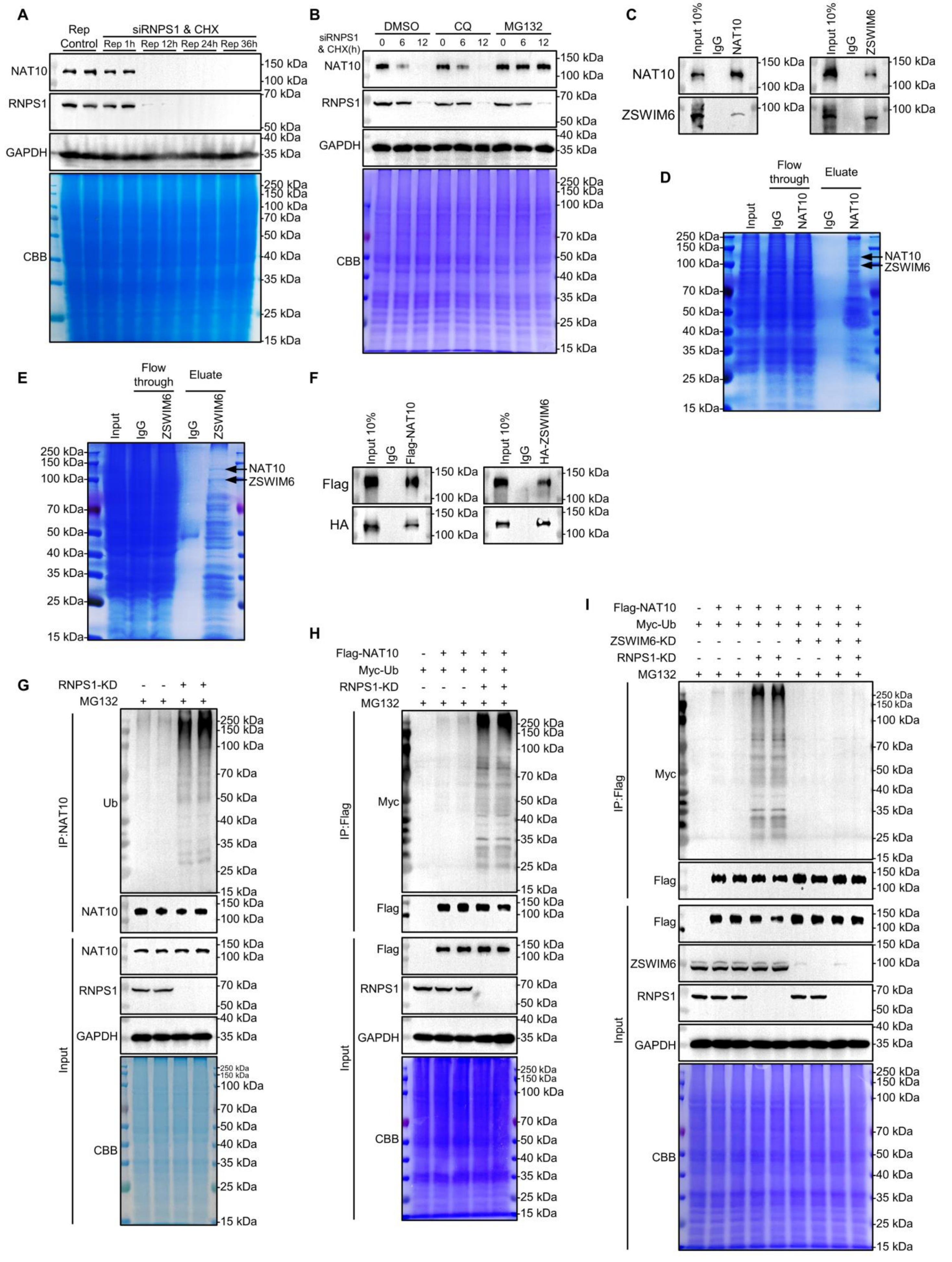
A
Rep Control
siRNPS1 & CHX
Rep 1h
Rep 12h
Rep 24h
Rep 36h
NAT10
RNPS1
GAPDH
CBB
B
DMSO
CQ
MG132
siRNPS1 & CHX(h)
0 6 12 0 6 12 0 6 12
NAT10
RNPS1
GAPDH
CBB
C
Input 10%
IgG
NAT10
ZSWIM6
D
Flow through
Eluate
Input
IgG
NAT10
ZSWIM6
E
Flow through
Eluate
Input
IgG
ZSWIM6
NAT10
F
Input 10%
IgG
Flag-NAT10
HA-ZSWIM6
Flag
HA
G
RNPS1-KD
MG132
IP:NAT10
Ub
NAT10
Input
RNPS1
GAPDH
CBB
H
Flag-NAT10
Myc-Ub
RNPS1-KD
MG132
IP:Flag
Myc
Flag
Input
RNPS1
GAPDH
CBB
I
Flag-NAT10
Myc-Ub
ZSWIM6-KD
RNPS1-KD
MG132
IP:Flag
Myc
Flag
Input
ZSWIM6
RNPS1
GAPDH
CBB
250 kDa
150 kDa
100 kDa
70 kDa
50 kDa
40 kDa
35 kDa
25 kDa
15 kDa

**Figure S10. RNPS1 ensures the stability of NAT10 by inhibiting the ubiquitination of NAT10**

(A) WB image and analysis showing NAT10 protein expression level in SCC-15 cells for serial RNPS1 knock-down and cycloheximide (CHX, 100 μg/mL) treatment durations. WB of GAPDH and Coomassie brilliant blue staining were used as loading control.

(B) WB showing expression level in cells treated with 100 μg/mL cycloheximide (CHX) for the indicated durations in the presence of RNPS1 knock-down with 20 μM chloroquine (CQ) or 10 μM MG132. WB of GAPDH and Coomassie brilliant blue staining were used as loading control.

(C) Association of endogenous NAT10 with ZSWIM6 in SCC-15 by co-IP with anti-NAT10 antibody or anti-ZSWIM6 antibody after RNPS1 knock-down and MG132 treatment. Anti-IgG antibody was used as a negative control.

(D and E) Protein samples from different steps of protein immunoprecipitation by NAT10 antibody(D), ZSWIM6 antibody (E) and IgG antibody from SCC-15 after RNPS1 knock-down and MG132 treatment were separated by SDS-PAGE and stained with Coomassie brilliant blue.

(F) Co-IP assay of Flag-NAT10 and HA-ZSWIM6 in 293T cells, which were transiently co-transfected with Flag-NAT10 and HA-ZSWIM6 plasmids and treated with RNPS1 knock-down and MG132.

(G) The WB images of NAT10 ubiquitination in SCC-15 cells after RNPS1 knock-down and MG132 (10 μM) treatment. GAPDH and CBB staining were used as loading control.

(H) The ubiquitination level of Flag-NAT10 analyzed by precipitating Flag-NAT10 with an anti-Flag antibody followed by WB using anti-Myc antibody in 293T cells transfected with Flag-NAT10 and Myc-Ub, after RNPS1 knock-down and MG132 (10 μM) treatment as indicated. GAPDH and CBB staining were used as loading control.

(I) The WB images shows the Myc-Ubiquitin enrichment level of Flag-NAT10 under different conditions including MG132 (10 μM) treatment, RNPS1 knock-down and/or ZSWIM6 knock-down. GAPDH and CBB staining were used as loading control.

**Figure S11**

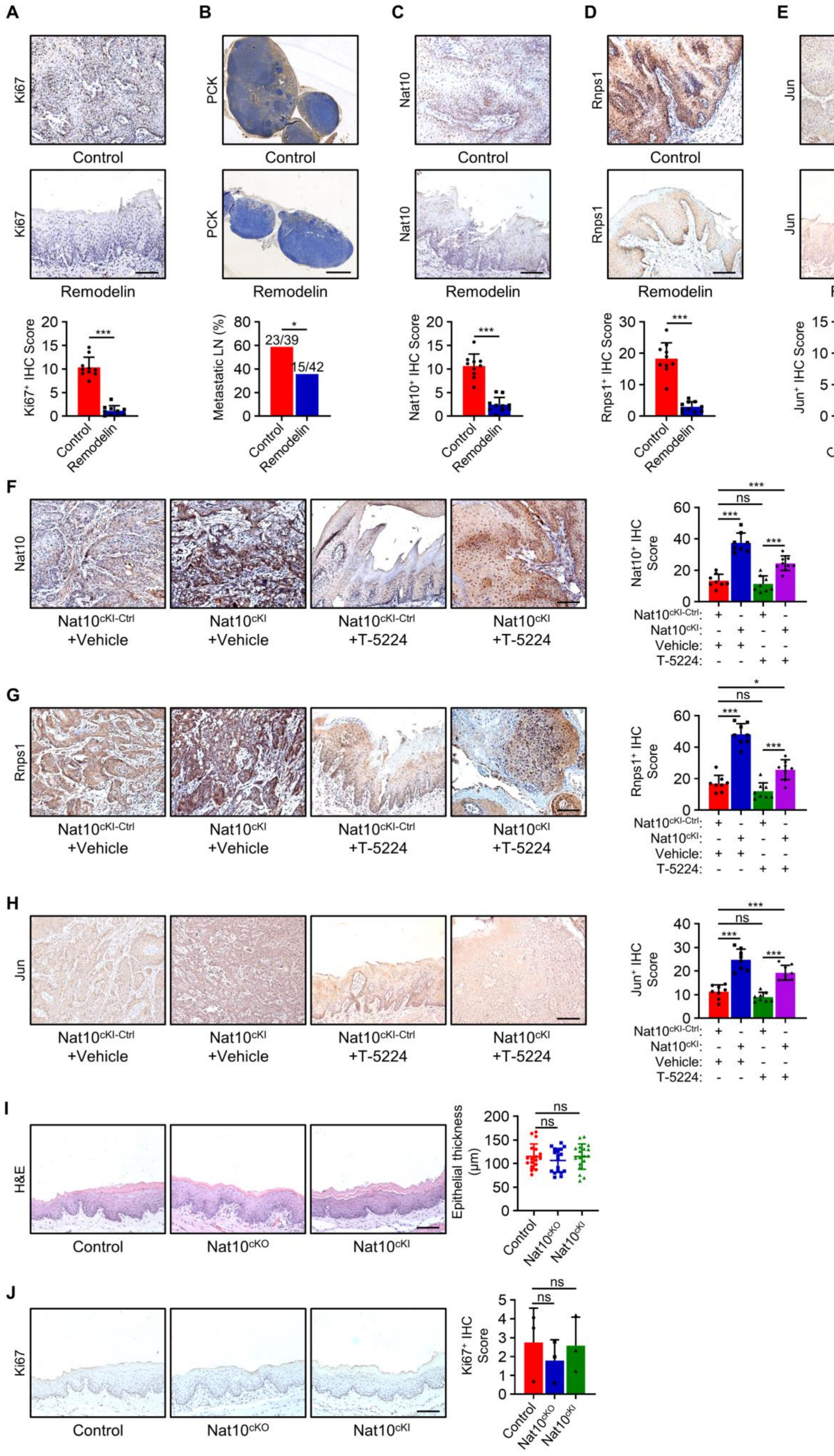

A
B
C
D
E
Ki67
Control
Remodelin
PCK
Nat10
Rnps1
Jun
Ki67⁺ IHC Score
***
Metastatic LN (%)
*
23/39
15/42
Nat10⁺ IHC Score
Rnps1⁺ IHC Score
Jun⁺ IHC Score
F
G
H
Nat10cKI-Ctrl +Vehicle
Nat10cKI +Vehicle
Nat10cKI-Ctrl +T-5224
Nat10cKI +T-5224
Nat10⁺ IHC Score
Rnps1⁺ IHC Score
Jun⁺ IHC Score
ns
Nat10cKI-Ctrl:
Nat10cKI:
Vehicle:
T-5224:
I
H&E
Control
Nat10cKO
Nat10cKI
Epithelial thickness (μm)
J
Ki67⁺ IHC Score

**Figure S11. Nat10 drives chemically-induced HNSCC tumorigenesis, related to Figure 6 and Figure 7**

(A-E) Representative IHC staining and IHC Score of Ki67 (A), Nat10 (C), Rnps1(D) and Jun (E) in HNSCC and PCK (B) in lymph node. The scale bar of (B) is 500 μm and the rest are 100 μm. n = 10 mice in each group. Data are represented as mean ± SD. *p < 0.05 and ***p < 0.001 in (A) and (C-E) by unpaired Student's t test. *p < 0.05 in (B) by Chi-square test.

(F-H) Representative IHC staining and IHC Score of Nat10 (F), Rnps1(G) and Jun (H) in HNSCC between different treatment groups. Scale bars are 100 μm. Data are represented as mean ± SD. *p < 0.05 and ***p < 0.001 by One-way ANOVA.

(I) Representative H&E staining of oral epithelium and quantification of oral epithelial thickness. Mice were not fed with 4NQO drinking water and six random locations of each mouse were selected for epithelial thickness measurement. n = 3 mice for each group. Scale bar, 100 μm. Data are represented as mean ± SD. p values by One-way ANOVA.

(J) Representative IHC staining and IHC Score of Ki67 in normal oral epithelium between Control, NAT10-KO and NAT10-KI mice groups. n = 3 mice for each group. Scale bar, 100 μm. Data are represented as mean ± SD. Data are represented as mean ± SD. p values by One-way ANOVA.

**NB replicates**

**Fig. 6 L**

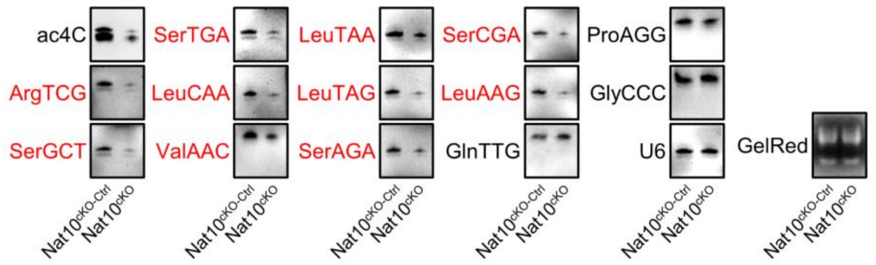


**Fig. S6 B**

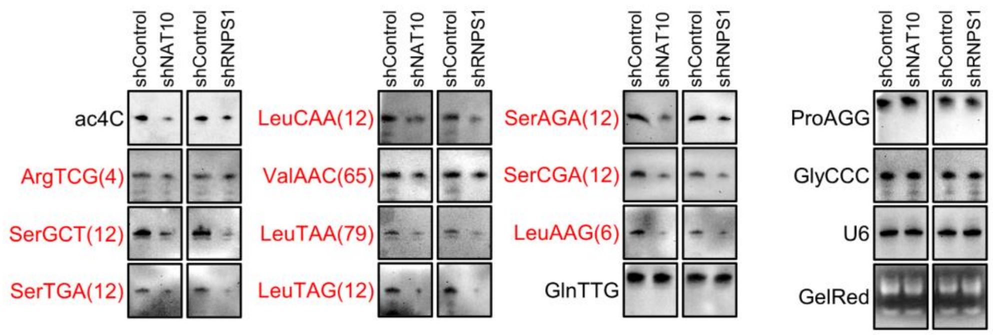

### WB replicates

**Fig. 2 B**

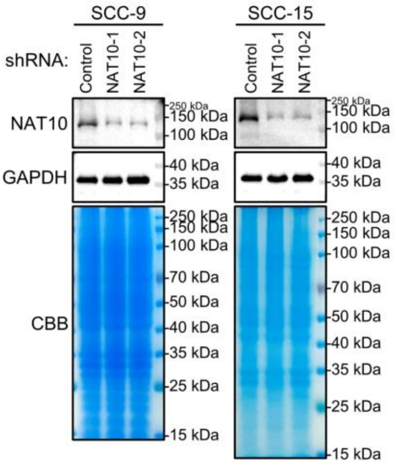


**Fig. 3 A**

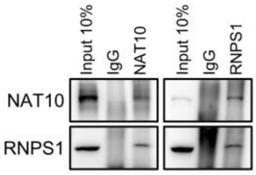


**Fig. 3 G**

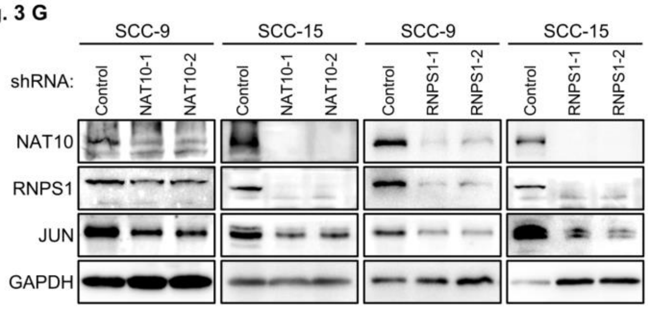


**Fig. S2 E**

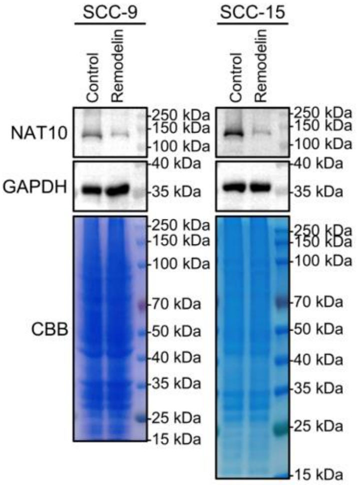


**Fig. S3 F**

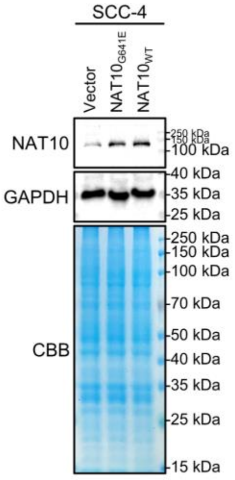


**Fig. S7 E**

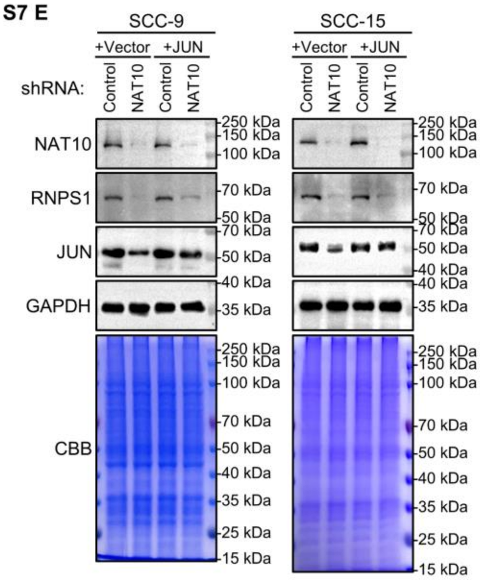


**Fig. S8 F**

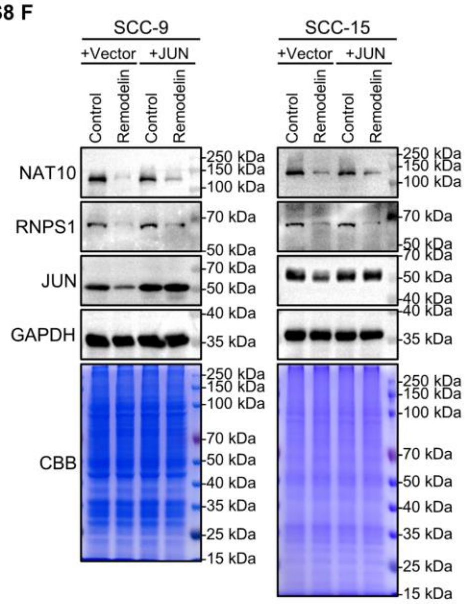


**Fig. S10 B**

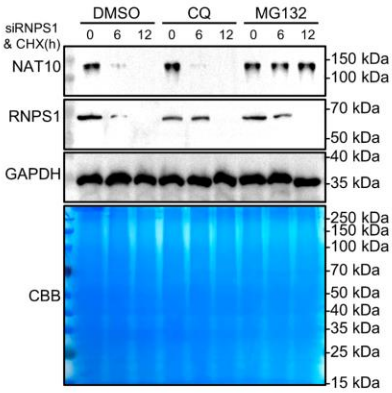

Table S1. related to Figure 3

| | | | | shControl | | | | | |
|---|---|---|---|---|---|---|---|---|---|
| Accessi | Gene Na | Descript | Mass | Score | Matches | Sequenc | emPAI | Coverage | |
| sp\|P3558 | MYH10 | Myosin-1 | 229827 | 3390 | 173(114) | 88(65) | 2.26 | 44% | P35580_ |
| tr\|Q1KLZ( | PS1TP5I | HCG159 | 42052 | 1014 | 44(35) | 10(9) | 2.35 | 39% | Q1KLZ0_ |
| sp\|Q9H0/ | NAT10 | RNA cyti | 116569 | 596 | 41(23) | 25(17) | 0.79 | 28% | Q9H0A0_ |
| tr\|Q05CW | NAT10 | NAT10 p | 63029 | 383 | 24(14) | 13(11) | 1.04 | 27% | Q05CW7_ |
| tr\|A0A2R | ACTB | Actin, cy | 8839 | 355 | 11(11) | 3(3) | 6.33 | 50% | A0A2R8Y |
| tr\|A0A07 | TRAJ56 | T cell rec | 2220 | 309 | 100(47) | 2(1) | 1.81 | 100% | A0A075B |
| sp\|P0CG3 | POTEI | POTE ar | 122858 | 272 | 16(10) | 5(2) | 0.14 | 4% | P0CG38_ |
| tr\|V9HVZ | HEL-S-1 | Glyceral | 36201 | 228 | 12(6) | 10(5) | 0.55 | 36% | V9HVZ4_ |
| sp\|P7852 | PRKDC | DNA-dep | 473749 | 206 | 41(7) | 39(7) | 0.09 | 11% | P78527_ |
| tr\|A0A08 | CLTC | Clathrin I | 193703 | 190 | 17(8) | 14(7) | 0.22 | 11% | A0A087W |
| tr\|B3KPS3 | | Tubulin a | 46725 | 183 | 10(4) | 9(4) | 0.51 | 30% | B3KPS3_ |
| tr\|Q6IPN( | EEF1A1 | Elongatic | 50433 | 180 | 9(7) | 6(6) | 0.46 | 17% | Q6IPN6_ |
| tr\|A0A0S2 | KRT6A | HCG203 | 60293 | 170 | 19(9) | 14(8) | 0.61 | 25% | A0A0S2Z |
| tr\|B4DKV | | cDNA FL | 55983 | 140 | 18(6) | 12(5) | 0.49 | 24% | B4DKV4_ |
| sp\|P4932 | FASN | Fatty aci | 275877 | 115 | 23(6) | 21(6) | 0.09 | 11% | P49327_ |
| tr\|B4E2J5 | | cDNA FL | 21684 | 115 | 6(3) | 5(3) | 1.06 | 29% | B4E2J5_u |
| tr\|A0A02 | ANXA2 | Annexin | 38808 | 102 | 5(3) | 5(3) | 0.39 | 20% | A0A024R |
| sp\|P8160 | DCD | Dermcidi | 11391 | 93 | 2(2) | 1(1) | 0.30 | 10% | P81605_ |
| tr\|B4DL32 | | cDNA FL | 27409 | 87 | 7(2) | 6(2) | 0.41 | 28% | B4DL32_ |
| tr\|A0A38 | | Tubulin b | 49893 | 83 | 8(3) | 7(3) | 0.29 | 19% | A0A384N |
| tr\|Q1466( | | Radiated | 30933 | 82 | 5(3) | 3(3) | 0.36 | 7% | Q14666_ |
| tr\|A0A08 | SPTBN1 | Spectrin | 275457 | 80 | 14(3) | 12(3) | 0.06 | 6% | A0A087W |
| tr\|Q6NVC | SLC25A5 | SLC25A5 | 35499 | 77 | 10(4) | 5(4) | 0.56 | 17% | Q6NVC0_ |
| tr\|H7C0A | ARPC4-T | ARPC4-T | 19558 | 72 | 6(2) | 3(1) | 0.17 | 17% | H7C0A3_ |
| sp\|Q1514 | PLEC | Plectin C | 533462 | 70 | 15(2) | 15(2) | 0.02 | 3% | Q15149_ |
| tr\|D3DU9 | RNPS1 | RNA bin | 34188 | 68 | 3(2) | 3(2) | 0.20 | 11% | D3DU92_ |
| sp\|Q142( | DYNC1H | Cytoplas | 534809 | 66 | 25(3) | 23(3) | 0.02 | 6% | Q14204_ |
| tr\|G3V1A | CFL1 | Cofilin 1 | 17029 | 59 | 4(1) | 3(1) | 0.20 | 26% | G3V1A4_ |
| tr\|A0A02 | DLST | Dihydroli | 49067 | 53 | 2(2) | 1(1) | 0.07 | 1% | A0A024R |
| tr\|A0A02 | ACTR3 | ARP3 ac | 47797 | 53 | 5(2) | 5(2) | 0.14 | 15% | A0A024R |
| tr\|A0A02 | CLPX | ClpX cas | 69922 | 52 | 3(1) | 3(1) | 0.05 | 5% | A0A024R |
| tr\|V9HW2 | HEL-S-1 | ATP synt | 59828 | 51 | 4(1) | 2(1) | 0.05 | 3% | V9HW26_ |
| tr\|A0A08 | ALB | Serum al | 46430 | 50 | 3(1) | 1(1) | 0.07 | 3% | A0A087W |
| tr\|Q7L4M | | KRT8 pr | 30802 | 50 | 3(2) | 3(2) | 0.23 | 12% | Q7L4M3_ |
| sp\|P6116 | ACTR2 | Actin-rel | 45017 | 48 | 6(2) | 6(2) | 0.24 | 15% | P61160_ |
| tr\|A8K7F6 | | cDNA FL | 46321 | 48 | 4(2) | 4(2) | 0.15 | 11% | A8K7F6_ |
| sp\|Q9HC | ZSWIM6 | Zinc fing | 135096 | 46 | 1(1) | 1(1) | 0.02 | 0% | Q9HCJ5_ |
| sp\|P0510 | S100A8 | Protein S | 10885 | 46 | 1(1) | 1(1) | 0.32 | 11% | P05109_ |
| tr\|A0A24 | | 40S ribo | 17049 | 45 | 1(1) | 1(1) | 0.20 | 11% | A0A248R |
| tr\|A0A0B | LYZ | Lysozym | 15661 | 43 | 1(1) | 1(1) | 0.22 | 8% | A0A0B4J |
| tr\|A0A02 | PKM2 | Pyruvate | 58538 | 42 | 7(1) | 6(1) | 0.18 | 13% | A0A024R |
| tr\|A0A02 | SLC25A3 | Solute ca | 40389 | 41 | 2(1) | 2(1) | 0.08 | 7% | A0A024R |
| tr\|A0A38 | | Epididym | 36494 | 40 | 2(1) | 2(1) | 0.09 | 6% | A0A384N |
| tr\|A0A0A | GSN | Gelsolin | 82759 | 40 | 5(0) | 5(0) | 0.08 | 7% | A0A0A0N |
| tr\|A0A1B( | DIRAS2 | GTP-bin | 12703 | 39 | 2(1) | 1(1) | 0.27 | 9% | A0A1B0G |
| tr\|A8K13 | | cDNA FL | 165961 | 37 | 4(1) | 4(1) | 0.02 | 3% | A8K134_ |
| tr\|A0A02 | IMPA1 | Inositol-1 | 30568 | 37 | 2(2) | 1(1) | 0.11 | 2% | A0A024R |
| tr\|A4D27 | ARPC1B | Actin-rel | 41722 | 36 | 7(1) | 6(1) | 0.08 | 23% | A4D275_ |
| tr\|A1A50 | PRSS3 | PRSS3 p | 27040 | 36 | 1(1) | 1(1) | 0.12 | 5% | A1A508_ |
| tr\|A0A02 | DHCR7 | 7-dehydr | 55195 | 35 | 2(1) | 2(1) | 0.06 | 4% | A0A024R |
| tr\|A0A02 | hCG_19 | HCG199 | 189761 | 33 | 12(2) | 12(2) | 0.03 | 9% | A0A024R |
| tr\|L0R5A1 | CSF2RB | Alternati | 11752 | 33 | 5(2) | 1(1) | 0.29 | 7% | L0R5A1_ |
| tr\|E5RGS | CHRAC1 | Chromat | 6260 | 32 | 2(1) | 1(1) | 0.58 | 14% | E5RGS9_ |
| tr\|B4DQP | | cDNA FL | 30377 | 31 | 3(1) | 2(1) | 0.11 | 12% | B4DQP1_ |
| tr\|Q0VAS | HIST1H4 | Histone I | 11364 | 31 | 1(1) | 1(1) | 0.30 | 9% | Q0VAS5_ |

| | | | | | | | | | |
|---|---|---|---|---|---|---|---|---|---|
| sp\|P1640 | HIST1H1 | Histone I | 22336 | 30 | 85(0) | 2(0) | 0.15 | 9% | P16402_u |
| tr\|B0I1S9\| | MYO1B | MYO1B \ | 132928 | 30 | 4(0) | 4(0) | 0.02 | 3% | B0I1S9_u |
| tr\|Q6MZU | DKFZp68 | Uncharac | 51864 | 30 | 3(0) | 2(0) | 0.06 | 3% | Q6MZU6 |
| tr\|C9JFM7 | RAB43 | Ras-relat | 9491 | 28 | 3(0) | 2(0) | 0.36 | 25% | C9JFM7_u |
| tr\|A0A024 | hCG_203 | HCG203 | 81003 | 28 | 3(1) | 3(1) | 0.04 | 4% | A0A024R |
| tr\|A0A024 | VARS | VARS OS | 141642 | 28 | 1(0) | 1(0) | 0.02 | 0% | A0A024R |
| tr\|M0QW | SARS2 | Serine--t | 58602 | 26 | 3(1) | 2(1) | 0.06 | 1% | M0QWZ7 |
| tr\|A0A0K0 | HEL-S-3 | Epididym | 41498 | 25 | 3(0) | 3(0) | 0.08 | 9% | A0A0K0K |
| sp\|O1502 | MAST4 | Microtub | 286146 | 23 | 9(0) | 6(0) | 0.01 | 1% | O15021_u |
| tr\|A0A2X0 | TRIO | TRIO (Fr | 344633 | 23 | 6(0) | 5(0) | 0.01 | 1% | A0A2X0S |
| tr\|Q53R19 | ARPC2 | Arp2/3 cc | 34426 | 21 | 7(0) | 7(0) | 0.10 | 25% | Q53R19_ |
| sp\|P4945 | CENPF | Centrom | 360442 | 21 | 7(0) | 6(0) | 0.01 | 1% | P49454_u |
| tr\|A0A2R8 | AUTS2 | Autism s | 17502 | 20 | 1(0) | 1(0) | 0.19 | 5% | A0A2R8Y |
| tr\|H0YAQ | KHDRBS | KH doma | 21539 | 20 | 1(0) | 1(0) | 0.16 | 5% | H0YAQ3_ |
| tr\|A0A024 | PPP1R1 | Protein p | 85171 | 20 | 2(0) | 2(0) | 0.04 | 2% | A0A024R |
| tr\|E3Q1J2 | HLA-B | MHC clas | 31790 | 20 | 1(0) | 1(0) | 0.10 | 5% | E3Q1J2_u |
| tr\|A0A0S2 | BTRC | Beta-tran | 70248 | 19 | 1(0) | 1(0) | 0.05 | 1% | A0A0S2Z |
| sp\|O9480 | STK10 | Serine/th | 112749 | 18 | 3(0) | 3(0) | 0.03 | 3% | O94804_u |
| tr\|A0A0S2 | CYBB | Cytochro | 66206 | 18 | 2(0) | 1(0) | 0.05 | 1% | A0A0S2Z |
| tr\|V9HWC | HEL-S-1 | Epididym | 25133 | 17 | 2(0) | 2(0) | 0.13 | 7% | V9HWC7 |
| tr\|D3DP6 | LYPD1 | LY6/PLA | 9970 | 16 | 3(0) | 1(0) | 0.35 | 11% | D3DP69_ |
| sp\|Q86VH | KIF27 | Kinesin-l | 161324 | 15 | 6(0) | 4(0) | 0.02 | 2% | Q86VH2_ |

IgG

| Accessi | Gene Na | Descript | Mass | Score | Matches | Sequenc | emPAI | Coverage | |
|---|---|---|---|---|---|---|---|---|---|
| sp\|P3558 | MYH10 | Myosin-1 | 229827 | 7837 | 324(222) | 98(90) | 6.18 | 50% | P35580_ |
| sp\|Q1381 | SPTAN1 | Spectrin | 285163 | 4224 | 199(137) | 103(85) | 2.74 | 48% | Q13813_ |
| tr\|Q1KLZ( | PS1TP5I | HCG159 | 42052 | 2927 | 163(108) | 18(15) | 11.13 | 61% | Q1KLZ0_ |
| tr\|B2ZZ89 | SPTBN1 | Spectrin | 275237 | 2482 | 154(88) | 96(74) | 1.53 | 50% | B2ZZ89_ |
| sp\|P7852 | PRKDC | DNA-dep | 473749 | 2070 | 128(65) | 92(58) | 0.62 | 26% | P78527_ |
| tr\|A0A2R | ACTB | Actin, cy | 17631 | 1371 | 66(44) | 6(6) | 10.61 | 70% | A0A2R8Y |
| sp\|Q1420 | DYNC1H | Cytoplas | 534809 | 1236 | 94(51) | 79(47) | 0.40 | 21% | Q14204_ |
| sp\|Q0966 | AHNAK | Neurobla | 629213 | 1145 | 117(55) | 94(49) | 0.36 | 22% | Q09666_ |
| sp\|Q562F | ACTBL2 | Beta-acti | 42318 | 996 | 66(30) | 6(3) | 0.69 | 16% | Q562R1_ |
| tr\|A0A08 | CLTC | Clathrin I | 193703 | 939 | 39(27) | 31(23) | 0.60 | 24% | A0A087W |
| tr\|A0A2R | ACTB | Actin, cy | 8839 | 745 | 51(25) | 8(3) | 9.21 | 65% | A0A2R8Y |
| tr\|A0A02 | hCG_19 | HCG199 | 189761 | 644 | 43(27) | 36(25) | 0.56 | 27% | A0A024R |
| sp\|Q1514 | PLEC | Plectin C | 533462 | 583 | 57(25) | 55(24) | 0.19 | 13% | Q15149_ |
| sp\|P4932 | FASN | Fatty aci | 275877 | 571 | 40(22) | 33(18) | 0.33 | 18% | P49327_ |
| tr\|A0A0S | KRT6A | HCG203 | 60293 | 558 | 37(22) | 21(17) | 1.60 | 37% | A0A0S2Z |
| sp\|Q7Z40 | MYH14 | Myosin-1 | 228701 | 556 | 43(25) | 19(13) | 0.24 | 10% | Q7Z406_ |
| tr\|A0A07 | TRAJ56 | T cell rec | 2220 | 516 | 90(73) | 1(1) | 1.81 | 38% | A0A075B |
| tr\|B4DKV | | cDNA FL | 55983 | 506 | 33(20) | 18(15) | 1.50 | 33% | B4DKV4_ |
| tr\|A0A1W | | Uncharac | 77022 | 476 | 34(17) | 15(12) | 0.79 | 29% | A0A1W2I |
| tr\|Q5TZZ | ANXA1 | Annexin | 38918 | 435 | 12(11) | 9(8) | 0.92 | 38% | Q5TZZ9_ |
| tr\|Q6IPN | EEF1A1 | Elongatic | 50433 | 382 | 23(16) | 8(7) | 0.88 | 25% | Q6IPN6_ |
| tr\|B4DL3 | | cDNA FL | 27409 | 345 | 16(12) | 10(8) | 2.15 | 39% | B4DL32_ |
| tr\|V9HVZ | HEL-S-1 | Glyceral | 36201 | 335 | 15(8) | 11(7) | 0.85 | 47% | V9HVZ4_ |
| sp\|P6116 | ACTR2 | Actin-rel | 45017 | 327 | 18(11) | 12(9) | 1.03 | 31% | P61160_ |
| tr\|B3KPS | | Tubulin a | 46725 | 290 | 15(7) | 10(7) | 0.85 | 32% | B3KPS3_ |
| tr\|B4E2J5 | | cDNA FL | 21684 | 269 | 13(9) | 4(4) | 0.78 | 26% | B4E2J5_ |
| tr\|Q8IWP | | Tubulin b | 50177 | 253 | 15(9) | 9(9) | 0.77 | 25% | Q8IWP6_ |
| tr\|A0A38 | | Fascin O | 55123 | 232 | 13(8) | 9(6) | 0.59 | 22% | A0A384N |
| tr\|A0A08 | TJP1 | Tight jun | 188076 | 225 | 29(10) | 22(10) | 0.21 | 17% | A0A087X |
| tr\|E1NZA | PRIC295 | Peroxiso | 294953 | 221 | 19(11) | 19(11) | 0.14 | 8% | E1NZA1_ |
| sp\|P1592 | DSP | Desmopl | 334021 | 206 | 22(7) | 19(7) | 0.09 | 7% | P15924_ |
| tr\|F8WCF | ARPC4- | Actin-rel | 21159 | 204 | 9(8) | 5(5) | 1.09 | 33% | F8WCF6_ |
| tr\|Q60FE | FLNA | Filamin-A | 280790 | 193 | 25(8) | 24(8) | 0.12 | 12% | Q60FE5_ |
| tr\|A0A02 | ACTR3 | ARP3 ac | 47797 | 185 | 18(7) | 12(6) | 0.49 | 32% | A0A024R |
| tr\|A0A02 | ANXA2 | Annexin | 38808 | 183 | 14(7) | 9(6) | 0.63 | 32% | A0A024R |
| tr\|B7Z2X | | cDNA FL | 78026 | 180 | 9(7) | 6(5) | 0.28 | 9% | B7Z2X4_ |
| tr\|A0A02 | JUP | Junction | 82416 | 178 | 4(3) | 3(2) | 0.12 | 6% | A0A024R |
| tr\|Q1466 | | Radiated | 30933 | 163 | 12(6) | 8(5) | 0.66 | 29% | Q14666_ |
| tr\|Q6I9V | SLC25A | Epididym | 33073 | 151 | 10(6) | 6(5) | 0.61 | 23% | Q6I9V5_ |
| tr\|A0A02 | HSP90A | Heat sho | 83554 | 142 | 9(5) | 8(4) | 0.21 | 13% | A0A024R |
| sp\|P2262 | HNRNPA | Heteroge | 37464 | 135 | 7(3) | 5(2) | 0.18 | 19% | P22626_ |
| tr\|Q53R1 | ARPC2 | Arp2/3 c | 34426 | 133 | 11(5) | 9(5) | 0.74 | 29% | Q53R19_ |
| tr\|Q7L4M | | KRT8 pr | 30802 | 132 | 5(4) | 4(3) | 0.36 | 15% | Q7L4M3_ |
| tr\|Q0P51 | SLC2A1 | Solute ca | 54361 | 131 | 8(5) | 6(4) | 0.27 | 10% | Q0P512_ |
| tr\|B2R4D | | Actin-rel | 20735 | 131 | 8(5) | 5(5) | 1.12 | 28% | B2R4D5_ |
| tr\|J3KT12 | EIF4A1 | Eukaryot | 39065 | 125 | 8(6) | 7(6) | 0.77 | 22% | J3KT12_ |
| tr\|A0A02 | HIST1H2 | Histone I | 13928 | 116 | 3(1) | 2(1) | 0.25 | 19% | A0A024C |
| tr\|A0A02 | SLC25A | Solute ca | 40389 | 112 | 9(4) | 6(4) | 0.37 | 22% | A0A024R |
| sp\|P5362 | COPA | Coatome | 139797 | 111 | 17(4) | 14(4) | 0.10 | 14% | P53621_ |
| sp\|Q6P2 | PRPF8 | Pre-mRN | 274738 | 109 | 11(4) | 11(4) | 0.06 | 5% | Q6P2Q9_ |
| tr\|G3V1A | CFL1 | Cofilin 1 | 17029 | 107 | 6(4) | 4(3) | 0.72 | 35% | G3V1A4_ |
| tr\|B4DE5 | | cDNA FL | 62862 | 104 | 11(3) | 6(2) | 0.17 | 14% | B4DE59_ |
| tr\|A0A02 | NUMA1 | Nuclear | 237456 | 103 | 13(2) | 13(2) | 0.06 | 8% | A0A024R |
| tr\|G3V1V | MYL6 | Myosin li | 18311 | 98 | 4(4) | 4(4) | 0.97 | 31% | G3V1V0_ |
| sp\|Q9ULE | FAM184 | Protein F | 121882 | 93 | 8(3) | 4(1) | 0.05 | 3% | Q9ULE4_ |

| | | | | | | | | | |
|---|---|---|---|---|---|---|---|---|---|
| sp\|O7536 | FLNB | Filamin-E | 280157 | 90 | 21(3) | 20(3) | 0.06 | 9% | O75369_ |
| tr\|A0A02 | CORO1C | Coronin | 53899 | 89 | 6(2) | 5(2) | 0.13 | 12% | A0A024R |
| tr\|A0A14 | | Testicula | 34465 | 84 | 8(3) | 8(3) | 0.58 | 33% | A0A140V |
| tr\|B2R5W | HNRNPC | Heteroge | 31986 | 82 | 8(3) | 6(3) | 0.35 | 18% | B2R5W2_ |
| tr\|A0A02 | PKM2 | Pyruvate | 58538 | 81 | 7(3) | 6(3) | 0.24 | 15% | A0A024R |
| tr\|B4DDH | | cDNA FL | 51214 | 80 | 5(3) | 4(3) | 0.21 | 12% | B4DDH8_ |
| tr\|B3KSQ | | cDNA FL | 37956 | 77 | 5(2) | 4(1) | 0.09 | 11% | B3KSQ1_ |
| tr\|B0I1S9\| | MYO1B | MYO1B | 132928 | 76 | 15(6) | 14(6) | 0.16 | 13% | B0I1S9_u |
| tr\|A0A02 | HUWE1 | HECT, U | 484651 | 72 | 11(4) | 11(4) | 0.03 | 3% | A0A024R |
| tr\|A0A02 | hCG_20 | HCG203 | 81003 | 70 | 5(2) | 5(2) | 0.13 | 7% | A0A024R |
| sp\|Q58FF | HSP90A | Putative | 58855 | 69 | 3(3) | 2(2) | 0.11 | 4% | Q58FF6_ |
| tr\|H3BPE | MACF1 | Microtub | 861649 | 69 | 31(2) | 29(2) | 0.02 | 4% | H3BPE1_ |
| sp\|P0DO | | Immunog | 49925 | 66 | 10(3) | 2(1) | 0.07 | 3% | P0DOX5_ |
| sp\|Q5D8 | FLG2 | Filaggrin | 249296 | 60 | 3(1) | 2(1) | 0.01 | 0% | Q5D862_ |
| tr\|B2RE4 | | cDNA, FI | 69405 | 60 | 5(3) | 5(3) | 0.15 | 12% | B2RE46_ |
| tr\|V9HW | HEL-S-1 | ATP synt | 59828 | 58 | 6(2) | 5(1) | 0.05 | 9% | V9HW26_ |
| tr\|B2R7I0 | | cDNA, FI | 34344 | 56 | 1(1) | 1(1) | 0.10 | 5% | B2R7I0_u |
| tr\|H3BMZ | HACD3 | Very-long | 16267 | 55 | 1(1) | 1(1) | 0.21 | 10% | H3BMZ1_ |
| tr\|A0A0A | DHCR24 | Delta(24) | 62205 | 55 | 2(1) | 2(1) | 0.11 | 4% | A0A0A0N |
| sp\|P2135 | NF1 | Neurofib | 322760 | 55 | 4(1) | 4(1) | 0.02 | 1% | P21359_ |
| tr\|Q6E43 | | Activated | 8649 | 54 | 1(1) | 1(1) | 0.40 | 15% | Q6E433_ |
| tr\|B1ALC | ARPC5 | Actin-rel | 14855 | 54 | 2(1) | 2(1) | 0.23 | 23% | B1ALC0_ |
| tr\|M0R1V | UBA52 | Ubiquitin | 7128 | 52 | 2(1) | 2(1) | 1.25 | 34% | M0R1V7_ |
| sp\|P1227 | PIP | Prolactin | 16847 | 52 | 2(1) | 2(1) | 0.20 | 13% | P12273_ |
| tr\|A0A02 | PKP1 | Plakophi | 81757 | 51 | 2(1) | 2(1) | 0.04 | 3% | A0A024R |
| tr\|H0YKU | TMOD3 | Tropomo | 20946 | 51 | 2(2) | 2(2) | 0.35 | 12% | H0YKU1_ |
| tr\|V9HW | HEL-S-1 | Epididym | 25133 | 50 | 5(1) | 3(1) | 0.13 | 12% | V9HWC7 |
| tr\|A0A0S | HM13 | Histocon | 41747 | 49 | 8(2) | 5(1) | 0.08 | 17% | A0A0S2Z |
| sp\|P8160 | DCD | Dermcidi | 11391 | 49 | 3(1) | 2(1) | 0.69 | 12% | P81605_ |
| tr\|A0A08 | ALB | Serum al | 46430 | 49 | 3(2) | 2(1) | 0.07 | 6% | A0A087V |
| sp\|Q0821 | DHX9 | ATP-dep | 142181 | 49 | 6(2) | 6(2) | 0.05 | 6% | Q08211_ |
| tr\|A0A02 | DHCR7 | 7-dehydr | 55195 | 49 | 5(1) | 4(1) | 0.19 | 11% | A0A024R |
| tr\|B3KTV | | cDNA FL | 68166 | 49 | 4(3) | 4(3) | 0.15 | 8% | B3KTV0_ |
| sp\|P1640 | HIST1H1 | Histone | 22336 | 47 | 92(1) | 3(1) | 0.15 | 14% | P16402_ |
| tr\|A0A02 | hCG_27 | HCG271 | 232294 | 46 | 8(1) | 8(1) | 0.03 | 3% | A0A024C |
| tr\|A0A02 | ENO1 | Enolase | 47481 | 46 | 1(1) | 1(1) | 0.07 | 3% | A0A024R |
| tr\|A0A02 | CTTN | Cortactin | 57603 | 45 | 3(1) | 3(1) | 0.06 | 7% | A0A024R |
| tr\|C9JPM | ARF4 | ADP-ribo | 14601 | 45 | 3(1) | 3(1) | 0.23 | 22% | C9JPM4_ |
| tr\|A0A02 | RPL6 | 60S ribo | 32765 | 44 | 4(1) | 4(1) | 0.21 | 16% | A0A024R |
| tr\|B3KSW | CDIPT | CDP-dia | 20575 | 43 | 1(1) | 1(1) | 0.16 | 7% | B3KSW0_ |
| tr\|A0A02 | DLST | Dihydroli | 49067 | 43 | 2(2) | 1(1) | 0.07 | 1% | A0A024R |
| sp\|P3115 | S100A7 | Protein S | 11578 | 43 | 1(1) | 1(1) | 0.30 | 10% | P31151_ |
| tr\|A0A0A | PRDX1 | Peroxire | 10727 | 43 | 3(1) | 3(1) | 0.74 | 44% | A0A0A0N |
| tr\|A0A1B | DIRAS2 | GTP-bin | 12703 | 42 | 2(1) | 1(1) | 0.27 | 9% | A0A1B0G |
| tr\|Q6IPH | RPL14 | RPL14 p | 23886 | 42 | 2(1) | 2(1) | 0.14 | 10% | Q6IPH7_ |
| tr\|L0R5A1 | CSF2RB | Alternati | 11752 | 42 | 5(3) | 1(1) | 0.29 | 7% | L0R5A1_ |
| tr\|A0A0S | ATP2A2 | ATPase | 116336 | 42 | 4(1) | 4(1) | 0.03 | 4% | A0A0S2Z |
| tr\|A0A0B | LYZ | Lysozym | 15661 | 41 | 1(1) | 1(1) | 0.22 | 8% | A0A0B4J |
| tr\|A8K48 | | Peptidyl- | 18230 | 41 | 4(1) | 3(1) | 0.41 | 18% | A8K486_ |
| tr\|A0A02 | VDAC2 | Voltage- | 35029 | 41 | 3(1) | 3(1) | 0.09 | 14% | A0A024C |
| tr\|A0A02 | CAPZA1 | F-actin-c | 33073 | 40 | 2(1) | 2(1) | 0.10 | 9% | A0A024R |
| sp\|Q5T74 | KPRP | Keratino | 67172 | 40 | 3(2) | 2(1) | 0.05 | 2% | Q5T749_ |
| sp\|P0DM | HSPA1A | Heat sho | 70294 | 40 | 3(1) | 3(1) | 0.10 | 5% | P0DMV8_ |
| tr\|H7C3J | SAMD11 | Sterile al | 57548 | 39 | 2(1) | 1(1) | 0.06 | 1% | H7C3J6_ |
| tr\|C9JFM | RAB43 | Ras-rela | 9491 | 39 | 3(0) | 2(0) | 0.36 | 25% | C9JFM7_ |
| sp\|Q0241 | DSG1 | Desmogl | 114702 | 39 | 3(1) | 3(1) | 0.03 | 4% | Q02413_ |
| tr\|A0A02 | IMPA1 | Inositol-1 | 30568 | 38 | 2(2) | 1(1) | 0.11 | 2% | A0A024R |
| tr\|J3QRN | MYO1D | Unconve | 111903 | 38 | 4(1) | 4(1) | 0.03 | 4% | J3QRN6_ |

| | | | | | | | | | |
|---|---|---|---|---|---|---|---|---|---|
| tr\|U3KPS! | TPI1 | Trioseph | 4245 | 37 | 1(1) | 1(1) | 0.88 | 36% | U3KPS5_ |
| tr\|A0A28( | ASAP2 | Arf-GAP | 37064 | 37 | 7(2) | 2(1) | 0.09 | 5% | A0A286Y |
| tr\|A4FU7: | SNRNP2 | SNRNP2 | 217585 | 37 | 11(1) | 10(1) | 0.03 | 6% | A4FU77_ |
| sp\|Q5T4S | UBR4 | E3 ubiqu | 580547 | 36 | 6(1) | 6(1) | 0.01 | 1% | Q5T4S7_ |
| tr\|M0QW | SARS2 | Serine--t | 58602 | 36 | 2(1) | 1(1) | 0.06 | 1% | M0QWZ7 |
| tr\|A0A14( | HLA-DOI | HLA clas | 20866 | 35 | 2(1) | 2(1) | 0.16 | 8% | A0A140T |
| tr\|A0A38 | | Epididym | 49517 | 35 | 2(1) | 2(1) | 0.07 | 5% | A0A384N |
| tr\|Q53YD | EEF1G | EEF1G p | 50429 | 34 | 5(1) | 5(1) | 0.07 | 13% | Q53YD7_ |
| tr\|J3KQE! | RAN | GTP-bin | 27027 | 34 | 2(1) | 2(1) | 0.12 | 8% | J3KQE5_ |
| tr\|F8VPD | CAD | CAD pro | 238097 | 34 | 7(1) | 7(1) | 0.01 | 4% | F8VPD4_ |
| tr\|A0A1W | HNRNPU | Heteroge | 81283 | 34 | 2(1) | 2(1) | 0.04 | 3% | A0A1W2I |
| sp\|Q6P1/ | LPCAT3 | Lysopho | 56511 | 33 | 3(1) | 3(1) | 0.06 | 6% | Q6P1A2_ |
| tr\|A0A38 | | Epididym | 96246 | 33 | 1(1) | 1(1) | 0.03 | 1% | A0A384N |
| sp\|Q5T75 | XP32 | Skin-spe | 28557 | 33 | 2(1) | 1(1) | 0.12 | 3% | Q5T750_ |
| tr\|J3KTA4 | DDX5 | Probable | 69557 | 33 | 3(1) | 3(1) | 0.05 | 4% | J3KTA4_ |
| sp\|P6285 | RPS25 | 40S ribo | 13791 | 33 | 1(1) | 1(1) | 0.25 | 8% | P62851_ |
| tr\|A0A1W | | N90-VRC | 13625 | 33 | 2(1) | 1(1) | 0.25 | 15% | A0A1W6I |
| sp\|Q9Y5E | SUPT16I | FACT co | 120409 | 32 | 2(1) | 2(1) | 0.03 | 2% | Q9Y5B9_ |
| tr\|J3KPX7 | PHB2 | Prohibitir | 33219 | 32 | 2(1) | 2(1) | 0.10 | 7% | J3KPX7_ |
| tr\|E5RGS! | CHRAC1 | Chromat | 6260 | 32 | 2(1) | 1(1) | 0.58 | 14% | E5RGS9_ |
| sp\|P0C0S | HIST1H2 | Histone I | 14083 | 31 | 3(0) | 2(0) | 0.24 | 21% | P0C0S8_ |
| tr\|A0A0A | IARS | Isoleucin | 143509 | 31 | 1(1) | 1(1) | 0.02 | 0% | A0A0A0N |
| tr\|A0A0K( | HEL-S-2 | Epididym | 16017 | 31 | 2(1) | 2(1) | 0.21 | 19% | A0A0K0K |
| tr\|A0A1B( | RBMXL1 | RNA-bin | 16783 | 31 | 1(0) | 1(0) | 0.20 | 8% | A0A1B0G |
| tr\|A0A02 | DOLPP1 | Dolichyl | 27127 | 30 | 1(1) | 1(1) | 0.12 | 5% | A0A024R |
| tr\|B4DQP | | cDNA FL | 30377 | 30 | 2(1) | 1(1) | 0.11 | 3% | B4DQP1_ |
| tr\|D0PNI1 | YWHAZ | Epididym | 27899 | 29 | 2(0) | 2(0) | 0.12 | 9% | D0PNI1_ |
| tr\|A0A49 | | Unchara | 79206 | 29 | 1(0) | 1(0) | 0.04 | 1% | A0A494C |
| tr\|Q5K63 | | SCCA2/S | 44677 | 29 | 3(1) | 3(1) | 0.07 | 6% | Q5K634_ |
| tr\|A0A02 | hCG_20( | DDX39B | 49416 | 28 | 4(0) | 3(0) | 0.14 | 7% | A0A024R |
| tr\|Q6ZTZ | | cDNA FL | 22481 | 28 | 2(0) | 1(0) | 0.15 | 4% | Q6ZTZ2_ |
| tr\|A0A02 | TMEM33 | Transme | 28302 | 28 | 2(0) | 2(0) | 0.12 | 6% | A0A024R |
| tr\|E9PR3( | FAU | 40S ribo | 10898 | 27 | 1(0) | 1(0) | 0.32 | 10% | E9PR30_ |
| tr\|A0A38 | | Epididym | 39026 | 27 | 2(1) | 2(1) | 0.08 | 6% | A0A384N |
| sp\|P1640 | HIST1H1 | Histone I | 22566 | 27 | 1(0) | 1(0) | 0.15 | 4% | P16401_ |
| tr\|A0A02 | DERL1 | Derlin OS | 28896 | 26 | 1(0) | 1(0) | 0.12 | 4% | A0A024R |
| tr\|B7Z3V | | Sodium/p | 113791 | 25 | 5(0) | 5(0) | 0.03 | 5% | B7Z3V1_ |
| tr\|Q6NZ5 | RPL27A | Ribosom | 16582 | 25 | 2(0) | 2(0) | 0.20 | 14% | Q6NZ52_ |
| tr\|A0A02 | RPS9 | 40S ribo | 22635 | 25 | 1(0) | 1(0) | 0.15 | 4% | A0A024R |
| tr\|C9JNW | RPL24 | 60S ribo | 17646 | 25 | 1(0) | 1(0) | 0.19 | 8% | C9JNW5_ |
| tr\|S4R2X2 | SFXN1 | Siderofle | 19962 | 24 | 1(0) | 1(0) | 0.17 | 7% | S4R2X2_ |
| tr\|A0A02 | HNRPK | Heteroge | 51472 | 23 | 1(0) | 1(0) | 0.06 | 3% | A0A024R |
| tr\|B4DZX! | | cDNA FL | 19688 | 22 | 2(0) | 1(0) | 0.17 | 6% | B4DZX5_ |
| tr\|A8K58 | | cDNA FL | 39578 | 21 | 5(0) | 3(0) | 0.08 | 7% | A8K588_ |
| tr\|B2RU0( | EIF4G1 | eIF4G1 p | 176807 | 21 | 3(0) | 3(0) | 0.02 | 2% | B2RU06_ |
| tr\|A0A02 | RECQL | ATP-dep | 74436 | 21 | 1(0) | 1(0) | 0.04 | 1% | A0A024R |
| tr\|A2J1N8 | | Rheumat | 10316 | 21 | 2(0) | 1(0) | 0.34 | 14% | A2J1N8_ |
| sp\|A2RUF | CCDC14 | Coiled-cc | 166448 | 20 | 3(0) | 3(0) | 0.02 | 2% | A2RUR9_ |
| tr\|A0A14( | | Testicula | 57499 | 20 | 1(0) | 1(0) | 0.06 | 1% | A0A140V |
| tr\|A0A2Z( | DBN1 | Drebrin / | 76890 | 19 | 4(0) | 4(0) | 0.04 | 10% | A0A2Z6A |
| tr\|Q5VVC | RPL11 | 60S ribo | 19240 | 19 | 1(0) | 1(0) | 0.18 | 8% | Q5VVC8_ |
| tr\|Q0VAS | HIST1H4 | Histone I | 11364 | 19 | 1(0) | 1(0) | 0.30 | 9% | Q0VAS5_ |
| tr\|B4DR6( | | cDNA FL | 119789 | 19 | 4(1) | 2(1) | 0.03 | 1% | B4DR60_ |
| sp\|A6NKI | TMEM2C | Transme | 64345 | 19 | 1(0) | 1(0) | 0.05 | 1% | A6NKL6_ |
| tr\|A0A02 | NNT | Nicotinar | 114564 | 17 | 3(0) | 2(0) | 0.03 | 1% | A0A024R |
| tr\|F4MH5 | UTY | Ubiquitou | 115990 | 16 | 2(0) | 1(0) | 0.03 | 1% | F4MH59_ |
| tr\|G3V1J( | WDR55 | WD repe | 24828 | 15 | 4(0) | 2(0) | 0.13 | 4% | G3V1J0_ |
| sp\|Q86VI | KIF27 | Kinesin-l | 161324 | 15 | 5(0) | 3(0) | 0.02 | 1% | Q86VH2_ |

| | | | | | | | | | |
|---|---|---|---|---|---|---|---|---|---|
| tr\|A0A2R8 | LRRC9 | Leucine- | 173242 | 15 | 4(0) | 2(0) | 0.02 | 1% | A0A2R8Y |
| tr\|A0A0S2 | L3MBTL2 | Lmbt-like | 72762 | 15 | 2(0) | 1(0) | 0.05 | 1% | A0A0S2Z |

shNAT10

| Accessi | Gene Na | Descript | Mass | Score | Matches | Sequenc | emPAI | Coverage | |
|---|---|---|---|---|---|---|---|---|---|
| sp\|P3558 | MYH10 | Myosin-1 | 229827 | 2179 | 124(69) | 70(49) | 1.39 | 37% | P35580_u |
| tr\|Q1KLZ( | PS1TP5I | HCG159 | 42052 | 888 | 49(31) | 10(9) | 2.35 | 39% | Q1KLZ0_u |
| tr\|A0A0S2 | KRT6A | HCG203 | 60293 | 513 | 35(21) | 21(15) | 1.34 | 37% | A0A0S2Z |
| tr\|B4DKV4 | | cDNA FL | 55983 | 409 | 30(16) | 17(11) | 1.10 | 32% | B4DKV4_ |
| tr\|A0A2R8 | ACTB | Actin, cyt | 8839 | 401 | 14(10) | 3(3) | 9.21 | 50% | A0A2R8Y |
| tr\|A0A075 | TRAJ56 | T cell rec | 2220 | 327 | 106(48) | 1(1) | 1.81 | 38% | A0A075B |
| sp\|Q9H0A | NAT10 | RNA cyti | 116569 | 324 | 27(12) | 18(10) | 0.39 | 21% | Q9H0A0_ |
| tr\|B4DL32 | | cDNA FL | 27409 | 304 | 21(10) | 10(6) | 1.81 | 40% | B4DL32_u |
| sp\|P0CG3 | POTEI | POTE ar | 122858 | 302 | 16(10) | 5(3) | 0.17 | 4% | P0CG38_ |
| sp\|Q1514 | PLEC | Plectin C | 533462 | 290 | 39(11) | 38(11) | 0.10 | 10% | Q15149_u |
| tr\|B4E2J5 | | cDNA FL | 21684 | 245 | 12(7) | 6(4) | 0.78 | 35% | B4E2J5_u |
| sp\|Q0966 | AHNAK | Neurobla | 629213 | 201 | 64(10) | 53(9) | 0.06 | 12% | Q09666_u |
| tr\|Q6IPN( | EEF1A1 | Elongatic | 50433 | 177 | 9(7) | 6(5) | 0.37 | 17% | Q6IPN6_u |
| sp\|P7852 | PRKDC | DNA-dep | 473749 | 173 | 26(8) | 25(8) | 0.06 | 7% | P78527_u |
| tr\|Q05CW | NAT10 | NAT10 p | 63029 | 166 | 14(6) | 9(5) | 0.36 | 19% | Q05CW7_ |
| tr\|V9HVZ | HEL-S-1 | Glyceral | 36201 | 156 | 8(7) | 6(5) | 0.69 | 27% | V9HVZ4_ |
| tr\|A0A08 | SPTBN1 | Spectrin | 275457 | 155 | 19(6) | 19(6) | 0.10 | 9% | A0A087W |
| tr\|Q1466( | | Radiated | 30933 | 145 | 10(4) | 7(3) | 0.50 | 22% | Q14666_u |
| tr\|B4DE59 | | cDNA FL | 62862 | 121 | 13(4) | 8(3) | 0.17 | 15% | B4DE59_u |
| tr\|A0A08 | CLTC | Clathrin I | 193703 | 117 | 14(3) | 12(3) | 0.14 | 10% | A0A087W |
| tr\|A0A024 | SPTAN1 | Spectrin, | 282874 | 106 | 20(3) | 20(3) | 0.05 | 9% | A0A024R |
| sp\|Q0241 | DSG1 | Desmogl | 114702 | 105 | 6(3) | 5(3) | 0.09 | 7% | Q02413_u |
| tr\|A0A024 | ANXA2 | Annexin | 38808 | 85 | 7(2) | 6(2) | 0.18 | 23% | A0A024R |
| sp\|Q5D8( | FLG2 | Filaggrin | 249296 | 80 | 1(1) | 1(1) | 0.01 | 0% | Q5D862_ |
| sp\|Q1420 | DYNC1H | Cytoplas | 534809 | 79 | 25(4) | 23(4) | 0.03 | 6% | Q14204_u |
| tr\|Q147W | KRT1B | KRT1B p | 37557 | 79 | 3(3) | 3(3) | 0.29 | 9% | Q147W7_ |
| tr\|A0A024 | ACTN1 | Actinin, a | 103563 | 77 | 4(2) | 4(2) | 0.10 | 5% | A0A024R |
| tr\|B3KPS3 | | Tubulin a | 46725 | 71 | 8(2) | 7(2) | 0.23 | 23% | B3KPS3_u |
| tr\|H7C0A | ARPC4-T | ARPC4-T | 19558 | 64 | 4(3) | 3(2) | 0.37 | 17% | H7C0A3_ |
| tr\|V9HW2 | HEL-S-1 | ATP synt | 59828 | 59 | 3(1) | 1(1) | 0.05 | 2% | V9HW26_ |
| sp\|P0510 | S100A8 | Protein S | 10885 | 56 | 1(1) | 1(1) | 0.32 | 11% | P05109_u |
| tr\|Q6NVC | SLC25A5 | SLC25A5 | 35499 | 54 | 9(1) | 6(1) | 0.20 | 20% | Q6NVC0_ |
| tr\|A4D275 | ARPC1B | Actin-rela | 41722 | 53 | 2(1) | 2(1) | 0.08 | 7% | A4D275_ |
| tr\|B4DM6 | | Actin-rela | 19220 | 52 | 3(1) | 3(1) | 0.18 | 18% | B4DM63_ |
| sp\|P4932 | FASN | Fatty aci | 275877 | 51 | 13(2) | 13(2) | 0.02 | 6% | P49327_u |
| sp\|P6116 | ACTR2 | Actin-rela | 45017 | 51 | 7(2) | 6(2) | 0.24 | 19% | P61160_u |
| tr\|A0A0K( | HEL-S-3 | Epididym | 41498 | 51 | 2(2) | 2(2) | 0.17 | 6% | A0A0K0K |
| tr\|A0A024 | SLC25A3 | Solute ca | 40389 | 50 | 2(1) | 2(1) | 0.17 | 7% | A0A024R |
| tr\|A0A024 | hCG_19 | HCG199 | 189761 | 50 | 11(3) | 11(3) | 0.05 | 9% | A0A024R |
| sp\|P8160 | DCD | Dermcidi | 11391 | 49 | 2(2) | 1(1) | 0.30 | 10% | P81605_u |
| tr\|A0A0B4 | LYZ | Lysozym | 15661 | 49 | 1(1) | 1(1) | 0.22 | 8% | A0A0B4J |
| tr\|Q5TZZ | ANXA1 | Annexin | 38918 | 46 | 3(1) | 3(1) | 0.08 | 10% | Q5TZZ9_ |
| sp\|Q9HC | ZSWIM6 | Zinc fing | 135096 | 46 | 1(1) | 1(1) | 0.02 | 0% | Q9HCJ5_ |
| tr\|Q53R1 | ARPC2 | Arp2/3 c | 34426 | 45 | 5(1) | 5(1) | 0.10 | 16% | Q53R19_ |
| tr\|A0A0C | ALB | Serum al | 71177 | 44 | 9(2) | 7(1) | 0.05 | 10% | A0A0C4D |
| tr\|J3KSD8 | BLMH | Bleomyc | 30529 | 44 | 2(1) | 2(1) | 0.11 | 11% | J3KSD8_u |
| tr\|A0A024 | CLPX | ClpX cas | 69922 | 43 | 2(1) | 2(1) | 0.05 | 3% | A0A024R |
| tr\|H9ZYJ2 | TXN | Thioredo | 12015 | 41 | 1(1) | 1(1) | 0.29 | 12% | H9ZYJ2_u |
| tr\|Q0VAS | HIST1H4 | Histone H | 11364 | 41 | 1(1) | 1(1) | 0.30 | 9% | Q0VAS5_ |
| tr\|A0A1B( | DIRAS2 | GTP-bin | 12703 | 40 | 2(1) | 1(1) | 0.27 | 9% | A0A1B0G |
| tr\|Q0P51 | SLC2A1 | Solute ca | 54361 | 40 | 2(1) | 2(1) | 0.06 | 3% | Q0P512_ |
| tr\|L0R5A1 | CSF2RB | Alternati | 11752 | 39 | 5(3) | 1(1) | 0.29 | 7% | L0R5A1_u |
| sp\|Q5T74 | KPRP | Keratino | 67172 | 39 | 11(1) | 6(1) | 0.05 | 17% | Q5T749_u |
| tr\|Q8IWP | | Tubulin b | 50177 | 38 | 8(1) | 5(1) | 0.07 | 14% | Q8IWP6_ |
| tr\|A0A384 | | Epididym | 36494 | 38 | 3(1) | 3(1) | 0.19 | 9% | A0A384N |

| | | | | | | | | | |
|---|---|---|---|---|---|---|---|---|---|
| tr\|A0A024 | ACTR3 | ARP3 ac | 47797 | 37 | 6(1) | 5(1) | 0.07 | 10% | A0A024R |
| sp\|P3115 | S100A7 | Protein S | 11578 | 36 | 3(1) | 3(1) | 0.30 | 22% | P31151_u |
| tr\|B2CIS9 | CASP14 | Caspase | 27947 | 36 | 5(1) | 4(1) | 0.12 | 18% | B2CIS9_u |
| tr\|A8K7F6 | | cDNA FL | 46321 | 36 | 4(1) | 4(1) | 0.07 | 12% | A8K7F6_u |
| tr\|A8K008 | | Uncharac | 52248 | 36 | 3(1) | 2(1) | 0.06 | 3% | A8K008_u |
| tr\|B4DQP | | cDNA FL | 30377 | 35 | 2(1) | 1(1) | 0.11 | 3% | B4DQP1_ |
| tr\|V9HWC | HEL-S-1 | Epididym | 25133 | 34 | 3(1) | 2(1) | 0.13 | 7% | V9HWC7 |
| tr\|Q7L4M | | KRT8 pro | 30802 | 33 | 4(1) | 3(1) | 0.11 | 11% | Q7L4M3_ |
| tr\|A0A024 | IMPA1 | Inositol-1 | 30568 | 32 | 2(1) | 1(1) | 0.11 | 2% | A0A024R |
| sp\|Q5T75 | XP32 | Skin-spe | 28557 | 32 | 2(1) | 1(1) | 0.12 | 3% | Q5T750_u |
| tr\|J3KPX7 | PHB2 | Prohibitin | 33219 | 32 | 1(1) | 1(1) | 0.10 | 3% | J3KPX7_u |
| tr\|B2R5W | HNRNPC | Heteroge | 31986 | 31 | 2(1) | 2(1) | 0.10 | 6% | B2R5W2_ |
| sp\|Q5T89 | ERCC6L | DNA exc | 178955 | 31 | 1(0) | 1(0) | 0.02 | 0% | Q5T890_u |
| tr\|A0A140 | | Testicula | 34465 | 28 | 4(1) | 3(1) | 0.20 | 11% | A0A140V |
| tr\|A0A024 | DLST | Dihydroli | 49067 | 26 | 1(1) | 1(1) | 0.07 | 1% | A0A024R |
| tr\|A0A140 | | Testicula | 62232 | 26 | 1(0) | 1(0) | 0.05 | 2% | A0A140V |
| tr\|H7C4W | FSTL1 | Follistatin | 2420 | 26 | 2(0) | 1(0) | 1.61 | 40% | H7C4W4_ |
| sp\|H7BZ5 | CROCC2 | Putative | 186900 | 24 | 2(0) | 2(0) | 0.02 | 1% | H7BZ55_u |
| tr\|M0QW | SARS2 | Serine--t | 58602 | 23 | 3(1) | 2(1) | 0.06 | 1% | M0QWZ7 |
| tr\|A0A024 | PKM2 | Pyruvate | 58538 | 21 | 4(0) | 4(0) | 0.06 | 10% | A0A024R |
| sp\|Q2TAC | KIF19 | Kinesin-li | 111834 | 19 | 1(0) | 1(0) | 0.03 | 1% | Q2TAC6_ |
| tr\|Q5T8U | SURF4 | Surfeit 4 | 21342 | 19 | 1(0) | 1(0) | 0.16 | 6% | Q5T8U5_ |
| tr\|D3DP6 | LYPD1 | LY6/PLA | 9970 | 19 | 3(0) | 1(0) | 0.35 | 11% | D3DP69_ |
| tr\|A0A1W | HNRNPU | Heteroge | 81283 | 19 | 3(0) | 3(0) | 0.04 | 6% | A0A1W2I |
| sp\|Q6UVI | CSPG4 | Chondro | 251067 | 18 | 5(0) | 4(0) | 0.01 | 2% | Q6UVK1_ |
| tr\|A0A087 | GALC | Galactoc | 45641 | 18 | 4(0) | 4(0) | 0.07 | 5% | A0A087W |
| tr\|G3V498 | DLST | Dihydroli | 4159 | 17 | 1(0) | 1(0) | 0.91 | 35% | G3V498_u |
| sp\|P6122 | ABCE1 | ATP-bind | 68240 | 15 | 2(0) | 1(0) | 0.05 | 2% | P61221_u |
| tr\|B3KTV0 | | cDNA FL | 68166 | 15 | 1(0) | 1(0) | 0.05 | 1% | B3KTV0_u |
| tr\|A0A024 | hCG_203 | HCG203 | 81003 | 15 | 2(0) | 2(0) | 0.04 | 3% | A0A024R |
| tr\|L0R5B2 | GPR68 | Alternativ | 13728 | 14 | 5(0) | 1(0) | 0.25 | 8% | L0R5B2_u |
| tr\|A0A024 | DHCR7 | 7-dehydr | 55195 | 14 | 3(0) | 3(0) | 0.06 | 6% | A0A024R |

**Group shControl after exclusion of Group IgG and Group shNAT10**
**Accession**
tr|A0A384NYT8|A0A384NYT8_HUMAN
tr|D3DU92|D3DU92_HUMAN
tr|A0A248RGE3|A0A248RGE3_HUMAN
tr|A0A0A0MS51|A0A0A0MS51_HUMAN
tr|A8K134|A8K134_HUMAN
tr|A1A508|A1A508_HUMAN
tr|Q6MZU6|Q6MZU6_HUMAN
tr|A0A024RCN6|A0A024RCN6_HUMAN
sp|O15021|MAST4_HUMAN
tr|A0A2X0SFJ0|A0A2X0SFJ0_HUMAN
sp|P49454|CENPF_HUMAN
tr|A0A2R8Y516|A0A2R8Y516_HUMAN
tr|H0YAQ3|H0YAQ3_HUMAN
tr|A0A024R4S2|A0A024R4S2_HUMAN
tr|E3Q1J2|E3Q1J2_HUMAN
tr|A0A0S2Z4P6|A0A0S2Z4P6_HUMAN
sp|O94804|STK10_HUMAN
tr|A0A0S2Z3S6|A0A0S2Z3S6_HUMAN

Table S2. related to Figure 5

| gene_id | gene_syml | P_value(sh | TR_shNAT: | TR_shNAT: | TR_shNAT: | average_TR_shNAT10/shControl | |
|---|---|---|---|---|---|---|---|
| 10657 | KHDRBS1 | 1.15E-08 | 0.56564 | 0.564225 | 0.568332 | 0.566066 | down |
| 55735 | DNAJC11 | 1.17E-08 | 0.502422 | 0.504224 | 0.493637 | 0.500094 | down |
| 55226 | NAT10 | 2.07E-08 | 0.495863 | 0.488737 | 0.499416 | 0.494672 | down |
| 81 | ACTN4 | 2.33E-08 | 0.521863 | 0.515101 | 0.527179 | 0.521381 | down |
| 372 | ARCN1 | 3.32E-08 | 0.478717 | 0.492239 | 0.493624 | 0.488193 | down |
| 2152 | F3 | 4.01E-08 | 0.533375 | 0.53993 | 0.539767 | 0.537691 | down |
| 92017 | SNX29 | 5.71E-08 | 0.438847 | 0.442914 | 0.436297 | 0.439352 | down |
| 80854 | SETD7 | 6.29E-08 | 0.570763 | 0.574829 | 0.567917 | 0.57117 | down |
| 10015 | PDCD6IP | 6.66E-08 | 0.562049 | 0.550427 | 0.55236 | 0.554945 | down |
| 80139 | ZNF703 | 9.95E-08 | 0.525128 | 0.513205 | 0.512388 | 0.516907 | down |
| 9180 | OSMR | 1.13E-07 | 0.483833 | 0.493444 | 0.503257 | 0.493511 | down |
| 2734 | GLG1 | 1.2E-07 | 0.419247 | 0.439676 | 0.432179 | 0.430367 | down |
| 22974 | TPX2 | 1.24E-07 | 0.500131 | 0.488944 | 0.506534 | 0.498536 | down |
| 399979 | SNX19 | 1.44E-07 | 0.490161 | 0.508062 | 0.499116 | 0.499113 | down |
| 6575 | SLC20A2 | 1.57E-07 | 0.493238 | 0.496636 | 0.484508 | 0.491461 | down |
| 79939 | SLC35E1 | 1.64E-07 | 0.559624 | 0.546959 | 0.55392 | 0.553501 | down |
| 79709 | COLGALT1 | 1.68E-07 | 0.601716 | 0.606891 | 0.612352 | 0.606987 | down |
| 11073 | TOPBP1 | 1.7E-07 | 0.559631 | 0.553911 | 0.56786 | 0.560467 | down |
| 6873 | TAF2 | 2.2E-07 | 0.513222 | 0.513913 | 0.528676 | 0.518604 | down |
| 9688 | NUP93 | 2.24E-07 | 0.53931 | 0.558406 | 0.543845 | 0.547187 | down |
| 121260 | SLC15A4 | 2.31E-07 | 0.554932 | 0.550198 | 0.553104 | 0.552744 | down |
| 55573 | CDV3 | 2.42E-07 | 0.512134 | 0.502795 | 0.495761 | 0.503563 | down |
| 22820 | COPG1 | 2.43E-07 | 0.481195 | 0.494462 | 0.468738 | 0.481465 | down |
| 4841 | NONO | 2.64E-07 | 0.505484 | 0.508046 | 0.494411 | 0.502647 | down |
| 10743 | RAI1 | 2.65E-07 | 0.505256 | 0.487551 | 0.492105 | 0.494971 | down |
| 3609 | ILF3 | 2.66E-07 | 0.430403 | 0.459958 | 0.444716 | 0.445026 | down |
| 1398 | CRK | 2.66E-07 | 0.557132 | 0.559838 | 0.55023 | 0.555734 | down |
| 1001 | CDH3 | 2.77E-07 | 0.532492 | 0.55042 | 0.530206 | 0.537706 | down |
| 130074 | FAM168B | 2.81E-07 | 0.433904 | 0.435908 | 0.457447 | 0.44242 | down |
| 5426 | POLE | 2.87E-07 | 0.470168 | 0.452666 | 0.441696 | 0.454843 | down |
| 3098 | HK1 | 3E-07 | 0.438087 | 0.440908 | 0.431102 | 0.436699 | down |
| 351 | APP | 3.25E-07 | 0.477521 | 0.491671 | 0.491764 | 0.486985 | down |
| 9805 | SCRN1 | 3.58E-07 | 0.47035 | 0.4819 | 0.485878 | 0.479376 | down |
| 8553 | BHLHE40 | 3.71E-07 | 0.477179 | 0.458497 | 0.460938 | 0.465538 | down |
| 84669 | USP32 | 3.82E-07 | 0.468261 | 0.465431 | 0.455595 | 0.463095 | down |
| 9516 | LITAF | 3.82E-07 | 0.56876 | 0.54928 | 0.557678 | 0.558573 | down |
| 10992 | SF3B2 | 3.89E-07 | 0.601497 | 0.579283 | 0.59132 | 0.5907 | down |
| 9221 | NOLC1 | 3.94E-07 | 0.441987 | 0.41704 | 0.434895 | 0.431307 | down |
| 22864 | R3HDM2 | 4.04E-07 | 0.393523 | 0.413745 | 0.407972 | 0.40508 | down |
| 10487 | CAP1 | 4.06E-07 | 0.650773 | 0.648933 | 0.645247 | 0.648318 | down |
| 9918 | NCAPD2 | 4.16E-07 | 0.436664 | 0.439575 | 0.433503 | 0.436581 | down |
| 6709 | SPTAN1 | 4.25E-07 | 0.405031 | 0.3932 | 0.400619 | 0.399617 | down |
| 7290 | HIRA | 4.78E-07 | 0.527184 | 0.555192 | 0.532933 | 0.538437 | down |
| 9392 | TGFBRAP1 | 4.87E-07 | 0.478288 | 0.47462 | 0.503566 | 0.485491 | down |
| 9794 | MAML1 | 4.91E-07 | 0.49606 | 0.519046 | 0.510011 | 0.508372 | down |
| 476 | ATP1A1 | 4.95E-07 | 0.525961 | 0.528831 | 0.537088 | 0.530627 | down |
| 950 | SCARB2 | 5.32E-07 | 0.504032 | 0.49804 | 0.527325 | 0.509799 | down |
| 6662 | SOX9 | 5.34E-07 | 0.467851 | 0.457383 | 0.488511 | 0.471249 | down |
| 22905 | EPN2 | 5.54E-07 | 0.520369 | 0.516648 | 0.50895 | 0.515322 | down |
| 206358 | SLC36A1 | 5.56E-07 | 0.505034 | 0.495905 | 0.525838 | 0.508926 | down |
| 9791 | PTDSS1 | 5.56E-07 | 0.486211 | 0.49061 | 0.461733 | 0.479518 | down |
| 23180 | RFTN1 | 5.69E-07 | 0.580327 | 0.562106 | 0.554212 | 0.565548 | down |
| 1642 | DDB1 | 5.74E-07 | 0.592851 | 0.600568 | 0.610897 | 0.601439 | down |
| 1786 | DNMT1 | 5.95E-07 | 0.405748 | 0.42393 | 0.435008 | 0.421562 | down |
| 83931 | STK40 | 6.04E-07 | 0.61947 | 0.61876 | 0.597592 | 0.611941 | down |
| 4170 | MCL1 | 6.58E-07 | 0.491945 | 0.518295 | 0.487604 | 0.499281 | down |

| | | | | | | | |
|---|---|---|---|---|---|---|---|
| 9448 | MAP4K4 | 6.58E-07 | 0.404207 | 0.404151 | 0.42435 | 0.410903 | down |
| 84447 | SYVN1 | 6.75E-07 | 0.52291 | 0.531554 | 0.550145 | 0.53487 | down |
| 2683 | B4GALT1 | 6.76E-07 | 0.501813 | 0.509528 | 0.494824 | 0.502055 | down |
| 9846 | GAB2 | 6.78E-07 | 0.446237 | 0.47446 | 0.47093 | 0.463876 | down |
| 200081 | TXLNA | 6.88E-07 | 0.505817 | 0.518583 | 0.538103 | 0.520834 | down |
| 23313 | KIAA0930 | 7.01E-07 | 0.654601 | 0.674755 | 0.668285 | 0.66588 | down |
| 23279 | NUP160 | 7.16E-07 | 0.555963 | 0.571248 | 0.586505 | 0.571239 | down |
| 4134 | MAP4 | 7.19E-07 | 0.468613 | 0.437488 | 0.465983 | 0.457361 | down |
| 29102 | DROSHA | 7.19E-07 | 0.524924 | 0.528375 | 0.534674 | 0.529325 | down |
| 55690 | PACS1 | 7.47E-07 | 0.550168 | 0.536333 | 0.552457 | 0.546319 | down |
| 8663 | EIF3C | 7.48E-07 | 0.428541 | 0.395685 | 0.419344 | 0.414523 | down |
| 54471 | MIEF1 | 7.94E-07 | 0.594056 | 0.621135 | 0.595263 | 0.603485 | down |
| 2196 | FAT2 | 8.08E-07 | 0.434616 | 0.430471 | 0.423123 | 0.429403 | down |
| 9870 | AREL1 | 8.11E-07 | 0.501751 | 0.504953 | 0.52136 | 0.509355 | down |
| 3840 | KPNA4 | 8.11E-07 | 0.548662 | 0.575026 | 0.567838 | 0.563842 | down |
| 6774 | STAT3 | 8.16E-07 | 0.515633 | 0.54452 | 0.540874 | 0.533676 | down |
| 55884 | WSB2 | 8.16E-07 | 0.652452 | 0.655039 | 0.662873 | 0.656788 | down |
| 3190 | HNRNPK | 8.66E-07 | 0.627051 | 0.627339 | 0.649198 | 0.634529 | down |
| 833 | CARS | 8.91E-07 | 0.568209 | 0.568784 | 0.547524 | 0.561506 | down |
| 831 | CAST | 8.95E-07 | 0.65799 | 0.630913 | 0.64496 | 0.644621 | down |
| 3326 | HSP90AB1 | 9.09E-07 | 0.494648 | 0.511485 | 0.498503 | 0.501545 | down |
| 7204 | TRIO | 9.14E-07 | 0.389683 | 0.359236 | 0.397552 | 0.382157 | down |
| 8531 | YBX3 | 9.66E-07 | 0.525702 | 0.501427 | 0.519263 | 0.515464 | down |
| 5832 | ALDH18A1 | 9.73E-07 | 0.60633 | 0.625733 | 0.602089 | 0.611384 | down |
| 3916 | LAMP1 | 9.77E-07 | 0.572668 | 0.56858 | 0.553358 | 0.564869 | down |
| 79188 | TMEM43 | 9.8E-07 | 0.546937 | 0.556941 | 0.563342 | 0.55574 | down |
| 9531 | BAG3 | 1.02E-06 | 0.547039 | 0.523045 | 0.513366 | 0.527817 | down |
| 51291 | GMIP | 1.07E-06 | 0.580598 | 0.552425 | 0.574787 | 0.56927 | down |
| 10915 | TCERG1 | 1.08E-06 | 0.557784 | 0.569654 | 0.590109 | 0.572516 | down |
| 222484 | LNX2 | 1.14E-06 | 0.529092 | 0.550935 | 0.527364 | 0.535797 | down |
| 9131 | AIFM1 | 1.14E-06 | 0.583522 | 0.60012 | 0.593234 | 0.592292 | down |
| 84959 | UBASH3B | 1.15E-06 | 0.515802 | 0.539226 | 0.523667 | 0.526232 | down |
| 10565 | ARFGEF1 | 1.15E-06 | 0.438935 | 0.418544 | 0.429569 | 0.429016 | down |
| 113251 | LARP4 | 1.17E-06 | 0.616293 | 0.60664 | 0.614388 | 0.61244 | down |
| 22836 | RHOBTB3 | 1.2E-06 | 0.455021 | 0.480603 | 0.48917 | 0.474931 | down |
| 55102 | ATG2B | 1.25E-06 | 0.379337 | 0.423904 | 0.400505 | 0.401249 | down |
| 6319 | SCD | 1.26E-06 | 0.430674 | 0.44593 | 0.445498 | 0.440701 | down |
| 23731 | TMEM245 | 1.26E-06 | 0.565634 | 0.584897 | 0.566037 | 0.572189 | down |
| 4233 | MET | 1.28E-06 | 0.472321 | 0.503637 | 0.500458 | 0.492139 | down |
| 5514 | PPP1R10 | 1.29E-06 | 0.482495 | 0.446143 | 0.476833 | 0.46849 | down |
| 55677 | IWS1 | 1.3E-06 | 0.545753 | 0.547337 | 0.544845 | 0.545978 | down |
| 11261 | CHP1 | 1.32E-06 | 0.492035 | 0.494652 | 0.507153 | 0.497947 | down |
| 2177 | FANCD2 | 1.33E-06 | 0.405582 | 0.439024 | 0.445292 | 0.429966 | down |
| 160428 | ALDH1L2 | 1.37E-06 | 0.371935 | 0.39448 | 0.395596 | 0.387337 | down |
| 55765 | INAVA | 1.37E-06 | 0.595054 | 0.611424 | 0.598307 | 0.601595 | down |
| 9265 | CYTH3 | 1.39E-06 | 0.529886 | 0.547256 | 0.562445 | 0.546529 | down |
| 10484 | SEC23A | 1.41E-06 | 0.556521 | 0.577289 | 0.576374 | 0.570061 | down |
| 27250 | PDCD4 | 1.43E-06 | 0.593561 | 0.599924 | 0.588629 | 0.594038 | down |
| 2821 | GPI | 1.45E-06 | 0.583267 | 0.593589 | 0.609549 | 0.595468 | down |
| 23019 | CNOT1 | 1.46E-06 | 0.328435 | 0.346547 | 0.343322 | 0.339435 | down |
| 4301 | AFDN | 1.48E-06 | 0.398544 | 0.411939 | 0.428075 | 0.412853 | down |
| 196463 | PLBD2 | 1.51E-06 | 0.461513 | 0.447599 | 0.453473 | 0.454195 | down |
| 2617 | GARS | 1.51E-06 | 0.576298 | 0.605961 | 0.571405 | 0.584555 | down |
| 25777 | SUN2 | 1.52E-06 | 0.483148 | 0.505063 | 0.509957 | 0.499389 | down |
| 56850 | GRIPAP1 | 1.56E-06 | 0.446094 | 0.458842 | 0.47191 | 0.458949 | down |
| 7126 | TNFAIP1 | 1.56E-06 | 0.548552 | 0.516601 | 0.527745 | 0.530966 | down |
| 164 | AP1G1 | 1.57E-06 | 0.433587 | 0.443527 | 0.456334 | 0.444483 | down |
| 7045 | TGFBI | 1.73E-06 | 0.550147 | 0.555544 | 0.518581 | 0.541424 | down |

| | | | | | | | |
|---|---|---|---|---|---|---|---|
| 6567 | SLC16A2 | 1.75E-06 | 0.538703 | 0.519583 | 0.549516 | 0.535934 | down |
| 1839 | HBEGF | 1.77E-06 | 0.60451 | 0.598079 | 0.578258 | 0.593616 | down |
| 2782 | GNB1 | 1.86E-06 | 0.571433 | 0.567583 | 0.59627 | 0.578429 | down |
| 1308 | COL17A1 | 1.93E-06 | 0.446476 | 0.451192 | 0.415792 | 0.43782 | down |
| 4522 | MTHFD1 | 1.93E-06 | 0.565539 | 0.603634 | 0.575609 | 0.581594 | down |
| 2048 | EPHB2 | 1.98E-06 | 0.527634 | 0.515887 | 0.513273 | 0.518931 | down |
| 1729 | DIAPH1 | 2E-06 | 0.447777 | 0.443464 | 0.467922 | 0.453054 | down |
| 3480 | IGF1R | 2.01E-06 | 0.426568 | 0.413218 | 0.386448 | 0.408745 | down |
| 25921 | ZDHHC5 | 2.02E-06 | 0.538296 | 0.531047 | 0.542216 | 0.537187 | down |
| 2962 | GTF2F1 | 2.03E-06 | 0.55533 | 0.538089 | 0.530533 | 0.541317 | down |
| 54765 | TRIM44 | 2.03E-06 | 0.435368 | 0.453784 | 0.481542 | 0.456898 | down |
| 8658 | TNKS | 2.04E-06 | 0.469496 | 0.460954 | 0.488784 | 0.473078 | down |
| 4172 | MCM3 | 2.06E-06 | 0.555274 | 0.516999 | 0.546022 | 0.539432 | down |
| 16 | AARS | 2.06E-06 | 0.517956 | 0.530472 | 0.527805 | 0.525411 | down |
| 25792 | CIZ1 | 2.1E-06 | 0.554057 | 0.518688 | 0.551769 | 0.541505 | down |
| 64866 | CDCP1 | 2.11E-06 | 0.400468 | 0.408495 | 0.43807 | 0.415678 | down |
| 87 | ACTN1 | 2.14E-06 | 0.51678 | 0.514132 | 0.495604 | 0.508839 | down |
| 23176 | 44447 | 2.15E-06 | 0.533595 | 0.521239 | 0.511847 | 0.522227 | down |
| 51474 | LIMA1 | 2.16E-06 | 0.641323 | 0.655815 | 0.67362 | 0.656919 | down |
| 3728 | JUP | 2.17E-06 | 0.577111 | 0.570218 | 0.54817 | 0.565166 | down |
| 92597 | MOB1B | 2.2E-06 | 0.638847 | 0.631212 | 0.634726 | 0.634928 | down |
| 5306 | PITPNA | 2.24E-06 | 0.610038 | 0.62414 | 0.604258 | 0.612812 | down |
| 10057 | ABCC5 | 2.26E-06 | 0.546599 | 0.550279 | 0.531361 | 0.542747 | down |
| 57590 | WDFY1 | 2.26E-06 | 0.501086 | 0.49649 | 0.515526 | 0.504368 | down |
| 23385 | NCSTN | 2.28E-06 | 0.51938 | 0.544812 | 0.544535 | 0.536242 | down |
| 1453 | CSNK1D | 2.29E-06 | 0.565589 | 0.567555 | 0.529641 | 0.554262 | down |
| 7128 | TNFAIP3 | 2.3E-06 | 0.504889 | 0.475081 | 0.510089 | 0.496687 | down |
| 57003 | CCDC47 | 2.32E-06 | 0.572547 | 0.585284 | 0.5888 | 0.58221 | down |
| 23310 | NCAPD3 | 2.32E-06 | 0.460294 | 0.482371 | 0.499673 | 0.480779 | down |
| 8678 | BECN1 | 2.36E-06 | 0.631179 | 0.645202 | 0.610005 | 0.628795 | down |
| 6772 | STAT1 | 2.37E-06 | 0.480565 | 0.490333 | 0.498323 | 0.48974 | down |
| 7994 | KAT6A | 2.48E-06 | 0.315976 | 0.357518 | 0.303023 | 0.325506 | down |
| 663 | BNIP2 | 2.49E-06 | 0.583022 | 0.582 | 0.605344 | 0.590122 | down |
| 22984 | PDCD11 | 2.51E-06 | 0.428691 | 0.398032 | 0.416862 | 0.414528 | down |
| 11198 | SUPT16H | 2.51E-06 | 0.49791 | 0.515953 | 0.526055 | 0.513306 | down |
| 6789 | STK4 | 2.52E-06 | 0.460416 | 0.50053 | 0.455211 | 0.472052 | down |
| 51631 | LUC7L2 | 2.57E-06 | 0.347171 | 0.288634 | 0.316393 | 0.317399 | down |
| 11100 | HNRNPUL | 2.58E-06 | 0.51603 | 0.495321 | 0.501179 | 0.504177 | down |
| 50488 | MINK1 | 2.59E-06 | 0.503179 | 0.459318 | 0.502034 | 0.488177 | down |
| 205 | AK4 | 2.62E-06 | 0.648116 | 0.648593 | 0.657833 | 0.651514 | down |
| 116496 | FAM129A | 2.63E-06 | 0.37053 | 0.37085 | 0.408835 | 0.383405 | down |
| 57498 | KIDINS220 | 2.65E-06 | 0.380692 | 0.419782 | 0.380996 | 0.393823 | down |
| 11128 | POLR3A | 2.66E-06 | 0.56456 | 0.545843 | 0.580873 | 0.563759 | down |
| 3918 | LAMC2 | 2.72E-06 | 0.47363 | 0.514106 | 0.505697 | 0.497811 | down |
| 4928 | NUP98 | 2.73E-06 | 0.427806 | 0.428106 | 0.430232 | 0.428715 | down |
| 10097 | ACTR2 | 2.77E-06 | 0.615722 | 0.627255 | 0.623006 | 0.621994 | down |
| 10525 | HYOU1 | 2.78E-06 | 0.495523 | 0.463295 | 0.480355 | 0.479725 | down |
| 58490 | RPRD1B | 2.81E-06 | 0.495404 | 0.525211 | 0.511015 | 0.510544 | down |
| 27327 | TNRC6A | 2.86E-06 | 0.362477 | 0.403444 | 0.394478 | 0.3868 | down |
| 23247 | KIAA0556 | 2.86E-06 | 0.458171 | 0.414291 | 0.448277 | 0.440246 | down |
| 8880 | FUBP1 | 2.87E-06 | 0.552761 | 0.555372 | 0.579064 | 0.562399 | down |
| 27301 | APEX2 | 2.87E-06 | 0.66053 | 0.641794 | 0.662639 | 0.654987 | down |
| 11196 | SEC23IP | 2.9E-06 | 0.565258 | 0.574219 | 0.569138 | 0.569538 | down |
| 1500 | CTNND1 | 2.91E-06 | 0.401785 | 0.376387 | 0.414348 | 0.397507 | down |
| 23518 | R3HDM1 | 2.94E-06 | 0.393998 | 0.44267 | 0.397309 | 0.411326 | down |
| 1490 | CTGF | 2.95E-06 | 0.562769 | 0.568438 | 0.54894 | 0.560049 | down |
| 3875 | KRT18 | 2.95E-06 | 0.627943 | 0.621712 | 0.597708 | 0.615788 | down |
| 261734 | NPHP4 | 3.01E-06 | 0.602902 | 0.641786 | 0.625864 | 0.623517 | down |

| | | | | | | | |
|---|---|---|---|---|---|---|---|
| 157922 | CAMSAP1 | 3.1E-06 | 0.468487 | 0.475138 | 0.42676 | 0.456795 | down |
| 3551 | IKBKB | 3.14E-06 | 0.564666 | 0.521355 | 0.53457 | 0.540197 | down |
| 5829 | PXN | 3.21E-06 | 0.563951 | 0.544178 | 0.56326 | 0.557129 | down |
| 8202 | NCOA3 | 3.21E-06 | 0.419382 | 0.406537 | 0.453147 | 0.426356 | down |
| 5066 | PAM | 3.3E-06 | 0.472247 | 0.479871 | 0.51994 | 0.490686 | down |
| 7259 | TSPYL1 | 3.34E-06 | 0.371586 | 0.391285 | 0.428358 | 0.397076 | down |
| 23221 | RHOBTB2 | 3.35E-06 | 0.524722 | 0.492996 | 0.499226 | 0.505648 | down |
| 1737 | DLAT | 3.37E-06 | 0.547188 | 0.555748 | 0.57491 | 0.559282 | down |
| 5894 | RAF1 | 3.37E-06 | 0.575534 | 0.596625 | 0.563933 | 0.578697 | down |
| 7248 | TSC1 | 3.42E-06 | 0.465359 | 0.468434 | 0.468652 | 0.467481 | down |
| 57492 | ARID1B | 3.45E-06 | 0.368501 | 0.369637 | 0.345616 | 0.361252 | down |
| 5786 | PTPRA | 3.48E-06 | 0.542073 | 0.573371 | 0.584639 | 0.566694 | down |
| 22883 | CLSTN1 | 3.48E-06 | 0.436463 | 0.464894 | 0.424822 | 0.44206 | down |
| 25937 | WWTR1 | 3.48E-06 | 0.560052 | 0.563135 | 0.597247 | 0.573478 | down |
| 9779 | TBC1D5 | 3.49E-06 | 0.446257 | 0.433606 | 0.453054 | 0.444306 | down |
| 4152 | MBD1 | 3.5E-06 | 0.615753 | 0.628513 | 0.618455 | 0.620907 | down |
| 55252 | ASXL2 | 3.57E-06 | 0.4043 | 0.388132 | 0.398677 | 0.397036 | down |
| 9948 | WDR1 | 3.59E-06 | 0.598487 | 0.59037 | 0.568745 | 0.585867 | down |
| 51621 | KLF13 | 3.66E-06 | 0.533972 | 0.534726 | 0.4892 | 0.519299 | down |
| 23633 | KPNA6 | 3.68E-06 | 0.56279 | 0.586144 | 0.572145 | 0.573693 | down |
| 31 | ACACA | 3.7E-06 | 0.379452 | 0.378989 | 0.381705 | 0.380049 | down |
| 7414 | VCL | 3.71E-06 | 0.422614 | 0.431086 | 0.447051 | 0.433584 | down |
| 9733 | SART3 | 3.71E-06 | 0.402199 | 0.426613 | 0.437064 | 0.421959 | down |
| 5754 | PTK7 | 3.76E-06 | 0.640909 | 0.636598 | 0.636498 | 0.638002 | down |
| 55159 | RFWD3 | 3.78E-06 | 0.585504 | 0.609292 | 0.582538 | 0.592445 | down |
| 4047 | LSS | 3.81E-06 | 0.548528 | 0.538615 | 0.520111 | 0.535751 | down |
| 9962 | SLC23A2 | 3.84E-06 | 0.431672 | 0.418264 | 0.460499 | 0.436812 | down |
| 9746 | CLSTN3 | 3.84E-06 | 0.576443 | 0.566063 | 0.535548 | 0.559351 | down |
| 9690 | UBE3C | 3.93E-06 | 0.434377 | 0.436397 | 0.451523 | 0.440766 | down |
| 57636 | ARHGAP23 | 4E-06 | 0.597835 | 0.57159 | 0.597301 | 0.588908 | down |
| 9777 | TM9SF4 | 4.03E-06 | 0.443734 | 0.470139 | 0.424968 | 0.44628 | down |
| 64065 | PERP | 4.03E-06 | 0.6223 | 0.636627 | 0.625014 | 0.62798 | down |
| 7159 | TP53BP2 | 4.03E-06 | 0.47627 | 0.43373 | 0.469582 | 0.459861 | down |
| 6749 | SSRP1 | 4.06E-06 | 0.482121 | 0.501875 | 0.488596 | 0.490864 | down |
| 51593 | SRRT | 4.07E-06 | 0.458655 | 0.407343 | 0.441343 | 0.43578 | down |
| 23394 | ADNP | 4.1E-06 | 0.59047 | 0.584612 | 0.574752 | 0.583278 | down |
| 2113 | ETS1 | 4.12E-06 | 0.424554 | 0.454758 | 0.466721 | 0.448678 | down |
| 51366 | UBR5 | 4.13E-06 | 0.381467 | 0.435568 | 0.411649 | 0.409561 | down |
| 1975 | EIF4B | 4.14E-06 | 0.604049 | 0.626854 | 0.581819 | 0.604241 | down |
| 5328 | PLAU | 4.15E-06 | 0.605954 | 0.640736 | 0.626273 | 0.624321 | down |
| 9967 | THRAP3 | 4.2E-06 | 0.450558 | 0.449322 | 0.444971 | 0.448284 | down |
| 960 | CD44 | 4.24E-06 | 0.490078 | 0.468317 | 0.509154 | 0.489183 | down |
| 51604 | PIGT | 4.3E-06 | 0.633567 | 0.660655 | 0.647905 | 0.647376 | down |
| 2137 | EXTL3 | 4.31E-06 | 0.601583 | 0.605628 | 0.590039 | 0.599083 | down |
| 595 | CCND1 | 4.32E-06 | 0.423411 | 0.428261 | 0.424271 | 0.425314 | down |
| 8943 | AP3D1 | 4.33E-06 | 0.450868 | 0.430679 | 0.401555 | 0.427701 | down |
| 144455 | E2F7 | 4.39E-06 | 0.39734 | 0.400278 | 0.415296 | 0.404305 | down |
| 57062 | DDX24 | 4.41E-06 | 0.518589 | 0.559035 | 0.539851 | 0.539158 | down |
| 2317 | FLNB | 4.43E-06 | 0.537569 | 0.517326 | 0.498498 | 0.517798 | down |
| 4952 | OCRL | 4.43E-06 | 0.39389 | 0.418998 | 0.43311 | 0.415333 | down |
| 23060 | ZNF609 | 4.44E-06 | 0.424629 | 0.430059 | 0.423558 | 0.426082 | down |
| 5660 | PSAP | 4.53E-06 | 0.6351 | 0.664683 | 0.625627 | 0.641803 | down |
| 6891 | TAP2 | 4.53E-06 | 0.458098 | 0.435777 | 0.470794 | 0.454889 | down |
| 154810 | AMOTL1 | 4.55E-06 | 0.364639 | 0.36998 | 0.393304 | 0.375974 | down |
| 10130 | PDIA6 | 4.55E-06 | 0.62387 | 0.615147 | 0.626861 | 0.621959 | down |
| 10838 | ZNF275 | 4.59E-06 | 0.402595 | 0.459727 | 0.414305 | 0.425543 | down |
| 6051 | RNPEP | 4.6E-06 | 0.57977 | 0.567158 | 0.603555 | 0.583495 | down |
| 23076 | RRP1B | 4.61E-06 | 0.445409 | 0.440072 | 0.436993 | 0.440825 | down |

| | | | | | | | |
|---|---|---|---|---|---|---|---|
| 10523 | CHERP | 4.62E-06 | 0.487153 | 0.446479 | 0.450688 | 0.46144 | down |
| 387521 | TMEM189 | 4.63E-06 | 0.615252 | 0.64432 | 0.630429 | 0.63 | down |
| 339448 | C1orf174 | 4.63E-06 | 0.609483 | 0.573851 | 0.595977 | 0.593104 | down |
| 30844 | EHD4 | 4.65E-06 | 0.532915 | 0.547981 | 0.53601 | 0.538968 | down |
| 5170 | PDPK1 | 4.7E-06 | 0.414007 | 0.451486 | 0.451173 | 0.438888 | down |
| 6046 | BRD2 | 4.71E-06 | 0.566844 | 0.54557 | 0.559158 | 0.557191 | down |
| 54828 | BCAS3 | 4.71E-06 | 0.577627 | 0.620779 | 0.583343 | 0.593916 | down |
| 7468 | NSD2 | 4.74E-06 | 0.482229 | 0.512539 | 0.487286 | 0.494018 | down |
| 10645 | CAMKK2 | 4.75E-06 | 0.587435 | 0.618471 | 0.613916 | 0.606608 | down |
| 10618 | TGOLN2 | 4.76E-06 | 0.412643 | 0.430592 | 0.43085 | 0.424695 | down |
| 5708 | PSMD2 | 4.77E-06 | 0.587059 | 0.603771 | 0.619847 | 0.603559 | down |
| 51729 | WBP11 | 4.78E-06 | 0.515497 | 0.503906 | 0.540389 | 0.51993 | down |
| 9100 | USP10 | 4.79E-06 | 0.513762 | 0.547642 | 0.535254 | 0.532219 | down |
| 56919 | DHX33 | 4.84E-06 | 0.410251 | 0.405599 | 0.409989 | 0.408613 | down |
| 4940 | OAS3 | 4.87E-06 | 0.433366 | 0.441546 | 0.476117 | 0.450343 | down |
| 23082 | PPRC1 | 4.88E-06 | 0.513974 | 0.546523 | 0.524015 | 0.528171 | down |
| 23636 | NUP62 | 4.88E-06 | 0.582314 | 0.605527 | 0.604655 | 0.597499 | down |
| 51061 | TXNDC11 | 4.88E-06 | 0.585425 | 0.59165 | 0.593489 | 0.590188 | down |
| 8476 | CDC42BPA | 4.89E-06 | 0.37735 | 0.428732 | 0.410719 | 0.4056 | down |
| 54994 | GID8 | 4.99E-06 | 0.638838 | 0.632007 | 0.64614 | 0.638995 | down |
| 103 | ADAR | 4.99E-06 | 0.397374 | 0.422519 | 0.417736 | 0.412543 | down |
| 55785 | FGD6 | 5E-06 | 0.640655 | 0.659869 | 0.653008 | 0.651177 | down |
| 7798 | LUZP1 | 5E-06 | 0.441116 | 0.447194 | 0.462378 | 0.450229 | down |
| 3675 | ITGA3 | 5.02E-06 | 0.640953 | 0.614356 | 0.62367 | 0.626326 | down |
| 51752 | ERAP1 | 5.03E-06 | 0.446105 | 0.443724 | 0.417966 | 0.435932 | down |
| 26060 | APPL1 | 5.04E-06 | 0.56976 | 0.554729 | 0.557483 | 0.560657 | down |
| 1717 | DHCR7 | 5.05E-06 | 0.670957 | 0.651685 | 0.644715 | 0.655785 | down |
| 23608 | MKRN1 | 5.08E-06 | 0.655667 | 0.631624 | 0.632407 | 0.639899 | down |
| 23435 | TARDBP | 5.09E-06 | 0.636395 | 0.630315 | 0.642233 | 0.636315 | down |
| 6950 | TCP1 | 5.1E-06 | 0.665851 | 0.651121 | 0.628055 | 0.648343 | down |
| 84515 | MCM8 | 5.14E-06 | 0.523927 | 0.500557 | 0.553559 | 0.526014 | down |
| 118987 | PDZD8 | 5.17E-06 | 0.42221 | 0.40698 | 0.398403 | 0.409198 | down |
| 7296 | TXNRD1 | 5.17E-06 | 0.561112 | 0.60436 | 0.587194 | 0.584222 | down |
| 54733 | SLC35F2 | 5.18E-06 | 0.494533 | 0.537063 | 0.540092 | 0.523896 | down |
| 57680 | CHD8 | 5.19E-06 | 0.343771 | 0.349666 | 0.395613 | 0.363016 | down |
| 149986 | LSM14B | 5.23E-06 | 0.446974 | 0.50717 | 0.467996 | 0.474047 | down |
| 9665 | MARF1 | 5.24E-06 | 0.412642 | 0.398399 | 0.421171 | 0.410737 | down |
| 11187 | PKP3 | 5.24E-06 | 0.563733 | 0.573697 | 0.542547 | 0.559992 | down |
| 6678 | SPARC | 5.25E-06 | 0.610754 | 0.637713 | 0.645841 | 0.631436 | down |
| 9219 | MTA2 | 5.36E-06 | 0.562467 | 0.515668 | 0.547141 | 0.541759 | down |
| 7267 | TTC3 | 5.36E-06 | 0.363556 | 0.382493 | 0.408248 | 0.384766 | down |
| 4088 | SMAD3 | 5.43E-06 | 0.523923 | 0.539076 | 0.558033 | 0.540344 | down |
| 23001 | WDFY3 | 5.46E-06 | 0.304306 | 0.336605 | 0.369337 | 0.33675 | down |
| 9788 | MTSS1 | 5.46E-06 | 0.527239 | 0.480062 | 0.478852 | 0.495384 | down |
| 8451 | CUL4A | 5.49E-06 | 0.530096 | 0.544018 | 0.559196 | 0.544437 | down |
| 7453 | WARS | 5.51E-06 | 0.591563 | 0.559791 | 0.547932 | 0.566428 | down |
| 382 | ARF6 | 5.54E-06 | 0.610876 | 0.627523 | 0.644679 | 0.627692 | down |
| 3996 | LLGL1 | 5.56E-06 | 0.579873 | 0.571416 | 0.54005 | 0.56378 | down |
| 3694 | ITGB6 | 5.58E-06 | 0.515373 | 0.538135 | 0.494908 | 0.516139 | down |
| 150696 | PROM2 | 5.59E-06 | 0.635511 | 0.602484 | 0.615514 | 0.617836 | down |
| 339287 | MSL1 | 5.62E-06 | 0.561925 | 0.536058 | 0.577589 | 0.558524 | down |
| 334 | APLP2 | 5.66E-06 | 0.56031 | 0.562831 | 0.550016 | 0.557719 | down |
| 10552 | ARPC1A | 5.67E-06 | 0.622783 | 0.605115 | 0.593352 | 0.607084 | down |
| 28964 | GIT1 | 5.75E-06 | 0.652682 | 0.618985 | 0.608977 | 0.626881 | down |
| 23380 | SRGAP2 | 5.76E-06 | 0.412933 | 0.418737 | 0.429837 | 0.420502 | down |
| 219 | ALDH1B1 | 5.84E-06 | 0.479936 | 0.513803 | 0.508975 | 0.500905 | down |
| 118 | ADD1 | 5.84E-06 | 0.519957 | 0.559019 | 0.52329 | 0.534088 | down |
| 9188 | DDX21 | 5.85E-06 | 0.54101 | 0.532121 | 0.561309 | 0.544813 | down |

| | | | | | | | |
|---|---|---|---|---|---|---|---|
| 4140 | MARK3 | 5.89E-06 | 0.631809 | 0.592825 | 0.605647 | 0.610094 | down |
| 6872 | TAF1 | 5.9E-06 | 0.419974 | 0.395759 | 0.416503 | 0.410745 | down |
| 6730 | SRP68 | 5.94E-06 | 0.553867 | 0.581671 | 0.604718 | 0.580085 | down |
| 2120 | ETV6 | 6.1E-06 | 0.520256 | 0.522689 | 0.548236 | 0.530394 | down |
| 55661 | DDX27 | 6.15E-06 | 0.545641 | 0.514383 | 0.49523 | 0.518418 | down |
| 23637 | RABGAP1 | 6.17E-06 | 0.551349 | 0.525214 | 0.519531 | 0.532031 | down |
| 2119 | ETV5 | 6.21E-06 | 0.600764 | 0.582096 | 0.59394 | 0.592267 | down |
| 23196 | FAM120A | 6.24E-06 | 0.451251 | 0.466792 | 0.490452 | 0.469498 | down |
| 47 | ACLY | 6.28E-06 | 0.408379 | 0.431877 | 0.416102 | 0.418786 | down |
| 10509 | SEMA4B | 6.34E-06 | 0.630548 | 0.626617 | 0.605187 | 0.620784 | down |
| 4323 | MMP14 | 6.35E-06 | 0.541916 | 0.490366 | 0.516239 | 0.516174 | down |
| 27339 | PRPF19 | 6.39E-06 | 0.635975 | 0.637118 | 0.618914 | 0.630669 | down |
| 221496 | LEMD2 | 6.41E-06 | 0.579764 | 0.60575 | 0.562164 | 0.58256 | down |
| 79139 | DERL1 | 6.42E-06 | 0.571813 | 0.532425 | 0.55703 | 0.553756 | down |
| 1958 | EGR1 | 6.43E-06 | 0.626435 | 0.594602 | 0.605831 | 0.608956 | down |
| 7430 | EZR | 6.44E-06 | 0.509537 | 0.531437 | 0.520972 | 0.520649 | down |
| 22992 | KDM2A | 6.46E-06 | 0.44911 | 0.488156 | 0.507685 | 0.48165 | down |
| 4609 | MYC | 6.46E-06 | 0.64967 | 0.610687 | 0.620414 | 0.626924 | down |
| 5859 | QARS | 6.49E-06 | 0.549414 | 0.552298 | 0.556471 | 0.552728 | down |
| 64327 | LMBR1 | 6.49E-06 | 0.662264 | 0.632227 | 0.647976 | 0.647489 | down |
| 23511 | NUP188 | 6.51E-06 | 0.463116 | 0.475628 | 0.442242 | 0.460328 | down |
| 54888 | NSUN2 | 6.52E-06 | 0.48244 | 0.517714 | 0.535708 | 0.511954 | down |
| 84747 | UNC119B | 6.52E-06 | 0.523124 | 0.553068 | 0.516598 | 0.53093 | down |
| 11214 | AKAP13 | 6.53E-06 | 0.347645 | 0.39979 | 0.366489 | 0.371308 | down |
| 51780 | KDM3B | 6.53E-06 | 0.396042 | 0.439937 | 0.391683 | 0.409221 | down |
| 26574 | AATF | 6.59E-06 | 0.604301 | 0.6009 | 0.567785 | 0.590995 | down |
| 23053 | ZSWIM8 | 6.6E-06 | 0.528134 | 0.566823 | 0.521335 | 0.538764 | down |
| 8570 | KHSRP | 6.66E-06 | 0.516531 | 0.496289 | 0.460861 | 0.491227 | down |
| 26020 | LRP10 | 6.66E-06 | 0.620471 | 0.592128 | 0.598834 | 0.603811 | down |
| 57410 | SCYL1 | 6.7E-06 | 0.616877 | 0.609816 | 0.596769 | 0.607821 | down |
| 2976 | GTF3C2 | 6.73E-06 | 0.609771 | 0.580782 | 0.560624 | 0.583726 | down |
| 79733 | E2F8 | 6.74E-06 | 0.411757 | 0.386613 | 0.372795 | 0.390388 | down |
| 26509 | MYOF | 6.74E-06 | 0.369567 | 0.394138 | 0.383288 | 0.382331 | down |
| 29882 | ANAPC2 | 6.76E-06 | 0.601469 | 0.616542 | 0.575098 | 0.597703 | down |
| 3069 | HDLBP | 6.76E-06 | 0.407609 | 0.430062 | 0.387271 | 0.408314 | down |
| 404093 | CUEDC1 | 6.8E-06 | 0.606366 | 0.599976 | 0.639081 | 0.615141 | down |
| 54206 | ERRFI1 | 6.9E-06 | 0.54524 | 0.542943 | 0.588393 | 0.558859 | down |
| 824 | CAPN2 | 6.92E-06 | 0.540469 | 0.540654 | 0.512323 | 0.531149 | down |
| 23020 | SNRNP200 | 6.92E-06 | 0.44017 | 0.445068 | 0.424069 | 0.436436 | down |
| 55157 | DARS2 | 6.93E-06 | 0.580448 | 0.573139 | 0.596323 | 0.583303 | down |
| 5257 | PHKB | 6.94E-06 | 0.513288 | 0.56503 | 0.536304 | 0.538207 | down |
| 85456 | TNKS1BP1 | 6.95E-06 | 0.422301 | 0.397981 | 0.424407 | 0.414897 | down |
| 1762 | DMWD | 6.95E-06 | 0.604746 | 0.623731 | 0.63773 | 0.622069 | down |
| 1889 | ECE1 | 6.98E-06 | 0.512413 | 0.49879 | 0.496493 | 0.502565 | down |
| 145567 | TTC7B | 7.02E-06 | 0.568699 | 0.513745 | 0.541306 | 0.54125 | down |
| 51573 | GDE1 | 7.03E-06 | 0.602842 | 0.575748 | 0.605952 | 0.594847 | down |
| 79134 | TMEM185I | 7.06E-06 | 0.563309 | 0.571577 | 0.52447 | 0.553119 | down |
| 3185 | HNRNPF | 7.12E-06 | 0.495138 | 0.546182 | 0.533752 | 0.525024 | down |
| 5362 | PLXNA2 | 7.12E-06 | 0.461177 | 0.453591 | 0.476825 | 0.463864 | down |
| 10664 | CTCF | 7.12E-06 | 0.499503 | 0.450402 | 0.439343 | 0.463083 | down |
| 8621 | CDK13 | 7.18E-06 | 0.497585 | 0.517115 | 0.47645 | 0.49705 | down |
| 3852 | KRT5 | 7.19E-06 | 0.512904 | 0.48923 | 0.493714 | 0.498616 | down |
| 170954 | PPP1R18 | 7.19E-06 | 0.619026 | 0.591344 | 0.569894 | 0.593421 | down |
| 5326 | PLAGL2 | 7.2E-06 | 0.479134 | 0.426657 | 0.451434 | 0.452408 | down |
| 3927 | LASP1 | 7.25E-06 | 0.532279 | 0.513035 | 0.531019 | 0.525444 | down |
| 57479 | PRR12 | 7.26E-06 | 0.513168 | 0.514368 | 0.485925 | 0.504487 | down |
| 8565 | YARS | 7.31E-06 | 0.592335 | 0.603335 | 0.58166 | 0.592443 | down |
| 56270 | WDR45B | 7.31E-06 | 0.619321 | 0.64556 | 0.619273 | 0.628051 | down |

| | | | | | | | |
|---|---|---|---|---|---|---|---|
| 80018 | NAA25 | 7.33E-06 | 0.482683 | 0.536754 | 0.516733 | 0.512057 | down |
| 23505 | TMEM131 | 7.39E-06 | 0.335222 | 0.357194 | 0.338506 | 0.343641 | down |
| 867 | CBL | 7.39E-06 | 0.370476 | 0.389765 | 0.383346 | 0.381196 | down |
| 51164 | DCTN4 | 7.41E-06 | 0.589359 | 0.564685 | 0.591238 | 0.581761 | down |
| 5430 | POLR2A | 7.42E-06 | 0.415518 | 0.439222 | 0.392725 | 0.415821 | down |
| 23534 | TNPO3 | 7.43E-06 | 0.439743 | 0.444287 | 0.464112 | 0.449381 | down |
| 26031 | OSBPL3 | 7.46E-06 | 0.458533 | 0.495815 | 0.480526 | 0.478291 | down |
| 6837 | MED22 | 7.5E-06 | 0.633582 | 0.651467 | 0.634406 | 0.639818 | down |
| 4053 | LTBP2 | 7.51E-06 | 0.520363 | 0.48159 | 0.512281 | 0.504745 | down |
| 58525 | WIZ | 7.58E-06 | 0.560014 | 0.548588 | 0.571368 | 0.55999 | down |
| 1979 | EIF4EBP2 | 7.59E-06 | 0.485129 | 0.443073 | 0.496026 | 0.474743 | down |
| 8661 | EIF3A | 7.65E-06 | 0.332574 | 0.35889 | 0.3465 | 0.345988 | down |
| 3005 | H1F0 | 7.65E-06 | 0.624819 | 0.590812 | 0.628307 | 0.614646 | down |
| 1503 | CTPS1 | 7.73E-06 | 0.571183 | 0.540553 | 0.549699 | 0.553812 | down |
| 1431 | CS | 7.75E-06 | 0.417437 | 0.448088 | 0.449261 | 0.438262 | down |
| 11338 | U2AF2 | 7.75E-06 | 0.634202 | 0.614093 | 0.62456 | 0.624285 | down |
| 161291 | TMEM30B | 7.82E-06 | 0.547597 | 0.566309 | 0.580216 | 0.564707 | down |
| 149371 | EXOC8 | 7.82E-06 | 0.512216 | 0.558756 | 0.520964 | 0.530645 | down |
| 6924 | ELOA | 7.84E-06 | 0.611539 | 0.610394 | 0.565193 | 0.595709 | down |
| 27102 | EIF2AK1 | 7.86E-06 | 0.590298 | 0.534053 | 0.570605 | 0.564986 | down |
| 1459 | CSNK2A2 | 7.86E-06 | 0.582998 | 0.586175 | 0.565139 | 0.578104 | down |
| 9813 | EFCAB14 | 7.86E-06 | 0.490317 | 0.521235 | 0.535308 | 0.51562 | down |
| 4478 | MSN | 7.87E-06 | 0.450535 | 0.411969 | 0.462725 | 0.441743 | down |
| 9703 | KIAA0100 | 7.93E-06 | 0.482628 | 0.532313 | 0.506393 | 0.507111 | down |
| 11044 | TENT4A | 7.94E-06 | 0.454754 | 0.451872 | 0.408347 | 0.438324 | down |
| 80790 | CMIP | 7.95E-06 | 0.499757 | 0.444879 | 0.463449 | 0.469362 | down |
| 3837 | KPNB1 | 8E-06 | 0.478259 | 0.498768 | 0.491866 | 0.489631 | down |
| 5048 | PAFAH1B1 | 8E-06 | 0.558481 | 0.559243 | 0.596436 | 0.571387 | down |
| 79837 | PIP4K2C | 8.09E-06 | 0.595588 | 0.56813 | 0.579765 | 0.581161 | down |
| 25929 | GEMIN5 | 8.11E-06 | 0.488736 | 0.514879 | 0.542284 | 0.5153 | down |
| 54467 | ANKIB1 | 8.13E-06 | 0.545517 | 0.533838 | 0.567841 | 0.549065 | down |
| 8500 | PPFIA1 | 8.17E-06 | 0.531887 | 0.533166 | 0.544084 | 0.536379 | down |
| 23367 | LARP1 | 8.2E-06 | 0.427296 | 0.419994 | 0.393349 | 0.413546 | down |
| 51144 | HSD17B12 | 8.25E-06 | 0.525281 | 0.549521 | 0.525736 | 0.533513 | down |
| 28996 | HIPK2 | 8.26E-06 | 0.386529 | 0.425566 | 0.426934 | 0.41301 | down |
| 4781 | NFIB | 8.3E-06 | 0.651824 | 0.648593 | 0.641649 | 0.647355 | down |
| 5796 | PTPRK | 8.31E-06 | 0.489855 | 0.474011 | 0.5279 | 0.497256 | down |
| 143 | PARP4 | 8.31E-06 | 0.431271 | 0.421321 | 0.390476 | 0.414356 | down |
| 23516 | SLC39A14 | 8.34E-06 | 0.417435 | 0.451535 | 0.455885 | 0.441618 | down |
| 55893 | ZNF395 | 8.35E-06 | 0.557914 | 0.569007 | 0.602239 | 0.576387 | down |
| 6832 | SUPV3L1 | 8.37E-06 | 0.596728 | 0.636849 | 0.610815 | 0.614798 | down |
| 55841 | WWC3 | 8.44E-06 | 0.564318 | 0.510344 | 0.513704 | 0.529455 | down |
| 57619 | SHROOM3 | 8.51E-06 | 0.508493 | 0.487609 | 0.444809 | 0.480303 | down |
| 55626 | AMBRA1 | 8.55E-06 | 0.550097 | 0.559431 | 0.538778 | 0.549435 | down |
| 29940 | DSE | 8.57E-06 | 0.525271 | 0.555606 | 0.513328 | 0.531401 | down |
| 10771 | ZMYND11 | 8.66E-06 | 0.506106 | 0.527251 | 0.566636 | 0.533331 | down |
| 8614 | STC2 | 8.73E-06 | 0.467574 | 0.482758 | 0.505618 | 0.485317 | down |
| 4763 | NF1 | 8.76E-06 | 0.532978 | 0.557639 | 0.561288 | 0.550635 | down |
| 10238 | DCAF7 | 8.79E-06 | 0.55239 | 0.554343 | 0.544028 | 0.550254 | down |
| 23061 | TBC1D9B | 8.79E-06 | 0.606624 | 0.629469 | 0.575697 | 0.60393 | down |
| 1849 | DUSP7 | 8.86E-06 | 0.486709 | 0.485861 | 0.539152 | 0.503907 | down |
| 83752 | LONP2 | 8.93E-06 | 0.569562 | 0.571676 | 0.56361 | 0.568283 | down |
| 1200 | TPP1 | 8.95E-06 | 0.405402 | 0.424971 | 0.430661 | 0.420345 | down |
| 29117 | BRD7 | 8.96E-06 | 0.510296 | 0.564909 | 0.51293 | 0.529379 | down |
| 160760 | PPTC7 | 8.99E-06 | 0.441385 | 0.476918 | 0.418183 | 0.445495 | down |
| 90007 | MIDN | 9E-06 | 0.502684 | 0.454554 | 0.460266 | 0.472501 | down |
| 1314 | COPA | 9E-06 | 0.447579 | 0.477555 | 0.434844 | 0.453326 | down |
| 22870 | PPP6R1 | 9.05E-06 | 0.58519 | 0.566075 | 0.544402 | 0.565223 | down |

| | | | | | | | |
|---|---|---|---|---|---|---|---|
| 359948 | IRF2BP2 | 9.09E-06 | 0.615477 | 0.586173 | 0.604612 | 0.602087 | down |
| 9289 | ADGRG1 | 9.23E-06 | 0.54979 | 0.498223 | 0.500461 | 0.516158 | down |
| 22928 | SEPHS2 | 9.24E-06 | 0.565745 | 0.605256 | 0.572313 | 0.581105 | down |
| 81607 | NECTIN4 | 9.3E-06 | 0.459978 | 0.465518 | 0.471715 | 0.465737 | down |
| 9475 | ROCK2 | 9.35E-06 | 0.496167 | 0.506144 | 0.482362 | 0.494891 | down |
| 146691 | TOM1L2 | 9.38E-06 | 0.623699 | 0.64129 | 0.609074 | 0.624688 | down |
| 22841 | RAB11FIP2 | 9.39E-06 | 0.53208 | 0.543379 | 0.564589 | 0.546683 | down |
| 5934 | RBL2 | 9.4E-06 | 0.444979 | 0.479606 | 0.492675 | 0.47242 | down |
| 6294 | SAFB | 9.42E-06 | 0.354298 | 0.324321 | 0.328292 | 0.335637 | down |
| 23300 | ATMIN | 9.51E-06 | 0.611996 | 0.622509 | 0.615372 | 0.616626 | down |
| 10163 | WASF2 | 9.53E-06 | 0.507513 | 0.471647 | 0.491724 | 0.490295 | down |
| 8609 | KLF7 | 9.55E-06 | 0.395393 | 0.401589 | 0.386113 | 0.394365 | down |
| 57153 | SLC44A2 | 9.56E-06 | 0.626172 | 0.571084 | 0.592052 | 0.596436 | down |
| 4173 | MCM4 | 9.56E-06 | 0.508042 | 0.54026 | 0.557266 | 0.53519 | down |
| 59338 | PLEKHA1 | 9.63E-06 | 0.531984 | 0.527536 | 0.539875 | 0.533132 | down |
| 130535 | KCTD18 | 9.64E-06 | 0.488646 | 0.485028 | 0.483164 | 0.485613 | down |
| 25 | ABL1 | 9.66E-06 | 0.519121 | 0.508569 | 0.504523 | 0.510738 | down |
| 64324 | NSD1 | 9.69E-06 | 0.323993 | 0.372654 | 0.362732 | 0.353126 | down |
| 9819 | TSC22D2 | 9.7E-06 | 0.491739 | 0.499776 | 0.488662 | 0.493393 | down |
| 9898 | UBAP2L | 9.73E-06 | 0.50844 | 0.45541 | 0.465751 | 0.476534 | down |
| 79979 | TRMT2B | 9.79E-06 | 0.559061 | 0.587528 | 0.543853 | 0.563481 | down |
| 7082 | TJP1 | 9.79E-06 | 0.38178 | 0.397327 | 0.428962 | 0.40269 | down |
| 23095 | KIF1B | 9.79E-06 | 0.441607 | 0.474895 | 0.442015 | 0.452839 | down |
| 1981 | EIF4G1 | 9.82E-06 | 0.483859 | 0.466514 | 0.469183 | 0.473185 | down |
| 23175 | LPIN1 | 9.82E-06 | 0.457169 | 0.43697 | 0.427259 | 0.440466 | down |
| 79784 | MYH14 | 9.83E-06 | 0.353979 | 0.325877 | 0.293727 | 0.324528 | down |
| 23509 | POFUT1 | 9.84E-06 | 0.619433 | 0.573309 | 0.607965 | 0.600235 | down |
| 6929 | TCF3 | 9.84E-06 | 0.593915 | 0.54229 | 0.558815 | 0.565006 | down |
| 65123 | INTS3 | 9.87E-06 | 0.46422 | 0.434421 | 0.497066 | 0.465236 | down |
| 64062 | RBM26 | 9.9E-06 | 0.560284 | 0.568354 | 0.56862 | 0.565753 | down |
| 163 | AP2B1 | 9.92E-06 | 0.484127 | 0.489528 | 0.529809 | 0.501154 | down |
| 7171 | TPM4 | 9.93E-06 | 0.541128 | 0.503103 | 0.549228 | 0.531153 | down |
| 79811 | SLTM | 9.95E-06 | 0.496484 | 0.466627 | 0.441388 | 0.468166 | down |
| 23264 | ZC3H7B | 1.01E-05 | 0.508381 | 0.477966 | 0.54495 | 0.510432 | down |
| 9826 | ARHGEF11 | 1.01E-05 | 0.45469 | 0.42191 | 0.384132 | 0.420244 | down |
| 6812 | STXBP1 | 1.01E-05 | 0.514099 | 0.465592 | 0.468742 | 0.482811 | down |
| 9261 | MAPKAPK2 | 1.01E-05 | 0.62604 | 0.635611 | 0.598694 | 0.620115 | down |
| 6646 | SOAT1 | 1.01E-05 | 0.569987 | 0.564092 | 0.587109 | 0.573729 | down |
| 387680 | WASHC2A | 1.02E-05 | 0.517216 | 0.542566 | 0.55162 | 0.537134 | down |
| 5589 | PRKCSH | 1.02E-05 | 0.496966 | 0.503412 | 0.471957 | 0.490778 | down |
| 4645 | MYO5B | 1.02E-05 | 0.425893 | 0.444541 | 0.463465 | 0.444633 | down |
| 9927 | MFN2 | 1.03E-05 | 0.512827 | 0.552137 | 0.513786 | 0.52625 | down |
| 113419 | TEX261 | 1.03E-05 | 0.573303 | 0.565926 | 0.613469 | 0.584233 | down |
| 4288 | MKI67 | 1.03E-05 | 0.329029 | 0.337097 | 0.31634 | 0.327488 | down |
| 10468 | FST | 1.03E-05 | 0.588141 | 0.594506 | 0.60972 | 0.597456 | down |
| 5836 | PYGL | 1.03E-05 | 0.546897 | 0.556148 | 0.586815 | 0.563287 | down |
| 79956 | ERMP1 | 1.04E-05 | 0.511225 | 0.562465 | 0.529049 | 0.534247 | down |
| 1639 | DCTN1 | 1.04E-05 | 0.530368 | 0.499257 | 0.496429 | 0.508685 | down |
| 5584 | PRKCI | 1.04E-05 | 0.554866 | 0.525547 | 0.578076 | 0.552829 | down |
| 7107 | GPR137B | 1.04E-05 | 0.582479 | 0.624234 | 0.627096 | 0.61127 | down |
| 3099 | HK2 | 1.05E-05 | 0.406745 | 0.404056 | 0.394393 | 0.401731 | down |
| 8888 | MCM3AP | 1.05E-05 | 0.462095 | 0.465195 | 0.519796 | 0.482362 | down |
| 9730 | DCAF1 | 1.06E-05 | 0.451174 | 0.397843 | 0.402588 | 0.417202 | down |
| 57459 | GATAD2B | 1.06E-05 | 0.450139 | 0.466579 | 0.476556 | 0.464425 | down |
| 146050 | ZSCAN29 | 1.06E-05 | 0.484723 | 0.471735 | 0.496686 | 0.484382 | down |
| 57659 | ZBTB4 | 1.06E-05 | 0.496188 | 0.487034 | 0.550198 | 0.51114 | down |
| 9875 | URB1 | 1.07E-05 | 0.456585 | 0.489025 | 0.462651 | 0.469421 | down |
| 11167 | FSTL1 | 1.08E-05 | 0.544155 | 0.567519 | 0.591248 | 0.567641 | down |

| | | | | | | | |
|---|---|---|---|---|---|---|---|
| 55718 | POLR3E | 1.09E-05 | 0.558211 | 0.550426 | 0.555356 | 0.554664 | down |
| 9319 | TRIP13 | 1.09E-05 | 0.622075 | 0.656164 | 0.658698 | 0.645645 | down |
| 10397 | NDRG1 | 1.09E-05 | 0.552278 | 0.601868 | 0.566219 | 0.573455 | down |
| 1595 | CYP51A1 | 1.09E-05 | 0.526618 | 0.545533 | 0.508184 | 0.526779 | down |
| 22803 | XRN2 | 1.09E-05 | 0.505741 | 0.546894 | 0.531454 | 0.52803 | down |
| 9683 | N4BP1 | 1.1E-05 | 0.539214 | 0.552435 | 0.530924 | 0.540858 | down |
| 392 | ARHGAP1 | 1.1E-05 | 0.514809 | 0.502158 | 0.481948 | 0.499638 | down |
| 55752 | 44450 | 1.11E-05 | 0.431686 | 0.495698 | 0.485395 | 0.470926 | down |
| 4017 | LOXL2 | 1.11E-05 | 0.493253 | 0.471643 | 0.5223 | 0.495732 | down |
| 9961 | MVP | 1.11E-05 | 0.536834 | 0.494071 | 0.480686 | 0.503863 | down |
| 79602 | ADIPOR2 | 1.11E-05 | 0.521922 | 0.549399 | 0.543485 | 0.538269 | down |
| 4779 | NFE2L1 | 1.12E-05 | 0.503043 | 0.532005 | 0.512578 | 0.515875 | down |
| 5546 | PRCC | 1.12E-05 | 0.655577 | 0.636535 | 0.635569 | 0.64256 | down |
| 1718 | DHCR24 | 1.12E-05 | 0.566882 | 0.583392 | 0.570501 | 0.573591 | down |
| 64750 | SMURF2 | 1.13E-05 | 0.653759 | 0.623112 | 0.658742 | 0.645204 | down |
| 55665 | URGCP | 1.13E-05 | 0.54977 | 0.538672 | 0.490321 | 0.526254 | down |
| 57508 | INTS2 | 1.14E-05 | 0.411508 | 0.43016 | 0.363048 | 0.401572 | down |
| 285203 | EOGT | 1.14E-05 | 0.533047 | 0.546287 | 0.580078 | 0.553137 | down |
| 140838 | NANP | 1.15E-05 | 0.525726 | 0.571679 | 0.585903 | 0.561103 | down |
| 3655 | ITGA6 | 1.15E-05 | 0.39341 | 0.415202 | 0.418363 | 0.408992 | down |
| 11083 | DIDO1 | 1.16E-05 | 0.460968 | 0.407314 | 0.435423 | 0.434568 | down |
| 23199 | GSE1 | 1.16E-05 | 0.52638 | 0.489837 | 0.505962 | 0.507393 | down |
| 4771 | NF2 | 1.16E-05 | 0.431164 | 0.41984 | 0.487356 | 0.44612 | down |
| 6830 | SUPT6H | 1.16E-05 | 0.458143 | 0.412829 | 0.409926 | 0.426966 | down |
| 9124 | PDLIM1 | 1.16E-05 | 0.624259 | 0.67081 | 0.653479 | 0.649516 | down |
| 10036 | CHAF1A | 1.17E-05 | 0.512827 | 0.497014 | 0.489165 | 0.499669 | down |
| 60685 | ZFAND3 | 1.17E-05 | 0.515379 | 0.569212 | 0.550509 | 0.545033 | down |
| 5236 | PGM1 | 1.17E-05 | 0.635302 | 0.661326 | 0.684091 | 0.660239 | down |
| 55717 | WDR11 | 1.17E-05 | 0.469147 | 0.489066 | 0.536967 | 0.498393 | down |
| 8720 | MBTPS1 | 1.18E-05 | 0.508844 | 0.564862 | 0.560739 | 0.544815 | down |
| 56907 | SPIRE1 | 1.18E-05 | 0.413936 | 0.424683 | 0.429386 | 0.422668 | down |
| 10939 | AFG3L2 | 1.18E-05 | 0.596327 | 0.547756 | 0.601267 | 0.581783 | down |
| 9972 | NUP153 | 1.18E-05 | 0.388997 | 0.405503 | 0.412973 | 0.402491 | down |
| 4124 | MAN2A1 | 1.19E-05 | 0.499849 | 0.499773 | 0.477414 | 0.492345 | down |
| 7317 | UBA1 | 1.19E-05 | 0.584055 | 0.610308 | 0.548761 | 0.581041 | down |
| 5034 | P4HB | 1.19E-05 | 0.624608 | 0.63479 | 0.589237 | 0.616212 | down |
| 3187 | HNRNPH1 | 1.19E-05 | 0.562617 | 0.589095 | 0.587334 | 0.579682 | down |
| 3914 | LAMB3 | 1.19E-05 | 0.508006 | 0.497783 | 0.459191 | 0.488326 | down |
| 23463 | ICMT | 1.2E-05 | 0.650597 | 0.630974 | 0.616177 | 0.632583 | down |
| 9726 | ZNF646 | 1.21E-05 | 0.52018 | 0.483355 | 0.524832 | 0.509456 | down |
| 50628 | GEMIN4 | 1.22E-05 | 0.510393 | 0.549164 | 0.561788 | 0.540448 | down |
| 23139 | MAST2 | 1.22E-05 | 0.540811 | 0.524913 | 0.516911 | 0.527545 | down |
| 64682 | ANAPC1 | 1.22E-05 | 0.373477 | 0.433244 | 0.42751 | 0.41141 | down |
| 54978 | SLC35F6 | 1.22E-05 | 0.519233 | 0.488937 | 0.46354 | 0.49057 | down |
| 58191 | CXCL16 | 1.22E-05 | 0.578538 | 0.56482 | 0.575746 | 0.573035 | down |
| 5058 | PAK1 | 1.22E-05 | 0.598088 | 0.592246 | 0.640047 | 0.610127 | down |
| 4798 | NFRKB | 1.23E-05 | 0.482019 | 0.4735 | 0.449427 | 0.468315 | down |
| 4293 | MAP3K9 | 1.24E-05 | 0.447407 | 0.450738 | 0.396934 | 0.431693 | down |
| 8224 | SYN3 | 1.24E-05 | 0.447239 | 0.492672 | 0.426829 | 0.45558 | down |
| 54887 | UHRF1BP1 | 1.24E-05 | 0.481662 | 0.500885 | 0.533688 | 0.505412 | down |
| 7374 | UNG | 1.24E-05 | 0.601458 | 0.622516 | 0.564122 | 0.596032 | down |
| 8496 | PPFIBP1 | 1.24E-05 | 0.42417 | 0.459942 | 0.465982 | 0.450031 | down |
| 9076 | CLDN1 | 1.24E-05 | 0.55403 | 0.561423 | 0.605188 | 0.573547 | down |
| 144348 | ZNF664 | 1.24E-05 | 0.467216 | 0.489792 | 0.526048 | 0.494352 | down |
| 57583 | TMEM181 | 1.24E-05 | 0.494067 | 0.474263 | 0.517242 | 0.495191 | down |
| 2969 | GTF2I | 1.25E-05 | 0.517843 | 0.546827 | 0.551818 | 0.538829 | down |
| 10732 | TCFL5 | 1.25E-05 | 0.644821 | 0.587753 | 0.623728 | 0.618767 | down |
| 6241 | RRM2 | 1.25E-05 | 0.576316 | 0.576929 | 0.612526 | 0.58859 | down |

| | | | | | | | |
|---|---|---|---|---|---|---|---|
| 5083 | PAX9 | 1.26E-05 | 0.561995 | 0.584786 | 0.612267 | 0.586349 | down |
| 55291 | PPP6R3 | 1.26E-05 | 0.542781 | 0.567846 | 0.594565 | 0.568397 | down |
| 6574 | SLC20A1 | 1.26E-05 | 0.479495 | 0.505311 | 0.507166 | 0.497324 | down |
| 2033 | EP300 | 1.26E-05 | 0.364418 | 0.384663 | 0.425115 | 0.391399 | down |
| 79834 | PEAK1 | 1.27E-05 | 0.398267 | 0.423341 | 0.420991 | 0.414199 | down |
| 26005 | C2CD3 | 1.27E-05 | 0.371624 | 0.381606 | 0.421597 | 0.391609 | down |
| 11325 | DDX42 | 1.28E-05 | 0.429587 | 0.4453 | 0.423699 | 0.432862 | down |
| 51479 | ANKFY1 | 1.28E-05 | 0.413822 | 0.473025 | 0.413162 | 0.433336 | down |
| 51280 | GOLM1 | 1.28E-05 | 0.464872 | 0.475524 | 0.508213 | 0.48287 | down |
| 23339 | VPS39 | 1.28E-05 | 0.431462 | 0.419355 | 0.4864 | 0.445739 | down |
| 57658 | CALCOCO | 1.29E-05 | 0.523339 | 0.515517 | 0.572147 | 0.537001 | down |
| 23450 | SF3B3 | 1.29E-05 | 0.457663 | 0.493306 | 0.447098 | 0.466023 | down |
| 10521 | DDX17 | 1.29E-05 | 0.457546 | 0.475358 | 0.501606 | 0.47817 | down |
| 7862 | BRPF1 | 1.29E-05 | 0.447427 | 0.488533 | 0.47835 | 0.471437 | down |
| 146754 | DNAH2 | 1.29E-05 | 0.379121 | 0.416842 | 0.42 | 0.405321 | down |
| 2800 | GOLGA1 | 1.29E-05 | 0.54231 | 0.518066 | 0.576322 | 0.545566 | down |
| 23360 | FNBP4 | 1.3E-05 | 0.492855 | 0.495611 | 0.448585 | 0.479017 | down |
| 23420 | NOMO1 | 1.3E-05 | 0.519538 | 0.54123 | 0.510769 | 0.523846 | down |
| 25939 | SAMHD1 | 1.3E-05 | 0.562343 | 0.556211 | 0.584279 | 0.567611 | down |
| 4076 | CAPRIN1 | 1.31E-05 | 0.545506 | 0.588974 | 0.597987 | 0.577489 | down |
| 10106 | CTDSP2 | 1.31E-05 | 0.510857 | 0.453984 | 0.496438 | 0.487093 | down |
| 220 | ALDH1A3 | 1.31E-05 | 0.426948 | 0.466358 | 0.451531 | 0.448279 | down |
| 22879 | MON1B | 1.32E-05 | 0.578584 | 0.586049 | 0.557185 | 0.573939 | down |
| 1040 | CDS1 | 1.32E-05 | 0.620864 | 0.593306 | 0.648438 | 0.620869 | down |
| 71 | ACTG1 | 1.32E-05 | 0.620471 | 0.649141 | 0.615976 | 0.62853 | down |
| 8985 | PLOD3 | 1.33E-05 | 0.586306 | 0.606854 | 0.560578 | 0.584579 | down |
| 84706 | GPT2 | 1.33E-05 | 0.655906 | 0.663654 | 0.631678 | 0.650413 | down |
| 1997 | ELF1 | 1.33E-05 | 0.563444 | 0.546832 | 0.601344 | 0.57054 | down |
| 5747 | PTK2 | 1.33E-05 | 0.474772 | 0.487777 | 0.49119 | 0.48458 | down |
| 54625 | PARP14 | 1.34E-05 | 0.395696 | 0.457811 | 0.414928 | 0.422812 | down |
| 8394 | PIP5K1A | 1.35E-05 | 0.550522 | 0.528752 | 0.566857 | 0.54871 | down |
| 59349 | KLHL12 | 1.36E-05 | 0.505668 | 0.514916 | 0.54342 | 0.521335 | down |
| 10342 | TFG | 1.36E-05 | 0.570617 | 0.573057 | 0.623036 | 0.588903 | down |
| 8555 | CDC14B | 1.37E-05 | 0.433436 | 0.487553 | 0.462362 | 0.461117 | down |
| 3192 | HNRNPU | 1.37E-05 | 0.469084 | 0.480633 | 0.487552 | 0.47909 | down |
| 23 | ABCF1 | 1.37E-05 | 0.557448 | 0.551256 | 0.506573 | 0.538425 | down |
| 55041 | PLEKHB2 | 1.41E-05 | 0.548597 | 0.535716 | 0.531494 | 0.538602 | down |
| 4343 | MOV10 | 1.42E-05 | 0.629693 | 0.576944 | 0.589256 | 0.598631 | down |
| 51429 | SNX9 | 1.42E-05 | 0.563252 | 0.581781 | 0.589595 | 0.578209 | down |
| 376497 | SLC27A1 | 1.43E-05 | 0.576132 | 0.554776 | 0.604828 | 0.578579 | down |
| 8648 | NCOA1 | 1.43E-05 | 0.406721 | 0.380923 | 0.457699 | 0.415114 | down |
| 10902 | BRD8 | 1.44E-05 | 0.533694 | 0.470829 | 0.514034 | 0.506186 | down |
| 2932 | GSK3B | 1.44E-05 | 0.569674 | 0.533613 | 0.540461 | 0.547916 | down |
| 5073 | PARN | 1.44E-05 | 0.549982 | 0.561001 | 0.552242 | 0.554408 | down |
| 51421 | AMOTL2 | 1.45E-05 | 0.447697 | 0.447511 | 0.482251 | 0.459153 | down |
| 8085 | KMT2D | 1.45E-05 | 0.486689 | 0.450042 | 0.448793 | 0.461841 | down |
| 8729 | GBF1 | 1.45E-05 | 0.464104 | 0.442138 | 0.41416 | 0.440134 | down |
| 23028 | KDM1A | 1.46E-05 | 0.44035 | 0.475468 | 0.423775 | 0.446531 | down |
| 4691 | NCL | 1.47E-05 | 0.537884 | 0.561875 | 0.572418 | 0.557393 | down |
| 10236 | HNRNPR | 1.47E-05 | 0.503578 | 0.497218 | 0.552776 | 0.517857 | down |
| 4548 | MTR | 1.47E-05 | 0.424207 | 0.432439 | 0.448983 | 0.43521 | down |
| 3038 | HAS3 | 1.47E-05 | 0.501182 | 0.543769 | 0.504861 | 0.516604 | down |
| 50848 | F11R | 1.48E-05 | 0.624305 | 0.65935 | 0.671416 | 0.65169 | down |
| 4430 | MYO1B | 1.48E-05 | 0.529182 | 0.538779 | 0.584467 | 0.550809 | down |
| 23230 | VPS13A | 1.48E-05 | 0.665188 | 0.622623 | 0.630649 | 0.639487 | down |
| 4642 | MYO1D | 1.48E-05 | 0.465098 | 0.419927 | 0.455828 | 0.446951 | down |
| 23365 | ARHGEF12 | 1.49E-05 | 0.407653 | 0.447785 | 0.423713 | 0.426384 | down |
| 9092 | SART1 | 1.49E-05 | 0.438205 | 0.370058 | 0.379329 | 0.395864 | down |

| | | | | | | | |
|---|---|---|---|---|---|---|---|
| 5774 | PTPN3 | 1.49E-05 | 0.391055 | 0.420077 | 0.398155 | 0.403096 | down |
| 23765 | IL17RA | 1.49E-05 | 0.563994 | 0.522918 | 0.500546 | 0.529153 | down |
| 9209 | LRRFIP2 | 1.5E-05 | 0.592349 | 0.59496 | 0.602701 | 0.59667 | down |
| 85440 | DOCK7 | 1.5E-05 | 0.532486 | 0.504304 | 0.535363 | 0.524051 | down |
| 2801 | GOLGA2 | 1.5E-05 | 0.446851 | 0.412945 | 0.404243 | 0.421346 | down |
| 613 | BCR | 1.5E-05 | 0.560176 | 0.544426 | 0.581419 | 0.562007 | down |
| 51199 | NIN | 1.51E-05 | 0.419505 | 0.37692 | 0.381568 | 0.392664 | down |
| 55699 | IARS2 | 1.51E-05 | 0.525615 | 0.570298 | 0.586118 | 0.560677 | down |
| 27044 | SND1 | 1.52E-05 | 0.584813 | 0.585062 | 0.543763 | 0.571213 | down |
| 63897 | HEATR6 | 1.52E-05 | 0.442483 | 0.468616 | 0.454272 | 0.455124 | down |
| 9904 | RBM19 | 1.52E-05 | 0.54967 | 0.498024 | 0.474968 | 0.507554 | down |
| 9761 | MLEC | 1.52E-05 | 0.527071 | 0.532296 | 0.565109 | 0.541492 | down |
| 7011 | TEP1 | 1.53E-05 | 0.5328 | 0.523238 | 0.514196 | 0.523411 | down |
| 11140 | CDC37 | 1.53E-05 | 0.650117 | 0.60556 | 0.614262 | 0.623313 | down |
| 55122 | AKIRIN2 | 1.54E-05 | 0.651003 | 0.621763 | 0.671189 | 0.647985 | down |
| 1316 | KLF6 | 1.55E-05 | 0.499995 | 0.506575 | 0.511825 | 0.506131 | down |
| 55852 | TEX2 | 1.55E-05 | 0.548228 | 0.570232 | 0.505372 | 0.541277 | down |
| 8675 | STX16 | 1.55E-05 | 0.439359 | 0.510128 | 0.481669 | 0.477052 | down |
| 790 | CAD | 1.55E-05 | 0.504716 | 0.512823 | 0.446559 | 0.488033 | down |
| 4238 | MFAP3 | 1.55E-05 | 0.611011 | 0.587667 | 0.624157 | 0.607612 | down |
| 23193 | GANAB | 1.56E-05 | 0.574582 | 0.570555 | 0.581204 | 0.575447 | down |
| 9601 | PDIA4 | 1.56E-05 | 0.600695 | 0.627704 | 0.65487 | 0.627756 | down |
| 987 | LRBA | 1.57E-05 | 0.417278 | 0.359552 | 0.437945 | 0.404925 | down |
| 651746 | ANKRD33E | 1.58E-05 | 0.393036 | 0.425393 | 0.430698 | 0.416375 | down |
| 4926 | NUMA1 | 1.58E-05 | 0.40021 | 0.421603 | 0.396641 | 0.406151 | down |
| 7871 | SLMAP | 1.59E-05 | 0.546522 | 0.537305 | 0.555389 | 0.546405 | down |
| 55571 | CNOT11 | 1.59E-05 | 0.63339 | 0.619964 | 0.585376 | 0.61291 | down |
| 10919 | EHMT2 | 1.59E-05 | 0.542348 | 0.583313 | 0.577771 | 0.567811 | down |
| 10498 | CARM1 | 1.59E-05 | 0.599629 | 0.615554 | 0.585923 | 0.600369 | down |
| 5663 | PSEN1 | 1.59E-05 | 0.656106 | 0.648614 | 0.682228 | 0.662316 | down |
| 3097 | HIVEP2 | 1.6E-05 | 0.391979 | 0.324132 | 0.323638 | 0.346583 | down |
| 6654 | SOS1 | 1.6E-05 | 0.383119 | 0.396975 | 0.41045 | 0.396848 | down |
| 25942 | SIN3A | 1.6E-05 | 0.401487 | 0.427387 | 0.376707 | 0.40186 | down |
| 203197 | TMEM268 | 1.6E-05 | 0.494465 | 0.469934 | 0.546437 | 0.503612 | down |
| 9343 | EFTUD2 | 1.6E-05 | 0.597873 | 0.597099 | 0.616547 | 0.60384 | down |
| 9555 | H2AFY | 1.61E-05 | 0.612782 | 0.629317 | 0.619821 | 0.62064 | down |
| 55253 | TYW1 | 1.62E-05 | 0.448827 | 0.431968 | 0.507096 | 0.46263 | down |
| 6242 | RTKN | 1.63E-05 | 0.61813 | 0.591401 | 0.608553 | 0.606028 | down |
| 537 | ATP6AP1 | 1.63E-05 | 0.650863 | 0.663661 | 0.666756 | 0.660426 | down |
| 29927 | SEC61A1 | 1.63E-05 | 0.524358 | 0.56412 | 0.537066 | 0.541848 | down |
| 10483 | SEC23B | 1.64E-05 | 0.551073 | 0.587375 | 0.555697 | 0.564715 | down |
| 2805 | GOT1 | 1.64E-05 | 0.599761 | 0.585887 | 0.592506 | 0.592718 | down |
| 55214 | P3H2 | 1.64E-05 | 0.561944 | 0.49439 | 0.550149 | 0.535494 | down |
| 10848 | PPP1R13L | 1.64E-05 | 0.639789 | 0.57888 | 0.599243 | 0.605971 | down |
| 23223 | RRP12 | 1.64E-05 | 0.535757 | 0.521849 | 0.47585 | 0.511152 | down |
| 811 | CALR | 1.65E-05 | 0.57623 | 0.546684 | 0.550564 | 0.557826 | down |
| 2806 | GOT2 | 1.65E-05 | 0.597808 | 0.621775 | 0.558519 | 0.592701 | down |
| 182 | JAG1 | 1.65E-05 | 0.470018 | 0.54014 | 0.526349 | 0.512169 | down |
| 22954 | TRIM32 | 1.65E-05 | 0.536704 | 0.533057 | 0.539187 | 0.536316 | down |
| 2036 | EPB41L1 | 1.66E-05 | 0.49734 | 0.481292 | 0.505383 | 0.494672 | down |
| 201626 | PDE12 | 1.66E-05 | 0.509779 | 0.543014 | 0.498725 | 0.517173 | down |
| 51132 | RLIM | 1.67E-05 | 0.589569 | 0.62859 | 0.648128 | 0.622095 | down |
| 55870 | ASH1L | 1.67E-05 | 0.348178 | 0.353666 | 0.409159 | 0.370334 | down |
| 27125 | AFF4 | 1.67E-05 | 0.476467 | 0.533593 | 0.53365 | 0.51457 | down |
| 23673 | STX12 | 1.68E-05 | 0.600958 | 0.599493 | 0.63688 | 0.612444 | down |
| 10723 | SLC12A7 | 1.68E-05 | 0.579972 | 0.550466 | 0.583981 | 0.571473 | down |
| 4627 | MYH9 | 1.68E-05 | 0.307887 | 0.313155 | 0.291402 | 0.304148 | down |
| 80012 | PHC3 | 1.68E-05 | 0.383798 | 0.420231 | 0.429231 | 0.411087 | down |

| | | | | | | | |
|---|---|---|---|---|---|---|---|
| 57178 | ZMIZ1 | 1.69E-05 | 0.487597 | 0.429896 | 0.443885 | 0.453793 | down |
| 6602 | SMARCD1 | 1.69E-05 | 0.45016 | 0.447336 | 0.471565 | 0.456354 | down |
| 23211 | ZC3H4 | 1.7E-05 | 0.459742 | 0.436425 | 0.450506 | 0.448891 | down |
| 11052 | CPSF6 | 1.7E-05 | 0.669513 | 0.640997 | 0.62655 | 0.645687 | down |
| 56829 | ZC3HAV1 | 1.71E-05 | 0.411062 | 0.479941 | 0.429413 | 0.440138 | down |
| 2114 | ETS2 | 1.71E-05 | 0.453777 | 0.501699 | 0.515291 | 0.490256 | down |
| 81608 | FIP1L1 | 1.72E-05 | 0.536676 | 0.56933 | 0.521092 | 0.542366 | down |
| 55696 | RBM22 | 1.72E-05 | 0.594041 | 0.612295 | 0.603372 | 0.603236 | down |
| 1660 | DHX9 | 1.73E-05 | 0.476477 | 0.471691 | 0.526958 | 0.491709 | down |
| 6272 | SORT1 | 1.74E-05 | 0.495916 | 0.54302 | 0.496167 | 0.511701 | down |
| 160 | AP2A1 | 1.77E-05 | 0.584707 | 0.612188 | 0.567725 | 0.588207 | down |
| 54700 | RRN3 | 1.77E-05 | 0.586159 | 0.635917 | 0.603415 | 0.608497 | down |
| 56894 | AGPAT3 | 1.77E-05 | 0.557734 | 0.556989 | 0.578606 | 0.564443 | down |
| 7052 | TGM2 | 1.77E-05 | 0.530798 | 0.540989 | 0.523206 | 0.531664 | down |
| 7150 | TOP1 | 1.78E-05 | 0.46615 | 0.509123 | 0.509455 | 0.494909 | down |
| 80279 | CDK5RAP3 | 1.78E-05 | 0.578143 | 0.558536 | 0.603165 | 0.579948 | down |
| 3936 | LCP1 | 1.78E-05 | 0.561371 | 0.614163 | 0.597744 | 0.591092 | down |
| 5770 | PTPN1 | 1.79E-05 | 0.540288 | 0.504925 | 0.56109 | 0.535434 | down |
| 8812 | CCNK | 1.8E-05 | 0.516351 | 0.445287 | 0.470941 | 0.477526 | down |
| 54902 | TTC19 | 1.8E-05 | 0.583434 | 0.549138 | 0.596543 | 0.576372 | down |
| 1663 | DDX11 | 1.82E-05 | 0.544433 | 0.519113 | 0.540192 | 0.534579 | down |
| 55683 | KANSL3 | 1.82E-05 | 0.52752 | 0.513125 | 0.494729 | 0.511791 | down |
| 11113 | CIT | 1.85E-05 | 0.325787 | 0.375743 | 0.321366 | 0.340965 | down |
| 9646 | CTR9 | 1.85E-05 | 0.596775 | 0.600934 | 0.585629 | 0.594446 | down |
| 116840 | CNTROB | 1.86E-05 | 0.597737 | 0.594247 | 0.64283 | 0.611605 | down |
| 2799 | GNS | 1.87E-05 | 0.467794 | 0.479389 | 0.462172 | 0.469785 | down |
| 117246 | FTSJ3 | 1.87E-05 | 0.471162 | 0.430038 | 0.386479 | 0.429227 | down |
| 7525 | YES1 | 1.88E-05 | 0.532401 | 0.526332 | 0.558739 | 0.539157 | down |
| 9110 | MTMR4 | 1.88E-05 | 0.473167 | 0.517729 | 0.541473 | 0.510789 | down |
| 5768 | QSOX1 | 1.88E-05 | 0.632136 | 0.57192 | 0.604557 | 0.602871 | down |
| 23254 | KAZN | 1.89E-05 | 0.508279 | 0.483535 | 0.503413 | 0.498409 | down |
| 64787 | EPS8L2 | 1.89E-05 | 0.667117 | 0.627442 | 0.644412 | 0.646324 | down |
| 26000 | TBC1D10B | 1.9E-05 | 0.577695 | 0.548467 | 0.521129 | 0.549097 | down |
| 10531 | PITRM1 | 1.9E-05 | 0.513028 | 0.53367 | 0.460642 | 0.502447 | down |
| 2131 | EXT1 | 1.9E-05 | 0.542422 | 0.507244 | 0.507683 | 0.519116 | down |
| 3030 | HADHA | 1.91E-05 | 0.598979 | 0.588343 | 0.607977 | 0.598433 | down |
| 8848 | TSC22D1 | 1.92E-05 | 0.650447 | 0.639106 | 0.631009 | 0.640187 | down |
| 9858 | PPP1R26 | 1.92E-05 | 0.459212 | 0.431576 | 0.438001 | 0.44293 | down |
| 81831 | NETO2 | 1.92E-05 | 0.571658 | 0.580726 | 0.532171 | 0.561518 | down |
| 79755 | ZNF750 | 1.93E-05 | 0.493037 | 0.43392 | 0.448717 | 0.458558 | down |
| 10160 | FARP1 | 1.93E-05 | 0.482347 | 0.500384 | 0.509589 | 0.49744 | down |
| 55689 | YEATS2 | 1.93E-05 | 0.396985 | 0.475747 | 0.393213 | 0.421982 | down |
| 25820 | ARIH1 | 1.94E-05 | 0.495038 | 0.489019 | 0.550991 | 0.511683 | down |
| 9567 | GTPBP1 | 1.94E-05 | 0.523019 | 0.445155 | 0.487542 | 0.485239 | down |
| 2956 | MSH6 | 1.94E-05 | 0.496911 | 0.545861 | 0.562713 | 0.535162 | down |
| 4853 | NOTCH2 | 1.94E-05 | 0.421648 | 0.425759 | 0.407606 | 0.418338 | down |
| 51341 | ZBTB7A | 1.95E-05 | 0.593899 | 0.562439 | 0.52199 | 0.559443 | down |
| 89970 | RSPRY1 | 1.95E-05 | 0.630275 | 0.659836 | 0.671603 | 0.653905 | down |
| 29775 | CARD10 | 1.95E-05 | 0.573081 | 0.496652 | 0.530386 | 0.533373 | down |
| 23268 | DNMBP | 1.96E-05 | 0.346521 | 0.370813 | 0.418714 | 0.378683 | down |
| 3915 | LAMC1 | 1.97E-05 | 0.462596 | 0.484074 | 0.483636 | 0.476769 | down |
| 6949 | TCOF1 | 1.97E-05 | 0.476224 | 0.4076 | 0.417056 | 0.433627 | down |
| 267 | AMFR | 1.98E-05 | 0.52165 | 0.550153 | 0.535649 | 0.535817 | down |
| 51582 | AZIN1 | 1.99E-05 | 0.55342 | 0.585347 | 0.597604 | 0.578791 | down |
| 1108 | CHD4 | 2.01E-05 | 0.418483 | 0.463986 | 0.401923 | 0.428131 | down |
| 7851 | MALL | 2.01E-05 | 0.591346 | 0.619697 | 0.619112 | 0.610052 | down |
| 124245 | ZC3H18 | 2.01E-05 | 0.537591 | 0.510153 | 0.53116 | 0.526302 | down |
| 284119 | CAVIN1 | 2.01E-05 | 0.490433 | 0.489407 | 0.477928 | 0.485923 | down |

| | | | | | | | |
|---|---|---|---|---|---|---|---|
| 55922 | NKRF | 2.01E-05 | 0.627447 | 0.634622 | 0.665263 | 0.642444 | down |
| 23191 | CYFIP1 | 2.02E-05 | 0.493982 | 0.535924 | 0.511784 | 0.513897 | down |
| 6548 | SLC9A1 | 2.03E-05 | 0.536815 | 0.472776 | 0.486263 | 0.498618 | down |
| 8662 | EIF3B | 2.03E-05 | 0.624255 | 0.590896 | 0.584457 | 0.599869 | down |
| 3313 | HSPA9 | 2.03E-05 | 0.571045 | 0.568923 | 0.567376 | 0.569115 | down |
| 24149 | ZNF318 | 2.03E-05 | 0.452387 | 0.420852 | 0.431847 | 0.435029 | down |
| 1956 | EGFR | 2.03E-05 | 0.335254 | 0.354657 | 0.36911 | 0.353007 | down |
| 2802 | GOLGA3 | 2.04E-05 | 0.431849 | 0.501109 | 0.48909 | 0.474016 | down |
| 116447 | TOP1MT | 2.05E-05 | 0.556503 | 0.517338 | 0.478151 | 0.51733 | down |
| 56243 | KIAA1217 | 2.06E-05 | 0.387452 | 0.338456 | 0.370845 | 0.365585 | down |
| 2186 | BPTF | 2.07E-05 | 0.411578 | 0.455062 | 0.441905 | 0.436181 | down |
| 5494 | PPM1A | 2.07E-05 | 0.57767 | 0.581499 | 0.602204 | 0.587125 | down |
| 1107 | CHD3 | 2.08E-05 | 0.434366 | 0.433301 | 0.409979 | 0.425882 | down |
| 283149 | BCL9L | 2.08E-05 | 0.531839 | 0.468141 | 0.528616 | 0.509532 | down |
| 1778 | DYNC1H1 | 2.08E-05 | 0.343644 | 0.381127 | 0.365019 | 0.363263 | down |
| 29954 | POMT2 | 2.09E-05 | 0.614498 | 0.588783 | 0.56816 | 0.59048 | down |
| 7458 | EIF4H | 2.09E-05 | 0.489211 | 0.458606 | 0.520559 | 0.489459 | down |
| 80209 | PROSER1 | 2.1E-05 | 0.391333 | 0.387711 | 0.42492 | 0.401321 | down |
| 7127 | TNFAIP2 | 2.1E-05 | 0.509094 | 0.5449 | 0.512352 | 0.522116 | down |
| 23352 | UBR4 | 2.1E-05 | 0.358231 | 0.376659 | 0.376232 | 0.370374 | down |
| 79098 | C1orf116 | 2.1E-05 | 0.516968 | 0.484211 | 0.465444 | 0.488874 | down |
| 65985 | AACS | 2.1E-05 | 0.5432 | 0.598313 | 0.597674 | 0.579729 | down |
| 5609 | MAP2K7 | 2.11E-05 | 0.602167 | 0.542781 | 0.578779 | 0.574575 | down |
| 81567 | TXNDC5 | 2.11E-05 | 0.636368 | 0.67438 | 0.610917 | 0.640555 | down |
| 123169 | LEO1 | 2.11E-05 | 0.629198 | 0.625277 | 0.581704 | 0.61206 | down |
| 23370 | ARHGEF18 | 2.11E-05 | 0.53542 | 0.499931 | 0.490007 | 0.508453 | down |
| 56893 | UBQLN4 | 2.12E-05 | 0.583132 | 0.614208 | 0.559168 | 0.585503 | down |
| 23225 | NUP210 | 2.12E-05 | 0.52198 | 0.468478 | 0.474624 | 0.48836 | down |
| 3096 | HIVEP1 | 2.12E-05 | 0.320992 | 0.329061 | 0.315895 | 0.321982 | down |
| 1627 | DBN1 | 2.13E-05 | 0.499426 | 0.53084 | 0.45633 | 0.495532 | down |
| 6692 | SPINT1 | 2.13E-05 | 0.654689 | 0.613335 | 0.590241 | 0.619422 | down |
| 8243 | SMC1A | 2.13E-05 | 0.421907 | 0.371258 | 0.372741 | 0.388635 | down |
| 2768 | GNA12 | 2.13E-05 | 0.611407 | 0.62986 | 0.598284 | 0.613184 | down |
| 54928 | IMPAD1 | 2.13E-05 | 0.578553 | 0.586868 | 0.594513 | 0.586644 | down |
| 22873 | DZIP1 | 2.14E-05 | 0.54224 | 0.523567 | 0.565399 | 0.543735 | down |
| 22994 | CEP131 | 2.14E-05 | 0.410496 | 0.412676 | 0.359371 | 0.394181 | down |
| 1739 | DLG1 | 2.14E-05 | 0.618765 | 0.619067 | 0.602187 | 0.61334 | down |
| 23306 | NEMP1 | 2.14E-05 | 0.559653 | 0.525196 | 0.58282 | 0.555889 | down |
| 54946 | SLC41A3 | 2.15E-05 | 0.648565 | 0.641678 | 0.680705 | 0.656983 | down |
| 488 | ATP2A2 | 2.16E-05 | 0.396983 | 0.453851 | 0.409476 | 0.420103 | down |
| 54861 | SNRK | 2.17E-05 | 0.464951 | 0.544713 | 0.512734 | 0.507466 | down |
| 10042 | HMGXB4 | 2.17E-05 | 0.566629 | 0.548931 | 0.594929 | 0.570163 | down |
| 5594 | MAPK1 | 2.18E-05 | 0.487631 | 0.453735 | 0.490441 | 0.477269 | down |
| 8289 | ARID1A | 2.18E-05 | 0.402308 | 0.413782 | 0.396882 | 0.404324 | down |
| 257397 | TAB3 | 2.19E-05 | 0.484316 | 0.432575 | 0.414262 | 0.443718 | down |
| 57189 | KIAA1147 | 2.2E-05 | 0.383175 | 0.428843 | 0.40107 | 0.404362 | down |
| 5654 | HTRA1 | 2.21E-05 | 0.64602 | 0.612888 | 0.639139 | 0.632682 | down |
| 10291 | SF3A1 | 2.22E-05 | 0.505695 | 0.42847 | 0.443601 | 0.459256 | down |
| 728642 | CDK11A | 2.23E-05 | 0.502471 | 0.509822 | 0.477488 | 0.496593 | down |
| 23536 | ADAT1 | 2.23E-05 | 0.66593 | 0.652642 | 0.616448 | 0.645006 | down |
| 84132 | USP42 | 2.23E-05 | 0.513096 | 0.54422 | 0.505618 | 0.520978 | down |
| 81839 | VANGL1 | 2.24E-05 | 0.567172 | 0.595579 | 0.607643 | 0.590131 | down |
| 23169 | SLC35D1 | 2.24E-05 | 0.456832 | 0.519007 | 0.443994 | 0.473278 | down |
| 51400 | PPME1 | 2.24E-05 | 0.552497 | 0.543711 | 0.562687 | 0.552965 | down |
| 1745 | DLX1 | 2.25E-05 | 0.669309 | 0.64141 | 0.621359 | 0.644026 | down |
| 57610 | RANBP10 | 2.25E-05 | 0.483722 | 0.527723 | 0.544304 | 0.518583 | down |
| 84985 | FAM83A | 2.25E-05 | 0.629033 | 0.592122 | 0.596149 | 0.605768 | down |
| 91369 | ANKRD40 | 2.25E-05 | 0.531204 | 0.572123 | 0.583133 | 0.562153 | down |

| | | | | | | | |
|---|---|---|---|---|---|---|---|
| 148789 | B3GALNT2 | 2.26E-05 | 0.438431 | 0.449722 | 0.425294 | 0.437816 | down |
| 6382 | SDC1 | 2.28E-05 | 0.615057 | 0.574658 | 0.596045 | 0.595253 | down |
| 821 | CANX | 2.29E-05 | 0.539091 | 0.578734 | 0.576023 | 0.564616 | down |
| 4175 | MCM6 | 2.29E-05 | 0.61007 | 0.625706 | 0.558034 | 0.597937 | down |
| 3184 | HNRNPD | 2.29E-05 | 0.557257 | 0.513004 | 0.581001 | 0.550421 | down |
| 57606 | SLAIN2 | 2.29E-05 | 0.526256 | 0.519036 | 0.516605 | 0.520632 | down |
| 26058 | GIGYF2 | 2.29E-05 | 0.446356 | 0.451152 | 0.424853 | 0.440787 | down |
| 6522 | SLC4A2 | 2.3E-05 | 0.667899 | 0.648894 | 0.635742 | 0.650845 | down |
| 30849 | PIK3R4 | 2.3E-05 | 0.49375 | 0.44145 | 0.448869 | 0.461357 | down |
| 8189 | SYMPK | 2.31E-05 | 0.553444 | 0.554262 | 0.504228 | 0.537311 | down |
| 55666 | NPLOC4 | 2.32E-05 | 0.676479 | 0.631779 | 0.626118 | 0.644792 | down |
| 23214 | XPO6 | 2.32E-05 | 0.534496 | 0.557444 | 0.56762 | 0.553187 | down |
| 221092 | HNRNPUL | 2.33E-05 | 0.463187 | 0.462795 | 0.435771 | 0.453918 | down |
| 10724 | OGA | 2.33E-05 | 0.474694 | 0.515906 | 0.482523 | 0.491041 | down |
| 4591 | TRIM37 | 2.33E-05 | 0.607403 | 0.605103 | 0.587919 | 0.600142 | down |
| 2720 | GLB1 | 2.34E-05 | 0.570997 | 0.567602 | 0.604423 | 0.581007 | down |
| 2316 | FLNA | 2.34E-05 | 0.531885 | 0.531743 | 0.504385 | 0.522671 | down |
| 11091 | WDR5 | 2.36E-05 | 0.592922 | 0.639237 | 0.575517 | 0.602559 | down |
| 153562 | MARVELD | 2.36E-05 | 0.419236 | 0.499312 | 0.419481 | 0.44601 | down |
| 1355 | COX15 | 2.36E-05 | 0.560531 | 0.553761 | 0.607353 | 0.573882 | down |
| 84717 | HDGFL2 | 2.37E-05 | 0.395082 | 0.347103 | 0.371529 | 0.371238 | down |
| 899 | CCNF | 2.37E-05 | 0.54353 | 0.567691 | 0.577999 | 0.563073 | down |
| 23649 | POLA2 | 2.37E-05 | 0.644498 | 0.629797 | 0.642439 | 0.638911 | down |
| 10075 | HUWE1 | 2.37E-05 | 0.369703 | 0.389217 | 0.382223 | 0.380381 | down |
| 64779 | MTHFSD | 2.37E-05 | 0.626727 | 0.607117 | 0.61843 | 0.617425 | down |
| 348995 | NUP43 | 2.38E-05 | 0.634998 | 0.643925 | 0.676307 | 0.651743 | down |
| 8672 | EIF4G3 | 2.39E-05 | 0.439246 | 0.445098 | 0.441042 | 0.441795 | down |
| 57805 | CCAR2 | 2.39E-05 | 0.551227 | 0.557097 | 0.498426 | 0.535584 | down |
| 4839 | NOP2 | 2.4E-05 | 0.678193 | 0.631297 | 0.660771 | 0.656754 | down |
| 23528 | ZNF281 | 2.4E-05 | 0.462175 | 0.474922 | 0.518753 | 0.485283 | down |
| 9019 | MPZL1 | 2.4E-05 | 0.514253 | 0.559868 | 0.58826 | 0.554127 | down |
| 3930 | LBR | 2.41E-05 | 0.644085 | 0.67733 | 0.632199 | 0.651205 | down |
| 171023 | ASXL1 | 2.41E-05 | 0.673468 | 0.632264 | 0.677323 | 0.661018 | down |
| 6829 | SUPT5H | 2.42E-05 | 0.490899 | 0.487332 | 0.508482 | 0.495571 | down |
| 9590 | AKAP12 | 2.42E-05 | 0.396003 | 0.426361 | 0.434156 | 0.41884 | down |
| 55140 | ELP3 | 2.42E-05 | 0.546355 | 0.549485 | 0.572621 | 0.556154 | down |
| 25831 | HECTD1 | 2.42E-05 | 0.490194 | 0.444678 | 0.520347 | 0.485073 | down |
| 55209 | SETD5 | 2.43E-05 | 0.427002 | 0.462028 | 0.496207 | 0.461745 | down |
| 37 | ACADVL | 2.44E-05 | 0.636946 | 0.641675 | 0.613008 | 0.630543 | down |
| 11336 | EXOC3 | 2.44E-05 | 0.597996 | 0.58778 | 0.620392 | 0.602056 | down |
| 5738 | PTGFRN | 2.45E-05 | 0.425412 | 0.455616 | 0.413705 | 0.431578 | down |
| 9968 | MED12 | 2.45E-05 | 0.408422 | 0.385626 | 0.407845 | 0.400631 | down |
| 2778 | GNAS | 2.46E-05 | 0.662887 | 0.672622 | 0.618604 | 0.651371 | down |
| 3309 | HSPA5 | 2.48E-05 | 0.600222 | 0.612472 | 0.607876 | 0.606856 | down |
| 1457 | CSNK2A1 | 2.48E-05 | 0.512249 | 0.469117 | 0.508208 | 0.496525 | down |
| 6047 | RNF4 | 2.49E-05 | 0.585768 | 0.540018 | 0.598222 | 0.574669 | down |
| 3303 | HSPA1A | 2.49E-05 | 0.414716 | 0.446423 | 0.413618 | 0.424919 | down |
| 64430 | PCNX4 | 2.5E-05 | 0.613611 | 0.630846 | 0.651291 | 0.631916 | down |
| 8148 | TAF15 | 2.5E-05 | 0.465481 | 0.454426 | 0.447463 | 0.45579 | down |
| 26470 | SEZ6L2 | 2.51E-05 | 0.639767 | 0.581616 | 0.585484 | 0.602289 | down |
| 9877 | ZC3H11A | 2.51E-05 | 0.515933 | 0.568282 | 0.576971 | 0.553729 | down |
| 9123 | SLC16A3 | 2.51E-05 | 0.681358 | 0.633973 | 0.655251 | 0.656861 | down |
| 643669 | CCER2 | 2.52E-05 | 0.575523 | 0.602566 | 0.567912 | 0.582 | down |
| 55671 | PPP4R3A | 2.52E-05 | 0.588558 | 0.598439 | 0.605093 | 0.597363 | down |
| 10528 | NOP56 | 2.53E-05 | 0.566509 | 0.520004 | 0.550853 | 0.545789 | down |
| 8140 | SLC7A5 | 2.53E-05 | 0.552381 | 0.53536 | 0.505983 | 0.531242 | down |
| 2022 | ENG | 2.54E-05 | 0.638795 | 0.622925 | 0.590896 | 0.617539 | down |
| 10956 | OS9 | 2.54E-05 | 0.630117 | 0.586209 | 0.610284 | 0.60887 | down |

| | | | | | | | |
|---|---|---|---|---|---|---|---|
| 9578 | CDC42BPE | 2.55E-05 | 0.453956 | 0.488703 | 0.446323 | 0.462994 | down |
| 55740 | ENAH | 2.55E-05 | 0.596272 | 0.57008 | 0.565474 | 0.577276 | down |
| 64110 | MAGEF1 | 2.55E-05 | 0.648285 | 0.631246 | 0.683283 | 0.654271 | down |
| 55362 | TMEM63B | 2.56E-05 | 0.518553 | 0.513458 | 0.535431 | 0.522481 | down |
| 6541 | SLC7A1 | 2.57E-05 | 0.454382 | 0.481864 | 0.425732 | 0.453993 | down |
| 54101 | RIPK4 | 2.57E-05 | 0.54087 | 0.566515 | 0.509672 | 0.539019 | down |
| 9816 | URB2 | 2.58E-05 | 0.488077 | 0.509185 | 0.508189 | 0.501817 | down |
| 5791 | PTPRE | 2.58E-05 | 0.456036 | 0.397058 | 0.366197 | 0.40643 | down |
| 23286 | WWC1 | 2.62E-05 | 0.476529 | 0.55275 | 0.524329 | 0.517869 | down |
| 22880 | MORC2 | 2.63E-05 | 0.450556 | 0.453074 | 0.458013 | 0.453881 | down |
| 22990 | PCNX1 | 2.63E-05 | 0.42018 | 0.463741 | 0.454555 | 0.446159 | down |
| 57466 | SCAF4 | 2.64E-05 | 0.45147 | 0.369502 | 0.401698 | 0.407556 | down |
| 5226 | PGD | 2.64E-05 | 0.636424 | 0.612661 | 0.611876 | 0.620321 | down |
| 57148 | RALGAPB | 2.66E-05 | 0.40466 | 0.451093 | 0.433728 | 0.429827 | down |
| 64210 | MMS19 | 2.67E-05 | 0.631441 | 0.638098 | 0.685524 | 0.651688 | down |
| 55627 | SMPD4 | 2.67E-05 | 0.59271 | 0.573732 | 0.563385 | 0.576609 | down |
| 53905 | DUOX1 | 2.67E-05 | 0.552039 | 0.4687 | 0.514208 | 0.511649 | down |
| 6389 | SDHA | 2.67E-05 | 0.681531 | 0.642483 | 0.645672 | 0.656562 | down |
| 375056 | MIA3 | 2.68E-05 | 0.406894 | 0.42305 | 0.40943 | 0.413125 | down |
| 2752 | GLUL | 2.7E-05 | 0.487136 | 0.545375 | 0.475039 | 0.502517 | down |
| 444 | ASPH | 2.71E-05 | 0.58617 | 0.631417 | 0.646948 | 0.621512 | down |
| 894 | CCND2 | 2.71E-05 | 0.521608 | 0.544308 | 0.546706 | 0.537541 | down |
| 2935 | GSPT1 | 2.72E-05 | 0.610303 | 0.593576 | 0.557776 | 0.587218 | down |
| 23270 | TSPYL4 | 2.72E-05 | 0.518809 | 0.569809 | 0.52153 | 0.536716 | down |
| 7916 | PRRC2A | 2.72E-05 | 0.530187 | 0.472834 | 0.473782 | 0.492268 | down |
| 3156 | HMGCR | 2.72E-05 | 0.621847 | 0.609384 | 0.64734 | 0.62619 | down |
| 55751 | TMEM184C | 2.74E-05 | 0.510978 | 0.542403 | 0.511701 | 0.521694 | down |
| 23344 | ESYT1 | 2.74E-05 | 0.528481 | 0.583134 | 0.542975 | 0.55153 | down |
| 65124 | SOWAHC | 2.75E-05 | 0.515339 | 0.467794 | 0.546349 | 0.509827 | down |
| 1655 | DDX5 | 2.75E-05 | 0.622582 | 0.633781 | 0.663991 | 0.640118 | down |
| 55031 | USP47 | 2.76E-05 | 0.538453 | 0.553247 | 0.605429 | 0.56571 | down |
| 4097 | MAFG | 2.76E-05 | 0.582313 | 0.588709 | 0.634465 | 0.601829 | down |
| 201163 | FLCN | 2.77E-05 | 0.63298 | 0.641136 | 0.576934 | 0.617017 | down |
| 10594 | PRPF8 | 2.77E-05 | 0.395645 | 0.413218 | 0.3795 | 0.396121 | down |
| 91748 | ELMSAN1 | 2.77E-05 | 0.388164 | 0.424817 | 0.472648 | 0.428543 | down |
| 51460 | SFMBT1 | 2.78E-05 | 0.440213 | 0.52525 | 0.461882 | 0.475782 | down |
| 817 | CAMK2D | 2.78E-05 | 0.399812 | 0.404614 | 0.454529 | 0.419652 | down |
| 4221 | MEN1 | 2.78E-05 | 0.593431 | 0.589954 | 0.595745 | 0.593043 | down |
| 5394 | EXOSC10 | 2.78E-05 | 0.536068 | 0.504512 | 0.519971 | 0.520183 | down |
| 161424 | NOP9 | 2.79E-05 | 0.580827 | 0.568195 | 0.568036 | 0.572353 | down |
| 80218 | NAA50 | 2.79E-05 | 0.556168 | 0.578546 | 0.617651 | 0.584122 | down |
| 56254 | RNF20 | 2.8E-05 | 0.530878 | 0.540782 | 0.513917 | 0.528526 | down |
| 23173 | METAP1 | 2.81E-05 | 0.611013 | 0.625014 | 0.668719 | 0.634915 | down |
| 96764 | TGS1 | 2.81E-05 | 0.571377 | 0.553673 | 0.538157 | 0.554403 | down |
| 317 | APAF1 | 2.81E-05 | 0.499687 | 0.569051 | 0.498604 | 0.522448 | down |
| 7068 | THRB | 2.82E-05 | 0.485331 | 0.433549 | 0.460767 | 0.459882 | down |
| 5116 | PCNT | 2.82E-05 | 0.377188 | 0.371185 | 0.375581 | 0.374651 | down |
| 56252 | YLPM1 | 2.83E-05 | 0.434217 | 0.389241 | 0.387037 | 0.403498 | down |
| 8027 | STAM | 2.83E-05 | 0.538107 | 0.54725 | 0.603209 | 0.562856 | down |
| 84919 | PPP1R15B | 2.84E-05 | 0.434175 | 0.459223 | 0.480778 | 0.458059 | down |
| 2335 | FN1 | 2.85E-05 | 0.389072 | 0.33935 | 0.322465 | 0.350296 | down |
| 4070 | TACSTD2 | 2.86E-05 | 0.602068 | 0.624914 | 0.560959 | 0.595981 | down |
| 23609 | MKRN2 | 2.86E-05 | 0.571064 | 0.543606 | 0.519614 | 0.544762 | down |
| 26747 | NUFIP1 | 2.87E-05 | 0.574144 | 0.591151 | 0.63658 | 0.600625 | down |
| 23768 | FLRT2 | 2.87E-05 | 0.462059 | 0.535311 | 0.455989 | 0.484453 | down |
| 2733 | GLE1 | 2.87E-05 | 0.522409 | 0.544895 | 0.600731 | 0.556011 | down |
| 5818 | NECTIN1 | 2.88E-05 | 0.588839 | 0.527736 | 0.527748 | 0.548108 | down |
| 8602 | NOP14 | 2.9E-05 | 0.511309 | 0.545766 | 0.464879 | 0.507318 | down |

| | | | | | | | |
|---|---|---|---|---|---|---|---|
| 3269 | HRH1 | 2.92E-05 | 0.604194 | 0.64592 | 0.646338 | 0.632151 | down |
| 10949 | HNRNPA0 | 2.92E-05 | 0.474682 | 0.462458 | 0.439708 | 0.458949 | down |
| 23774 | BRD1 | 2.92E-05 | 0.442424 | 0.458833 | 0.458663 | 0.453307 | down |
| 23014 | FBXO21 | 2.93E-05 | 0.505787 | 0.516207 | 0.568495 | 0.530163 | down |
| 8826 | IQGAP1 | 2.94E-05 | 0.467732 | 0.51752 | 0.552283 | 0.512512 | down |
| 9675 | TTI1 | 2.95E-05 | 0.489421 | 0.568183 | 0.537863 | 0.531823 | down |
| 1196 | CLK2 | 2.95E-05 | 0.603382 | 0.587852 | 0.545764 | 0.578999 | down |
| 342371 | ATXN1L | 2.95E-05 | 0.376446 | 0.45581 | 0.45185 | 0.428036 | down |
| 219988 | PATL1 | 2.96E-05 | 0.440516 | 0.412042 | 0.466143 | 0.439567 | down |
| 55206 | SBNO1 | 2.98E-05 | 0.424118 | 0.430764 | 0.469697 | 0.441526 | down |
| 79658 | ARHGAP1( | 2.98E-05 | 0.517472 | 0.543529 | 0.519184 | 0.526728 | down |
| 23216 | TBC1D1 | 2.98E-05 | 0.548394 | 0.596109 | 0.592559 | 0.579021 | down |
| 23524 | SRRM2 | 2.98E-05 | 0.381528 | 0.408265 | 0.355109 | 0.381634 | down |
| 7464 | CORO2A | 2.99E-05 | 0.591474 | 0.559337 | 0.599175 | 0.583329 | down |
| 79026 | AHNAK | 2.99E-05 | 0.391989 | 0.4348 | 0.440989 | 0.422593 | down |
| 7461 | CLIP2 | 3E-05 | 0.494166 | 0.505499 | 0.518167 | 0.505944 | down |
| 23165 | NUP205 | 3.01E-05 | 0.404391 | 0.458983 | 0.450382 | 0.437919 | down |
| 64771 | C6orf106 | 3.01E-05 | 0.688832 | 0.663972 | 0.643197 | 0.665334 | down |
| 81848 | SPRY4 | 3.01E-05 | 0.534981 | 0.506016 | 0.494048 | 0.511682 | down |
| 79874 | RABEP2 | 3.01E-05 | 0.647406 | 0.589956 | 0.623282 | 0.620215 | down |
| 8061 | FOSL1 | 3.02E-05 | 0.656273 | 0.618947 | 0.66451 | 0.646577 | down |
| 8874 | ARHGEF7 | 3.02E-05 | 0.484565 | 0.512999 | 0.491951 | 0.496505 | down |
| 5781 | PTPN11 | 3.02E-05 | 0.45673 | 0.493151 | 0.514592 | 0.488158 | down |
| 64207 | IRF2BPL | 3.03E-05 | 0.596978 | 0.534392 | 0.547456 | 0.559609 | down |
| 7410 | VAV2 | 3.04E-05 | 0.52932 | 0.50148 | 0.54678 | 0.52586 | down |
| 10413 | YAP1 | 3.04E-05 | 0.444349 | 0.493003 | 0.477695 | 0.471682 | down |
| 63922 | CHTF18 | 3.05E-05 | 0.547403 | 0.468473 | 0.464696 | 0.493524 | down |
| 92822 | ZNF276 | 3.06E-05 | 0.632332 | 0.648743 | 0.584356 | 0.621811 | down |
| 1495 | CTNNA1 | 3.08E-05 | 0.553019 | 0.485107 | 0.491916 | 0.510014 | down |
| 84875 | PARP10 | 3.08E-05 | 0.616483 | 0.622422 | 0.650079 | 0.629661 | down |
| 1832 | DSP | 3.08E-05 | 0.368144 | 0.328446 | 0.349589 | 0.348726 | down |
| 10384 | BTN3A3 | 3.09E-05 | 0.522481 | 0.507711 | 0.58542 | 0.538537 | down |
| 55902 | ACSS2 | 3.09E-05 | 0.615219 | 0.640537 | 0.584882 | 0.613546 | down |
| 1687 | GSDME | 3.09E-05 | 0.651031 | 0.616951 | 0.652199 | 0.64006 | down |
| 490 | ATP2B1 | 3.1E-05 | 0.574226 | 0.522125 | 0.51406 | 0.536803 | down |
| 5591 | PRKDC | 3.1E-05 | 0.337499 | 0.388627 | 0.367966 | 0.364697 | down |
| 55066 | PDPR | 3.1E-05 | 0.429174 | 0.409697 | 0.435828 | 0.4249 | down |
| 2195 | FAT1 | 3.11E-05 | 0.300319 | 0.319771 | 0.293378 | 0.30449 | down |
| 80830 | APOL6 | 3.11E-05 | 0.547239 | 0.561618 | 0.572541 | 0.560466 | down |
| 5338 | PLD2 | 3.12E-05 | 0.623588 | 0.550535 | 0.573777 | 0.582633 | down |
| 283638 | CEP170B | 3.12E-05 | 0.59603 | 0.526319 | 0.527571 | 0.549973 | down |
| 91782 | CHMP7 | 3.14E-05 | 0.560375 | 0.581318 | 0.594165 | 0.57862 | down |
| 162 | AP1B1 | 3.14E-05 | 0.652343 | 0.592406 | 0.590396 | 0.611715 | down |
| 84823 | LMNB2 | 3.14E-05 | 0.502477 | 0.490072 | 0.477807 | 0.490118 | down |
| 9775 | EIF4A3 | 3.15E-05 | 0.663083 | 0.695088 | 0.639214 | 0.665795 | down |
| 29894 | CPSF1 | 3.16E-05 | 0.601311 | 0.578851 | 0.550938 | 0.577033 | down |
| 8936 | WASF1 | 3.16E-05 | 0.504857 | 0.547057 | 0.456157 | 0.50269 | down |
| 718 | C3 | 3.17E-05 | 0.482383 | 0.507652 | 0.454889 | 0.481641 | down |
| 7037 | TFRC | 3.17E-05 | 0.591611 | 0.625887 | 0.640182 | 0.619227 | down |
| 23348 | DOCK9 | 3.17E-05 | 0.346841 | 0.417238 | 0.394187 | 0.386089 | down |
| 23530 | NNT | 3.18E-05 | 0.512182 | 0.533109 | 0.512331 | 0.519208 | down |
| 27242 | TNFRSF21 | 3.18E-05 | 0.499885 | 0.480059 | 0.519951 | 0.499965 | down |
| 9146 | HGS | 3.19E-05 | 0.638563 | 0.627069 | 0.58604 | 0.617224 | down |
| 3376 | IARS | 3.19E-05 | 0.528918 | 0.536786 | 0.585138 | 0.550281 | down |
| 51232 | CRIM1 | 3.19E-05 | 0.461122 | 0.491631 | 0.513898 | 0.488884 | down |
| 55341 | LSG1 | 3.2E-05 | 0.61055 | 0.61425 | 0.601276 | 0.608692 | down |
| 79697 | RIOX1 | 3.2E-05 | 0.577328 | 0.584528 | 0.609999 | 0.590619 | down |
| 23526 | ARHGAP4! | 3.21E-05 | 0.611114 | 0.646048 | 0.604066 | 0.620409 | down |

| | | | | | | | |
|---|---|---|---|---|---|---|---|
| 112939 | NACC1 | 3.21E-05 | 0.6646 | 0.596272 | 0.603693 | 0.621522 | down |
| 8467 | SMARCA5 | 3.21E-05 | 0.653861 | 0.676686 | 0.616686 | 0.649077 | down |
| 22882 | ZHX2 | 3.21E-05 | 0.503438 | 0.533869 | 0.555897 | 0.531068 | down |
| 4154 | MBNL1 | 3.22E-05 | 0.592918 | 0.627488 | 0.631381 | 0.617262 | down |
| 22794 | CASC3 | 3.22E-05 | 0.63506 | 0.57829 | 0.567203 | 0.593518 | down |
| 23271 | CAMSAP2 | 3.23E-05 | 0.511316 | 0.531269 | 0.559143 | 0.533909 | down |
| 3321 | IGSF3 | 3.24E-05 | 0.424705 | 0.438357 | 0.479253 | 0.447438 | down |
| 9883 | POM121 | 3.24E-05 | 0.528599 | 0.534341 | 0.500616 | 0.521185 | down |
| 5927 | KDM5A | 3.25E-05 | 0.377803 | 0.472253 | 0.414668 | 0.421574 | down |
| 54862 | CC2D1A | 3.26E-05 | 0.598003 | 0.522782 | 0.568319 | 0.563035 | down |
| 558 | AXL | 3.27E-05 | 0.475602 | 0.450535 | 0.450326 | 0.458821 | down |
| 9632 | SEC24C | 3.27E-05 | 0.55808 | 0.552297 | 0.501018 | 0.537132 | down |
| 3304 | HSPA1B | 3.29E-05 | 0.472775 | 0.426927 | 0.407558 | 0.435753 | down |
| 23513 | SCRIB | 3.3E-05 | 0.585046 | 0.592359 | 0.553658 | 0.577021 | down |
| 10801 | 44448 | 3.3E-05 | 0.63213 | 0.587196 | 0.605813 | 0.60838 | down |
| 56895 | AGPAT4 | 3.3E-05 | 0.528439 | 0.561584 | 0.480189 | 0.523404 | down |
| 79068 | FTO | 3.31E-05 | 0.322227 | 0.41548 | 0.377071 | 0.371593 | down |
| 23085 | ERC1 | 3.32E-05 | 0.355451 | 0.394846 | 0.395227 | 0.381841 | down |
| 7917 | BAG6 | 3.33E-05 | 0.566278 | 0.571429 | 0.592266 | 0.576657 | down |
| 5981 | RFC1 | 3.33E-05 | 0.505457 | 0.537816 | 0.56915 | 0.537474 | down |
| 146223 | CMTM4 | 3.33E-05 | 0.499945 | 0.430836 | 0.464572 | 0.465118 | down |
| 55720 | TSR1 | 3.34E-05 | 0.522766 | 0.515436 | 0.49527 | 0.511157 | down |
| 26130 | GAPVD1 | 3.34E-05 | 0.477915 | 0.492506 | 0.458545 | 0.476322 | down |
| 81610 | FAM83D | 3.34E-05 | 0.685198 | 0.657161 | 0.640702 | 0.66102 | down |
| 5054 | SERPINE1 | 3.35E-05 | 0.555578 | 0.546536 | 0.604798 | 0.568971 | down |
| 9793 | CKAP5 | 3.36E-05 | 0.362624 | 0.387903 | 0.42863 | 0.393052 | down |
| 223 | ALDH9A1 | 3.37E-05 | 0.658234 | 0.586948 | 0.630101 | 0.625094 | down |
| 23143 | LRCH1 | 3.37E-05 | 0.412054 | 0.413938 | 0.45962 | 0.428537 | down |
| 29035 | C16orf72 | 3.38E-05 | 0.621344 | 0.588856 | 0.573841 | 0.59468 | down |
| 6714 | SRC | 3.38E-05 | 0.582772 | 0.57404 | 0.522433 | 0.559749 | down |
| 6509 | SLC1A4 | 3.39E-05 | 0.599664 | 0.572633 | 0.562415 | 0.578238 | down |
| 677 | ZFP36L1 | 3.4E-05 | 0.556841 | 0.473562 | 0.513624 | 0.514676 | down |
| 55622 | TTC27 | 3.41E-05 | 0.575909 | 0.625345 | 0.644472 | 0.615242 | down |
| 10444 | ZER1 | 3.41E-05 | 0.630762 | 0.572675 | 0.605288 | 0.602908 | down |
| 2064 | ERBB2 | 3.41E-05 | 0.634161 | 0.614457 | 0.575835 | 0.608151 | down |
| 9921 | RNF10 | 3.41E-05 | 0.553842 | 0.552959 | 0.521022 | 0.542608 | down |
| 57521 | RPTOR | 3.41E-05 | 0.48015 | 0.53669 | 0.509937 | 0.508925 | down |
| 3949 | LDLR | 3.41E-05 | 0.513967 | 0.548541 | 0.483303 | 0.515271 | down |
| 125950 | RAVER1 | 3.42E-05 | 0.62241 | 0.566938 | 0.552654 | 0.580668 | down |
| 146909 | KIF18B | 3.44E-05 | 0.514137 | 0.508723 | 0.55134 | 0.524734 | down |
| 11060 | WWP2 | 3.45E-05 | 0.511443 | 0.482484 | 0.47413 | 0.489352 | down |
| 54487 | DGCR8 | 3.45E-05 | 0.465946 | 0.552891 | 0.483902 | 0.500913 | down |
| 3566 | IL4R | 3.45E-05 | 0.639833 | 0.575861 | 0.619207 | 0.611634 | down |
| 3975 | LHX1 | 3.47E-05 | 0.518865 | 0.441107 | 0.524162 | 0.494711 | down |
| 9416 | DDX23 | 3.51E-05 | 0.574885 | 0.540439 | 0.549942 | 0.555089 | down |
| 22978 | NT5C2 | 3.51E-05 | 0.614116 | 0.59406 | 0.653001 | 0.620392 | down |
| 3954 | LETM1 | 3.51E-05 | 0.460505 | 0.479491 | 0.478254 | 0.47275 | down |
| 2017 | CTTN | 3.52E-05 | 0.446005 | 0.44806 | 0.411464 | 0.435176 | down |
| 50650 | ARHGEF3 | 3.52E-05 | 0.466449 | 0.518309 | 0.530278 | 0.505012 | down |
| 55125 | CEP192 | 3.53E-05 | 0.489863 | 0.459576 | 0.48593 | 0.478456 | down |
| 8939 | FUBP3 | 3.54E-05 | 0.450594 | 0.495689 | 0.509664 | 0.485316 | down |
| 83737 | ITCH | 3.55E-05 | 0.550025 | 0.617371 | 0.585213 | 0.584203 | down |
| 59342 | SCPEP1 | 3.56E-05 | 0.563813 | 0.584345 | 0.638935 | 0.595697 | down |
| 124739 | USP43 | 3.56E-05 | 0.468672 | 0.560336 | 0.50302 | 0.510676 | down |
| 4864 | NPC1 | 3.58E-05 | 0.462904 | 0.46938 | 0.455292 | 0.462526 | down |
| 151636 | DTX3L | 3.59E-05 | 0.438607 | 0.50157 | 0.485107 | 0.475095 | down |
| 9885 | OSBPL2 | 3.61E-05 | 0.478208 | 0.468829 | 0.478566 | 0.475201 | down |
| 7342 | UBP1 | 3.62E-05 | 0.54758 | 0.535522 | 0.498231 | 0.527111 | down |

| | | | | | | | |
|---|---|---|---|---|---|---|---|
| 4854 | NOTCH3 | 3.62E-05 | 0.500362 | 0.435649 | 0.44759 | 0.461201 | down |
| 11260 | XPOT | 3.63E-05 | 0.562655 | 0.633609 | 0.614413 | 0.603559 | down |
| 142 | PARP1 | 3.65E-05 | 0.486421 | 0.509824 | 0.510525 | 0.502257 | down |
| 9682 | KDM4A | 3.65E-05 | 0.417417 | 0.46116 | 0.456874 | 0.44515 | down |
| 8473 | OGT | 3.65E-05 | 0.589628 | 0.56632 | 0.636596 | 0.597515 | down |
| 23154 | NCDN | 3.67E-05 | 0.578285 | 0.592773 | 0.542666 | 0.571242 | down |
| 54617 | INO80 | 3.67E-05 | 0.425518 | 0.483745 | 0.444686 | 0.451317 | down |
| 51111 | KMT5B | 3.67E-05 | 0.578746 | 0.526871 | 0.568799 | 0.558139 | down |
| 10989 | IMMT | 3.67E-05 | 0.632436 | 0.643275 | 0.629187 | 0.634966 | down |
| 27153 | ZNF777 | 3.68E-05 | 0.58867 | 0.571121 | 0.543984 | 0.567925 | down |
| 7249 | TSC2 | 3.7E-05 | 0.529911 | 0.49271 | 0.474736 | 0.499119 | down |
| 23049 | SMG1 | 3.7E-05 | 0.384062 | 0.401173 | 0.393471 | 0.392902 | down |
| 51552 | RAB14 | 3.71E-05 | 0.670985 | 0.644082 | 0.683542 | 0.666203 | down |
| 124454 | EARS2 | 3.73E-05 | 0.595984 | 0.607967 | 0.58191 | 0.595287 | down |
| 10299 | 44261 | 3.73E-05 | 0.565468 | 0.534043 | 0.512778 | 0.53743 | down |
| 55623 | THUMPD1 | 3.75E-05 | 0.576488 | 0.597269 | 0.584113 | 0.585957 | down |
| 55379 | LRRC59 | 3.75E-05 | 0.557703 | 0.512544 | 0.524313 | 0.53152 | down |
| 9500 | MAGED1 | 3.76E-05 | 0.562076 | 0.577055 | 0.495669 | 0.544933 | down |
| 84687 | PPP1R9B | 3.76E-05 | 0.464296 | 0.498707 | 0.475078 | 0.479361 | down |
| 6426 | SRSF1 | 3.77E-05 | 0.61939 | 0.642467 | 0.632629 | 0.631496 | down |
| 9831 | ZNF623 | 3.77E-05 | 0.4952 | 0.452758 | 0.527304 | 0.491754 | down |
| 5879 | RAC1 | 3.78E-05 | 0.627756 | 0.653831 | 0.693864 | 0.658484 | down |
| 2130 | EWSR1 | 3.78E-05 | 0.529324 | 0.489933 | 0.506226 | 0.508494 | down |
| 5469 | MED1 | 3.8E-05 | 0.402406 | 0.442976 | 0.384061 | 0.409814 | down |
| 5725 | PTBP1 | 3.8E-05 | 0.548824 | 0.597349 | 0.560671 | 0.568948 | down |
| 93624 | TADA2B | 3.8E-05 | 0.514901 | 0.506649 | 0.508068 | 0.509873 | down |
| 8603 | FAM193A | 3.81E-05 | 0.420035 | 0.442992 | 0.476911 | 0.446646 | down |
| 84440 | RAB11FIP4 | 3.81E-05 | 0.533549 | 0.491463 | 0.511063 | 0.512025 | down |
| 5780 | PTPN9 | 3.81E-05 | 0.545001 | 0.478975 | 0.538296 | 0.520757 | down |
| 84656 | GLYR1 | 3.82E-05 | 0.572461 | 0.556735 | 0.512635 | 0.547277 | down |
| 54961 | SSH3 | 3.82E-05 | 0.688555 | 0.629989 | 0.624522 | 0.647689 | down |
| 780 | DDR1 | 3.82E-05 | 0.643891 | 0.576858 | 0.580347 | 0.600365 | down |
| 23272 | FAM208A | 3.83E-05 | 0.665807 | 0.652745 | 0.658766 | 0.659106 | down |
| 19 | ABCA1 | 3.83E-05 | 0.625 | 0.627564 | 0.579575 | 0.610713 | down |
| 166647 | ADGRA3 | 3.85E-05 | 0.495152 | 0.522673 | 0.49291 | 0.503578 | down |
| 3716 | JAK1 | 3.86E-05 | 0.425884 | 0.468685 | 0.469741 | 0.45477 | down |
| 23131 | GPATCH8 | 3.86E-05 | 0.423733 | 0.330313 | 0.41496 | 0.389669 | down |
| 7739 | ZNF185 | 3.87E-05 | 0.552454 | 0.483963 | 0.504398 | 0.513605 | down |
| 65095 | KRI1 | 3.87E-05 | 0.466722 | 0.447822 | 0.413855 | 0.4428 | down |
| 80230 | RUFY1 | 3.87E-05 | 0.567939 | 0.621151 | 0.612787 | 0.600625 | down |
| 1938 | EEF2 | 3.88E-05 | 0.544043 | 0.551426 | 0.50108 | 0.532183 | down |
| 56288 | PARD3 | 3.88E-05 | 0.532637 | 0.495019 | 0.530911 | 0.519522 | down |
| 1432 | MAPK14 | 3.9E-05 | 0.408958 | 0.460121 | 0.420882 | 0.429987 | down |
| 9128 | PRPF4 | 3.91E-05 | 0.51668 | 0.505083 | 0.544744 | 0.522169 | down |
| 23382 | AHCYL2 | 3.92E-05 | 0.436698 | 0.370923 | 0.402682 | 0.403434 | down |
| 54386 | TERF2IP | 3.92E-05 | 0.68133 | 0.633739 | 0.656774 | 0.657281 | down |
| 6185 | RPN2 | 3.93E-05 | 0.626979 | 0.646802 | 0.660184 | 0.644655 | down |
| 7009 | TMBIM6 | 3.93E-05 | 0.661206 | 0.635708 | 0.679149 | 0.658687 | down |
| 1277 | COL1A1 | 3.93E-05 | 0.478774 | 0.469988 | 0.411474 | 0.453412 | down |
| 222658 | KCTD20 | 3.94E-05 | 0.662315 | 0.592096 | 0.607238 | 0.620549 | down |
| 7528 | YY1 | 3.94E-05 | 0.463592 | 0.536999 | 0.496651 | 0.499081 | down |
| 9513 | FXR2 | 3.96E-05 | 0.636593 | 0.600926 | 0.589876 | 0.609132 | down |
| 92154 | MTSS1L | 3.96E-05 | 0.604576 | 0.53295 | 0.55989 | 0.565806 | down |
| 51114 | ZDHHC9 | 3.96E-05 | 0.630459 | 0.624436 | 0.645955 | 0.633617 | down |
| 24137 | KIF4A | 3.98E-05 | 0.559964 | 0.597238 | 0.600908 | 0.586037 | down |
| 4774 | NFIA | 4E-05 | 0.458385 | 0.492496 | 0.486668 | 0.479183 | down |
| 55756 | INTS9 | 4E-05 | 0.581985 | 0.587565 | 0.597898 | 0.589149 | down |
| 116985 | ARAP1 | 4.03E-05 | 0.600691 | 0.642533 | 0.594245 | 0.61249 | down |

| | | | | | | | |
|---|---|---|---|---|---|---|---|
| 9462 | RASAL2 | 4.03E-05 | 0.461036 | 0.456643 | 0.445297 | 0.454326 | down |
| 23246 | BOP1 | 4.03E-05 | 0.665572 | 0.677755 | 0.632642 | 0.658656 | down |
| 79966 | SCD5 | 4.04E-05 | 0.542246 | 0.612201 | 0.559948 | 0.571465 | down |
| 1523 | CUX1 | 4.04E-05 | 0.680245 | 0.651366 | 0.663532 | 0.665048 | down |
| 10869 | USP19 | 4.04E-05 | 0.600125 | 0.603189 | 0.531994 | 0.578436 | down |
| 3691 | ITGB4 | 4.06E-05 | 0.560741 | 0.550259 | 0.496939 | 0.53598 | down |
| 10497 | UNC13B | 4.07E-05 | 0.386576 | 0.338148 | 0.39328 | 0.372668 | down |
| 3705 | ITPK1 | 4.07E-05 | 0.650565 | 0.633066 | 0.640064 | 0.641232 | down |
| 55163 | PNPO | 4.07E-05 | 0.560617 | 0.582434 | 0.552756 | 0.565269 | down |
| 8975 | USP13 | 4.07E-05 | 0.54007 | 0.606406 | 0.567562 | 0.571346 | down |
| 51755 | CDK12 | 4.08E-05 | 0.335773 | 0.372566 | 0.400144 | 0.369494 | down |
| 132884 | EVC2 | 4.1E-05 | 0.504762 | 0.490139 | 0.454633 | 0.483178 | down |
| 6732 | SRPK1 | 4.1E-05 | 0.557008 | 0.575095 | 0.579186 | 0.57043 | down |
| 8295 | TRRAP | 4.1E-05 | 0.363 | 0.394031 | 0.390685 | 0.382572 | down |
| 5562 | PRKAA1 | 4.11E-05 | 0.603744 | 0.626172 | 0.663913 | 0.631276 | down |
| 7005 | TEAD3 | 4.11E-05 | 0.552941 | 0.60117 | 0.610452 | 0.588188 | down |
| 2273 | FHL1 | 4.11E-05 | 0.566214 | 0.554293 | 0.623957 | 0.581488 | down |
| 4756 | NEO1 | 4.12E-05 | 0.380297 | 0.365657 | 0.375575 | 0.373843 | down |
| 5496 | PPM1G | 4.13E-05 | 0.546623 | 0.506474 | 0.478159 | 0.510418 | down |
| 1793 | DOCK1 | 4.15E-05 | 0.384737 | 0.407788 | 0.41395 | 0.402158 | down |
| 10614 | HEXIM1 | 4.15E-05 | 0.47734 | 0.497735 | 0.5049 | 0.493325 | down |
| 9659 | PDE4DIP | 4.16E-05 | 0.500193 | 0.480113 | 0.49802 | 0.492775 | down |
| 10270 | AKAP8 | 4.16E-05 | 0.420152 | 0.494701 | 0.459899 | 0.458251 | down |
| 79572 | ATP13A3 | 4.17E-05 | 0.515481 | 0.543958 | 0.563578 | 0.541006 | down |
| 10495 | ENOX2 | 4.17E-05 | 0.622219 | 0.584187 | 0.656875 | 0.621094 | down |
| 9371 | KIF3B | 4.18E-05 | 0.429131 | 0.493768 | 0.439105 | 0.454001 | down |
| 5428 | POLG | 4.24E-05 | 0.471053 | 0.555599 | 0.494713 | 0.507122 | down |
| 9208 | LRRFIP1 | 4.24E-05 | 0.440929 | 0.432509 | 0.421626 | 0.431688 | down |
| 3710 | ITPR3 | 4.25E-05 | 0.482691 | 0.47972 | 0.456854 | 0.473088 | down |
| 64778 | FNDC3B | 4.25E-05 | 0.497238 | 0.496354 | 0.508603 | 0.500732 | down |
| 30846 | EHD2 | 4.26E-05 | 0.522724 | 0.522016 | 0.478717 | 0.507819 | down |
| 6892 | TAPBP | 4.26E-05 | 0.408378 | 0.445898 | 0.396585 | 0.416954 | down |
| 2355 | FOSL2 | 4.27E-05 | 0.462616 | 0.463299 | 0.538078 | 0.487998 | down |
| 29978 | UBQLN2 | 4.27E-05 | 0.5219 | 0.548624 | 0.473228 | 0.514584 | down |
| 57551 | TAOK1 | 4.28E-05 | 0.455195 | 0.483453 | 0.487368 | 0.475338 | down |
| 8539 | API5 | 4.28E-05 | 0.560986 | 0.631363 | 0.630203 | 0.607517 | down |
| 10241 | CALCOCO | 4.3E-05 | 0.621641 | 0.61733 | 0.656719 | 0.631897 | down |
| 253943 | YTHDF3 | 4.3E-05 | 0.566984 | 0.582539 | 0.616635 | 0.588719 | down |
| 9052 | GPRC5A | 4.33E-05 | 0.489269 | 0.518256 | 0.479372 | 0.495632 | down |
| 55742 | PARVA | 4.33E-05 | 0.604754 | 0.529635 | 0.580833 | 0.571741 | down |
| 4141 | MARS | 4.33E-05 | 0.565568 | 0.60511 | 0.590744 | 0.587141 | down |
| 23035 | PHLPP2 | 4.35E-05 | 0.39006 | 0.441082 | 0.37774 | 0.402961 | down |
| 865 | CBFB | 4.36E-05 | 0.466988 | 0.507826 | 0.535417 | 0.50341 | down |
| 2000 | ELF4 | 4.38E-05 | 0.568729 | 0.531389 | 0.519357 | 0.539825 | down |
| 5311 | PKD2 | 4.38E-05 | 0.500508 | 0.44125 | 0.458065 | 0.466608 | down |
| 7415 | VCP | 4.39E-05 | 0.600997 | 0.570657 | 0.532103 | 0.567919 | down |
| 2069 | EREG | 4.39E-05 | 0.531424 | 0.582829 | 0.567226 | 0.560493 | down |
| 10983 | CCNI | 4.4E-05 | 0.53018 | 0.55558 | 0.506234 | 0.530665 | down |
| 23013 | SPEN | 4.4E-05 | 0.325878 | 0.376655 | 0.343172 | 0.348568 | down |
| 7534 | YWHAZ | 4.41E-05 | 0.626585 | 0.656596 | 0.685879 | 0.656353 | down |
| 79977 | GRHL2 | 4.42E-05 | 0.496866 | 0.57715 | 0.508032 | 0.52735 | down |
| 3664 | IRF6 | 4.42E-05 | 0.56623 | 0.5638 | 0.543008 | 0.55768 | down |
| 5305 | PIP4K2A | 4.43E-05 | 0.582312 | 0.594061 | 0.624459 | 0.600277 | down |
| 85415 | RHPN2 | 4.43E-05 | 0.518416 | 0.513042 | 0.592437 | 0.541299 | down |
| 55612 | FERMT1 | 4.44E-05 | 0.482021 | 0.508964 | 0.48808 | 0.493022 | down |
| 57649 | PHF12 | 4.45E-05 | 0.53241 | 0.552551 | 0.579906 | 0.554956 | down |
| 23212 | RRS1 | 4.48E-05 | 0.544275 | 0.54662 | 0.489704 | 0.526866 | down |
| 11176 | BAZ2A | 4.49E-05 | 0.464072 | 0.501124 | 0.450181 | 0.471793 | down |

| | | | | | | | |
|---|---|---|---|---|---|---|---|
| 1488 | CTBP2 | 4.5E-05 | 0.620054 | 0.667451 | 0.614017 | 0.633841 | down |
| 1E+08 | NBPF20 | 4.52E-05 | 0.269254 | 0.304688 | 0.358893 | 0.310945 | down |
| 8816 | DCAF5 | 4.52E-05 | 0.53532 | 0.509531 | 0.581049 | 0.541966 | down |
| 254225 | RNF169 | 4.57E-05 | 0.334397 | 0.353622 | 0.360726 | 0.349582 | down |
| 54623 | PAF1 | 4.57E-05 | 0.572803 | 0.550924 | 0.63208 | 0.585269 | down |
| 7186 | TRAF2 | 4.58E-05 | 0.595571 | 0.564826 | 0.581211 | 0.580536 | down |
| 140710 | SOGA1 | 4.58E-05 | 0.517751 | 0.430175 | 0.516256 | 0.488061 | down |
| 6117 | RPA1 | 4.58E-05 | 0.589449 | 0.632526 | 0.603464 | 0.60848 | down |
| 9780 | PIEZO1 | 4.6E-05 | 0.572227 | 0.557589 | 0.554252 | 0.561356 | down |
| 4067 | LYN | 4.6E-05 | 0.590205 | 0.566417 | 0.619652 | 0.592091 | down |
| 57169 | ZNFX1 | 4.61E-05 | 0.489234 | 0.485704 | 0.457341 | 0.477426 | down |
| 84749 | USP30 | 4.65E-05 | 0.586261 | 0.538151 | 0.577014 | 0.567142 | down |
| 4363 | ABCC1 | 4.65E-05 | 0.522797 | 0.422482 | 0.468181 | 0.471153 | down |
| 5007 | OSBP | 4.65E-05 | 0.415576 | 0.491711 | 0.490628 | 0.465972 | down |
| 29072 | SETD2 | 4.66E-05 | 0.402146 | 0.414278 | 0.43828 | 0.418235 | down |
| 22828 | SCAF8 | 4.71E-05 | 0.458062 | 0.545121 | 0.499422 | 0.500868 | down |
| 8754 | ADAM9 | 4.72E-05 | 0.621879 | 0.662375 | 0.639241 | 0.641165 | down |
| 6793 | STK10 | 4.74E-05 | 0.537298 | 0.519817 | 0.481148 | 0.512754 | down |
| 23089 | PEG10 | 4.74E-05 | 0.448036 | 0.493324 | 0.480996 | 0.474119 | down |
| 3065 | HDAC1 | 4.75E-05 | 0.517862 | 0.583211 | 0.571295 | 0.557456 | down |
| 256364 | EML3 | 4.75E-05 | 0.682801 | 0.6357 | 0.663289 | 0.660597 | down |
| 1288 | COL4A6 | 4.76E-05 | 0.481964 | 0.451225 | 0.457957 | 0.463715 | down |
| 4811 | NID1 | 4.77E-05 | 0.368037 | 0.344019 | 0.360022 | 0.357359 | down |
| 9830 | TRIM14 | 4.79E-05 | 0.526258 | 0.540271 | 0.573171 | 0.546567 | down |
| 23499 | MACF1 | 4.79E-05 | 0.340874 | 0.366483 | 0.364109 | 0.357155 | down |
| 8428 | STK24 | 4.79E-05 | 0.576706 | 0.574061 | 0.576927 | 0.575898 | down |
| 23476 | BRD4 | 4.81E-05 | 0.481743 | 0.463998 | 0.448795 | 0.464845 | down |
| 7532 | YWHAG | 4.81E-05 | 0.508666 | 0.497284 | 0.470869 | 0.492273 | down |
| 3720 | JARID2 | 4.83E-05 | 0.366765 | 0.398319 | 0.468212 | 0.411099 | down |
| 27333 | GOLIM4 | 4.84E-05 | 0.39726 | 0.378342 | 0.440332 | 0.405311 | down |
| 2194 | FASN | 4.84E-05 | 0.484148 | 0.461048 | 0.426197 | 0.457131 | down |
| 23252 | OTUD3 | 4.85E-05 | 0.395286 | 0.369813 | 0.41803 | 0.394376 | down |
| 957 | ENTPD5 | 4.87E-05 | 0.56868 | 0.504215 | 0.52606 | 0.532985 | down |
| 55561 | CDC42BPG | 4.92E-05 | 0.575179 | 0.522392 | 0.547695 | 0.548422 | down |
| 23200 | ATP11B | 4.94E-05 | 0.599734 | 0.645576 | 0.677421 | 0.64091 | down |
| 164091 | PAQR7 | 4.94E-05 | 0.542224 | 0.582468 | 0.621203 | 0.581965 | down |
| 64083 | GOLPH3 | 4.97E-05 | 0.453003 | 0.506848 | 0.466846 | 0.475566 | down |
| 10955 | SERINC3 | 4.97E-05 | 0.450201 | 0.484628 | 0.485388 | 0.473406 | down |
| 389136 | VGLL3 | 4.97E-05 | 0.443233 | 0.500247 | 0.535509 | 0.492996 | down |
| 4628 | MYH10 | 4.99E-05 | 0.346643 | 0.361872 | 0.315732 | 0.341416 | down |
| 3182 | HNRNPAB | 4.99E-05 | 0.658283 | 0.628461 | 0.583486 | 0.62341 | down |
| 10611 | PDLIM5 | 5E-05 | 0.678399 | 0.626221 | 0.653742 | 0.652788 | down |
| 10929 | SRSF8 | 5E-05 | 0.625169 | 0.634306 | 0.566723 | 0.608733 | down |
| 26043 | UBXN7 | 5.01E-05 | 0.475807 | 0.543446 | 0.523519 | 0.514257 | down |
| 254428 | SLC41A1 | 5.02E-05 | 0.463831 | 0.416381 | 0.408407 | 0.429539 | down |
| 91039 | DPP9 | 5.02E-05 | 0.585553 | 0.613837 | 0.576497 | 0.591962 | down |
| 9919 | SEC16A | 5.02E-05 | 0.477858 | 0.481349 | 0.442197 | 0.467135 | down |
| 7343 | UBTF | 5.02E-05 | 0.54902 | 0.490393 | 0.477234 | 0.505549 | down |
| 92521 | SPECC1 | 5.04E-05 | 0.571501 | 0.518472 | 0.599437 | 0.563137 | down |
| 6091 | ROBO1 | 5.04E-05 | 0.39885 | 0.386814 | 0.490054 | 0.425239 | down |
| 23318 | TUT4 | 5.04E-05 | 0.371009 | 0.42981 | 0.427627 | 0.409482 | down |
| 835 | CASP2 | 5.05E-05 | 0.398191 | 0.443087 | 0.376411 | 0.405896 | down |
| 8242 | KDM5C | 5.06E-05 | 0.54072 | 0.529989 | 0.494252 | 0.521654 | down |
| 3383 | ICAM1 | 5.1E-05 | 0.597913 | 0.560033 | 0.561024 | 0.57299 | down |
| 55761 | TTC17 | 5.11E-05 | 0.532915 | 0.600476 | 0.513389 | 0.548927 | down |
| 26135 | SERBP1 | 5.12E-05 | 0.612928 | 0.654442 | 0.666447 | 0.644606 | down |
| 5347 | PLK1 | 5.13E-05 | 0.673037 | 0.661567 | 0.61557 | 0.650058 | down |
| 23760 | PITPNB | 5.13E-05 | 0.647348 | 0.627041 | 0.689017 | 0.654469 | down |

| | | | | | | | |
|---|---|---|---|---|---|---|---|
| 23099 | ZBTB43 | 5.14E-05 | 0.446054 | 0.383685 | 0.474114 | 0.434618 | down |
| 7071 | KLF10 | 5.15E-05 | 0.429104 | 0.474007 | 0.498113 | 0.467074 | down |
| 23167 | EFR3A | 5.15E-05 | 0.592978 | 0.608511 | 0.623431 | 0.608307 | down |
| 84079 | ANKRD27 | 5.16E-05 | 0.541819 | 0.593562 | 0.615798 | 0.583726 | down |
| 57720 | GPR107 | 5.17E-05 | 0.460287 | 0.45466 | 0.454437 | 0.456461 | down |
| 81844 | TRIM56 | 5.19E-05 | 0.376463 | 0.414593 | 0.376863 | 0.389307 | down |
| 23355 | VPS8 | 5.2E-05 | 0.66113 | 0.690509 | 0.614499 | 0.655379 | down |
| 80146 | UXS1 | 5.2E-05 | 0.609511 | 0.661298 | 0.61323 | 0.628013 | down |
| 1E+08 | POM121C | 5.21E-05 | 0.528992 | 0.453192 | 0.441569 | 0.474585 | down |
| 6418 | SET | 5.23E-05 | 0.60838 | 0.63912 | 0.64371 | 0.630404 | down |
| 7407 | VARS | 5.24E-05 | 0.590672 | 0.579884 | 0.546246 | 0.572267 | down |
| 7706 | TRIM25 | 5.26E-05 | 0.496174 | 0.528548 | 0.51777 | 0.514164 | down |
| 3897 | L1CAM | 5.26E-05 | 0.534248 | 0.507402 | 0.455201 | 0.498951 | down |
| 26057 | ANKRD17 | 5.27E-05 | 0.422974 | 0.468635 | 0.433949 | 0.441853 | down |
| 50807 | ASAP1 | 5.27E-05 | 0.441933 | 0.417448 | 0.497617 | 0.452332 | down |
| 79066 | METTL16 | 5.28E-05 | 0.469276 | 0.50386 | 0.570754 | 0.51463 | down |
| 1522 | CTSZ | 5.28E-05 | 0.666965 | 0.603746 | 0.608198 | 0.626303 | down |
| 1785 | DNM2 | 5.28E-05 | 0.611478 | 0.584471 | 0.55942 | 0.585123 | down |
| 6239 | RREB1 | 5.28E-05 | 0.400169 | 0.419795 | 0.434728 | 0.418231 | down |
| 5253 | PHF2 | 5.29E-05 | 0.492036 | 0.451038 | 0.447516 | 0.46353 | down |
| 54842 | MFSD6 | 5.29E-05 | 0.415089 | 0.421532 | 0.470238 | 0.43562 | down |
| 9922 | IQSEC1 | 5.29E-05 | 0.505273 | 0.449881 | 0.453937 | 0.469697 | down |
| 999 | CDH1 | 5.3E-05 | 0.410371 | 0.406575 | 0.405936 | 0.407627 | down |
| 7328 | UBE2H | 5.3E-05 | 0.507996 | 0.530988 | 0.491769 | 0.510251 | down |
| 9810 | RNF40 | 5.31E-05 | 0.500938 | 0.392071 | 0.454962 | 0.449324 | down |
| 25865 | PRKD2 | 5.31E-05 | 0.627638 | 0.588057 | 0.600925 | 0.60554 | down |
| 150465 | TTL | 5.31E-05 | 0.593429 | 0.586618 | 0.649091 | 0.609713 | down |
| 22837 | COBLL1 | 5.32E-05 | 0.505013 | 0.46918 | 0.550072 | 0.508088 | down |
| 11276 | SYNRG | 5.33E-05 | 0.485916 | 0.505715 | 0.516616 | 0.502749 | down |
| 23317 | DNAJC13 | 5.36E-05 | 0.556893 | 0.562512 | 0.59229 | 0.570565 | down |
| 8165 | AKAP1 | 5.36E-05 | 0.571243 | 0.552338 | 0.592189 | 0.571923 | down |
| 23379 | ICE1 | 5.37E-05 | 0.441331 | 0.486502 | 0.518672 | 0.482168 | down |
| 48 | ACO1 | 5.39E-05 | 0.488476 | 0.49157 | 0.533593 | 0.504546 | down |
| 53637 | S1PR5 | 5.39E-05 | 0.612341 | 0.617783 | 0.583671 | 0.604598 | down |
| 27183 | VPS4A | 5.41E-05 | 0.647745 | 0.603749 | 0.613925 | 0.621806 | down |
| 55700 | MAP7D1 | 5.41E-05 | 0.565554 | 0.474568 | 0.528532 | 0.522885 | down |
| 7039 | TGFA | 5.41E-05 | 0.608637 | 0.649021 | 0.654701 | 0.637453 | down |
| 23250 | ATP11A | 5.42E-05 | 0.377423 | 0.385328 | 0.35662 | 0.373124 | down |
| 124637 | CYB5D1 | 5.42E-05 | 0.577194 | 0.539797 | 0.578769 | 0.565253 | down |
| 8846 | ALKBH1 | 5.43E-05 | 0.572468 | 0.53346 | 0.503857 | 0.536595 | down |
| 54820 | NDE1 | 5.45E-05 | 0.549596 | 0.551134 | 0.61215 | 0.57096 | down |
| 823 | CAPN1 | 5.45E-05 | 0.673407 | 0.605776 | 0.606956 | 0.628713 | down |
| 57634 | EP400 | 5.46E-05 | 0.37421 | 0.419413 | 0.391836 | 0.395153 | down |
| 1847 | DUSP5 | 5.48E-05 | 0.544726 | 0.554681 | 0.581579 | 0.560329 | down |
| 51379 | CRLF3 | 5.48E-05 | 0.561793 | 0.609461 | 0.610148 | 0.593801 | down |
| 26155 | NOC2L | 5.49E-05 | 0.601609 | 0.585561 | 0.570159 | 0.585776 | down |
| 151354 | FAM84A | 5.49E-05 | 0.570464 | 0.575071 | 0.521263 | 0.555599 | down |
| 7162 | TPBG | 5.5E-05 | 0.59971 | 0.574751 | 0.535104 | 0.569855 | down |
| 2318 | FLNC | 5.51E-05 | 0.512991 | 0.511846 | 0.477886 | 0.500908 | down |
| 26065 | LSM14A | 5.51E-05 | 0.606444 | 0.677596 | 0.659141 | 0.647727 | down |
| 57724 | EPG5 | 5.51E-05 | 0.403146 | 0.331897 | 0.397107 | 0.377383 | down |
| 57592 | ZNF687 | 5.52E-05 | 0.548046 | 0.588358 | 0.528696 | 0.555033 | down |
| 5792 | PTPRF | 5.52E-05 | 0.481937 | 0.516485 | 0.472996 | 0.490473 | down |
| 5834 | PYGB | 5.53E-05 | 0.637924 | 0.635571 | 0.610843 | 0.628113 | down |
| 63893 | UBE2O | 5.53E-05 | 0.519631 | 0.490403 | 0.442343 | 0.484126 | down |
| 4281 | MID1 | 5.55E-05 | 0.458518 | 0.376229 | 0.479115 | 0.437954 | down |
| 1265 | CNN2 | 5.56E-05 | 0.592809 | 0.5951 | 0.534054 | 0.573988 | down |
| 64145 | RBSN | 5.57E-05 | 0.519577 | 0.535197 | 0.510344 | 0.521706 | down |

| | | | | | | | |
|---|---|---|---|---|---|---|---|
| 112574 | SNX18 | 5.58E-05 | 0.521491 | 0.53663 | 0.569759 | 0.542626 | down |
| 79869 | CPSF7 | 5.58E-05 | 0.607412 | 0.565575 | 0.58043 | 0.584472 | down |
| 2624 | GATA2 | 5.59E-05 | 0.561508 | 0.575946 | 0.511763 | 0.549739 | down |
| 345778 | MTX3 | 5.61E-05 | 0.392487 | 0.457754 | 0.463014 | 0.437752 | down |
| 9887 | SMG7 | 5.63E-05 | 0.444689 | 0.501078 | 0.495296 | 0.480354 | down |
| 8439 | NSMAF | 5.65E-05 | 0.515626 | 0.561896 | 0.542757 | 0.540093 | down |
| 10768 | AHCYL1 | 5.65E-05 | 0.502747 | 0.521541 | 0.542493 | 0.52226 | down |
| 6711 | SPTBN1 | 5.67E-05 | 0.367453 | 0.38151 | 0.366284 | 0.371749 | down |
| 51449 | PCYOX1 | 5.69E-05 | 0.52548 | 0.572289 | 0.569446 | 0.555738 | down |
| 54566 | EPB41L4B | 5.71E-05 | 0.626967 | 0.652116 | 0.624142 | 0.634409 | down |
| 55007 | FAM118A | 5.71E-05 | 0.565696 | 0.629039 | 0.638301 | 0.611012 | down |
| 55810 | FOXJ2 | 5.72E-05 | 0.483857 | 0.461542 | 0.510788 | 0.485395 | down |
| 8396 | PIP4K2B | 5.72E-05 | 0.533843 | 0.56271 | 0.524899 | 0.540484 | down |
| 29966 | STRN3 | 5.73E-05 | 0.59849 | 0.536787 | 0.592886 | 0.576054 | down |
| 23057 | NMNAT2 | 5.74E-05 | 0.48976 | 0.492153 | 0.580048 | 0.520654 | down |
| 23005 | MAPKBP1 | 5.74E-05 | 0.538184 | 0.618244 | 0.565573 | 0.574 | down |
| 11059 | WWP1 | 5.76E-05 | 0.567968 | 0.603756 | 0.639342 | 0.603689 | down |
| 1267 | CNP | 5.78E-05 | 0.518157 | 0.564166 | 0.512263 | 0.531529 | down |
| 4957 | ODF2 | 5.79E-05 | 0.43112 | 0.439278 | 0.432198 | 0.434199 | down |
| 5621 | PRNP | 5.8E-05 | 0.634938 | 0.667571 | 0.683408 | 0.661972 | down |
| 80208 | SPG11 | 5.81E-05 | 0.520806 | 0.544661 | 0.519719 | 0.528395 | down |
| 7286 | TUFT1 | 5.85E-05 | 0.477897 | 0.525454 | 0.490949 | 0.4981 | down |
| 5293 | PIK3CD | 5.85E-05 | 0.423008 | 0.426518 | 0.406992 | 0.418839 | down |
| 26133 | TRPC4AP | 5.86E-05 | 0.602159 | 0.627671 | 0.67475 | 0.63486 | down |
| 23780 | APOL2 | 5.86E-05 | 0.524143 | 0.600955 | 0.568379 | 0.564492 | down |
| 54845 | ESRP1 | 5.86E-05 | 0.538418 | 0.555086 | 0.623683 | 0.572396 | down |
| 23645 | PPP1R15A | 5.88E-05 | 0.637488 | 0.548581 | 0.596211 | 0.594093 | down |
| 11123 | RCAN3 | 5.9E-05 | 0.547985 | 0.597145 | 0.527074 | 0.557401 | down |
| 11183 | MAP4K5 | 5.91E-05 | 0.577001 | 0.569427 | 0.622554 | 0.58966 | down |
| 6840 | SVIL | 5.92E-05 | 0.395285 | 0.376702 | 0.30292 | 0.358302 | down |
| 64784 | CRTC3 | 5.93E-05 | 0.494718 | 0.491641 | 0.456489 | 0.480949 | down |
| 10725 | NFAT5 | 5.93E-05 | 0.386205 | 0.419262 | 0.440971 | 0.415479 | down |
| 81788 | NUAK2 | 5.94E-05 | 0.540974 | 0.461426 | 0.496546 | 0.499649 | down |
| 871 | SERPINH1 | 5.94E-05 | 0.663068 | 0.706607 | 0.629199 | 0.666291 | down |
| 3619 | INCENP | 5.95E-05 | 0.341055 | 0.396853 | 0.376113 | 0.37134 | down |
| 2081 | ERN1 | 5.95E-05 | 0.422428 | 0.463927 | 0.421035 | 0.435797 | down |
| 54434 | SSH1 | 5.95E-05 | 0.463819 | 0.470879 | 0.514225 | 0.482974 | down |
| 6653 | SORL1 | 5.96E-05 | 0.424619 | 0.315155 | 0.324502 | 0.354758 | down |
| 54904 | NSD3 | 5.98E-05 | 0.480881 | 0.514909 | 0.529685 | 0.508491 | down |
| 2005 | ELK4 | 5.98E-05 | 0.434951 | 0.429606 | 0.494793 | 0.453117 | down |
| 2975 | GTF3C1 | 5.98E-05 | 0.460505 | 0.494602 | 0.457918 | 0.471008 | down |
| 11231 | SEC63 | 6E-05 | 0.624636 | 0.602735 | 0.623594 | 0.616988 | down |
| 23130 | ATG2A | 6E-05 | 0.571868 | 0.502107 | 0.505789 | 0.526588 | down |
| 284613 | CYB561D1 | 6E-05 | 0.666041 | 0.647115 | 0.598268 | 0.637141 | down |
| 5604 | MAP2K1 | 6.02E-05 | 0.672875 | 0.653895 | 0.614416 | 0.647062 | down |
| 5493 | PPL | 6.03E-05 | 0.457647 | 0.386328 | 0.418248 | 0.420741 | down |
| 5214 | PFKP | 6.03E-05 | 0.564758 | 0.594738 | 0.53823 | 0.565909 | down |
| 7078 | TIMP3 | 6.04E-05 | 0.531575 | 0.546308 | 0.573664 | 0.550516 | down |
| 54838 | WBP1L | 6.04E-05 | 0.581572 | 0.617209 | 0.652483 | 0.617088 | down |
| 64757 | 44256 | 6.04E-05 | 0.622078 | 0.690839 | 0.651739 | 0.654886 | down |
| 163882 | CNST | 6.06E-05 | 0.590682 | 0.633758 | 0.651546 | 0.625329 | down |
| 23052 | ENDOD1 | 6.08E-05 | 0.439458 | 0.486109 | 0.490101 | 0.47189 | down |
| 80315 | CPEB4 | 6.08E-05 | 0.418364 | 0.418198 | 0.451836 | 0.429466 | down |
| 9031 | BAZ1B | 6.11E-05 | 0.385102 | 0.467037 | 0.458573 | 0.436904 | down |
| 9231 | DLG5 | 6.12E-05 | 0.424271 | 0.434839 | 0.398334 | 0.419148 | down |
| 23332 | CLASP1 | 6.12E-05 | 0.325156 | 0.348919 | 0.42393 | 0.366001 | down |
| 29123 | ANKRD11 | 6.14E-05 | 0.420324 | 0.439858 | 0.413541 | 0.424575 | down |
| 128239 | IQGAP3 | 6.14E-05 | 0.443716 | 0.47079 | 0.391691 | 0.435399 | down |

| | | | | | | | |
|---|---|---|---|---|---|---|---|
| 53944 | CSNK1G1 | 6.14E-05 | 0.535563 | 0.532427 | 0.573582 | 0.547191 | down |
| 10749 | KIF1C | 6.15E-05 | 0.597847 | 0.523972 | 0.513945 | 0.545255 | down |
| 27020 | NPTN | 6.15E-05 | 0.668021 | 0.615679 | 0.639463 | 0.641054 | down |
| 79711 | IPO4 | 6.16E-05 | 0.624284 | 0.626252 | 0.551319 | 0.600618 | down |
| 4790 | NFKB1 | 6.16E-05 | 0.470563 | 0.494479 | 0.553747 | 0.506263 | down |
| 60436 | TGIF2 | 6.17E-05 | 0.484981 | 0.573992 | 0.556789 | 0.538588 | down |
| 147179 | WIPF2 | 6.18E-05 | 0.485732 | 0.495383 | 0.54295 | 0.508022 | down |
| 84168 | ANTXR1 | 6.21E-05 | 0.463316 | 0.440489 | 0.40879 | 0.437532 | down |
| 143384 | CACUL1 | 6.24E-05 | 0.482087 | 0.573063 | 0.547046 | 0.534065 | down |
| 23063 | WAPL | 6.25E-05 | 0.439111 | 0.508379 | 0.503416 | 0.483635 | down |
| 10527 | IPO7 | 6.26E-05 | 0.609751 | 0.660079 | 0.682797 | 0.650876 | down |
| 54778 | RNF111 | 6.26E-05 | 0.527409 | 0.540689 | 0.468578 | 0.512225 | down |
| 6195 | RPS6KA1 | 6.26E-05 | 0.654473 | 0.656434 | 0.677471 | 0.662793 | down |
| 192111 | PGAM5 | 6.28E-05 | 0.6293 | 0.595348 | 0.611862 | 0.61217 | down |
| 2590 | GALNT2 | 6.29E-05 | 0.546681 | 0.485541 | 0.493996 | 0.508739 | down |
| 50809 | HP1BP3 | 6.29E-05 | 0.486045 | 0.5186 | 0.528709 | 0.511118 | down |
| 83999 | KREMEN1 | 6.29E-05 | 0.590164 | 0.629733 | 0.636678 | 0.618859 | down |
| 646 | BNC1 | 6.31E-05 | 0.461526 | 0.503191 | 0.49324 | 0.485985 | down |
| 65979 | PHACTR4 | 6.32E-05 | 0.510408 | 0.568503 | 0.492812 | 0.523908 | down |
| 7074 | TIAM1 | 6.33E-05 | 0.433085 | 0.429639 | 0.395455 | 0.419393 | down |
| 55727 | BTBD7 | 6.36E-05 | 0.489644 | 0.497307 | 0.479752 | 0.488901 | down |
| 84726 | PRRC2B | 6.36E-05 | 0.455157 | 0.455271 | 0.465354 | 0.458594 | down |
| 55705 | IPO9 | 6.37E-05 | 0.487251 | 0.539833 | 0.499028 | 0.508704 | down |
| 57222 | ERGIC1 | 6.39E-05 | 0.598929 | 0.531721 | 0.550562 | 0.560404 | down |
| 4651 | MYO10 | 6.39E-05 | 0.372147 | 0.401921 | 0.35028 | 0.374783 | down |
| 57211 | ADGRG6 | 6.4E-05 | 0.596186 | 0.598223 | 0.605403 | 0.599938 | down |
| 2235 | FECH | 6.4E-05 | 0.5355 | 0.619469 | 0.533925 | 0.562965 | down |
| 23022 | PALLD | 6.4E-05 | 0.540039 | 0.554604 | 0.550186 | 0.548277 | down |
| 4299 | AFF1 | 6.41E-05 | 0.401174 | 0.430923 | 0.437219 | 0.423105 | down |
| 11234 | HPS5 | 6.42E-05 | 0.535649 | 0.575345 | 0.566229 | 0.559074 | down |
| 5422 | POLA1 | 6.43E-05 | 0.413636 | 0.506231 | 0.509189 | 0.476352 | down |
| 9663 | LPIN2 | 6.44E-05 | 0.382455 | 0.476325 | 0.465174 | 0.441318 | down |
| 10615 | SPAG5 | 6.44E-05 | 0.622294 | 0.628651 | 0.642118 | 0.631021 | down |
| 7703 | PCGF2 | 6.45E-05 | 0.542352 | 0.45205 | 0.435866 | 0.476756 | down |
| 127544 | RNF19B | 6.45E-05 | 0.616692 | 0.663065 | 0.588802 | 0.622853 | down |
| 6778 | STAT6 | 6.46E-05 | 0.612374 | 0.545159 | 0.556031 | 0.571188 | down |
| 7874 | USP7 | 6.47E-05 | 0.492329 | 0.542139 | 0.52336 | 0.519276 | down |
| 29855 | UBN1 | 6.47E-05 | 0.43764 | 0.451915 | 0.538963 | 0.476172 | down |
| 84925 | DIRC2 | 6.48E-05 | 0.512063 | 0.477456 | 0.580807 | 0.523442 | down |
| 286128 | ZFP41 | 6.48E-05 | 0.510229 | 0.576046 | 0.561791 | 0.549355 | down |
| 9444 | QKI | 6.51E-05 | 0.541306 | 0.555215 | 0.499251 | 0.531924 | down |
| 22874 | PLEKHA6 | 6.53E-05 | 0.476343 | 0.495252 | 0.453037 | 0.474877 | down |
| 135228 | CD109 | 6.53E-05 | 0.538926 | 0.558368 | 0.6107 | 0.569331 | down |
| 10938 | EHD1 | 6.54E-05 | 0.627534 | 0.619377 | 0.566989 | 0.604634 | down |
| 2534 | FYN | 6.54E-05 | 0.502861 | 0.57751 | 0.554277 | 0.544883 | down |
| 64319 | FBRS | 6.55E-05 | 0.616391 | 0.538635 | 0.620293 | 0.591773 | down |
| 203259 | FAM219A | 6.55E-05 | 0.559906 | 0.604114 | 0.567314 | 0.577111 | down |
| 64132 | XYLT2 | 6.57E-05 | 0.586923 | 0.516686 | 0.491562 | 0.531724 | down |
| 390916 | NUDT19 | 6.59E-05 | 0.501458 | 0.573332 | 0.497997 | 0.524262 | down |
| 79180 | EFHD2 | 6.61E-05 | 0.698854 | 0.649338 | 0.627664 | 0.658619 | down |
| 4919 | ROR1 | 6.61E-05 | 0.401639 | 0.491021 | 0.398438 | 0.430366 | down |
| 55095 | SAMD4B | 6.65E-05 | 0.548836 | 0.56806 | 0.598192 | 0.571696 | down |
| 22897 | CEP164 | 6.65E-05 | 0.563736 | 0.547418 | 0.558129 | 0.556428 | down |
| 6733 | SRPK2 | 6.65E-05 | 0.507681 | 0.519824 | 0.557846 | 0.52845 | down |
| 3064 | HTT | 6.65E-05 | 0.466659 | 0.478657 | 0.437798 | 0.461038 | down |
| 493 | ATP2B4 | 6.66E-05 | 0.384772 | 0.376074 | 0.449294 | 0.40338 | down |
| 55187 | VPS13D | 6.67E-05 | 0.315634 | 0.309087 | 0.349101 | 0.324607 | down |
| 5610 | EIF2AK2 | 6.67E-05 | 0.493121 | 0.531561 | 0.55366 | 0.526114 | down |

| | | | | | | | |
|---|---|---|---|---|---|---|---|
| 390980 | ZNF805 | 6.67E-05 | 0.57704 | 0.509903 | 0.478803 | 0.521916 | down |
| 93487 | MAPK1IP1 | 6.72E-05 | 0.53149 | 0.457358 | 0.561867 | 0.516905 | down |
| 29956 | CERS2 | 6.74E-05 | 0.588072 | 0.623021 | 0.639763 | 0.616952 | down |
| 23093 | TTLL5 | 6.76E-05 | 0.475261 | 0.41504 | 0.493494 | 0.461265 | down |
| 9709 | HERPUD1 | 6.76E-05 | 0.637126 | 0.646128 | 0.630538 | 0.637931 | down |
| 84077 | C3orf20 | 6.77E-05 | 0.365385 | 0.250248 | 0.336336 | 0.317323 | down |
| 54832 | VPS13C | 6.77E-05 | 0.346863 | 0.466579 | 0.441038 | 0.41816 | down |
| 5255 | PHKA1 | 6.83E-05 | 0.53166 | 0.4937 | 0.519108 | 0.514823 | down |
| 109 | ADCY3 | 6.83E-05 | 0.465162 | 0.567614 | 0.487807 | 0.506861 | down |
| 11143 | KAT7 | 6.83E-05 | 0.496037 | 0.445523 | 0.485388 | 0.475649 | down |
| 57506 | MAVS | 6.84E-05 | 0.573028 | 0.639699 | 0.595039 | 0.602589 | down |
| 6667 | SP1 | 6.86E-05 | 0.459402 | 0.538139 | 0.510024 | 0.502522 | down |
| 60 | ACTB | 6.9E-05 | 0.59341 | 0.581677 | 0.530792 | 0.568626 | down |
| 3191 | HNRNPL | 6.9E-05 | 0.683809 | 0.652237 | 0.617858 | 0.651301 | down |
| 79039 | DDX54 | 6.91E-05 | 0.591748 | 0.542875 | 0.529988 | 0.554871 | down |
| 23203 | PMPCA | 6.93E-05 | 0.651418 | 0.602286 | 0.584363 | 0.612689 | down |
| 5211 | PFKL | 6.94E-05 | 0.592583 | 0.622568 | 0.600164 | 0.605105 | down |
| 138050 | HGSNAT | 6.95E-05 | 0.374284 | 0.446339 | 0.419622 | 0.413415 | down |
| 57648 | KIAA1522 | 6.99E-05 | 0.604019 | 0.551973 | 0.528388 | 0.56146 | down |
| 64837 | KLC2 | 7.01E-05 | 0.666566 | 0.58294 | 0.618919 | 0.622808 | down |
| 5049 | PAFAH1B2 | 7.03E-05 | 0.561273 | 0.619057 | 0.629553 | 0.603294 | down |
| 23140 | ZZEF1 | 7.09E-05 | 0.420299 | 0.398127 | 0.394422 | 0.404283 | down |
| 9929 | JOSD1 | 7.09E-05 | 0.553568 | 0.636234 | 0.627674 | 0.605826 | down |
| 254102 | EHBP1L1 | 7.1E-05 | 0.52641 | 0.508953 | 0.493896 | 0.509753 | down |
| 23283 | CSTF2T | 7.1E-05 | 0.603558 | 0.626665 | 0.667351 | 0.632525 | down |
| 8914 | TIMELESS | 7.1E-05 | 0.493161 | 0.542524 | 0.479024 | 0.504903 | down |
| 57534 | MIB1 | 7.12E-05 | 0.502199 | 0.567476 | 0.524798 | 0.531491 | down |
| 1968 | EIF2S3 | 7.13E-05 | 0.473447 | 0.516646 | 0.4917 | 0.493931 | down |
| 8853 | ASAP2 | 7.16E-05 | 0.449552 | 0.515845 | 0.470815 | 0.478737 | down |
| 54920 | DUS2 | 7.17E-05 | 0.633621 | 0.653176 | 0.6102 | 0.632332 | down |
| 2185 | PTK2B | 7.17E-05 | 0.514571 | 0.493463 | 0.432229 | 0.480088 | down |
| 4034 | LRCH4 | 7.2E-05 | 0.670818 | 0.619958 | 0.68402 | 0.658265 | down |
| 157769 | FAM91A1 | 7.2E-05 | 0.536563 | 0.60581 | 0.615929 | 0.586101 | down |
| 9698 | PUM1 | 7.2E-05 | 0.510676 | 0.507956 | 0.550824 | 0.523152 | down |
| 7020 | TFAP2A | 7.22E-05 | 0.572568 | 0.524563 | 0.496525 | 0.531219 | down |
| 10469 | TIMM44 | 7.23E-05 | 0.658736 | 0.607744 | 0.570961 | 0.61248 | down |
| 55703 | POLR3B | 7.23E-05 | 0.48866 | 0.417455 | 0.447174 | 0.451097 | down |
| 55147 | RBM23 | 7.24E-05 | 0.615289 | 0.612598 | 0.536419 | 0.588102 | down |
| 2192 | FBLN1 | 7.25E-05 | 0.653886 | 0.685783 | 0.648925 | 0.662865 | down |
| 4133 | MAP2 | 7.26E-05 | 0.479829 | 0.399793 | 0.487395 | 0.455672 | down |
| 4802 | NFYC | 7.27E-05 | 0.571071 | 0.545103 | 0.611535 | 0.575903 | down |
| 92105 | INTS4 | 7.27E-05 | 0.646993 | 0.701857 | 0.623014 | 0.657288 | down |
| 8907 | AP1M1 | 7.3E-05 | 0.658121 | 0.667913 | 0.632241 | 0.652759 | down |
| 1209 | CLPTM1 | 7.3E-05 | 0.671727 | 0.587909 | 0.594678 | 0.618105 | down |
| 79751 | SLC25A22 | 7.3E-05 | 0.658412 | 0.614235 | 0.573069 | 0.615239 | down |
| 22822 | PHLDA1 | 7.3E-05 | 0.504818 | 0.505396 | 0.502582 | 0.504265 | down |
| 9824 | ARHGAP11 | 7.31E-05 | 0.5633 | 0.553372 | 0.639069 | 0.585247 | down |
| 84172 | POLR1B | 7.33E-05 | 0.547701 | 0.496701 | 0.496253 | 0.513552 | down |
| 6385 | SDC4 | 7.33E-05 | 0.589958 | 0.589634 | 0.666014 | 0.615202 | down |
| 7084 | TK2 | 7.38E-05 | 0.569728 | 0.566077 | 0.551029 | 0.562278 | down |
| 84327 | ZBED3 | 7.41E-05 | 0.290944 | 0.40154 | 0.345746 | 0.346077 | down |
| 220064 | LTO1 | 7.42E-05 | 0.658414 | 0.632068 | 0.687929 | 0.65947 | down |
| 55218 | EXD2 | 7.44E-05 | 0.630675 | 0.644107 | 0.569405 | 0.614729 | down |
| 57533 | TBC1D14 | 7.45E-05 | 0.582762 | 0.590483 | 0.595905 | 0.589717 | down |
| 10155 | TRIM28 | 7.46E-05 | 0.627669 | 0.634337 | 0.599004 | 0.620337 | down |
| 9993 | DGCR2 | 7.47E-05 | 0.549293 | 0.549177 | 0.584189 | 0.560886 | down |
| 254042 | METAP1D | 7.47E-05 | 0.483677 | 0.495694 | 0.576561 | 0.518644 | down |
| 6311 | ATXN2 | 7.48E-05 | 0.460677 | 0.37613 | 0.398627 | 0.411811 | down |

| | | | | | | | |
|---|---|---|---|---|---|---|---|
| 10905 | MAN1A2 | 7.49E-05 | 0.513092 | 0.557844 | 0.485143 | 0.518693 | down |
| 10013 | HDAC6 | 7.5E-05 | 0.508025 | 0.586461 | 0.519494 | 0.537993 | down |
| 399665 | FAM102A | 7.5E-05 | 0.65131 | 0.607767 | 0.658293 | 0.639124 | down |
| 140809 | SRXN1 | 7.51E-05 | 0.582402 | 0.552977 | 0.57706 | 0.570813 | down |
| 3892 | KRT86 | 7.51E-05 | 0.654148 | 0.572687 | 0.644196 | 0.623677 | down |
| 1.01E+08 | LOC10106 | 7.53E-05 | 0.504151 | 0.527047 | 0.534798 | 0.521999 | down |
| 22849 | CPEB3 | 7.53E-05 | 0.525422 | 0.481498 | 0.447564 | 0.484828 | down |
| 6293 | VPS52 | 7.58E-05 | 0.471663 | 0.56984 | 0.514036 | 0.518513 | down |
| 51386 | EIF3L | 7.59E-05 | 0.589163 | 0.622411 | 0.617739 | 0.609771 | down |
| 64854 | USP46 | 7.61E-05 | 0.555597 | 0.547286 | 0.592793 | 0.565225 | down |
| 7546 | ZIC2 | 7.63E-05 | 0.566853 | 0.503515 | 0.462839 | 0.511069 | down |
| 23361 | ZNF629 | 7.64E-05 | 0.559021 | 0.472971 | 0.45995 | 0.497314 | down |
| 64393 | ZMAT3 | 7.68E-05 | 0.589781 | 0.636667 | 0.631955 | 0.619468 | down |
| 23054 | NCOA6 | 7.68E-05 | 0.431503 | 0.472982 | 0.446494 | 0.450326 | down |
| 27252 | KLHL20 | 7.69E-05 | 0.56616 | 0.59572 | 0.639776 | 0.600552 | down |
| 23431 | AP4E1 | 7.69E-05 | 0.384695 | 0.511191 | 0.444003 | 0.44663 | down |
| 221687 | RNF182 | 7.69E-05 | 0.629333 | 0.601421 | 0.572927 | 0.601227 | down |
| 116984 | ARAP2 | 7.7E-05 | 0.494174 | 0.565736 | 0.47983 | 0.513246 | down |
| 284058 | KANSL1 | 7.7E-05 | 0.446503 | 0.483533 | 0.474648 | 0.468228 | down |
| 23523 | CABIN1 | 7.71E-05 | 0.60246 | 0.516407 | 0.546064 | 0.554977 | down |
| 51 | ACOX1 | 7.71E-05 | 0.510997 | 0.489364 | 0.499293 | 0.499885 | down |
| 4784 | NFIX | 7.72E-05 | 0.578785 | 0.514034 | 0.527948 | 0.540256 | down |
| 5291 | PIK3CB | 7.72E-05 | 0.491666 | 0.514166 | 0.500573 | 0.502135 | down |
| 90338 | ZNF160 | 7.73E-05 | 0.485193 | 0.433354 | 0.527365 | 0.481971 | down |
| 7280 | TUBB2A | 7.73E-05 | 0.687431 | 0.609946 | 0.675651 | 0.657676 | down |
| 85455 | DISP2 | 7.74E-05 | 0.608183 | 0.561226 | 0.621883 | 0.597097 | down |
| 26049 | FAM169A | 7.75E-05 | 0.57384 | 0.643525 | 0.589198 | 0.602188 | down |
| 148304 | C1orf74 | 7.76E-05 | 0.484341 | 0.431363 | 0.505781 | 0.473829 | down |
| 2475 | MTOR | 7.77E-05 | 0.406741 | 0.421525 | 0.388626 | 0.405631 | down |
| 60412 | EXOC4 | 7.78E-05 | 0.440714 | 0.493815 | 0.529395 | 0.487974 | down |
| 10726 | NUDC | 7.78E-05 | 0.583949 | 0.620394 | 0.557664 | 0.587336 | down |
| 84260 | TCHP | 7.79E-05 | 0.419488 | 0.436694 | 0.453227 | 0.43647 | down |
| 6259 | RYK | 7.8E-05 | 0.579987 | 0.618573 | 0.55747 | 0.585343 | down |
| 80144 | FRAS1 | 7.8E-05 | 0.371042 | 0.3954 | 0.41518 | 0.393874 | down |
| 6838 | SURF6 | 7.81E-05 | 0.534817 | 0.4754 | 0.468175 | 0.492797 | down |
| 6470 | SHMT1 | 7.81E-05 | 0.634682 | 0.67163 | 0.677543 | 0.661285 | down |
| 984 | CDK11B | 7.82E-05 | 0.61711 | 0.558513 | 0.57862 | 0.584748 | down |
| 4967 | OGDH | 7.83E-05 | 0.544743 | 0.537036 | 0.504729 | 0.528836 | down |
| 5287 | PIK3C2B | 7.83E-05 | 0.356634 | 0.452024 | 0.351296 | 0.386651 | down |
| 22930 | RAB3GAP1 | 7.85E-05 | 0.397526 | 0.472858 | 0.461016 | 0.4438 | down |
| 11180 | WDR6 | 7.87E-05 | 0.586372 | 0.512737 | 0.492312 | 0.530474 | down |
| 23107 | MRPS27 | 7.88E-05 | 0.48598 | 0.476548 | 0.548203 | 0.503577 | down |
| 10318 | TNIP1 | 7.9E-05 | 0.622998 | 0.584477 | 0.587665 | 0.59838 | down |
| 5880 | RAC2 | 7.91E-05 | 0.678242 | 0.632365 | 0.673368 | 0.661325 | down |
| 6651 | SON | 7.95E-05 | 0.621351 | 0.626868 | 0.661156 | 0.636459 | down |
| 9451 | EIF2AK3 | 7.96E-05 | 0.381814 | 0.467923 | 0.497383 | 0.44904 | down |
| 1605 | DAG1 | 7.96E-05 | 0.464961 | 0.494908 | 0.481856 | 0.480575 | down |
| 113 | ADCY7 | 8.05E-05 | 0.478705 | 0.435833 | 0.409745 | 0.441428 | down |
| 1591 | CYP24A1 | 8.05E-05 | 0.594574 | 0.556422 | 0.568734 | 0.573244 | down |
| 9711 | RUBCN | 8.06E-05 | 0.504656 | 0.51837 | 0.478556 | 0.500528 | down |
| 9414 | TJP2 | 8.07E-05 | 0.530634 | 0.54879 | 0.504185 | 0.52787 | down |
| 79903 | NAA60 | 8.07E-05 | 0.685415 | 0.648442 | 0.615032 | 0.64963 | down |
| 375484 | SIMC1 | 8.08E-05 | 0.54664 | 0.558959 | 0.550699 | 0.552099 | down |
| 54545 | MTMR12 | 8.09E-05 | 0.513464 | 0.588579 | 0.581609 | 0.561217 | down |
| 7508 | XPC | 8.11E-05 | 0.437187 | 0.391208 | 0.504268 | 0.444221 | down |
| 56886 | UGGT1 | 8.12E-05 | 0.436044 | 0.485913 | 0.541361 | 0.487773 | down |
| 9739 | SETD1A | 8.15E-05 | 0.583196 | 0.520014 | 0.519109 | 0.540773 | down |
| 146059 | CDAN1 | 8.15E-05 | 0.578155 | 0.616306 | 0.643365 | 0.612609 | down |

| | | | | | | | |
|---|---|---|---|---|---|---|---|
| 23187 | PHLDB1 | 8.2E-05 | 0.528193 | 0.455257 | 0.559452 | 0.514301 | down |
| 7915 | ALDH5A1 | 8.21E-05 | 0.630452 | 0.608596 | 0.60089 | 0.613313 | down |
| 23621 | BACE1 | 8.22E-05 | 0.550693 | 0.560925 | 0.516446 | 0.542688 | down |
| 1284 | COL4A2 | 8.23E-05 | 0.43298 | 0.398773 | 0.509352 | 0.447035 | down |
| 23333 | DPY19L1 | 8.23E-05 | 0.550849 | 0.522369 | 0.612828 | 0.562015 | down |
| 23171 | GPD1L | 8.25E-05 | 0.429725 | 0.477246 | 0.485649 | 0.464207 | down |
| 1407 | CRY1 | 8.26E-05 | 0.536782 | 0.575067 | 0.56 | 0.557283 | down |
| 55755 | CDK5RAP2 | 8.27E-05 | 0.371865 | 0.476769 | 0.447346 | 0.431993 | down |
| 1952 | CELSR2 | 8.27E-05 | 0.486264 | 0.460006 | 0.450061 | 0.465444 | down |
| 81565 | NDEL1 | 8.28E-05 | 0.608049 | 0.521152 | 0.559539 | 0.562913 | down |
| 64761 | PARP12 | 8.28E-05 | 0.632653 | 0.562009 | 0.629124 | 0.607929 | down |
| 63891 | RNF123 | 8.28E-05 | 0.608331 | 0.584593 | 0.563419 | 0.585448 | down |
| 1186 | CLCN7 | 8.3E-05 | 0.645133 | 0.64547 | 0.564671 | 0.618425 | down |
| 8776 | MTMR1 | 8.31E-05 | 0.496152 | 0.492943 | 0.584588 | 0.524561 | down |
| 23389 | MED13L | 8.33E-05 | 0.350179 | 0.392906 | 0.44308 | 0.395388 | down |
| 23481 | PES1 | 8.35E-05 | 0.570905 | 0.53864 | 0.529139 | 0.546228 | down |
| 51586 | MED15 | 8.38E-05 | 0.605854 | 0.576945 | 0.570046 | 0.584282 | down |
| 57661 | PHRF1 | 8.44E-05 | 0.489588 | 0.481997 | 0.480198 | 0.483928 | down |
| 96459 | FNIP1 | 8.45E-05 | 0.51235 | 0.549561 | 0.575079 | 0.545663 | down |
| 23170 | TTLL12 | 8.47E-05 | 0.661506 | 0.667261 | 0.613478 | 0.647415 | down |
| 23248 | RPRD2 | 8.47E-05 | 0.401694 | 0.371399 | 0.432165 | 0.401753 | down |
| 25885 | POLR1A | 8.47E-05 | 0.409433 | 0.474495 | 0.441804 | 0.44191 | down |
| 286262 | TPRN | 8.48E-05 | 0.683916 | 0.649187 | 0.637283 | 0.656795 | down |
| 23446 | SLC44A1 | 8.5E-05 | 0.433314 | 0.471889 | 0.471162 | 0.458788 | down |
| 463 | ZFHX3 | 8.5E-05 | 0.414705 | 0.456763 | 0.398626 | 0.423365 | down |
| 23243 | ANKRD28 | 8.5E-05 | 0.509493 | 0.566831 | 0.482852 | 0.519725 | down |
| 25778 | DSTYK | 8.51E-05 | 0.53112 | 0.547247 | 0.558534 | 0.545634 | down |
| 283373 | ANKRD52 | 8.51E-05 | 0.505367 | 0.515756 | 0.574999 | 0.532041 | down |
| 64397 | ZNF106 | 8.54E-05 | 0.362571 | 0.448164 | 0.440865 | 0.4172 | down |
| 9394 | HS6ST1 | 8.54E-05 | 0.586786 | 0.509666 | 0.505591 | 0.534014 | down |
| 114885 | OSBPL11 | 8.54E-05 | 0.572476 | 0.637491 | 0.601362 | 0.603776 | down |
| 55809 | TRERF1 | 8.55E-05 | 0.358471 | 0.454119 | 0.321934 | 0.378175 | down |
| 1303 | COL12A1 | 8.57E-05 | 0.28 | 0.311333 | 0.400733 | 0.330689 | down |
| 4524 | MTHFR | 8.57E-05 | 0.63631 | 0.668796 | 0.639846 | 0.648317 | down |
| 9368 | SLC9A3R1 | 8.57E-05 | 0.651927 | 0.604613 | 0.573068 | 0.609869 | down |
| 128637 | TBC1D20 | 8.58E-05 | 0.694301 | 0.654713 | 0.627595 | 0.658869 | down |
| 3482 | IGF2R | 8.6E-05 | 0.349986 | 0.402586 | 0.345088 | 0.365887 | down |
| 84678 | KDM2B | 8.61E-05 | 0.480921 | 0.546622 | 0.585401 | 0.537648 | down |
| 4851 | NOTCH1 | 8.62E-05 | 0.470612 | 0.498368 | 0.494318 | 0.487766 | down |
| 5813 | PURA | 8.62E-05 | 0.691188 | 0.620221 | 0.671215 | 0.660875 | down |
| 159090 | FAM122B | 8.62E-05 | 0.594751 | 0.639642 | 0.587496 | 0.607296 | down |
| 26015 | RPAP1 | 8.64E-05 | 0.490088 | 0.506939 | 0.55143 | 0.516152 | down |
| 155435 | RBM33 | 8.65E-05 | 0.446518 | 0.485505 | 0.457474 | 0.463166 | down |
| 1824 | DSC2 | 8.65E-05 | 0.511281 | 0.517666 | 0.604843 | 0.544597 | down |
| 60528 | ELAC2 | 8.65E-05 | 0.609952 | 0.596004 | 0.540958 | 0.582305 | down |
| 4171 | MCM2 | 8.66E-05 | 0.573451 | 0.607559 | 0.589308 | 0.590106 | down |
| 6597 | SMARCA4 | 8.66E-05 | 0.378809 | 0.397284 | 0.348515 | 0.374869 | down |
| 4082 | MARCKS | 8.68E-05 | 0.559319 | 0.583735 | 0.509279 | 0.550778 | down |
| 9851 | KIAA0753 | 8.68E-05 | 0.465496 | 0.521127 | 0.531927 | 0.506183 | down |
| 51665 | ASB1 | 8.7E-05 | 0.663818 | 0.601539 | 0.63428 | 0.633212 | down |
| 57473 | ZNF512B | 8.71E-05 | 0.611742 | 0.590127 | 0.622697 | 0.608189 | down |
| 57602 | USP36 | 8.72E-05 | 0.498006 | 0.480633 | 0.521106 | 0.499915 | down |
| 9988 | DMTF1 | 8.73E-05 | 0.606357 | 0.619942 | 0.654616 | 0.626972 | down |
| 23215 | PRRC2C | 8.74E-05 | 0.406595 | 0.47074 | 0.477503 | 0.451613 | down |
| 5530 | PPP3CA | 8.76E-05 | 0.464926 | 0.522668 | 0.549209 | 0.512268 | down |
| 22877 | MLXIP | 8.77E-05 | 0.564808 | 0.547344 | 0.514794 | 0.542315 | down |
| 7175 | TPR | 8.77E-05 | 0.482956 | 0.46423 | 0.499152 | 0.482112 | down |
| 64857 | PLEKHG2 | 8.78E-05 | 0.572771 | 0.473706 | 0.504354 | 0.516944 | down |

| | | | | | | | |
|---|---|---|---|---|---|---|---|
| 92140 | MTDH | 8.79E-05 | 0.574079 | 0.648428 | 0.621403 | 0.614637 | down |
| 2521 | FUS | 8.81E-05 | 0.479466 | 0.435766 | 0.533017 | 0.48275 | down |
| 9645 | MICAL2 | 8.86E-05 | 0.420068 | 0.443813 | 0.416654 | 0.426845 | down |
| 90580 | TIMM29 | 8.87E-05 | 0.695385 | 0.62587 | 0.612222 | 0.644492 | down |
| 1454 | CSNK1E | 8.87E-05 | 0.6171 | 0.586665 | 0.551876 | 0.585214 | down |
| 196527 | ANO6 | 8.87E-05 | 0.43774 | 0.478586 | 0.484998 | 0.467108 | down |
| 153090 | DAB2IP | 8.89E-05 | 0.56582 | 0.504301 | 0.56284 | 0.54432 | down |
| 6868 | ADAM17 | 8.9E-05 | 0.56832 | 0.577372 | 0.657694 | 0.601129 | down |
| 9792 | SERTAD2 | 8.91E-05 | 0.538675 | 0.557538 | 0.534547 | 0.543587 | down |
| 143458 | LDLRAD3 | 8.91E-05 | 0.513791 | 0.550859 | 0.586698 | 0.550449 | down |
| 624 | BDKRB2 | 8.91E-05 | 0.425782 | 0.385081 | 0.399043 | 0.403302 | down |
| 81619 | TSPAN14 | 8.93E-05 | 0.535323 | 0.584602 | 0.530649 | 0.550192 | down |
| 5954 | RCN1 | 8.93E-05 | 0.683122 | 0.643495 | 0.656797 | 0.661138 | down |
| 23307 | FKBP15 | 8.93E-05 | 0.480363 | 0.446346 | 0.468853 | 0.465187 | down |
| 11216 | AKAP10 | 8.94E-05 | 0.596818 | 0.632822 | 0.662565 | 0.630735 | down |
| 8091 | HMGA2 | 8.97E-05 | 0.559371 | 0.581831 | 0.65375 | 0.598318 | down |
| 3667 | IRS1 | 8.97E-05 | 0.498909 | 0.542891 | 0.538947 | 0.526916 | down |
| 2011 | MARK2 | 9E-05 | 0.542516 | 0.465711 | 0.500114 | 0.502781 | down |
| 55128 | TRIM68 | 9.03E-05 | 0.594329 | 0.679087 | 0.621454 | 0.631623 | down |
| 22985 | ACIN1 | 9.05E-05 | 0.510289 | 0.437734 | 0.465549 | 0.471191 | down |
| 10243 | GPHN | 9.06E-05 | 0.520396 | 0.595507 | 0.565915 | 0.560606 | down |
| 374819 | LRRC37A3 | 9.07E-05 | 0.475765 | 0.513296 | 0.566979 | 0.51868 | down |
| 57458 | TMCC3 | 9.12E-05 | 0.515276 | 0.492331 | 0.577589 | 0.528399 | down |
| 79595 | SAP130 | 9.14E-05 | 0.463539 | 0.478595 | 0.469121 | 0.470418 | down |
| 8518 | ELP1 | 9.15E-05 | 0.608402 | 0.656735 | 0.566299 | 0.610479 | down |
| 80314 | EPC1 | 9.18E-05 | 0.604424 | 0.641767 | 0.667407 | 0.637866 | down |
| 5830 | PEX5 | 9.21E-05 | 0.596137 | 0.542383 | 0.550332 | 0.562951 | down |
| 9747 | TCAF1 | 9.23E-05 | 0.464984 | 0.530039 | 0.445435 | 0.480153 | down |
| 10514 | MYBBP1A | 9.24E-05 | 0.498595 | 0.484344 | 0.438706 | 0.473882 | down |
| 4212 | MEIS2 | 9.27E-05 | 0.614625 | 0.507671 | 0.539627 | 0.553974 | down |
| 64411 | ARAP3 | 9.27E-05 | 0.562379 | 0.499544 | 0.603754 | 0.555226 | down |
| 283489 | CHAMP1 | 9.27E-05 | 0.418401 | 0.393411 | 0.471425 | 0.427746 | down |
| 5318 | PKP2 | 9.28E-05 | 0.458475 | 0.540082 | 0.519563 | 0.50604 | down |
| 57645 | POGK | 9.28E-05 | 0.503327 | 0.548733 | 0.589312 | 0.547124 | down |
| 6801 | STRN | 9.28E-05 | 0.571172 | 0.600976 | 0.645509 | 0.605886 | down |
| 23155 | CLCC1 | 9.33E-05 | 0.526684 | 0.502848 | 0.551667 | 0.527066 | down |
| 79654 | HECTD3 | 9.35E-05 | 0.606518 | 0.531245 | 0.57253 | 0.570098 | down |
| 7048 | TGFBR2 | 9.35E-05 | 0.465751 | 0.558804 | 0.52401 | 0.516188 | down |
| 10395 | DLC1 | 9.37E-05 | 0.431482 | 0.442157 | 0.449203 | 0.440947 | down |
| 55322 | C5orf22 | 9.38E-05 | 0.564835 | 0.597065 | 0.632676 | 0.598192 | down |
| 2261 | FGFR3 | 9.38E-05 | 0.692719 | 0.636778 | 0.653768 | 0.661088 | down |
| 10162 | LPCAT3 | 9.39E-05 | 0.627828 | 0.63773 | 0.656071 | 0.640543 | down |
| 79693 | YRDC | 9.44E-05 | 0.615781 | 0.620783 | 0.596957 | 0.611174 | down |
| 9517 | SPTLC2 | 9.44E-05 | 0.623871 | 0.587511 | 0.537924 | 0.583102 | down |
| 23397 | NCAPH | 9.47E-05 | 0.594924 | 0.640598 | 0.58496 | 0.606827 | down |
| 65260 | COA7 | 9.51E-05 | 0.586923 | 0.667235 | 0.634502 | 0.629553 | down |
| 23358 | USP24 | 9.53E-05 | 0.394611 | 0.387112 | 0.396024 | 0.392582 | down |
| 23008 | KLHDC10 | 9.54E-05 | 0.474395 | 0.491191 | 0.492098 | 0.485895 | down |
| 2673 | GFPT1 | 9.59E-05 | 0.603534 | 0.611573 | 0.673805 | 0.629637 | down |
| 133619 | PRRC1 | 9.59E-05 | 0.522587 | 0.596214 | 0.597853 | 0.572218 | down |
| 23648 | SSBP3 | 9.59E-05 | 0.52504 | 0.596227 | 0.603647 | 0.574972 | down |
| 220972 | 44263 | 9.62E-05 | 0.452206 | 0.530381 | 0.447652 | 0.476746 | down |
| 217 | ALDH2 | 9.63E-05 | 0.531591 | 0.586077 | 0.526066 | 0.547911 | down |
| 89795 | NAV3 | 9.66E-05 | 0.418162 | 0.387764 | 0.395449 | 0.400458 | down |
| 6650 | CAPN15 | 9.7E-05 | 0.690807 | 0.647031 | 0.621353 | 0.653064 | down |
| 29763 | PACSIN3 | 9.77E-05 | 0.552495 | 0.51491 | 0.525699 | 0.531035 | down |
| 90102 | PHLDB2 | 9.78E-05 | 0.556308 | 0.594951 | 0.561822 | 0.571027 | down |
| 26504 | CNNM4 | 9.8E-05 | 0.601006 | 0.48539 | 0.533624 | 0.540007 | down |

| | | | | | | | |
|---|---|---|---|---|---|---|---|
| 9917 | FAM20B | 9.82E-05 | 0.623146 | 0.648344 | 0.610241 | 0.627243 | down |
| 6560 | SLC12A4 | 9.83E-05 | 0.623354 | 0.658637 | 0.564386 | 0.615459 | down |
| 90678 | LRSAM1 | 9.83E-05 | 0.525982 | 0.495576 | 0.588693 | 0.53675 | down |
| 10499 | NCOA2 | 9.85E-05 | 0.423418 | 0.468263 | 0.434766 | 0.442149 | down |
| 9928 | KIF14 | 9.87E-05 | 0.489577 | 0.537782 | 0.457969 | 0.495109 | down |
| 9580 | SOX13 | 9.89E-05 | 0.596584 | 0.487125 | 0.540689 | 0.541466 | down |
| 9631 | NUP155 | 9.93E-05 | 0.608828 | 0.624627 | 0.648056 | 0.62717 | down |
| 7813 | EVI5 | 9.95E-05 | 0.4131 | 0.348729 | 0.449557 | 0.403796 | down |
| 10928 | RALBP1 | 9.98E-05 | 0.516069 | 0.56676 | 0.491769 | 0.524866 | down |
| 27097 | TAF5L | 9.99E-05 | 0.623977 | 0.662218 | 0.583742 | 0.623312 | down |
| 54867 | TMEM214 | 9.99E-05 | 0.575824 | 0.626666 | 0.562352 | 0.588281 | down |
| 51663 | ZFR | 0.0001 | 0.572569 | 0.604029 | 0.66463 | 0.613743 | down |
| 80179 | MYO19 | 0.0001 | 0.632819 | 0.637148 | 0.644409 | 0.638126 | down |
| 23075 | SWAP70 | 0.0001 | 0.418133 | 0.479933 | 0.507625 | 0.468564 | down |
| 9731 | CEP104 | 0.000101 | 0.477535 | 0.496332 | 0.46686 | 0.480242 | down |
| 9491 | PSMF1 | 0.000101 | 0.639677 | 0.604194 | 0.622408 | 0.622093 | down |
| 23363 | OBSL1 | 0.000101 | 0.613815 | 0.565462 | 0.523297 | 0.567525 | down |
| 5717 | PSMD11 | 0.000101 | 0.632788 | 0.682592 | 0.65886 | 0.65808 | down |
| 4775 | NFATC3 | 0.000101 | 0.478087 | 0.493086 | 0.531109 | 0.500761 | down |
| 140775 | SMCR8 | 0.000101 | 0.412332 | 0.461243 | 0.454505 | 0.442693 | down |
| 10099 | TSPAN3 | 0.000101 | 0.572063 | 0.663715 | 0.588985 | 0.608254 | down |
| 57496 | MRTFB | 0.000101 | 0.414405 | 0.462459 | 0.475222 | 0.450695 | down |
| 64434 | NOM1 | 0.000102 | 0.353019 | 0.453338 | 0.465821 | 0.424059 | down |
| 5331 | PLCB3 | 0.000102 | 0.491329 | 0.489457 | 0.443894 | 0.474894 | down |
| 1955 | MEGF9 | 0.000102 | 0.533337 | 0.527036 | 0.456147 | 0.505507 | down |
| 8650 | NUMB | 0.000102 | 0.479575 | 0.517927 | 0.559842 | 0.519115 | down |
| 1.08E+08 | LOC10798 | 0.000102 | 0.562844 | 0.643156 | 0.574139 | 0.59338 | down |
| 7840 | ALMS1 | 0.000102 | 0.464139 | 0.382788 | 0.467472 | 0.438133 | down |
| 2706 | GJB2 | 0.000102 | 0.616149 | 0.619899 | 0.665705 | 0.633917 | down |
| 57727 | NCOA5 | 0.000103 | 0.514666 | 0.617201 | 0.52735 | 0.553072 | down |
| 56262 | LRRC8A | 0.000103 | 0.567449 | 0.566551 | 0.561282 | 0.565094 | down |
| 6599 | SMARCC1 | 0.000103 | 0.430706 | 0.48078 | 0.530433 | 0.48064 | down |
| 7298 | TYMS | 0.000103 | 0.668182 | 0.632683 | 0.650698 | 0.650521 | down |
| 9736 | USP34 | 0.000103 | 0.439219 | 0.48349 | 0.486725 | 0.469811 | down |
| 4214 | MAP3K1 | 0.000103 | 0.472006 | 0.490816 | 0.413853 | 0.458891 | down |
| 1478 | CSTF2 | 0.000103 | 0.670448 | 0.577409 | 0.641539 | 0.629799 | down |
| 23456 | ABCB10 | 0.000104 | 0.596409 | 0.626791 | 0.6707 | 0.6313 | down |
| 8030 | CCDC6 | 0.000104 | 0.530968 | 0.55106 | 0.598339 | 0.560122 | down |
| 7094 | TLN1 | 0.000104 | 0.505707 | 0.453608 | 0.432451 | 0.463922 | down |
| 210 | ALAD | 0.000105 | 0.664683 | 0.635965 | 0.690909 | 0.663852 | down |
| 57221 | ARFGEF3 | 0.000105 | 0.421587 | 0.372014 | 0.399445 | 0.397682 | down |
| 90317 | ZNF616 | 0.000105 | 0.582712 | 0.633048 | 0.586617 | 0.600792 | down |
| 23383 | MAU2 | 0.000105 | 0.539376 | 0.458632 | 0.548376 | 0.515461 | down |
| 3838 | KPNA2 | 0.000105 | 0.585121 | 0.679178 | 0.621816 | 0.628705 | down |
| 1857 | DVL3 | 0.000105 | 0.626934 | 0.542707 | 0.545106 | 0.571582 | down |
| 2314 | FLII | 0.000105 | 0.503908 | 0.552554 | 0.481624 | 0.512695 | down |
| 55806 | HR | 0.000105 | 0.57619 | 0.57685 | 0.513443 | 0.555495 | down |
| 54751 | FBLIM1 | 0.000105 | 0.659731 | 0.60281 | 0.668192 | 0.643577 | down |
| 1387 | CREBBP | 0.000105 | 0.420097 | 0.46545 | 0.377839 | 0.421129 | down |
| 8924 | HERC2 | 0.000106 | 0.399257 | 0.501373 | 0.466949 | 0.455859 | down |
| 64087 | MCCC2 | 0.000106 | 0.545761 | 0.562562 | 0.584508 | 0.564277 | down |
| 271 | AMPD2 | 0.000106 | 0.557549 | 0.583067 | 0.562383 | 0.567666 | down |
| 4905 | NSF | 0.000106 | 0.420662 | 0.437419 | 0.466807 | 0.441629 | down |
| 25909 | AHCTF1 | 0.000106 | 0.551333 | 0.591645 | 0.624581 | 0.589186 | down |
| 5213 | PFKM | 0.000106 | 0.537928 | 0.522731 | 0.483613 | 0.514757 | down |
| 22853 | LMTK2 | 0.000106 | 0.407806 | 0.409194 | 0.427259 | 0.414753 | down |
| 8502 | PKP4 | 0.000106 | 0.53623 | 0.560296 | 0.464874 | 0.520467 | down |
| 4641 | MYO1C | 0.000106 | 0.539277 | 0.547437 | 0.485492 | 0.524069 | down |

| | | | | | | | |
|---|---|---|---|---|---|---|---|
| 636 | BICD1 | 0.000106 | 0.365448 | 0.388406 | 0.404805 | 0.38622 | down |
| 23338 | JADE2 | 0.000107 | 0.599233 | 0.570136 | 0.643153 | 0.604174 | down |
| 4735 | 44441 | 0.000107 | 0.620594 | 0.645119 | 0.688048 | 0.651253 | down |
| 23164 | MPRIP | 0.000107 | 0.478372 | 0.51583 | 0.483595 | 0.492599 | down |
| 3831 | KLC1 | 0.000107 | 0.516396 | 0.50936 | 0.571761 | 0.532506 | down |
| 55114 | ARHGAP17 | 0.000107 | 0.479638 | 0.430077 | 0.525743 | 0.478486 | down |
| 55361 | PI4K2A | 0.000108 | 0.571913 | 0.632093 | 0.532532 | 0.578846 | down |
| 3836 | KPNA1 | 0.000108 | 0.553248 | 0.578502 | 0.564159 | 0.565303 | down |
| 8315 | BRAP | 0.000108 | 0.492379 | 0.474549 | 0.524041 | 0.49699 | down |
| 4077 | NBR1 | 0.000108 | 0.472608 | 0.494305 | 0.567582 | 0.511498 | down |
| 64759 | TNS3 | 0.000108 | 0.389739 | 0.426347 | 0.371343 | 0.395809 | down |
| 9695 | EDEM1 | 0.000108 | 0.384654 | 0.458607 | 0.430243 | 0.424501 | down |
| 1030 | CDKN2B | 0.000108 | 0.449605 | 0.40557 | 0.482814 | 0.445996 | down |
| 7922 | SLC39A7 | 0.000108 | 0.511133 | 0.480144 | 0.558594 | 0.516623 | down |
| 284129 | SLC26A11 | 0.000108 | 0.624635 | 0.6177 | 0.583572 | 0.608636 | down |
| 81539 | SLC38A1 | 0.000109 | 0.46539 | 0.556793 | 0.506348 | 0.50951 | down |
| 66008 | TRAK2 | 0.000109 | 0.49726 | 0.528672 | 0.593503 | 0.539811 | down |
| 10847 | SRCAP | 0.000109 | 0.484062 | 0.457731 | 0.453162 | 0.464985 | down |
| 8925 | HERC1 | 0.000109 | 0.360563 | 0.40572 | 0.314304 | 0.360196 | down |
| 6809 | STX3 | 0.000109 | 0.512994 | 0.500298 | 0.526424 | 0.513239 | down |
| 55729 | ATF7IP | 0.000109 | 0.356566 | 0.392945 | 0.372999 | 0.37417 | down |
| 5566 | PRKACA | 0.000109 | 0.560685 | 0.446108 | 0.493817 | 0.500204 | down |
| 23303 | KIF13B | 0.000109 | 0.360863 | 0.405347 | 0.316829 | 0.361013 | down |
| 146705 | TEPSIN | 0.00011 | 0.674193 | 0.635777 | 0.652477 | 0.654149 | down |
| 146057 | TTBK2 | 0.00011 | 0.496296 | 0.546932 | 0.486578 | 0.509936 | down |
| 152503 | SH3D19 | 0.00011 | 0.500857 | 0.552843 | 0.502687 | 0.518796 | down |
| 91523 | PCED1B | 0.00011 | 0.522155 | 0.459742 | 0.529705 | 0.503868 | down |
| 81706 | PPP1R14C | 0.00011 | 0.649485 | 0.585996 | 0.635003 | 0.623494 | down |
| 4836 | NMT1 | 0.000111 | 0.624303 | 0.680651 | 0.68082 | 0.661925 | down |
| 85450 | ITPRIP | 0.000111 | 0.508953 | 0.525232 | 0.561452 | 0.531879 | down |
| 9862 | MED24 | 0.000111 | 0.650623 | 0.583862 | 0.583474 | 0.605986 | down |
| 51599 | LSR | 0.000111 | 0.643489 | 0.625354 | 0.578422 | 0.615755 | down |
| 57646 | USP28 | 0.000111 | 0.433879 | 0.430629 | 0.420122 | 0.42821 | down |
| 8897 | MTMR3 | 0.000111 | 0.502412 | 0.397497 | 0.490605 | 0.463504 | down |
| 113146 | AHNAK2 | 0.000112 | 0.39603 | 0.403874 | 0.376095 | 0.392 | down |
| 10644 | IGF2BP2 | 0.000112 | 0.497217 | 0.538012 | 0.485051 | 0.50676 | down |
| 55193 | PBRM1 | 0.000112 | 0.366074 | 0.336105 | 0.364043 | 0.355407 | down |
| 114991 | ZNF618 | 0.000112 | 0.472518 | 0.536 | 0.479443 | 0.495987 | down |
| 22980 | TCF25 | 0.000112 | 0.690312 | 0.631443 | 0.652959 | 0.658238 | down |
| 5351 | PLOD1 | 0.000113 | 0.668621 | 0.592429 | 0.572705 | 0.611252 | down |
| 8567 | MADD | 0.000113 | 0.459501 | 0.56105 | 0.49556 | 0.505371 | down |
| 22848 | AAK1 | 0.000113 | 0.539568 | 0.524297 | 0.536905 | 0.53359 | down |
| 23070 | CMTR1 | 0.000114 | 0.556892 | 0.623488 | 0.592481 | 0.590954 | down |
| 60485 | SAV1 | 0.000114 | 0.663152 | 0.696885 | 0.639843 | 0.666626 | down |
| 114112 | TXNRD3 | 0.000114 | 0.698124 | 0.621381 | 0.661189 | 0.660231 | down |
| 79968 | WDR76 | 0.000114 | 0.625987 | 0.6376 | 0.705027 | 0.656204 | down |
| 221656 | KDM1B | 0.000114 | 0.525877 | 0.520561 | 0.566792 | 0.537743 | down |
| 1399 | CRKL | 0.000114 | 0.435575 | 0.504756 | 0.432504 | 0.457612 | down |
| 1994 | ELAVL1 | 0.000114 | 0.666362 | 0.665991 | 0.644783 | 0.659045 | down |
| 112936 | VPS26B | 0.000115 | 0.665113 | 0.563464 | 0.605348 | 0.611308 | down |
| 25819 | NOCT | 0.000115 | 0.584656 | 0.547147 | 0.586837 | 0.57288 | down |
| 57584 | ARHGAP21 | 0.000115 | 0.371 | 0.42804 | 0.344819 | 0.381286 | down |
| 9382 | COG1 | 0.000115 | 0.56231 | 0.488542 | 0.46627 | 0.505707 | down |
| 9776 | ATG13 | 0.000115 | 0.654384 | 0.664799 | 0.622654 | 0.647279 | down |
| 4035 | LRP1 | 0.000115 | 0.535276 | 0.430693 | 0.411928 | 0.459299 | down |
| 23065 | EMC1 | 0.000116 | 0.50068 | 0.467723 | 0.446553 | 0.471652 | down |
| 23220 | DTX4 | 0.000116 | 0.621112 | 0.589295 | 0.608684 | 0.606364 | down |
| 94005 | PIGS | 0.000116 | 0.631733 | 0.597628 | 0.623288 | 0.61755 | down |

| | | | | | | | |
|---|---|---|---|---|---|---|---|
| 10454 | TAB1 | 0.000116 | 0.617944 | 0.628434 | 0.674108 | 0.640162 | down |
| 22976 | PAXIP1 | 0.000117 | 0.485534 | 0.514238 | 0.552429 | 0.5174 | down |
| 26523 | AGO1 | 0.000117 | 0.404236 | 0.430859 | 0.443001 | 0.426032 | down |
| 4924 | NUCB1 | 0.000117 | 0.655249 | 0.560587 | 0.612341 | 0.609392 | down |
| 3660 | IRF2 | 0.000117 | 0.597329 | 0.586965 | 0.561941 | 0.582078 | down |
| 26064 | RAI14 | 0.000117 | 0.558147 | 0.613277 | 0.590592 | 0.587339 | down |
| 10564 | ARFGEF2 | 0.000117 | 0.430142 | 0.448234 | 0.421849 | 0.433408 | down |
| 9611 | NCOR1 | 0.000117 | 0.395252 | 0.489495 | 0.486587 | 0.457111 | down |
| 2034 | EPAS1 | 0.000117 | 0.423882 | 0.459973 | 0.449955 | 0.444603 | down |
| 5611 | DNAJC3 | 0.000118 | 0.504938 | 0.598951 | 0.579689 | 0.561193 | down |
| 2305 | FOXM1 | 0.000118 | 0.627619 | 0.608956 | 0.70192 | 0.646165 | down |
| 7559 | ZNF12 | 0.000118 | 0.559037 | 0.523083 | 0.611401 | 0.564507 | down |
| 2909 | ARHGAP3! | 0.000118 | 0.378908 | 0.395587 | 0.470475 | 0.41499 | down |
| 4507 | MTAP | 0.000119 | 0.626838 | 0.68275 | 0.675042 | 0.661544 | down |
| 4216 | MAP3K4 | 0.000119 | 0.413674 | 0.373885 | 0.470225 | 0.419261 | down |
| 8797 | TNFRSF10/ | 0.00012 | 0.563368 | 0.644027 | 0.570161 | 0.592519 | down |
| 79443 | FYCO1 | 0.00012 | 0.358692 | 0.411456 | 0.404513 | 0.391554 | down |
| 1659 | DHX8 | 0.00012 | 0.446264 | 0.529043 | 0.435304 | 0.470204 | down |
| 6609 | SMPD1 | 0.00012 | 0.660096 | 0.65421 | 0.640835 | 0.651714 | down |
| 2990 | GUSB | 0.00012 | 0.605088 | 0.678667 | 0.643371 | 0.642375 | down |
| 124565 | SLC38A10 | 0.00012 | 0.550516 | 0.44533 | 0.543109 | 0.512985 | down |
| 55898 | UNC45A | 0.00012 | 0.578894 | 0.552367 | 0.589537 | 0.573599 | down |
| 54737 | MPHOSPH | 0.000121 | 0.601645 | 0.628465 | 0.655566 | 0.628559 | down |
| 1743 | DLST | 0.000121 | 0.503987 | 0.602127 | 0.565631 | 0.557248 | down |
| 147040 | KCTD11 | 0.000121 | 0.636967 | 0.701653 | 0.612485 | 0.650369 | down |
| 83858 | ATAD3B | 0.000121 | 0.565979 | 0.646957 | 0.633345 | 0.615427 | down |
| 85462 | FHDC1 | 0.000121 | 0.436053 | 0.51005 | 0.515315 | 0.487139 | down |
| 9138 | ARHGEF1 | 0.000121 | 0.56751 | 0.57607 | 0.512514 | 0.552031 | down |
| 64795 | RMND5A | 0.000121 | 0.520899 | 0.591454 | 0.492343 | 0.534899 | down |
| 23186 | RCOR1 | 0.000121 | 0.466224 | 0.529052 | 0.545647 | 0.513641 | down |
| 7764 | ZNF217 | 0.000122 | 0.351128 | 0.445132 | 0.458609 | 0.41829 | down |
| 79691 | QTRT2 | 0.000122 | 0.412759 | 0.435049 | 0.480537 | 0.442782 | down |
| 221937 | FOXK1 | 0.000122 | 0.51342 | 0.544038 | 0.470007 | 0.509155 | down |
| 23151 | GRAMD4 | 0.000123 | 0.441692 | 0.479998 | 0.527892 | 0.483194 | down |
| 6453 | ITSN1 | 0.000123 | 0.52465 | 0.435677 | 0.532465 | 0.497597 | down |
| 54790 | TET2 | 0.000123 | 0.467941 | 0.431229 | 0.506867 | 0.468679 | down |
| 81846 | SBF2 | 0.000123 | 0.492653 | 0.503936 | 0.521625 | 0.506071 | down |
| 23133 | PHF8 | 0.000123 | 0.423121 | 0.464798 | 0.393754 | 0.427225 | down |
| 23042 | PDXDC1 | 0.000124 | 0.414726 | 0.519571 | 0.426468 | 0.453588 | down |
| 64764 | CREB3L2 | 0.000124 | 0.378382 | 0.422935 | 0.388175 | 0.396497 | down |
| 5411 | PNN | 0.000124 | 0.582547 | 0.634508 | 0.614914 | 0.610656 | down |
| 55275 | VPS53 | 0.000124 | 0.454015 | 0.502946 | 0.381169 | 0.446043 | down |
| 8863 | PER3 | 0.000124 | 0.469417 | 0.449582 | 0.451014 | 0.456671 | down |
| 30011 | SH3KBP1 | 0.000124 | 0.648683 | 0.690081 | 0.65836 | 0.665708 | down |
| 126133 | ALDH16A1 | 0.000125 | 0.575814 | 0.520433 | 0.588258 | 0.561502 | down |
| 25782 | RAB3GAP2 | 0.000125 | 0.460876 | 0.578868 | 0.528152 | 0.522632 | down |
| 148867 | SLC30A7 | 0.000125 | 0.369898 | 0.427846 | 0.423274 | 0.407006 | down |
| 144165 | PRICKLE1 | 0.000125 | 0.54411 | 0.527088 | 0.457143 | 0.509447 | down |
| 114823 | LENG8 | 0.000125 | 0.493066 | 0.469127 | 0.521273 | 0.494488 | down |
| 5019 | OXCT1 | 0.000125 | 0.572415 | 0.597342 | 0.619886 | 0.596548 | down |
| 4850 | CNOT4 | 0.000125 | 0.392867 | 0.464206 | 0.41826 | 0.425111 | down |
| 583 | BBS2 | 0.000126 | 0.531541 | 0.482098 | 0.46076 | 0.491467 | down |
| 9592 | IER2 | 0.000126 | 0.502366 | 0.502769 | 0.512524 | 0.505886 | down |
| 1182 | CLCN3 | 0.000126 | 0.534747 | 0.509883 | 0.528983 | 0.524537 | down |
| 8408 | ULK1 | 0.000126 | 0.596864 | 0.527368 | 0.522344 | 0.548859 | down |
| 6844 | VAMP2 | 0.000126 | 0.705248 | 0.60958 | 0.642326 | 0.652385 | down |
| 84343 | HPS3 | 0.000126 | 0.597519 | 0.670243 | 0.577978 | 0.615246 | down |
| 79789 | CLMN | 0.000127 | 0.605499 | 0.586065 | 0.598127 | 0.596563 | down |

| | | | | | | | |
|---|---|---|---|---|---|---|---|
| 10081 | PDCD7 | 0.000127 | 0.638589 | 0.59891 | 0.554468 | 0.597322 | down |
| 57693 | ZNF317 | 0.000127 | 0.406018 | 0.466786 | 0.407428 | 0.426744 | down |
| 3340 | NDST1 | 0.000127 | 0.555262 | 0.540376 | 0.580461 | 0.5587 | down |
| 79710 | MORC4 | 0.000127 | 0.444504 | 0.526036 | 0.561668 | 0.510736 | down |
| 92345 | NAF1 | 0.000127 | 0.585959 | 0.623764 | 0.660055 | 0.623259 | down |
| 10140 | TOB1 | 0.000128 | 0.606533 | 0.541691 | 0.552258 | 0.566827 | down |
| 253725 | WASHC2C | 0.000128 | 0.538211 | 0.509719 | 0.464387 | 0.504106 | down |
| 285704 | RGMB | 0.000128 | 0.539163 | 0.569983 | 0.60342 | 0.570855 | down |
| 57205 | ATP10D | 0.000129 | 0.481227 | 0.485085 | 0.501806 | 0.489373 | down |
| 23122 | CLASP2 | 0.00013 | 0.507758 | 0.61894 | 0.574021 | 0.566906 | down |
| 582 | BBS1 | 0.00013 | 0.548045 | 0.509004 | 0.529118 | 0.528722 | down |
| 5295 | PIK3R1 | 0.00013 | 0.478793 | 0.496076 | 0.529698 | 0.501522 | down |
| 323 | APBB2 | 0.00013 | 0.469821 | 0.489837 | 0.446993 | 0.468884 | down |
| 1.01E+08 | NBPF19 | 0.00013 | 0.408583 | 0.326191 | 0.321088 | 0.351954 | down |
| 286827 | TRIM59 | 0.00013 | 0.654725 | 0.604855 | 0.595486 | 0.618355 | down |
| 54407 | SLC38A2 | 0.00013 | 0.618987 | 0.681728 | 0.666279 | 0.655665 | down |
| 123283 | TARSL2 | 0.000131 | 0.583774 | 0.587458 | 0.604489 | 0.591907 | down |
| 6942 | TCF20 | 0.000131 | 0.38814 | 0.432455 | 0.49277 | 0.437788 | down |
| 90381 | TICRR | 0.000131 | 0.457465 | 0.478859 | 0.498825 | 0.478383 | down |
| 79990 | PLEKHH3 | 0.000131 | 0.624108 | 0.597513 | 0.576696 | 0.599439 | down |
| 4668 | NAGA | 0.000131 | 0.686625 | 0.603284 | 0.593527 | 0.627812 | down |
| 26298 | EHF | 0.000131 | 0.400474 | 0.470187 | 0.517194 | 0.462619 | down |
| 9489 | PGS1 | 0.000131 | 0.578289 | 0.640205 | 0.642958 | 0.620484 | down |
| 129293 | TRABD2A | 0.000131 | 0.674465 | 0.657204 | 0.582465 | 0.638045 | down |
| 1736 | DKC1 | 0.000131 | 0.594173 | 0.628173 | 0.551315 | 0.59122 | down |
| 2035 | EPB41 | 0.000131 | 0.516332 | 0.595429 | 0.581265 | 0.564342 | down |
| 9871 | SEC24D | 0.000131 | 0.545546 | 0.652129 | 0.586427 | 0.594701 | down |
| 54919 | DNAAF5 | 0.000131 | 0.603435 | 0.59981 | 0.533614 | 0.578953 | down |
| 57217 | TTC7A | 0.000131 | 0.619872 | 0.585183 | 0.64943 | 0.618162 | down |
| 85021 | REPS1 | 0.000132 | 0.521685 | 0.585344 | 0.576288 | 0.561106 | down |
| 9529 | BAG5 | 0.000132 | 0.530121 | 0.5451 | 0.564492 | 0.546571 | down |
| 22872 | SEC31A | 0.000132 | 0.541196 | 0.524939 | 0.557134 | 0.54109 | down |
| 57553 | MICAL3 | 0.000132 | 0.535925 | 0.444357 | 0.529161 | 0.503148 | down |
| 7056 | THBD | 0.000132 | 0.451179 | 0.525218 | 0.492028 | 0.489475 | down |
| 8237 | USP11 | 0.000132 | 0.623061 | 0.60905 | 0.615127 | 0.615746 | down |
| 4646 | MYO6 | 0.000133 | 0.482849 | 0.51641 | 0.595562 | 0.531607 | down |
| 80728 | ARHGAP3( | 0.000133 | 0.570504 | 0.519819 | 0.583472 | 0.557932 | down |
| 4678 | NASP | 0.000133 | 0.589738 | 0.519828 | 0.537302 | 0.548956 | down |
| 23197 | FAF2 | 0.000133 | 0.500298 | 0.562527 | 0.513705 | 0.52551 | down |
| 27229 | TUBGCP4 | 0.000134 | 0.516753 | 0.541835 | 0.536421 | 0.531669 | down |
| 79877 | DCAKD | 0.000134 | 0.5313 | 0.626491 | 0.621035 | 0.592942 | down |
| 3976 | LIF | 0.000134 | 0.548449 | 0.508547 | 0.572483 | 0.54316 | down |
| 5198 | PFAS | 0.000135 | 0.430284 | 0.473352 | 0.360745 | 0.42146 | down |
| 22884 | WDR37 | 0.000135 | 0.495518 | 0.537977 | 0.575209 | 0.536234 | down |
| 200734 | SPRED2 | 0.000135 | 0.604134 | 0.510144 | 0.591523 | 0.5686 | down |
| 55568 | GALNT10 | 0.000135 | 0.515144 | 0.51444 | 0.554627 | 0.52807 | down |
| 57326 | PBXIP1 | 0.000136 | 0.562291 | 0.554577 | 0.478268 | 0.531712 | down |
| 54926 | UBE2R2 | 0.000136 | 0.597407 | 0.560703 | 0.545659 | 0.567923 | down |
| 23392 | ECPAS | 0.000136 | 0.494143 | 0.54367 | 0.603759 | 0.54719 | down |
| 9520 | NPEPPS | 0.000136 | 0.622798 | 0.59488 | 0.621556 | 0.613078 | down |
| 4599 | MX1 | 0.000136 | 0.600023 | 0.615458 | 0.591969 | 0.602483 | down |
| 55167 | MSL2 | 0.000136 | 0.424907 | 0.479988 | 0.544162 | 0.483019 | down |
| 93109 | TMEM44 | 0.000136 | 0.596365 | 0.629559 | 0.594865 | 0.60693 | down |
| 56957 | OTUD7B | 0.000136 | 0.4079 | 0.496982 | 0.46236 | 0.455747 | down |
| 1788 | DNMT3A | 0.000136 | 0.426517 | 0.500498 | 0.496288 | 0.474434 | down |
| 2309 | FOXO3 | 0.000136 | 0.433282 | 0.479087 | 0.529642 | 0.48067 | down |
| 4800 | NFYA | 0.000137 | 0.501048 | 0.555639 | 0.548014 | 0.5349 | down |
| 10186 | LHFPL6 | 0.000137 | 0.604876 | 0.647981 | 0.591183 | 0.61468 | down |

| | | | | | | | |
|---|---|---|---|---|---|---|---|
| 23654 | PLXNB2 | 0.000138 | 0.595189 | 0.630473 | 0.56548 | 0.597047 | down |
| 6625 | SNRNP70 | 0.000138 | 0.577852 | 0.468416 | 0.545853 | 0.530707 | down |
| 21 | ABCA3 | 0.000138 | 0.612938 | 0.633862 | 0.56358 | 0.60346 | down |
| 8824 | CES2 | 0.000139 | 0.641829 | 0.649793 | 0.679061 | 0.656894 | down |
| 113235 | SLC46A1 | 0.000139 | 0.64204 | 0.593751 | 0.530437 | 0.588743 | down |
| 2729 | GCLC | 0.000139 | 0.560036 | 0.539586 | 0.514611 | 0.538078 | down |
| 152006 | RNF38 | 0.000139 | 0.473527 | 0.529929 | 0.470641 | 0.491365 | down |
| 5099 | PCDH7 | 0.000139 | 0.436175 | 0.511514 | 0.510575 | 0.486088 | down |
| 84918 | LRP11 | 0.000139 | 0.506677 | 0.561388 | 0.531841 | 0.533302 | down |
| 80017 | DGLUCY | 0.000139 | 0.644137 | 0.595371 | 0.61346 | 0.617656 | down |
| 9203 | ZMYM3 | 0.000139 | 0.603886 | 0.592228 | 0.684363 | 0.626825 | down |
| 23048 | FNBP1 | 0.00014 | 0.529884 | 0.538446 | 0.473584 | 0.513971 | down |
| 57216 | VANGL2 | 0.00014 | 0.401046 | 0.443211 | 0.493762 | 0.446006 | down |
| 1798 | DPAGT1 | 0.00014 | 0.574912 | 0.579933 | 0.559093 | 0.571312 | down |
| 26037 | SIPA1L1 | 0.00014 | 0.404351 | 0.462037 | 0.495269 | 0.453886 | down |
| 65264 | UBE2Z | 0.00014 | 0.673475 | 0.582884 | 0.66786 | 0.641406 | down |
| 54870 | QRICH1 | 0.00014 | 0.600465 | 0.668743 | 0.572755 | 0.613988 | down |
| 6721 | SREBF2 | 0.00014 | 0.622347 | 0.584312 | 0.524372 | 0.577011 | down |
| 7077 | TIMP2 | 0.00014 | 0.5806 | 0.591153 | 0.532534 | 0.568095 | down |
| 51307 | FAM53C | 0.000141 | 0.573932 | 0.580023 | 0.667365 | 0.607107 | down |
| 83595 | SOX7 | 0.000141 | 0.612026 | 0.516375 | 0.497485 | 0.541962 | down |
| 151963 | MB21D2 | 0.000141 | 0.599228 | 0.671813 | 0.575046 | 0.615363 | down |
| 1783 | DYNC1LI2 | 0.000141 | 0.518605 | 0.556586 | 0.54361 | 0.5396 | down |
| 79707 | NOL9 | 0.000141 | 0.492302 | 0.480066 | 0.480055 | 0.484141 | down |
| 694 | BTG1 | 0.000141 | 0.615754 | 0.681462 | 0.662994 | 0.653403 | down |
| 3959 | LGALS3BP | 0.000141 | 0.690916 | 0.610461 | 0.590968 | 0.630782 | down |
| 23163 | GGA3 | 0.000141 | 0.505393 | 0.533867 | 0.620001 | 0.553087 | down |
| 9802 | DAZAP2 | 0.000141 | 0.643257 | 0.572386 | 0.59665 | 0.604098 | down |
| 84433 | CARD11 | 0.000142 | 0.498414 | 0.480755 | 0.440221 | 0.47313 | down |
| 10452 | TOMM40 | 0.000142 | 0.693912 | 0.60713 | 0.637992 | 0.646345 | down |
| 1654 | DDX3X | 0.000142 | 0.527801 | 0.586274 | 0.584402 | 0.566159 | down |
| 84236 | RHBDD1 | 0.000142 | 0.457028 | 0.465427 | 0.575162 | 0.499206 | down |
| 57020 | VPS35L | 0.000142 | 0.50591 | 0.608948 | 0.509624 | 0.541494 | down |
| 26091 | HERC4 | 0.000142 | 0.633792 | 0.668839 | 0.671697 | 0.658109 | down |
| 23094 | SIPA1L3 | 0.000142 | 0.426353 | 0.429502 | 0.443074 | 0.432977 | down |
| 23086 | EXPH5 | 0.000142 | 0.47876 | 0.520137 | 0.472046 | 0.490314 | down |
| 9373 | PLAA | 0.000142 | 0.641071 | 0.676074 | 0.664451 | 0.660532 | down |
| 605 | BCL7A | 0.000142 | 0.546378 | 0.573141 | 0.597518 | 0.572346 | down |
| 23309 | SIN3B | 0.000143 | 0.644866 | 0.651511 | 0.643494 | 0.646624 | down |
| 63916 | ELMO2 | 0.000143 | 0.50512 | 0.507162 | 0.569569 | 0.527284 | down |
| 122830 | NAA30 | 0.000143 | 0.538794 | 0.580046 | 0.560821 | 0.559887 | down |
| 90204 | ZSWIM1 | 0.000143 | 0.501478 | 0.443576 | 0.536615 | 0.49389 | down |
| 22911 | WDR47 | 0.000143 | 0.600406 | 0.62008 | 0.675794 | 0.632093 | down |
| 54438 | GFOD1 | 0.000144 | 0.417813 | 0.46482 | 0.38519 | 0.422608 | down |
| 80829 | ZFP91 | 0.000144 | 0.52928 | 0.562401 | 0.567435 | 0.553039 | down |
| 54726 | OTUD4 | 0.000144 | 0.467445 | 0.55762 | 0.54976 | 0.524942 | down |
| 55069 | TMEM248 | 0.000144 | 0.62335 | 0.654454 | 0.603647 | 0.62715 | down |
| 116064 | LRRC58 | 0.000144 | 0.49657 | 0.54248 | 0.573288 | 0.537446 | down |
| 729830 | FAM160A1 | 0.000144 | 0.437536 | 0.508145 | 0.409173 | 0.451618 | down |
| 23062 | GGA2 | 0.000144 | 0.605647 | 0.682693 | 0.58366 | 0.624 | down |
| 54815 | GATAD2A | 0.000145 | 0.452926 | 0.465754 | 0.401379 | 0.440019 | down |
| 6683 | SPAST | 0.000145 | 0.537159 | 0.529212 | 0.509325 | 0.525232 | down |
| 54583 | EGLN1 | 0.000145 | 0.503157 | 0.577271 | 0.529492 | 0.53664 | down |
| 2101 | ESRRA | 0.000145 | 0.590943 | 0.588914 | 0.646864 | 0.608907 | down |
| 5966 | REL | 0.000145 | 0.431617 | 0.513811 | 0.492189 | 0.479206 | down |
| 24144 | TFIP11 | 0.000145 | 0.655431 | 0.583913 | 0.543451 | 0.594265 | down |
| 140885 | SIRPA | 0.000145 | 0.574817 | 0.557935 | 0.50005 | 0.544267 | down |
| 171017 | ZNF384 | 0.000146 | 0.515766 | 0.530905 | 0.580545 | 0.542405 | down |

| | | | | | | | |
|---|---|---|---|---|---|---|---|
| 80223 | RAB11FIP1 | 0.000146 | 0.459819 | 0.508477 | 0.4852 | 0.484499 | down |
| 26993 | AKAP8L | 0.000146 | 0.559386 | 0.55907 | 0.47005 | 0.529502 | down |
| 29 | ABR | 0.000146 | 0.568576 | 0.582935 | 0.511039 | 0.554183 | down |
| 1E+08 | ARMCX4 | 0.000146 | 0.38257 | 0.466949 | 0.38585 | 0.41179 | down |
| 57462 | MYORG | 0.000146 | 0.533451 | 0.481732 | 0.600188 | 0.538457 | down |
| 9610 | RIN1 | 0.000146 | 0.69663 | 0.625147 | 0.677692 | 0.66649 | down |
| 64426 | SUDS3 | 0.000147 | 0.603625 | 0.691277 | 0.624295 | 0.639733 | down |
| 5297 | PI4KA | 0.000147 | 0.484288 | 0.552392 | 0.502582 | 0.513087 | down |
| 11252 | PACSIN2 | 0.000147 | 0.477116 | 0.526608 | 0.526218 | 0.509981 | down |
| 84720 | PIGO | 0.000147 | 0.618717 | 0.519268 | 0.618508 | 0.585498 | down |
| 9716 | AQR | 0.000148 | 0.616259 | 0.695508 | 0.674643 | 0.662137 | down |
| 64782 | AEN | 0.000148 | 0.59437 | 0.589084 | 0.65417 | 0.612541 | down |
| 25829 | TMEM184I | 0.000149 | 0.529011 | 0.484832 | 0.507258 | 0.507034 | down |
| 57585 | CRAMP1 | 0.000149 | 0.540229 | 0.481169 | 0.493332 | 0.50491 | down |
| 9743 | ARHGAP3: | 0.000149 | 0.324164 | 0.370773 | 0.426169 | 0.373702 | down |
| 120 | ADD3 | 0.00015 | 0.589446 | 0.659624 | 0.655111 | 0.634727 | down |
| 4041 | LRP5 | 0.00015 | 0.621218 | 0.683182 | 0.590765 | 0.631722 | down |
| 9564 | BCAR1 | 0.00015 | 0.6785 | 0.60639 | 0.581448 | 0.622113 | down |
| 571 | BACH1 | 0.000151 | 0.483163 | 0.568989 | 0.592983 | 0.548379 | down |
| 23405 | DICER1 | 0.000151 | 0.393979 | 0.50787 | 0.458824 | 0.453557 | down |
| 84619 | ZGPAT | 0.000151 | 0.541521 | 0.444395 | 0.438972 | 0.474963 | down |
| 11328 | FKBP9 | 0.000151 | 0.482596 | 0.521482 | 0.467746 | 0.490608 | down |
| 2517 | FUCA1 | 0.000151 | 0.615377 | 0.684859 | 0.665376 | 0.655204 | down |
| 10112 | KIF20A | 0.000151 | 0.564309 | 0.59933 | 0.572845 | 0.578828 | down |
| 29979 | UBQLN1 | 0.000151 | 0.539355 | 0.617837 | 0.602866 | 0.586686 | down |
| 10250 | SRRM1 | 0.000152 | 0.42544 | 0.381827 | 0.412046 | 0.406438 | down |
| 23328 | SASH1 | 0.000152 | 0.389774 | 0.412933 | 0.407256 | 0.403321 | down |
| 10554 | AGPAT1 | 0.000152 | 0.636275 | 0.593402 | 0.648428 | 0.626035 | down |
| 1376 | CPT2 | 0.000152 | 0.678053 | 0.571859 | 0.632324 | 0.627412 | down |
| 257364 | SNX33 | 0.000153 | 0.525203 | 0.564616 | 0.638895 | 0.576238 | down |
| 9202 | ZMYM4 | 0.000153 | 0.473795 | 0.56314 | 0.533606 | 0.523513 | down |
| 5339 | PLEC | 0.000153 | 0.59318 | 0.56488 | 0.548136 | 0.568732 | down |
| 9790 | BMS1 | 0.000153 | 0.578064 | 0.534443 | 0.509969 | 0.540825 | down |
| 729438 | CASTOR2 | 0.000153 | 0.56369 | 0.519663 | 0.526739 | 0.536697 | down |
| 54476 | RNF216 | 0.000153 | 0.611127 | 0.640258 | 0.620657 | 0.624014 | down |
| 80727 | TTYH3 | 0.000153 | 0.644934 | 0.562331 | 0.637784 | 0.615016 | down |
| 26505 | CNNM3 | 0.000154 | 0.574614 | 0.537803 | 0.588512 | 0.566976 | down |
| 54989 | ZNF770 | 0.000154 | 0.55524 | 0.609703 | 0.62119 | 0.595378 | down |
| 9696 | CROCC | 0.000154 | 0.395221 | 0.346263 | 0.353012 | 0.364832 | down |
| 1829 | DSG2 | 0.000154 | 0.488194 | 0.55733 | 0.555767 | 0.533764 | down |
| 4001 | LMNB1 | 0.000154 | 0.661881 | 0.653524 | 0.637 | 0.650802 | down |
| 55526 | DHTKD1 | 0.000154 | 0.549131 | 0.637817 | 0.539515 | 0.575488 | down |
| 10766 | TOB2 | 0.000155 | 0.546461 | 0.515516 | 0.579108 | 0.547028 | down |
| 56478 | EIF4ENIF1 | 0.000155 | 0.526798 | 0.608672 | 0.597759 | 0.577743 | down |
| 1E+08 | ZNF316 | 0.000155 | 0.508744 | 0.496999 | 0.495287 | 0.500343 | down |
| 55030 | FBXO34 | 0.000156 | 0.578938 | 0.491021 | 0.558365 | 0.542774 | down |
| 84133 | ZNRF3 | 0.000156 | 0.534652 | 0.505936 | 0.528099 | 0.522896 | down |
| 157638 | FAM84B | 0.000156 | 0.635071 | 0.69745 | 0.654465 | 0.662329 | down |
| 9785 | DHX38 | 0.000156 | 0.544136 | 0.502807 | 0.466198 | 0.50438 | down |
| 152485 | ZNF827 | 0.000157 | 0.38411 | 0.48414 | 0.400537 | 0.422929 | down |
| 10137 | RBM12 | 0.000157 | 0.633868 | 0.525758 | 0.592262 | 0.583963 | down |
| 8503 | PIK3R3 | 0.000157 | 0.637275 | 0.621436 | 0.553641 | 0.604117 | down |
| 27347 | STK39 | 0.000157 | 0.589587 | 0.537375 | 0.566479 | 0.56448 | down |
| 54884 | RETSAT | 0.000157 | 0.507546 | 0.544466 | 0.473836 | 0.508616 | down |
| 8945 | BTRC | 0.000158 | 0.526117 | 0.480033 | 0.554301 | 0.52015 | down |
| 55326 | AGPAT5 | 0.000158 | 0.645818 | 0.560201 | 0.642214 | 0.616078 | down |
| 8514 | KCNAB2 | 0.000159 | 0.49375 | 0.493394 | 0.580674 | 0.522606 | down |
| 27436 | EML4 | 0.000159 | 0.497672 | 0.55331 | 0.570672 | 0.540551 | down |

| | | | | | | | |
|---|---|---|---|---|---|---|---|
| 7205 | TRIP6 | 0.000159 | 0.612977 | 0.642211 | 0.671962 | 0.642383 | down |
| 399687 | MYO18A | 0.000159 | 0.355203 | 0.417429 | 0.404854 | 0.392495 | down |
| 81566 | CSRNP2 | 0.000159 | 0.636012 | 0.572221 | 0.646765 | 0.618332 | down |
| 3636 | INPPL1 | 0.00016 | 0.574157 | 0.584209 | 0.566827 | 0.575064 | down |
| 7804 | LRP8 | 0.00016 | 0.544861 | 0.431221 | 0.508165 | 0.494749 | down |
| 10426 | TUBGCP3 | 0.00016 | 0.544195 | 0.520921 | 0.518638 | 0.527918 | down |
| 375033 | PEAR1 | 0.00016 | 0.643148 | 0.618512 | 0.589019 | 0.616893 | down |
| 11108 | PRDM4 | 0.00016 | 0.506678 | 0.545312 | 0.550529 | 0.534173 | down |
| 9931 | HELZ | 0.000161 | 0.451642 | 0.52881 | 0.419796 | 0.466749 | down |
| 256987 | SERINC5 | 0.000161 | 0.354783 | 0.456505 | 0.4454 | 0.418896 | down |
| 8542 | APOL1 | 0.000161 | 0.63814 | 0.640515 | 0.69313 | 0.657262 | down |
| 2059 | EPS8 | 0.000161 | 0.676891 | 0.643608 | 0.597987 | 0.639496 | down |
| 10276 | NET1 | 0.000161 | 0.501625 | 0.565049 | 0.531449 | 0.532708 | down |
| 93621 | MRFAP1 | 0.000161 | 0.578171 | 0.577249 | 0.564541 | 0.57332 | down |
| 10391 | CORO2B | 0.000161 | 0.527755 | 0.580589 | 0.485385 | 0.531243 | down |
| 85379 | KIAA1671 | 0.000161 | 0.348537 | 0.366847 | 0.315243 | 0.343543 | down |
| 25943 | C20orf194 | 0.000162 | 0.469422 | 0.44818 | 0.400618 | 0.439407 | down |
| 83786 | FRMD8 | 0.000162 | 0.546173 | 0.55203 | 0.521467 | 0.53989 | down |
| 80724 | ACAD10 | 0.000163 | 0.561306 | 0.511451 | 0.458941 | 0.510566 | down |
| 83607 | AMMECR1 | 0.000163 | 0.623648 | 0.533411 | 0.608247 | 0.588435 | down |
| 10019 | SH2B3 | 0.000163 | 0.453553 | 0.493672 | 0.496081 | 0.481102 | down |
| 8626 | TP63 | 0.000163 | 0.500108 | 0.479421 | 0.585591 | 0.521706 | down |
| 205564 | SENP5 | 0.000163 | 0.516929 | 0.52646 | 0.550402 | 0.531264 | down |
| 85377 | MICALL1 | 0.000164 | 0.683359 | 0.622875 | 0.592048 | 0.632761 | down |
| 54462 | CCSER2 | 0.000164 | 0.540139 | 0.469754 | 0.562006 | 0.523966 | down |
| 84951 | TNS4 | 0.000165 | 0.561783 | 0.473715 | 0.488953 | 0.50815 | down |
| 8458 | TTF2 | 0.000165 | 0.463673 | 0.450027 | 0.486557 | 0.466752 | down |
| 55210 | ATAD3A | 0.000165 | 0.661165 | 0.602836 | 0.563056 | 0.609019 | down |
| 1039 | CDR2 | 0.000165 | 0.480598 | 0.484787 | 0.532158 | 0.499181 | down |
| 5887 | RAD23B | 0.000165 | 0.601621 | 0.612177 | 0.60418 | 0.605993 | down |
| 53838 | C11orf24 | 0.000165 | 0.645576 | 0.654521 | 0.682398 | 0.660832 | down |
| 115 | ADCY9 | 0.000166 | 0.540076 | 0.397468 | 0.474597 | 0.470714 | down |
| 338442 | HCAR2 | 0.000166 | 0.609933 | 0.617387 | 0.60033 | 0.609217 | down |
| 2132 | EXT2 | 0.000166 | 0.504819 | 0.534951 | 0.566088 | 0.535286 | down |
| 1969 | EPHA2 | 0.000166 | 0.620606 | 0.545832 | 0.527212 | 0.56455 | down |
| 121665 | SPPL3 | 0.000167 | 0.670529 | 0.640044 | 0.57207 | 0.627548 | down |
| 4298 | MLLT1 | 0.000167 | 0.581627 | 0.5283 | 0.502266 | 0.537398 | down |
| 64855 | FAM129B | 0.000167 | 0.671941 | 0.587343 | 0.596851 | 0.618712 | down |
| 2923 | PDIA3 | 0.000167 | 0.68368 | 0.647719 | 0.641358 | 0.657586 | down |
| 8660 | IRS2 | 0.000167 | 0.52863 | 0.466994 | 0.524198 | 0.506607 | down |
| 55746 | NUP133 | 0.000167 | 0.508733 | 0.576078 | 0.494671 | 0.526494 | down |
| 4209 | MEF2D | 0.000168 | 0.528257 | 0.48648 | 0.532563 | 0.515767 | down |
| 10769 | PLK2 | 0.000168 | 0.591692 | 0.628955 | 0.59891 | 0.606519 | down |
| 3673 | ITGA2 | 0.000168 | 0.396347 | 0.422187 | 0.476638 | 0.431724 | down |
| 255738 | PCSK9 | 0.000169 | 0.557938 | 0.546043 | 0.569216 | 0.557733 | down |
| 79867 | TCTN2 | 0.000169 | 0.538891 | 0.472937 | 0.492506 | 0.501445 | down |
| 80196 | RNF34 | 0.000169 | 0.575595 | 0.636417 | 0.636152 | 0.616055 | down |
| 3913 | LAMB2 | 0.000169 | 0.486248 | 0.507373 | 0.427882 | 0.473834 | down |
| 9778 | KIAA0232 | 0.000169 | 0.442233 | 0.528818 | 0.464272 | 0.478441 | down |
| 317762 | CCDC85C | 0.000169 | 0.594213 | 0.659303 | 0.599017 | 0.617511 | down |
| 55738 | ARFGAP1 | 0.00017 | 0.57543 | 0.68202 | 0.634588 | 0.630679 | down |
| 545 | ATR | 0.00017 | 0.494604 | 0.507865 | 0.531818 | 0.511429 | down |
| 83892 | KCTD10 | 0.00017 | 0.619082 | 0.668424 | 0.681968 | 0.656491 | down |
| 293 | SLC25A6 | 0.00017 | 0.632935 | 0.617201 | 0.534403 | 0.594846 | down |
| 55294 | FBXW7 | 0.000171 | 0.483451 | 0.481925 | 0.481391 | 0.482256 | down |
| 56889 | TM9SF3 | 0.000171 | 0.579504 | 0.61534 | 0.657098 | 0.617314 | down |
| 127933 | UHMK1 | 0.000171 | 0.578004 | 0.614682 | 0.615384 | 0.60269 | down |
| 53349 | ZFYVE1 | 0.000171 | 0.709722 | 0.62028 | 0.6305 | 0.6535 | down |

| | | | | | | | |
|---|---|---|---|---|---|---|---|
| 79364 | ZXDC | 0.000172 | 0.617985 | 0.526032 | 0.520532 | 0.55485 | down |
| 9113 | LATS1 | 0.000172 | 0.416841 | 0.46627 | 0.432321 | 0.438477 | down |
| 6780 | STAU1 | 0.000172 | 0.549639 | 0.572917 | 0.539441 | 0.553999 | down |
| 10147 | SUGP2 | 0.000172 | 0.560121 | 0.604943 | 0.534671 | 0.566578 | down |
| 64601 | VPS16 | 0.000173 | 0.627813 | 0.650736 | 0.555133 | 0.611227 | down |
| 23168 | RTF1 | 0.000173 | 0.622621 | 0.578609 | 0.617731 | 0.60632 | down |
| 9975 | NR1D2 | 0.000173 | 0.543745 | 0.568877 | 0.600598 | 0.571074 | down |
| 80024 | SLC8B1 | 0.000173 | 0.6663 | 0.622961 | 0.564281 | 0.617847 | down |
| 55731 | FAM222B | 0.000173 | 0.472103 | 0.426063 | 0.417029 | 0.438398 | down |
| 1027 | CDKN1B | 0.000174 | 0.532598 | 0.634761 | 0.529198 | 0.565519 | down |
| 54788 | DNAJB12 | 0.000174 | 0.624059 | 0.549647 | 0.583345 | 0.585683 | down |
| 1388 | ATF6B | 0.000174 | 0.573195 | 0.501983 | 0.595076 | 0.556751 | down |
| 254065 | BRWD3 | 0.000175 | 0.369136 | 0.412244 | 0.515782 | 0.432387 | down |
| 57787 | MARK4 | 0.000175 | 0.505578 | 0.610806 | 0.538531 | 0.551638 | down |
| 126298 | IRGQ | 0.000176 | 0.505595 | 0.619541 | 0.523038 | 0.549391 | down |
| 8607 | RUVBL1 | 0.000176 | 0.58434 | 0.648138 | 0.584088 | 0.605522 | down |
| 53981 | CPSF2 | 0.000177 | 0.553765 | 0.660367 | 0.574681 | 0.596271 | down |
| 2140 | EYA3 | 0.000177 | 0.627123 | 0.672191 | 0.590468 | 0.629927 | down |
| 51564 | HDAC7 | 0.000177 | 0.623239 | 0.689245 | 0.586119 | 0.632868 | down |
| 5976 | UPF1 | 0.000177 | 0.461125 | 0.479854 | 0.415498 | 0.452159 | down |
| 6256 | RXRA | 0.000178 | 0.637598 | 0.613583 | 0.621092 | 0.624091 | down |
| 91056 | AP5B1 | 0.000178 | 0.66498 | 0.597638 | 0.601397 | 0.621338 | down |
| 8227 | AKAP17A | 0.000178 | 0.445293 | 0.393813 | 0.509513 | 0.44954 | down |
| 50 | ACO2 | 0.000179 | 0.573819 | 0.570975 | 0.506495 | 0.55043 | down |
| 92689 | FAM114A1 | 0.000179 | 0.53875 | 0.607903 | 0.65405 | 0.600234 | down |
| 8777 | MPDZ | 0.00018 | 0.464859 | 0.429418 | 0.458333 | 0.45087 | down |
| 6744 | ITPRID2 | 0.00018 | 0.609877 | 0.677467 | 0.681177 | 0.656174 | down |
| 5062 | PAK2 | 0.00018 | 0.467108 | 0.540792 | 0.496725 | 0.501542 | down |
| 3714 | JAG2 | 0.00018 | 0.539664 | 0.561949 | 0.488792 | 0.530135 | down |
| 55177 | RMDN3 | 0.000181 | 0.589829 | 0.538969 | 0.581253 | 0.570017 | down |
| 115207 | KCTD12 | 0.000181 | 0.495431 | 0.55473 | 0.575323 | 0.541828 | down |
| 54914 | FOCAD | 0.000181 | 0.434368 | 0.490358 | 0.427618 | 0.450782 | down |
| 23576 | DDAH1 | 0.000181 | 0.626927 | 0.660432 | 0.693127 | 0.660162 | down |
| 10721 | POLQ | 0.000182 | 0.482579 | 0.573702 | 0.514384 | 0.523555 | down |
| 5613 | PRKX | 0.000182 | 0.593191 | 0.657895 | 0.691394 | 0.647493 | down |
| 3992 | FADS1 | 0.000183 | 0.567059 | 0.596664 | 0.627434 | 0.597052 | down |
| 374654 | KIF7 | 0.000183 | 0.405215 | 0.473051 | 0.382581 | 0.420282 | down |
| 2548 | GAA | 0.000183 | 0.608556 | 0.652039 | 0.551055 | 0.603883 | down |
| 5937 | RBMS1 | 0.000183 | 0.591821 | 0.654828 | 0.641512 | 0.629387 | down |
| 6874 | TAF4 | 0.000183 | 0.595395 | 0.488095 | 0.533516 | 0.539002 | down |
| 9605 | VPS9D1 | 0.000183 | 0.592807 | 0.648652 | 0.565746 | 0.602402 | down |
| 27348 | TOR1B | 0.000183 | 0.459557 | 0.459149 | 0.566765 | 0.495157 | down |
| 162073 | ITPRIPL2 | 0.000184 | 0.386617 | 0.424555 | 0.390579 | 0.400584 | down |
| 80331 | DNAJC5 | 0.000184 | 0.575724 | 0.600463 | 0.532334 | 0.569507 | down |
| 23503 | ZFYVE26 | 0.000184 | 0.45829 | 0.473462 | 0.469516 | 0.467089 | down |
| 5087 | PBX1 | 0.000184 | 0.487466 | 0.535948 | 0.583993 | 0.535802 | down |
| 57670 | KIAA1549 | 0.000184 | 0.404613 | 0.456385 | 0.378559 | 0.413186 | down |
| 85457 | CIPC | 0.000184 | 0.547143 | 0.568974 | 0.610807 | 0.575641 | down |
| 56910 | STARD7 | 0.000184 | 0.542046 | 0.583284 | 0.569784 | 0.565038 | down |
| 642 | BLMH | 0.000184 | 0.618108 | 0.536349 | 0.551171 | 0.568543 | down |
| 10927 | SPIN1 | 0.000184 | 0.554068 | 0.62437 | 0.617319 | 0.598586 | down |
| 23334 | SZT2 | 0.000185 | 0.507848 | 0.545064 | 0.509814 | 0.520909 | down |
| 57505 | AARS2 | 0.000185 | 0.453051 | 0.487185 | 0.577269 | 0.505835 | down |
| 3983 | ABLIM1 | 0.000186 | 0.378857 | 0.414157 | 0.424054 | 0.405689 | down |
| 149076 | ZNF362 | 0.000186 | 0.565368 | 0.505665 | 0.540278 | 0.537104 | down |
| 6249 | CLIP1 | 0.000186 | 0.454651 | 0.538158 | 0.52072 | 0.50451 | down |
| 163590 | TOR1AIP2 | 0.000186 | 0.471266 | 0.561721 | 0.52192 | 0.518302 | down |
| 3911 | LAMA5 | 0.000186 | 0.566323 | 0.592154 | 0.534109 | 0.564195 | down |

| | | | | | | | |
|---|---|---|---|---|---|---|---|
| 84464 | SLX4 | 0.000187 | 0.427141 | 0.445146 | 0.465707 | 0.445998 | down |
| 3998 | LMAN1 | 0.000187 | 0.621776 | 0.606687 | 0.658149 | 0.628871 | down |
| 3703 | STT3A | 0.000187 | 0.61404 | 0.608289 | 0.662287 | 0.628205 | down |
| 23039 | XPO7 | 0.000188 | 0.496326 | 0.53176 | 0.568215 | 0.5321 | down |
| 55148 | UBR7 | 0.000188 | 0.561102 | 0.64122 | 0.670051 | 0.624124 | down |
| 653464 | SRGAP2C | 0.000188 | 0.592556 | 0.558885 | 0.540386 | 0.563942 | down |
| 54890 | ALKBH5 | 0.000189 | 0.621721 | 0.583091 | 0.540732 | 0.581848 | down |
| 8208 | CHAF1B | 0.000189 | 0.488334 | 0.605 | 0.527366 | 0.540233 | down |
| 55930 | MYO5C | 0.00019 | 0.465565 | 0.425623 | 0.421608 | 0.437598 | down |
| 64219 | PJA1 | 0.00019 | 0.66836 | 0.612135 | 0.694954 | 0.658483 | down |
| 6655 | SOS2 | 0.00019 | 0.367256 | 0.521699 | 0.437624 | 0.442193 | down |
| 2037 | EPB41L2 | 0.000191 | 0.550836 | 0.53089 | 0.42167 | 0.501132 | down |
| 137964 | GPAT4 | 0.000191 | 0.561919 | 0.658616 | 0.602577 | 0.607704 | down |
| 144983 | HNRNPA1 | 0.000191 | 0.553483 | 0.601798 | 0.617947 | 0.591076 | down |
| 79716 | NPEPL1 | 0.000191 | 0.627396 | 0.625525 | 0.537517 | 0.596813 | down |
| 114818 | KLHL29 | 0.000191 | 0.557736 | 0.519094 | 0.490893 | 0.522574 | down |
| 8507 | ENC1 | 0.000191 | 0.453094 | 0.50635 | 0.472896 | 0.477447 | down |
| 440275 | EIF2AK4 | 0.000191 | 0.461074 | 0.533875 | 0.530494 | 0.508481 | down |
| 9855 | FARP2 | 0.000191 | 0.505369 | 0.473458 | 0.433238 | 0.470688 | down |
| 9640 | ZNF592 | 0.000191 | 0.569466 | 0.645824 | 0.58984 | 0.60171 | down |
| 8795 | TNFRSF10I | 0.000192 | 0.518107 | 0.544094 | 0.530405 | 0.530869 | down |
| 9667 | SAFB2 | 0.000192 | 0.325469 | 0.308021 | 0.358615 | 0.330702 | down |
| 260425 | MAGI3 | 0.000192 | 0.464069 | 0.515152 | 0.593956 | 0.524392 | down |
| 91607 | SLFN11 | 0.000192 | 0.570468 | 0.666632 | 0.631871 | 0.62299 | down |
| 10181 | RBM5 | 0.000192 | 0.63692 | 0.562294 | 0.569176 | 0.589463 | down |
| 1487 | CTBP1 | 0.000192 | 0.626621 | 0.618152 | 0.5688 | 0.604524 | down |
| 9654 | TTLL4 | 0.000192 | 0.440268 | 0.535096 | 0.47561 | 0.483658 | down |
| 126917 | IFFO2 | 0.000193 | 0.485448 | 0.441743 | 0.519819 | 0.482337 | down |
| 23235 | SIK2 | 0.000194 | 0.533004 | 0.509779 | 0.510019 | 0.517601 | down |
| 23468 | CBX5 | 0.000194 | 0.522391 | 0.58718 | 0.563087 | 0.557553 | down |
| 26030 | PLEKHG3 | 0.000194 | 0.617189 | 0.657884 | 0.558244 | 0.611106 | down |
| 57224 | NHSL1 | 0.000194 | 0.486348 | 0.399251 | 0.399753 | 0.428451 | down |
| 55920 | RCC2 | 0.000194 | 0.454113 | 0.521713 | 0.429281 | 0.468369 | down |
| 7003 | TEAD1 | 0.000195 | 0.347795 | 0.419933 | 0.468532 | 0.412087 | down |
| 4249 | MGAT5 | 0.000195 | 0.393713 | 0.436499 | 0.437852 | 0.422688 | down |
| 51776 | MAP3K20 | 0.000196 | 0.568732 | 0.616421 | 0.586854 | 0.590669 | down |
| 90161 | HS6ST2 | 0.000196 | 0.564617 | 0.664397 | 0.654407 | 0.627807 | down |
| 79085 | SLC25A23 | 0.000196 | 0.575976 | 0.589009 | 0.587448 | 0.584144 | down |
| 92999 | ZBTB47 | 0.000197 | 0.580926 | 0.660861 | 0.65185 | 0.631212 | down |
| 10560 | SLC19A2 | 0.000197 | 0.595126 | 0.594728 | 0.524594 | 0.571483 | down |
| 11056 | DDX52 | 0.000197 | 0.589497 | 0.698504 | 0.657604 | 0.648535 | down |
| 9727 | RAB11FIP3 | 0.000198 | 0.565938 | 0.534357 | 0.605468 | 0.568588 | down |
| 2957 | GTF2A1 | 0.000198 | 0.545811 | 0.553518 | 0.601062 | 0.566797 | down |
| 5797 | PTPRM | 0.000198 | 0.479774 | 0.490422 | 0.439788 | 0.469994 | down |
| 23201 | FAM168A | 0.000198 | 0.519173 | 0.509049 | 0.526407 | 0.51821 | down |
| 27352 | SGSM3 | 0.000199 | 0.700654 | 0.64428 | 0.653642 | 0.666192 | down |
| 3054 | HCFC1 | 0.000199 | 0.473461 | 0.521803 | 0.460032 | 0.485099 | down |
| 23233 | EXOC6B | 0.000199 | 0.439525 | 0.563896 | 0.477818 | 0.493747 | down |
| 113178 | SCAMP4 | 0.000199 | 0.627685 | 0.629907 | 0.603361 | 0.620318 | down |
| 79080 | CCDC86 | 0.000199 | 0.664704 | 0.571288 | 0.607272 | 0.614421 | down |
| 65998 | C11orf95 | 0.000199 | 0.482032 | 0.415633 | 0.522789 | 0.473485 | down |
| 57572 | DOCK6 | 0.0002 | 0.602643 | 0.523835 | 0.580988 | 0.569155 | down |
| 23077 | MYCBP2 | 0.0002 | 0.360937 | 0.417225 | 0.405181 | 0.394448 | down |
| 57162 | PELI1 | 0.0002 | 0.531034 | 0.513808 | 0.478304 | 0.507715 | down |
| 57822 | GRHL3 | 0.000201 | 0.517329 | 0.485005 | 0.443018 | 0.481784 | down |
| 79827 | CLMP | 0.000201 | 0.566986 | 0.628173 | 0.627227 | 0.607462 | down |
| 9738 | CCP110 | 0.000201 | 0.623724 | 0.594048 | 0.656503 | 0.624758 | down |
| 200014 | CC2D1B | 0.000201 | 0.585626 | 0.636507 | 0.597276 | 0.60647 | down |

| | | | | | | | |
|---|---|---|---|---|---|---|---|
| 203427 | SLC25A43 | 0.000202 | 0.455894 | 0.542781 | 0.519408 | 0.506028 | down |
| 58499 | ZNF462 | 0.000203 | 0.409517 | 0.379398 | 0.408477 | 0.39913 | down |
| 25998 | IBTK | 0.000203 | 0.54421 | 0.612446 | 0.618764 | 0.591807 | down |
| 53373 | TPCN1 | 0.000204 | 0.609038 | 0.576792 | 0.65034 | 0.612057 | down |
| 29070 | CCDC113 | 0.000204 | 0.558479 | 0.562654 | 0.582567 | 0.5679 | down |
| 55917 | CTTNBP2N | 0.000205 | 0.419107 | 0.458894 | 0.543664 | 0.473888 | down |
| 8862 | APLN | 0.000205 | 0.618223 | 0.517686 | 0.512515 | 0.549475 | down |
| 64359 | NXN | 0.000205 | 0.51104 | 0.552103 | 0.492059 | 0.518401 | down |
| 57018 | CCNL1 | 0.000206 | 0.598019 | 0.633172 | 0.562879 | 0.598023 | down |
| 2117 | ETV3 | 0.000206 | 0.380956 | 0.499151 | 0.418354 | 0.43282 | down |
| 57541 | ZNF398 | 0.000206 | 0.550692 | 0.461083 | 0.52419 | 0.511988 | down |
| 85012 | TCEAL3 | 0.000207 | 0.648192 | 0.594586 | 0.706854 | 0.649877 | down |
| 90780 | PYGO2 | 0.000207 | 0.667306 | 0.570657 | 0.560828 | 0.599597 | down |
| 7424 | VEGFC | 0.000207 | 0.614367 | 0.647372 | 0.621135 | 0.627625 | down |
| 22998 | LIMCH1 | 0.000207 | 0.427241 | 0.413004 | 0.428674 | 0.422973 | down |
| 124930 | ANKRD13E | 0.000207 | 0.601733 | 0.590154 | 0.496488 | 0.562792 | down |
| 57555 | NLGN2 | 0.000207 | 0.585331 | 0.498578 | 0.536384 | 0.540098 | down |
| 54680 | ZNHIT6 | 0.000207 | 0.494238 | 0.584553 | 0.530411 | 0.536401 | down |
| 7297 | TYK2 | 0.000208 | 0.583966 | 0.637733 | 0.557013 | 0.592904 | down |
| 57609 | DIP2B | 0.000208 | 0.392571 | 0.457002 | 0.464498 | 0.438024 | down |
| 81603 | TRIM8 | 0.000208 | 0.562106 | 0.604723 | 0.55043 | 0.57242 | down |
| 23368 | PPP1R13B | 0.000208 | 0.530986 | 0.487133 | 0.511851 | 0.50999 | down |
| 10146 | G3BP1 | 0.000208 | 0.643354 | 0.702743 | 0.631275 | 0.659124 | down |
| 197131 | UBR1 | 0.000209 | 0.51953 | 0.568231 | 0.555439 | 0.547733 | down |
| 2086 | ERV3-1 | 0.000209 | 0.637088 | 0.666874 | 0.676632 | 0.660198 | down |
| 65125 | WNK1 | 0.00021 | 0.347532 | 0.416169 | 0.419878 | 0.394526 | down |
| 57109 | REXO4 | 0.000211 | 0.675717 | 0.619252 | 0.628289 | 0.641086 | down |
| 54906 | FAM208B | 0.000211 | 0.422903 | 0.539705 | 0.497283 | 0.48663 | down |
| 23277 | CLUH | 0.000211 | 0.535311 | 0.546448 | 0.513001 | 0.531587 | down |
| 80131 | LRRC8E | 0.000211 | 0.580801 | 0.576854 | 0.58103 | 0.579562 | down |
| 10985 | GCN1 | 0.000212 | 0.452981 | 0.471036 | 0.45553 | 0.459849 | down |
| 55243 | KIRREL1 | 0.000213 | 0.40225 | 0.485689 | 0.482721 | 0.456887 | down |
| 55119 | PRPF38B | 0.000213 | 0.50555 | 0.591059 | 0.567913 | 0.554841 | down |
| 79794 | C12orf49 | 0.000213 | 0.608121 | 0.670462 | 0.656583 | 0.645055 | down |
| 28514 | DLL1 | 0.000214 | 0.532339 | 0.390037 | 0.450704 | 0.457693 | down |
| 57488 | ESYT2 | 0.000214 | 0.53041 | 0.634423 | 0.575537 | 0.580123 | down |
| 94121 | SYTL4 | 0.000214 | 0.526139 | 0.561007 | 0.517424 | 0.534857 | down |
| 552889 | ATXN7L3B | 0.000214 | 0.643682 | 0.568519 | 0.652835 | 0.621679 | down |
| 473 | RERE | 0.000215 | 0.535506 | 0.490476 | 0.444056 | 0.490013 | down |
| 3678 | ITGA5 | 0.000215 | 0.598011 | 0.574082 | 0.571852 | 0.581315 | down |
| 23527 | ACAP2 | 0.000216 | 0.445219 | 0.512573 | 0.50578 | 0.487857 | down |
| 54916 | TMEM260 | 0.000216 | 0.530217 | 0.559247 | 0.611519 | 0.566994 | down |
| 57679 | ALS2 | 0.000216 | 0.53592 | 0.514249 | 0.612014 | 0.554061 | down |
| 24139 | EML2 | 0.000216 | 0.474268 | 0.582553 | 0.44933 | 0.50205 | down |
| 3658 | IREB2 | 0.000217 | 0.538307 | 0.600649 | 0.526592 | 0.555183 | down |
| 57493 | HEG1 | 0.000217 | 0.402118 | 0.449988 | 0.441195 | 0.4311 | down |
| 1822 | ATN1 | 0.000217 | 0.619804 | 0.503457 | 0.539402 | 0.554221 | down |
| 8498 | RANBP3 | 0.000217 | 0.638147 | 0.589701 | 0.587518 | 0.605122 | down |
| 6238 | RRBP1 | 0.000218 | 0.431099 | 0.364035 | 0.389932 | 0.395022 | down |
| 157285 | PRAG1 | 0.000218 | 0.518646 | 0.576944 | 0.54727 | 0.54762 | down |
| 6421 | SFPQ | 0.000218 | 0.520061 | 0.517422 | 0.445505 | 0.494329 | down |
| 23580 | CDC42EP4 | 0.000218 | 0.525075 | 0.584071 | 0.627202 | 0.578783 | down |
| 9725 | TMEM63A | 0.000218 | 0.620471 | 0.501464 | 0.596346 | 0.57276 | down |
| 8427 | ZNF282 | 0.000218 | 0.68473 | 0.611733 | 0.60987 | 0.635444 | down |
| 89781 | HPS4 | 0.000218 | 0.538705 | 0.59026 | 0.467038 | 0.532001 | down |
| 9043 | SPAG9 | 0.000219 | 0.445537 | 0.551995 | 0.487091 | 0.494874 | down |
| 1140 | CHRNB1 | 0.000219 | 0.592783 | 0.57876 | 0.644757 | 0.605433 | down |
| 64129 | TINAGL1 | 0.000219 | 0.69332 | 0.61746 | 0.609712 | 0.640164 | down |

| | | | | | | | |
|---|---|---|---|---|---|---|---|
| 2926 | GRSF1 | 0.000219 | 0.604992 | 0.677533 | 0.632906 | 0.638477 | down |
| 405 | ARNT | 0.000219 | 0.491268 | 0.538719 | 0.559751 | 0.529913 | down |
| 10114 | HIPK3 | 0.000219 | 0.542117 | 0.641317 | 0.633023 | 0.605485 | down |
| 255967 | PAN3 | 0.00022 | 0.523131 | 0.586013 | 0.47271 | 0.527285 | down |
| 23064 | SETX | 0.00022 | 0.480382 | 0.489053 | 0.462305 | 0.477247 | down |
| 84193 | SETD3 | 0.00022 | 0.463852 | 0.563788 | 0.58489 | 0.53751 | down |
| 399512 | SLC25A35 | 0.000221 | 0.687465 | 0.676243 | 0.58888 | 0.650863 | down |
| 8019 | BRD3 | 0.000221 | 0.583563 | 0.491341 | 0.618614 | 0.564506 | down |
| 607 | BCL9 | 0.000221 | 0.463843 | 0.517037 | 0.561705 | 0.514195 | down |
| 26010 | SPATS2L | 0.000222 | 0.581227 | 0.571851 | 0.597445 | 0.583507 | down |
| 8761 | PABPC4 | 0.000223 | 0.517304 | 0.544245 | 0.474129 | 0.511893 | down |
| 3845 | KRAS | 0.000223 | 0.543906 | 0.608898 | 0.573191 | 0.575332 | down |
| 9320 | TRIP12 | 0.000224 | 0.533876 | 0.590496 | 0.582576 | 0.568983 | down |
| 26098 | EDRF1 | 0.000224 | 0.479955 | 0.485054 | 0.50006 | 0.488356 | down |
| 84901 | NFATC2IP | 0.000225 | 0.595429 | 0.662199 | 0.659015 | 0.638881 | down |
| 10389 | SCML2 | 0.000225 | 0.52623 | 0.516495 | 0.56482 | 0.535848 | down |
| 57472 | CNOT6 | 0.000226 | 0.555218 | 0.611668 | 0.580619 | 0.582502 | down |
| 80381 | CD276 | 0.000226 | 0.690773 | 0.581422 | 0.630784 | 0.634326 | down |
| 4804 | NGFR | 0.000226 | 0.635077 | 0.529317 | 0.576888 | 0.580427 | down |
| 126353 | MISP | 0.000227 | 0.602741 | 0.486412 | 0.56898 | 0.552711 | down |
| 55719 | SLF2 | 0.000227 | 0.50011 | 0.62882 | 0.520124 | 0.549685 | down |
| 63925 | ZNF335 | 0.000227 | 0.603723 | 0.49056 | 0.464969 | 0.519751 | down |
| 23466 | CBX6 | 0.000227 | 0.516852 | 0.532055 | 0.50004 | 0.516316 | down |
| 29967 | LRP12 | 0.000227 | 0.488305 | 0.589689 | 0.57239 | 0.550128 | down |
| 4644 | MYO5A | 0.000227 | 0.492444 | 0.403284 | 0.485986 | 0.460571 | down |
| 5528 | PPP2R5D | 0.000227 | 0.633325 | 0.53689 | 0.60999 | 0.593402 | down |
| 23299 | BICD2 | 0.000227 | 0.418631 | 0.482348 | 0.421274 | 0.440751 | down |
| 63950 | DMRTA2 | 0.000228 | 0.56338 | 0.534124 | 0.475924 | 0.524476 | down |
| 6491 | STIL | 0.000228 | 0.602398 | 0.580568 | 0.598836 | 0.593934 | down |
| 23384 | SPECC1L | 0.000229 | 0.491503 | 0.426826 | 0.43631 | 0.451547 | down |
| 55796 | MBNL3 | 0.000229 | 0.668076 | 0.624143 | 0.604082 | 0.6321 | down |
| 151887 | CCDC80 | 0.00023 | 0.517117 | 0.620845 | 0.537927 | 0.55863 | down |
| 440193 | CCDC88C | 0.00023 | 0.379073 | 0.374758 | 0.453733 | 0.402522 | down |
| 8324 | FZD7 | 0.00023 | 0.625 | 0.550388 | 0.565179 | 0.580189 | down |
| 7057 | THBS1 | 0.000231 | 0.387497 | 0.456095 | 0.400477 | 0.41469 | down |
| 4193 | MDM2 | 0.000231 | 0.579082 | 0.646659 | 0.672128 | 0.632623 | down |
| 23293 | SMG6 | 0.000231 | 0.56684 | 0.547483 | 0.508676 | 0.540999 | down |
| 7030 | TFE3 | 0.000232 | 0.699445 | 0.594044 | 0.599829 | 0.631106 | down |
| 4026 | LPP | 0.000232 | 0.527568 | 0.589017 | 0.613226 | 0.576603 | down |
| 3839 | KPNA3 | 0.000232 | 0.636488 | 0.71705 | 0.638173 | 0.663904 | down |
| 23301 | EHBP1 | 0.000232 | 0.564179 | 0.658005 | 0.593526 | 0.605237 | down |
| 55357 | TBC1D2 | 0.000232 | 0.606821 | 0.610987 | 0.566933 | 0.594914 | down |
| 699 | BUB1 | 0.000232 | 0.660879 | 0.693253 | 0.645324 | 0.666485 | down |
| 1497 | CTNS | 0.000232 | 0.652312 | 0.705706 | 0.604905 | 0.654308 | down |
| 55833 | UBAP2 | 0.000232 | 0.391412 | 0.510276 | 0.503416 | 0.468368 | down |
| 57674 | RNF213 | 0.000233 | 0.388819 | 0.410735 | 0.406689 | 0.402081 | down |
| 55778 | ZNF839 | 0.000233 | 0.577464 | 0.672604 | 0.679182 | 0.643084 | down |
| 4670 | HNRNPM | 0.000233 | 0.498267 | 0.51692 | 0.428916 | 0.481368 | down |
| 2531 | KDSR | 0.000233 | 0.588385 | 0.578991 | 0.625215 | 0.59753 | down |
| 51512 | GTSE1 | 0.000234 | 0.640884 | 0.625132 | 0.640529 | 0.635515 | down |
| 60481 | ELOVL5 | 0.000234 | 0.662626 | 0.618622 | 0.666193 | 0.649147 | down |
| 2004 | ELK3 | 0.000234 | 0.52259 | 0.574819 | 0.553061 | 0.550157 | down |
| 199953 | TMEM201 | 0.000234 | 0.603281 | 0.559856 | 0.63569 | 0.599609 | down |
| 9181 | ARHGEF2 | 0.000235 | 0.571936 | 0.505943 | 0.523137 | 0.533672 | down |
| 129642 | MBOAT2 | 0.000235 | 0.541498 | 0.550341 | 0.637402 | 0.576413 | down |
| 3607 | FOXK2 | 0.000235 | 0.5661 | 0.638313 | 0.586125 | 0.596846 | down |
| 200894 | ARL13B | 0.000236 | 0.566635 | 0.641623 | 0.604123 | 0.604127 | down |
| 6720 | SREBF1 | 0.000236 | 0.62754 | 0.618905 | 0.614352 | 0.620266 | down |

| | | | | | | | |
|---|---|---|---|---|---|---|---|
| 80004 | ESRP2 | 0.000236 | 0.574313 | 0.532877 | 0.640991 | 0.582727 | down |
| 25930 | PTPN23 | 0.000237 | 0.627964 | 0.543147 | 0.536058 | 0.569056 | down |
| 26469 | PTPN18 | 0.000237 | 0.675634 | 0.610955 | 0.630723 | 0.639104 | down |
| 9054 | NFS1 | 0.000237 | 0.630119 | 0.672209 | 0.558901 | 0.62041 | down |
| 55225 | RAVER2 | 0.000237 | 0.569003 | 0.531032 | 0.518037 | 0.539357 | down |
| 93380 | MMGT1 | 0.000237 | 0.525068 | 0.591923 | 0.580668 | 0.565886 | down |
| 3092 | HIP1 | 0.000238 | 0.438938 | 0.445101 | 0.434418 | 0.439486 | down |
| 2308 | FOXO1 | 0.000239 | 0.389964 | 0.546962 | 0.468641 | 0.468523 | down |
| 861 | RUNX1 | 0.000239 | 0.545231 | 0.460666 | 0.523882 | 0.509926 | down |
| 6787 | NEK4 | 0.000239 | 0.517731 | 0.458772 | 0.576342 | 0.517615 | down |
| 157506 | RDH10 | 0.000239 | 0.538167 | 0.522216 | 0.479843 | 0.513409 | down |
| 23259 | DDHD2 | 0.00024 | 0.445481 | 0.542169 | 0.470364 | 0.486005 | down |
| 8879 | SGPL1 | 0.000241 | 0.544194 | 0.574248 | 0.503938 | 0.540793 | down |
| 25966 | C2CD2 | 0.000241 | 0.469259 | 0.504269 | 0.553965 | 0.509164 | down |
| 22850 | ADNP2 | 0.000241 | 0.569899 | 0.568368 | 0.45842 | 0.532229 | down |
| 91754 | NEK9 | 0.000242 | 0.492076 | 0.448986 | 0.485803 | 0.475622 | down |
| 5074 | PAWR | 0.000242 | 0.498495 | 0.521152 | 0.613457 | 0.544368 | down |
| 197259 | MLKL | 0.000242 | 0.671511 | 0.583677 | 0.637484 | 0.63089 | down |
| 1.03E+08 | LOC10272 | 0.000243 | 0.573349 | 0.445131 | 0.486835 | 0.501771 | down |
| 160851 | DGKH | 0.000243 | 0.49378 | 0.539226 | 0.615495 | 0.5495 | down |
| 27 | ABL2 | 0.000244 | 0.458908 | 0.511116 | 0.49007 | 0.486698 | down |
| 79065 | ATG9A | 0.000244 | 0.625525 | 0.610022 | 0.508584 | 0.581377 | down |
| 10277 | UBE4B | 0.000244 | 0.363477 | 0.421931 | 0.414832 | 0.40008 | down |
| 56938 | ARNTL2 | 0.000244 | 0.572618 | 0.647233 | 0.659506 | 0.626452 | down |
| 11273 | ATXN2L | 0.000244 | 0.590426 | 0.470917 | 0.501254 | 0.520866 | down |
| 57678 | GPAM | 0.000244 | 0.389264 | 0.510894 | 0.529463 | 0.47654 | down |
| 58513 | EPS15L1 | 0.000244 | 0.587654 | 0.529708 | 0.519955 | 0.545772 | down |
| 1742 | DLG4 | 0.000246 | 0.65074 | 0.626007 | 0.652792 | 0.64318 | down |
| 9354 | UBE4A | 0.000246 | 0.42801 | 0.556554 | 0.506303 | 0.496955 | down |
| 57523 | NYNRIN | 0.000246 | 0.569116 | 0.483607 | 0.437018 | 0.49658 | down |
| 6446 | SGK1 | 0.000247 | 0.530686 | 0.52545 | 0.528038 | 0.528058 | down |
| 1911 | PHC1 | 0.000247 | 0.562092 | 0.536095 | 0.521122 | 0.53977 | down |
| 58476 | TP53INP2 | 0.000247 | 0.570302 | 0.513588 | 0.499984 | 0.527958 | down |
| 3487 | IGFBP4 | 0.000248 | 0.671837 | 0.635981 | 0.664136 | 0.657318 | down |
| 114793 | FMNL2 | 0.000248 | 0.468078 | 0.498985 | 0.589004 | 0.518689 | down |
| 11215 | AKAP11 | 0.000249 | 0.492899 | 0.538993 | 0.575854 | 0.535915 | down |
| 7702 | ZNF143 | 0.000249 | 0.549052 | 0.65781 | 0.591463 | 0.599441 | down |
| 10802 | SEC24A | 0.000249 | 0.538311 | 0.643902 | 0.625702 | 0.602638 | down |
| 6624 | FSCN1 | 0.00025 | 0.650512 | 0.616 | 0.563604 | 0.610038 | down |
| 9677 | PPIP5K1 | 0.00025 | 0.361301 | 0.479189 | 0.473251 | 0.437914 | down |
| 23678 | SGK3 | 0.00025 | 0.357646 | 0.404403 | 0.422545 | 0.394865 | down |
| 9456 | HOMER1 | 0.000251 | 0.582378 | 0.61806 | 0.570387 | 0.590275 | down |
| 7498 | XDH | 0.000251 | 0.53852 | 0.513066 | 0.462243 | 0.50461 | down |
| 54872 | PIGG | 0.000252 | 0.533489 | 0.544174 | 0.584677 | 0.554113 | down |
| 55636 | CHD7 | 0.000252 | 0.410984 | 0.512407 | 0.461765 | 0.461719 | down |
| 3801 | KIFC3 | 0.000252 | 0.634891 | 0.560932 | 0.500555 | 0.565459 | down |
| 23357 | ANGEL1 | 0.000252 | 0.534918 | 0.605562 | 0.633908 | 0.591463 | down |
| 10602 | CDC42EP3 | 0.000253 | 0.529021 | 0.594995 | 0.528851 | 0.550956 | down |
| 29128 | UHRF1 | 0.000253 | 0.418269 | 0.442841 | 0.346109 | 0.402407 | down |
| 6734 | SRPRA | 0.000254 | 0.442601 | 0.478371 | 0.517363 | 0.479445 | down |
| 54439 | RBM27 | 0.000254 | 0.595359 | 0.63208 | 0.700926 | 0.642788 | down |
| 9920 | KBTBD11 | 0.000254 | 0.334786 | 0.470895 | 0.471925 | 0.425869 | down |
| 7841 | MOGS | 0.000254 | 0.672503 | 0.670481 | 0.56885 | 0.637278 | down |
| 2886 | GRB7 | 0.000254 | 0.698275 | 0.650153 | 0.629388 | 0.659272 | down |
| 9841 | ZBTB24 | 0.000254 | 0.573931 | 0.497333 | 0.445359 | 0.505541 | down |
| 9569 | GTF2IRD1 | 0.000254 | 0.576448 | 0.487424 | 0.584429 | 0.549434 | down |
| 1948 | EFNB2 | 0.000255 | 0.615079 | 0.531818 | 0.527202 | 0.558033 | down |
| 2175 | FANCA | 0.000255 | 0.662359 | 0.615489 | 0.64254 | 0.640129 | down |

| | | | | | | | |
|---|---|---|---|---|---|---|---|
| 55205 | ZNF532 | 0.000256 | 0.423624 | 0.433462 | 0.531118 | 0.462735 | down |
| 112 | ADCY6 | 0.000257 | 0.615224 | 0.537524 | 0.635232 | 0.595994 | down |
| 25946 | ZNF385A | 0.000257 | 0.678883 | 0.559586 | 0.614371 | 0.617613 | down |
| 23683 | PRKD3 | 0.000258 | 0.539401 | 0.611745 | 0.621899 | 0.591015 | down |
| 55230 | USP40 | 0.000258 | 0.411286 | 0.514737 | 0.478889 | 0.468304 | down |
| 4125 | MAN2B1 | 0.000258 | 0.631104 | 0.619157 | 0.519546 | 0.589936 | down |
| 1185 | CLCN6 | 0.000259 | 0.506676 | 0.443854 | 0.390698 | 0.447076 | down |
| 84498 | FAM120B | 0.000259 | 0.537173 | 0.442628 | 0.44327 | 0.474357 | down |
| 10367 | MICU1 | 0.000259 | 0.701037 | 0.626629 | 0.618021 | 0.648562 | down |
| 9873 | FCHSD2 | 0.000259 | 0.546303 | 0.650371 | 0.53018 | 0.575618 | down |
| 311 | ANXA11 | 0.00026 | 0.687771 | 0.580348 | 0.640136 | 0.636085 | down |
| 9187 | SLC24A1 | 0.00026 | 0.502191 | 0.380068 | 0.366478 | 0.416246 | down |
| 26986 | PABPC1 | 0.00026 | 0.640011 | 0.679888 | 0.641704 | 0.653868 | down |
| 64744 | SMAP2 | 0.00026 | 0.614989 | 0.587372 | 0.699342 | 0.633901 | down |
| 5709 | PSMD3 | 0.00026 | 0.630678 | 0.582985 | 0.598192 | 0.603951 | down |
| 11129 | CLASRP | 0.00026 | 0.631537 | 0.587415 | 0.549114 | 0.589355 | down |
| 1184 | CLCN5 | 0.000261 | 0.449846 | 0.425305 | 0.471475 | 0.448875 | down |
| 85476 | GFM1 | 0.000261 | 0.485905 | 0.595491 | 0.573019 | 0.551472 | down |
| 58506 | SCAF1 | 0.000261 | 0.645042 | 0.597434 | 0.517159 | 0.586545 | down |
| 10762 | NUP50 | 0.000261 | 0.54018 | 0.630746 | 0.582991 | 0.584639 | down |
| 9644 | SH3PXD2A | 0.000261 | 0.459015 | 0.537335 | 0.48512 | 0.493824 | down |
| 27340 | UTP20 | 0.000262 | 0.319009 | 0.387691 | 0.396753 | 0.367818 | down |
| 81609 | SNX27 | 0.000262 | 0.587485 | 0.521776 | 0.608112 | 0.572458 | down |
| 6792 | CDKL5 | 0.000262 | 0.34359 | 0.347473 | 0.341891 | 0.344318 | down |
| 9656 | MDC1 | 0.000262 | 0.47245 | 0.409531 | 0.454343 | 0.445441 | down |
| 9880 | ZBTB39 | 0.000262 | 0.482814 | 0.527051 | 0.463056 | 0.490974 | down |
| 22838 | RNF44 | 0.000263 | 0.555033 | 0.48315 | 0.584883 | 0.541022 | down |
| 10745 | PHTF1 | 0.000263 | 0.616765 | 0.594864 | 0.705491 | 0.63904 | down |
| 9545 | RAB3D | 0.000264 | 0.494604 | 0.543784 | 0.487481 | 0.508623 | down |
| 4089 | SMAD4 | 0.000264 | 0.613628 | 0.556191 | 0.641135 | 0.603651 | down |
| 221477 | C6orf89 | 0.000264 | 0.554228 | 0.61592 | 0.622227 | 0.597459 | down |
| 5817 | PVR | 0.000265 | 0.53446 | 0.566826 | 0.48248 | 0.527922 | down |
| 51667 | NUB1 | 0.000265 | 0.682043 | 0.581082 | 0.675885 | 0.646337 | down |
| 64900 | LPIN3 | 0.000266 | 0.490287 | 0.621106 | 0.560285 | 0.557226 | down |
| 3984 | LIMK1 | 0.000267 | 0.647356 | 0.573664 | 0.555648 | 0.592223 | down |
| 4820 | NKTR | 0.000268 | 0.432849 | 0.485679 | 0.472972 | 0.463833 | down |
| 79776 | ZFHX4 | 0.000268 | 0.507752 | 0.524484 | 0.447607 | 0.493281 | down |
| 63898 | SH2D4A | 0.000269 | 0.602496 | 0.530669 | 0.629668 | 0.587611 | down |
| 23326 | USP22 | 0.000269 | 0.439131 | 0.518152 | 0.45852 | 0.471934 | down |
| 9619 | ABCG1 | 0.00027 | 0.454154 | 0.541176 | 0.379285 | 0.458205 | down |
| 84273 | NOA1 | 0.00027 | 0.620936 | 0.675459 | 0.570944 | 0.622446 | down |
| 84859 | LRCH3 | 0.00027 | 0.472813 | 0.479482 | 0.442968 | 0.465088 | down |
| 10409 | BASP1 | 0.00027 | 0.630009 | 0.594036 | 0.557924 | 0.59399 | down |
| 55388 | MCM10 | 0.00027 | 0.403361 | 0.505362 | 0.401184 | 0.436636 | down |
| 1183 | CLCN4 | 0.000271 | 0.513525 | 0.43496 | 0.430675 | 0.45972 | down |
| 85378 | TUBGCP6 | 0.000271 | 0.518175 | 0.596877 | 0.588319 | 0.56779 | down |
| 55249 | YY1AP1 | 0.000271 | 0.540896 | 0.469385 | 0.572476 | 0.527586 | down |
| 5481 | PPID | 0.000271 | 0.614201 | 0.668105 | 0.574114 | 0.618807 | down |
| 5989 | RFX1 | 0.000271 | 0.545911 | 0.542373 | 0.4343 | 0.507528 | down |
| 286077 | FAM83H | 0.000272 | 0.584361 | 0.547547 | 0.496289 | 0.542732 | down |
| 3685 | ITGAV | 0.000272 | 0.604386 | 0.670001 | 0.638386 | 0.637591 | down |
| 80232 | WDR26 | 0.000272 | 0.460466 | 0.428552 | 0.453267 | 0.447428 | down |
| 9314 | KLF4 | 0.000272 | 0.623456 | 0.553594 | 0.559111 | 0.57872 | down |
| 80216 | ALPK1 | 0.000274 | 0.614379 | 0.653325 | 0.628585 | 0.632096 | down |
| 57460 | PPM1H | 0.000274 | 0.615733 | 0.633816 | 0.539084 | 0.596211 | down |
| 10630 | PDPN | 0.000274 | 0.497306 | 0.490021 | 0.587262 | 0.524863 | down |
| 6764 | ST5 | 0.000275 | 0.545234 | 0.563645 | 0.636111 | 0.581663 | down |
| 79628 | SH3TC2 | 0.000276 | 0.579832 | 0.549108 | 0.55235 | 0.56043 | down |

| | | | | | | | |
|---|---|---|---|---|---|---|---|
| 10195 | ALG3 | 0.000277 | 0.625252 | 0.708602 | 0.625877 | 0.653243 | down |
| 8527 | DGKD | 0.000278 | 0.482857 | 0.543208 | 0.429988 | 0.485351 | down |
| 9411 | ARHGAP29 | 0.000278 | 0.622925 | 0.669601 | 0.700295 | 0.664274 | down |
| 79567 | RIPOR1 | 0.000278 | 0.630257 | 0.483197 | 0.536816 | 0.55009 | down |
| 9697 | TRAM2 | 0.000278 | 0.495371 | 0.440406 | 0.43693 | 0.457569 | down |
| 27336 | HTATSF1 | 0.000278 | 0.546418 | 0.63446 | 0.613151 | 0.59801 | down |
| 63967 | CLSPN | 0.000278 | 0.5517 | 0.67926 | 0.607193 | 0.612718 | down |
| 10166 | SLC25A15 | 0.000279 | 0.703758 | 0.611449 | 0.654346 | 0.656517 | down |
| 57154 | SMURF1 | 0.000279 | 0.475682 | 0.412761 | 0.500951 | 0.463131 | down |
| 9742 | IFT140 | 0.000279 | 0.492303 | 0.570491 | 0.461084 | 0.507959 | down |
| 114876 | OSBPL1A | 0.00028 | 0.555491 | 0.626945 | 0.646024 | 0.609487 | down |
| 481 | ATP1B1 | 0.00028 | 0.584559 | 0.680057 | 0.639999 | 0.634872 | down |
| 9329 | GTF3C4 | 0.000281 | 0.482631 | 0.581119 | 0.541127 | 0.534959 | down |
| 642273 | FAM110C | 0.000281 | 0.591869 | 0.512082 | 0.480781 | 0.528244 | down |
| 6002 | RGS12 | 0.000281 | 0.620634 | 0.673743 | 0.697129 | 0.663835 | down |
| 7617 | ZNF66 | 0.000281 | 0.434838 | 0.44719 | 0.519463 | 0.467164 | down |
| 55654 | TMEM127 | 0.000282 | 0.542663 | 0.581816 | 0.654557 | 0.593012 | down |
| 58489 | ABHD17C | 0.000282 | 0.527859 | 0.546073 | 0.518155 | 0.530696 | down |
| 22906 | TRAK1 | 0.000283 | 0.49077 | 0.57971 | 0.514242 | 0.528241 | down |
| 57470 | LRRC47 | 0.000283 | 0.48416 | 0.580417 | 0.485966 | 0.516848 | down |
| 51594 | NBAS | 0.000284 | 0.455794 | 0.530756 | 0.524421 | 0.503657 | down |
| 57630 | SH3RF1 | 0.000284 | 0.562236 | 0.50888 | 0.551657 | 0.540924 | down |
| 643680 | MS4A4E | 0.000284 | 0.712825 | 0.601783 | 0.6164 | 0.643669 | down |
| 57522 | SRGAP1 | 0.000284 | 0.474608 | 0.417103 | 0.479833 | 0.457181 | down |
| 59343 | SENP2 | 0.000285 | 0.497247 | 0.527883 | 0.509771 | 0.511634 | down |
| 197258 | FUK | 0.000285 | 0.643613 | 0.603967 | 0.649686 | 0.632422 | down |
| 26524 | LATS2 | 0.000285 | 0.539822 | 0.490114 | 0.452772 | 0.494236 | down |
| 255743 | NPNT | 0.000285 | 0.456687 | 0.532879 | 0.49222 | 0.493929 | down |
| 132660 | LIN54 | 0.000285 | 0.553542 | 0.543638 | 0.539127 | 0.545436 | down |
| 9889 | ZBED4 | 0.000285 | 0.518944 | 0.439576 | 0.424567 | 0.461029 | down |
| 55914 | ERBIN | 0.000285 | 0.631792 | 0.708492 | 0.629904 | 0.656729 | down |
| 9620 | CELSR1 | 0.000285 | 0.536862 | 0.49443 | 0.470949 | 0.500747 | down |
| 10066 | SCAMP2 | 0.000286 | 0.650297 | 0.572403 | 0.589808 | 0.604169 | down |
| 84255 | SLC37A3 | 0.000286 | 0.596547 | 0.520613 | 0.656892 | 0.591351 | down |
| 84164 | ASCC2 | 0.000287 | 0.581217 | 0.52413 | 0.537946 | 0.547764 | down |
| 4302 | MLLT6 | 0.000287 | 0.571052 | 0.518817 | 0.619535 | 0.569801 | down |
| 59274 | TLNRD1 | 0.000288 | 0.690696 | 0.693475 | 0.615517 | 0.666563 | down |
| 5187 | PER1 | 0.000289 | 0.639505 | 0.643098 | 0.527975 | 0.603526 | down |
| 3725 | JUN | 0.000289 | 0.616684 | 0.615513 | 0.546134 | 0.592777 | down |
| 5337 | PLD1 | 0.00029 | 0.5024 | 0.471032 | 0.558358 | 0.510597 | down |
| 2181 | ACSL3 | 0.00029 | 0.561184 | 0.642644 | 0.546274 | 0.583367 | down |
| 140735 | DYNLL2 | 0.00029 | 0.580123 | 0.631193 | 0.613183 | 0.608166 | down |
| 1287 | COL4A5 | 0.000291 | 0.463451 | 0.529907 | 0.498391 | 0.49725 | down |
| 9905 | SGSM2 | 0.000291 | 0.641575 | 0.640049 | 0.596216 | 0.625947 | down |
| 340527 | NHSL2 | 0.000292 | 0.351297 | 0.316918 | 0.368511 | 0.345575 | down |
| 22862 | FNDC3A | 0.000292 | 0.553943 | 0.589182 | 0.642826 | 0.595317 | down |
| 59 | ACTA2 | 0.000293 | 0.712668 | 0.636719 | 0.649307 | 0.666231 | down |
| 116843 | SLC18B1 | 0.000295 | 0.622021 | 0.631534 | 0.6669 | 0.640152 | down |
| 57567 | ZNF319 | 0.000295 | 0.684321 | 0.643586 | 0.603737 | 0.643881 | down |
| 80335 | WDR82 | 0.000296 | 0.538046 | 0.577073 | 0.601 | 0.57204 | down |
| 4163 | MCC | 0.000296 | 0.43643 | 0.528014 | 0.461804 | 0.475416 | down |
| 9639 | ARHGEF10 | 0.000297 | 0.44008 | 0.538588 | 0.46043 | 0.479699 | down |
| 55625 | ZDHHC7 | 0.000297 | 0.518758 | 0.512682 | 0.417373 | 0.482937 | down |
| 2633 | GBP1 | 0.000297 | 0.700147 | 0.661393 | 0.61872 | 0.660087 | down |
| 4055 | LTBR | 0.000297 | 0.700454 | 0.614951 | 0.623705 | 0.64637 | down |
| 1656 | DDX6 | 0.000297 | 0.391843 | 0.471505 | 0.464115 | 0.442488 | down |
| 5783 | PTPN13 | 0.000298 | 0.411297 | 0.524372 | 0.481258 | 0.472309 | down |
| 9825 | SPATA2 | 0.000298 | 0.613879 | 0.647023 | 0.550528 | 0.60381 | down |

| | | | | | | | |
|---|---|---|---|---|---|---|---|
| 5641 | LGMN | 0.000299 | 0.621429 | 0.721962 | 0.646978 | 0.663456 | down |
| 1.01E+08 | OCLN | 0.0003 | 0.477171 | 0.511307 | 0.416424 | 0.468301 | down |
| 23112 | TNRC6B | 0.000301 | 0.382782 | 0.452505 | 0.523455 | 0.452914 | down |
| 9334 | B4GALT5 | 0.000301 | 0.399365 | 0.476317 | 0.463396 | 0.44636 | down |
| 3508 | IGHMBP2 | 0.000301 | 0.615086 | 0.61036 | 0.517021 | 0.580822 | down |
| 9064 | MAP3K6 | 0.000302 | 0.579087 | 0.466657 | 0.483218 | 0.509654 | down |
| 8322 | FZD4 | 0.000303 | 0.433898 | 0.39189 | 0.485449 | 0.437079 | down |
| 51585 | PCF11 | 0.000304 | 0.518877 | 0.541752 | 0.629803 | 0.563478 | down |
| 3693 | ITGB5 | 0.000304 | 0.483303 | 0.570256 | 0.472547 | 0.508702 | down |
| 5914 | RARA | 0.000304 | 0.598868 | 0.514607 | 0.494625 | 0.536033 | down |
| 6198 | RPS6KB1 | 0.000304 | 0.476175 | 0.581228 | 0.50181 | 0.519737 | down |
| 6867 | TACC1 | 0.000304 | 0.454041 | 0.432432 | 0.523132 | 0.469868 | down |
| 7405 | UVRAG | 0.000304 | 0.434082 | 0.474984 | 0.465517 | 0.458194 | down |
| 23291 | FBXW11 | 0.000305 | 0.581151 | 0.582489 | 0.683308 | 0.61565 | down |
| 57186 | RALGAPA2 | 0.000305 | 0.303211 | 0.400597 | 0.339806 | 0.347871 | down |
| 79922 | MRM1 | 0.000305 | 0.638023 | 0.604241 | 0.708446 | 0.650237 | down |
| 54552 | GNL3L | 0.000305 | 0.532892 | 0.596489 | 0.60257 | 0.577317 | down |
| 9306 | SOCS6 | 0.000305 | 0.673768 | 0.692087 | 0.632296 | 0.66605 | down |
| 91746 | YTHDC1 | 0.000306 | 0.538305 | 0.627854 | 0.52501 | 0.563723 | down |
| 6429 | SRSF4 | 0.000306 | 0.49665 | 0.554154 | 0.511384 | 0.52073 | down |
| 80320 | SP6 | 0.000306 | 0.597737 | 0.513758 | 0.544423 | 0.551973 | down |
| 138199 | CARNMT1 | 0.000306 | 0.558349 | 0.557522 | 0.59828 | 0.571384 | down |
| 79672 | FN3KRP | 0.000307 | 0.617438 | 0.556702 | 0.536798 | 0.570313 | down |
| 11124 | FAF1 | 0.000307 | 0.541619 | 0.591025 | 0.676172 | 0.602938 | down |
| 9480 | ONECUT2 | 0.000307 | 0.342491 | 0.34375 | 0.3344 | 0.340214 | down |
| 152002 | XXYLT1 | 0.000307 | 0.588357 | 0.655406 | 0.61622 | 0.619994 | down |
| 55971 | BAIAP2L1 | 0.000308 | 0.584834 | 0.57705 | 0.646749 | 0.602878 | down |
| 9044 | BTAF1 | 0.000308 | 0.416931 | 0.497699 | 0.493062 | 0.46923 | down |
| 1808 | DPYSL2 | 0.000308 | 0.397845 | 0.509155 | 0.449483 | 0.452161 | down |
| 29994 | BAZ2B | 0.000308 | 0.556861 | 0.493016 | 0.581709 | 0.543862 | down |
| 3267 | AGFG1 | 0.000309 | 0.589796 | 0.680085 | 0.626132 | 0.632004 | down |
| 1845 | DUSP3 | 0.000309 | 0.614907 | 0.688547 | 0.695916 | 0.666457 | down |
| 10184 | LHFPL2 | 0.000309 | 0.386407 | 0.421082 | 0.52547 | 0.44432 | down |
| 84622 | ZNF594 | 0.00031 | 0.492103 | 0.540607 | 0.506148 | 0.512952 | down |
| 149013 | NBPF12 | 0.00031 | 0.38424 | 0.342751 | 0.494069 | 0.40702 | down |
| 23395 | LARS2 | 0.00031 | 0.564224 | 0.565853 | 0.569483 | 0.56652 | down |
| 123606 | NIPA1 | 0.00031 | 0.499724 | 0.576086 | 0.561957 | 0.545922 | down |
| 6507 | SLC1A3 | 0.000312 | 0.569184 | 0.586281 | 0.656448 | 0.603971 | down |
| 84444 | DOT1L | 0.000312 | 0.641388 | 0.557253 | 0.556471 | 0.585037 | down |
| 80351 | TNKS2 | 0.000312 | 0.545864 | 0.610791 | 0.637227 | 0.59796 | down |
| 122786 | FRMD6 | 0.000313 | 0.503858 | 0.611588 | 0.548956 | 0.554801 | down |
| 57647 | DHX37 | 0.000313 | 0.561392 | 0.563223 | 0.49809 | 0.540902 | down |
| 84632 | AFAP1L2 | 0.000313 | 0.457821 | 0.544923 | 0.603732 | 0.535492 | down |
| 83855 | KLF16 | 0.000314 | 0.655561 | 0.586834 | 0.623875 | 0.62209 | down |
| 5317 | PKP1 | 0.000314 | 0.674915 | 0.556348 | 0.544356 | 0.591873 | down |
| 51104 | ABHD17B | 0.000315 | 0.561135 | 0.623541 | 0.514002 | 0.566226 | down |
| 23522 | KAT6B | 0.000315 | 0.301003 | 0.360991 | 0.479688 | 0.380561 | down |
| 22887 | FOXJ3 | 0.000315 | 0.457878 | 0.544612 | 0.540297 | 0.514263 | down |
| 79801 | SHCBP1 | 0.000316 | 0.543148 | 0.601662 | 0.634338 | 0.593049 | down |
| 4791 | NFKB2 | 0.000316 | 0.490675 | 0.605286 | 0.501311 | 0.532424 | down |
| 9139 | CBFA2T2 | 0.000316 | 0.457002 | 0.588608 | 0.536486 | 0.527365 | down |
| 57089 | ENTPD7 | 0.000316 | 0.499506 | 0.493036 | 0.566763 | 0.519768 | down |
| 4286 | MITF | 0.000317 | 0.407043 | 0.507813 | 0.558295 | 0.49105 | down |
| 23229 | ARHGEF9 | 0.000317 | 0.671391 | 0.638298 | 0.649741 | 0.653143 | down |
| 253143 | PRR14L | 0.000317 | 0.319034 | 0.330802 | 0.306768 | 0.318868 | down |
| 18 | ABAT | 0.000317 | 0.448434 | 0.499738 | 0.330827 | 0.426333 | down |
| 56987 | BBX | 0.000318 | 0.443627 | 0.530693 | 0.502811 | 0.492377 | down |
| 55748 | CNDP2 | 0.000318 | 0.602005 | 0.517795 | 0.557008 | 0.558936 | down |

| | | | | | | | |
|---|---|---|---|---|---|---|---|
| 5903 | RANBP2 | 0.000319 | 0.610914 | 0.639223 | 0.628846 | 0.626328 | down |
| 155061 | ZNF746 | 0.000319 | 0.640969 | 0.528598 | 0.501491 | 0.557019 | down |
| 57834 | CYP4F11 | 0.000321 | 0.52046 | 0.572267 | 0.637675 | 0.576801 | down |
| 55238 | SLC38A7 | 0.000321 | 0.572598 | 0.553398 | 0.589448 | 0.571814 | down |
| 202 | CRYBG1 | 0.000321 | 0.36756 | 0.407318 | 0.467601 | 0.41416 | down |
| 85406 | DNAJC14 | 0.000322 | 0.544479 | 0.651155 | 0.54771 | 0.581115 | down |
| 79651 | RHBDF2 | 0.000323 | 0.671767 | 0.562101 | 0.535387 | 0.589752 | down |
| 23034 | SAMD4A | 0.000324 | 0.473447 | 0.516293 | 0.418534 | 0.469425 | down |
| 1.03E+08 | LOC10272 | 0.000326 | 0.506127 | 0.595498 | 0.608989 | 0.570205 | down |
| 57698 | SHTN1 | 0.000326 | 0.418716 | 0.456372 | 0.553477 | 0.476188 | down |
| 79591 | ARMH3 | 0.000326 | 0.555319 | 0.528324 | 0.627765 | 0.570469 | down |
| 27161 | AGO2 | 0.000327 | 0.41923 | 0.471052 | 0.46942 | 0.453234 | down |
| 3423 | IDS | 0.000327 | 0.580121 | 0.604416 | 0.658958 | 0.614498 | down |
| 9053 | MAP7 | 0.000328 | 0.552344 | 0.441337 | 0.591672 | 0.528451 | down |
| 5193 | PEX12 | 0.000328 | 0.510525 | 0.465553 | 0.507347 | 0.494475 | down |
| 80014 | WWC2 | 0.000328 | 0.423641 | 0.445239 | 0.564817 | 0.477899 | down |
| 125919 | ZNF543 | 0.000329 | 0.427497 | 0.527976 | 0.491293 | 0.482255 | down |
| 8737 | RIPK1 | 0.000329 | 0.41006 | 0.525535 | 0.535687 | 0.490427 | down |
| 1606 | DGKA | 0.00033 | 0.590889 | 0.498323 | 0.581161 | 0.556791 | down |
| 2539 | G6PD | 0.00033 | 0.684993 | 0.581855 | 0.583428 | 0.616759 | down |
| 5603 | MAPK13 | 0.000331 | 0.632575 | 0.698593 | 0.648169 | 0.659779 | down |
| 84629 | TNRC18 | 0.000331 | 0.536382 | 0.424696 | 0.453815 | 0.471631 | down |
| 10479 | SLC9A6 | 0.000331 | 0.516074 | 0.594011 | 0.561139 | 0.557075 | down |
| 91010 | FMNL3 | 0.000332 | 0.56381 | 0.518841 | 0.435419 | 0.506023 | down |
| 9282 | MED14 | 0.000332 | 0.41762 | 0.464904 | 0.506009 | 0.462844 | down |
| 6497 | SKI | 0.000334 | 0.531773 | 0.459101 | 0.451969 | 0.480948 | down |
| 51322 | WAC | 0.000334 | 0.518953 | 0.588376 | 0.6108 | 0.57271 | down |
| 6601 | SMARCC2 | 0.000335 | 0.520141 | 0.375312 | 0.449002 | 0.448151 | down |
| 3638 | INSIG1 | 0.000335 | 0.649413 | 0.592411 | 0.582859 | 0.608228 | down |
| 79944 | L2HGDH | 0.000335 | 0.58089 | 0.638673 | 0.660852 | 0.626805 | down |
| 83903 | HASPIN | 0.000336 | 0.542443 | 0.549609 | 0.47695 | 0.523001 | down |
| 81559 | TRIM11 | 0.000336 | 0.645025 | 0.563398 | 0.59543 | 0.601284 | down |
| 83860 | TAF3 | 0.000338 | 0.520658 | 0.508873 | 0.413792 | 0.481108 | down |
| 27154 | BRPF3 | 0.000338 | 0.451752 | 0.441634 | 0.574229 | 0.489205 | down |
| 10007 | GNPDA1 | 0.000338 | 0.633077 | 0.636953 | 0.714548 | 0.661526 | down |
| 5442 | POLRMT | 0.000338 | 0.675207 | 0.634894 | 0.681613 | 0.663904 | down |
| 56897 | WRNIP1 | 0.000338 | 0.598215 | 0.650868 | 0.579546 | 0.609543 | down |
| 26122 | EPC2 | 0.000339 | 0.484008 | 0.610215 | 0.51488 | 0.536368 | down |
| 387893 | KMT5A | 0.000339 | 0.695005 | 0.606557 | 0.585657 | 0.629073 | down |
| 10188 | TNK2 | 0.000339 | 0.670944 | 0.65678 | 0.590535 | 0.63942 | down |
| 195828 | ZNF367 | 0.000339 | 0.518909 | 0.587075 | 0.517503 | 0.541162 | down |
| 388115 | CCDC9B | 0.000339 | 0.471415 | 0.450993 | 0.494749 | 0.472386 | down |
| 642475 | MROH6 | 0.000339 | 0.653848 | 0.690353 | 0.585463 | 0.643221 | down |
| 1362 | CPD | 0.00034 | 0.521731 | 0.613887 | 0.613767 | 0.583128 | down |
| 115509 | ZNF689 | 0.000341 | 0.693155 | 0.693665 | 0.597638 | 0.661486 | down |
| 200424 | TET3 | 0.000341 | 0.338462 | 0.399906 | 0.3807 | 0.373022 | down |
| 4335 | MNT | 0.000341 | 0.547011 | 0.520444 | 0.510154 | 0.52587 | down |
| 30850 | CDR2L | 0.000341 | 0.512002 | 0.593738 | 0.555649 | 0.553796 | down |
| 9612 | NCOR2 | 0.000342 | 0.560116 | 0.456524 | 0.498778 | 0.505139 | down |
| 768 | CA9 | 0.000342 | 0.659407 | 0.563897 | 0.52694 | 0.583414 | down |
| 3656 | IRAK2 | 0.000342 | 0.43336 | 0.55891 | 0.446532 | 0.479601 | down |
| 84953 | MICALCL | 0.000343 | 0.525041 | 0.504668 | 0.517766 | 0.515825 | down |
| 5576 | PRKAR2A | 0.000343 | 0.560972 | 0.668177 | 0.605826 | 0.611658 | down |
| 26093 | CCDC9 | 0.000343 | 0.55967 | 0.407713 | 0.510262 | 0.492548 | down |
| 2145 | EZH1 | 0.000343 | 0.405716 | 0.48311 | 0.465651 | 0.451492 | down |
| 1021 | CDK6 | 0.000343 | 0.378181 | 0.494993 | 0.440977 | 0.43805 | down |
| 131544 | CRYBG3 | 0.000343 | 0.324705 | 0.38053 | 0.379464 | 0.361566 | down |
| 54507 | ADAMTSL4 | 0.000344 | 0.644935 | 0.589411 | 0.550863 | 0.59507 | down |

| | | | | | | | |
|---|---|---|---|---|---|---|---|
| 55904 | KMT2E | 0.000345 | 0.395024 | 0.494503 | 0.47412 | 0.454549 | down |
| 346389 | MACC1 | 0.000346 | 0.534393 | 0.53648 | 0.603346 | 0.558073 | down |
| 8412 | BCAR3 | 0.000346 | 0.543191 | 0.590303 | 0.501037 | 0.544843 | down |
| 10061 | ABCF2 | 0.000347 | 0.585709 | 0.684436 | 0.577134 | 0.61576 | down |
| 2012 | EMP1 | 0.000347 | 0.588361 | 0.555877 | 0.570882 | 0.571707 | down |
| 57706 | DENND1A | 0.000348 | 0.516117 | 0.505884 | 0.544226 | 0.522076 | down |
| 55324 | ABCF3 | 0.000348 | 0.582906 | 0.623836 | 0.671483 | 0.626075 | down |
| 57448 | BIRC6 | 0.000349 | 0.405212 | 0.454358 | 0.405518 | 0.421696 | down |
| 54464 | XRN1 | 0.00035 | 0.624985 | 0.60394 | 0.652277 | 0.627067 | down |
| 57587 | CFAP97 | 0.00035 | 0.488591 | 0.58311 | 0.617119 | 0.56294 | down |
| 9759 | HDAC4 | 0.00035 | 0.538468 | 0.443987 | 0.421984 | 0.468146 | down |
| 116115 | ZNF526 | 0.00035 | 0.650055 | 0.654816 | 0.579036 | 0.627969 | down |
| 85451 | UNK | 0.00035 | 0.598087 | 0.682221 | 0.609354 | 0.629887 | down |
| 124997 | WDR81 | 0.000351 | 0.549842 | 0.605723 | 0.546476 | 0.567347 | down |
| 25959 | KANK2 | 0.000351 | 0.519169 | 0.464611 | 0.456204 | 0.479995 | down |
| 2906 | GRIN2D | 0.000351 | 0.470163 | 0.616336 | 0.559211 | 0.54857 | down |
| 57688 | ZSWIM6 | 0.000352 | 0.527955 | 0.461944 | 0.543216 | 0.511038 | down |
| 80020 | FOXRED2 | 0.000352 | 0.569085 | 0.524172 | 0.587874 | 0.560377 | down |
| 4303 | FOXO4 | 0.000352 | 0.533144 | 0.5629 | 0.538044 | 0.544696 | down |
| 5209 | PFKFB3 | 0.000353 | 0.50485 | 0.472718 | 0.466418 | 0.481329 | down |
| 144501 | KRT80 | 0.000353 | 0.563416 | 0.457825 | 0.583079 | 0.534774 | down |
| 6477 | SIAH1 | 0.000354 | 0.557777 | 0.650065 | 0.636372 | 0.614738 | down |
| 57532 | NUFIP2 | 0.000354 | 0.408186 | 0.52568 | 0.461741 | 0.465202 | down |
| 54893 | MTMR10 | 0.000354 | 0.45368 | 0.422277 | 0.466571 | 0.447509 | down |
| 3799 | KIF5B | 0.000354 | 0.541472 | 0.618398 | 0.632817 | 0.597563 | down |
| 9923 | ZBTB40 | 0.000355 | 0.448202 | 0.46757 | 0.471867 | 0.462546 | down |
| 64848 | YTHDC2 | 0.000356 | 0.512811 | 0.614459 | 0.514058 | 0.547109 | down |
| 5495 | PPM1B | 0.000356 | 0.511593 | 0.52022 | 0.486796 | 0.506203 | down |
| 9213 | XPR1 | 0.000356 | 0.580117 | 0.645979 | 0.65342 | 0.626505 | down |
| 10123 | ARL4C | 0.000358 | 0.591591 | 0.497395 | 0.482546 | 0.523844 | down |
| 389333 | PROB1 | 0.000358 | 0.718057 | 0.612103 | 0.608228 | 0.646129 | down |
| 58487 | CREBZF | 0.000358 | 0.579299 | 0.50635 | 0.553667 | 0.546439 | down |
| 30000 | TNPO2 | 0.000359 | 0.679227 | 0.625358 | 0.56336 | 0.622648 | down |
| 56122 | PCDHB14 | 0.000359 | 0.346163 | 0.41858 | 0.422577 | 0.395773 | down |
| 9344 | TAOK2 | 0.00036 | 0.530852 | 0.479363 | 0.536893 | 0.515702 | down |
| 201562 | HACD2 | 0.00036 | 0.576207 | 0.647142 | 0.626086 | 0.616478 | down |
| 121551 | BTBD11 | 0.000361 | 0.600157 | 0.601882 | 0.657409 | 0.619816 | down |
| 23224 | SYNE2 | 0.000362 | 0.312409 | 0.346798 | 0.407612 | 0.355607 | down |
| 8379 | MAD1L1 | 0.000362 | 0.539341 | 0.520099 | 0.531484 | 0.530308 | down |
| 760 | CA2 | 0.000362 | 0.670062 | 0.655722 | 0.646784 | 0.657522 | down |
| 10505 | SEMA4F | 0.000362 | 0.609841 | 0.686142 | 0.567998 | 0.621327 | down |
| 5724 | PTAFR | 0.000363 | 0.645191 | 0.687335 | 0.597243 | 0.643256 | down |
| 259173 | ALS2CL | 0.000366 | 0.686024 | 0.63312 | 0.650343 | 0.656496 | down |
| 994 | CDC25B | 0.000366 | 0.572175 | 0.544345 | 0.488821 | 0.535113 | down |
| 4059 | BCAM | 0.000366 | 0.716043 | 0.668195 | 0.605243 | 0.663161 | down |
| 4123 | MAN2C1 | 0.000367 | 0.652317 | 0.591367 | 0.560589 | 0.601424 | down |
| 64285 | RHBDF1 | 0.000368 | 0.590918 | 0.554661 | 0.506333 | 0.550637 | down |
| 540 | ATP7B | 0.000368 | 0.37264 | 0.310778 | 0.46526 | 0.382893 | down |
| 79147 | FKRP | 0.000369 | 0.544168 | 0.599822 | 0.61266 | 0.58555 | down |
| 23189 | KANK1 | 0.000369 | 0.451655 | 0.454849 | 0.513919 | 0.473474 | down |
| 6773 | STAT2 | 0.00037 | 0.346729 | 0.438926 | 0.348851 | 0.378169 | down |
| 1855 | DVL1 | 0.00037 | 0.652298 | 0.621393 | 0.647644 | 0.640445 | down |
| 84874 | ZNF514 | 0.00037 | 0.537717 | 0.610526 | 0.573277 | 0.57384 | down |
| 83879 | CDCA7 | 0.000371 | 0.63897 | 0.561763 | 0.662333 | 0.621022 | down |
| 9757 | KMT2B | 0.000371 | 0.553265 | 0.49325 | 0.605264 | 0.550593 | down |
| 9439 | MED23 | 0.000372 | 0.516759 | 0.416291 | 0.390765 | 0.441272 | down |
| 64864 | RFX7 | 0.000372 | 0.529756 | 0.503488 | 0.485383 | 0.506209 | down |
| 83692 | CD99L2 | 0.000372 | 0.647108 | 0.599348 | 0.546127 | 0.597528 | down |

| | | | | | | | |
|---|---|---|---|---|---|---|---|
| 83394 | PITPNM3 | 0.000373 | 0.509931 | 0.455555 | 0.508133 | 0.491206 | down |
| 23195 | MDN1 | 0.000373 | 0.44057 | 0.356969 | 0.371723 | 0.389754 | down |
| 7586 | ZKSCAN1 | 0.000374 | 0.428946 | 0.457721 | 0.428299 | 0.438322 | down |
| 54414 | SIAE | 0.000375 | 0.692096 | 0.635651 | 0.659951 | 0.662566 | down |
| 9894 | TELO2 | 0.000375 | 0.587514 | 0.631862 | 0.558744 | 0.592707 | down |
| 51514 | DTL | 0.000375 | 0.517759 | 0.635492 | 0.642582 | 0.598611 | down |
| 6645 | SNTB2 | 0.000376 | 0.557014 | 0.649064 | 0.613366 | 0.606481 | down |
| 727910 | TLCD2 | 0.000376 | 0.525305 | 0.604861 | 0.556116 | 0.562094 | down |
| 580 | BARD1 | 0.000376 | 0.596204 | 0.724235 | 0.649047 | 0.656495 | down |
| 55691 | FRMD4A | 0.000377 | 0.506458 | 0.523246 | 0.391055 | 0.473586 | down |
| 56123 | PCDHB13 | 0.000377 | 0.506182 | 0.476205 | 0.355601 | 0.445996 | down |
| 51559 | NT5DC3 | 0.000377 | 0.57177 | 0.546971 | 0.426641 | 0.515127 | down |
| 5420 | PODXL | 0.000378 | 0.399578 | 0.457976 | 0.480819 | 0.446124 | down |
| 2342 | FNTB | 0.000378 | 0.267096 | 0.142639 | 0.261511 | 0.223749 | down |
| 353116 | RILPL1 | 0.00038 | 0.510211 | 0.479703 | 0.636752 | 0.542222 | down |
| 7571 | ZNF23 | 0.000381 | 0.648187 | 0.618205 | 0.560839 | 0.609077 | down |
| 8506 | CNTNAP1 | 0.000381 | 0.373425 | 0.433463 | 0.400538 | 0.402475 | down |
| 5094 | PCBP2 | 0.000382 | 0.704064 | 0.659731 | 0.618085 | 0.660627 | down |
| 7536 | SF1 | 0.000382 | 0.650991 | 0.520825 | 0.56694 | 0.579586 | down |
| 91 | ACVR1B | 0.000382 | 0.535706 | 0.611847 | 0.491453 | 0.546335 | down |
| 157 | GRK3 | 0.000382 | 0.593212 | 0.708449 | 0.586346 | 0.629336 | down |
| 9240 | PNMA1 | 0.000383 | 0.538693 | 0.589271 | 0.520327 | 0.54943 | down |
| 9169 | SCAF11 | 0.000384 | 0.591969 | 0.637581 | 0.569783 | 0.599778 | down |
| 27094 | KCNMB3 | 0.000384 | 0.390147 | 0.394534 | 0.31483 | 0.366503 | down |
| 55421 | NCBP3 | 0.000385 | 0.549584 | 0.607814 | 0.630576 | 0.595991 | down |
| 8851 | CDK5R1 | 0.000386 | 0.494181 | 0.52656 | 0.456348 | 0.492363 | down |
| 8867 | SYNJ1 | 0.000388 | 0.506453 | 0.479914 | 0.548002 | 0.511456 | down |
| 91862 | MARVELD3 | 0.00039 | 0.623834 | 0.635406 | 0.613381 | 0.624207 | down |
| 54020 | SLC37A1 | 0.000391 | 0.545333 | 0.604143 | 0.576932 | 0.575469 | down |
| 1662 | DDX10 | 0.000391 | 0.587636 | 0.594769 | 0.653196 | 0.611867 | down |
| 5089 | PBX2 | 0.000392 | 0.536807 | 0.524606 | 0.602318 | 0.554577 | down |
| 1.08E+08 | LOC10798 | 0.000393 | 0.422251 | 0.340431 | 0.527998 | 0.430227 | down |
| 50618 | ITSN2 | 0.000393 | 0.518453 | 0.604544 | 0.527489 | 0.550162 | down |
| 4236 | MFAP1 | 0.000394 | 0.699169 | 0.594031 | 0.589544 | 0.627581 | down |
| 1856 | DVL2 | 0.000396 | 0.597401 | 0.567725 | 0.672726 | 0.612617 | down |
| 8974 | P4HA2 | 0.000396 | 0.657516 | 0.606541 | 0.58075 | 0.614936 | down |
| 114825 | PWWP2A | 0.000397 | 0.671121 | 0.636351 | 0.67571 | 0.661061 | down |
| 9651 | PLCH2 | 0.000397 | 0.54966 | 0.504673 | 0.469669 | 0.508001 | down |
| 6821 | SUOX | 0.000398 | 0.640736 | 0.557918 | 0.616246 | 0.604967 | down |
| 26146 | TRAF3IP1 | 0.000398 | 0.45545 | 0.442276 | 0.47649 | 0.458072 | down |
| 85403 | EAF1 | 0.000399 | 0.407757 | 0.493738 | 0.545703 | 0.482399 | down |
| 84162 | KIAA1109 | 0.000399 | 0.488433 | 0.46306 | 0.538816 | 0.496769 | down |
| 3141 | HLCS | 0.000399 | 0.559792 | 0.529521 | 0.457376 | 0.515563 | down |
| 114790 | STK11IP | 0.0004 | 0.571831 | 0.646637 | 0.622882 | 0.613783 | down |
| 5865 | RAB3B | 0.0004 | 0.447141 | 0.529942 | 0.455155 | 0.477413 | down |
| 254887 | ZDHHC23 | 0.0004 | 0.451289 | 0.458242 | 0.581313 | 0.496948 | down |
| 6768 | ST14 | 0.000401 | 0.680005 | 0.628728 | 0.597266 | 0.635333 | down |
| 283358 | B4GALNT3 | 0.000402 | 0.6 | 0.502754 | 0.521742 | 0.541499 | down |
| 54883 | CWC25 | 0.000402 | 0.513994 | 0.649413 | 0.58747 | 0.583626 | down |
| 5912 | RAP2B | 0.000403 | 0.551476 | 0.596094 | 0.562567 | 0.570046 | down |
| 9372 | ZFYVE9 | 0.000403 | 0.530528 | 0.430554 | 0.514263 | 0.491782 | down |
| 144100 | PLEKHA7 | 0.000404 | 0.588841 | 0.525523 | 0.599748 | 0.571371 | down |
| 56997 | COQ8A | 0.000404 | 0.665006 | 0.657577 | 0.635496 | 0.652693 | down |
| 22847 | ZNF507 | 0.000404 | 0.461719 | 0.517615 | 0.566023 | 0.515119 | down |
| 643338 | C15orf62 | 0.000405 | 0.4 | 0.319051 | 0.234432 | 0.317828 | down |
| 6502 | SKP2 | 0.000405 | 0.668349 | 0.648069 | 0.624983 | 0.647134 | down |
| 56255 | TMX4 | 0.000405 | 0.516372 | 0.507628 | 0.631845 | 0.551949 | down |
| 85441 | HELZ2 | 0.000405 | 0.579813 | 0.621563 | 0.546806 | 0.582727 | down |

| | | | | | | | |
|---|---|---|---|---|---|---|---|
| 2580 | GAK | 0.000406 | 0.565136 | 0.543175 | 0.491817 | 0.533376 | down |
| 55213 | RCBTB1 | 0.000406 | 0.608011 | 0.61303 | 0.668894 | 0.629979 | down |
| 9267 | CYTH1 | 0.000406 | 0.624891 | 0.517877 | 0.550668 | 0.564479 | down |
| 57510 | XPO5 | 0.000406 | 0.440514 | 0.484554 | 0.524629 | 0.483233 | down |
| 205717 | USF3 | 0.000408 | 0.33442 | 0.423613 | 0.476879 | 0.411637 | down |
| 83742 | MARVELD: | 0.000408 | 0.667301 | 0.654314 | 0.57249 | 0.631368 | down |
| 8906 | AP1G2 | 0.000408 | 0.654629 | 0.597309 | 0.651927 | 0.634622 | down |
| 143888 | KDELC2 | 0.000408 | 0.493576 | 0.469101 | 0.529888 | 0.497522 | down |
| 139285 | AMER1 | 0.00041 | 0.395225 | 0.369053 | 0.464646 | 0.409642 | down |
| 8312 | AXIN1 | 0.00041 | 0.596616 | 0.506005 | 0.540621 | 0.547747 | down |
| 60312 | AFAP1 | 0.00041 | 0.515641 | 0.521415 | 0.432338 | 0.489798 | down |
| 390010 | NKX1-2 | 0.000411 | 0.446097 | 0.474797 | 0.572486 | 0.497793 | down |
| 818 | CAMK2G | 0.000412 | 0.581394 | 0.681558 | 0.671573 | 0.644842 | down |
| 5361 | PLXNA1 | 0.000412 | 0.560325 | 0.569856 | 0.587492 | 0.572558 | down |
| 11190 | CEP250 | 0.000413 | 0.466162 | 0.293006 | 0.372475 | 0.377215 | down |
| 6938 | TCF12 | 0.000414 | 0.481387 | 0.537648 | 0.502965 | 0.507334 | down |
| 23157 | 44445 | 0.000415 | 0.540082 | 0.550559 | 0.659048 | 0.583229 | down |
| 9817 | KEAP1 | 0.000417 | 0.61928 | 0.659071 | 0.586161 | 0.621504 | down |
| 9258 | MFHAS1 | 0.000417 | 0.43577 | 0.43408 | 0.582113 | 0.483988 | down |
| 23245 | ASTN2 | 0.000417 | 0.601075 | 0.568792 | 0.497678 | 0.555848 | down |
| 23386 | NUDCD3 | 0.000417 | 0.666144 | 0.669677 | 0.648919 | 0.66158 | down |
| 84858 | ZNF503 | 0.000418 | 0.61286 | 0.668373 | 0.527557 | 0.60293 | down |
| 23335 | WDR7 | 0.000419 | 0.450472 | 0.548889 | 0.533804 | 0.511055 | down |
| 84256 | FLYWCH1 | 0.00042 | 0.679934 | 0.639033 | 0.655516 | 0.658161 | down |
| 57552 | NCEH1 | 0.00042 | 0.567513 | 0.53865 | 0.525155 | 0.543772 | down |
| 55646 | LYAR | 0.00042 | 0.673094 | 0.6565 | 0.593865 | 0.641153 | down |
| 9828 | ARHGEF17 | 0.000422 | 0.651554 | 0.590767 | 0.530435 | 0.590918 | down |
| 64218 | SEMA4A | 0.000422 | 0.624992 | 0.57579 | 0.679102 | 0.626628 | down |
| 23369 | PUM2 | 0.000425 | 0.610638 | 0.700096 | 0.67716 | 0.662632 | down |
| 161 | AP2A2 | 0.000426 | 0.627178 | 0.569479 | 0.474366 | 0.557008 | down |
| 2889 | RAPGEF1 | 0.000427 | 0.546206 | 0.543182 | 0.433694 | 0.507694 | down |
| 9897 | WASHC5 | 0.000427 | 0.63807 | 0.652491 | 0.654413 | 0.648325 | down |
| 55127 | HEATR1 | 0.000427 | 0.435234 | 0.49659 | 0.535653 | 0.489159 | down |
| 113201 | CASC4 | 0.000428 | 0.555536 | 0.632954 | 0.577968 | 0.588819 | down |
| 5523 | PPP2R3A | 0.000429 | 0.525679 | 0.62862 | 0.637066 | 0.597122 | down |
| 158586 | ZXDB | 0.000429 | 0.526042 | 0.498322 | 0.427576 | 0.48398 | down |
| 1846 | DUSP4 | 0.00043 | 0.593061 | 0.51357 | 0.563034 | 0.556555 | down |
| 79585 | CORO7 | 0.00043 | 0.623606 | 0.563985 | 0.478399 | 0.55533 | down |
| 9909 | DENND4B | 0.00043 | 0.688307 | 0.594958 | 0.643102 | 0.642122 | down |
| 285590 | SH3PXD2E | 0.00043 | 0.454392 | 0.389044 | 0.463761 | 0.435732 | down |
| 283450 | HECTD4 | 0.00043 | 0.413073 | 0.374462 | 0.390847 | 0.392794 | down |
| 5580 | PRKCD | 0.000431 | 0.585651 | 0.659758 | 0.53097 | 0.592126 | down |
| 29890 | RBM15B | 0.000431 | 0.615401 | 0.596149 | 0.560409 | 0.590653 | down |
| 65083 | NOL6 | 0.000432 | 0.512812 | 0.586441 | 0.595038 | 0.564764 | down |
| 2286 | FKBP2 | 0.000432 | 0.615806 | 0.659799 | 0.603872 | 0.626492 | down |
| 26088 | GGA1 | 0.000433 | 0.700435 | 0.602943 | 0.63349 | 0.645623 | down |
| 7515 | XRCC1 | 0.000433 | 0.621656 | 0.561019 | 0.514378 | 0.565684 | down |
| 55198 | APPL2 | 0.000433 | 0.521734 | 0.584885 | 0.644965 | 0.583861 | down |
| 219771 | CCNY | 0.000434 | 0.62968 | 0.698739 | 0.611782 | 0.646734 | down |
| 6603 | SMARCD2 | 0.000434 | 0.63859 | 0.708916 | 0.643853 | 0.663787 | down |
| 8994 | LIMD1 | 0.000435 | 0.539833 | 0.654781 | 0.555489 | 0.583368 | down |
| 60625 | DHX35 | 0.000436 | 0.575786 | 0.513892 | 0.540295 | 0.543324 | down |
| 4824 | NKX3-1 | 0.000436 | 0.545017 | 0.444332 | 0.51761 | 0.502319 | down |
| 22898 | DENND3 | 0.000437 | 0.590663 | 0.717849 | 0.608127 | 0.63888 | down |
| 23038 | WDTC1 | 0.000437 | 0.627322 | 0.503723 | 0.588066 | 0.573037 | down |
| 23067 | SETD1B | 0.000438 | 0.533781 | 0.408636 | 0.397443 | 0.44662 | down |
| 23255 | MTCL1 | 0.00044 | 0.403592 | 0.477625 | 0.421906 | 0.434374 | down |
| 23240 | TMEM131I | 0.00044 | 0.49408 | 0.551429 | 0.56899 | 0.538166 | down |

| | | | | | | | |
|---|---|---|---|---|---|---|---|
| 2026 | ENO2 | 0.00044 | 0.621152 | 0.576091 | 0.609809 | 0.602351 | down |
| 51347 | TAOK3 | 0.000441 | 0.597338 | 0.66949 | 0.636541 | 0.634456 | down |
| 23650 | TRIM29 | 0.000441 | 0.687316 | 0.599681 | 0.668065 | 0.651687 | down |
| 8050 | PDHX | 0.000442 | 0.695196 | 0.598222 | 0.670926 | 0.654781 | down |
| 343578 | ARHGAP4( | 0.000442 | 0.527531 | 0.610002 | 0.475552 | 0.537695 | down |
| 752 | FMNL1 | 0.000442 | 0.608283 | 0.608284 | 0.605464 | 0.607343 | down |
| 7716 | VEZF1 | 0.000443 | 0.453384 | 0.581863 | 0.56566 | 0.533636 | down |
| 84640 | USP38 | 0.000445 | 0.600798 | 0.629414 | 0.556277 | 0.595496 | down |
| 60558 | GUF1 | 0.000446 | 0.494922 | 0.599826 | 0.6241 | 0.572949 | down |
| 126308 | MOB3A | 0.000447 | 0.498213 | 0.475944 | 0.51571 | 0.496622 | down |
| 4205 | MEF2A | 0.000448 | 0.629756 | 0.565244 | 0.619947 | 0.604982 | down |
| 647135 | SRGAP2B | 0.000449 | 0.567127 | 0.646527 | 0.603186 | 0.605614 | down |
| 2767 | GNA11 | 0.000449 | 0.563811 | 0.48474 | 0.513344 | 0.520632 | down |
| 10970 | CKAP4 | 0.00045 | 0.573975 | 0.52482 | 0.445211 | 0.514669 | down |
| 57673 | BEND3 | 0.00045 | 0.36559 | 0.475552 | 0.455547 | 0.43223 | down |
| 204851 | HIPK1 | 0.000451 | 0.358724 | 0.468188 | 0.497902 | 0.441605 | down |
| 2200 | FBN1 | 0.000452 | 0.393939 | 0.429668 | 0.571429 | 0.465012 | down |
| 201164 | PLD6 | 0.000452 | 0.432789 | 0.564423 | 0.48965 | 0.49562 | down |
| 6668 | SP2 | 0.000452 | 0.575396 | 0.486474 | 0.5867 | 0.549523 | down |
| 65249 | ZSWIM4 | 0.000454 | 0.593362 | 0.66325 | 0.555726 | 0.604113 | down |
| 84939 | MUM1 | 0.000456 | 0.532633 | 0.612054 | 0.475539 | 0.540075 | down |
| 79960 | JADE1 | 0.000457 | 0.574441 | 0.55392 | 0.544884 | 0.557748 | down |
| 283578 | TMED8 | 0.000457 | 0.571573 | 0.617206 | 0.682925 | 0.623901 | down |
| 375449 | MAST4 | 0.000457 | 0.472222 | 0.502176 | 0.46092 | 0.478439 | down |
| 389432 | SAMD5 | 0.000457 | 0.574701 | 0.627841 | 0.654785 | 0.619109 | down |
| 51092 | SIDT2 | 0.000457 | 0.616086 | 0.552815 | 0.667606 | 0.612169 | down |
| 101 | ADAM8 | 0.000458 | 0.403498 | 0.438742 | 0.571086 | 0.471109 | down |
| 7014 | TERF2 | 0.000458 | 0.537364 | 0.581136 | 0.661275 | 0.593258 | down |
| 93643 | TJAP1 | 0.000459 | 0.684459 | 0.545847 | 0.563497 | 0.597934 | down |
| 81857 | MED25 | 0.00046 | 0.674062 | 0.577812 | 0.655762 | 0.635879 | down |
| 51592 | TRIM33 | 0.000461 | 0.483582 | 0.612122 | 0.628278 | 0.574661 | down |
| 56652 | TWNK | 0.000464 | 0.558115 | 0.612055 | 0.59871 | 0.589627 | down |
| 9013 | TAF1C | 0.000465 | 0.715951 | 0.598384 | 0.58809 | 0.634142 | down |
| 54629 | MINDY2 | 0.000466 | 0.626404 | 0.533997 | 0.617362 | 0.592588 | down |
| 55628 | ZNF407 | 0.000466 | 0.351545 | 0.437463 | 0.450835 | 0.413281 | down |
| 9658 | ZNF516 | 0.000467 | 0.426461 | 0.342874 | 0.450932 | 0.406755 | down |
| 114990 | VASN | 0.000468 | 0.562786 | 0.496063 | 0.401408 | 0.486752 | down |
| 7109 | TRAPPC10 | 0.000468 | 0.43701 | 0.425331 | 0.464549 | 0.442297 | down |
| 9351 | SLC9A3R2 | 0.000469 | 0.65826 | 0.531553 | 0.552851 | 0.580888 | down |
| 150275 | CCDC117 | 0.00047 | 0.488485 | 0.596733 | 0.589585 | 0.558268 | down |
| 55585 | UBE2Q1 | 0.000471 | 0.625361 | 0.563444 | 0.645815 | 0.61154 | down |
| 55697 | VAC14 | 0.000471 | 0.59371 | 0.680489 | 0.593121 | 0.62244 | down |
| 124359 | CDYL2 | 0.000472 | 0.565323 | 0.408163 | 0.470073 | 0.481187 | down |
| 23087 | TRIM35 | 0.000472 | 0.603976 | 0.54485 | 0.500855 | 0.549894 | down |
| 23108 | RAP1GAP2 | 0.000472 | 0.495047 | 0.600092 | 0.567387 | 0.554176 | down |
| 10256 | CNKSR1 | 0.000473 | 0.615021 | 0.510402 | 0.521524 | 0.548982 | down |
| 57478 | USP31 | 0.000473 | 0.647573 | 0.506563 | 0.617933 | 0.590689 | down |
| 688 | KLF5 | 0.000473 | 0.479474 | 0.57059 | 0.553258 | 0.53444 | down |
| 2180 | ACSL1 | 0.000475 | 0.535171 | 0.619566 | 0.556278 | 0.570338 | down |
| 54517 | PUS7 | 0.000476 | 0.544315 | 0.580047 | 0.517301 | 0.547221 | down |
| 11343 | MGLL | 0.000477 | 0.536665 | 0.532366 | 0.620325 | 0.563119 | down |
| 91949 | COG7 | 0.000479 | 0.602911 | 0.671742 | 0.654543 | 0.643065 | down |
| 1282 | COL4A1 | 0.000479 | 0.325019 | 0.377577 | 0.491514 | 0.398037 | down |
| 84181 | CHD6 | 0.00048 | 0.349615 | 0.434557 | 0.432439 | 0.405537 | down |
| 79073 | TMEM109 | 0.00048 | 0.586206 | 0.609792 | 0.644354 | 0.61345 | down |
| 8225 | GTPBP6 | 0.00048 | 0.624668 | 0.672525 | 0.537381 | 0.611525 | down |
| 29785 | CYP2S1 | 0.00048 | 0.571962 | 0.452086 | 0.527771 | 0.517273 | down |
| 23125 | CAMTA2 | 0.000481 | 0.648047 | 0.66881 | 0.554509 | 0.623789 | down |

| | | | | | | | |
|---|---|---|---|---|---|---|---|
| 1025 | CDK9 | 0.000481 | 0.6787 | 0.639685 | 0.543486 | 0.620623 | down |
| 5021 | OXTR | 0.000481 | 0.486322 | 0.555099 | 0.494029 | 0.511817 | down |
| 140688 | NOL4L | 0.000482 | 0.426965 | 0.59907 | 0.46962 | 0.498552 | down |
| 54897 | CASZ1 | 0.000483 | 0.59532 | 0.50243 | 0.451246 | 0.516332 | down |
| 54877 | ZCCHC2 | 0.000484 | 0.403135 | 0.459557 | 0.54848 | 0.470391 | down |
| 64224 | HERPUD2 | 0.000484 | 0.543834 | 0.529319 | 0.542745 | 0.538633 | down |
| 201232 | SLC16A13 | 0.000486 | 0.645852 | 0.496684 | 0.491632 | 0.544723 | down |
| 55116 | TMEM39B | 0.000486 | 0.561872 | 0.678144 | 0.668316 | 0.636111 | down |
| 6877 | TAF5 | 0.000487 | 0.571727 | 0.601714 | 0.446721 | 0.540054 | down |
| 54863 | TOR4A | 0.000489 | 0.716761 | 0.642721 | 0.608648 | 0.656043 | down |
| 51109 | RDH11 | 0.000489 | 0.546723 | 0.629388 | 0.612222 | 0.596111 | down |
| 90957 | DHX57 | 0.000489 | 0.500678 | 0.511452 | 0.621529 | 0.544553 | down |
| 51163 | DBR1 | 0.00049 | 0.599444 | 0.554399 | 0.688926 | 0.614256 | down |
| 5814 | PURB | 0.00049 | 0.476052 | 0.578717 | 0.54217 | 0.532313 | down |
| 23051 | ZHX3 | 0.00049 | 0.546706 | 0.570091 | 0.569722 | 0.562173 | down |
| 24148 | PRPF6 | 0.000492 | 0.542276 | 0.618474 | 0.53317 | 0.56464 | down |
| 144108 | SPTY2D1 | 0.000492 | 0.600614 | 0.583336 | 0.650449 | 0.611466 | down |
| 81570 | CLPB | 0.000492 | 0.585677 | 0.556678 | 0.462464 | 0.534939 | down |
| 23759 | PPIL2 | 0.000493 | 0.640471 | 0.627314 | 0.645997 | 0.637927 | down |
| 9679 | FAM53B | 0.000494 | 0.522436 | 0.560851 | 0.595882 | 0.559723 | down |
| 83696 | TRAPPC9 | 0.000494 | 0.568955 | 0.638298 | 0.620223 | 0.609159 | down |
| 6397 | SEC14L1 | 0.000495 | 0.459544 | 0.46399 | 0.536155 | 0.486563 | down |
| 4126 | MANBA | 0.000497 | 0.521759 | 0.385589 | 0.448062 | 0.451803 | down |
| 64780 | MICAL1 | 0.000498 | 0.620483 | 0.587456 | 0.704534 | 0.637491 | down |
| 10643 | IGF2BP3 | 0.000498 | 0.650175 | 0.709471 | 0.57483 | 0.644825 | down |
| 23141 | ANKLE2 | 0.000499 | 0.509077 | 0.596918 | 0.598632 | 0.568209 | down |
| 4488 | MSX2 | 0.000499 | 0.653291 | 0.532159 | 0.590352 | 0.591934 | down |
| 7026 | NR2F2 | 0.000499 | 0.66995 | 0.693005 | 0.582625 | 0.648526 | down |
| 26985 | AP3M1 | 0.000501 | 0.617946 | 0.611711 | 0.715655 | 0.648437 | down |
| 55672 | NBPF1 | 0.000502 | 0.416572 | 0.52945 | 0.584659 | 0.510227 | down |
| 10152 | ABI2 | 0.000503 | 0.516877 | 0.646374 | 0.549961 | 0.57107 | down |
| 7022 | TFAP2C | 0.000505 | 0.516437 | 0.590503 | 0.48843 | 0.53179 | down |
| 51278 | IER5 | 0.000507 | 0.663716 | 0.552707 | 0.539777 | 0.5854 | down |
| 89796 | NAV1 | 0.000508 | 0.362812 | 0.44971 | 0.345137 | 0.385886 | down |
| 9101 | USP8 | 0.000509 | 0.628639 | 0.694574 | 0.674194 | 0.665802 | down |
| 5905 | RANGAP1 | 0.00051 | 0.641541 | 0.607099 | 0.573604 | 0.607415 | down |
| 9815 | GIT2 | 0.000511 | 0.622383 | 0.535661 | 0.538221 | 0.565422 | down |
| 57210 | SLC45A4 | 0.000512 | 0.662 | 0.566761 | 0.544725 | 0.591162 | down |
| 22839 | DLGAP4 | 0.000512 | 0.568977 | 0.52493 | 0.477133 | 0.52368 | down |
| 83593 | RASSF5 | 0.000515 | 0.514848 | 0.502268 | 0.554067 | 0.523728 | down |
| 27043 | PELP1 | 0.000515 | 0.640622 | 0.48562 | 0.515609 | 0.547283 | down |
| 9853 | RUSC2 | 0.000515 | 0.637035 | 0.668143 | 0.523139 | 0.609439 | down |
| 55240 | STEAP3 | 0.000516 | 0.571092 | 0.590923 | 0.565364 | 0.575793 | down |
| 54509 | RHOF | 0.000518 | 0.638348 | 0.649575 | 0.570882 | 0.619601 | down |
| 84441 | MAML2 | 0.000519 | 0.59522 | 0.496549 | 0.444508 | 0.512093 | down |
| 22861 | NLRP1 | 0.00052 | 0.538241 | 0.600785 | 0.577703 | 0.572243 | down |
| 1830 | DSG3 | 0.000522 | 0.539363 | 0.543254 | 0.581719 | 0.554779 | down |
| 200958 | MUC20 | 0.000522 | 0.7248 | 0.624324 | 0.619895 | 0.656339 | down |
| 23417 | MLYCD | 0.000523 | 0.631543 | 0.482263 | 0.607852 | 0.573886 | down |
| 57514 | ARHGAP31 | 0.000523 | 0.546424 | 0.439394 | 0.539546 | 0.508455 | down |
| 89890 | KBTBD6 | 0.000523 | 0.510092 | 0.599373 | 0.596169 | 0.568545 | down |
| 5154 | PDGFA | 0.000525 | 0.585474 | 0.649203 | 0.53297 | 0.589216 | down |
| 197320 | ZNF778 | 0.000527 | 0.355082 | 0.427774 | 0.423169 | 0.402009 | down |
| 55051 | NRDE2 | 0.000528 | 0.543372 | 0.65268 | 0.486522 | 0.560858 | down |
| 284098 | PIGW | 0.000528 | 0.51269 | 0.517262 | 0.473395 | 0.501116 | down |
| 1294 | COL7A1 | 0.000529 | 0.5 | 0.361111 | 0.375 | 0.412037 | down |
| 9844 | ELMO1 | 0.000529 | 0.359696 | 0.413209 | 0.514482 | 0.429129 | down |
| 8916 | HERC3 | 0.000529 | 0.484314 | 0.542 | 0.442029 | 0.489448 | down |

| | | | | | | | |
|---|---|---|---|---|---|---|---|
| 169611 | OLFML2A | 0.000529 | 0.561863 | 0.516473 | 0.49289 | 0.523742 | down |
| 22821 | RASA3 | 0.000531 | 0.579821 | 0.554608 | 0.426269 | 0.520233 | down |
| 23258 | DENND5A | 0.000532 | 0.528202 | 0.554636 | 0.539433 | 0.540757 | down |
| 8021 | NUP214 | 0.000533 | 0.514948 | 0.605424 | 0.475409 | 0.531927 | down |
| 1859 | DYRK1A | 0.000533 | 0.517064 | 0.655794 | 0.644275 | 0.605711 | down |
| 10513 | APPBP2 | 0.000533 | 0.671718 | 0.583802 | 0.665856 | 0.640459 | down |
| 2908 | NR3C1 | 0.000533 | 0.616951 | 0.630907 | 0.58869 | 0.612183 | down |
| 1795 | DOCK3 | 0.000534 | 0.521429 | 0.571089 | 0.57862 | 0.557046 | down |
| 26873 | OPLAH | 0.000536 | 0.478636 | 0.575869 | 0.442229 | 0.498911 | down |
| 140901 | STK35 | 0.000536 | 0.508269 | 0.604534 | 0.533406 | 0.548736 | down |
| 201475 | RAB12 | 0.000537 | 0.545675 | 0.535446 | 0.616395 | 0.565839 | down |
| 117584 | RFFL | 0.000537 | 0.532392 | 0.536826 | 0.621954 | 0.563724 | down |
| 9797 | TATDN2 | 0.000537 | 0.608136 | 0.538784 | 0.524081 | 0.557 | down |
| 57827 | C6orf47 | 0.000538 | 0.579136 | 0.43452 | 0.536291 | 0.516649 | down |
| 9859 | CEP170 | 0.000538 | 0.566482 | 0.692512 | 0.592726 | 0.61724 | down |
| 246 | ALOX15 | 0.000539 | 0.581402 | 0.582745 | 0.674914 | 0.61302 | down |
| 23274 | CLEC16A | 0.00054 | 0.482022 | 0.529353 | 0.467829 | 0.493068 | down |
| 57537 | SORCS2 | 0.00054 | 0.446271 | 0.417632 | 0.541628 | 0.46851 | down |
| 90268 | OTULIN | 0.000541 | 0.566503 | 0.638998 | 0.584301 | 0.596601 | down |
| 29997 | NOP53 | 0.000541 | 0.607108 | 0.556519 | 0.519928 | 0.561185 | down |
| 9829 | DNAJC6 | 0.000542 | 0.47261 | 0.412607 | 0.591551 | 0.492256 | down |
| 3631 | INPP4A | 0.000543 | 0.522231 | 0.546408 | 0.425219 | 0.497953 | down |
| 8218 | CLTCL1 | 0.000543 | 0.638112 | 0.495 | 0.586824 | 0.573312 | down |
| 80301 | PLEKHO2 | 0.000543 | 0.662949 | 0.654347 | 0.564692 | 0.62733 | down |
| 285440 | CYP4V2 | 0.000543 | 0.570506 | 0.414931 | 0.547247 | 0.510895 | down |
| 23033 | DOP1A | 0.000544 | 0.545142 | 0.583211 | 0.496257 | 0.541537 | down |
| 9990 | SLC12A6 | 0.000545 | 0.525881 | 0.504861 | 0.531425 | 0.520722 | down |
| 163049 | ZNF791 | 0.000547 | 0.605595 | 0.550853 | 0.682416 | 0.612955 | down |
| 81573 | ANKRD13( | 0.000547 | 0.615347 | 0.693814 | 0.630624 | 0.646595 | down |
| 57798 | GATAD1 | 0.000547 | 0.686101 | 0.587798 | 0.581943 | 0.618614 | down |
| 4084 | MXD1 | 0.000548 | 0.671557 | 0.511887 | 0.551567 | 0.578337 | down |
| 286410 | ATP11C | 0.000549 | 0.635501 | 0.718607 | 0.636928 | 0.663679 | down |
| 10658 | CELF1 | 0.000549 | 0.534998 | 0.600633 | 0.497272 | 0.544301 | down |
| 10763 | NES | 0.000549 | 0.471717 | 0.406349 | 0.252234 | 0.376767 | down |
| 4650 | MYO9B | 0.000549 | 0.463532 | 0.497762 | 0.432242 | 0.464512 | down |
| 3268 | AGFG2 | 0.00055 | 0.565857 | 0.447186 | 0.614499 | 0.542514 | down |
| 285362 | SUMF1 | 0.00055 | 0.707124 | 0.576695 | 0.617184 | 0.633668 | down |
| 8239 | USP9X | 0.00055 | 0.467965 | 0.541379 | 0.553889 | 0.521078 | down |
| 80789 | INTS5 | 0.00055 | 0.635013 | 0.662522 | 0.668978 | 0.655504 | down |
| 5467 | PPARD | 0.00055 | 0.595707 | 0.556056 | 0.61918 | 0.590314 | down |
| 23081 | KDM4C | 0.000551 | 0.577918 | 0.687102 | 0.528082 | 0.597701 | down |
| 2650 | GCNT1 | 0.000551 | 0.645984 | 0.57663 | 0.688542 | 0.637052 | down |
| 158158 | RASEF | 0.000552 | 0.527452 | 0.508506 | 0.5285 | 0.521486 | down |
| 51696 | HECA | 0.000555 | 0.554475 | 0.666694 | 0.672078 | 0.631082 | down |
| 84247 | RTL6 | 0.000556 | 0.66839 | 0.547708 | 0.582074 | 0.599391 | down |
| 79670 | TUT7 | 0.000556 | 0.618894 | 0.556702 | 0.666311 | 0.613969 | down |
| 3833 | KIFC1 | 0.000556 | 0.58567 | 0.707908 | 0.636175 | 0.643251 | down |
| 9969 | MED13 | 0.000557 | 0.409629 | 0.452248 | 0.504712 | 0.455529 | down |
| 84126 | ATRIP | 0.000557 | 0.575 | 0.63408 | 0.667894 | 0.625658 | down |
| 342897 | NCCRP1 | 0.000558 | 0.537825 | 0.568859 | 0.507623 | 0.538102 | down |
| 5310 | PKD1 | 0.00056 | 0.580677 | 0.582034 | 0.520236 | 0.560982 | down |
| 11168 | PSIP1 | 0.00056 | 0.595086 | 0.688473 | 0.681104 | 0.654888 | down |
| 26115 | TANC2 | 0.00056 | 0.361629 | 0.430442 | 0.322302 | 0.371458 | down |
| 221178 | SPATA13 | 0.00056 | 0.501253 | 0.371506 | 0.436364 | 0.436374 | down |
| 51150 | SDF4 | 0.000561 | 0.692663 | 0.657947 | 0.599744 | 0.650118 | down |
| 56996 | SLC12A9 | 0.000562 | 0.57126 | 0.709918 | 0.642824 | 0.641334 | down |
| 64839 | FBXL17 | 0.000563 | 0.448709 | 0.591641 | 0.538636 | 0.526329 | down |
| 84166 | NLRC5 | 0.000563 | 0.587626 | 0.631751 | 0.618381 | 0.612586 | down |

| | | | | | | | |
|---|---|---|---|---|---|---|---|
| 2040 | STOM | 0.000564 | 0.622689 | 0.68612 | 0.688511 | 0.665773 | down |
| 374378 | GALNT18 | 0.000566 | 0.547402 | 0.682846 | 0.64092 | 0.623723 | down |
| 54108 | CHRAC1 | 0.000566 | 0.630125 | 0.603384 | 0.650606 | 0.628038 | down |
| 523 | ATP6V1A | 0.000567 | 0.43269 | 0.570604 | 0.462848 | 0.488714 | down |
| 3300 | DNAJB2 | 0.000568 | 0.665724 | 0.518976 | 0.596411 | 0.593704 | down |
| 23670 | CEMIP2 | 0.000569 | 0.475107 | 0.403392 | 0.4166 | 0.4317 | down |
| 5652 | PRSS8 | 0.00057 | 0.616068 | 0.55002 | 0.646757 | 0.604281 | down |
| 9926 | LPGAT1 | 0.000572 | 0.46894 | 0.60879 | 0.598519 | 0.558749 | down |
| 55784 | MCTP2 | 0.000573 | 0.370062 | 0.521377 | 0.520961 | 0.4708 | down |
| 1789 | DNMT3B | 0.000574 | 0.498494 | 0.516275 | 0.587211 | 0.533993 | down |
| 130507 | UBR3 | 0.000574 | 0.577681 | 0.491678 | 0.626022 | 0.565127 | down |
| 55604 | CARMIL1 | 0.000574 | 0.403022 | 0.425694 | 0.491244 | 0.439987 | down |
| 85369 | STRIP1 | 0.000574 | 0.620076 | 0.628885 | 0.726193 | 0.658385 | down |
| 22856 | CHSY1 | 0.000575 | 0.41586 | 0.43807 | 0.379857 | 0.411262 | down |
| 116113 | FOXP4 | 0.000575 | 0.680516 | 0.53118 | 0.550353 | 0.587349 | down |
| 284371 | ZNF841 | 0.000575 | 0.458876 | 0.433447 | 0.451741 | 0.448021 | down |
| 4643 | MYO1E | 0.000577 | 0.542043 | 0.44392 | 0.492562 | 0.492842 | down |
| 22893 | BAHD1 | 0.000578 | 0.559188 | 0.554699 | 0.650499 | 0.588129 | down |
| 11011 | TLK2 | 0.000578 | 0.569122 | 0.544847 | 0.532671 | 0.54888 | down |
| 54878 | DPP8 | 0.00058 | 0.549778 | 0.481894 | 0.410358 | 0.480677 | down |
| 4040 | LRP6 | 0.000581 | 0.3714 | 0.495938 | 0.407041 | 0.424793 | down |
| 79640 | C22orf46 | 0.000581 | 0.466098 | 0.399775 | 0.339131 | 0.401668 | down |
| 23185 | LARP4B | 0.000582 | 0.40277 | 0.491581 | 0.479991 | 0.458114 | down |
| 79613 | TANGO6 | 0.000582 | 0.691816 | 0.574139 | 0.573835 | 0.613263 | down |
| 85461 | TANC1 | 0.000583 | 0.447388 | 0.52235 | 0.432497 | 0.467412 | down |
| 847 | CAT | 0.000583 | 0.5164 | 0.606401 | 0.675045 | 0.599282 | down |
| 7158 | TP53BP1 | 0.000584 | 0.346492 | 0.477895 | 0.382936 | 0.402441 | down |
| 10371 | SEMA3A | 0.000584 | 0.513981 | 0.60534 | 0.558098 | 0.55914 | down |
| 84886 | C1orf198 | 0.000585 | 0.553158 | 0.644208 | 0.630176 | 0.609181 | down |
| 10844 | TUBGCP2 | 0.000585 | 0.637404 | 0.65758 | 0.628487 | 0.641157 | down |
| 9924 | PAN2 | 0.000587 | 0.3448 | 0.398997 | 0.435067 | 0.392954 | down |
| 10620 | ARID3B | 0.000587 | 0.597008 | 0.525508 | 0.490937 | 0.537818 | down |
| 4038 | LRP4 | 0.00059 | 0.460784 | 0.404029 | 0.404249 | 0.423021 | down |
| 1893 | ECM1 | 0.000591 | 0.552384 | 0.61499 | 0.574306 | 0.58056 | down |
| 2256 | FGF11 | 0.000592 | 0.629722 | 0.640038 | 0.566871 | 0.61221 | down |
| 7760 | ZNF213 | 0.000592 | 0.683723 | 0.581351 | 0.572957 | 0.612677 | down |
| 10482 | NXF1 | 0.000592 | 0.557849 | 0.467834 | 0.54507 | 0.523584 | down |
| 54764 | ZRANB1 | 0.000592 | 0.509686 | 0.635112 | 0.528935 | 0.557911 | down |
| 1371 | CPOX | 0.000592 | 0.525662 | 0.541174 | 0.517064 | 0.527967 | down |
| 3516 | RBPJ | 0.000593 | 0.404906 | 0.541516 | 0.529795 | 0.492073 | down |
| 474170 | LRRC37A2 | 0.000594 | 0.477428 | 0.533978 | 0.473547 | 0.494984 | down |
| 7791 | ZYX | 0.000596 | 0.711373 | 0.579093 | 0.59614 | 0.628869 | down |
| 375790 | AGRN | 0.000598 | 0.55859 | 0.557776 | 0.514525 | 0.54363 | down |
| 2289 | FKBP5 | 0.000599 | 0.390592 | 0.584291 | 0.473639 | 0.482841 | down |
| 154881 | KCTD7 | 0.000599 | 0.617914 | 0.549112 | 0.534448 | 0.567158 | down |
| 1.02E+08 | LOC10192 | 0.0006 | 0.690826 | 0.60228 | 0.541576 | 0.61156 | down |
| 9836 | LCMT2 | 0.000601 | 0.59906 | 0.71163 | 0.653619 | 0.65477 | down |
| 3787 | KCNS1 | 0.000602 | 0.516745 | 0.57772 | 0.509426 | 0.53463 | down |
| 6712 | SPTBN2 | 0.000603 | 0.523591 | 0.561724 | 0.448204 | 0.511173 | down |
| 5819 | NECTIN2 | 0.000604 | 0.57455 | 0.511081 | 0.533863 | 0.539831 | down |
| 4891 | SLC11A2 | 0.000604 | 0.582885 | 0.611495 | 0.726066 | 0.640149 | down |
| 23327 | NEDD4L | 0.000606 | 0.554858 | 0.592251 | 0.559777 | 0.568962 | down |
| 7058 | THBS2 | 0.000607 | 0.407221 | 0.445581 | 0.387224 | 0.413342 | down |
| 64645 | MFSD14A | 0.000608 | 0.592242 | 0.614748 | 0.679656 | 0.628882 | down |
| 353274 | ZNF445 | 0.000609 | 0.329768 | 0.395781 | 0.473427 | 0.399659 | down |
| 401548 | SNX30 | 0.00061 | 0.466053 | 0.480669 | 0.43727 | 0.461331 | down |
| 10079 | ATP9A | 0.000613 | 0.401852 | 0.465723 | 0.468597 | 0.44539 | down |
| 9943 | OXSR1 | 0.000614 | 0.409877 | 0.508175 | 0.530111 | 0.482721 | down |

| | | | | | | | |
|---|---|---|---|---|---|---|---|
| 55215 | FANCI | 0.000615 | 0.560364 | 0.662852 | 0.655015 | 0.626077 | down |
| 7779 | SLC30A1 | 0.000615 | 0.545757 | 0.62732 | 0.561748 | 0.578275 | down |
| 9581 | PREPL | 0.000616 | 0.568027 | 0.621791 | 0.688421 | 0.626079 | down |
| 1122 | CHML | 0.000616 | 0.56059 | 0.624306 | 0.466077 | 0.550324 | down |
| 9469 | CHST3 | 0.000616 | 0.472842 | 0.529193 | 0.495252 | 0.499096 | down |
| 6314 | ATXN7 | 0.000618 | 0.360635 | 0.413184 | 0.427283 | 0.400367 | down |
| 2896 | GRN | 0.000618 | 0.653222 | 0.595117 | 0.516649 | 0.588329 | down |
| 4486 | MST1R | 0.000619 | 0.619485 | 0.665756 | 0.679374 | 0.654872 | down |
| 85446 | ZFHX2 | 0.000619 | 0.351852 | 0.426316 | 0.407407 | 0.395192 | down |
| 123720 | WHAMM | 0.000619 | 0.479349 | 0.598886 | 0.5808 | 0.553012 | down |
| 79958 | DENND1C | 0.000621 | 0.312043 | 0.467891 | 0.490644 | 0.423526 | down |
| 54972 | TMEM132/ | 0.000622 | 0.621834 | 0.653968 | 0.582912 | 0.619571 | down |
| 80169 | CTC1 | 0.000624 | 0.53756 | 0.464489 | 0.541655 | 0.514568 | down |
| 28232 | SLCO3A1 | 0.000627 | 0.601806 | 0.617107 | 0.466518 | 0.56181 | down |
| 678 | ZFP36L2 | 0.000629 | 0.558562 | 0.519639 | 0.477995 | 0.518732 | down |
| 60496 | AASDHPP' | 0.000629 | 0.61263 | 0.665527 | 0.707322 | 0.661826 | down |
| 56342 | PPAN | 0.000631 | 0.690049 | 0.554514 | 0.675914 | 0.640159 | down |
| 79832 | QSER1 | 0.000632 | 0.550818 | 0.63431 | 0.692723 | 0.62595 | down |
| 114879 | OSBPL5 | 0.000633 | 0.418942 | 0.521268 | 0.37035 | 0.436853 | down |
| 9441 | MED26 | 0.000633 | 0.597112 | 0.507642 | 0.525952 | 0.543569 | down |
| 2072 | ERCC4 | 0.000634 | 0.436948 | 0.473726 | 0.463423 | 0.458032 | down |
| 23761 | PISD | 0.000634 | 0.696937 | 0.679488 | 0.593323 | 0.656583 | down |
| 9980 | DOP1B | 0.000635 | 0.408494 | 0.533965 | 0.421384 | 0.454614 | down |
| 54822 | TRPM7 | 0.000635 | 0.586922 | 0.648142 | 0.651973 | 0.629012 | down |
| 1954 | MEGF8 | 0.000636 | 0.495409 | 0.559141 | 0.599703 | 0.551418 | down |
| 26160 | IFT172 | 0.000638 | 0.54471 | 0.521692 | 0.406637 | 0.491013 | down |
| 2960 | GTF2E1 | 0.000642 | 0.651437 | 0.588703 | 0.599378 | 0.613173 | down |
| 2263 | FGFR2 | 0.000643 | 0.544092 | 0.598749 | 0.550964 | 0.564602 | down |
| 56606 | SLC2A9 | 0.000647 | 0.615056 | 0.695683 | 0.631202 | 0.647314 | down |
| 2533 | FYB1 | 0.000649 | 0.606781 | 0.523367 | 0.688267 | 0.606139 | down |
| 152137 | CCDC50 | 0.00065 | 0.62238 | 0.592206 | 0.69997 | 0.638185 | down |
| 79571 | GCC1 | 0.00065 | 0.444183 | 0.523123 | 0.488058 | 0.485121 | down |
| 10765 | KDM5B | 0.000653 | 0.396863 | 0.562575 | 0.514651 | 0.491363 | down |
| 10868 | USP20 | 0.000654 | 0.583041 | 0.607854 | 0.578253 | 0.589716 | down |
| 219833 | C11orf45 | 0.000657 | 0.666534 | 0.539848 | 0.624997 | 0.61046 | down |
| 55250 | ELP2 | 0.000658 | 0.684768 | 0.591304 | 0.577251 | 0.617774 | down |
| 23265 | EXOC7 | 0.000659 | 0.542723 | 0.640281 | 0.629495 | 0.604166 | down |
| 9949 | AMMECR1 | 0.00066 | 0.48243 | 0.608514 | 0.497256 | 0.5294 | down |
| 9818 | NUP58 | 0.00066 | 0.585011 | 0.621416 | 0.683748 | 0.630058 | down |
| 79813 | EHMT1 | 0.000661 | 0.502342 | 0.656407 | 0.577181 | 0.578643 | down |
| 64768 | IPPK | 0.000661 | 0.538981 | 0.564471 | 0.638462 | 0.580638 | down |
| 9440 | MED17 | 0.000662 | 0.634845 | 0.656993 | 0.668947 | 0.653595 | down |
| 91351 | DDX60L | 0.000662 | 0.634707 | 0.662585 | 0.695373 | 0.664222 | down |
| 114826 | SMYD4 | 0.000664 | 0.5445 | 0.647499 | 0.627331 | 0.606444 | down |
| 4008 | LMO7 | 0.000665 | 0.394865 | 0.517518 | 0.55625 | 0.489544 | down |
| 3713 | IVL | 0.000665 | 0.406912 | 0.394999 | 0.36481 | 0.388907 | down |
| 11099 | PTPN21 | 0.000666 | 0.364689 | 0.442085 | 0.489932 | 0.432235 | down |
| 57190 | SELENON | 0.000667 | 0.645536 | 0.64154 | 0.589152 | 0.625409 | down |
| 90 | ACVR1 | 0.000667 | 0.534854 | 0.639751 | 0.535862 | 0.570156 | down |
| 9706 | ULK2 | 0.000667 | 0.480401 | 0.559312 | 0.627551 | 0.555755 | down |
| 84456 | L3MBTL3 | 0.000668 | 0.463251 | 0.428359 | 0.559897 | 0.483836 | down |
| 11054 | OGFR | 0.000668 | 0.540427 | 0.398708 | 0.455229 | 0.464788 | down |
| 1820 | ARID3A | 0.000669 | 0.611717 | 0.561571 | 0.465429 | 0.546239 | down |
| 342357 | ZKSCAN2 | 0.000669 | 0.508786 | 0.412773 | 0.58171 | 0.50109 | down |
| 1848 | DUSP6 | 0.000669 | 0.665163 | 0.597263 | 0.641668 | 0.634698 | down |
| 57613 | FAM234B | 0.00067 | 0.400416 | 0.519957 | 0.582367 | 0.500913 | down |
| 81558 | FAM117A | 0.000671 | 0.526781 | 0.617689 | 0.604511 | 0.582994 | down |
| 219699 | UNC5B | 0.000671 | 0.486081 | 0.535811 | 0.526209 | 0.516034 | down |

| | | | | | | | |
|---|---|---|---|---|---|---|---|
| 284110 | GSDMA | 0.000671 | 0.643038 | 0.521907 | 0.615972 | 0.593639 | down |
| 79269 | DCAF10 | 0.000674 | 0.538074 | 0.53455 | 0.630462 | 0.567695 | down |
| 64689 | GORASP1 | 0.000676 | 0.627402 | 0.662329 | 0.699229 | 0.662987 | down |
| 23387 | SIK3 | 0.000679 | 0.552748 | 0.409744 | 0.51953 | 0.494008 | down |
| 6400 | SEL1L | 0.000679 | 0.369373 | 0.439678 | 0.440938 | 0.416663 | down |
| 55764 | IFT122 | 0.000681 | 0.604926 | 0.485105 | 0.410721 | 0.500251 | down |
| 79888 | LPCAT1 | 0.000681 | 0.545194 | 0.520142 | 0.504474 | 0.52327 | down |
| 5581 | PRKCE | 0.000684 | 0.443185 | 0.538733 | 0.504764 | 0.495561 | down |
| 63971 | KIF13A | 0.000686 | 0.585308 | 0.554328 | 0.580073 | 0.573236 | down |
| 55536 | CDCA7L | 0.000687 | 0.592869 | 0.716681 | 0.575479 | 0.628343 | down |
| 8976 | WASL | 0.000687 | 0.53648 | 0.659197 | 0.618972 | 0.604883 | down |
| 5371 | PML | 0.000688 | 0.686967 | 0.617732 | 0.657429 | 0.654043 | down |
| 1746 | DLX2 | 0.000688 | 0.639106 | 0.47149 | 0.519868 | 0.543488 | down |
| 54892 | NCAPG2 | 0.000689 | 0.47206 | 0.51449 | 0.34857 | 0.44504 | down |
| 9197 | SLC33A1 | 0.00069 | 0.536566 | 0.640809 | 0.552803 | 0.576726 | down |
| 84986 | ARHGAP1! | 0.000691 | 0.539454 | 0.580769 | 0.693585 | 0.604603 | down |
| 121536 | AEBP2 | 0.000691 | 0.469218 | 0.522038 | 0.640131 | 0.543796 | down |
| 64420 | SUSD1 | 0.000693 | 0.650727 | 0.606084 | 0.489399 | 0.58207 | down |
| 730755 | KRTAP2-3 | 0.000695 | 0.630284 | 0.69237 | 0.614632 | 0.645762 | down |
| 399474 | TMEM200I | 0.000696 | 0.584685 | 0.647165 | 0.588247 | 0.606699 | down |
| 54812 | AFTPH | 0.000698 | 0.574525 | 0.616409 | 0.639576 | 0.61017 | down |
| 7799 | PRDM2 | 0.000701 | 0.561655 | 0.529466 | 0.538955 | 0.543359 | down |
| 84890 | ADO | 0.000703 | 0.702642 | 0.568641 | 0.628664 | 0.633316 | down |
| 51588 | PIAS4 | 0.000704 | 0.607469 | 0.609978 | 0.598671 | 0.605373 | down |
| 10314 | LANCL1 | 0.000704 | 0.61106 | 0.644625 | 0.731689 | 0.662458 | down |
| 57455 | REXO1 | 0.000704 | 0.654421 | 0.595247 | 0.558115 | 0.602594 | down |
| 79183 | TTPAL | 0.000705 | 0.461751 | 0.574377 | 0.568418 | 0.534849 | down |
| 9673 | SLC25A44 | 0.000706 | 0.526104 | 0.604247 | 0.622878 | 0.58441 | down |
| 84766 | CRACR2A | 0.000706 | 0.487664 | 0.549934 | 0.452344 | 0.496647 | down |
| 4719 | NDUFS1 | 0.000708 | 0.580712 | 0.723698 | 0.677805 | 0.660739 | down |
| 26119 | LDLRAP1 | 0.000708 | 0.652431 | 0.534288 | 0.564763 | 0.583827 | down |
| 57132 | CHMP1B | 0.000709 | 0.601605 | 0.64728 | 0.565653 | 0.604846 | down |
| 5519 | PPP2R1B | 0.000711 | 0.565666 | 0.49965 | 0.562992 | 0.542769 | down |
| 1E+08 | ZNF605 | 0.000712 | 0.574425 | 0.71781 | 0.660365 | 0.650867 | down |
| 7156 | TOP3A | 0.000715 | 0.553691 | 0.614665 | 0.493555 | 0.55397 | down |
| 55701 | ARHGEF4C | 0.000716 | 0.541801 | 0.39376 | 0.515376 | 0.483645 | down |
| 23218 | NBEAL2 | 0.000716 | 0.668108 | 0.634946 | 0.550351 | 0.617802 | down |
| 55144 | LRRC8D | 0.000716 | 0.599444 | 0.735274 | 0.656497 | 0.663738 | down |
| 208 | AKT2 | 0.000716 | 0.724041 | 0.646698 | 0.612515 | 0.661085 | down |
| 114884 | OSBPL10 | 0.000719 | 0.405451 | 0.550963 | 0.545063 | 0.500492 | down |
| 23091 | ZC3H13 | 0.000719 | 0.509566 | 0.592892 | 0.432812 | 0.511757 | down |
| 57343 | ZNF304 | 0.000721 | 0.377097 | 0.403781 | 0.42905 | 0.403309 | down |
| 25836 | NIPBL | 0.000725 | 0.569794 | 0.680618 | 0.624173 | 0.624862 | down |
| 357 | SHROOM2 | 0.000727 | 0.36321 | 0.394918 | 0.445269 | 0.401132 | down |
| 79699 | ZYG11B | 0.000728 | 0.587191 | 0.53446 | 0.676812 | 0.599487 | down |
| 150726 | FBXO41 | 0.000729 | 0.657709 | 0.516092 | 0.587109 | 0.58697 | down |
| 55823 | VPS11 | 0.000731 | 0.53932 | 0.594257 | 0.563888 | 0.565822 | down |
| 55603 | TENT5A | 0.000732 | 0.418868 | 0.482754 | 0.609136 | 0.503586 | down |
| 23231 | SEL1L3 | 0.000735 | 0.566235 | 0.620187 | 0.544571 | 0.576998 | down |
| 8449 | DHX16 | 0.000738 | 0.558386 | 0.639784 | 0.574389 | 0.590853 | down |
| 1962 | EHHADH | 0.000739 | 0.511881 | 0.442368 | 0.566001 | 0.50675 | down |
| 535 | ATP6V0A1 | 0.000741 | 0.547266 | 0.458135 | 0.5497 | 0.518367 | down |
| 57719 | ANO8 | 0.000742 | 0.49098 | 0.516175 | 0.517477 | 0.508211 | down |
| 347344 | ZNF81 | 0.000743 | 0.504321 | 0.439537 | 0.5125 | 0.485453 | down |
| 57126 | CD177 | 0.000743 | 0.674293 | 0.660766 | 0.605093 | 0.646717 | down |
| 51701 | NLK | 0.000743 | 0.482644 | 0.553953 | 0.493367 | 0.509988 | down |
| 8445 | DYRK2 | 0.000747 | 0.515067 | 0.543478 | 0.571384 | 0.54331 | down |
| 9882 | TBC1D4 | 0.000747 | 0.447317 | 0.556532 | 0.497274 | 0.500374 | down |

| | | | | | | | |
|---|---|---|---|---|---|---|---|
| 5929 | RBBP5 | 0.000747 | 0.546 | 0.646225 | 0.5221 | 0.571442 | down |
| 1374 | CPT1A | 0.000748 | 0.607345 | 0.691027 | 0.669258 | 0.655877 | down |
| 23308 | ICOSLG | 0.000749 | 0.574812 | 0.594951 | 0.460184 | 0.543316 | down |
| 8349 | HIST2H2BI | 0.00075 | 0.564651 | 0.618493 | 0.65658 | 0.613241 | down |
| 54521 | WDR44 | 0.00075 | 0.557314 | 0.574526 | 0.699017 | 0.610286 | down |
| 29841 | GRHL1 | 0.00075 | 0.48669 | 0.56 | 0.539774 | 0.528821 | down |
| 3418 | IDH2 | 0.000752 | 0.651027 | 0.587855 | 0.599379 | 0.612753 | down |
| 23294 | ANKS1A | 0.000754 | 0.432347 | 0.521615 | 0.416146 | 0.456703 | down |
| 11138 | TBC1D8 | 0.000755 | 0.583412 | 0.53543 | 0.705189 | 0.60801 | down |
| 4796 | TONSL | 0.000756 | 0.70655 | 0.649486 | 0.591738 | 0.649258 | down |
| 144699 | FBXL14 | 0.000757 | 0.495305 | 0.5371 | 0.565409 | 0.532605 | down |
| 64167 | ERAP2 | 0.000757 | 0.574574 | 0.58778 | 0.716489 | 0.626281 | down |
| 150864 | FAM117B | 0.000758 | 0.533156 | 0.399898 | 0.523115 | 0.485389 | down |
| 79890 | RIN3 | 0.000758 | 0.6981 | 0.624256 | 0.568291 | 0.630216 | down |
| 146433 | IL34 | 0.000759 | 0.356474 | 0.419333 | 0.360087 | 0.378631 | down |
| 9061 | PAPSS1 | 0.000763 | 0.642751 | 0.557682 | 0.636345 | 0.612259 | down |
| 84461 | NEURL4 | 0.000766 | 0.618766 | 0.637248 | 0.500246 | 0.58542 | down |
| 6433 | SFSWAP | 0.000768 | 0.545891 | 0.509582 | 0.667527 | 0.574333 | down |
| 2002 | ELK1 | 0.00077 | 0.679861 | 0.639798 | 0.651665 | 0.657108 | down |
| 8884 | SLC5A6 | 0.00077 | 0.635916 | 0.599151 | 0.603949 | 0.613005 | down |
| 23592 | LEMD3 | 0.000772 | 0.656427 | 0.593807 | 0.681569 | 0.643934 | down |
| 114803 | MYSM1 | 0.000773 | 0.596437 | 0.642135 | 0.723624 | 0.654065 | down |
| 89958 | SAPCD2 | 0.000773 | 0.718435 | 0.591806 | 0.641523 | 0.650588 | down |
| 9681 | DEPDC5 | 0.000774 | 0.51634 | 0.546968 | 0.667742 | 0.577017 | down |
| 374655 | ZNF710 | 0.000775 | 0.595949 | 0.539574 | 0.546415 | 0.560646 | down |
| 8440 | NCK2 | 0.000777 | 0.727 | 0.567392 | 0.608398 | 0.634263 | down |
| 27092 | CACNG4 | 0.000779 | 0.575397 | 0.557692 | 0.695373 | 0.609487 | down |
| 5727 | PTCH1 | 0.00078 | 0.581779 | 0.468329 | 0.467611 | 0.505906 | down |
| 51703 | ACSL5 | 0.00078 | 0.566199 | 0.610983 | 0.653854 | 0.610345 | down |
| 22941 | SHANK2 | 0.00078 | 0.334512 | 0.487289 | 0.434652 | 0.418818 | down |
| 55617 | TASP1 | 0.000782 | 0.54435 | 0.688077 | 0.576107 | 0.602845 | down |
| 23137 | SMC5 | 0.000782 | 0.64754 | 0.530192 | 0.665289 | 0.61434 | down |
| 54540 | FAM193B | 0.000782 | 0.647109 | 0.631451 | 0.620617 | 0.633059 | down |
| 148362 | BROX | 0.000783 | 0.526963 | 0.654084 | 0.598659 | 0.593236 | down |
| 57512 | GPR158 | 0.000784 | 0.44527 | 0.471518 | 0.59617 | 0.504319 | down |
| 54805 | CNNM2 | 0.000787 | 0.561084 | 0.515211 | 0.539233 | 0.538509 | down |
| 57605 | PITPNM2 | 0.000789 | 0.618041 | 0.559774 | 0.639899 | 0.605905 | down |
| 7182 | NR2C2 | 0.00079 | 0.388468 | 0.47591 | 0.43942 | 0.434599 | down |
| 79829 | NAA40 | 0.000791 | 0.517222 | 0.594066 | 0.642759 | 0.584682 | down |
| 8263 | F8A1 | 0.000792 | 0.703984 | 0.591147 | 0.688782 | 0.661304 | down |
| 90333 | ZNF468 | 0.000796 | 0.650272 | 0.574097 | 0.666656 | 0.630342 | down |
| 2664 | GDI1 | 0.000796 | 0.730031 | 0.616419 | 0.633492 | 0.65998 | down |
| 5256 | PHKA2 | 0.000797 | 0.426065 | 0.502337 | 0.464403 | 0.464268 | down |
| 54894 | RNF43 | 0.000797 | 0.555718 | 0.479034 | 0.567394 | 0.534048 | down |
| 5093 | PCBP1 | 0.000798 | 0.701711 | 0.660436 | 0.620731 | 0.660959 | down |
| 414 | ARSD | 0.000801 | 0.495134 | 0.634232 | 0.648883 | 0.59275 | down |
| 3912 | LAMB1 | 0.000802 | 0.484576 | 0.493779 | 0.408949 | 0.462434 | down |
| 79660 | PPP1R3B | 0.000804 | 0.535331 | 0.43987 | 0.491843 | 0.489015 | down |
| 643314 | KIAA0754 | 0.000804 | 0.378508 | 0.324074 | 0.408877 | 0.370486 | down |
| 222584 | FAM83B | 0.000805 | 0.557882 | 0.639797 | 0.69077 | 0.629483 | down |
| 4215 | MAP3K3 | 0.000807 | 0.670704 | 0.527666 | 0.626545 | 0.608305 | down |
| 283431 | GAS2L3 | 0.000807 | 0.457603 | 0.611195 | 0.417974 | 0.495591 | down |
| 9881 | TRANK1 | 0.000808 | 0.298039 | 0.384793 | 0.377858 | 0.353564 | down |
| 23181 | DIP2A | 0.000809 | 0.602809 | 0.533217 | 0.436867 | 0.524298 | down |
| 114548 | NLRP3 | 0.000809 | 0.493479 | 0.329832 | 0.46616 | 0.429824 | down |
| 83959 | SLC4A11 | 0.000812 | 0.692238 | 0.595102 | 0.528406 | 0.605249 | down |
| 6499 | SKIV2L | 0.000812 | 0.617016 | 0.594365 | 0.549206 | 0.586862 | down |
| 22904 | SBNO2 | 0.000813 | 0.63579 | 0.608208 | 0.547074 | 0.597024 | down |

| | | | | | | | |
|---|---|---|---|---|---|---|---|
| 84450 | ZNF512 | 0.000813 | 0.65013 | 0.541364 | 0.524218 | 0.571904 | down |
| 5208 | PFKFB2 | 0.000814 | 0.438002 | 0.482123 | 0.476961 | 0.465695 | down |
| 9400 | RECQL5 | 0.000815 | 0.540478 | 0.581222 | 0.53303 | 0.551577 | down |
| 22981 | NINL | 0.000816 | 0.386502 | 0.403379 | 0.476555 | 0.422145 | down |
| 63926 | ANKEF1 | 0.000816 | 0.649519 | 0.594698 | 0.691785 | 0.645334 | down |
| 7049 | TGFBR3 | 0.000817 | 0.479167 | 0.621262 | 0.468652 | 0.523027 | down |
| 55131 | RBM28 | 0.000818 | 0.572189 | 0.597972 | 0.467463 | 0.545875 | down |
| 1540 | CYLD | 0.00082 | 0.65176 | 0.664583 | 0.573652 | 0.629998 | down |
| 133686 | NADK2 | 0.000822 | 0.65169 | 0.65683 | 0.592793 | 0.633771 | down |
| 4277 | MICB | 0.000822 | 0.566862 | 0.701603 | 0.55641 | 0.608292 | down |
| 3987 | LIMS1 | 0.000824 | 0.631431 | 0.576208 | 0.612132 | 0.606591 | down |
| 84301 | DDI2 | 0.000824 | 0.343414 | 0.405038 | 0.398879 | 0.382444 | down |
| 284615 | ANKRD34/ | 0.000825 | 0.538652 | 0.620347 | 0.684435 | 0.614478 | down |
| 23566 | LPAR3 | 0.000827 | 0.527785 | 0.642079 | 0.67262 | 0.614161 | down |
| 9874 | TLK1 | 0.000827 | 0.523844 | 0.64861 | 0.635236 | 0.602563 | down |
| 29960 | MRM2 | 0.000827 | 0.662186 | 0.63671 | 0.676331 | 0.658409 | down |
| 79705 | LRRK1 | 0.00083 | 0.563817 | 0.500745 | 0.394145 | 0.486236 | down |
| 51363 | CHST15 | 0.000835 | 0.529765 | 0.612922 | 0.468621 | 0.537103 | down |
| 57482 | KIAA1211 | 0.000836 | 0.542257 | 0.356406 | 0.532861 | 0.477175 | down |
| 4297 | KMT2A | 0.000837 | 0.317323 | 0.396512 | 0.358696 | 0.35751 | down |
| 8303 | SNN | 0.000837 | 0.640344 | 0.636341 | 0.61434 | 0.630342 | down |
| 9749 | PHACTR2 | 0.000837 | 0.652266 | 0.660085 | 0.659717 | 0.657356 | down |
| 80312 | TET1 | 0.000837 | 0.397878 | 0.365079 | 0.494545 | 0.419168 | down |
| 4799 | NFX1 | 0.00084 | 0.623606 | 0.658718 | 0.544459 | 0.608927 | down |
| 5365 | PLXNB3 | 0.00084 | 0.579408 | 0.688604 | 0.602721 | 0.623578 | down |
| 55054 | ATG16L1 | 0.000843 | 0.508554 | 0.558766 | 0.615092 | 0.560804 | down |
| 57187 | THOC2 | 0.000843 | 0.643129 | 0.631901 | 0.570694 | 0.615241 | down |
| 56946 | EMSY | 0.000846 | 0.522783 | 0.496189 | 0.490996 | 0.503322 | down |
| 27134 | TJP3 | 0.000848 | 0.603175 | 0.571681 | 0.619914 | 0.598257 | down |
| 55589 | BMP2K | 0.000848 | 0.424704 | 0.447095 | 0.509298 | 0.460365 | down |
| 134957 | STXBP5 | 0.00085 | 0.291397 | 0.328242 | 0.48351 | 0.367716 | down |
| 8535 | CBX4 | 0.000851 | 0.576651 | 0.46087 | 0.6385 | 0.558674 | down |
| 22902 | RUFY3 | 0.000852 | 0.637727 | 0.501929 | 0.625964 | 0.58854 | down |
| 152185 | SPICE1 | 0.000853 | 0.565319 | 0.701671 | 0.671119 | 0.646036 | down |
| 22846 | VASH1 | 0.000854 | 0.733175 | 0.640142 | 0.621214 | 0.664844 | down |
| 153572 | IRX2 | 0.000855 | 0.587143 | 0.655738 | 0.492929 | 0.578603 | down |
| 374383 | NCR3LG1 | 0.000856 | 0.429174 | 0.396384 | 0.414815 | 0.413458 | down |
| 10512 | SEMA3C | 0.000859 | 0.555662 | 0.619538 | 0.651663 | 0.608955 | down |
| 5045 | FURIN | 0.000862 | 0.664477 | 0.620178 | 0.560482 | 0.615046 | down |
| 84947 | SERAC1 | 0.000863 | 0.573809 | 0.560874 | 0.540817 | 0.5585 | down |
| 10142 | AKAP9 | 0.000865 | 0.60309 | 0.68386 | 0.690658 | 0.659203 | down |
| 23198 | PSME4 | 0.000867 | 0.57978 | 0.650445 | 0.674069 | 0.634765 | down |
| 5549 | PRELP | 0.000868 | 0.509671 | 0.530357 | 0.639716 | 0.559915 | down |
| 29068 | ZBTB44 | 0.000869 | 0.537124 | 0.684547 | 0.643708 | 0.621793 | down |
| 93183 | PIGM | 0.00087 | 0.617514 | 0.655004 | 0.587686 | 0.620068 | down |
| 169714 | QSOX2 | 0.00087 | 0.575114 | 0.578799 | 0.546728 | 0.56688 | down |
| 9925 | ZBTB5 | 0.000872 | 0.564907 | 0.526285 | 0.602767 | 0.564653 | down |
| 143098 | MPP7 | 0.000874 | 0.45813 | 0.554195 | 0.617046 | 0.543124 | down |
| 10432 | RBM14 | 0.000876 | 0.573261 | 0.594866 | 0.639394 | 0.602507 | down |
| 7023 | TFAP4 | 0.000876 | 0.6742 | 0.563642 | 0.487223 | 0.575022 | down |
| 9842 | PLEKHM1 | 0.000877 | 0.687983 | 0.641921 | 0.65818 | 0.662694 | down |
| 23347 | SMCHD1 | 0.000877 | 0.580058 | 0.582309 | 0.5685 | 0.576956 | down |
| 8940 | TOP3B | 0.000877 | 0.679461 | 0.514153 | 0.558366 | 0.583993 | down |
| 84129 | ACAD11 | 0.000878 | 0.669038 | 0.567213 | 0.653992 | 0.630081 | down |
| 29907 | SNX15 | 0.00088 | 0.701794 | 0.571527 | 0.633303 | 0.635541 | down |
| 3643 | INSR | 0.000881 | 0.3574 | 0.479206 | 0.408152 | 0.414919 | down |
| 23126 | POGZ | 0.000882 | 0.378642 | 0.455209 | 0.436008 | 0.423287 | down |
| 1871 | E2F3 | 0.000883 | 0.495063 | 0.58595 | 0.496299 | 0.52577 | down |

| | | | | | | | |
|---|---|---|---|---|---|---|---|
| 23677 | SH3BP4 | 0.000885 | 0.48642 | 0.596453 | 0.45759 | 0.513488 | down |
| 200205 | IBA57 | 0.000886 | 0.569388 | 0.708799 | 0.681484 | 0.653224 | down |
| 10681 | GNB5 | 0.000886 | 0.693472 | 0.626712 | 0.673038 | 0.664407 | down |
| 24147 | FJX1 | 0.000887 | 0.637538 | 0.697349 | 0.585241 | 0.640043 | down |
| 55722 | CEP72 | 0.000889 | 0.529088 | 0.597417 | 0.532841 | 0.553115 | down |
| 2997 | GYS1 | 0.000889 | 0.68333 | 0.631051 | 0.611443 | 0.641941 | down |
| 131566 | DCBLD2 | 0.000889 | 0.565508 | 0.631748 | 0.503961 | 0.567072 | down |
| 659 | BMPR2 | 0.000891 | 0.560088 | 0.551071 | 0.681517 | 0.597559 | down |
| 4000 | LMNA | 0.000892 | 0.596203 | 0.574308 | 0.537914 | 0.569475 | down |
| 80055 | PGAP1 | 0.000894 | 0.443876 | 0.532684 | 0.546875 | 0.507812 | down |
| 64599 | GIGYF1 | 0.000894 | 0.528917 | 0.59574 | 0.485366 | 0.536674 | down |
| 3880 | KRT19 | 0.000894 | 0.679897 | 0.65195 | 0.54964 | 0.627162 | down |
| 7988 | ZNF212 | 0.000895 | 0.680186 | 0.593943 | 0.600412 | 0.624847 | down |
| 4849 | CNOT3 | 0.000895 | 0.664243 | 0.545899 | 0.502664 | 0.570935 | down |
| 55251 | PCMTD2 | 0.000897 | 0.666667 | 0.555696 | 0.653317 | 0.625227 | down |
| 5784 | PTPN14 | 0.000897 | 0.462534 | 0.514719 | 0.604412 | 0.527221 | down |
| 26094 | DCAF4 | 0.000906 | 0.718117 | 0.599553 | 0.54947 | 0.62238 | down |
| 116987 | AGAP1 | 0.000906 | 0.525087 | 0.464715 | 0.512879 | 0.500894 | down |
| 51308 | REEP2 | 0.000908 | 0.624571 | 0.6763 | 0.686549 | 0.662473 | down |
| 9766 | SUSD6 | 0.000908 | 0.502369 | 0.504801 | 0.463805 | 0.490325 | down |
| 51435 | SCARA3 | 0.000909 | 0.629944 | 0.728077 | 0.597044 | 0.651688 | down |
| 375346 | STIMATE | 0.00091 | 0.641294 | 0.660121 | 0.591222 | 0.630879 | down |
| 10401 | PIAS3 | 0.00091 | 0.61985 | 0.518141 | 0.679613 | 0.605868 | down |
| 3556 | IL1RAP | 0.000911 | 0.628727 | 0.636251 | 0.702615 | 0.655865 | down |
| 54799 | MBTD1 | 0.000911 | 0.536691 | 0.483275 | 0.619842 | 0.546603 | down |
| 22927 | HABP4 | 0.000912 | 0.709122 | 0.555221 | 0.640641 | 0.634995 | down |
| 9807 | IP6K1 | 0.000914 | 0.728313 | 0.568015 | 0.628446 | 0.641591 | down |
| 130271 | PLEKHH2 | 0.000918 | 0.457366 | 0.404003 | 0.479322 | 0.446897 | down |
| 403313 | PLPP6 | 0.000922 | 0.507773 | 0.527099 | 0.466163 | 0.500345 | down |
| 222950 | NYAP1 | 0.000923 | 0.320497 | 0.40431 | 0.491256 | 0.405354 | down |
| 51804 | SIX4 | 0.000923 | 0.48339 | 0.539747 | 0.561301 | 0.528146 | down |
| 7626 | ZNF75D | 0.000924 | 0.576329 | 0.528127 | 0.423378 | 0.509278 | down |
| 868 | CBLB | 0.000928 | 0.556852 | 0.610714 | 0.628571 | 0.598713 | down |
| 51317 | PHF21A | 0.000932 | 0.57146 | 0.389712 | 0.536502 | 0.499225 | down |
| 10150 | MBNL2 | 0.000934 | 0.574903 | 0.656842 | 0.671092 | 0.634279 | down |
| 80210 | ARMC9 | 0.000935 | 0.494086 | 0.640853 | 0.540086 | 0.558342 | down |
| 23324 | MAN2B2 | 0.000937 | 0.442189 | 0.478338 | 0.484251 | 0.46826 | down |
| 55094 | GPATCH1 | 0.000937 | 0.703532 | 0.623092 | 0.540163 | 0.622263 | down |
| 4113 | MAGEB2 | 0.000941 | 0.527168 | 0.619904 | 0.635057 | 0.594043 | down |
| 3339 | HSPG2 | 0.000941 | 0.479633 | 0.539266 | 0.454933 | 0.491277 | down |
| 4810 | NHS | 0.000942 | 0.308271 | 0.384247 | 0.423181 | 0.371899 | down |
| 800 | CALD1 | 0.000942 | 0.529893 | 0.637813 | 0.664855 | 0.610853 | down |
| 55787 | TXLNG | 0.000946 | 0.589854 | 0.673723 | 0.63275 | 0.632109 | down |
| 7466 | WFS1 | 0.000947 | 0.596335 | 0.554955 | 0.584459 | 0.578583 | down |
| 6878 | TAF6 | 0.000948 | 0.73873 | 0.595395 | 0.58152 | 0.638548 | down |
| 11275 | KLHL2 | 0.000951 | 0.47332 | 0.423882 | 0.619542 | 0.505581 | down |
| 9223 | MAGI1 | 0.000952 | 0.348022 | 0.348188 | 0.324253 | 0.340154 | down |
| 4155 | MBP | 0.000955 | 0.533197 | 0.585871 | 0.539829 | 0.552966 | down |
| 3198 | HOXA1 | 0.000956 | 0.449437 | 0.581989 | 0.548926 | 0.526784 | down |
| 222389 | BEND7 | 0.000957 | 0.546174 | 0.615581 | 0.406831 | 0.522862 | down |
| 196528 | ARID2 | 0.000957 | 0.551824 | 0.666739 | 0.606622 | 0.608395 | down |
| 57695 | USP37 | 0.000958 | 0.373223 | 0.457135 | 0.604061 | 0.478139 | down |
| 9361 | LONP1 | 0.00096 | 0.600738 | 0.689482 | 0.535653 | 0.608625 | down |
| 1004 | CDH6 | 0.000964 | 0.25 | 0.5 | 0.333333 | 0.361111 | down |
| 9772 | TMEM94 | 0.000966 | 0.615602 | 0.536057 | 0.623343 | 0.591667 | down |
| 80824 | DUSP16 | 0.000968 | 0.523627 | 0.540411 | 0.440651 | 0.501563 | down |
| 5447 | POR | 0.000968 | 0.640404 | 0.56181 | 0.59917 | 0.600461 | down |
| 54454 | ATAD2B | 0.000968 | 0.551282 | 0.636395 | 0.448052 | 0.545243 | down |

| | | | | | | | |
|---|---|---|---|---|---|---|---|
| 55274 | PHF10 | 0.000968 | 0.593134 | 0.626811 | 0.734714 | 0.651553 | down |
| 84725 | PLEKHA8 | 0.000971 | 0.547158 | 0.528415 | 0.637663 | 0.571079 | down |
| 170690 | ADAMTS1( | 0.000973 | 0.444712 | 0.624396 | 0.521463 | 0.53019 | down |
| 1295 | COL8A1 | 0.000974 | 0.350131 | 0.45283 | 0.559633 | 0.454198 | down |
| 11245 | GPR176 | 0.000974 | 0.629541 | 0.645387 | 0.548661 | 0.607863 | down |
| 222229 | LRWD1 | 0.000975 | 0.53192 | 0.70976 | 0.592316 | 0.611332 | down |
| 57717 | PCDHB16 | 0.000975 | 0.325704 | 0.426421 | 0.28134 | 0.344488 | down |
| 144717 | PHETA1 | 0.000977 | 0.729793 | 0.649555 | 0.585755 | 0.655034 | down |
| 4629 | MYH11 | 0.000979 | 0.484848 | 0.412941 | 0.314685 | 0.404158 | down |
| 115752 | DIS3L | 0.000987 | 0.511773 | 0.513947 | 0.686439 | 0.57072 | down |
| 55346 | TCP11L1 | 0.000987 | 0.534308 | 0.622828 | 0.668966 | 0.608701 | down |
| 4998 | ORC1 | 0.000988 | 0.630265 | 0.60374 | 0.535179 | 0.589728 | down |
| 23764 | MAFF | 0.000989 | 0.51871 | 0.650621 | 0.585416 | 0.584916 | down |
| 6461 | SHB | 0.000991 | 0.634414 | 0.681608 | 0.510116 | 0.608713 | down |
| 57474 | ZNF490 | 0.000993 | 0.452479 | 0.448523 | 0.530924 | 0.477309 | down |
| 84902 | CEP89 | 0.000996 | 0.577811 | 0.48137 | 0.641083 | 0.566755 | down |
| 90355 | C5orf30 | 0.000997 | 0.593957 | 0.698139 | 0.684872 | 0.658989 | down |
| 64746 | ACBD3 | 0.000997 | 0.498957 | 0.616197 | 0.548123 | 0.554426 | down |

Table S3. related to Figure 5

| gene_id | gene_syml | P_value(sh | TR_shRNPS | TR_shRNPS | TR_shRNPS | average_TR_shRNPS1/shControl | |
|---|---|---|---|---|---|---|---|
| 57153 | SLC44A2 | 5.3E-08 | 0.611297 | 0.606943 | 0.610845 | 0.609695 | down |
| 3875 | KRT18 | 7.88E-08 | 0.629371 | 0.644462 | 0.641097 | 0.63831 | down |
| 8570 | KHSRP | 1.03E-07 | 0.664937 | 0.659824 | 0.657752 | 0.660838 | down |
| 54206 | ERRFI1 | 1.32E-07 | 0.672913 | 0.656907 | 0.669011 | 0.666277 | down |
| 2821 | GPI | 1.42E-07 | 0.61218 | 0.598998 | 0.602978 | 0.604718 | down |
| 83931 | STK40 | 2.01E-07 | 0.542484 | 0.538144 | 0.53558 | 0.538736 | down |
| 64132 | XYLT2 | 2.67E-07 | 0.570799 | 0.581386 | 0.579079 | 0.577088 | down |
| 9746 | CLSTN3 | 2.9E-07 | 0.563804 | 0.559928 | 0.551989 | 0.558574 | down |
| 1958 | EGR1 | 3.79E-07 | 0.328533 | 0.341449 | 0.335693 | 0.335225 | down |
| 200958 | MUC20 | 4.13E-07 | 0.557773 | 0.532039 | 0.549552 | 0.546455 | down |
| 50488 | MINK1 | 4.88E-07 | 0.584071 | 0.561365 | 0.574967 | 0.573468 | down |
| 3675 | ITGA3 | 4.98E-07 | 0.595408 | 0.61916 | 0.609389 | 0.607986 | down |
| 57636 | ARHGAP2: | 5.57E-07 | 0.578398 | 0.576445 | 0.556947 | 0.570597 | down |
| 59 | ACTA2 | 5.72E-07 | 0.575634 | 0.594601 | 0.580243 | 0.583493 | down |
| 2118 | ETV4 | 6.17E-07 | 0.649217 | 0.673256 | 0.659811 | 0.660762 | down |
| 1364 | CLDN4 | 6.42E-07 | 0.523968 | 0.523302 | 0.494651 | 0.513974 | down |
| 8189 | SYMPK | 6.66E-07 | 0.514717 | 0.513554 | 0.52732 | 0.518531 | down |
| 57462 | MYORG | 9.18E-07 | 0.598196 | 0.613591 | 0.623403 | 0.61173 | down |
| 1786 | DNMT1 | 1.48E-06 | 0.612354 | 0.613701 | 0.595363 | 0.607139 | down |
| 23061 | TBC1D9B | 1.53E-06 | 0.650962 | 0.662575 | 0.639596 | 0.651044 | down |
| 23264 | ZC3H7B | 1.6E-06 | 0.53876 | 0.530929 | 0.560791 | 0.543493 | down |
| 81 | ACTN4 | 1.67E-06 | 0.657591 | 0.677479 | 0.65404 | 0.663037 | down |
| 29954 | POMT2 | 1.68E-06 | 0.581764 | 0.593231 | 0.61123 | 0.595408 | down |
| 4854 | NOTCH3 | 1.69E-06 | 0.404909 | 0.446866 | 0.429823 | 0.427199 | down |
| 79143 | MBOAT7 | 1.77E-06 | 0.572955 | 0.560624 | 0.564889 | 0.566156 | down |
| 255783 | INAFM1 | 1.81E-06 | 0.644744 | 0.652462 | 0.623821 | 0.640342 | down |
| 91289 | LMF2 | 2.1E-06 | 0.583403 | 0.59804 | 0.608613 | 0.596685 | down |
| 10081 | PDCD7 | 2.11E-06 | 0.633146 | 0.658995 | 0.661454 | 0.651198 | down |
| 9918 | NCAPD2 | 2.19E-06 | 0.582069 | 0.561496 | 0.552576 | 0.56538 | down |
| 6929 | TCF3 | 2.35E-06 | 0.597654 | 0.566084 | 0.582647 | 0.582129 | down |
| 3996 | LLGL1 | 2.54E-06 | 0.548091 | 0.572065 | 0.543085 | 0.554414 | down |
| 10163 | WASF2 | 2.55E-06 | 0.575932 | 0.604787 | 0.578321 | 0.586347 | down |
| 54471 | MIEF1 | 2.64E-06 | 0.641392 | 0.639644 | 0.611468 | 0.630835 | down |
| 1453 | CSNK1D | 2.65E-06 | 0.651156 | 0.652118 | 0.65961 | 0.654295 | down |
| 9221 | NOLC1 | 2.78E-06 | 0.600953 | 0.639728 | 0.623371 | 0.621351 | down |
| 1497 | CTNS | 2.83E-06 | 0.617987 | 0.647407 | 0.641315 | 0.63557 | down |
| 23170 | TTLL12 | 2.9E-06 | 0.583744 | 0.568687 | 0.561873 | 0.571435 | down |
| 7517 | XRCC3 | 3E-06 | 0.538673 | 0.542203 | 0.554408 | 0.545095 | down |
| 4609 | MYC | 3.05E-06 | 0.474728 | 0.468972 | 0.489094 | 0.477598 | down |
| 57680 | CHD8 | 3.08E-06 | 0.660113 | 0.652055 | 0.627787 | 0.646651 | down |
| 10743 | RAI1 | 3.14E-06 | 0.62692 | 0.593665 | 0.59467 | 0.605085 | down |
| 9401 | RECQL4 | 3.22E-06 | 0.606867 | 0.630069 | 0.634154 | 0.623697 | down |
| 2196 | FAT2 | 3.25E-06 | 0.612504 | 0.627206 | 0.605559 | 0.61509 | down |
| 55226 | NAT10 | 3.26E-06 | 0.634256 | 0.604641 | 0.611793 | 0.616897 | down |
| 84678 | KDM2B | 3.31E-06 | 0.558538 | 0.597068 | 0.592434 | 0.58268 | down |
| 643669 | CCER2 | 3.46E-06 | 0.602414 | 0.595566 | 0.60092 | 0.599633 | down |
| 37 | ACADVL | 3.67E-06 | 0.522557 | 0.481405 | 0.484046 | 0.496003 | down |
| 23492 | CBX7 | 3.74E-06 | 0.483509 | 0.466047 | 0.476139 | 0.475232 | down |
| 10554 | AGPAT1 | 3.76E-06 | 0.49364 | 0.528171 | 0.484151 | 0.501987 | down |
| 79102 | RNF26 | 3.85E-06 | 0.555338 | 0.558545 | 0.542196 | 0.552026 | down |
| 23082 | PPRC1 | 4.26E-06 | 0.497025 | 0.519938 | 0.511812 | 0.509592 | down |
| 23218 | NBEAL2 | 4.67E-06 | 0.450405 | 0.425769 | 0.415108 | 0.430427 | down |
| 790 | CAD | 4.69E-06 | 0.517225 | 0.503683 | 0.501605 | 0.507504 | down |
| 80178 | TEDC2 | 4.7E-06 | 0.548779 | 0.52879 | 0.506218 | 0.527929 | down |
| 92017 | SNX29 | 4.75E-06 | 0.621737 | 0.596553 | 0.60455 | 0.607613 | down |
| 57659 | ZBTB4 | 4.8E-06 | 0.637923 | 0.665214 | 0.658701 | 0.653946 | down |

| | | | | | | | |
|---|---|---|---|---|---|---|---|
| 22883 | CLSTN1 | 4.81E-06 | 0.622306 | 0.62043 | 0.618795 | 0.620511 | down |
| 65123 | INTS3 | 4.85E-06 | 0.56962 | 0.522929 | 0.555795 | 0.549448 | down |
| 4771 | NF2 | 4.88E-06 | 0.627473 | 0.597472 | 0.608266 | 0.61107 | down |
| 57572 | DOCK6 | 4.99E-06 | 0.511262 | 0.521149 | 0.518754 | 0.517055 | down |
| 7468 | NSD2 | 5E-06 | 0.583581 | 0.580621 | 0.564771 | 0.576325 | down |
| 4150 | MAZ | 5.03E-06 | 0.632967 | 0.619377 | 0.649128 | 0.633824 | down |
| 8514 | KCNAB2 | 5.12E-06 | 0.530827 | 0.509939 | 0.527601 | 0.522789 | down |
| 51006 | SLC35C2 | 5.15E-06 | 0.566494 | 0.543111 | 0.551679 | 0.553762 | down |
| 79709 | COLGALT1 | 5.33E-06 | 0.675956 | 0.639003 | 0.651279 | 0.655413 | down |
| 2353 | FOS | 5.33E-06 | 0.440682 | 0.426344 | 0.394945 | 0.420657 | down |
| 84058 | WDR54 | 5.36E-06 | 0.604504 | 0.647406 | 0.633986 | 0.628632 | down |
| 6046 | BRD2 | 5.36E-06 | 0.593089 | 0.622354 | 0.581321 | 0.598922 | down |
| 2707 | GJB3 | 5.38E-06 | 0.619855 | 0.642205 | 0.604201 | 0.622087 | down |
| 5514 | PPP1R10 | 5.44E-06 | 0.592426 | 0.554375 | 0.600018 | 0.582273 | down |
| 27301 | APEX2 | 5.44E-06 | 0.658023 | 0.637861 | 0.613091 | 0.636325 | down |
| 382 | ARF6 | 5.45E-06 | 0.645736 | 0.674929 | 0.668071 | 0.662912 | down |
| 114794 | ELFN2 | 5.49E-06 | 0.350685 | 0.282857 | 0.336471 | 0.323338 | down |
| 5863 | RGL2 | 5.57E-06 | 0.59145 | 0.584936 | 0.571025 | 0.58247 | down |
| 55544 | RBM38 | 5.63E-06 | 0.631433 | 0.643292 | 0.653741 | 0.642822 | down |
| 84231 | TRAF7 | 5.67E-06 | 0.584873 | 0.561248 | 0.534336 | 0.560152 | down |
| 7851 | MALL | 5.78E-06 | 0.438743 | 0.448568 | 0.470663 | 0.452658 | down |
| 387521 | TMEM189 | 5.78E-06 | 0.629237 | 0.652327 | 0.648277 | 0.64328 | down |
| 22905 | EPN2 | 5.81E-06 | 0.612024 | 0.632568 | 0.63628 | 0.626957 | down |
| 112939 | NACC1 | 5.81E-06 | 0.676582 | 0.653186 | 0.649558 | 0.659775 | down |
| 58525 | WIZ | 5.93E-06 | 0.607229 | 0.628759 | 0.613551 | 0.616513 | down |
| 79581 | SLC52A2 | 5.95E-06 | 0.656197 | 0.666096 | 0.634745 | 0.652346 | down |
| 2686 | GGT7 | 5.99E-06 | 0.630476 | 0.642226 | 0.611047 | 0.627917 | down |
| 10869 | USP19 | 6.22E-06 | 0.565737 | 0.586358 | 0.594279 | 0.582125 | down |
| 16 | AARS | 6.3E-06 | 0.680504 | 0.666178 | 0.640568 | 0.662416 | down |
| 441478 | NRARP | 6.49E-06 | 0.474891 | 0.473839 | 0.474804 | 0.474511 | down |
| 80149 | ZC3H12A | 6.73E-06 | 0.416078 | 0.442709 | 0.458372 | 0.439053 | down |
| 5786 | PTPRA | 6.85E-06 | 0.581328 | 0.569645 | 0.609534 | 0.586836 | down |
| 975 | CD81 | 6.94E-06 | 0.60679 | 0.573191 | 0.60992 | 0.596634 | down |
| 55643 | BTBD2 | 6.95E-06 | 0.615665 | 0.613207 | 0.569163 | 0.599345 | down |
| 84720 | PIGO | 6.97E-06 | 0.524458 | 0.543389 | 0.524846 | 0.530898 | down |
| 622 | BDH1 | 7.02E-06 | 0.634286 | 0.653743 | 0.656444 | 0.648158 | down |
| 9826 | ARHGEF11 | 7.21E-06 | 0.587454 | 0.560776 | 0.552155 | 0.566795 | down |
| 7040 | TGFB1 | 7.26E-06 | 0.636433 | 0.611288 | 0.62711 | 0.624943 | down |
| 64218 | SEMA4A | 7.26E-06 | 0.558472 | 0.520966 | 0.500036 | 0.526491 | down |
| 78999 | LRFN4 | 7.3E-06 | 0.552861 | 0.560532 | 0.53181 | 0.548401 | down |
| 4668 | NAGA | 7.32E-06 | 0.607435 | 0.613477 | 0.587461 | 0.602791 | down |
| 79651 | RHBDF2 | 7.36E-06 | 0.554804 | 0.53554 | 0.530916 | 0.54042 | down |
| 8061 | FOSL1 | 7.49E-06 | 0.449261 | 0.423245 | 0.423568 | 0.432025 | down |
| 79134 | TMEM185I | 7.5E-06 | 0.498686 | 0.53447 | 0.494703 | 0.509286 | down |
| 27339 | PRPF19 | 7.52E-06 | 0.588661 | 0.609572 | 0.585441 | 0.594558 | down |
| 2783 | GNB2 | 7.72E-06 | 0.6485 | 0.658265 | 0.61959 | 0.642119 | down |
| 2177 | FANCD2 | 7.85E-06 | 0.609668 | 0.631425 | 0.654515 | 0.631869 | down |
| 25921 | ZDHHC5 | 7.85E-06 | 0.629803 | 0.660477 | 0.672134 | 0.654138 | down |
| 113452 | TMEM54 | 8.22E-06 | 0.687089 | 0.651036 | 0.659162 | 0.665762 | down |
| 2734 | GLG1 | 8.26E-06 | 0.608822 | 0.587444 | 0.623711 | 0.606659 | down |
| 5351 | PLOD1 | 8.35E-06 | 0.679 | 0.634339 | 0.647071 | 0.65347 | down |
| 23523 | CABIN1 | 8.53E-06 | 0.61992 | 0.57697 | 0.605355 | 0.600748 | down |
| 27352 | SGSM3 | 8.54E-06 | 0.63253 | 0.629976 | 0.62061 | 0.627705 | down |
| 56850 | GRIPAP1 | 8.8E-06 | 0.616377 | 0.614621 | 0.633605 | 0.621534 | down |
| 55670 | PEX26 | 8.82E-06 | 0.53907 | 0.590326 | 0.545502 | 0.558299 | down |
| 22984 | PDCD11 | 8.92E-06 | 0.581123 | 0.618412 | 0.561981 | 0.587172 | down |
| 79922 | MRM1 | 9.04E-06 | 0.527873 | 0.52578 | 0.564867 | 0.539507 | down |
| 1001 | CDH3 | 9.07E-06 | 0.673374 | 0.636137 | 0.634825 | 0.648112 | down |

| | | | | | | | |
|---|---|---|---|---|---|---|---|
| 51073 | MRPL4 | 9.07E-06 | 0.67771 | 0.654621 | 0.659982 | 0.664104 | down |
| 8862 | APLN | 9.26E-06 | 0.320114 | 0.348603 | 0.315718 | 0.328145 | down |
| 6575 | SLC20A2 | 9.36E-06 | 0.596106 | 0.554511 | 0.595673 | 0.582096 | down |
| 55735 | DNAJC11 | 9.5E-06 | 0.678391 | 0.655619 | 0.633828 | 0.655946 | down |
| 7703 | PCGF2 | 9.5E-06 | 0.618237 | 0.56827 | 0.604782 | 0.597096 | down |
| 9726 | ZNF646 | 9.6E-06 | 0.532154 | 0.513456 | 0.545009 | 0.530206 | down |
| 22992 | KDM2A | 9.96E-06 | 0.616038 | 0.627257 | 0.651277 | 0.631524 | down |
| 9777 | TM9SF4 | 1.01E-05 | 0.623035 | 0.575738 | 0.616305 | 0.605026 | down |
| 4221 | MEN1 | 1.02E-05 | 0.559191 | 0.585398 | 0.558442 | 0.567677 | down |
| 55128 | TRIM68 | 1.03E-05 | 0.643704 | 0.621263 | 0.645438 | 0.636802 | down |
| 56996 | SLC12A9 | 1.04E-05 | 0.515833 | 0.567143 | 0.523512 | 0.535496 | down |
| 3728 | JUP | 1.04E-05 | 0.607762 | 0.606287 | 0.642647 | 0.618899 | down |
| 162 | AP1B1 | 1.05E-05 | 0.663467 | 0.629104 | 0.617737 | 0.636769 | down |
| 7290 | HIRA | 1.05E-05 | 0.567968 | 0.624759 | 0.602135 | 0.598287 | down |
| 8704 | B4GALT2 | 1.06E-05 | 0.648771 | 0.623597 | 0.606188 | 0.626185 | down |
| 23310 | NCAPD3 | 1.06E-05 | 0.650787 | 0.679852 | 0.667158 | 0.665932 | down |
| 5296 | PIK3R2 | 1.08E-05 | 0.594346 | 0.62355 | 0.570035 | 0.595977 | down |
| 8703 | B4GALT3 | 1.08E-05 | 0.572576 | 0.589468 | 0.583512 | 0.581852 | down |
| 2064 | ERBB2 | 1.08E-05 | 0.611857 | 0.611267 | 0.577023 | 0.600049 | down |
| 4343 | MOV10 | 1.09E-05 | 0.636446 | 0.632097 | 0.657512 | 0.642018 | down |
| 10904 | BLCAP | 1.09E-05 | 0.581728 | 0.61007 | 0.636924 | 0.609574 | down |
| 5606 | MAP2K3 | 1.09E-05 | 0.641105 | 0.679441 | 0.636673 | 0.652407 | down |
| 92154 | MTSS1L | 1.09E-05 | 0.471774 | 0.446901 | 0.462401 | 0.460359 | down |
| 823 | CAPN1 | 1.12E-05 | 0.647669 | 0.63207 | 0.618898 | 0.632879 | down |
| 3927 | LASP1 | 1.13E-05 | 0.637601 | 0.675974 | 0.652694 | 0.655423 | down |
| 26020 | LRP10 | 1.13E-05 | 0.657744 | 0.684266 | 0.636976 | 0.659662 | down |
| 79784 | MYH14 | 1.14E-05 | 0.450968 | 0.443609 | 0.40728 | 0.433952 | down |
| 29882 | ANAPC2 | 1.15E-05 | 0.667168 | 0.642617 | 0.614965 | 0.641583 | down |
| 112 | ADCY6 | 1.17E-05 | 0.548035 | 0.534182 | 0.577705 | 0.553307 | down |
| 6692 | SPINT1 | 1.19E-05 | 0.666317 | 0.66488 | 0.649887 | 0.660361 | down |
| 113829 | SLC35A4 | 1.19E-05 | 0.578457 | 0.583388 | 0.615222 | 0.592356 | down |
| 8720 | MBTPS1 | 1.2E-05 | 0.59658 | 0.603412 | 0.62369 | 0.607894 | down |
| 9580 | SOX13 | 1.22E-05 | 0.597146 | 0.570859 | 0.539119 | 0.569041 | down |
| 8924 | HERC2 | 1.22E-05 | 0.52892 | 0.524554 | 0.472747 | 0.50874 | down |
| 56894 | AGPAT3 | 1.22E-05 | 0.512428 | 0.510492 | 0.5194 | 0.514107 | down |
| 10945 | KDELR1 | 1.23E-05 | 0.62115 | 0.672854 | 0.642321 | 0.645442 | down |
| 4296 | MAP3K11 | 1.26E-05 | 0.607549 | 0.593599 | 0.59112 | 0.597423 | down |
| 22879 | MON1B | 1.26E-05 | 0.559737 | 0.595151 | 0.566712 | 0.573867 | down |
| 55665 | URGCP | 1.27E-05 | 0.628303 | 0.628909 | 0.5836 | 0.613604 | down |
| 117246 | FTSJ3 | 1.28E-05 | 0.656959 | 0.608154 | 0.636535 | 0.633883 | down |
| 9703 | KIAA0100 | 1.29E-05 | 0.573954 | 0.608905 | 0.595064 | 0.592641 | down |
| 4294 | MAP3K10 | 1.3E-05 | 0.550756 | 0.504233 | 0.483429 | 0.512806 | down |
| 8888 | MCM3AP | 1.3E-05 | 0.597299 | 0.577353 | 0.582922 | 0.585858 | down |
| 6723 | SRM | 1.31E-05 | 0.542024 | 0.547565 | 0.53696 | 0.542183 | down |
| 3486 | IGFBP3 | 1.32E-05 | 0.555131 | 0.515486 | 0.577 | 0.549206 | down |
| 65249 | ZSWIM4 | 1.34E-05 | 0.587328 | 0.599976 | 0.619936 | 0.602414 | down |
| 6812 | STXBP1 | 1.35E-05 | 0.568708 | 0.576218 | 0.626561 | 0.590495 | down |
| 359948 | IRF2BP2 | 1.36E-05 | 0.651961 | 0.609216 | 0.619141 | 0.626773 | down |
| 682 | BSG | 1.36E-05 | 0.62328 | 0.597649 | 0.615317 | 0.612082 | down |
| 23193 | GANAB | 1.36E-05 | 0.582543 | 0.582229 | 0.603383 | 0.589385 | down |
| 23199 | GSE1 | 1.37E-05 | 0.645719 | 0.680359 | 0.632945 | 0.653007 | down |
| 2261 | FGFR3 | 1.37E-05 | 0.505274 | 0.472081 | 0.471164 | 0.48284 | down |
| 27112 | FAM155B | 1.37E-05 | 0.517506 | 0.497083 | 0.456823 | 0.49047 | down |
| 3959 | LGALS3BP | 1.37E-05 | 0.570128 | 0.525597 | 0.559759 | 0.551828 | down |
| 10523 | CHERP | 1.4E-05 | 0.618748 | 0.570606 | 0.597829 | 0.595728 | down |
| 51729 | WBP11 | 1.4E-05 | 0.645665 | 0.605537 | 0.65337 | 0.634857 | down |
| 929 | CD14 | 1.41E-05 | 0.55115 | 0.537897 | 0.570035 | 0.553027 | down |
| 54862 | CC2D1A | 1.44E-05 | 0.674813 | 0.623644 | 0.631956 | 0.643471 | down |

| | | | | | | | |
|---|---|---|---|---|---|---|---|
| 51621 | KLF13 | 1.45E-05 | 0.608427 | 0.635939 | 0.644622 | 0.629663 | down |
| 51593 | SRRT | 1.47E-05 | 0.60054 | 0.58816 | 0.645427 | 0.611376 | down |
| 10938 | EHD1 | 1.54E-05 | 0.573267 | 0.596981 | 0.582371 | 0.584206 | down |
| 196463 | PLBD2 | 1.54E-05 | 0.59169 | 0.635089 | 0.647363 | 0.624714 | down |
| 171023 | ASXL1 | 1.56E-05 | 0.669831 | 0.679785 | 0.642755 | 0.664124 | down |
| 60 | ACTB | 1.56E-05 | 0.598731 | 0.65257 | 0.616701 | 0.622667 | down |
| 10131 | TRAP1 | 1.56E-05 | 0.630975 | 0.623405 | 0.601806 | 0.618729 | down |
| 23247 | KIAA0556 | 1.57E-05 | 0.569987 | 0.593376 | 0.624759 | 0.596041 | down |
| 6662 | SOX9 | 1.58E-05 | 0.506886 | 0.53009 | 0.477648 | 0.504875 | down |
| 9600 | PITPNM1 | 1.58E-05 | 0.644637 | 0.592637 | 0.598895 | 0.612056 | down |
| 23509 | POFUT1 | 1.6E-05 | 0.677305 | 0.643321 | 0.678763 | 0.666463 | down |
| 10848 | PPP1R13L | 1.64E-05 | 0.637454 | 0.639253 | 0.622621 | 0.63311 | down |
| 4940 | OAS3 | 1.66E-05 | 0.512406 | 0.568475 | 0.564177 | 0.548353 | down |
| 5430 | POLR2A | 1.67E-05 | 0.564881 | 0.56778 | 0.552789 | 0.561816 | down |
| 9013 | TAF1C | 1.67E-05 | 0.588546 | 0.537889 | 0.566416 | 0.564284 | down |
| 64866 | CDCP1 | 1.69E-05 | 0.632632 | 0.6183 | 0.671955 | 0.640963 | down |
| 51291 | GMIP | 1.73E-05 | 0.672538 | 0.664725 | 0.621795 | 0.653019 | down |
| 5585 | PKN1 | 1.74E-05 | 0.607368 | 0.623483 | 0.60574 | 0.612197 | down |
| 1.01E+08 | INAFM2 | 1.74E-05 | 0.528657 | 0.567739 | 0.579416 | 0.558604 | down |
| 29803 | REPIN1 | 1.74E-05 | 0.494666 | 0.548695 | 0.556596 | 0.533319 | down |
| 6874 | TAF4 | 1.77E-05 | 0.547126 | 0.582835 | 0.615245 | 0.581735 | down |
| 2192 | FBLN1 | 1.8E-05 | 0.565459 | 0.604783 | 0.593769 | 0.588004 | down |
| 81848 | SPRY4 | 1.8E-05 | 0.604914 | 0.596116 | 0.625149 | 0.608726 | down |
| 23400 | ATP13A2 | 1.81E-05 | 0.526469 | 0.559372 | 0.498851 | 0.52823 | down |
| 83959 | SLC4A11 | 1.82E-05 | 0.43006 | 0.493185 | 0.469468 | 0.464238 | down |
| 57140 | RNPEPL1 | 1.83E-05 | 0.518604 | 0.494827 | 0.514215 | 0.509215 | down |
| 53373 | TPCN1 | 1.84E-05 | 0.444025 | 0.505242 | 0.481505 | 0.476924 | down |
| 113 | ADCY7 | 1.84E-05 | 0.438813 | 0.475811 | 0.4355 | 0.450042 | down |
| 1802 | DPH2 | 1.86E-05 | 0.612659 | 0.637453 | 0.590476 | 0.613529 | down |
| 25 | ABL1 | 1.87E-05 | 0.638735 | 0.683289 | 0.664219 | 0.662081 | down |
| 3654 | IRAK1 | 1.87E-05 | 0.677153 | 0.679471 | 0.630127 | 0.66225 | down |
| 197258 | FUK | 1.87E-05 | 0.479484 | 0.457155 | 0.47085 | 0.469163 | down |
| 283149 | BCL9L | 1.88E-05 | 0.583486 | 0.534642 | 0.569921 | 0.562683 | down |
| 23211 | ZC3H4 | 1.9E-05 | 0.494326 | 0.485151 | 0.443127 | 0.474201 | down |
| 56666 | PANX2 | 1.91E-05 | 0.631659 | 0.641155 | 0.587647 | 0.620154 | down |
| 1500 | CTNND1 | 1.99E-05 | 0.612633 | 0.639283 | 0.65524 | 0.635719 | down |
| 57222 | ERGIC1 | 1.99E-05 | 0.667243 | 0.646456 | 0.684251 | 0.665983 | down |
| 2806 | GOT2 | 2E-05 | 0.663656 | 0.657494 | 0.671479 | 0.664209 | down |
| 6891 | TAP2 | 2.01E-05 | 0.560117 | 0.567514 | 0.560409 | 0.56268 | down |
| 140688 | NOL4L | 2.02E-05 | 0.596908 | 0.576284 | 0.61808 | 0.597091 | down |
| 23765 | IL17RA | 2.02E-05 | 0.649493 | 0.622286 | 0.636556 | 0.636112 | down |
| 124222 | PAQR4 | 2.03E-05 | 0.469647 | 0.511331 | 0.513567 | 0.498182 | down |
| 7127 | TNFAIP2 | 2.05E-05 | 0.494885 | 0.528026 | 0.512062 | 0.511658 | down |
| 5768 | QSOX1 | 2.05E-05 | 0.607534 | 0.537779 | 0.552439 | 0.565918 | down |
| 1489 | CTF1 | 2.06E-05 | 0.541336 | 0.548623 | 0.525775 | 0.538578 | down |
| 22898 | DENND3 | 2.06E-05 | 0.618079 | 0.676393 | 0.642457 | 0.645643 | down |
| 57662 | CAMSAP3 | 2.06E-05 | 0.63508 | 0.621958 | 0.67073 | 0.642589 | down |
| 8906 | AP1G2 | 2.13E-05 | 0.582344 | 0.622935 | 0.589103 | 0.598127 | down |
| 91304 | TMEM259 | 2.14E-05 | 0.537544 | 0.542088 | 0.498852 | 0.526161 | down |
| 219 | ALDH1B1 | 2.18E-05 | 0.481062 | 0.499157 | 0.559463 | 0.513227 | down |
| 23139 | MAST2 | 2.19E-05 | 0.613707 | 0.608302 | 0.569457 | 0.597155 | down |
| 55691 | FRMD4A | 2.22E-05 | 0.473493 | 0.527203 | 0.480748 | 0.493815 | down |
| 64207 | IRF2BPL | 2.23E-05 | 0.643359 | 0.603522 | 0.589926 | 0.612269 | down |
| 22864 | R3HDM2 | 2.23E-05 | 0.535732 | 0.533323 | 0.601627 | 0.556894 | down |
| 651746 | ANKRD33E | 2.26E-05 | 0.445387 | 0.477723 | 0.481897 | 0.468336 | down |
| 192111 | PGAM5 | 2.27E-05 | 0.63192 | 0.646289 | 0.596238 | 0.624816 | down |
| 8986 | RPS6KA4 | 2.28E-05 | 0.684554 | 0.639025 | 0.667618 | 0.663733 | down |
| 1277 | COL1A1 | 2.29E-05 | 0.620744 | 0.573953 | 0.600949 | 0.598548 | down |

| | | | | | | | |
|---|---|---|---|---|---|---|---|
| 10120 | ACTR1B | 2.29E-05 | 0.672252 | 0.666168 | 0.621449 | 0.65329 | down |
| 81037 | CLPTM1L | 2.3E-05 | 0.649362 | 0.67602 | 0.65938 | 0.661588 | down |
| 150696 | PROM2 | 2.32E-05 | 0.528003 | 0.557477 | 0.597779 | 0.561087 | down |
| 6603 | SMARCD2 | 2.34E-05 | 0.573223 | 0.584872 | 0.62235 | 0.593482 | down |
| 1999 | ELF3 | 2.35E-05 | 0.653299 | 0.639516 | 0.628674 | 0.640496 | down |
| 29102 | DROSHA | 2.35E-05 | 0.606782 | 0.552036 | 0.590173 | 0.582997 | down |
| 6625 | SNRNP70 | 2.39E-05 | 0.624675 | 0.592776 | 0.606064 | 0.607839 | down |
| 10295 | BCKDK | 2.39E-05 | 0.518683 | 0.564905 | 0.559206 | 0.547598 | down |
| 58191 | CXCL16 | 2.42E-05 | 0.497818 | 0.515357 | 0.477265 | 0.496813 | down |
| 1981 | EIF4G1 | 2.43E-05 | 0.630885 | 0.653413 | 0.617026 | 0.633775 | down |
| 8140 | SLC7A5 | 2.44E-05 | 0.615056 | 0.658369 | 0.624779 | 0.632735 | down |
| 154810 | AMOTL1 | 2.46E-05 | 0.568817 | 0.585661 | 0.592694 | 0.582391 | down |
| 25929 | GEMIN5 | 2.48E-05 | 0.641408 | 0.632471 | 0.638537 | 0.637472 | down |
| 85456 | TNKS1BP1 | 2.48E-05 | 0.618599 | 0.673713 | 0.625578 | 0.639297 | down |
| 55841 | WWC3 | 2.49E-05 | 0.665293 | 0.599719 | 0.616292 | 0.627101 | down |
| 124565 | SLC38A10 | 2.51E-05 | 0.565219 | 0.510166 | 0.565265 | 0.546883 | down |
| 23163 | GGA3 | 2.51E-05 | 0.613913 | 0.657162 | 0.667217 | 0.646097 | down |
| 124454 | EARS2 | 2.54E-05 | 0.596139 | 0.644688 | 0.627994 | 0.622941 | down |
| 2022 | ENG | 2.56E-05 | 0.638324 | 0.620244 | 0.591498 | 0.616689 | down |
| 8558 | CDK10 | 2.61E-05 | 0.52164 | 0.542878 | 0.549946 | 0.538155 | down |
| 3551 | IKBKB | 2.67E-05 | 0.567285 | 0.540705 | 0.498649 | 0.535546 | down |
| 79180 | EFHD2 | 2.68E-05 | 0.64865 | 0.618031 | 0.581726 | 0.616136 | down |
| 1718 | DHCR24 | 2.73E-05 | 0.599167 | 0.647197 | 0.591576 | 0.612647 | down |
| 126282 | TNFAIP8L1 | 2.74E-05 | 0.540235 | 0.60458 | 0.602361 | 0.582392 | down |
| 90007 | MIDN | 2.76E-05 | 0.494699 | 0.523073 | 0.447682 | 0.488484 | down |
| 6242 | RTKN | 2.78E-05 | 0.634769 | 0.663914 | 0.594164 | 0.630949 | down |
| 56919 | DHX33 | 2.81E-05 | 0.617758 | 0.660897 | 0.615118 | 0.631258 | down |
| 23180 | RFTN1 | 2.82E-05 | 0.603948 | 0.647841 | 0.581259 | 0.611016 | down |
| 1490 | CTGF | 2.85E-05 | 0.528041 | 0.574874 | 0.583073 | 0.561996 | down |
| 31 | ACACA | 2.85E-05 | 0.587619 | 0.64135 | 0.634654 | 0.621208 | down |
| 60436 | TGIF2 | 2.86E-05 | 0.548728 | 0.620934 | 0.585063 | 0.584908 | down |
| 9054 | NFS1 | 2.87E-05 | 0.662976 | 0.657074 | 0.676065 | 0.665372 | down |
| 80279 | CDK5RAP3 | 2.87E-05 | 0.579505 | 0.59792 | 0.602495 | 0.593307 | down |
| 1454 | CSNK1E | 2.88E-05 | 0.57772 | 0.640732 | 0.619844 | 0.612766 | down |
| 1663 | DDX11 | 2.88E-05 | 0.547167 | 0.543433 | 0.526092 | 0.538897 | down |
| 2137 | EXTL3 | 2.89E-05 | 0.613008 | 0.654987 | 0.608585 | 0.625526 | down |
| 5287 | PIK3C2B | 2.89E-05 | 0.5057 | 0.525303 | 0.56059 | 0.530531 | down |
| 5442 | POLRMT | 2.91E-05 | 0.604555 | 0.605909 | 0.586271 | 0.598912 | down |
| 10106 | CTDSP2 | 2.92E-05 | 0.605507 | 0.592613 | 0.640351 | 0.612824 | down |
| 226 | ALDOA | 2.95E-05 | 0.676326 | 0.674778 | 0.622351 | 0.657818 | down |
| 30000 | TNPO2 | 2.95E-05 | 0.650221 | 0.657519 | 0.619843 | 0.642528 | down |
| 5119 | CHMP1A | 2.97E-05 | 0.661257 | 0.606699 | 0.620642 | 0.629533 | down |
| 11188 | NISCH | 2.98E-05 | 0.655837 | 0.596319 | 0.589223 | 0.613793 | down |
| 5883 | RAD9A | 2.99E-05 | 0.60999 | 0.630546 | 0.571096 | 0.603877 | down |
| 64319 | FBRS | 3.05E-05 | 0.66006 | 0.612528 | 0.635618 | 0.636069 | down |
| 1.03E+08 | HOTS | 3.05E-05 | 0.467918 | 0.474956 | 0.524487 | 0.48912 | down |
| 4301 | AFDN | 3.08E-05 | 0.637298 | 0.653133 | 0.691205 | 0.660545 | down |
| 4041 | LRP5 | 3.11E-05 | 0.499006 | 0.55817 | 0.501203 | 0.51946 | down |
| 220 | ALDH1A3 | 3.12E-05 | 0.599771 | 0.616484 | 0.571179 | 0.595811 | down |
| 613 | BCR | 3.16E-05 | 0.638857 | 0.648709 | 0.635136 | 0.6409 | down |
| 9961 | MVP | 3.17E-05 | 0.653071 | 0.661184 | 0.600192 | 0.638149 | down |
| 3860 | KRT13 | 3.18E-05 | 0.644199 | 0.59705 | 0.600479 | 0.613909 | down |
| 54765 | TRIM44 | 3.19E-05 | 0.625349 | 0.674588 | 0.662788 | 0.654242 | down |
| 26005 | C2CD3 | 3.19E-05 | 0.574198 | 0.604213 | 0.573508 | 0.583973 | down |
| 10645 | CAMKK2 | 3.21E-05 | 0.590417 | 0.644332 | 0.641806 | 0.625518 | down |
| 93487 | MAPK1IP1 | 3.23E-05 | 0.680631 | 0.650015 | 0.661511 | 0.664052 | down |
| 6837 | MED22 | 3.26E-05 | 0.621857 | 0.590276 | 0.590267 | 0.6008 | down |
| 6650 | CAPN15 | 3.27E-05 | 0.592806 | 0.596609 | 0.547438 | 0.578951 | down |

| | | | | | | | |
|---|---|---|---|---|---|---|---|
| 5609 | MAP2K7 | 3.28E-05 | 0.650135 | 0.698568 | 0.640814 | 0.663172 | down |
| 29775 | CARD10 | 3.28E-05 | 0.629646 | 0.649431 | 0.650799 | 0.643292 | down |
| 4363 | ABCC1 | 3.29E-05 | 0.60699 | 0.558331 | 0.56966 | 0.578327 | down |
| 54756 | IL17RD | 3.29E-05 | 0.575325 | 0.60375 | 0.617451 | 0.598842 | down |
| 78994 | PRR14 | 3.3E-05 | 0.539734 | 0.475451 | 0.487159 | 0.500781 | down |
| 9048 | ARTN | 3.31E-05 | 0.665041 | 0.667896 | 0.606452 | 0.646463 | down |
| 7994 | KAT6A | 3.32E-05 | 0.630487 | 0.64052 | 0.587612 | 0.61954 | down |
| 9610 | RIN1 | 3.35E-05 | 0.543143 | 0.550391 | 0.511255 | 0.534929 | down |
| 51114 | ZDHHC9 | 3.38E-05 | 0.609451 | 0.559878 | 0.601431 | 0.590253 | down |
| 4191 | MDH2 | 3.42E-05 | 0.619573 | 0.565753 | 0.599165 | 0.59483 | down |
| 4967 | OGDH | 3.43E-05 | 0.533488 | 0.523266 | 0.593523 | 0.550092 | down |
| 25987 | TSKU | 3.44E-05 | 0.645305 | 0.645065 | 0.667271 | 0.652547 | down |
| 29920 | PYCR2 | 3.44E-05 | 0.615188 | 0.660478 | 0.66118 | 0.645615 | down |
| 51604 | PIGT | 3.45E-05 | 0.661931 | 0.623561 | 0.688897 | 0.65813 | down |
| 9905 | SGSM2 | 3.45E-05 | 0.553483 | 0.589856 | 0.583045 | 0.575461 | down |
| 9123 | SLC16A3 | 3.47E-05 | 0.642017 | 0.620736 | 0.603339 | 0.622031 | down |
| 339287 | MSL1 | 3.48E-05 | 0.593248 | 0.658668 | 0.645375 | 0.63243 | down |
| 23357 | ANGEL1 | 3.52E-05 | 0.660436 | 0.645934 | 0.64547 | 0.650613 | down |
| 23246 | BOP1 | 3.56E-05 | 0.643631 | 0.653177 | 0.598751 | 0.631853 | down |
| 55040 | EPN3 | 3.6E-05 | 0.594028 | 0.509202 | 0.552429 | 0.551886 | down |
| 57553 | MICAL3 | 3.61E-05 | 0.643367 | 0.618334 | 0.595231 | 0.618977 | down |
| 55627 | SMPD4 | 3.61E-05 | 0.647814 | 0.597104 | 0.610017 | 0.618312 | down |
| 27158 | NDOR1 | 3.62E-05 | 0.598974 | 0.532297 | 0.579547 | 0.570273 | down |
| 23645 | PPP1R15A | 3.63E-05 | 0.590909 | 0.574691 | 0.625023 | 0.596874 | down |
| 9727 | RAB11FIP3 | 3.63E-05 | 0.549453 | 0.566111 | 0.58372 | 0.566428 | down |
| 9524 | TECR | 3.63E-05 | 0.572253 | 0.59973 | 0.633016 | 0.601666 | down |
| 91010 | FMNL3 | 3.65E-05 | 0.562404 | 0.612147 | 0.540736 | 0.571762 | down |
| 50628 | GEMIN4 | 3.66E-05 | 0.559538 | 0.528049 | 0.556086 | 0.547891 | down |
| 23005 | MAPKBP1 | 3.72E-05 | 0.477342 | 0.559585 | 0.518168 | 0.518365 | down |
| 399979 | SNX19 | 3.72E-05 | 0.58458 | 0.552456 | 0.612848 | 0.583295 | down |
| 1.12E+08 | LOC11226 | 3.73E-05 | 0.431123 | 0.448935 | 0.482454 | 0.454171 | down |
| 84875 | PARP10 | 3.74E-05 | 0.563819 | 0.528263 | 0.480319 | 0.524134 | down |
| 7739 | ZNF185 | 3.76E-05 | 0.663263 | 0.617223 | 0.647792 | 0.642759 | down |
| 5818 | NECTIN1 | 3.79E-05 | 0.587875 | 0.576297 | 0.546154 | 0.570109 | down |
| 7317 | UBA1 | 3.79E-05 | 0.611144 | 0.622666 | 0.648285 | 0.627365 | down |
| 9688 | NUP93 | 3.79E-05 | 0.59282 | 0.666838 | 0.626247 | 0.628635 | down |
| 80727 | TTYH3 | 3.82E-05 | 0.499125 | 0.515095 | 0.579332 | 0.531184 | down |
| 54620 | FBXL19 | 3.82E-05 | 0.583091 | 0.55059 | 0.607246 | 0.580309 | down |
| 25865 | PRKD2 | 3.84E-05 | 0.637332 | 0.625766 | 0.637828 | 0.633642 | down |
| 9788 | MTSS1 | 3.9E-05 | 0.653693 | 0.619493 | 0.669736 | 0.647641 | down |
| 6464 | SHC1 | 3.92E-05 | 0.633106 | 0.633196 | 0.664184 | 0.643495 | down |
| 83986 | FAM234A | 3.94E-05 | 0.576852 | 0.550633 | 0.490034 | 0.539173 | down |
| 6778 | STAT6 | 3.94E-05 | 0.508005 | 0.425731 | 0.439306 | 0.457681 | down |
| 7039 | TGFA | 3.96E-05 | 0.610157 | 0.661374 | 0.647718 | 0.63975 | down |
| 4796 | TONSL | 3.97E-05 | 0.466106 | 0.548758 | 0.51055 | 0.508472 | down |
| 23214 | XPO6 | 3.97E-05 | 0.55458 | 0.553349 | 0.590381 | 0.566103 | down |
| 10749 | KIF1C | 3.99E-05 | 0.647139 | 0.657211 | 0.618318 | 0.640889 | down |
| 3851 | KRT4 | 4.01E-05 | 0.507545 | 0.449001 | 0.403497 | 0.453348 | down |
| 30846 | EHD2 | 4.05E-05 | 0.545581 | 0.612088 | 0.55788 | 0.57185 | down |
| 6872 | TAF1 | 4.07E-05 | 0.630708 | 0.67231 | 0.656923 | 0.653314 | down |
| 23130 | ATG2A | 4.07E-05 | 0.54411 | 0.494497 | 0.522414 | 0.52034 | down |
| 26155 | NOC2L | 4.1E-05 | 0.652244 | 0.67321 | 0.648518 | 0.65799 | down |
| 51421 | AMOTL2 | 4.1E-05 | 0.563901 | 0.623428 | 0.617135 | 0.601488 | down |
| 26504 | CNNM4 | 4.13E-05 | 0.671053 | 0.603222 | 0.638184 | 0.637487 | down |
| 85414 | SLC45A3 | 4.13E-05 | 0.510016 | 0.520525 | 0.540153 | 0.523565 | down |
| 10397 | NDRG1 | 4.14E-05 | 0.665754 | 0.669152 | 0.641046 | 0.658651 | down |
| 4478 | MSN | 4.14E-05 | 0.654085 | 0.648859 | 0.681174 | 0.661373 | down |
| 2048 | EPHB2 | 4.14E-05 | 0.658197 | 0.695317 | 0.624546 | 0.659353 | down |

| | | | | | | | |
|---|---|---|---|---|---|---|---|
| 63893 | UBE2O | 4.17E-05 | 0.626704 | 0.691637 | 0.635492 | 0.651278 | down |
| 595 | CCND1 | 4.18E-05 | 0.590818 | 0.660489 | 0.624268 | 0.625191 | down |
| 8751 | ADAM15 | 4.18E-05 | 0.666168 | 0.599491 | 0.617632 | 0.627764 | down |
| 11151 | CORO1A | 4.21E-05 | 0.659672 | 0.641502 | 0.666179 | 0.655784 | down |
| 124245 | ZC3H18 | 4.22E-05 | 0.594775 | 0.585732 | 0.617136 | 0.599214 | down |
| 1366 | CLDN7 | 4.22E-05 | 0.668728 | 0.646277 | 0.603076 | 0.63936 | down |
| 161145 | TMEM229I | 4.23E-05 | 0.439125 | 0.400376 | 0.413205 | 0.417569 | down |
| 9810 | RNF40 | 4.25E-05 | 0.659194 | 0.590192 | 0.639975 | 0.629787 | down |
| 8224 | SYN3 | 4.25E-05 | 0.465656 | 0.501366 | 0.518609 | 0.49521 | down |
| 5894 | RAF1 | 4.28E-05 | 0.645296 | 0.605962 | 0.612565 | 0.621275 | down |
| 4804 | NGFR | 4.28E-05 | 0.478755 | 0.435132 | 0.430477 | 0.448121 | down |
| 170463 | SSBP4 | 4.3E-05 | 0.53696 | 0.54823 | 0.472214 | 0.519135 | down |
| 23344 | ESYT1 | 4.32E-05 | 0.592639 | 0.6366 | 0.601408 | 0.610216 | down |
| 9783 | RIMS3 | 4.33E-05 | 0.471429 | 0.363798 | 0.414475 | 0.416567 | down |
| 10298 | PAK4 | 4.33E-05 | 0.584141 | 0.590115 | 0.532862 | 0.569039 | down |
| 23268 | DNMBP | 4.35E-05 | 0.683224 | 0.618421 | 0.666366 | 0.656004 | down |
| 9962 | SLC23A2 | 4.39E-05 | 0.597894 | 0.583091 | 0.658071 | 0.613019 | down |
| 6405 | SEMA3F | 4.49E-05 | 0.491272 | 0.495586 | 0.474566 | 0.487141 | down |
| 9052 | GPRC5A | 4.5E-05 | 0.494561 | 0.547067 | 0.504461 | 0.515363 | down |
| 83590 | TMUB1 | 4.53E-05 | 0.578581 | 0.635831 | 0.576592 | 0.597001 | down |
| 2548 | GAA | 4.56E-05 | 0.522749 | 0.563102 | 0.535373 | 0.540408 | down |
| 57492 | ARID1B | 4.59E-05 | 0.659087 | 0.631198 | 0.639192 | 0.643159 | down |
| 286262 | TPRN | 4.6E-05 | 0.631245 | 0.605395 | 0.566077 | 0.600906 | down |
| 9927 | MFN2 | 4.6E-05 | 0.645118 | 0.69811 | 0.649377 | 0.664202 | down |
| 9392 | TGFBRAP1 | 4.63E-05 | 0.621251 | 0.676106 | 0.61893 | 0.638762 | down |
| 718 | C3 | 4.67E-05 | 0.516326 | 0.528159 | 0.476032 | 0.506839 | down |
| 115 | ADCY9 | 4.67E-05 | 0.516496 | 0.511886 | 0.536162 | 0.521515 | down |
| 7916 | PRRC2A | 4.68E-05 | 0.599826 | 0.552918 | 0.562404 | 0.571716 | down |
| 8525 | DGKZ | 4.69E-05 | 0.623162 | 0.594075 | 0.579403 | 0.59888 | down |
| 64837 | KLC2 | 4.7E-05 | 0.658721 | 0.669914 | 0.634794 | 0.654476 | down |
| 6653 | SORL1 | 4.72E-05 | 0.582748 | 0.529511 | 0.518383 | 0.543547 | down |
| 9779 | TBC1D5 | 4.73E-05 | 0.679209 | 0.662175 | 0.62771 | 0.656365 | down |
| 7625 | ZNF74 | 4.78E-05 | 0.621333 | 0.577952 | 0.639011 | 0.612765 | down |
| 284615 | ANKRD34/ | 4.81E-05 | 0.60545 | 0.610955 | 0.553678 | 0.590028 | down |
| 57479 | PRR12 | 4.81E-05 | 0.556967 | 0.515785 | 0.544935 | 0.539229 | down |
| 23001 | WDFY3 | 4.82E-05 | 0.581434 | 0.554554 | 0.549963 | 0.561984 | down |
| 8509 | NDST2 | 4.83E-05 | 0.626651 | 0.623691 | 0.609089 | 0.61981 | down |
| 7706 | TRIM25 | 4.86E-05 | 0.491209 | 0.527682 | 0.50352 | 0.50747 | down |
| 79650 | USB1 | 4.87E-05 | 0.505137 | 0.513959 | 0.5319 | 0.516999 | down |
| 1200 | TPP1 | 4.87E-05 | 0.584054 | 0.603318 | 0.635409 | 0.607594 | down |
| 5880 | RAC2 | 4.87E-05 | 0.617503 | 0.594487 | 0.667613 | 0.626534 | down |
| 64210 | MMS19 | 4.88E-05 | 0.677832 | 0.666849 | 0.621039 | 0.65524 | down |
| 2194 | FASN | 4.89E-05 | 0.523209 | 0.565586 | 0.511269 | 0.533355 | down |
| 899 | CCNF | 4.92E-05 | 0.645903 | 0.678878 | 0.673885 | 0.666222 | down |
| 9761 | MLEC | 4.93E-05 | 0.617207 | 0.668706 | 0.633228 | 0.639714 | down |
| 200734 | SPRED2 | 4.96E-05 | 0.645013 | 0.573728 | 0.593558 | 0.6041 | down |
| 1979 | EIF4EBP2 | 4.96E-05 | 0.638809 | 0.645278 | 0.669891 | 0.651326 | down |
| 10999 | SLC27A4 | 4.99E-05 | 0.636783 | 0.555072 | 0.579381 | 0.590412 | down |
| 23450 | SF3B3 | 5.06E-05 | 0.632129 | 0.670587 | 0.648908 | 0.650541 | down |
| 3321 | IGSF3 | 5.12E-05 | 0.613154 | 0.650655 | 0.626939 | 0.630249 | down |
| 8091 | HMGA2 | 5.14E-05 | 0.613089 | 0.627004 | 0.628449 | 0.622847 | down |
| 10312 | TCIRG1 | 5.14E-05 | 0.560981 | 0.620968 | 0.544373 | 0.575441 | down |
| 90780 | PYGO2 | 5.15E-05 | 0.629559 | 0.614358 | 0.639921 | 0.627946 | down |
| 57521 | RPTOR | 5.16E-05 | 0.534716 | 0.542287 | 0.504071 | 0.527025 | down |
| 51257 | 44257 | 5.16E-05 | 0.577699 | 0.547036 | 0.58352 | 0.569418 | down |
| 4784 | NFIX | 5.19E-05 | 0.614546 | 0.61937 | 0.598094 | 0.61067 | down |
| 23339 | VPS39 | 5.2E-05 | 0.651911 | 0.629347 | 0.645558 | 0.642272 | down |
| 10036 | CHAF1A | 5.2E-05 | 0.629082 | 0.618101 | 0.619902 | 0.622362 | down |

| | | | | | | | |
|---|---|---|---|---|---|---|---|
| 57787 | MARK4 | 5.2E-05 | 0.595004 | 0.602143 | 0.649037 | 0.615395 | down |
| 83715 | ESPN | 5.21E-05 | 0.502066 | 0.56626 | 0.493713 | 0.52068 | down |
| 11264 | PXMP4 | 5.22E-05 | 0.600282 | 0.584681 | 0.640222 | 0.608395 | down |
| 84747 | UNC119B | 5.26E-05 | 0.623285 | 0.597498 | 0.590692 | 0.603825 | down |
| 64755 | C16orf58 | 5.26E-05 | 0.567635 | 0.487163 | 0.514054 | 0.522951 | down |
| 11180 | WDR6 | 5.28E-05 | 0.59014 | 0.499365 | 0.563069 | 0.550858 | down |
| 5211 | PFKL | 5.3E-05 | 0.562024 | 0.606862 | 0.569466 | 0.579451 | down |
| 3659 | IRF1 | 5.3E-05 | 0.52469 | 0.554669 | 0.507418 | 0.528925 | down |
| 23191 | CYFIP1 | 5.3E-05 | 0.605007 | 0.59119 | 0.626008 | 0.607402 | down |
| 23513 | SCRIB | 5.37E-05 | 0.62093 | 0.564812 | 0.53519 | 0.573644 | down |
| 2039 | DMTN | 5.38E-05 | 0.499551 | 0.514304 | 0.4576 | 0.490485 | down |
| 9780 | PIEZO1 | 5.4E-05 | 0.59706 | 0.617785 | 0.548488 | 0.587778 | down |
| 103 | ADAR | 5.42E-05 | 0.649986 | 0.691303 | 0.652841 | 0.66471 | down |
| 83999 | KREMEN1 | 5.45E-05 | 0.567255 | 0.596472 | 0.623653 | 0.595793 | down |
| 4798 | NFRKB | 5.46E-05 | 0.654093 | 0.636869 | 0.641335 | 0.644099 | down |
| 894 | CCND2 | 5.47E-05 | 0.590486 | 0.657263 | 0.594541 | 0.614097 | down |
| 1E+08 | WASHC1 | 5.51E-05 | 0.672878 | 0.620266 | 0.673138 | 0.655427 | down |
| 79007 | DBNDD1 | 5.52E-05 | 0.505144 | 0.553215 | 0.556257 | 0.538205 | down |
| 80024 | SLC8B1 | 5.55E-05 | 0.550816 | 0.525075 | 0.535522 | 0.537137 | down |
| 79751 | SLC25A22 | 5.57E-05 | 0.513322 | 0.442699 | 0.502395 | 0.486139 | down |
| 283638 | CEP170B | 5.58E-05 | 0.67195 | 0.593555 | 0.643743 | 0.636416 | down |
| 81622 | UNC93B1 | 5.6E-05 | 0.601832 | 0.624977 | 0.575477 | 0.600762 | down |
| 89958 | SAPCD2 | 5.61E-05 | 0.534749 | 0.603315 | 0.515783 | 0.551283 | down |
| 23129 | PLXND1 | 5.63E-05 | 0.706308 | 0.662168 | 0.629165 | 0.66588 | down |
| 92799 | SHKBP1 | 5.64E-05 | 0.546608 | 0.637054 | 0.577873 | 0.587178 | down |
| 3141 | HLCS | 5.74E-05 | 0.583068 | 0.616077 | 0.614314 | 0.604486 | down |
| 8242 | KDM5C | 5.81E-05 | 0.624285 | 0.656489 | 0.610841 | 0.630538 | down |
| 29927 | SEC61A1 | 5.83E-05 | 0.63877 | 0.693788 | 0.664165 | 0.665574 | down |
| 9968 | MED12 | 5.85E-05 | 0.47803 | 0.568791 | 0.527752 | 0.524857 | down |
| 9776 | ATG13 | 5.85E-05 | 0.604378 | 0.633311 | 0.610314 | 0.616001 | down |
| 253980 | KCTD13 | 5.92E-05 | 0.551416 | 0.61059 | 0.579172 | 0.580393 | down |
| 3159 | HMGA1 | 5.93E-05 | 0.598138 | 0.545153 | 0.577065 | 0.573452 | down |
| 79567 | RIPOR1 | 5.95E-05 | 0.643381 | 0.563606 | 0.590944 | 0.59931 | down |
| 9917 | FAM20B | 5.97E-05 | 0.630923 | 0.626436 | 0.604986 | 0.620782 | down |
| 23780 | APOL2 | 5.99E-05 | 0.589818 | 0.617517 | 0.551049 | 0.586128 | down |
| 23380 | SRGAP2 | 6E-05 | 0.569551 | 0.598046 | 0.651572 | 0.60639 | down |
| 5326 | PLAGL2 | 6E-05 | 0.50746 | 0.587735 | 0.586446 | 0.560547 | down |
| 284613 | CYB561D1 | 6E-05 | 0.475181 | 0.513681 | 0.406169 | 0.46501 | down |
| 57533 | TBC1D14 | 6.03E-05 | 0.601349 | 0.628733 | 0.567065 | 0.599049 | down |
| 27242 | TNFRSF21 | 6.04E-05 | 0.592937 | 0.609816 | 0.637118 | 0.61329 | down |
| 6714 | SRC | 6.05E-05 | 0.508878 | 0.553768 | 0.597352 | 0.553333 | down |
| 55561 | CDC42BPG | 6.07E-05 | 0.580426 | 0.506676 | 0.561342 | 0.549481 | down |
| 113235 | SLC46A1 | 6.09E-05 | 0.669968 | 0.657707 | 0.620638 | 0.649438 | down |
| 3949 | LDLR | 6.1E-05 | 0.598371 | 0.625971 | 0.575844 | 0.600062 | down |
| 118738 | ZNF488 | 6.12E-05 | 0.481257 | 0.454263 | 0.538238 | 0.491253 | down |
| 957 | ENTPD5 | 6.12E-05 | 0.647787 | 0.61144 | 0.592149 | 0.617125 | down |
| 84985 | FAM83A | 6.13E-05 | 0.667812 | 0.596397 | 0.594187 | 0.619465 | down |
| 23254 | KAZN | 6.15E-05 | 0.557958 | 0.578841 | 0.574378 | 0.570392 | down |
| 56905 | C15orf39 | 6.22E-05 | 0.532128 | 0.508587 | 0.464252 | 0.501656 | down |
| 3303 | HSPA1A | 6.23E-05 | 0.5478 | 0.58004 | 0.524896 | 0.550912 | down |
| 50626 | CYHR1 | 6.23E-05 | 0.64574 | 0.606614 | 0.561379 | 0.604578 | down |
| 2720 | GLB1 | 6.23E-05 | 0.641716 | 0.654406 | 0.701579 | 0.6659 | down |
| 7866 | IFRD2 | 6.25E-05 | 0.675244 | 0.668102 | 0.602703 | 0.648683 | down |
| 8837 | CFLAR | 6.33E-05 | 0.630434 | 0.614026 | 0.646787 | 0.630416 | down |
| 9020 | MAP3K14 | 6.33E-05 | 0.585595 | 0.638289 | 0.609851 | 0.611245 | down |
| 26233 | FBXL6 | 6.34E-05 | 0.550651 | 0.582508 | 0.507859 | 0.547006 | down |
| 826 | CAPNS1 | 6.35E-05 | 0.638223 | 0.612698 | 0.671218 | 0.640713 | down |
| 8936 | WASF1 | 6.38E-05 | 0.625551 | 0.615385 | 0.654705 | 0.63188 | down |

| | | | | | | | |
|---|---|---|---|---|---|---|---|
| 10057 | ABCC5 | 6.4E-05 | 0.619529 | 0.576963 | 0.527656 | 0.574716 | down |
| 9700 | ESPL1 | 6.42E-05 | 0.61178 | 0.571561 | 0.526455 | 0.569932 | down |
| 254428 | SLC41A1 | 6.42E-05 | 0.568505 | 0.598268 | 0.650909 | 0.605894 | down |
| 57805 | CCAR2 | 6.49E-05 | 0.651085 | 0.706779 | 0.630911 | 0.662925 | down |
| 8824 | CES2 | 6.5E-05 | 0.595847 | 0.615723 | 0.600725 | 0.604098 | down |
| 23352 | UBR4 | 6.51E-05 | 0.496049 | 0.519375 | 0.520065 | 0.51183 | down |
| 23286 | WWC1 | 6.53E-05 | 0.565404 | 0.637843 | 0.642204 | 0.61515 | down |
| 54662 | TBC1D13 | 6.55E-05 | 0.63627 | 0.635445 | 0.670965 | 0.64756 | down |
| 4193 | MDM2 | 6.56E-05 | 0.633082 | 0.665027 | 0.625101 | 0.64107 | down |
| 9725 | TMEM63A | 6.59E-05 | 0.499455 | 0.509026 | 0.51072 | 0.5064 | down |
| 4302 | MLLT6 | 6.63E-05 | 0.514396 | 0.558967 | 0.57761 | 0.550324 | down |
| 127281 | PRXL2B | 6.66E-05 | 0.649275 | 0.623761 | 0.658871 | 0.643969 | down |
| 149473 | CCDC24 | 6.67E-05 | 0.656275 | 0.644469 | 0.609864 | 0.636869 | down |
| 79036 | KXD1 | 6.69E-05 | 0.595313 | 0.57658 | 0.624908 | 0.598933 | down |
| 126133 | ALDH16A1 | 6.7E-05 | 0.608089 | 0.584552 | 0.593839 | 0.595493 | down |
| 10626 | TRIM16 | 6.73E-05 | 0.606909 | 0.661613 | 0.625702 | 0.631408 | down |
| 51337 | THEM6 | 6.78E-05 | 0.542237 | 0.580569 | 0.606155 | 0.57632 | down |
| 27315 | PGAP2 | 6.82E-05 | 0.626813 | 0.654134 | 0.597637 | 0.626195 | down |
| 1627 | DBN1 | 6.89E-05 | 0.551105 | 0.493479 | 0.466793 | 0.503792 | down |
| 374819 | LRRC37A3 | 6.94E-05 | 0.627624 | 0.695275 | 0.676334 | 0.666411 | down |
| 84988 | PPP1R16A | 6.94E-05 | 0.596763 | 0.54759 | 0.521549 | 0.555301 | down |
| 55066 | PDPR | 6.96E-05 | 0.550012 | 0.581903 | 0.595837 | 0.575917 | down |
| 51400 | PPME1 | 6.96E-05 | 0.63012 | 0.638232 | 0.565159 | 0.61117 | down |
| 84335 | AKT1S1 | 7.02E-05 | 0.638601 | 0.656306 | 0.648666 | 0.647857 | down |
| 80347 | COASY | 7.02E-05 | 0.585673 | 0.634876 | 0.657643 | 0.626064 | down |
| 58986 | TMEM8A | 7.08E-05 | 0.604542 | 0.595688 | 0.552296 | 0.584175 | down |
| 81603 | TRIM8 | 7.09E-05 | 0.5116 | 0.54049 | 0.552758 | 0.534949 | down |
| 113130 | CDCA5 | 7.13E-05 | 0.655696 | 0.642282 | 0.574809 | 0.624263 | down |
| 54623 | PAF1 | 7.2E-05 | 0.651537 | 0.628899 | 0.704822 | 0.661753 | down |
| 6844 | VAMP2 | 7.21E-05 | 0.48417 | 0.551365 | 0.496324 | 0.51062 | down |
| 2036 | EPB41L1 | 7.23E-05 | 0.656096 | 0.683355 | 0.657147 | 0.665533 | down |
| 54461 | FBXW5 | 7.23E-05 | 0.614894 | 0.6196 | 0.615257 | 0.616584 | down |
| 80346 | REEP4 | 7.25E-05 | 0.628098 | 0.623662 | 0.569487 | 0.607082 | down |
| 22874 | PLEKHA6 | 7.29E-05 | 0.504834 | 0.505687 | 0.590888 | 0.533803 | down |
| 7407 | VARS | 7.33E-05 | 0.561892 | 0.568955 | 0.488456 | 0.539768 | down |
| 10723 | SLC12A7 | 7.34E-05 | 0.644274 | 0.596414 | 0.653555 | 0.631414 | down |
| 56893 | UBQLN4 | 7.35E-05 | 0.671157 | 0.656139 | 0.638487 | 0.655261 | down |
| 112936 | VPS26B | 7.37E-05 | 0.639022 | 0.60436 | 0.648018 | 0.630466 | down |
| 1431 | CS | 7.38E-05 | 0.597625 | 0.654854 | 0.669464 | 0.640648 | down |
| 5425 | POLD2 | 7.41E-05 | 0.624924 | 0.608274 | 0.600072 | 0.61109 | down |
| 55163 | PNPO | 7.42E-05 | 0.600917 | 0.605454 | 0.57837 | 0.594914 | down |
| 130074 | FAM168B | 7.42E-05 | 0.586728 | 0.663598 | 0.654611 | 0.634979 | down |
| 1376 | CPT2 | 7.44E-05 | 0.581773 | 0.605183 | 0.626855 | 0.604604 | down |
| 92840 | REEP6 | 7.44E-05 | 0.556393 | 0.623768 | 0.558787 | 0.57965 | down |
| 25777 | SUN2 | 7.45E-05 | 0.562987 | 0.593408 | 0.521481 | 0.559292 | down |
| 221092 | HNRNPUL | 7.47E-05 | 0.666464 | 0.626303 | 0.62834 | 0.640369 | down |
| 6051 | RNPEP | 7.5E-05 | 0.664665 | 0.694655 | 0.637377 | 0.665566 | down |
| 23363 | OBSL1 | 7.51E-05 | 0.596457 | 0.66737 | 0.59205 | 0.618626 | down |
| 125950 | RAVER1 | 7.52E-05 | 0.623874 | 0.598255 | 0.635495 | 0.619208 | down |
| 3315 | HSPB1 | 7.53E-05 | 0.644855 | 0.686869 | 0.637044 | 0.656256 | down |
| 2872 | MKNK2 | 7.55E-05 | 0.599776 | 0.686108 | 0.634938 | 0.640274 | down |
| 1846 | DUSP4 | 7.58E-05 | 0.576949 | 0.577094 | 0.548184 | 0.567409 | down |
| 9898 | UBAP2L | 7.6E-05 | 0.63291 | 0.640237 | 0.704878 | 0.659341 | down |
| 10514 | MYBBP1A | 7.62E-05 | 0.473736 | 0.490749 | 0.425935 | 0.463474 | down |
| 8641 | PCDHGB4 | 7.65E-05 | 0.545411 | 0.516189 | 0.506735 | 0.522778 | down |
| 51479 | ANKFY1 | 7.72E-05 | 0.642027 | 0.687177 | 0.607283 | 0.645495 | down |
| 9592 | IER2 | 7.74E-05 | 0.422588 | 0.497156 | 0.405654 | 0.441799 | down |
| 4298 | MLLT1 | 7.75E-05 | 0.589951 | 0.60741 | 0.582495 | 0.593285 | down |

| | | | | | | | |
|---|---|---|---|---|---|---|---|
| 1.08E+08 | LOC10798 | 7.78E-05 | 0.140519 | 0.25221 | 0.212951 | 0.201894 | down |
| 11044 | TENT4A | 7.79E-05 | 0.623043 | 0.68358 | 0.670218 | 0.658947 | down |
| 140710 | SOGA1 | 7.81E-05 | 0.617987 | 0.668991 | 0.626523 | 0.637834 | down |
| 10454 | TAB1 | 7.86E-05 | 0.573098 | 0.510765 | 0.602923 | 0.562262 | down |
| 23154 | NCDN | 7.86E-05 | 0.626916 | 0.617068 | 0.621659 | 0.621881 | down |
| 5253 | PHF2 | 7.89E-05 | 0.630141 | 0.681757 | 0.624619 | 0.645506 | down |
| 6949 | TCOF1 | 7.92E-05 | 0.62113 | 0.695617 | 0.638572 | 0.651773 | down |
| 9816 | URB2 | 7.95E-05 | 0.530713 | 0.570574 | 0.62242 | 0.574569 | down |
| 130535 | KCTD18 | 7.97E-05 | 0.627675 | 0.636062 | 0.639111 | 0.634283 | down |
| 4957 | ODF2 | 8.06E-05 | 0.564861 | 0.509406 | 0.510675 | 0.528314 | down |
| 26015 | RPAP1 | 8.09E-05 | 0.598875 | 0.505531 | 0.53116 | 0.545188 | down |
| 9830 | TRIM14 | 8.17E-05 | 0.468058 | 0.454432 | 0.549703 | 0.490731 | down |
| 10528 | NOP56 | 8.23E-05 | 0.654558 | 0.652969 | 0.647137 | 0.651555 | down |
| 9670 | IPO13 | 8.29E-05 | 0.611806 | 0.643202 | 0.614422 | 0.623143 | down |
| 24139 | EML2 | 8.29E-05 | 0.627653 | 0.56558 | 0.619013 | 0.604082 | down |
| 56829 | ZC3HAV1 | 8.31E-05 | 0.580212 | 0.582251 | 0.496497 | 0.552986 | down |
| 65998 | C11orf95 | 8.32E-05 | 0.584225 | 0.632919 | 0.573488 | 0.596878 | down |
| 116985 | ARAP1 | 8.34E-05 | 0.588645 | 0.656209 | 0.569628 | 0.604827 | down |
| 10160 | FARP1 | 8.35E-05 | 0.516871 | 0.580801 | 0.59551 | 0.564394 | down |
| 26007 | TKFC | 8.38E-05 | 0.598782 | 0.657808 | 0.591251 | 0.615947 | down |
| 4088 | SMAD3 | 8.51E-05 | 0.617857 | 0.574561 | 0.636858 | 0.609759 | down |
| 2919 | CXCL1 | 8.54E-05 | 0.638359 | 0.617135 | 0.627812 | 0.627769 | down |
| 80209 | PROSER1 | 8.69E-05 | 0.569475 | 0.586932 | 0.605798 | 0.587402 | down |
| 10919 | EHMT2 | 8.71E-05 | 0.568238 | 0.597579 | 0.498465 | 0.55476 | down |
| 91748 | ELMSAN1 | 8.74E-05 | 0.558782 | 0.600045 | 0.64482 | 0.601216 | down |
| 91433 | RCCD1 | 8.8E-05 | 0.528199 | 0.567893 | 0.624398 | 0.573497 | down |
| 2906 | GRIN2D | 8.83E-05 | 0.43985 | 0.496296 | 0.425026 | 0.453724 | down |
| 83475 | DOHH | 8.87E-05 | 0.559838 | 0.624708 | 0.653396 | 0.612647 | down |
| 8243 | SMC1A | 8.88E-05 | 0.630889 | 0.708916 | 0.638467 | 0.659424 | down |
| 128239 | IQGAP3 | 8.91E-05 | 0.538621 | 0.542497 | 0.506042 | 0.529053 | down |
| 201176 | ARHGAP27 | 9.02E-05 | 0.568738 | 0.602389 | 0.647331 | 0.606153 | down |
| 115290 | FBXO17 | 9.02E-05 | 0.626941 | 0.685228 | 0.631774 | 0.647981 | down |
| 64682 | ANAPC1 | 9.06E-05 | 0.613654 | 0.629392 | 0.678172 | 0.640406 | down |
| 219931 | TPCN2 | 9.1E-05 | 0.625348 | 0.66136 | 0.58197 | 0.622893 | down |
| 9828 | ARHGEF17 | 9.11E-05 | 0.505545 | 0.451493 | 0.446806 | 0.467948 | down |
| 254102 | EHBP1L1 | 9.11E-05 | 0.603375 | 0.533606 | 0.572463 | 0.569815 | down |
| 261734 | NPHP4 | 9.12E-05 | 0.591566 | 0.524461 | 0.609423 | 0.57515 | down |
| 7128 | TNFAIP3 | 9.13E-05 | 0.654575 | 0.572825 | 0.58976 | 0.60572 | down |
| 6548 | SLC9A1 | 9.14E-05 | 0.664993 | 0.614893 | 0.68394 | 0.654609 | down |
| 29941 | PKN3 | 9.15E-05 | 0.564603 | 0.582536 | 0.605219 | 0.58412 | down |
| 7023 | TFAP4 | 9.2E-05 | 0.587597 | 0.540624 | 0.588792 | 0.572338 | down |
| 4627 | MYH9 | 9.22E-05 | 0.537492 | 0.591737 | 0.528072 | 0.552434 | down |
| 57139 | RGL3 | 9.25E-05 | 0.536587 | 0.507077 | 0.596344 | 0.546669 | down |
| 10594 | PRPF8 | 9.25E-05 | 0.552583 | 0.585159 | 0.557544 | 0.565095 | down |
| 10195 | ALG3 | 9.37E-05 | 0.545797 | 0.574024 | 0.550138 | 0.556653 | down |
| 156 | GRK2 | 9.53E-05 | 0.628328 | 0.676405 | 0.610753 | 0.638495 | down |
| 54820 | NDE1 | 9.61E-05 | 0.646602 | 0.690084 | 0.603501 | 0.646729 | down |
| 1E+08 | ZNF316 | 9.64E-05 | 0.470596 | 0.500365 | 0.490947 | 0.487303 | down |
| 23223 | RRP12 | 9.71E-05 | 0.654116 | 0.645356 | 0.568946 | 0.622806 | down |
| 63891 | RNF123 | 9.74E-05 | 0.615841 | 0.541095 | 0.582397 | 0.579778 | down |
| 51599 | LSR | 9.74E-05 | 0.620587 | 0.658843 | 0.630358 | 0.636596 | down |
| 8502 | PKP4 | 9.78E-05 | 0.606872 | 0.668482 | 0.609981 | 0.628445 | down |
| 11100 | HNRNPUL | 9.8E-05 | 0.63394 | 0.70367 | 0.662375 | 0.666662 | down |
| 64857 | PLEKHG2 | 9.82E-05 | 0.566768 | 0.468588 | 0.516434 | 0.517263 | down |
| 55095 | SAMD4B | 9.82E-05 | 0.579628 | 0.574274 | 0.633553 | 0.595818 | down |
| 2186 | BPTF | 0.0001 | 0.59909 | 0.64961 | 0.621916 | 0.623539 | down |
| 7052 | TGM2 | 0.0001 | 0.563803 | 0.553115 | 0.458115 | 0.525011 | down |
| 83879 | CDCA7 | 0.000101 | 0.590658 | 0.642319 | 0.650345 | 0.627774 | down |

| | | | | | | | |
|---|---|---|---|---|---|---|---|
| 79711 | IPO4 | 0.000101 | 0.66027 | 0.607063 | 0.574476 | 0.613936 | down |
| 57610 | RANBP10 | 0.000101 | 0.635731 | 0.627438 | 0.655523 | 0.639564 | down |
| 6821 | SUOX | 0.000104 | 0.52528 | 0.546054 | 0.626754 | 0.566029 | down |
| 55362 | TMEM63B | 0.000104 | 0.663468 | 0.618182 | 0.573508 | 0.618386 | down |
| 23225 | NUP210 | 0.000104 | 0.584958 | 0.523462 | 0.474951 | 0.52779 | down |
| 760 | CA2 | 0.000105 | 0.604062 | 0.557393 | 0.56231 | 0.574588 | down |
| 587 | BCAT2 | 0.000105 | 0.648133 | 0.66274 | 0.651014 | 0.653962 | down |
| 376497 | SLC27A1 | 0.000106 | 0.604639 | 0.564351 | 0.507202 | 0.558731 | down |
| 9110 | MTMR4 | 0.000107 | 0.616847 | 0.657383 | 0.691587 | 0.655272 | down |
| 3566 | IL4R | 0.000107 | 0.575683 | 0.623296 | 0.622834 | 0.607271 | down |
| 9647 | PPM1F | 0.000108 | 0.591933 | 0.573759 | 0.51685 | 0.560848 | down |
| 2768 | GNA12 | 0.000108 | 0.62796 | 0.703802 | 0.664033 | 0.665265 | down |
| 55116 | TMEM39B | 0.000109 | 0.549097 | 0.596202 | 0.565853 | 0.570384 | down |
| 131583 | FAM43A | 0.000109 | 0.636169 | 0.643895 | 0.590079 | 0.623381 | down |
| 23524 | SRRM2 | 0.00011 | 0.516687 | 0.548726 | 0.547765 | 0.537726 | down |
| 23038 | WDTC1 | 0.00011 | 0.55035 | 0.506854 | 0.521056 | 0.526087 | down |
| 81844 | TRIM56 | 0.00011 | 0.4971 | 0.545522 | 0.575026 | 0.539216 | down |
| 64324 | NSD1 | 0.000111 | 0.610279 | 0.612043 | 0.625947 | 0.61609 | down |
| 57506 | MAVS | 0.000111 | 0.547107 | 0.633221 | 0.59601 | 0.592113 | down |
| 1382 | CRABP2 | 0.000111 | 0.624275 | 0.543392 | 0.56858 | 0.578749 | down |
| 5198 | PFAS | 0.000112 | 0.457421 | 0.483012 | 0.436027 | 0.45882 | down |
| 5426 | POLE | 0.000113 | 0.573353 | 0.526422 | 0.617764 | 0.572513 | down |
| 8603 | FAM193A | 0.000113 | 0.575991 | 0.595361 | 0.615653 | 0.595668 | down |
| 84687 | PPP1R9B | 0.000114 | 0.439649 | 0.509924 | 0.531698 | 0.493757 | down |
| 6878 | TAF6 | 0.000114 | 0.631037 | 0.530386 | 0.53988 | 0.567101 | down |
| 55739 | NAXD | 0.000114 | 0.591365 | 0.598932 | 0.501097 | 0.563798 | down |
| 6574 | SLC20A1 | 0.000114 | 0.631112 | 0.693148 | 0.634571 | 0.652944 | down |
| 6494 | SIPA1 | 0.000115 | 0.593072 | 0.596234 | 0.496733 | 0.562013 | down |
| 64900 | LPIN3 | 0.000115 | 0.529235 | 0.521769 | 0.511513 | 0.520839 | down |
| 79085 | SLC25A23 | 0.000116 | 0.550029 | 0.564185 | 0.61533 | 0.576515 | down |
| 1889 | ECE1 | 0.000116 | 0.658844 | 0.657499 | 0.589121 | 0.635155 | down |
| 64855 | FAM129B | 0.000116 | 0.555557 | 0.569168 | 0.55451 | 0.559745 | down |
| 8729 | GBF1 | 0.000117 | 0.6114 | 0.646559 | 0.547803 | 0.601921 | down |
| 7867 | MAPKAPK: | 0.000117 | 0.570869 | 0.660438 | 0.624701 | 0.618669 | down |
| 84878 | ZBTB45 | 0.000117 | 0.541372 | 0.637181 | 0.608055 | 0.595536 | down |
| 147179 | WIPF2 | 0.000117 | 0.567799 | 0.635181 | 0.643082 | 0.615354 | down |
| 607 | BCL9 | 0.000118 | 0.590511 | 0.645133 | 0.59886 | 0.611501 | down |
| 23125 | CAMTA2 | 0.000118 | 0.486326 | 0.519104 | 0.496617 | 0.500683 | down |
| 51341 | ZBTB7A | 0.000118 | 0.664635 | 0.615731 | 0.672457 | 0.650941 | down |
| 3954 | LETM1 | 0.000118 | 0.578253 | 0.602521 | 0.603418 | 0.594731 | down |
| 5871 | MAP4K2 | 0.000118 | 0.645224 | 0.642049 | 0.62399 | 0.637088 | down |
| 164091 | PAQR7 | 0.000118 | 0.599365 | 0.567282 | 0.571661 | 0.579436 | down |
| 3038 | HAS3 | 0.000118 | 0.37042 | 0.501485 | 0.375605 | 0.415837 | down |
| 58506 | SCAF1 | 0.000119 | 0.643583 | 0.622965 | 0.572856 | 0.613135 | down |
| 6829 | SUPT5H | 0.000119 | 0.6113 | 0.6415 | 0.603318 | 0.618706 | down |
| 192683 | SCAMP5 | 0.000119 | 0.504471 | 0.560559 | 0.582838 | 0.549289 | down |
| 1847 | DUSP5 | 0.00012 | 0.515418 | 0.613586 | 0.584896 | 0.5713 | down |
| 8733 | GPAA1 | 0.00012 | 0.650916 | 0.623348 | 0.572107 | 0.615457 | down |
| 23270 | TSPYL4 | 0.00012 | 0.615454 | 0.605569 | 0.624827 | 0.615283 | down |
| 1798 | DPAGT1 | 0.00012 | 0.523484 | 0.507409 | 0.600898 | 0.54393 | down |
| 5591 | PRKDC | 0.000121 | 0.523234 | 0.582576 | 0.555076 | 0.553629 | down |
| 90933 | TRIM41 | 0.000121 | 0.664779 | 0.580217 | 0.584381 | 0.609792 | down |
| 55114 | ARHGAP1˜ | 0.000122 | 0.612254 | 0.654557 | 0.629006 | 0.631939 | down |
| 5770 | PTPN1 | 0.000123 | 0.616544 | 0.645963 | 0.653003 | 0.638504 | down |
| 51629 | SLC25A39 | 0.000124 | 0.618503 | 0.602611 | 0.662878 | 0.627997 | down |
| 122416 | ANKRD9 | 0.000124 | 0.605802 | 0.620393 | 0.570553 | 0.598916 | down |
| 55650 | PIGV | 0.000124 | 0.626307 | 0.677026 | 0.608564 | 0.637299 | down |
| 28232 | SLCO3A1 | 0.000125 | 0.579349 | 0.568392 | 0.530712 | 0.559484 | down |

| | | | | | | | |
|---|---|---|---|---|---|---|---|
| 63925 | ZNF335 | 0.000125 | 0.627123 | 0.608134 | 0.530182 | 0.58848 | down |
| 51307 | FAM53C | 0.000126 | 0.51554 | 0.606737 | 0.578563 | 0.566947 | down |
| 9665 | MARF1 | 0.000127 | 0.578551 | 0.642708 | 0.61553 | 0.612263 | down |
| 10318 | TNIP1 | 0.000127 | 0.619701 | 0.644407 | 0.633683 | 0.632597 | down |
| 166647 | ADGRA3 | 0.000127 | 0.603877 | 0.609332 | 0.590814 | 0.601341 | down |
| 26470 | SEZ6L2 | 0.000127 | 0.660636 | 0.585017 | 0.667137 | 0.637597 | down |
| 54507 | ADAMTSL4 | 0.000128 | 0.525849 | 0.421199 | 0.532231 | 0.493093 | down |
| 56895 | AGPAT4 | 0.000128 | 0.474747 | 0.421866 | 0.537041 | 0.477885 | down |
| 9921 | RNF10 | 0.000128 | 0.674339 | 0.690168 | 0.622686 | 0.662398 | down |
| 121665 | SPPL3 | 0.000129 | 0.650897 | 0.59049 | 0.55991 | 0.600432 | down |
| 79050 | NOC4L | 0.000129 | 0.66182 | 0.632067 | 0.595591 | 0.629826 | down |
| 9875 | URB1 | 0.000129 | 0.570314 | 0.672345 | 0.600918 | 0.614526 | down |
| 6541 | SLC7A1 | 0.000129 | 0.61566 | 0.670279 | 0.63614 | 0.640693 | down |
| 22822 | PHLDA1 | 0.000129 | 0.532581 | 0.621698 | 0.530373 | 0.561551 | down |
| 4774 | NFIA | 0.000131 | 0.591295 | 0.634778 | 0.596764 | 0.607612 | down |
| 2068 | ERCC2 | 0.000131 | 0.631191 | 0.640227 | 0.638986 | 0.636801 | down |
| 57111 | RAB25 | 0.000131 | 0.667393 | 0.673929 | 0.614133 | 0.651818 | down |
| 6768 | ST14 | 0.000131 | 0.570318 | 0.560509 | 0.499689 | 0.543505 | down |
| 144501 | KRT80 | 0.000131 | 0.545642 | 0.486069 | 0.473232 | 0.501648 | down |
| 1183 | CLCN4 | 0.000132 | 0.588378 | 0.617422 | 0.567452 | 0.591084 | down |
| 284352 | PPP1R37 | 0.000133 | 0.648474 | 0.588972 | 0.635677 | 0.624374 | down |
| 5724 | PTAFR | 0.000133 | 0.5446 | 0.581786 | 0.557406 | 0.561264 | down |
| 81857 | MED25 | 0.000134 | 0.635745 | 0.593449 | 0.647782 | 0.625658 | down |
| 80775 | TMEM177 | 0.000134 | 0.634466 | 0.680707 | 0.651765 | 0.655646 | down |
| 64748 | PLPPR2 | 0.000134 | 0.584607 | 0.512839 | 0.558749 | 0.552065 | down |
| 56970 | ATXN7L3 | 0.000134 | 0.602545 | 0.585055 | 0.595903 | 0.594501 | down |
| 340385 | ZNF517 | 0.000135 | 0.483022 | 0.569251 | 0.527272 | 0.526515 | down |
| 84726 | PRRC2B | 0.000135 | 0.557092 | 0.613485 | 0.600815 | 0.590464 | down |
| 1388 | ATF6B | 0.000135 | 0.578535 | 0.532433 | 0.586941 | 0.56597 | down |
| 29 | ABR | 0.000136 | 0.569823 | 0.591883 | 0.53549 | 0.565732 | down |
| 9883 | POM121 | 0.000136 | 0.639455 | 0.563474 | 0.526593 | 0.576507 | down |
| 311 | ANXA11 | 0.000137 | 0.677341 | 0.633637 | 0.655999 | 0.655659 | down |
| 23399 | CTDNEP1 | 0.000137 | 0.636639 | 0.544909 | 0.549115 | 0.576888 | down |
| 4171 | MCM2 | 0.000137 | 0.680466 | 0.66501 | 0.592128 | 0.645868 | down |
| 63922 | CHTF18 | 0.000137 | 0.661696 | 0.568735 | 0.561183 | 0.597205 | down |
| 90060 | CCDC120 | 0.000137 | 0.672866 | 0.620327 | 0.668338 | 0.653844 | down |
| 1952 | CELSR2 | 0.000138 | 0.581206 | 0.595513 | 0.537747 | 0.571489 | down |
| 317762 | CCDC85C | 0.000139 | 0.610785 | 0.641872 | 0.541942 | 0.5982 | down |
| 1848 | DUSP6 | 0.000139 | 0.562387 | 0.54083 | 0.573472 | 0.558896 | down |
| 54863 | TOR4A | 0.000139 | 0.572264 | 0.498107 | 0.535546 | 0.535306 | down |
| 259173 | ALS2CL | 0.000139 | 0.565133 | 0.574477 | 0.548023 | 0.562544 | down |
| 56262 | LRRC8A | 0.00014 | 0.615952 | 0.673212 | 0.658733 | 0.649299 | down |
| 115704 | EVI5L | 0.00014 | 0.574042 | 0.520586 | 0.589147 | 0.561258 | down |
| 6787 | NEK4 | 0.00014 | 0.599029 | 0.622256 | 0.667796 | 0.629694 | down |
| 23203 | PMPCA | 0.000141 | 0.657401 | 0.640387 | 0.593205 | 0.630331 | down |
| 210 | ALAD | 0.000141 | 0.607063 | 0.552936 | 0.65085 | 0.603616 | down |
| 94005 | PIGS | 0.000141 | 0.530952 | 0.613942 | 0.598761 | 0.581218 | down |
| 10432 | RBM14 | 0.000142 | 0.638979 | 0.577214 | 0.546939 | 0.587711 | down |
| 1107 | CHD3 | 0.000142 | 0.635754 | 0.689192 | 0.640624 | 0.65519 | down |
| 6624 | FSCN1 | 0.000143 | 0.577304 | 0.582881 | 0.537516 | 0.5659 | down |
| 2002 | ELK1 | 0.000144 | 0.59937 | 0.643544 | 0.574264 | 0.605726 | down |
| 3373 | HYAL1 | 0.000145 | 0.680065 | 0.614745 | 0.580401 | 0.62507 | down |
| 283373 | ANKRD52 | 0.000145 | 0.657836 | 0.663241 | 0.66379 | 0.661623 | down |
| 2030 | SLC29A1 | 0.000145 | 0.610088 | 0.531612 | 0.58604 | 0.575913 | down |
| 808 | CALM3 | 0.000146 | 0.655788 | 0.625763 | 0.642561 | 0.641371 | down |
| 677 | ZFP36L1 | 0.000147 | 0.644029 | 0.664474 | 0.644801 | 0.651101 | down |
| 23648 | SSBP3 | 0.000147 | 0.621313 | 0.558509 | 0.632984 | 0.604269 | down |
| 5655 | KLK10 | 0.000147 | 0.566728 | 0.523732 | 0.525575 | 0.538678 | down |

| | | | | | | | |
|---|---|---|---|---|---|---|---|
| 51222 | ZNF219 | 0.000147 | 0.607689 | 0.677645 | 0.572493 | 0.619276 | down |
| 90204 | ZSWIM1 | 0.000148 | 0.41933 | 0.431948 | 0.516769 | 0.456016 | down |
| 23308 | ICOSLG | 0.000148 | 0.534109 | 0.514727 | 0.459432 | 0.502756 | down |
| 51236 | HGH1 | 0.000149 | 0.600386 | 0.645752 | 0.619279 | 0.621806 | down |
| 84717 | HDGFL2 | 0.00015 | 0.632842 | 0.648718 | 0.569548 | 0.617036 | down |
| 11060 | WWP2 | 0.00015 | 0.640822 | 0.629997 | 0.620553 | 0.630457 | down |
| 64118 | DUS1L | 0.000152 | 0.690148 | 0.643744 | 0.613905 | 0.649266 | down |
| 9203 | ZMYM3 | 0.000152 | 0.52537 | 0.587466 | 0.621126 | 0.577987 | down |
| 10712 | FAM189B | 0.000152 | 0.577016 | 0.540247 | 0.592256 | 0.56984 | down |
| 5331 | PLCB3 | 0.000152 | 0.536101 | 0.605772 | 0.604448 | 0.582107 | down |
| 144100 | PLEKHA7 | 0.000152 | 0.560935 | 0.635083 | 0.57828 | 0.591433 | down |
| 56926 | NCLN | 0.000152 | 0.655609 | 0.700142 | 0.632088 | 0.662613 | down |
| 84287 | ZDHHC16 | 0.000152 | 0.635003 | 0.596564 | 0.637729 | 0.623098 | down |
| 1.01E+08 | TIMM23B | 0.000153 | 0.678956 | 0.589833 | 0.669516 | 0.646102 | down |
| 64856 | VWA1 | 0.000153 | 0.687414 | 0.674613 | 0.593428 | 0.651819 | down |
| 114818 | KLHL29 | 0.000153 | 0.56661 | 0.632791 | 0.536464 | 0.578622 | down |
| 10025 | MED16 | 0.000153 | 0.570352 | 0.4772 | 0.452346 | 0.499966 | down |
| 3732 | CD82 | 0.000154 | 0.643537 | 0.596846 | 0.539035 | 0.593139 | down |
| 7259 | TSPYL1 | 0.000154 | 0.610028 | 0.693695 | 0.643997 | 0.64924 | down |
| 955 | ENTPD6 | 0.000154 | 0.609192 | 0.574443 | 0.638253 | 0.607296 | down |
| 254531 | LPCAT4 | 0.000155 | 0.439614 | 0.487644 | 0.52269 | 0.483316 | down |
| 116988 | AGAP3 | 0.000155 | 0.565972 | 0.633129 | 0.579742 | 0.592948 | down |
| 285440 | CYP4V2 | 0.000155 | 0.456956 | 0.482299 | 0.585941 | 0.508399 | down |
| 93109 | TMEM44 | 0.000156 | 0.682858 | 0.600667 | 0.598312 | 0.627279 | down |
| 57449 | PLEKHG5 | 0.000156 | 0.424495 | 0.374536 | 0.520964 | 0.439998 | down |
| 8295 | TRRAP | 0.000157 | 0.514352 | 0.599048 | 0.546078 | 0.553159 | down |
| 10769 | PLK2 | 0.000157 | 0.593394 | 0.649963 | 0.592551 | 0.611969 | down |
| 79098 | C1orf116 | 0.000157 | 0.580677 | 0.650163 | 0.600595 | 0.610478 | down |
| 23406 | COTL1 | 0.000158 | 0.674942 | 0.645912 | 0.607516 | 0.64279 | down |
| 4645 | MYO5B | 0.000158 | 0.465848 | 0.577075 | 0.478369 | 0.507097 | down |
| 582 | BBS1 | 0.000159 | 0.567666 | 0.529763 | 0.552708 | 0.550046 | down |
| 1.05E+08 | LOC10537 | 0.000159 | 0.353702 | 0.397968 | 0.36144 | 0.371036 | down |
| 25966 | C2CD2 | 0.000159 | 0.574859 | 0.642035 | 0.60277 | 0.606555 | down |
| 79789 | CLMN | 0.000159 | 0.632295 | 0.572632 | 0.548485 | 0.584471 | down |
| 3304 | HSPA1B | 0.000159 | 0.593967 | 0.655968 | 0.539667 | 0.596534 | down |
| 79084 | WDR77 | 0.00016 | 0.707804 | 0.608218 | 0.643053 | 0.653025 | down |
| 124930 | ANKRD13E | 0.00016 | 0.563153 | 0.525882 | 0.556055 | 0.548363 | down |
| 9922 | IQSEC1 | 0.00016 | 0.599269 | 0.542749 | 0.588431 | 0.576816 | down |
| 55224 | ETNK2 | 0.00016 | 0.604479 | 0.681313 | 0.579862 | 0.621885 | down |
| 2318 | FLNC | 0.000161 | 0.650022 | 0.699093 | 0.640421 | 0.663179 | down |
| 9294 | S1PR2 | 0.000161 | 0.505337 | 0.61601 | 0.562839 | 0.561395 | down |
| 149076 | ZNF362 | 0.000162 | 0.633338 | 0.682804 | 0.635835 | 0.650659 | down |
| 57498 | KIDINS220 | 0.000162 | 0.671083 | 0.70158 | 0.619792 | 0.664152 | down |
| 55160 | ARHGEF1C | 0.000162 | 0.541343 | 0.534989 | 0.550754 | 0.542362 | down |
| 8651 | SOCS1 | 0.000163 | 0.440558 | 0.559497 | 0.424636 | 0.474897 | down |
| 91828 | EXOC3L4 | 0.000163 | 0.272727 | 0.370968 | 0.406349 | 0.350015 | down |
| 2975 | GTF3C1 | 0.000163 | 0.55434 | 0.627967 | 0.543212 | 0.575173 | down |
| 116113 | FOXP4 | 0.000164 | 0.604006 | 0.548876 | 0.536108 | 0.562997 | down |
| 55684 | RABL6 | 0.000165 | 0.562944 | 0.585077 | 0.523171 | 0.557064 | down |
| 64781 | CERK | 0.000166 | 0.620556 | 0.702796 | 0.643673 | 0.655675 | down |
| 23348 | DOCK9 | 0.000166 | 0.592303 | 0.648122 | 0.623452 | 0.621292 | down |
| 64847 | SPATA20 | 0.000166 | 0.613604 | 0.541898 | 0.596969 | 0.584157 | down |
| 1160 | CKMT2 | 0.000167 | 0.51378 | 0.533129 | 0.55273 | 0.533213 | down |
| 9825 | SPATA2 | 0.000167 | 0.503719 | 0.606638 | 0.50955 | 0.539969 | down |
| 4800 | NFYA | 0.000167 | 0.669482 | 0.699523 | 0.622115 | 0.663707 | down |
| 57473 | ZNF512B | 0.000168 | 0.587933 | 0.678408 | 0.647335 | 0.637892 | down |
| 9706 | ULK2 | 0.000168 | 0.614962 | 0.66856 | 0.63825 | 0.640591 | down |
| 29894 | CPSF1 | 0.000168 | 0.617286 | 0.666392 | 0.551639 | 0.611772 | down |

| | | | | | | | |
|---|---|---|---|---|---|---|---|
| 22846 | VASH1 | 0.000168 | 0.636752 | 0.588043 | 0.554204 | 0.593 | down |
| 11113 | CIT | 0.000169 | 0.584807 | 0.632384 | 0.608713 | 0.608635 | down |
| 57148 | RALGAPB | 0.000169 | 0.649892 | 0.709567 | 0.639594 | 0.666351 | down |
| 134549 | SHROOM1 | 0.00017 | 0.623772 | 0.568089 | 0.522503 | 0.571455 | down |
| 84725 | PLEKHA8 | 0.000172 | 0.672144 | 0.650166 | 0.59425 | 0.638853 | down |
| 10391 | CORO2B | 0.000173 | 0.631412 | 0.538431 | 0.592215 | 0.587353 | down |
| 55756 | INTS9 | 0.000173 | 0.622159 | 0.589826 | 0.69569 | 0.635892 | down |
| 23654 | PLXNB2 | 0.000173 | 0.614677 | 0.641683 | 0.616664 | 0.624341 | down |
| 4303 | FOXO4 | 0.000174 | 0.542691 | 0.489826 | 0.479194 | 0.503904 | down |
| 23277 | CLUH | 0.000175 | 0.527727 | 0.608011 | 0.570711 | 0.568816 | down |
| 7862 | BRPF1 | 0.000176 | 0.627273 | 0.721722 | 0.638729 | 0.662575 | down |
| 9491 | PSMF1 | 0.000177 | 0.678609 | 0.638144 | 0.639587 | 0.652113 | down |
| 399512 | SLC25A35 | 0.000177 | 0.589265 | 0.519883 | 0.45938 | 0.522843 | down |
| 64777 | RMND5B | 0.000177 | 0.622348 | 0.509422 | 0.566398 | 0.566056 | down |
| 8208 | CHAF1B | 0.000178 | 0.635664 | 0.697871 | 0.663587 | 0.665707 | down |
| 6398 | SECTM1 | 0.000179 | 0.433035 | 0.484854 | 0.53756 | 0.48515 | down |
| 23381 | SMG5 | 0.000179 | 0.617498 | 0.674439 | 0.620594 | 0.63751 | down |
| 10509 | SEMA4B | 0.00018 | 0.638673 | 0.620612 | 0.528066 | 0.595783 | down |
| 163732 | CITED4 | 0.00018 | 0.53022 | 0.497119 | 0.432369 | 0.486569 | down |
| 643680 | MS4A4E | 0.00018 | 0.684654 | 0.583856 | 0.595185 | 0.621232 | down |
| 57585 | CRAMP1 | 0.000181 | 0.639817 | 0.643507 | 0.628202 | 0.637175 | down |
| 283358 | B4GALNT3 | 0.000183 | 0.503448 | 0.492419 | 0.391164 | 0.462344 | down |
| 642475 | MROH6 | 0.000183 | 0.475554 | 0.564973 | 0.472561 | 0.504363 | down |
| 3664 | IRF6 | 0.000184 | 0.620416 | 0.683546 | 0.626107 | 0.643356 | down |
| 7621 | ZNF70 | 0.000184 | 0.488226 | 0.416448 | 0.52958 | 0.478085 | down |
| 7020 | TFAP2A | 0.000184 | 0.596744 | 0.621448 | 0.596721 | 0.604971 | down |
| 124583 | CANT1 | 0.000184 | 0.617783 | 0.636362 | 0.634681 | 0.629608 | down |
| 65095 | KRI1 | 0.000185 | 0.648705 | 0.656442 | 0.590407 | 0.631851 | down |
| 84836 | ABHD14B | 0.000185 | 0.609843 | 0.491836 | 0.592074 | 0.564584 | down |
| 81876 | RAB1B | 0.000185 | 0.656309 | 0.633096 | 0.691331 | 0.660245 | down |
| 23522 | KAT6B | 0.000185 | 0.55968 | 0.633123 | 0.659798 | 0.617534 | down |
| 9605 | VPS9D1 | 0.000186 | 0.590615 | 0.655261 | 0.57701 | 0.607629 | down |
| 55652 | SLC48A1 | 0.000186 | 0.444985 | 0.516356 | 0.504546 | 0.488629 | down |
| 5792 | PTPRF | 0.000187 | 0.610392 | 0.665827 | 0.611686 | 0.629302 | down |
| 9679 | FAM53B | 0.000188 | 0.508621 | 0.587121 | 0.513387 | 0.536376 | down |
| 9759 | HDAC4 | 0.000188 | 0.556909 | 0.597904 | 0.537892 | 0.564235 | down |
| 221927 | BRAT1 | 0.000188 | 0.657515 | 0.561055 | 0.607495 | 0.608688 | down |
| 7108 | TM7SF2 | 0.000188 | 0.661459 | 0.624683 | 0.573269 | 0.619803 | down |
| 11083 | DIDO1 | 0.000189 | 0.580438 | 0.63886 | 0.588349 | 0.602549 | down |
| 54919 | DNAAF5 | 0.00019 | 0.611872 | 0.665103 | 0.559815 | 0.612263 | down |
| 7629 | ZNF76 | 0.00019 | 0.638869 | 0.628669 | 0.573234 | 0.61359 | down |
| 291 | SLC25A4 | 0.000191 | 0.575433 | 0.616854 | 0.662176 | 0.618154 | down |
| 360 | AQP3 | 0.000191 | 0.484232 | 0.435376 | 0.421903 | 0.44717 | down |
| 7076 | TIMP1 | 0.000191 | 0.654157 | 0.591745 | 0.638046 | 0.627983 | down |
| 84444 | DOT1L | 0.000191 | 0.574891 | 0.583544 | 0.547829 | 0.568755 | down |
| 10013 | HDAC6 | 0.000192 | 0.559657 | 0.645807 | 0.53062 | 0.578695 | down |
| 1778 | DYNC1H1 | 0.000193 | 0.584688 | 0.674453 | 0.605823 | 0.621655 | down |
| 8218 | CLTCL1 | 0.000194 | 0.656874 | 0.5525 | 0.600185 | 0.603186 | down |
| 57555 | NLGN2 | 0.000194 | 0.592058 | 0.525217 | 0.538676 | 0.551984 | down |
| 22937 | SCAP | 0.000195 | 0.553505 | 0.585872 | 0.534283 | 0.557886 | down |
| 85455 | DISP2 | 0.000195 | 0.613462 | 0.534734 | 0.577174 | 0.575123 | down |
| 4123 | MAN2C1 | 0.000195 | 0.546755 | 0.532169 | 0.429921 | 0.502948 | down |
| 338442 | HCAR2 | 0.000195 | 0.563878 | 0.594705 | 0.557067 | 0.571883 | down |
| 6293 | VPS52 | 0.000195 | 0.590873 | 0.650265 | 0.573149 | 0.604762 | down |
| 2013 | EMP2 | 0.000196 | 0.59313 | 0.656824 | 0.588049 | 0.612668 | down |
| 10019 | SH2B3 | 0.000196 | 0.519676 | 0.56987 | 0.530921 | 0.540156 | down |
| 1839 | HBEGF | 0.000197 | 0.477473 | 0.604235 | 0.553114 | 0.544941 | down |
| 54552 | GNL3L | 0.000198 | 0.623844 | 0.693217 | 0.629258 | 0.648773 | down |

| | | | | | | | |
|---|---|---|---|---|---|---|---|
| 5493 | PPL | 0.000198 | 0.652058 | 0.646059 | 0.68193 | 0.660016 | down |
| 5738 | PTGFRN | 0.000199 | 0.648372 | 0.703277 | 0.630602 | 0.660751 | down |
| 57720 | GPR107 | 0.000199 | 0.640886 | 0.638058 | 0.605712 | 0.628219 | down |
| 55199 | FAM86C1 | 0.000199 | 0.573432 | 0.594178 | 0.61343 | 0.59368 | down |
| 9675 | TTI1 | 0.000201 | 0.563647 | 0.682757 | 0.615493 | 0.620633 | down |
| 85451 | UNK | 0.000201 | 0.648448 | 0.59236 | 0.524112 | 0.588307 | down |
| 64601 | VPS16 | 0.000202 | 0.663754 | 0.694639 | 0.616995 | 0.658463 | down |
| 84958 | SYTL1 | 0.000202 | 0.497006 | 0.533199 | 0.501277 | 0.510494 | down |
| 54434 | SSH1 | 0.000203 | 0.61219 | 0.672531 | 0.680547 | 0.655089 | down |
| 90120 | TMEM250 | 0.000204 | 0.583146 | 0.52838 | 0.572083 | 0.561203 | down |
| 146059 | CDAN1 | 0.000205 | 0.622828 | 0.595238 | 0.62206 | 0.613376 | down |
| 80381 | CD276 | 0.000205 | 0.614105 | 0.544041 | 0.624408 | 0.594185 | down |
| 79068 | FTO | 0.000207 | 0.611628 | 0.689082 | 0.604247 | 0.634986 | down |
| 8907 | AP1M1 | 0.000207 | 0.719796 | 0.613716 | 0.653288 | 0.662267 | down |
| 27013 | CNPPD1 | 0.000207 | 0.609842 | 0.599677 | 0.557068 | 0.588862 | down |
| 9620 | CELSR1 | 0.000207 | 0.56931 | 0.610872 | 0.576636 | 0.585606 | down |
| 1820 | ARID3A | 0.000208 | 0.71236 | 0.645212 | 0.640473 | 0.666015 | down |
| 1352 | COX10 | 0.000208 | 0.56799 | 0.662613 | 0.667346 | 0.63265 | down |
| 8555 | CDC14B | 0.000209 | 0.574865 | 0.689746 | 0.618419 | 0.627677 | down |
| 55238 | SLC38A7 | 0.000209 | 0.618569 | 0.662876 | 0.630731 | 0.637392 | down |
| 6556 | SLC11A1 | 0.00021 | 0.4 | 0.326531 | 0.375 | 0.367177 | down |
| 5524 | PTPA | 0.00021 | 0.629004 | 0.606113 | 0.5722 | 0.602439 | down |
| 8863 | PER3 | 0.00021 | 0.580716 | 0.482509 | 0.499129 | 0.520785 | down |
| 8612 | PLPP2 | 0.000211 | 0.622404 | 0.706291 | 0.60497 | 0.644555 | down |
| 7074 | TIAM1 | 0.000211 | 0.634462 | 0.585695 | 0.570313 | 0.596823 | down |
| 284110 | GSDMA | 0.000211 | 0.479715 | 0.412018 | 0.39615 | 0.429294 | down |
| 55201 | MAP1S | 0.000211 | 0.517484 | 0.506822 | 0.553365 | 0.52589 | down |
| 8402 | SLC25A11 | 0.000211 | 0.660678 | 0.690988 | 0.64212 | 0.664595 | down |
| 3913 | LAMB2 | 0.000211 | 0.541215 | 0.57244 | 0.504344 | 0.539333 | down |
| 56848 | SPHK2 | 0.000213 | 0.632586 | 0.613201 | 0.526263 | 0.590684 | down |
| 114822 | RHPN1 | 0.000213 | 0.397054 | 0.424343 | 0.401223 | 0.40754 | down |
| 9536 | PTGES | 0.000214 | 0.626337 | 0.70601 | 0.599704 | 0.644017 | down |
| 266655 | BRD3OS | 0.000214 | 0.606252 | 0.685483 | 0.6353 | 0.642345 | down |
| 3892 | KRT86 | 0.000214 | 0.361864 | 0.475336 | 0.469251 | 0.435484 | down |
| 23764 | MAFF | 0.000216 | 0.579722 | 0.595514 | 0.589077 | 0.588104 | down |
| 57190 | SELENON | 0.000216 | 0.519004 | 0.541272 | 0.498191 | 0.519489 | down |
| 55893 | ZNF395 | 0.000216 | 0.557489 | 0.65972 | 0.617916 | 0.611708 | down |
| 79415 | CYBC1 | 0.000218 | 0.639142 | 0.593788 | 0.537051 | 0.589993 | down |
| 5681 | PSKH1 | 0.00022 | 0.489626 | 0.503804 | 0.488466 | 0.493965 | down |
| 375346 | STIMATE | 0.000221 | 0.55209 | 0.611951 | 0.552327 | 0.572122 | down |
| 1468 | SLC25A10 | 0.000221 | 0.583124 | 0.581651 | 0.662956 | 0.609244 | down |
| 146909 | KIF18B | 0.000222 | 0.666714 | 0.689408 | 0.611173 | 0.655765 | down |
| 152992 | TRMT44 | 0.000223 | 0.631277 | 0.652619 | 0.628276 | 0.637391 | down |
| 11270 | NRM | 0.000225 | 0.602091 | 0.506233 | 0.550335 | 0.552886 | down |
| 80004 | ESRP2 | 0.000225 | 0.619326 | 0.576381 | 0.648733 | 0.614813 | down |
| 10782 | ZNF274 | 0.000226 | 0.54872 | 0.55292 | 0.640986 | 0.580875 | down |
| 10075 | HUWE1 | 0.000226 | 0.609865 | 0.637492 | 0.649076 | 0.632145 | down |
| 79990 | PLEKHH3 | 0.000227 | 0.592013 | 0.594847 | 0.60032 | 0.595727 | down |
| 752 | FMNL1 | 0.000227 | 0.454633 | 0.555894 | 0.533064 | 0.51453 | down |
| 1E+08 | POM121C | 0.000228 | 0.626089 | 0.588633 | 0.632582 | 0.615768 | down |
| 29924 | EPN1 | 0.000228 | 0.57423 | 0.500159 | 0.555453 | 0.543281 | down |
| 79867 | TCTN2 | 0.000229 | 0.568237 | 0.611269 | 0.550021 | 0.576509 | down |
| 6892 | TAPBP | 0.00023 | 0.507358 | 0.611286 | 0.555607 | 0.558084 | down |
| 54617 | INO80 | 0.000231 | 0.625278 | 0.643758 | 0.656982 | 0.642006 | down |
| 1969 | EPHA2 | 0.000232 | 0.613707 | 0.510983 | 0.491412 | 0.538701 | down |
| 3423 | IDS | 0.000232 | 0.558818 | 0.569016 | 0.558557 | 0.56213 | down |
| 57466 | SCAF4 | 0.000232 | 0.5893 | 0.612736 | 0.610501 | 0.604179 | down |
| 27154 | BRPF3 | 0.000233 | 0.628334 | 0.666576 | 0.675856 | 0.656922 | down |

| | | | | | | | |
|---|---|---|---|---|---|---|---|
| 55205 | ZNF532 | 0.000235 | 0.656657 | 0.638541 | 0.541108 | 0.612102 | down |
| 55806 | HR | 0.000236 | 0.658565 | 0.657017 | 0.597298 | 0.637627 | down |
| 8542 | APOL1 | 0.000236 | 0.581992 | 0.634538 | 0.690021 | 0.635517 | down |
| 8396 | PIP4K2B | 0.000237 | 0.593915 | 0.683235 | 0.574148 | 0.617099 | down |
| 57326 | PBXIP1 | 0.000238 | 0.644215 | 0.650715 | 0.583095 | 0.626008 | down |
| 7159 | TP53BP2 | 0.000238 | 0.652098 | 0.593263 | 0.679224 | 0.641528 | down |
| 92999 | ZBTB47 | 0.000239 | 0.59194 | 0.580971 | 0.574273 | 0.582395 | down |
| 83696 | TRAPPC9 | 0.000239 | 0.476888 | 0.593289 | 0.499495 | 0.523224 | down |
| 116986 | AGAP2 | 0.000241 | 0.661601 | 0.681712 | 0.614114 | 0.652475 | down |
| 8289 | ARID1A | 0.000245 | 0.563271 | 0.672701 | 0.629658 | 0.621877 | down |
| 10555 | AGPAT2 | 0.000245 | 0.666417 | 0.617889 | 0.625002 | 0.636436 | down |
| 51540 | SCLY | 0.000246 | 0.582555 | 0.651339 | 0.656057 | 0.629984 | down |
| 10847 | SRCAP | 0.000246 | 0.619534 | 0.649134 | 0.598117 | 0.622262 | down |
| 84892 | POMGNT2 | 0.000246 | 0.535295 | 0.522313 | 0.599129 | 0.552246 | down |
| 1059 | CENPB | 0.000247 | 0.629056 | 0.616911 | 0.534765 | 0.593577 | down |
| 51246 | SHISA5 | 0.000248 | 0.554745 | 0.49662 | 0.535358 | 0.528908 | down |
| 5002 | SLC22A18 | 0.000248 | 0.505922 | 0.480663 | 0.565013 | 0.5172 | down |
| 92305 | TMEM129 | 0.00025 | 0.595444 | 0.591701 | 0.5915 | 0.592882 | down |
| 9772 | TMEM94 | 0.000251 | 0.637905 | 0.552939 | 0.537731 | 0.576192 | down |
| 81490 | PTDSS2 | 0.000251 | 0.624315 | 0.64902 | 0.618851 | 0.630729 | down |
| 286077 | FAM83H | 0.000252 | 0.614433 | 0.636886 | 0.552272 | 0.601197 | down |
| 79977 | GRHL2 | 0.000252 | 0.625677 | 0.684367 | 0.582411 | 0.630819 | down |
| 493 | ATP2B4 | 0.000253 | 0.56188 | 0.676239 | 0.635514 | 0.624544 | down |
| 149175 | MANEAL | 0.000254 | 0.542984 | 0.621668 | 0.635419 | 0.600024 | down |
| 399687 | MYO18A | 0.000254 | 0.402306 | 0.493598 | 0.466706 | 0.454203 | down |
| 81619 | TSPAN14 | 0.000254 | 0.615049 | 0.617001 | 0.606846 | 0.612965 | down |
| 55809 | TRERF1 | 0.000254 | 0.502817 | 0.632123 | 0.50823 | 0.547724 | down |
| 57592 | ZNF687 | 0.000255 | 0.619776 | 0.687365 | 0.603703 | 0.636948 | down |
| 84971 | ATG4D | 0.000255 | 0.675338 | 0.644135 | 0.571858 | 0.630443 | down |
| 9909 | DENND4B | 0.000256 | 0.679735 | 0.582385 | 0.57776 | 0.613293 | down |
| 23140 | ZZEF1 | 0.000256 | 0.597821 | 0.660401 | 0.570915 | 0.609712 | down |
| 57657 | HCN3 | 0.000257 | 0.636364 | 0.585028 | 0.580583 | 0.600658 | down |
| 22845 | DOLK | 0.000257 | 0.588841 | 0.62665 | 0.675174 | 0.630222 | down |
| 159090 | FAM122B | 0.000257 | 0.54563 | 0.670073 | 0.619548 | 0.61175 | down |
| 9578 | CDC42BPB | 0.000257 | 0.680183 | 0.668985 | 0.602159 | 0.650442 | down |
| 79144 | PPDPF | 0.000259 | 0.56305 | 0.485417 | 0.569089 | 0.539185 | down |
| 404093 | CUEDC1 | 0.000259 | 0.563987 | 0.584584 | 0.482454 | 0.543675 | down |
| 10505 | SEMA4F | 0.000261 | 0.61981 | 0.665408 | 0.570121 | 0.618446 | down |
| 5613 | PRKX | 0.000261 | 0.528581 | 0.580146 | 0.653996 | 0.587575 | down |
| 399909 | PCNX3 | 0.000265 | 0.596543 | 0.639417 | 0.623203 | 0.619721 | down |
| 24147 | FJX1 | 0.000265 | 0.484695 | 0.524424 | 0.403585 | 0.470902 | down |
| 157638 | FAM84B | 0.000266 | 0.663267 | 0.672589 | 0.659233 | 0.66503 | down |
| 22897 | CEP164 | 0.000266 | 0.656424 | 0.565263 | 0.667432 | 0.629706 | down |
| 22994 | CEP131 | 0.000266 | 0.677327 | 0.684274 | 0.598812 | 0.653471 | down |
| 1140 | CHRNB1 | 0.000268 | 0.630319 | 0.599373 | 0.661667 | 0.630453 | down |
| 5875 | RABGGTA | 0.000269 | 0.523561 | 0.649658 | 0.530928 | 0.568049 | down |
| 257364 | SNX33 | 0.000269 | 0.578645 | 0.649652 | 0.695731 | 0.641343 | down |
| 9394 | HS6ST1 | 0.000269 | 0.655451 | 0.665885 | 0.622076 | 0.647804 | down |
| 9757 | KMT2B | 0.000269 | 0.54599 | 0.619444 | 0.659003 | 0.608146 | down |
| 23373 | CRTC1 | 0.00027 | 0.586975 | 0.518051 | 0.574338 | 0.559788 | down |
| 124359 | CDYL2 | 0.000271 | 0.613619 | 0.683136 | 0.564563 | 0.62044 | down |
| 8607 | RUVBL1 | 0.000271 | 0.661453 | 0.702754 | 0.609635 | 0.657948 | down |
| 23094 | SIPA1L3 | 0.000271 | 0.575321 | 0.56902 | 0.517901 | 0.554081 | down |
| 57605 | PITPNM2 | 0.000271 | 0.553723 | 0.549628 | 0.561667 | 0.555006 | down |
| 57619 | SHROOM3 | 0.000271 | 0.650947 | 0.631288 | 0.703024 | 0.661753 | down |
| 5467 | PPARD | 0.000271 | 0.55332 | 0.556855 | 0.619343 | 0.576506 | down |
| 64427 | TTC31 | 0.000272 | 0.526839 | 0.51217 | 0.505908 | 0.514972 | down |
| 5900 | RALGDS | 0.000272 | 0.529622 | 0.513432 | 0.532584 | 0.525212 | down |

| | | | | | | | |
|---|---|---|---|---|---|---|---|
| 7011 | TEP1 | 0.000272 | 0.402257 | 0.532936 | 0.424479 | 0.453224 | down |
| 1201 | CLN3 | 0.000272 | 0.662853 | 0.600635 | 0.615408 | 0.626298 | down |
| 55612 | FERMT1 | 0.000272 | 0.622899 | 0.685615 | 0.665304 | 0.65794 | down |
| 4055 | LTBR | 0.000275 | 0.686369 | 0.606283 | 0.652795 | 0.648482 | down |
| 342897 | NCCRP1 | 0.000276 | 0.465767 | 0.482464 | 0.380835 | 0.443022 | down |
| 4209 | MEF2D | 0.000276 | 0.591291 | 0.578852 | 0.643181 | 0.604441 | down |
| 56886 | UGGT1 | 0.000276 | 0.62111 | 0.680731 | 0.646045 | 0.649296 | down |
| 9889 | ZBED4 | 0.000278 | 0.58638 | 0.688367 | 0.602513 | 0.625753 | down |
| 126308 | MOB3A | 0.000279 | 0.420224 | 0.530849 | 0.499916 | 0.483663 | down |
| 109 | ADCY3 | 0.00028 | 0.566515 | 0.483791 | 0.521607 | 0.523971 | down |
| 2063 | NR2F6 | 0.00028 | 0.646206 | 0.630511 | 0.571706 | 0.616141 | down |
| 8131 | NPRL3 | 0.000281 | 0.602404 | 0.594367 | 0.674089 | 0.62362 | down |
| 23164 | MPRIP | 0.000281 | 0.589198 | 0.636551 | 0.57224 | 0.599329 | down |
| 51564 | HDAC7 | 0.000281 | 0.58361 | 0.640945 | 0.6379 | 0.620819 | down |
| 56997 | COQ8A | 0.000283 | 0.5386 | 0.603708 | 0.461064 | 0.534457 | down |
| 219855 | SLC37A2 | 0.000284 | 0.666306 | 0.597001 | 0.56649 | 0.609932 | down |
| 54976 | C20orf27 | 0.000284 | 0.646343 | 0.650656 | 0.630165 | 0.642388 | down |
| 1288 | COL4A6 | 0.000285 | 0.674381 | 0.643199 | 0.565282 | 0.627621 | down |
| 999 | CDH1 | 0.000285 | 0.598098 | 0.615352 | 0.606225 | 0.606558 | down |
| 79152 | FA2H | 0.000285 | 0.521189 | 0.351245 | 0.412712 | 0.428382 | down |
| 7538 | ZFP36 | 0.000286 | 0.592949 | 0.568233 | 0.502663 | 0.554615 | down |
| 374378 | GALNT18 | 0.000288 | 0.596064 | 0.587922 | 0.497323 | 0.560437 | down |
| 113178 | SCAMP4 | 0.000288 | 0.679944 | 0.691282 | 0.613108 | 0.661445 | down |
| 3703 | STT3A | 0.000288 | 0.664295 | 0.604994 | 0.693376 | 0.654222 | down |
| 8448 | DOC2A | 0.000288 | 0.498596 | 0.510552 | 0.591252 | 0.533467 | down |
| 64598 | MOSPD3 | 0.000289 | 0.542775 | 0.57881 | 0.673474 | 0.598353 | down |
| 6665 | SOX15 | 0.000289 | 0.644334 | 0.562835 | 0.612299 | 0.606489 | down |
| 5989 | RFX1 | 0.00029 | 0.609863 | 0.676274 | 0.625425 | 0.637187 | down |
| 25885 | POLR1A | 0.00029 | 0.516734 | 0.627284 | 0.561088 | 0.568368 | down |
| 22873 | DZIP1 | 0.00029 | 0.621958 | 0.718833 | 0.628672 | 0.656488 | down |
| 22904 | SBNO2 | 0.000291 | 0.541314 | 0.566882 | 0.485209 | 0.531135 | down |
| 3693 | ITGB5 | 0.000291 | 0.576361 | 0.585254 | 0.488589 | 0.550068 | down |
| 23403 | FBXO46 | 0.000291 | 0.48722 | 0.537204 | 0.40716 | 0.477195 | down |
| 3064 | HTT | 0.000293 | 0.613462 | 0.634331 | 0.564885 | 0.604226 | down |
| 8263 | F8A1 | 0.000294 | 0.649994 | 0.625288 | 0.648028 | 0.641103 | down |
| 64411 | ARAP3 | 0.000297 | 0.620433 | 0.525253 | 0.485941 | 0.543875 | down |
| 23526 | ARHGAP4! | 0.000299 | 0.612681 | 0.698578 | 0.6078 | 0.639686 | down |
| 6305 | SBF1 | 0.000299 | 0.528845 | 0.568121 | 0.45973 | 0.518899 | down |
| 23759 | PPIL2 | 0.0003 | 0.672574 | 0.639443 | 0.627516 | 0.646511 | down |
| 51684 | SUFU | 0.0003 | 0.693241 | 0.598365 | 0.613403 | 0.635003 | down |
| 602 | BCL3 | 0.000301 | 0.642437 | 0.574145 | 0.548345 | 0.588309 | down |
| 55705 | IPO9 | 0.000302 | 0.614687 | 0.674341 | 0.686781 | 0.658603 | down |
| 2017 | CTTN | 0.000302 | 0.674984 | 0.673923 | 0.587132 | 0.645346 | down |
| 5916 | RARG | 0.000302 | 0.626485 | 0.608728 | 0.671698 | 0.635637 | down |
| 11135 | CDC42EP1 | 0.000303 | 0.647112 | 0.604116 | 0.64715 | 0.632793 | down |
| 2921 | CXCL3 | 0.000304 | 0.670722 | 0.650617 | 0.655442 | 0.658927 | down |
| 220972 | 44263 | 0.000304 | 0.663805 | 0.622885 | 0.624388 | 0.637026 | down |
| 80223 | RAB11FIP1 | 0.000304 | 0.556291 | 0.624873 | 0.569499 | 0.583554 | down |
| 146223 | CMTM4 | 0.000307 | 0.448719 | 0.526491 | 0.543989 | 0.5064 | down |
| 10985 | GCN1 | 0.000307 | 0.523746 | 0.551516 | 0.457614 | 0.510959 | down |
| 22877 | MLXIP | 0.000307 | 0.668945 | 0.625394 | 0.561367 | 0.618569 | down |
| 79671 | NLRX1 | 0.000309 | 0.603398 | 0.516969 | 0.491354 | 0.53724 | down |
| 83933 | HDAC10 | 0.000309 | 0.608535 | 0.550107 | 0.537959 | 0.565534 | down |
| 51703 | ACSL5 | 0.000311 | 0.613597 | 0.517492 | 0.634707 | 0.588599 | down |
| 91949 | COG7 | 0.000311 | 0.672856 | 0.575913 | 0.672284 | 0.640351 | down |
| 8694 | DGAT1 | 0.000312 | 0.622273 | 0.617195 | 0.55752 | 0.598996 | down |
| 132014 | IL17RE | 0.000313 | 0.565502 | 0.61601 | 0.548053 | 0.576522 | down |
| 5293 | PIK3CD | 0.000315 | 0.512105 | 0.563099 | 0.538776 | 0.537993 | down |

| | | | | | | | |
|---|---|---|---|---|---|---|---|
| 80233 | FAAP100 | 0.000316 | 0.620059 | 0.501119 | 0.505001 | 0.54206 | down |
| 51780 | KDM3B | 0.000316 | 0.593857 | 0.686654 | 0.659849 | 0.646787 | down |
| 4047 | LSS | 0.000318 | 0.601567 | 0.70083 | 0.581046 | 0.627814 | down |
| 58489 | ABHD17C | 0.000319 | 0.554141 | 0.586046 | 0.536292 | 0.558827 | down |
| 1870 | E2F2 | 0.000319 | 0.574398 | 0.55612 | 0.633448 | 0.587989 | down |
| 25844 | YIPF3 | 0.00032 | 0.643006 | 0.647827 | 0.662777 | 0.651204 | down |
| 57634 | EP400 | 0.000321 | 0.559424 | 0.653993 | 0.606831 | 0.606749 | down |
| 6573 | SLC19A1 | 0.000323 | 0.5336 | 0.629442 | 0.488394 | 0.550479 | down |
| 56252 | YLPM1 | 0.000324 | 0.65992 | 0.555412 | 0.662228 | 0.625854 | down |
| 9260 | PDLIM7 | 0.000327 | 0.636175 | 0.56918 | 0.633589 | 0.612981 | down |
| 8985 | PLOD3 | 0.000328 | 0.667171 | 0.620017 | 0.565202 | 0.617463 | down |
| 5307 | PITX1 | 0.000329 | 0.66485 | 0.648682 | 0.621483 | 0.645005 | down |
| 54920 | DUS2 | 0.00033 | 0.523178 | 0.653741 | 0.585694 | 0.587538 | down |
| 84922 | FIZ1 | 0.00033 | 0.593761 | 0.503958 | 0.521094 | 0.539604 | down |
| 79733 | E2F8 | 0.00033 | 0.681459 | 0.618261 | 0.61414 | 0.637953 | down |
| 138050 | HGSNAT | 0.000331 | 0.596594 | 0.646047 | 0.657643 | 0.633428 | down |
| 8692 | HYAL2 | 0.000335 | 0.608007 | 0.504186 | 0.52726 | 0.546484 | down |
| 23659 | PLA2G15 | 0.000335 | 0.595525 | 0.662612 | 0.699206 | 0.652448 | down |
| 10868 | USP20 | 0.000339 | 0.525673 | 0.588745 | 0.521698 | 0.545372 | down |
| 2769 | GNA15 | 0.00034 | 0.69002 | 0.644949 | 0.664065 | 0.666345 | down |
| 2997 | GYS1 | 0.000341 | 0.579279 | 0.600011 | 0.48617 | 0.555153 | down |
| 64419 | MTMR14 | 0.000342 | 0.647443 | 0.676611 | 0.636592 | 0.653549 | down |
| 51278 | IER5 | 0.000342 | 0.62545 | 0.552438 | 0.49412 | 0.557336 | down |
| 66005 | CHID1 | 0.000342 | 0.624293 | 0.648386 | 0.5335 | 0.60206 | down |
| 8303 | SNN | 0.000344 | 0.56465 | 0.655003 | 0.635201 | 0.618285 | down |
| 25943 | C20orf194 | 0.000345 | 0.570979 | 0.607775 | 0.554572 | 0.577775 | down |
| 1471 | CST3 | 0.000346 | 0.70984 | 0.660186 | 0.622733 | 0.664253 | down |
| 8629 | JRK | 0.000348 | 0.426345 | 0.45147 | 0.497484 | 0.458433 | down |
| 151354 | FAM84A | 0.000348 | 0.523702 | 0.638803 | 0.564001 | 0.575502 | down |
| 57189 | KIAA1147 | 0.000349 | 0.588388 | 0.654164 | 0.651127 | 0.631226 | down |
| 11143 | KAT7 | 0.000349 | 0.678839 | 0.628854 | 0.688847 | 0.665513 | down |
| 23131 | GPATCH8 | 0.000351 | 0.613682 | 0.641339 | 0.583279 | 0.612767 | down |
| 6840 | SVIL | 0.000352 | 0.609357 | 0.630076 | 0.613914 | 0.617782 | down |
| 7094 | TLN1 | 0.000353 | 0.653315 | 0.645826 | 0.593517 | 0.630886 | down |
| 6871 | TADA2A | 0.000354 | 0.504138 | 0.572464 | 0.609432 | 0.562011 | down |
| 994 | CDC25B | 0.000354 | 0.601389 | 0.563783 | 0.574291 | 0.579821 | down |
| 8940 | TOP3B | 0.000355 | 0.569187 | 0.470733 | 0.438185 | 0.492702 | down |
| 23511 | NUP188 | 0.000355 | 0.551145 | 0.688671 | 0.590616 | 0.610144 | down |
| 54535 | CCHCR1 | 0.000356 | 0.621371 | 0.67814 | 0.622783 | 0.640765 | down |
| 10250 | SRRM1 | 0.000357 | 0.570646 | 0.669951 | 0.609553 | 0.616717 | down |
| 6721 | SREBF2 | 0.000357 | 0.642231 | 0.705405 | 0.634416 | 0.660684 | down |
| 5339 | PLEC | 0.000358 | 0.669396 | 0.648286 | 0.570075 | 0.629252 | down |
| 6253 | RTN2 | 0.000358 | 0.611263 | 0.486301 | 0.592316 | 0.563293 | down |
| 10067 | SCAMP3 | 0.000358 | 0.685461 | 0.616181 | 0.682313 | 0.661318 | down |
| 23107 | MRPS27 | 0.00036 | 0.603978 | 0.68934 | 0.705982 | 0.666433 | down |
| 124975 | GGT6 | 0.00036 | 0.632398 | 0.705916 | 0.586723 | 0.641679 | down |
| 5291 | PIK3CB | 0.000361 | 0.608299 | 0.682783 | 0.695863 | 0.662315 | down |
| 55187 | VPS13D | 0.000362 | 0.563786 | 0.574156 | 0.537468 | 0.55847 | down |
| 4628 | MYH10 | 0.000364 | 0.559193 | 0.639233 | 0.514204 | 0.570877 | down |
| 29123 | ANKRD11 | 0.000366 | 0.614533 | 0.687054 | 0.620066 | 0.640551 | down |
| 6564 | SLC15A1 | 0.000366 | 0.508117 | 0.606035 | 0.644744 | 0.586298 | down |
| 54101 | RIPK4 | 0.000367 | 0.634404 | 0.713928 | 0.583378 | 0.643903 | down |
| 4669 | NAGLU | 0.000367 | 0.67671 | 0.613906 | 0.688556 | 0.659724 | down |
| 9080 | CLDN9 | 0.000368 | 0.28306 | 0.426096 | 0.360816 | 0.356657 | down |
| 23779 | ARHGAP8 | 0.000368 | 0.711198 | 0.592975 | 0.598659 | 0.634277 | down |
| 4646 | MYO6 | 0.000369 | 0.629253 | 0.588198 | 0.704017 | 0.640489 | down |
| 2314 | FLII | 0.00037 | 0.639224 | 0.690986 | 0.617471 | 0.649227 | down |
| 23065 | EMC1 | 0.000371 | 0.647506 | 0.617626 | 0.571567 | 0.612233 | down |

| | | | | | | | |
|---|---|---|---|---|---|---|---|
| 284129 | SLC26A11 | 0.000371 | 0.637177 | 0.555284 | 0.572431 | 0.588298 | down |
| 23087 | TRIM35 | 0.000372 | 0.630138 | 0.702031 | 0.582747 | 0.638305 | down |
| 101 | ADAM8 | 0.000373 | 0.259853 | 0.339286 | 0.171443 | 0.256861 | down |
| 29785 | CYP2S1 | 0.000374 | 0.505071 | 0.40733 | 0.529894 | 0.480765 | down |
| 6904 | TBCD | 0.000375 | 0.601389 | 0.646729 | 0.567886 | 0.605334 | down |
| 1440 | CSF3 | 0.000376 | 0.311525 | 0.19122 | 0.249257 | 0.250668 | down |
| 79591 | ARMH3 | 0.000376 | 0.578781 | 0.617651 | 0.687189 | 0.627874 | down |
| 84876 | ORAI1 | 0.000376 | 0.621029 | 0.601941 | 0.667121 | 0.630031 | down |
| 57661 | PHRF1 | 0.000378 | 0.632408 | 0.596799 | 0.585243 | 0.604817 | down |
| 79369 | B3GNT4 | 0.000379 | 0.642857 | 0.547936 | 0.540979 | 0.577257 | down |
| 4035 | LRP1 | 0.000379 | 0.672767 | 0.547337 | 0.561912 | 0.594005 | down |
| 284023 | RNF227 | 0.00038 | 0.526168 | 0.49242 | 0.591559 | 0.536715 | down |
| 112399 | EGLN3 | 0.00038 | 0.684936 | 0.665835 | 0.620887 | 0.657219 | down |
| 79628 | SH3TC2 | 0.000382 | 0.54849 | 0.548772 | 0.554825 | 0.550695 | down |
| 23085 | ERC1 | 0.000382 | 0.619899 | 0.672834 | 0.657252 | 0.649995 | down |
| 22880 | MORC2 | 0.000383 | 0.643932 | 0.632372 | 0.599141 | 0.625148 | down |
| 26173 | INTS1 | 0.000384 | 0.506153 | 0.577041 | 0.500335 | 0.527843 | down |
| 84939 | MUM1 | 0.000384 | 0.570857 | 0.521816 | 0.433411 | 0.508695 | down |
| 10066 | SCAMP2 | 0.000384 | 0.612784 | 0.562391 | 0.609659 | 0.594945 | down |
| 80758 | PRR7 | 0.000384 | 0.47955 | 0.568807 | 0.457721 | 0.502026 | down |
| 5154 | PDGFA | 0.000385 | 0.546594 | 0.599065 | 0.542271 | 0.562643 | down |
| 710 | SERPING1 | 0.000386 | 0.520714 | 0.574808 | 0.560838 | 0.55212 | down |
| 51755 | CDK12 | 0.000386 | 0.594967 | 0.681959 | 0.677531 | 0.651486 | down |
| 125058 | TBC1D16 | 0.000387 | 0.609394 | 0.525949 | 0.490289 | 0.541877 | down |
| 115817 | DHRS1 | 0.000388 | 0.529745 | 0.465487 | 0.517483 | 0.504238 | down |
| 339123 | JMJD8 | 0.00039 | 0.610217 | 0.550972 | 0.676087 | 0.612425 | down |
| 1455 | CSNK1G2 | 0.00039 | 0.646255 | 0.71495 | 0.619617 | 0.660274 | down |
| 116039 | OSR2 | 0.000392 | 0.592004 | 0.554506 | 0.530901 | 0.559137 | down |
| 5338 | PLD2 | 0.000393 | 0.675751 | 0.675924 | 0.584273 | 0.645316 | down |
| 7343 | UBTF | 0.000394 | 0.627417 | 0.636621 | 0.652983 | 0.639007 | down |
| 1E+08 | ARMCX4 | 0.000395 | 0.4639 | 0.557062 | 0.534483 | 0.518482 | down |
| 84067 | FAM160A2 | 0.000395 | 0.647984 | 0.675987 | 0.554255 | 0.626076 | down |
| 3107 | HLA-C | 0.000395 | 0.695741 | 0.676801 | 0.611064 | 0.661202 | down |
| 5428 | POLG | 0.000397 | 0.599392 | 0.690417 | 0.674403 | 0.654737 | down |
| 23093 | TTLL5 | 0.000398 | 0.650718 | 0.685854 | 0.629527 | 0.655366 | down |
| 26012 | NSMF | 0.000398 | 0.595519 | 0.521744 | 0.621237 | 0.5795 | down |
| 1287 | COL4A5 | 0.000398 | 0.576384 | 0.604541 | 0.561955 | 0.58096 | down |
| 11346 | SYNPO | 0.000399 | 0.562825 | 0.599341 | 0.555573 | 0.57258 | down |
| 5540 | NPY4R | 0.000399 | 0.433069 | 0.29347 | 0.464 | 0.396846 | down |
| 57523 | NYNRIN | 0.0004 | 0.481067 | 0.526012 | 0.570533 | 0.525871 | down |
| 7297 | TYK2 | 0.000401 | 0.520051 | 0.653776 | 0.567159 | 0.580329 | down |
| 6239 | RREB1 | 0.000403 | 0.647097 | 0.630914 | 0.619336 | 0.632449 | down |
| 54890 | ALKBH5 | 0.000405 | 0.615792 | 0.681987 | 0.567985 | 0.621921 | down |
| 255057 | CBARP | 0.000405 | 0.576221 | 0.478665 | 0.614831 | 0.556572 | down |
| 26469 | PTPN18 | 0.000406 | 0.597462 | 0.524133 | 0.649501 | 0.590365 | down |
| 65083 | NOL6 | 0.000407 | 0.52371 | 0.536421 | 0.571125 | 0.543752 | down |
| 9742 | IFT140 | 0.000408 | 0.55848 | 0.585719 | 0.497111 | 0.547103 | down |
| 56257 | MEPCE | 0.00041 | 0.693413 | 0.614888 | 0.560627 | 0.622976 | down |
| 64285 | RHBDF1 | 0.000412 | 0.560642 | 0.598038 | 0.512342 | 0.557007 | down |
| 23138 | N4BP3 | 0.000412 | 0.549457 | 0.487443 | 0.402824 | 0.479908 | down |
| 5817 | PVR | 0.000413 | 0.67379 | 0.639712 | 0.594505 | 0.636003 | down |
| 1E+08 | ATP6V1FN | 0.000414 | 0.44 | 0.466667 | 0.584615 | 0.497094 | down |
| 5310 | PKD1 | 0.000414 | 0.532724 | 0.512389 | 0.460174 | 0.501762 | down |
| 3619 | INCENP | 0.000415 | 0.573441 | 0.657101 | 0.609076 | 0.613206 | down |
| 79724 | ZNF768 | 0.000415 | 0.634699 | 0.655052 | 0.659134 | 0.649628 | down |
| 201232 | SLC16A13 | 0.000416 | 0.551199 | 0.576426 | 0.453544 | 0.527056 | down |
| 83786 | FRMD8 | 0.000416 | 0.551588 | 0.661479 | 0.523136 | 0.578734 | down |
| 3911 | LAMA5 | 0.000417 | 0.627804 | 0.614733 | 0.543155 | 0.595231 | down |

| | | | | | | | |
|---|---|---|---|---|---|---|---|
| 9641 | IKBKE | 0.000419 | 0.542179 | 0.518994 | 0.641763 | 0.567645 | down |
| 116236 | ABHD15 | 0.000422 | 0.664247 | 0.651521 | 0.555321 | 0.623696 | down |
| 149013 | NBPF12 | 0.000425 | 0.56014 | 0.606052 | 0.571059 | 0.579084 | down |
| 20 | ABCA2 | 0.000426 | 0.474745 | 0.49027 | 0.40998 | 0.458332 | down |
| 10331 | B3GNT3 | 0.000427 | 0.622525 | 0.494311 | 0.540842 | 0.55256 | down |
| 7067 | THRA | 0.000429 | 0.581152 | 0.54941 | 0.482616 | 0.537726 | down |
| 84172 | POLR1B | 0.000429 | 0.674814 | 0.614918 | 0.65864 | 0.649458 | down |
| 11194 | ABCB8 | 0.00043 | 0.541066 | 0.551322 | 0.574195 | 0.555528 | down |
| 80851 | SH3BP5L | 0.000432 | 0.650161 | 0.559742 | 0.496641 | 0.568848 | down |
| 57822 | GRHL3 | 0.000434 | 0.608237 | 0.592328 | 0.526522 | 0.575695 | down |
| 84433 | CARD11 | 0.000435 | 0.62494 | 0.633327 | 0.544018 | 0.600762 | down |
| 84629 | TNRC18 | 0.000437 | 0.617235 | 0.598264 | 0.50813 | 0.574543 | down |
| 84256 | FLYWCH1 | 0.000438 | 0.662418 | 0.656171 | 0.643634 | 0.654074 | down |
| 54929 | TMEM161/ | 0.000438 | 0.639283 | 0.488658 | 0.534201 | 0.554047 | down |
| 23108 | RAP1GAP2 | 0.000438 | 0.529684 | 0.579941 | 0.564569 | 0.558065 | down |
| 59274 | TLNRD1 | 0.000438 | 0.618464 | 0.678873 | 0.562547 | 0.619961 | down |
| 10370 | CITED2 | 0.000438 | 0.618914 | 0.65488 | 0.692004 | 0.655266 | down |
| 80724 | ACAD10 | 0.00044 | 0.593037 | 0.556533 | 0.457023 | 0.535531 | down |
| 10188 | TNK2 | 0.00044 | 0.688849 | 0.610929 | 0.597996 | 0.632591 | down |
| 8884 | SLC5A6 | 0.000442 | 0.532411 | 0.5224 | 0.6329 | 0.562571 | down |
| 85441 | HELZ2 | 0.000442 | 0.567427 | 0.540636 | 0.527221 | 0.545095 | down |
| 9088 | PKMYT1 | 0.000445 | 0.697006 | 0.672557 | 0.572889 | 0.647484 | down |
| 5361 | PLXNA1 | 0.000445 | 0.564087 | 0.567095 | 0.559251 | 0.563478 | down |
| 4059 | BCAM | 0.000447 | 0.71208 | 0.661324 | 0.586795 | 0.6534 | down |
| 10497 | UNC13B | 0.000447 | 0.574504 | 0.478771 | 0.523625 | 0.525633 | down |
| 23761 | PISD | 0.000448 | 0.676131 | 0.575906 | 0.600543 | 0.617526 | down |
| 5986 | RFNG | 0.000448 | 0.674644 | 0.592878 | 0.684058 | 0.650527 | down |
| 7071 | KLF10 | 0.00045 | 0.552252 | 0.632872 | 0.68571 | 0.623611 | down |
| 1742 | DLG4 | 0.00045 | 0.680176 | 0.574404 | 0.680457 | 0.645012 | down |
| 90678 | LRSAM1 | 0.00045 | 0.512579 | 0.647562 | 0.593369 | 0.584503 | down |
| 23518 | R3HDM1 | 0.000451 | 0.606283 | 0.671404 | 0.703203 | 0.660297 | down |
| 5360 | PLTP | 0.000451 | 0.640787 | 0.595516 | 0.56076 | 0.599021 | down |
| 79654 | HECTD3 | 0.000451 | 0.663984 | 0.657763 | 0.629997 | 0.650581 | down |
| 1196 | CLK2 | 0.000451 | 0.591045 | 0.68892 | 0.570684 | 0.616883 | down |
| 5297 | PI4KA | 0.000452 | 0.636701 | 0.688266 | 0.578847 | 0.634605 | down |
| 124402 | UBALD1 | 0.000453 | 0.566538 | 0.653662 | 0.611666 | 0.610622 | down |
| 64744 | SMAP2 | 0.000453 | 0.611524 | 0.663211 | 0.70762 | 0.660785 | down |
| 57827 | C6orf47 | 0.000453 | 0.593852 | 0.63694 | 0.595319 | 0.608704 | down |
| 5371 | PML | 0.000454 | 0.618756 | 0.625389 | 0.701558 | 0.648568 | down |
| 57126 | CD177 | 0.000455 | 0.709155 | 0.600738 | 0.605728 | 0.63854 | down |
| 57538 | ALPK3 | 0.000456 | 0.566038 | 0.421296 | 0.463426 | 0.483587 | down |
| 3097 | HIVEP2 | 0.000456 | 0.612819 | 0.657272 | 0.596213 | 0.622101 | down |
| 22838 | RNF44 | 0.000457 | 0.579088 | 0.549787 | 0.677211 | 0.602029 | down |
| 81607 | NECTIN4 | 0.000458 | 0.569051 | 0.682388 | 0.684825 | 0.645421 | down |
| 9731 | CEP104 | 0.000459 | 0.640484 | 0.694067 | 0.618028 | 0.65086 | down |
| 23248 | RPRD2 | 0.000459 | 0.571971 | 0.625317 | 0.68854 | 0.628609 | down |
| 1509 | CTSD | 0.000459 | 0.689169 | 0.667551 | 0.635746 | 0.664155 | down |
| 6050 | RNH1 | 0.00046 | 0.676678 | 0.662101 | 0.569966 | 0.636248 | down |
| 114548 | NLRP3 | 0.000463 | 0.361538 | 0.255206 | 0.419408 | 0.345384 | down |
| 284348 | LYPD5 | 0.000463 | 0.487598 | 0.604364 | 0.545242 | 0.545735 | down |
| 375790 | AGRN | 0.000465 | 0.53357 | 0.53825 | 0.502106 | 0.524642 | down |
| 8795 | TNFRSF10I | 0.000466 | 0.649708 | 0.651411 | 0.662273 | 0.654464 | down |
| 3177 | SLC29A2 | 0.000468 | 0.557667 | 0.522005 | 0.458658 | 0.512777 | down |
| 80017 | DGLUCY | 0.000468 | 0.631931 | 0.587329 | 0.665835 | 0.628365 | down |
| 5190 | PEX6 | 0.000469 | 0.571292 | 0.530198 | 0.648434 | 0.583308 | down |
| 10659 | CELF2 | 0.00047 | 0.557522 | 0.687644 | 0.629119 | 0.624762 | down |
| 10603 | SH2B2 | 0.00047 | 0.4245 | 0.439659 | 0.518704 | 0.460954 | down |
| 80020 | FOXRED2 | 0.000471 | 0.49971 | 0.583089 | 0.619697 | 0.567499 | down |

| | | | | | | | |
|---|---|---|---|---|---|---|---|
| 79894 | ZNF672 | 0.000473 | 0.576054 | 0.556145 | 0.576049 | 0.569416 | down |
| 8567 | MADD | 0.000474 | 0.596326 | 0.629075 | 0.651638 | 0.62568 | down |
| 23151 | GRAMD4 | 0.000474 | 0.518659 | 0.586212 | 0.438589 | 0.514487 | down |
| 9649 | RALGPS1 | 0.000475 | 0.648381 | 0.583977 | 0.70512 | 0.645826 | down |
| 9839 | ZEB2 | 0.000475 | 0.21875 | 0.171429 | 0.2 | 0.196726 | down |
| 23646 | PLD3 | 0.000475 | 0.681998 | 0.574541 | 0.567659 | 0.608066 | down |
| 51752 | ERAP1 | 0.000476 | 0.618681 | 0.688502 | 0.617056 | 0.641413 | down |
| 9894 | TELO2 | 0.000477 | 0.603097 | 0.558302 | 0.530137 | 0.563845 | down |
| 79073 | TMEM109 | 0.000477 | 0.563787 | 0.608291 | 0.664982 | 0.612354 | down |
| 2195 | FAT1 | 0.000478 | 0.607481 | 0.665039 | 0.632685 | 0.635069 | down |
| 1.08E+08 | 1.08E+08 | 0.000479 | 0.536113 | 0.587561 | 0.680884 | 0.601519 | down |
| 208 | AKT2 | 0.000479 | 0.65829 | 0.603534 | 0.5477 | 0.603175 | down |
| 3714 | JAG2 | 0.00048 | 0.644581 | 0.687625 | 0.605873 | 0.646026 | down |
| 6621 | SNAPC4 | 0.00048 | 0.554984 | 0.477947 | 0.539394 | 0.524108 | down |
| 116028 | RMI2 | 0.00048 | 0.575617 | 0.689448 | 0.615329 | 0.626798 | down |
| 4355 | MPP2 | 0.000482 | 0.550502 | 0.58468 | 0.520368 | 0.55185 | down |
| 155435 | RBM33 | 0.000482 | 0.600809 | 0.647521 | 0.660004 | 0.636111 | down |
| 55065 | SLC52A1 | 0.000484 | 0.605959 | 0.505026 | 0.568138 | 0.559708 | down |
| 3054 | HCFC1 | 0.000487 | 0.563582 | 0.656733 | 0.577456 | 0.599257 | down |
| 7077 | TIMP2 | 0.000488 | 0.639517 | 0.652914 | 0.636713 | 0.643048 | down |
| 8425 | LTBP4 | 0.000494 | 0.505023 | 0.564168 | 0.454964 | 0.508052 | down |
| 6786 | STIM1 | 0.000495 | 0.670192 | 0.572685 | 0.574539 | 0.605805 | down |
| 387509 | GPR153 | 0.000496 | 0.375492 | 0.398969 | 0.500409 | 0.424957 | down |
| 57704 | GBA2 | 0.000497 | 0.670847 | 0.582204 | 0.591374 | 0.614808 | down |
| 57505 | AARS2 | 0.0005 | 0.492537 | 0.652168 | 0.542268 | 0.562325 | down |
| 283489 | CHAMP1 | 0.000501 | 0.522965 | 0.612271 | 0.667265 | 0.600834 | down |
| 259232 | NALCN | 0.000501 | 0.530833 | 0.508275 | 0.497627 | 0.512245 | down |
| 4324 | MMP15 | 0.000504 | 0.563907 | 0.546084 | 0.604892 | 0.571628 | down |
| 9351 | SLC9A3R2 | 0.000504 | 0.65361 | 0.574038 | 0.55085 | 0.592833 | down |
| 83985 | SPNS1 | 0.000505 | 0.685488 | 0.633863 | 0.613535 | 0.644296 | down |
| 256364 | EML3 | 0.000505 | 0.640864 | 0.660542 | 0.525645 | 0.609017 | down |
| 23102 | TBC1D2B | 0.000506 | 0.608108 | 0.623909 | 0.65719 | 0.629736 | down |
| 80208 | SPG11 | 0.000507 | 0.557646 | 0.701798 | 0.624645 | 0.62803 | down |
| 146705 | TEPSIN | 0.000508 | 0.642185 | 0.691074 | 0.621124 | 0.651461 | down |
| 1295 | COL8A1 | 0.000508 | 0.327068 | 0.460638 | 0.46281 | 0.416838 | down |
| 10948 | STARD3 | 0.000509 | 0.666463 | 0.588576 | 0.610786 | 0.621942 | down |
| 4641 | MYO1C | 0.000509 | 0.645534 | 0.72425 | 0.609449 | 0.659744 | down |
| 9920 | KBTBD11 | 0.000511 | 0.53402 | 0.463797 | 0.576299 | 0.524705 | down |
| 25930 | PTPN23 | 0.000511 | 0.591842 | 0.720129 | 0.606614 | 0.639528 | down |
| 84133 | ZNRF3 | 0.000512 | 0.65806 | 0.623427 | 0.642553 | 0.641347 | down |
| 23476 | BRD4 | 0.000517 | 0.618002 | 0.654839 | 0.703311 | 0.658717 | down |
| 84926 | SPRYD3 | 0.000519 | 0.594379 | 0.582958 | 0.617648 | 0.598329 | down |
| 150726 | FBXO41 | 0.000519 | 0.613952 | 0.551155 | 0.485574 | 0.550227 | down |
| 6820 | SULT2B1 | 0.000521 | 0.346711 | 0.357896 | 0.424387 | 0.376331 | down |
| 57693 | ZNF317 | 0.000522 | 0.562588 | 0.647482 | 0.607993 | 0.606021 | down |
| 57674 | RNF213 | 0.000525 | 0.476819 | 0.573056 | 0.538458 | 0.529444 | down |
| 7262 | PHLDA2 | 0.000528 | 0.649607 | 0.57326 | 0.534906 | 0.585925 | down |
| 10634 | GAS2L1 | 0.000529 | 0.563571 | 0.679186 | 0.535041 | 0.592599 | down |
| 79888 | LPCAT1 | 0.000529 | 0.54431 | 0.585043 | 0.502307 | 0.543886 | down |
| 27348 | TOR1B | 0.00053 | 0.60293 | 0.625804 | 0.713529 | 0.647421 | down |
| 57221 | ARFGEF3 | 0.00053 | 0.595223 | 0.67476 | 0.602048 | 0.62401 | down |
| 57537 | SORCS2 | 0.00053 | 0.56451 | 0.385221 | 0.497991 | 0.482574 | down |
| 9654 | TTLL4 | 0.000531 | 0.587117 | 0.610704 | 0.576915 | 0.591579 | down |
| 83937 | RASSF4 | 0.000535 | 0.482954 | 0.565631 | 0.637339 | 0.561975 | down |
| 388115 | CCDC9B | 0.000535 | 0.538848 | 0.572952 | 0.603826 | 0.571876 | down |
| 79877 | DCAKD | 0.00054 | 0.519121 | 0.652924 | 0.640963 | 0.604336 | down |
| 25829 | TMEM184I | 0.00054 | 0.664196 | 0.589763 | 0.683103 | 0.645687 | down |
| 4851 | NOTCH1 | 0.00054 | 0.619334 | 0.712823 | 0.66448 | 0.665546 | down |

| | | | | | | | |
|---|---|---|---|---|---|---|---|
| 4548 | MTR | 0.000543 | 0.626573 | 0.646473 | 0.717523 | 0.663523 | down |
| 3975 | LHX1 | 0.000546 | 0.454706 | 0.599653 | 0.553066 | 0.535809 | down |
| 8925 | HERC1 | 0.000547 | 0.49181 | 0.543246 | 0.563594 | 0.532883 | down |
| 211 | ALAS1 | 0.000551 | 0.606089 | 0.681438 | 0.705266 | 0.664264 | down |
| 2012 | EMP1 | 0.000551 | 0.643666 | 0.596367 | 0.650248 | 0.630094 | down |
| 201255 | LRRC45 | 0.000551 | 0.683494 | 0.575179 | 0.601784 | 0.620153 | down |
| 80789 | INTS5 | 0.000553 | 0.575091 | 0.586774 | 0.65107 | 0.604312 | down |
| 56652 | TWNK | 0.000555 | 0.545156 | 0.614991 | 0.614059 | 0.591402 | down |
| 23133 | PHF8 | 0.000555 | 0.508853 | 0.626741 | 0.553788 | 0.563128 | down |
| 2580 | GAK | 0.000559 | 0.568468 | 0.660928 | 0.557538 | 0.595645 | down |
| 6421 | SFPQ | 0.000561 | 0.619219 | 0.657498 | 0.578753 | 0.61849 | down |
| 9677 | PPIP5K1 | 0.000561 | 0.522865 | 0.560018 | 0.651347 | 0.578077 | down |
| 28231 | SLCO4A1 | 0.000561 | 0.478207 | 0.622218 | 0.477203 | 0.525876 | down |
| 84236 | RHBDD1 | 0.000563 | 0.506856 | 0.630908 | 0.616455 | 0.58474 | down |
| 7185 | TRAF1 | 0.000564 | 0.65812 | 0.566176 | 0.588135 | 0.604144 | down |
| 64115 | VSIR | 0.000564 | 0.484624 | 0.651732 | 0.579042 | 0.571799 | down |
| 23534 | TNPO3 | 0.000564 | 0.568634 | 0.678705 | 0.702673 | 0.650004 | down |
| 4133 | MAP2 | 0.000568 | 0.538958 | 0.551948 | 0.683571 | 0.591492 | down |
| 3297 | HSF1 | 0.000568 | 0.638732 | 0.564782 | 0.615189 | 0.606234 | down |
| 84948 | TIGD5 | 0.000571 | 0.690748 | 0.647419 | 0.631241 | 0.656469 | down |
| 254042 | METAP1D | 0.000573 | 0.639648 | 0.491223 | 0.606462 | 0.579111 | down |
| 1.08E+08 | LOC10798 | 0.000574 | 0.56909 | 0.460643 | 0.464415 | 0.49805 | down |
| 57801 | HES4 | 0.000574 | 0.635765 | 0.691652 | 0.563768 | 0.630395 | down |
| 155368 | METTL27 | 0.000574 | 0.50534 | 0.491622 | 0.61534 | 0.537434 | down |
| 1185 | CLCN6 | 0.000576 | 0.591012 | 0.577169 | 0.516404 | 0.561528 | down |
| 84464 | SLX4 | 0.000577 | 0.535375 | 0.520522 | 0.606593 | 0.554163 | down |
| 54842 | MFSD6 | 0.000578 | 0.568499 | 0.58488 | 0.680569 | 0.611316 | down |
| 27043 | PELP1 | 0.000578 | 0.648804 | 0.595872 | 0.553312 | 0.599329 | down |
| 3418 | IDH2 | 0.000578 | 0.676239 | 0.700569 | 0.612631 | 0.663146 | down |
| 91 | ACVR1B | 0.000578 | 0.61419 | 0.68591 | 0.686128 | 0.662076 | down |
| 1616 | DAXX | 0.000581 | 0.682088 | 0.605846 | 0.62846 | 0.638798 | down |
| 345643 | MCIDAS | 0.000582 | 0.533012 | 0.459596 | 0.514889 | 0.502499 | down |
| 10481 | HOXB13 | 0.000583 | 0.656392 | 0.59677 | 0.593424 | 0.615529 | down |
| 53838 | C11orf24 | 0.000585 | 0.621178 | 0.58108 | 0.72294 | 0.641732 | down |
| 148022 | TICAM1 | 0.000585 | 0.580536 | 0.509932 | 0.593772 | 0.561413 | down |
| 51559 | NT5DC3 | 0.000586 | 0.624735 | 0.65733 | 0.611813 | 0.631293 | down |
| 1605 | DAG1 | 0.000588 | 0.607544 | 0.726023 | 0.659341 | 0.664303 | down |
| 114884 | OSBPL10 | 0.00059 | 0.597413 | 0.618962 | 0.597384 | 0.604587 | down |
| 3014 | H2AFX | 0.00059 | 0.663195 | 0.616539 | 0.593667 | 0.624467 | down |
| 51435 | SCARA3 | 0.00059 | 0.585874 | 0.66161 | 0.605805 | 0.617763 | down |
| 9231 | DLG5 | 0.000591 | 0.615291 | 0.710263 | 0.661603 | 0.662386 | down |
| 28996 | HIPK2 | 0.000591 | 0.591996 | 0.647059 | 0.699041 | 0.646032 | down |
| 221 | ALDH3B1 | 0.000591 | 0.551209 | 0.603044 | 0.493367 | 0.549206 | down |
| 200895 | DHFR2 | 0.000594 | 0.569107 | 0.536049 | 0.613347 | 0.572834 | down |
| 3976 | LIF | 0.000595 | 0.543853 | 0.543526 | 0.659898 | 0.582426 | down |
| 51385 | ZNF589 | 0.000595 | 0.63874 | 0.599294 | 0.66747 | 0.635168 | down |
| 861 | RUNX1 | 0.000599 | 0.609812 | 0.661313 | 0.706643 | 0.659256 | down |
| 6942 | TCF20 | 0.000599 | 0.55537 | 0.598894 | 0.688005 | 0.61409 | down |
| 2011 | MARK2 | 0.0006 | 0.609313 | 0.580605 | 0.716074 | 0.635331 | down |
| 125111 | GJD3 | 0.0006 | 0.641347 | 0.516866 | 0.518853 | 0.559022 | down |
| 9545 | RAB3D | 0.0006 | 0.568064 | 0.666137 | 0.574736 | 0.602979 | down |
| 1954 | MEGF8 | 0.000601 | 0.485016 | 0.581811 | 0.525308 | 0.530712 | down |
| 583 | BBS2 | 0.000601 | 0.706579 | 0.621549 | 0.59138 | 0.639836 | down |
| 171017 | ZNF384 | 0.000605 | 0.646382 | 0.606272 | 0.697572 | 0.650075 | down |
| 79412 | KREMEN2 | 0.000609 | 0.489235 | 0.5915 | 0.488669 | 0.523135 | down |
| 9730 | DCAF1 | 0.000609 | 0.572706 | 0.712914 | 0.640027 | 0.641882 | down |
| 79814 | AGMAT | 0.000609 | 0.523845 | 0.631295 | 0.597359 | 0.584166 | down |
| 1282 | COL4A1 | 0.000611 | 0.55259 | 0.630169 | 0.554453 | 0.579071 | down |

| | | | | | | | |
|---|---|---|---|---|---|---|---|
| 7090 | TLE3 | 0.000612 | 0.640528 | 0.505424 | 0.576658 | 0.574204 | down |
| 56949 | XAB2 | 0.000612 | 0.677986 | 0.668577 | 0.623946 | 0.656837 | down |
| 3200 | HOXA3 | 0.000615 | 0.559789 | 0.564555 | 0.581278 | 0.568541 | down |
| 6794 | STK11 | 0.000616 | 0.62838 | 0.642646 | 0.557279 | 0.609435 | down |
| 136051 | ZNF786 | 0.000617 | 0.524332 | 0.648688 | 0.498111 | 0.557044 | down |
| 54682 | MANSC1 | 0.000618 | 0.519097 | 0.497308 | 0.43818 | 0.484862 | down |
| 1.03E+08 | LOC10272 | 0.000619 | 0.451657 | 0.524049 | 0.620294 | 0.532 | down |
| 388228 | SBK1 | 0.000626 | 0.34749 | 0.442708 | 0.360156 | 0.383451 | down |
| 399474 | TMEM200I | 0.000628 | 0.626474 | 0.622771 | 0.621528 | 0.623591 | down |
| 2264 | FGFR4 | 0.000628 | 0.548091 | 0.654191 | 0.501221 | 0.567834 | down |
| 396 | ARHGDIA | 0.000629 | 0.661885 | 0.552058 | 0.585929 | 0.599958 | down |
| 1609 | DGKQ | 0.000634 | 0.673097 | 0.616857 | 0.552061 | 0.614005 | down |
| 400916 | CHCHD10 | 0.00064 | 0.657021 | 0.613125 | 0.634952 | 0.635033 | down |
| 79695 | GALNT12 | 0.000642 | 0.671868 | 0.575102 | 0.575766 | 0.607579 | down |
| 8079 | MLF2 | 0.000643 | 0.636375 | 0.56484 | 0.640304 | 0.61384 | down |
| 9696 | CROCC | 0.000647 | 0.522141 | 0.485435 | 0.532377 | 0.513318 | down |
| 401145 | CCSER1 | 0.000648 | 0.509091 | 0.545455 | 0.518519 | 0.524355 | down |
| 51517 | NCKIPSD | 0.000648 | 0.630369 | 0.641406 | 0.553409 | 0.608395 | down |
| 8991 | SELENBP1 | 0.00065 | 0.445071 | 0.525728 | 0.524004 | 0.498268 | down |
| 8527 | DGKD | 0.000651 | 0.545588 | 0.594997 | 0.573217 | 0.571267 | down |
| 63892 | THADA | 0.000653 | 0.624963 | 0.595664 | 0.560174 | 0.5936 | down |
| 6597 | SMARCA4 | 0.000653 | 0.63631 | 0.625678 | 0.583074 | 0.615021 | down |
| 10609 | P3H4 | 0.000655 | 0.617804 | 0.592455 | 0.644647 | 0.618302 | down |
| 1.01E+08 | LOC10106 | 0.000655 | 0.432339 | 0.362374 | 0.45655 | 0.417088 | down |
| 114823 | LENG8 | 0.000656 | 0.374023 | 0.541142 | 0.540752 | 0.485306 | down |
| 28955 | DEXI | 0.000657 | 0.698635 | 0.646246 | 0.574113 | 0.639665 | down |
| 10392 | NOD1 | 0.000661 | 0.587057 | 0.68171 | 0.607315 | 0.625361 | down |
| 1822 | ATN1 | 0.000662 | 0.677785 | 0.57688 | 0.665066 | 0.63991 | down |
| 64236 | PDLIM2 | 0.000662 | 0.548719 | 0.571055 | 0.664405 | 0.594726 | down |
| 79735 | TBC1D17 | 0.000663 | 0.727759 | 0.58158 | 0.624569 | 0.644636 | down |
| 25974 | MMACHC | 0.000664 | 0.610297 | 0.653753 | 0.600572 | 0.62154 | down |
| 80169 | CTC1 | 0.000665 | 0.57963 | 0.531773 | 0.526487 | 0.545963 | down |
| 79932 | KIAA0319L | 0.000667 | 0.576463 | 0.652133 | 0.567399 | 0.598665 | down |
| 3092 | HIP1 | 0.00067 | 0.603882 | 0.643275 | 0.546544 | 0.5979 | down |
| 9924 | PAN2 | 0.000671 | 0.465185 | 0.502931 | 0.428522 | 0.465546 | down |
| 54853 | WDR55 | 0.000675 | 0.617786 | 0.68764 | 0.670649 | 0.658692 | down |
| 116987 | AGAP1 | 0.000675 | 0.579428 | 0.577939 | 0.475789 | 0.544385 | down |
| 11022 | TDRKH | 0.000677 | 0.717125 | 0.647378 | 0.574344 | 0.646282 | down |
| 7498 | XDH | 0.000677 | 0.552175 | 0.597605 | 0.674565 | 0.608115 | down |
| 1869 | E2F1 | 0.000677 | 0.634912 | 0.58189 | 0.529816 | 0.582206 | down |
| 5652 | PRSS8 | 0.000678 | 0.615684 | 0.586307 | 0.635861 | 0.612617 | down |
| 8506 | CNTNAP1 | 0.000679 | 0.45927 | 0.509298 | 0.46385 | 0.477473 | down |
| 2475 | MTOR | 0.000679 | 0.596618 | 0.707234 | 0.572864 | 0.625572 | down |
| 2026 | ENO2 | 0.000679 | 0.662013 | 0.596182 | 0.618088 | 0.625428 | down |
| 25766 | PRPF40B | 0.000681 | 0.555249 | 0.560459 | 0.469584 | 0.52843 | down |
| 53827 | FXYD5 | 0.000684 | 0.640087 | 0.555508 | 0.663778 | 0.619791 | down |
| 3482 | IGF2R | 0.000688 | 0.593496 | 0.689793 | 0.585034 | 0.622775 | down |
| 11322 | TMC6 | 0.000688 | 0.567562 | 0.682236 | 0.512798 | 0.587532 | down |
| 4335 | MNT | 0.00069 | 0.565262 | 0.541716 | 0.571095 | 0.559358 | down |
| 5029 | P2RY2 | 0.000693 | 0.649745 | 0.528549 | 0.645277 | 0.607857 | down |
| 23334 | SZT2 | 0.000698 | 0.503723 | 0.510397 | 0.571212 | 0.528444 | down |
| 284904 | SEC14L4 | 0.000698 | 0.554793 | 0.595982 | 0.575687 | 0.575487 | down |
| 22859 | ADGRL1 | 0.000702 | 0.486122 | 0.497223 | 0.593527 | 0.525624 | down |
| 51442 | VGLL1 | 0.000702 | 0.599999 | 0.521427 | 0.688522 | 0.603316 | down |
| 30850 | CDR2L | 0.000702 | 0.585042 | 0.675786 | 0.683575 | 0.648134 | down |
| 124997 | WDR81 | 0.000702 | 0.531914 | 0.639174 | 0.476757 | 0.549282 | down |
| 2117 | ETV3 | 0.000702 | 0.582199 | 0.689528 | 0.684334 | 0.65202 | down |
| 53905 | DUOX1 | 0.000703 | 0.44218 | 0.595968 | 0.558298 | 0.532149 | down |

| | | | | | | | |
|---|---|---|---|---|---|---|---|
| 345079 | SOWAHB | 0.000704 | 0.577483 | 0.681188 | 0.560593 | 0.606421 | down |
| 835 | CASP2 | 0.000704 | 0.501664 | 0.671039 | 0.54566 | 0.572788 | down |
| 29801 | ZDHHC8 | 0.000704 | 0.587554 | 0.612517 | 0.528564 | 0.576212 | down |
| 23013 | SPEN | 0.000707 | 0.591625 | 0.680971 | 0.677335 | 0.649977 | down |
| 79813 | EHMT1 | 0.000708 | 0.598258 | 0.704476 | 0.626474 | 0.643069 | down |
| 64145 | RBSN | 0.000713 | 0.731455 | 0.637752 | 0.612448 | 0.660552 | down |
| 283450 | HECTD4 | 0.000714 | 0.514156 | 0.591731 | 0.539663 | 0.548517 | down |
| 114897 | C1QTNF1 | 0.000715 | 0.506081 | 0.625826 | 0.634781 | 0.588896 | down |
| 9572 | NR1D1 | 0.000716 | 0.516856 | 0.475848 | 0.634662 | 0.542455 | down |
| 22906 | TRAK1 | 0.000723 | 0.551142 | 0.628995 | 0.655317 | 0.611818 | down |
| 92335 | STRADA | 0.000726 | 0.646281 | 0.571778 | 0.656876 | 0.624978 | down |
| 8310 | ACOX3 | 0.000727 | 0.579187 | 0.600059 | 0.608352 | 0.595866 | down |
| 8914 | TIMELESS | 0.000727 | 0.681316 | 0.704706 | 0.613448 | 0.66649 | down |
| 9739 | SETD1A | 0.000727 | 0.71849 | 0.564041 | 0.644861 | 0.642464 | down |
| 10256 | CNKSR1 | 0.000728 | 0.657954 | 0.608281 | 0.659319 | 0.641852 | down |
| 55230 | USP40 | 0.000728 | 0.475966 | 0.610875 | 0.570554 | 0.552465 | down |
| 11273 | ATXN2L | 0.000731 | 0.702014 | 0.585704 | 0.654945 | 0.647555 | down |
| 57706 | DENND1A | 0.000733 | 0.586343 | 0.575126 | 0.515211 | 0.558893 | down |
| 8994 | LIMD1 | 0.000734 | 0.638378 | 0.636669 | 0.609241 | 0.628096 | down |
| 1.03E+08 | LOC10272 | 0.000734 | 0.615354 | 0.63932 | 0.504097 | 0.586257 | down |
| 51063 | CALHM2 | 0.000737 | 0.644452 | 0.689473 | 0.628501 | 0.654142 | down |
| 9651 | PLCH2 | 0.000742 | 0.541935 | 0.616784 | 0.546932 | 0.56855 | down |
| 23250 | ATP11A | 0.000742 | 0.556725 | 0.663636 | 0.688032 | 0.636131 | down |
| 57602 | USP36 | 0.000742 | 0.59818 | 0.653593 | 0.613522 | 0.621765 | down |
| 9148 | NEURL1 | 0.000743 | 0.532965 | 0.563959 | 0.500066 | 0.53233 | down |
| 118813 | ZFYVE27 | 0.000751 | 0.562827 | 0.628977 | 0.571087 | 0.58763 | down |
| 9582 | APOBEC3E | 0.000751 | 0.678959 | 0.548281 | 0.652755 | 0.626665 | down |
| 154881 | KCTD7 | 0.000754 | 0.59657 | 0.663661 | 0.702824 | 0.654351 | down |
| 81570 | CLPB | 0.000758 | 0.609216 | 0.578279 | 0.580781 | 0.589425 | down |
| 9656 | MDC1 | 0.000758 | 0.61199 | 0.530415 | 0.594787 | 0.579064 | down |
| 23144 | ZC3H3 | 0.000758 | 0.686351 | 0.662414 | 0.555123 | 0.634629 | down |
| 55731 | FAM222B | 0.00076 | 0.619462 | 0.587664 | 0.582553 | 0.59656 | down |
| 57799 | RAB40C | 0.000761 | 0.620776 | 0.688085 | 0.596997 | 0.635286 | down |
| 4351 | MPI | 0.000767 | 0.718701 | 0.625672 | 0.62779 | 0.657388 | down |
| 6668 | SP2 | 0.000769 | 0.639707 | 0.575914 | 0.688345 | 0.634656 | down |
| 23457 | ABCB9 | 0.000771 | 0.550535 | 0.572451 | 0.439485 | 0.520824 | down |
| 79365 | BHLHE41 | 0.000772 | 0.572211 | 0.50447 | 0.504115 | 0.526932 | down |
| 4241 | MELTF | 0.000773 | 0.672482 | 0.680667 | 0.586881 | 0.646677 | down |
| 3795 | KHK | 0.000774 | 0.554765 | 0.584639 | 0.677037 | 0.60548 | down |
| 10227 | MFSD10 | 0.000775 | 0.651176 | 0.653326 | 0.554905 | 0.619802 | down |
| 84874 | ZNF514 | 0.000775 | 0.627214 | 0.657922 | 0.654938 | 0.646691 | down |
| 23505 | TMEM131 | 0.000775 | 0.530263 | 0.692418 | 0.626359 | 0.616347 | down |
| 162979 | ZNF296 | 0.000778 | 0.696738 | 0.639803 | 0.617811 | 0.651451 | down |
| 2896 | GRN | 0.00078 | 0.688893 | 0.649852 | 0.59567 | 0.644805 | down |
| 54438 | GFOD1 | 0.000782 | 0.612474 | 0.690682 | 0.580745 | 0.627967 | down |
| 199221 | DZIP1L | 0.000785 | 0.59 | 0.555698 | 0.661628 | 0.602442 | down |
| 5336 | PLCG2 | 0.000785 | 0.622087 | 0.66987 | 0.587768 | 0.626575 | down |
| 140700 | SAMD10 | 0.000786 | 0.433309 | 0.559292 | 0.590744 | 0.527782 | down |
| 8507 | ENC1 | 0.000789 | 0.570421 | 0.59717 | 0.518135 | 0.561909 | down |
| 79147 | FKRP | 0.00079 | 0.559772 | 0.624315 | 0.50994 | 0.564676 | down |
| 8078 | USP5 | 0.000791 | 0.680676 | 0.660024 | 0.5796 | 0.6401 | down |
| 23649 | POLA2 | 0.000792 | 0.697629 | 0.557145 | 0.643138 | 0.632637 | down |
| 23358 | USP24 | 0.000792 | 0.645381 | 0.715645 | 0.622453 | 0.661159 | down |
| 92714 | ARRDC1 | 0.000795 | 0.71138 | 0.620048 | 0.596588 | 0.642672 | down |
| 65263 | PYCR3 | 0.000797 | 0.624007 | 0.641149 | 0.548235 | 0.604464 | down |
| 64359 | NXN | 0.000798 | 0.622161 | 0.71416 | 0.585616 | 0.640646 | down |
| 147015 | DHRS13 | 0.000799 | 0.731268 | 0.635357 | 0.62891 | 0.665178 | down |
| 4267 | CD99 | 0.000801 | 0.600598 | 0.625955 | 0.577087 | 0.601213 | down |

| | | | | | | | |
|---|---|---|---|---|---|---|---|
| 57647 | DHX37 | 0.000803 | 0.654453 | 0.684063 | 0.575831 | 0.638116 | down |
| 10347 | ABCA7 | 0.000803 | 0.502687 | 0.454035 | 0.51812 | 0.491614 | down |
| 84675 | TRIM55 | 0.000803 | 0.639204 | 0.565468 | 0.514358 | 0.57301 | down |
| 55229 | PANK4 | 0.000803 | 0.596954 | 0.622606 | 0.654612 | 0.624724 | down |
| 83742 | MARVELD: | 0.000805 | 0.623143 | 0.679781 | 0.512671 | 0.605198 | down |
| 7048 | TGFBR2 | 0.000807 | 0.587549 | 0.732875 | 0.601607 | 0.640677 | down |
| 1543 | CYP1A1 | 0.000808 | 0.582895 | 0.428865 | 0.516291 | 0.509351 | down |
| 4249 | MGAT5 | 0.00081 | 0.617567 | 0.69419 | 0.61421 | 0.641989 | down |
| 25970 | SH2B1 | 0.000811 | 0.656849 | 0.538816 | 0.5476 | 0.581088 | down |
| 624 | BDKRB2 | 0.000815 | 0.62509 | 0.690674 | 0.596362 | 0.637375 | down |
| 8812 | CCNK | 0.000816 | 0.646908 | 0.563839 | 0.7087 | 0.639816 | down |
| 7760 | ZNF213 | 0.000816 | 0.595887 | 0.652462 | 0.541516 | 0.596622 | down |
| 51490 | SPOUT1 | 0.000816 | 0.629836 | 0.608687 | 0.616896 | 0.618473 | down |
| 114991 | ZNF618 | 0.000816 | 0.579044 | 0.616692 | 0.64177 | 0.612502 | down |
| 3880 | KRT19 | 0.000817 | 0.653689 | 0.652756 | 0.534092 | 0.613512 | down |
| 23307 | FKBP15 | 0.000817 | 0.67058 | 0.627206 | 0.645517 | 0.647768 | down |
| 23466 | CBX6 | 0.000817 | 0.626089 | 0.722829 | 0.602907 | 0.650608 | down |
| 8349 | HIST2H2BI | 0.000818 | 0.524261 | 0.566087 | 0.635142 | 0.575163 | down |
| 6497 | SKI | 0.000819 | 0.591149 | 0.576535 | 0.581943 | 0.583209 | down |
| 3725 | JUN | 0.000824 | 0.706576 | 0.681795 | 0.590227 | 0.659533 | down |
| 56172 | ANKH | 0.000825 | 0.672945 | 0.660685 | 0.665672 | 0.666434 | down |
| 8479 | HIRIP3 | 0.000826 | 0.641956 | 0.583417 | 0.655692 | 0.627022 | down |
| 85379 | KIAA1671 | 0.000827 | 0.524679 | 0.654132 | 0.578215 | 0.585676 | down |
| 90324 | CCDC97 | 0.00083 | 0.629161 | 0.643794 | 0.70948 | 0.660812 | down |
| 9415 | FADS2 | 0.000833 | 0.638571 | 0.64163 | 0.630769 | 0.63699 | down |
| 1460 | CSNK2B | 0.000835 | 0.62988 | 0.584291 | 0.708538 | 0.640903 | down |
| 10435 | CDC42EP2 | 0.000835 | 0.401103 | 0.313656 | 0.463524 | 0.392761 | down |
| 79443 | FYCO1 | 0.000837 | 0.56806 | 0.659498 | 0.589379 | 0.605646 | down |
| 7156 | TOP3A | 0.000839 | 0.637786 | 0.711585 | 0.62978 | 0.659717 | down |
| 26873 | OPLAH | 0.00084 | 0.550331 | 0.590521 | 0.539133 | 0.559995 | down |
| 4054 | LTBP3 | 0.000843 | 0.648449 | 0.650599 | 0.587054 | 0.628701 | down |
| 22885 | ABLIM3 | 0.000843 | 0.444715 | 0.55328 | 0.5701 | 0.522698 | down |
| 11023 | VAX1 | 0.000844 | 0.316667 | 0.395833 | 0.346667 | 0.353056 | down |
| 5187 | PER1 | 0.000847 | 0.697394 | 0.639969 | 0.543681 | 0.627015 | down |
| 91523 | PCED1B | 0.000849 | 0.445423 | 0.5045 | 0.575583 | 0.508502 | down |
| 9064 | MAP3K6 | 0.000853 | 0.648889 | 0.502611 | 0.460193 | 0.537231 | down |
| 26523 | AGO1 | 0.000859 | 0.577284 | 0.623489 | 0.56159 | 0.587454 | down |
| 5819 | NECTIN2 | 0.00086 | 0.64172 | 0.583111 | 0.678588 | 0.634473 | down |
| 3720 | JARID2 | 0.000862 | 0.649281 | 0.554608 | 0.638079 | 0.613989 | down |
| 64773 | PCED1A | 0.000865 | 0.661418 | 0.629504 | 0.590732 | 0.627218 | down |
| 1.05E+08 | LOC10536 | 0.000869 | 0.58094 | 0.60152 | 0.498626 | 0.560362 | down |
| 3633 | INPP5B | 0.00087 | 0.578463 | 0.569016 | 0.715208 | 0.620896 | down |
| 196403 | DTX3 | 0.00087 | 0.585721 | 0.523427 | 0.671764 | 0.593637 | down |
| 91107 | TRIM47 | 0.000871 | 0.619785 | 0.573628 | 0.640504 | 0.611306 | down |
| 28227 | PPP2R3B | 0.000871 | 0.597487 | 0.670482 | 0.577041 | 0.615003 | down |
| 3557 | IL1RN | 0.000871 | 0.683967 | 0.597465 | 0.64076 | 0.640731 | down |
| 7056 | THBD | 0.000874 | 0.534973 | 0.70164 | 0.556895 | 0.597836 | down |
| 55697 | VAC14 | 0.000875 | 0.640743 | 0.658731 | 0.647341 | 0.648938 | down |
| 4968 | OGG1 | 0.000879 | 0.541422 | 0.676706 | 0.511619 | 0.576582 | down |
| 64858 | DCLRE1B | 0.00088 | 0.57773 | 0.685596 | 0.693438 | 0.652255 | down |
| 23241 | PACS2 | 0.000883 | 0.666133 | 0.627866 | 0.608533 | 0.634177 | down |
| 56654 | NPDC1 | 0.000885 | 0.640334 | 0.611129 | 0.578987 | 0.61015 | down |
| 54921 | CHTF8 | 0.00089 | 0.713683 | 0.566759 | 0.628872 | 0.636438 | down |
| 1399 | CRKL | 0.000891 | 0.642984 | 0.686846 | 0.664033 | 0.664621 | down |
| 57186 | RALGAPA2 | 0.000895 | 0.480991 | 0.561728 | 0.563743 | 0.535488 | down |
| 84166 | NLRC5 | 0.000895 | 0.60884 | 0.676531 | 0.619085 | 0.634819 | down |
| 57719 | ANO8 | 0.000897 | 0.524345 | 0.563041 | 0.546568 | 0.544651 | down |
| 84858 | ZNF503 | 0.000899 | 0.587682 | 0.59945 | 0.691254 | 0.626129 | down |

| | | | | | | | |
|---|---|---|---|---|---|---|---|
| 50 | ACO2 | 0.0009 | 0.700738 | 0.622527 | 0.638366 | 0.653877 | down |
| 3614 | IMPDH1 | 0.000903 | 0.706421 | 0.616512 | 0.589117 | 0.63735 | down |
| 117195 | MRGPRX3 | 0.000908 | 0.656262 | 0.594692 | 0.654825 | 0.635259 | down |
| 57121 | LPAR5 | 0.000911 | 0.532008 | 0.619691 | 0.614538 | 0.588746 | down |
| 729830 | FAM160A1 | 0.000911 | 0.626058 | 0.730094 | 0.620534 | 0.658895 | down |
| 27175 | TUBG2 | 0.000912 | 0.642679 | 0.579072 | 0.603783 | 0.608511 | down |
| 8851 | CDK5R1 | 0.000912 | 0.430292 | 0.596781 | 0.399408 | 0.475494 | down |
| 90378 | SAMD1 | 0.000912 | 0.610232 | 0.598358 | 0.491108 | 0.566566 | down |
| 25959 | KANK2 | 0.000913 | 0.621027 | 0.638673 | 0.583706 | 0.614469 | down |
| 1.12E+08 | LOC11226 | 0.000916 | 0.552373 | 0.389925 | 0.517004 | 0.486434 | down |
| 54583 | EGLN1 | 0.000918 | 0.604029 | 0.697725 | 0.678313 | 0.660022 | down |
| 64760 | FAM160B2 | 0.000921 | 0.481629 | 0.650531 | 0.612636 | 0.581599 | down |
| 79132 | DHX58 | 0.000931 | 0.70047 | 0.616774 | 0.568442 | 0.628562 | down |
| 55920 | RCC2 | 0.000934 | 0.627503 | 0.714095 | 0.597982 | 0.646527 | down |
| 27153 | ZNF777 | 0.000934 | 0.643999 | 0.664208 | 0.536693 | 0.614966 | down |
| 6601 | SMARCC2 | 0.000939 | 0.677289 | 0.569171 | 0.629244 | 0.625235 | down |
| 4791 | NFKB2 | 0.000942 | 0.596047 | 0.651832 | 0.663422 | 0.6371 | down |
| 7423 | VEGFB | 0.000942 | 0.638722 | 0.668291 | 0.596211 | 0.634408 | down |
| 7405 | UVRAG | 0.000952 | 0.651618 | 0.613064 | 0.69 | 0.651561 | down |
| 221184 | CPNE2 | 0.000959 | 0.675251 | 0.659333 | 0.633585 | 0.656056 | down |
| 162427 | RETREG3 | 0.000959 | 0.532848 | 0.631981 | 0.613426 | 0.592751 | down |
| 29890 | RBM15B | 0.00096 | 0.632397 | 0.735092 | 0.625504 | 0.664331 | down |
| 9762 | LZTS3 | 0.000962 | 0.643612 | 0.617558 | 0.537472 | 0.599547 | down |
| 56006 | SMG9 | 0.000971 | 0.618725 | 0.723668 | 0.6344 | 0.658931 | down |
| 89910 | UBE3B | 0.000973 | 0.691905 | 0.618408 | 0.672213 | 0.660842 | down |
| 9817 | KEAP1 | 0.000974 | 0.654864 | 0.637566 | 0.583236 | 0.625222 | down |
| 153572 | IRX2 | 0.000976 | 0.63698 | 0.699944 | 0.616873 | 0.651266 | down |
| 284273 | ZADH2 | 0.000977 | 0.607724 | 0.645515 | 0.590349 | 0.614529 | down |
| 64114 | TMBIM1 | 0.000977 | 0.675491 | 0.618238 | 0.620198 | 0.637976 | down |
| 6712 | SPTBN2 | 0.000984 | 0.544811 | 0.646578 | 0.533568 | 0.574985 | down |
| 57455 | REXO1 | 0.000993 | 0.639934 | 0.656651 | 0.572917 | 0.623168 | down |
| 23255 | MTCL1 | 0.000996 | 0.519481 | 0.599873 | 0.522824 | 0.547393 | down |
| 727800 | RNF208 | 0.000999 | 0.311456 | 0.463193 | 0.360217 | 0.378289 | down |

Table S4. related to Figure 5

| RNC-seq shNAT10_vs_shControl | | | | RNC-seq shRNPS1_vs_shControl | | | |
|---|---|---|---|---|---|---|---|
| Ingenuity C | -log(p-valu | p-value | Ratio | Ingenuity C | -log(p-valu | p-value | Ratio |
| Molecular I | 17.6 | 2.51E-18 | 0.294 | Pancreatic | 9.76 | 1.74E-10 | 0.238 |
| Rac Signal | 13 | 1E-13 | 0.391 | Chronic My | 8.07 | 8.51E-09 | 0.234 |
| PTEN Sign | 12.4 | 3.98E-13 | 0.373 | Molecular I | 7.69 | 2.04E-08 | 0.135 |
| Actin Cytos | 12.2 | 6.31E-13 | 0.314 | Wnt/β-cate | 6.94 | 1.15E-07 | 0.179 |
| HGF Signa | 11.5 | 3.16E-12 | 0.379 | Glioma Sig | 6.73 | 1.86E-07 | 0.202 |
| Epithelial A | 11.4 | 3.98E-12 | 0.354 | Senescenc | 6.55 | 2.82E-07 | 0.145 |
| Chronic My | 11 | 1E-11 | 0.402 | Regulation | 6.39 | 4.07E-07 | 0.167 |
| Paxillin Sig | 10.8 | 1.58E-11 | 0.398 | PDGF Sigr | 5.9 | 1.26E-06 | 0.221 |
| Neuregulin | 10.7 | 2E-11 | 0.385 | IL-6 Signali | 5.88 | 1.32E-06 | 0.188 |
| FAK Signal | 10.7 | 2E-11 | 0.385 | RAR Activa | 5.71 | 1.95E-06 | 0.158 |
| Pancreatic | 10.5 | 3.16E-11 | 0.373 | Tight Junct | 5.65 | 2.24E-06 | 0.163 |
| Signaling b | 10.5 | 3.16E-11 | 0.291 | Germ Cell- | 5.52 | 3.02E-06 | 0.164 |
| PPARα/RX | 10.4 | 3.98E-11 | 0.321 | Endocanna | 5.52 | 3.02E-06 | 0.175 |
| EGF Signa | 10.3 | 5.01E-11 | 0.509 | RANK Sigr | 5.5 | 3.16E-06 | 0.209 |
| Integrin Sig | 10.3 | 5.01E-11 | 0.31 | EGF Signa | 5.26 | 5.5E-06 | 0.255 |
| Fcγ Recep | 9.85 | 1.41E-10 | 0.404 | Small Cell | 5.14 | 7.24E-06 | 0.198 |
| RAR Activa | 9.63 | 2.34E-10 | 0.311 | Ceramide S | 4.97 | 1.07E-05 | 0.2 |
| HER-2 Sigr | 9.63 | 2.34E-10 | 0.311 | Sertoli Cell | 4.82 | 1.51E-05 | 0.146 |
| Virus Entry | 9.55 | 2.82E-10 | 0.385 | Epithelial A | 4.74 | 1.82E-05 | 0.158 |
| Reelin Sigr | 9.41 | 3.89E-10 | 0.357 | Apelin End | 4.73 | 1.86E-05 | 0.165 |
| AMPK Sigr | 9.36 | 4.37E-10 | 0.289 | AMPK Sigr | 4.63 | 2.34E-05 | 0.136 |
| Insulin Rec | 9.29 | 5.13E-10 | 0.343 | iNOS Signa | 4.61 | 2.45E-05 | 0.255 |
| Sertoli Cell | 9.12 | 7.59E-10 | 0.301 | Autophagy | 4.53 | 2.95E-05 | 0.141 |
| Clathrin-me | 9 | 1E-09 | 0.306 | PPARα/RX | 4.51 | 3.09E-05 | 0.145 |
| Germ Cell- | 8.86 | 1.38E-09 | 0.316 | Signaling b | 4.47 | 3.39E-05 | 0.131 |
| HIPPO sigr | 8.74 | 1.82E-09 | 0.4 | Synaptoge | 4.47 | 3.39E-05 | 0.125 |
| Senescenc | 8.58 | 2.63E-09 | 0.266 | p53 Signali | 4.45 | 3.55E-05 | 0.184 |
| ILK Signali | 8.53 | 2.95E-09 | 0.298 | Telomeras | 4.44 | 3.63E-05 | 0.178 |
| MSP-RON | 8.48 | 3.31E-09 | 0.333 | Acute Myel | 4.33 | 4.68E-05 | 0.187 |
| IGF-1 Sign | 8.37 | 4.27E-09 | 0.365 | CD40 Sign | 4.22 | 6.03E-05 | 0.209 |
| Synaptoge | 8.25 | 5.62E-09 | 0.26 | Cell Cycle: | 4.15 | 7.08E-05 | 0.206 |
| Autophagy | 8.07 | 8.51E-09 | 0.286 | Non-Small | 4.15 | 7.08E-05 | 0.181 |
| Glioma Sig | 8.04 | 9.12E-09 | 0.339 | IL-1 Signali | 4.15 | 7.08E-05 | 0.181 |
| B Cell Rec | 8.02 | 9.55E-09 | 0.294 | Hereditary | 4.1 | 7.94E-05 | 0.155 |
| ERK/MAPK | 7.98 | 1.05E-08 | 0.285 | B Cell Rec | 4.05 | 8.91E-05 | 0.139 |
| PDGF Sigr | 7.97 | 1.07E-08 | 0.384 | RhoA Signa | 4.03 | 9.33E-05 | 0.161 |
| Protein Kin | 7.95 | 1.12E-08 | 0.241 | Acute Phas | 4 | 0.0001 | 0.141 |
| RhoGDI Si | 7.9 | 1.26E-08 | 0.284 | HER-2 Sigr | 3.98 | 0.000105 | 0.138 |
| IL-3 Signali | 7.8 | 1.58E-08 | 0.392 | ILK Signali | 3.9 | 0.000126 | 0.136 |
| Tight Junct | 7.72 | 1.91E-08 | 0.298 | Triacylglyc | 3.9 | 0.000126 | 0.234 |
| Regulation | 7.51 | 3.09E-08 | 0.287 | Lymphotox | 3.89 | 0.000129 | 0.218 |
| Thrombin S | 7.49 | 3.24E-08 | 0.281 | Role of NF | 3.88 | 0.000132 | 0.132 |
| Non-Small | 7.44 | 3.63E-08 | 0.362 | NGF Signa | 3.85 | 0.000141 | 0.161 |
| ErbB Signa | 7.44 | 3.63E-08 | 0.362 | Rac Signal | 3.83 | 0.000148 | 0.152 |
| Telomeras | 7.43 | 3.72E-08 | 0.346 | JAK/Stat S | 3.79 | 0.000162 | 0.183 |
| Regulation | 7.35 | 4.47E-08 | 0.286 | Actin Cytos | 3.78 | 0.000166 | 0.127 |
| Hepatic Fib | 7.02 | 9.55E-08 | 0.232 | ErbB2-ErbI | 3.76 | 0.000174 | 0.2 |
| Renin-Angi | 6.97 | 1.07E-07 | 0.325 | PTEN Sign | 3.75 | 0.000178 | 0.147 |
| JAK/Stat S | 6.77 | 1.7E-07 | 0.366 | Cardiac Hy | 3.72 | 0.000191 | 0.124 |
| RANK Sigr | 6.71 | 1.95E-07 | 0.352 | Leukocyte | 3.69 | 0.000204 | 0.135 |
| Glioblastom | 6.63 | 2.34E-07 | 0.287 | Hepatic Fib | 3.66 | 0.000219 | 0.134 |
| CXCR4 Sig | 6.55 | 2.82E-07 | 0.287 | HGF Signa | 3.65 | 0.000224 | 0.152 |
| FLT3 Signa | 6.47 | 3.39E-07 | 0.362 | Regulation | 3.62 | 0.00024 | 0.133 |
| Wnt/β-cate | 6.46 | 3.47E-07 | 0.283 | ErbB Signa | 3.62 | 0.00024 | 0.17 |
| Leukocyte | 6.46 | 3.47E-07 | 0.275 | PI3K Signa | 3.61 | 0.000245 | 0.147 |
| Role of Tis | 6.41 | 3.89E-07 | 0.319 | Endocanna | 3.61 | 0.000245 | 0.154 |

| | | | | | | | |
|---|---|---|---|---|---|---|---|
| p53 Signali | 6.4 | 3.98E-07 | 0.337 | Virus Entry | 3.59 | 0.000257 | 0.163 |
| PI3K/AKT S | 6.39 | 4.07E-07 | 0.271 | CDP-diacyl | 3.55 | 0.000282 | 0.318 |
| Endometria | 6.37 | 4.27E-07 | 0.4 | IL-15 Signa | 3.54 | 0.000288 | 0.182 |
| Estrogen R | 6.35 | 4.47E-07 | 0.228 | TNFR1 Sig | 3.49 | 0.000324 | 0.212 |
| Sumoylatio | 6.34 | 4.57E-07 | 0.33 | Toll-like Re | 3.48 | 0.000331 | 0.179 |
| Prolactin S | 6.25 | 5.62E-07 | 0.349 | Adrenomec | 3.48 | 0.000331 | 0.131 |
| Mouse Em | 6.23 | 5.89E-07 | 0.327 | IL-2 Signali | 3.44 | 0.000363 | 0.197 |
| ErbB2-ErbI | 6.22 | 6.03E-07 | 0.385 | Neuregulin | 3.43 | 0.000372 | 0.154 |
| NGF Signa | 6.21 | 6.17E-07 | 0.314 | Thyroid Ca | 3.42 | 0.00038 | 0.177 |
| IL-15 Produ | 6.18 | 6.61E-07 | 0.309 | Sphingosin | 3.38 | 0.000417 | 0.153 |
| NF-κB Acti | 6.17 | 6.76E-07 | 0.359 | Aryl Hydroc | 3.38 | 0.000417 | 0.138 |
| IL-7 Signali | 6.17 | 6.76E-07 | 0.359 | Huntington | 3.38 | 0.000417 | 0.117 |
| Axonal Gui | 6.12 | 7.59E-07 | 0.215 | Coronaviru | 3.35 | 0.000447 | 0.128 |
| Apelin End | 6.02 | 9.55E-07 | 0.295 | P2Y Purige | 3.34 | 0.000457 | 0.147 |
| Prostate Ca | 5.87 | 1.35E-06 | 0.312 | Glioblaston | 3.33 | 0.000468 | 0.135 |
| Ephrin Rec | 5.86 | 1.38E-06 | 0.264 | Clathrin-me | 3.32 | 0.000479 | 0.13 |
| VEGF Sign | 5.78 | 1.66E-06 | 0.323 | IL-10 Signa | 3.3 | 0.000501 | 0.181 |
| Unfolded p | 5.77 | 1.7E-06 | 0.333 | Phosphatic | 3.29 | 0.000513 | 0.292 |
| GNRH Sigr | 5.69 | 2.04E-06 | 0.266 | Hepatic Fib | 3.21 | 0.000617 | 0.105 |
| PI3K Signa | 5.67 | 2.14E-06 | 0.287 | Prostate Ca | 3.2 | 0.000631 | 0.152 |
| Small Cell | 5.61 | 2.45E-06 | 0.323 | TNFR2 Sig | 3.19 | 0.000646 | 0.25 |
| IL-8 Signali | 5.53 | 2.95E-06 | 0.256 | Role of Ma | 3.15 | 0.000708 | 0.111 |
| GM-CSF S | 5.53 | 2.95E-06 | 0.357 | TR/RXR Ac | 3.14 | 0.000724 | 0.167 |
| Caveolar-m | 5.45 | 3.55E-06 | 0.347 | Cyclins anc | 3.14 | 0.000724 | 0.167 |
| Ferroptosis | 5.44 | 3.63E-06 | 0.294 | CD27 Sign | 3.13 | 0.000741 | 0.193 |
| TR/RXR Ac | 5.43 | 3.72E-06 | 0.333 | IL-8 Signali | 3.09 | 0.000813 | 0.123 |
| Role of BR | 5.38 | 4.17E-06 | 0.338 | LPS-stimul | 3.09 | 0.000813 | 0.165 |
| Renal Cell | 5.38 | 4.17E-06 | 0.338 | CXCR4 Sig | 3.09 | 0.000813 | 0.132 |
| 3-phosphoi | 5.37 | 4.27E-06 | 0.256 | Cardiac Hy | 3.08 | 0.000832 | 0.0985 |
| Thrombopo | 5.32 | 4.79E-06 | 0.365 | Colorectal | 3.07 | 0.000851 | 0.115 |
| Cell Cycle: | 5.23 | 5.89E-06 | 0.353 | IL-17A Sigr | 3.06 | 0.000871 | 0.179 |
| Acute Myel | 5.16 | 6.92E-06 | 0.319 | Reelin Sigr | 3.04 | 0.000912 | 0.143 |
| Superpathv | 5.15 | 7.08E-06 | 0.245 | Integrin Sig | 3.03 | 0.000933 | 0.122 |
| Role of NF. | 4.96 | 1.1E-05 | 0.245 | Production | 3.02 | 0.000955 | 0.126 |
| 14-3-3-mec | 4.94 | 1.15E-05 | 0.283 | Melanoma | 3.02 | 0.000955 | 0.2 |
| p70S6K Si | 4.93 | 1.17E-05 | 0.28 | ERK/MAPK | 3 | 0.001 | 0.121 |
| Huntington | 4.92 | 1.2E-05 | 0.231 | April Media | 2.99 | 0.001023 | 0.214 |
| PAK Signa | 4.86 | 1.38E-05 | 0.288 | FAK Signal | 2.98 | 0.001047 | 0.145 |
| Endocanna | 4.86 | 1.38E-05 | 0.273 | MSP-RON | 2.97 | 0.001072 | 0.138 |
| IL-6 Signali | 4.85 | 1.41E-05 | 0.281 | Endometria | 2.94 | 0.001148 | 0.183 |
| RhoA Sign | 4.77 | 1.7E-05 | 0.282 | B Cell Activ | 2.92 | 0.001202 | 0.209 |
| IL-15 Signa | 4.69 | 2.04E-05 | 0.325 | Stearate Bi | 2.82 | 0.001514 | 0.189 |
| Amyloid Pr | 4.65 | 2.24E-05 | 0.373 | Gap Juncti | 2.8 | 0.001585 | 0.121 |
| VDR/RXR | 4.59 | 2.57E-05 | 0.321 | Superpathv | 2.79 | 0.001622 | 0.116 |
| Hereditary | 4.55 | 2.82E-05 | 0.268 | Gα12/13 S | 2.76 | 0.001738 | 0.135 |
| CDK5 Sign | 4.53 | 2.95E-05 | 0.286 | GNRH Sigr | 2.76 | 0.001738 | 0.122 |
| GP6 Signa | 4.53 | 2.95E-05 | 0.276 | Role of p14 | 2.75 | 0.001778 | 0.241 |
| UVA-Induc | 4.48 | 3.31E-05 | 0.296 | Tumor Micr | 2.69 | 0.002042 | 0.123 |
| IL-2 Signali | 4.47 | 3.39E-05 | 0.344 | STAT3 Pat | 2.69 | 0.002042 | 0.133 |
| Agrin Intera | 4.46 | 3.47E-05 | 0.329 | Endothelin- | 2.67 | 0.002138 | 0.12 |
| Endocanna | 4.45 | 3.55E-05 | 0.276 | Leptin Sign | 2.67 | 0.002138 | 0.162 |
| FGF Signa | 4.45 | 3.55E-05 | 0.31 | Melanocyte | 2.66 | 0.002188 | 0.149 |
| Insulin Sec | 4.43 | 3.72E-05 | 0.228 | 3-phosphoi | 2.66 | 0.002188 | 0.118 |
| Regulation | 4.42 | 3.8E-05 | 0.303 | RhoGDI Si | 2.64 | 0.002291 | 0.116 |
| CNTF Sign | 4.42 | 3.8E-05 | 0.351 | Gαq Signal | 2.62 | 0.002399 | 0.124 |
| Melanocyte | 4.4 | 3.98E-05 | 0.298 | HIPPO sigr | 2.61 | 0.002455 | 0.153 |
| LPS-stimul | 4.35 | 4.47E-05 | 0.306 | Bladder Ca | 2.6 | 0.002512 | 0.138 |
| Macropino | 4.31 | 4.9E-05 | 0.316 | Mitotic Role | 2.59 | 0.00257 | 0.167 |
| CD40 Sign | 4.29 | 5.13E-05 | 0.328 | Death Rec | 2.57 | 0.002692 | 0.146 |

| | | | |
|---|---|---|---|
| Ephrin B Si | 4.24 | 5.75E-05 | 0.319 |
| Death Rec | 4.22 | 6.03E-05 | 0.292 |
| TGF-β Sigr | 4.22 | 6.03E-05 | 0.292 |
| ErbB4 Sigr | 4.18 | 6.61E-05 | 0.324 |
| Notch Sign | 4.13 | 7.41E-05 | 0.395 |
| Aldosteron | 4.12 | 7.59E-05 | 0.25 |
| Production | 4.11 | 7.76E-05 | 0.241 |
| Colorectal | 4.11 | 7.76E-05 | 0.223 |
| Protein Ubi | 4.09 | 8.13E-05 | 0.222 |
| Cardiac Hy | 4.09 | 8.13E-05 | 0.225 |
| Gα12/13 S | 4.08 | 8.32E-05 | 0.263 |
| Gαq Signal | 4.07 | 8.51E-05 | 0.247 |
| Coronaviru | 4.05 | 8.91E-05 | 0.236 |
| Ephrin A Si | 4.03 | 9.33E-05 | 0.362 |
| Gap Juncti | 4.02 | 9.55E-05 | 0.237 |
| Natural Kill | 3.97 | 0.000107 | 0.236 |
| IL-9 Signali | 3.96 | 0.00011 | 0.4 |
| UVB-Induc | 3.95 | 0.000112 | 0.346 |
| Adipogene | 3.94 | 0.000115 | 0.259 |
| Dilated Car | 3.91 | 0.000123 | 0.253 |
| Pyridoxal 5 | 3.9 | 0.000126 | 0.318 |
| Regulation | 3.86 | 0.000138 | 0.24 |
| fMLP Signa | 3.86 | 0.000138 | 0.26 |
| Aryl Hydro | 3.76 | 0.000174 | 0.245 |
| Angiopoieti | 3.75 | 0.000178 | 0.299 |
| Fc Epsilon | 3.74 | 0.000182 | 0.265 |
| PPAR Sign | 3.73 | 0.000186 | 0.271 |
| P2Y Purige | 3.63 | 0.000234 | 0.256 |
| Cholecysto | 3.6 | 0.000251 | 0.261 |
| Lymphotox | 3.6 | 0.000251 | 0.327 |
| Semaphori | 3.59 | 0.000257 | 0.245 |
| Thyroid Ca | 3.57 | 0.000269 | 0.291 |
| April Media | 3.56 | 0.000275 | 0.357 |
| UVC-Induc | 3.54 | 0.000288 | 0.333 |
| Type II Dia | 3.53 | 0.000295 | 0.243 |
| Ceramide S | 3.47 | 0.000339 | 0.278 |
| Cardiac Hy | 3.44 | 0.000363 | 0.19 |
| B Cell Activ | 3.43 | 0.000372 | 0.349 |
| Oncostatin | 3.43 | 0.000372 | 0.349 |
| D-myo-inos | 3.4 | 0.000398 | 0.228 |
| tRNA Char | 3.38 | 0.000417 | 0.359 |
| Semaphori | 3.35 | 0.000447 | 0.306 |
| IL-17A Sigr | 3.33 | 0.000468 | 0.299 |
| ERK5 Sign | 3.32 | 0.000479 | 0.292 |
| 3-phosphoi | 3.31 | 0.00049 | 0.228 |
| mTOR Sigr | 3.3 | 0.000501 | 0.222 |
| Spliceoson | 3.25 | 0.000562 | 0.327 |
| Remodelin | 3.24 | 0.000575 | 0.294 |
| Androgen S | 3.19 | 0.000646 | 0.231 |
| IL-1 Signali | 3.16 | 0.000692 | 0.266 |
| Cyclins anc | 3.15 | 0.000708 | 0.274 |
| PEDF Sign | 3.15 | 0.000708 | 0.274 |
| Leptin Sign | 3.15 | 0.000708 | 0.284 |
| Estrogen-D | 3.14 | 0.000724 | 0.278 |
| Phosphatic | 3.14 | 0.000724 | 0.417 |
| Melanoma | 3.14 | 0.000724 | 0.32 |
| Apelin Pan | 3.07 | 0.000851 | 0.326 |
| D-myo-inos | 3.07 | 0.000851 | 0.226 |
| D-myo-inos | 3.07 | 0.000851 | 0.226 |

| | | | |
|---|---|---|---|
| Macropinoc | 2.57 | 0.002692 | 0.158 |
| HOTAIR R | 2.49 | 0.003236 | 0.123 |
| Thrombin S | 2.46 | 0.003467 | 0.114 |
| Renin-Angi | 2.45 | 0.003548 | 0.133 |
| PI3K/AKT S | 2.44 | 0.003631 | 0.116 |
| FAT10 Car | 2.44 | 0.003631 | 0.18 |
| IL-3 Signali | 2.43 | 0.003715 | 0.152 |
| Circadian F | 2.41 | 0.00389 | 0.212 |
| Role of BR | 2.38 | 0.004169 | 0.15 |
| Renal Cell | 2.38 | 0.004169 | 0.15 |
| CDK5 Sign | 2.35 | 0.004467 | 0.134 |
| BEX2 Sign | 2.34 | 0.004571 | 0.148 |
| Role of JAK | 2.34 | 0.004571 | 0.206 |
| Adipogene | 2.31 | 0.004898 | 0.126 |
| IL-4 Signali | 2.31 | 0.004898 | 0.141 |
| Systemic L | 2.3 | 0.005012 | 0.105 |
| Assembly c | 2.29 | 0.005129 | 0.333 |
| IGF-1 Sign | 2.25 | 0.005623 | 0.135 |
| Thrombopo | 2.24 | 0.005754 | 0.159 |
| 14-3-3-mec | 2.21 | 0.006166 | 0.126 |
| Role of Tis | 2.21 | 0.006166 | 0.129 |
| PEDF Sign | 2.21 | 0.006166 | 0.143 |
| Semaphori | 2.17 | 0.006761 | 0.119 |
| Insulin Rec | 2.16 | 0.006918 | 0.121 |
| Induction o | 2.15 | 0.007079 | 0.154 |
| PPAR Sign | 2.14 | 0.007244 | 0.131 |
| p38 MAPK | 2.14 | 0.007244 | 0.127 |
| Natural Kill | 2.14 | 0.007244 | 0.111 |
| IL-23 Signa | 2.13 | 0.007413 | 0.174 |
| Prolactin S | 2.13 | 0.007413 | 0.14 |
| Role of Ost | 2.12 | 0.007586 | 0.107 |
| ATM Signa | 2.11 | 0.007762 | 0.134 |
| Relaxin Sig | 2.11 | 0.007762 | 0.118 |
| GDNF Fam | 2.11 | 0.007762 | 0.145 |
| Neurotroph | 2.11 | 0.007762 | 0.145 |
| UVA-Induc | 2.08 | 0.008318 | 0.133 |
| Role of IL-1 | 2.06 | 0.00871 | 0.158 |
| VEGF Sign | 2.04 | 0.00912 | 0.131 |
| NF-κB Acti | 2.03 | 0.009333 | 0.141 |
| IL-7 Signali | 2.03 | 0.009333 | 0.141 |

| | | | |
|---|---|---|---|
| Opioid Sigr | 3.04 | 0.000912 | 0.207 |
| RAN Signa | 3.02 | 0.000955 | 0.471 |
| Systemic L | 3 | 0.001 | 0.206 |
| Growth Ho | 2.98 | 0.001047 | 0.282 |
| Acute Phas | 2.95 | 0.001122 | 0.222 |
| HIF1α Sigr | 2.95 | 0.001122 | 0.216 |
| Docosahex | 2.95 | 0.001122 | 0.342 |
| IL-4 Signali | 2.94 | 0.001148 | 0.261 |
| Regulation | 2.92 | 0.001202 | 0.264 |
| HOTAIR R | 2.92 | 0.001202 | 0.227 |
| Ovarian Ca | 2.89 | 0.001288 | 0.228 |
| Actin Nucle | 2.86 | 0.00138 | 0.258 |
| Adrenomec | 2.84 | 0.001445 | 0.216 |
| Glioma Inv | 2.82 | 0.001514 | 0.274 |
| CDP-diacyl | 2.81 | 0.001549 | 0.409 |
| Hepatic Fib | 2.8 | 0.001585 | 0.216 |
| Erythropoie | 2.79 | 0.001622 | 0.22 |
| Role of Ma | 2.79 | 0.001622 | 0.197 |
| Xenobiotic | 2.78 | 0.00166 | 0.201 |
| Regulation | 2.76 | 0.001738 | 0.241 |
| Wnt/Ca+ p | 2.73 | 0.001862 | 0.281 |
| NRF2-med | 2.71 | 0.00195 | 0.207 |
| α-Adrenerg | 2.71 | 0.00195 | 0.245 |
| FAT10 Car | 2.66 | 0.002188 | 0.3 |
| Sphingosin | 2.64 | 0.002291 | 0.237 |
| BEX2 Sign | 2.6 | 0.002512 | 0.259 |
| Neurotroph | 2.59 | 0.00257 | 0.263 |
| Cell Cycle | 2.56 | 0.002754 | 0.286 |
| Apoptosis | 2.5 | 0.003162 | 0.24 |
| Vitamin-C | 2.5 | 0.003162 | 0.375 |
| iNOS Signa | 2.49 | 0.003236 | 0.298 |
| Role of IL- | 2.48 | 0.003311 | 0.281 |
| Apelin Carc | 2.47 | 0.003388 | 0.242 |
| Sirtuin Sigr | 2.46 | 0.003467 | 0.195 |
| Iron homec | 2.42 | 0.003802 | 0.223 |
| Role of p1 | 2.42 | 0.003802 | 0.345 |
| Cancer Dru | 2.39 | 0.004074 | 0.276 |
| 4-1BB Sigr | 2.36 | 0.004365 | 0.324 |
| Role of JAI | 2.36 | 0.004365 | 0.324 |
| Role of JAI | 2.36 | 0.004365 | 0.36 |
| Role of JAI | 2.34 | 0.004571 | 0.261 |
| CCR3 Sigr | 2.34 | 0.004571 | 0.222 |
| Endoplasm | 2.32 | 0.004786 | 0.381 |
| IL-12 Signa | 2.29 | 0.005129 | 0.221 |
| ATM Signa | 2.27 | 0.00537 | 0.237 |
| Role of JAI | 2.23 | 0.005888 | 0.346 |
| Xenobiotic | 2.23 | 0.005888 | 0.217 |
| Cell Cycle: | 2.22 | 0.006026 | 0.28 |
| BMP signa | 2.21 | 0.006166 | 0.241 |
| Calcium Si | 2.15 | 0.007079 | 0.199 |
| Role of Oct | 2.14 | 0.007244 | 0.283 |
| PFKFB4 Si | 2.14 | 0.007244 | 0.283 |
| IL-23 Signa | 2.14 | 0.007244 | 0.283 |
| IL-10 Signa | 2.13 | 0.007413 | 0.25 |
| Necroptosi | 2.12 | 0.007586 | 0.21 |
| Role of Ost | 2.08 | 0.008318 | 0.196 |
| CD27 Sign | 2.07 | 0.008511 | 0.263 |
| Role of CH | 2.07 | 0.008511 | 0.263 |
| STAT3 Pat | 2.06 | 0.00871 | 0.215 |

| | | | |
|---|---|---|---|
| TNFR1 Sig | 2.06 | 0.00871 | 0.269 |

Table S5. related to Figure 5

| Gene Syml | shControl | shNAT10 a | shNAT10/shControl | |
|---|---|---|---|---|
| SHF | 4.425749 | 0.827645 | 0.187007 | down |
| ZNF282 | 4.88398 | 1.300318 | 0.266241 | down |
| CNTN5 | 6.21535 | 1.791456 | 0.288231 | down |
| CABP2 | 3.796707 | 1.242144 | 0.327164 | down |
| CX3CR1 | 3.35626 | 1.098439 | 0.327281 | down |
| KNDC1 | 2.963975 | 1.051419 | 0.354733 | down |
| MANEAL | 2.470626 | 0.899808 | 0.364202 | down |
| ZNF469 | 3.346232 | 1.285533 | 0.384173 | down |
| NEB | 2.114149 | 0.822729 | 0.389154 | down |
| APOLD1 | 2.493969 | 0.989418 | 0.396724 | down |
| SPHKAP | 3.045546 | 1.250414 | 0.410571 | down |
| FBP2 | 2.20345 | 0.939069 | 0.426181 | down |
| PLG | 2.088781 | 0.923509 | 0.442128 | down |
| SPZ1 | 3.548902 | 1.586177 | 0.446949 | down |
| MTTP | 3.639659 | 1.728097 | 0.474797 | down |
| TIMP2 | 0.771476 | 0.372195 | 0.482446 | down |
| SDC4 | 2.139825 | 1.04968 | 0.490545 | down |
| PUDP | 2.095482 | 1.028926 | 0.491021 | down |
| MFAP3 | 1.630508 | 0.818294 | 0.501864 | down |
| NOS2 | 2.503813 | 1.259048 | 0.502852 | down |
| SEMA6D | 1.781326 | 0.911506 | 0.511701 | down |
| ABHD13 | 1.729631 | 0.907002 | 0.524391 | down |
| KRT4 | 2.528086 | 1.342357 | 0.530978 | down |
| SPPL2B | 1.585254 | 0.84358 | 0.532142 | down |
| ZNF750 | 2.499557 | 1.341764 | 0.536801 | down |
| HPX | 1.801603 | 0.976309 | 0.541911 | down |
| TTC23L | 1.787451 | 0.970741 | 0.543086 | down |
| SRPX2 | 1.891184 | 1.032152 | 0.54577 | down |
| GPR162 | 2.208801 | 1.22826 | 0.556075 | down |
| PCDHB7 | 1.969438 | 1.097482 | 0.557257 | down |
| FCRL5 | 1.495462 | 0.842349 | 0.56327 | down |
| LAPTM4B | 2.159351 | 1.218231 | 0.564165 | down |
| PRXL2A | 2.082753 | 1.178434 | 0.565806 | down |
| HYI | 1.865319 | 1.062651 | 0.569688 | down |
| TMBIM1 | 1.639688 | 0.943929 | 0.575676 | down |
| FUT3 | 1.733304 | 1.011905 | 0.583801 | down |
| ERVMER34 | 1.504224 | 0.884757 | 0.588182 | down |
| FDFT1 | 1.584216 | 0.931923 | 0.588255 | down |
| L1CAM | 1.745017 | 1.033569 | 0.592297 | down |
| PRSS21 | 2.085428 | 1.249419 | 0.599119 | down |
| F10 | 1.729791 | 1.038104 | 0.600133 | down |
| KRT16 | 1.78061 | 1.071344 | 0.601672 | down |
| COL11A1 | 2.018794 | 1.220771 | 0.604703 | down |
| FAM102B | 1.783326 | 1.081163 | 0.606262 | down |
| TPRG1L | 1.415043 | 0.858818 | 0.606921 | down |
| NOLC1 | 2.231812 | 1.356815 | 0.607943 | down |
| ARL8B | 1.487098 | 0.905524 | 0.60892 | down |
| RAB10 | 1.609323 | 0.982483 | 0.610494 | down |
| OLR1 | 1.645124 | 1.009377 | 0.613557 | down |
| ART4 | 2.107053 | 1.292859 | 0.613586 | down |
| CLSTN3 | 1.847771 | 1.1395 | 0.616689 | down |
| FBXO41 | 1.857088 | 1.149401 | 0.618927 | down |
| PLEKHA7 | 1.495196 | 0.926421 | 0.619598 | down |
| EDNRA | 1.519494 | 0.941945 | 0.619907 | down |
| KRT23 | 1.498222 | 0.92904 | 0.620095 | down |
| GPR84 | 1.693992 | 1.050676 | 0.620237 | down |

| | | | | |
|---|---|---|---|---|
| MSMO1 | 1.361826 | 0.846236 | 0.621398 | down |
| TMEM222 | 1.395507 | 0.872685 | 0.625353 | down |
| NID1 | 1.636056 | 1.023506 | 0.625594 | down |
| TNFSF9 | 1.497548 | 0.937576 | 0.626074 | down |
| CLCN3 | 1.518827 | 0.953952 | 0.628085 | down |
| EPB41L4B | 0.972696 | 0.610971 | 0.628121 | down |
| LRRTM2 | 1.395444 | 0.881636 | 0.631796 | down |
| SH3BGR | 1.369268 | 0.866722 | 0.632982 | down |
| TNFRSF12/ | 1.48769 | 0.94459 | 0.634938 | down |
| SSR3 | 1.19891 | 0.761604 | 0.635247 | down |
| SERINC3 | 1.398802 | 0.889864 | 0.636161 | down |
| BORCS8 | 1.613252 | 1.026607 | 0.636359 | down |
| KRT76 | 1.779164 | 1.132832 | 0.636721 | down |
| VSIR | 1.248445 | 0.796067 | 0.637647 | down |
| KRT17 | 1.810094 | 1.154211 | 0.637653 | down |
| ACOT4 | 1.393891 | 0.890887 | 0.639137 | down |
| KRT13 | 2.016554 | 1.292138 | 0.640765 | down |
| RTN3 | 1.756528 | 1.125853 | 0.640954 | down |
| ITIH6 | 1.707737 | 1.09465 | 0.640995 | down |
| HM13 | 1.394813 | 0.895048 | 0.641697 | down |
| HERPUD2 | 1.475771 | 0.948312 | 0.642588 | down |
| CLN3 | 1.602291 | 1.030646 | 0.643232 | down |
| LCN2 | 1.601045 | 1.031482 | 0.644256 | down |
| HLA-C | 1.35337 | 0.872029 | 0.644339 | down |
| INPP5D | 1.058999 | 0.683447 | 0.645371 | down |
| SELENOI | 1.347702 | 0.870897 | 0.646208 | down |
| HMGCR | 1.84293 | 1.191231 | 0.646379 | down |
| CXCL16 | 1.547461 | 1.001155 | 0.646967 | down |
| GPSM3 | 1.662016 | 1.076063 | 0.647445 | down |
| GRIN2D | 1.899048 | 1.244749 | 0.65546 | down |
| SSR2 | 1.231094 | 0.807525 | 0.655941 | down |
| ADA2 | 1.617705 | 1.063446 | 0.65738 | down |
| SCD5 | 1.597024 | 1.050604 | 0.657851 | down |
| NDFIP1 | 1.488068 | 0.981481 | 0.659567 | down |
| SQLE | 1.480438 | 0.977445 | 0.66024 | down |
| SLC35F6 | 1.212065 | 0.80117 | 0.660996 | down |
| CD58 | 1.513695 | 1.004823 | 0.663821 | down |
| ADIPOR2 | 1.341249 | 0.891852 | 0.664941 | down |
| ACVR1 | 1.592964 | 1.060028 | 0.665444 | down |
| APOE | 1.352521 | 0.909173 | 0.672206 | down |
| SCD | 1.365224 | 0.918679 | 0.672914 | down |
| PLGRKT | 1.549742 | 1.043498 | 0.673337 | down |
| ABHD2 | 1.740889 | 1.172534 | 0.673526 | down |
| TMEM147 | 1.658798 | 1.117309 | 0.673566 | down |
| CFAP46 | 0.538765 | 0.363114 | 0.673975 | down |
| TMEM97 | 1.632101 | 1.101122 | 0.674665 | down |
| TMEM59 | 1.375178 | 0.928387 | 0.675103 | down |
| BEND4 | 1.885441 | 1.275374 | 0.676433 | down |
| ANKRA2 | 1.644609 | 1.114602 | 0.677731 | down |
| ARL6IP4 | 1.564623 | 1.061222 | 0.67826 | down |
| ATL1 | 1.180058 | 0.800685 | 0.678513 | down |
| ABCA12 | 1.511838 | 1.030066 | 0.681334 | down |
| REEP2 | 1.438393 | 0.980195 | 0.681452 | down |
| RNF26 | 0.697382 | 0.475572 | 0.68194 | down |
| TSPO | 1.491048 | 1.017587 | 0.682464 | down |
| STC1 | 1.257661 | 0.859884 | 0.683717 | down |
| SPRR3 | 1.677631 | 1.149007 | 0.684899 | down |
| JUN | 1.330773 | 0.913103 | 0.686145 | down |

| | | | | |
|---|---|---|---|---|
| SHB | 1.73747 | 1.192236 | 0.686191 | down |
| LAMTOR3 | 1.620503 | 1.112645 | 0.686605 | down |
| MUC1 | 1.360089 | 0.935333 | 0.6877 | down |
| LOXL2 | 1.45515 | 1.002309 | 0.688801 | down |
| TMEM40 | 1.2344 | 0.850466 | 0.688971 | down |
| ETV4 | 1.513889 | 1.043285 | 0.689142 | down |
| TERB2 | 1.460913 | 1.008279 | 0.690171 | down |
| PLEKHG2 | 1.19983 | 0.828914 | 0.690859 | down |
| ICAM1 | 1.295737 | 0.896493 | 0.691879 | down |
| B2M | 1.475936 | 1.022092 | 0.692504 | down |
| JPH2 | 1.53547 | 1.064174 | 0.693061 | down |
| PGS1 | 1.402972 | 0.973727 | 0.694046 | down |
| TMEM35B | 1.456385 | 1.010847 | 0.69408 | down |
| MED21 | 1.393552 | 0.967852 | 0.694522 | down |
| CD63 | 1.224143 | 0.851075 | 0.695242 | down |
| HLA-B | 1.449723 | 1.008126 | 0.695392 | down |
| ARMC10 | 1.228503 | 0.854547 | 0.6956 | down |
| SRD5A3 | 1.371604 | 0.955806 | 0.696853 | down |
| CIB4 | 4.344682 | 3.027893 | 0.696919 | down |
| CCDC183 | 1.708283 | 1.190714 | 0.697024 | down |
| TK2 | 1.537591 | 1.07207 | 0.69724 | down |
| PPM1J | 1.048614 | 0.731194 | 0.697295 | down |
| F8 | 1.595564 | 1.113535 | 0.697894 | down |
| NDUFB2 | 1.808065 | 1.261984 | 0.697975 | down |
| TTC34 | 1.846226 | 1.291761 | 0.699677 | down |
| ZBED8 | 1.640617 | 1.148098 | 0.699797 | down |
| LGALS3BP | 1.421921 | 0.996242 | 0.700631 | down |
| ZNF30 | 1.426455 | 0.999477 | 0.700672 | down |
| KRT14 | 1.529099 | 1.072725 | 0.70154 | down |
| DHCR7 | 1.432278 | 1.005337 | 0.701915 | down |
| C3 | 1.368629 | 0.960683 | 0.701931 | down |
| TPBG | 1.43003 | 1.004012 | 0.702091 | down |
| RP1L1 | 1.307461 | 0.918884 | 0.7028 | down |
| TMEM19 | 1.762263 | 1.23879 | 0.702955 | down |
| MMP1 | 1.392683 | 0.979018 | 0.702973 | down |
| RNF13 | 1.44844 | 1.018607 | 0.703244 | down |
| SOD2 | 1.358794 | 0.957271 | 0.704501 | down |
| LZTS3 | 1.233393 | 0.870447 | 0.705734 | down |
| ARL8A | 1.597061 | 1.12801 | 0.706304 | down |
| ZNF174 | 1.286413 | 0.90929 | 0.706842 | down |
| CLDN7 | 1.313999 | 0.931249 | 0.708713 | down |
| STK36 | 2.395521 | 1.698694 | 0.709112 | down |
| PLEC | 1.606593 | 1.139424 | 0.709218 | down |
| SLC2A1 | 1.378769 | 0.978381 | 0.709605 | down |
| MANSC1 | 1.263806 | 0.897697 | 0.710312 | down |
| TSPAN31 | 1.399601 | 0.994465 | 0.710534 | down |
| TRIM29 | 1.479648 | 1.051467 | 0.710619 | down |
| FGFBP1 | 1.426456 | 1.014387 | 0.711124 | down |
| FAAP24 | 1.585654 | 1.128646 | 0.711786 | down |
| ADGRG1 | 1.446019 | 1.030525 | 0.712664 | down |
| TPTE2 | 1.532367 | 1.09439 | 0.714183 | down |
| SLX1A | 1.408237 | 1.006421 | 0.714667 | down |
| SLC25A1 | 1.488707 | 1.064402 | 0.714984 | down |
| RAP2A | 1.298298 | 0.928371 | 0.715068 | down |
| PKP1 | 1.217714 | 0.871093 | 0.715351 | down |
| ATP6V0D2 | 1.541695 | 1.103256 | 0.715612 | down |
| GCSH | 1.288419 | 0.923737 | 0.716954 | down |
| SC5D | 1.358717 | 0.97476 | 0.717412 | down |

| | | | | |
|---|---|---|---|---|
| KRT1 | 1.781503 | 1.278161 | 0.717462 | down |
| MAPK8IP1 | 1.81266 | 1.302708 | 0.718672 | down |
| LRMDA | 1.386466 | 0.996732 | 0.718901 | down |
| PLPP1 | 1.306733 | 0.940075 | 0.719409 | down |
| ZDHHC7 | 1.240062 | 0.892202 | 0.719482 | down |
| RP1 | 1.512054 | 1.089137 | 0.720303 | down |
| GPR87 | 1.366448 | 0.986246 | 0.721759 | down |
| MREG | 1.092806 | 0.791098 | 0.723914 | down |
| TRPM4 | 1.660408 | 1.202482 | 0.724209 | down |
| ALDH1L2 | 1.352489 | 0.97978 | 0.724428 | down |
| OGDHL | 1.393787 | 1.011815 | 0.725947 | down |
| APP | 1.355308 | 0.984376 | 0.726312 | down |
| MRPL54 | 1.349362 | 0.980605 | 0.726718 | down |
| CEP250 | 1.169454 | 0.850283 | 0.727077 | down |
| HSDL1 | 1.402398 | 1.01997 | 0.727305 | down |
| SIRT1 | 1.646062 | 1.199219 | 0.728538 | down |
| CDSN | 1.328014 | 0.968202 | 0.72906 | down |
| TMEM150/ | 1.015958 | 0.74185 | 0.730197 | down |
| LAMB2 | 1.391394 | 1.017239 | 0.731093 | down |
| APOBEC3( | 1.188142 | 0.870872 | 0.73297 | down |
| TRIM22 | 1.468839 | 1.077993 | 0.733908 | down |
| TBRG1 | 1.243343 | 0.91332 | 0.734568 | down |
| ISOC2 | 1.508829 | 1.1086 | 0.734742 | down |
| OSTC | 1.302053 | 0.957799 | 0.735607 | down |
| BCKDHB | 1.365472 | 1.004593 | 0.735711 | down |
| HSPG2 | 1.420944 | 1.046042 | 0.73616 | down |
| DMWD | 1.312725 | 0.966759 | 0.736452 | down |
| ABAT | 1.44478 | 1.065542 | 0.737511 | down |
| BRI3 | 1.017024 | 0.750576 | 0.738012 | down |
| KRT15 | 1.50907 | 1.113941 | 0.738164 | down |
| SPINT2 | 1.45438 | 1.073726 | 0.738271 | down |
| IL10RB | 1.33468 | 0.98563 | 0.738477 | down |
| NIPSNAP3 | 1.397479 | 1.032115 | 0.738555 | down |
| SLC15A4 | 1.304752 | 0.963953 | 0.738802 | down |
| CLDN2 | 1.27425 | 0.941455 | 0.73883 | down |
| INS | 1.903939 | 1.409012 | 0.740051 | down |
| BAK1 | 1.248623 | 0.924059 | 0.740063 | down |
| TUBGCP4 | 1.44373 | 1.068747 | 0.740268 | down |
| ZNFX1 | 1.427137 | 1.056504 | 0.740296 | down |
| CMTM6 | 1.186005 | 0.878218 | 0.740484 | down |
| ITPRIPL1 | 1.501941 | 1.112444 | 0.740671 | down |
| SLC18A3 | 1.254059 | 0.928959 | 0.740762 | down |
| LMBRD1 | 1.331434 | 0.986731 | 0.741104 | down |
| PMFBP1 | 1.412918 | 1.047741 | 0.741544 | down |
| CFAP300 | 2.436094 | 1.806509 | 0.74156 | down |
| PAH | 1.089252 | 0.80798 | 0.741776 | down |
| ASAH1 | 1.300481 | 0.964873 | 0.741935 | down |
| APOC3 | 1.32461 | 0.983499 | 0.742482 | down |
| RAP1A | 1.408478 | 1.046662 | 0.743115 | down |
| SOX6 | 1.460098 | 1.085397 | 0.743372 | down |
| PAFAH2 | 1.217966 | 0.90578 | 0.743683 | down |
| LDAH | 1.361625 | 1.014225 | 0.744864 | down |
| MGST3 | 1.37794 | 1.027175 | 0.745443 | down |
| PSAP | 1.346283 | 1.003763 | 0.745581 | down |
| MME | 1.334346 | 0.996189 | 0.746575 | down |
| CLCN6 | 1.75135 | 1.307945 | 0.746821 | down |
| RAB5B | 1.401106 | 1.046403 | 0.746841 | down |
| SLC39A3 | 1.195745 | 0.893109 | 0.746906 | down |

| | | | | |
|---|---|---|---|---|
| SIGMAR1 | 1.430115 | 1.068299 | 0.747002 | down |
| TMX4 | 1.21167 | 0.905273 | 0.747128 | down |
| RAB4B | 1.281485 | 0.958488 | 0.747951 | down |
| TADA3 | 1.268859 | 0.94911 | 0.748003 | down |
| NDUFA3 | 1.445563 | 1.081406 | 0.748086 | down |
| PLXNB3 | 1.546252 | 1.156938 | 0.748221 | down |
| CLK4 | 1.503194 | 1.124894 | 0.748336 | down |
| PCNX1 | 1.547901 | 1.158664 | 0.748539 | down |
| GJB3 | 1.528261 | 1.14408 | 0.748616 | down |
| ESCO1 | 1.022334 | 0.765992 | 0.749259 | down |
| ZFP28 | 1.319206 | 0.988796 | 0.749539 | down |
| ADGRF4 | 1.22978 | 0.922358 | 0.750018 | down |
| PEG10 | 0.661465 | 0.496112 | 0.750019 | down |
| DHRS13 | 1.326918 | 0.995621 | 0.750325 | down |
| DHCR24 | 1.341213 | 1.00656 | 0.750485 | down |
| CHCHD7 | 1.200695 | 0.901261 | 0.750616 | down |
| CHADL | 1.32533 | 0.994858 | 0.750649 | down |
| ZNF585A | 1.119418 | 0.840348 | 0.750701 | down |
| RBP3 | 1.470356 | 1.103871 | 0.75075 | down |
| THBS2 | 1.501865 | 1.127846 | 0.750964 | down |
| ACSL1 | 1.323423 | 0.99389 | 0.750999 | down |
| C6orf203 | 1.333223 | 1.001851 | 0.751451 | down |
| ZNF205 | 1.301418 | 0.978516 | 0.751884 | down |
| MOS | 1.292261 | 0.972485 | 0.752545 | down |
| LAMP2 | 1.272548 | 0.957824 | 0.752682 | down |
| CCNL1 | 1.414252 | 1.064777 | 0.752891 | down |
| WWTR1 | 0.671941 | 0.506126 | 0.753229 | down |
| HLA-E | 1.415372 | 1.066475 | 0.753494 | down |
| FAM167B | 1.136459 | 0.857304 | 0.754365 | down |
| 44257 | 1.267452 | 0.956406 | 0.754589 | down |
| FBLN1 | 1.267888 | 0.957668 | 0.755326 | down |
| GBP5 | 1.332015 | 1.006819 | 0.755861 | down |
| ARL6IP5 | 1.328911 | 1.004623 | 0.755975 | down |
| ACP5 | 1.200585 | 0.907613 | 0.755976 | down |
| LIMK1 | 1.983731 | 1.500356 | 0.756331 | down |
| SDCBP | 1.224521 | 0.92641 | 0.756549 | down |
| SLC25A20 | 1.299282 | 0.983351 | 0.756842 | down |
| SLC1A4 | 1.129657 | 0.855035 | 0.756898 | down |
| ELOVL5 | 1.269853 | 0.961471 | 0.757151 | down |
| RAP2C | 1.342955 | 1.017127 | 0.75738 | down |
| CTSB | 1.33052 | 1.008133 | 0.757699 | down |
| KDELR3 | 1.186312 | 0.899374 | 0.758126 | down |
| SLC25A29 | 1.393704 | 1.057426 | 0.758716 | down |
| TNFRSF1A | 1.295997 | 0.983754 | 0.759071 | down |
| MPDU1 | 1.431166 | 1.087248 | 0.759694 | down |
| TIMP1 | 1.313899 | 0.998258 | 0.759768 | down |
| MLLT6 | 1.26579 | 0.962081 | 0.760063 | down |
| DENND1A | 1.698215 | 1.290884 | 0.760141 | down |
| RPUSD4 | 1.206923 | 0.917814 | 0.760458 | down |
| PYURF | 1.527474 | 1.161746 | 0.760567 | down |
| CELSR2 | 1.277885 | 0.972462 | 0.760993 | down |
| ACOX3 | 1.432677 | 1.09175 | 0.762035 | down |
| MTFR1L | 1.403434 | 1.06962 | 0.762145 | down |
| ITM2B | 1.278514 | 0.974467 | 0.762188 | down |
| RDH10 | 1.368147 | 1.043134 | 0.762443 | down |
| FBF1 | 1.190163 | 0.907527 | 0.762523 | down |
| GSTA4 | 0.983403 | 0.749987 | 0.762644 | down |
| HIGD2A | 1.287872 | 0.98229 | 0.762723 | down |

| | | | | |
|---|---|---|---|---|
| MAS1L | 1.339317 | 1.022318 | 0.763313 | down |
| GALNT14 | 1.30521 | 0.996833 | 0.763733 | down |
| PLCD3 | 1.342358 | 1.025272 | 0.763784 | down |
| LCOR | 1.236363 | 0.945013 | 0.764349 | down |
| PFN3 | 2.041987 | 1.561085 | 0.764493 | down |
| KRT8 | 1.491698 | 1.14044 | 0.764525 | down |
| ATP5MPL | 1.067289 | 0.816194 | 0.764736 | down |
| SLC35A2 | 1.19402 | 0.913128 | 0.764751 | down |
| TACSTD2 | 1.326442 | 1.014603 | 0.764906 | down |
| TNNI3K | 1.243471 | 0.951496 | 0.765194 | down |
| PLAU | 1.401515 | 1.07254 | 0.765272 | down |
| ACTR3B | 1.410532 | 1.079598 | 0.765384 | down |
| LSS | 1.316713 | 1.007806 | 0.765396 | down |
| PSEN1 | 1.276362 | 0.977312 | 0.765701 | down |
| PTDSS1 | 1.281396 | 0.981448 | 0.765921 | down |
| GBA | 1.382011 | 1.058689 | 0.766049 | down |
| WFS1 | 1.269956 | 0.973273 | 0.766383 | down |
| SCARB2 | 1.272205 | 0.975496 | 0.766776 | down |
| JMJD7 | 1.12069 | 0.859548 | 0.766981 | down |
| KDELR2 | 1.204403 | 0.924224 | 0.767371 | down |
| MCM9 | 1.492443 | 1.145603 | 0.767603 | down |
| S100A14 | 1.412408 | 1.084296 | 0.767693 | down |
| ATG101 | 1.087756 | 0.835463 | 0.768061 | down |
| CST3 | 1.253114 | 0.962984 | 0.768472 | down |
| SQOR | 1.285816 | 0.988169 | 0.768515 | down |
| CMAS | 1.303072 | 1.001527 | 0.768589 | down |
| MPZL2 | 1.217094 | 0.935529 | 0.768658 | down |
| ATP2B4 | 1.272687 | 0.978626 | 0.768945 | down |
| FGFR2 | 1.361032 | 1.046664 | 0.769022 | down |
| PINLYP | 1.478233 | 1.138965 | 0.770491 | down |
| TSPAN6 | 1.204558 | 0.928281 | 0.770641 | down |
| MRPL32 | 1.420191 | 1.094719 | 0.770825 | down |
| BPHL | 1.367308 | 1.054594 | 0.771292 | down |
| MLEC | 1.331823 | 1.027812 | 0.771733 | down |
| ITM2C | 1.310083 | 1.011123 | 0.771801 | down |
| CTSL | 1.215863 | 0.93843 | 0.771822 | down |
| SCPEP1 | 1.32466 | 1.022603 | 0.771974 | down |
| NCOA4 | 1.268176 | 0.97924 | 0.772164 | down |
| TMEM106I | 1.231252 | 0.951317 | 0.772642 | down |
| VKORC1L1 | 1.181388 | 0.913022 | 0.772838 | down |
| KRT5 | 1.635132 | 1.263813 | 0.772912 | down |
| AGR2 | 1.366829 | 1.056664 | 0.773077 | down |
| CPT1A | 1.222272 | 0.945442 | 0.773511 | down |
| DEGS1 | 1.326689 | 1.026453 | 0.773695 | down |
| CCDC88A | 1.665456 | 1.288561 | 0.773699 | down |
| C22orf23 | 1.382836 | 1.070524 | 0.774151 | down |
| ENDOG | 1.467187 | 1.137177 | 0.775073 | down |
| FAM178B | 1.330306 | 1.031396 | 0.775308 | down |
| EPN1 | 1.090113 | 0.845311 | 0.775434 | down |
| ATP6V0C | 1.285816 | 0.997186 | 0.775528 | down |
| IDH2 | 1.381553 | 1.071859 | 0.775836 | down |
| TMEM186 | 1.26498 | 0.981631 | 0.776005 | down |
| STX8 | 1.237182 | 0.961112 | 0.776856 | down |
| METTL11B | 1.3192 | 1.025127 | 0.777082 | down |
| TCHH | 1.51912 | 1.180508 | 0.7771 | down |
| PTTG1IP | 1.35775 | 1.055898 | 0.777682 | down |
| GINM1 | 1.2619 | 0.981546 | 0.777832 | down |
| CYP51A1 | 1.234348 | 0.960337 | 0.778011 | down |

| | | | | |
|---|---|---|---|---|
| KRT6A | 1.871433 | 1.457231 | 0.778671 | down |
| TMEM138 | 1.201909 | 0.935932 | 0.778704 | down |
| BANF1 | 1.227897 | 0.956433 | 0.77892 | down |
| MPC1 | 1.253124 | 0.976329 | 0.779116 | down |
| PRNP | 1.320534 | 1.028926 | 0.779174 | down |
| TFB1M | 1.292526 | 1.007374 | 0.779384 | down |
| TGOLN2 | 1.276082 | 0.994592 | 0.779411 | down |
| PI4K2A | 1.289674 | 1.005439 | 0.779607 | down |
| ITGAV | 1.32675 | 1.03436 | 0.77962 | down |
| TMEM258 | 1.142516 | 0.890975 | 0.779835 | down |
| DPAGT1 | 1.283273 | 1.00221 | 0.780979 | down |
| MCTP2 | 1.328804 | 1.03905 | 0.781944 | down |
| AURKAIP1 | 1.192909 | 0.933187 | 0.782278 | down |
| DNPEP | 1.27367 | 0.996471 | 0.782362 | down |
| TMEM199 | 1.048234 | 0.820404 | 0.782653 | down |
| HINT2 | 1.335393 | 1.04515 | 0.782654 | down |
| ABHD17A | 1.262745 | 0.988343 | 0.782694 | down |
| ZNF444 | 1.212238 | 0.948866 | 0.782739 | down |
| KAT14 | 1.28672 | 1.007424 | 0.782939 | down |
| COX7B | 1.165418 | 0.912497 | 0.782978 | down |
| MRPL51 | 1.169801 | 0.915994 | 0.783034 | down |
| WNT4 | 0.835744 | 0.654431 | 0.783052 | down |
| RRAS | 1.506182 | 1.179736 | 0.783263 | down |
| COX6C | 1.259552 | 0.986747 | 0.783411 | down |
| RETREG3 | 1.340907 | 1.050483 | 0.783413 | down |
| REEP6 | 1.143017 | 0.895703 | 0.783631 | down |
| FMC1 | 1.20261 | 0.942482 | 0.783697 | down |
| PNPLA6 | 1.283455 | 1.005857 | 0.78371 | down |
| SH3BP5 | 0.970169 | 0.760745 | 0.784136 | down |
| SMPDL3B | 1.247819 | 0.978837 | 0.784438 | down |
| LACTB | 1.366662 | 1.072067 | 0.784442 | down |
| SYVN1 | 1.339107 | 1.051267 | 0.785051 | down |
| TMPPE | 1.127961 | 0.885757 | 0.785273 | down |
| GNB2 | 1.364392 | 1.071634 | 0.78543 | down |
| SCARB1 | 1.122716 | 0.881857 | 0.785467 | down |
| RAB3D | 1.321786 | 1.038312 | 0.785538 | down |
| OR8B4 | 1.253614 | 0.985687 | 0.786276 | down |
| DDRGK1 | 1.314114 | 1.033763 | 0.786662 | down |
| CANT1 | 1.220526 | 0.96017 | 0.786685 | down |
| MICU1 | 1.247667 | 0.981621 | 0.786765 | down |
| TFRC | 1.376251 | 1.083015 | 0.786931 | down |
| TAP2 | 1.230135 | 0.968524 | 0.787331 | down |
| BCL2L1 | 1.218985 | 0.959786 | 0.787364 | down |
| PIGH | 1.446456 | 1.139094 | 0.787507 | down |
| EGFR | 1.295634 | 1.020526 | 0.787666 | down |
| PEX16 | 1.236126 | 0.974206 | 0.788112 | down |
| UQCR11 | 1.396182 | 1.100595 | 0.788289 | down |
| LRFN3 | 1.309478 | 1.032461 | 0.788452 | down |
| PBXIP1 | 1.322213 | 1.042807 | 0.788683 | down |
| TADA2A | 1.306239 | 1.030218 | 0.78869 | down |
| NDUFB1 | 1.31065 | 1.033715 | 0.788704 | down |
| CLIC3 | 1.383549 | 1.091543 | 0.788944 | down |
| OTOP2 | 0.59021 | 0.465809 | 0.789226 | down |
| RNF167 | 1.496483 | 1.181614 | 0.789594 | down |
| PRSS1 | 0.902178 | 0.712374 | 0.789616 | down |
| TM2D3 | 1.485827 | 1.173567 | 0.789841 | down |
| C4BPA | 1.082792 | 0.855285 | 0.789889 | down |
| CIB2 | 1.096121 | 0.866217 | 0.790257 | down |

| | | | | |
|---|---|---|---|---|
| CKMT1A | 1.264013 | 0.998963 | 0.79031 | down |
| GALNS | 1.266099 | 1.000632 | 0.790327 | down |
| ATP5MD | 1.414635 | 1.118184 | 0.79044 | down |
| NAAA | 1.22511 | 0.968472 | 0.790519 | down |
| TMCC3 | 1.288632 | 1.018872 | 0.790662 | down |
| HNRNPC | 1.369188 | 1.082569 | 0.790665 | down |
| RAB6A | 1.326793 | 1.049221 | 0.790795 | down |
| GGT6 | 1.289555 | 1.020923 | 0.791686 | down |
| SLC30A7 | 1.209361 | 0.957641 | 0.791857 | down |
| PLAGL2 | 0.842785 | 0.668011 | 0.792623 | down |
| MMAB | 1.290538 | 1.023133 | 0.792796 | down |
| CRP | 1.503114 | 1.191757 | 0.792859 | down |
| SOST | 1.428509 | 1.132941 | 0.793093 | down |
| ABHD16A | 1.327987 | 1.053429 | 0.793252 | down |
| TAX1BP1 | 1.255242 | 0.995762 | 0.793284 | down |
| GDPD3 | 1.259467 | 0.999299 | 0.79343 | down |
| C4A | 1.166317 | 0.92561 | 0.793618 | down |
| STBD1 | 1.131172 | 0.897937 | 0.793811 | down |
| ALDH5A1 | 1.301637 | 1.034108 | 0.794467 | down |
| IL6ST | 1.473973 | 1.17156 | 0.794831 | down |
| ATP6AP1 | 1.226558 | 0.975098 | 0.794987 | down |
| VWA5A | 2.05077 | 1.630353 | 0.794995 | down |
| PTPRB | 0.6876 | 0.546652 | 0.795015 | down |
| ZDHHC13 | 1.195459 | 0.950513 | 0.795103 | down |
| CD14 | 1.295093 | 1.029744 | 0.795112 | down |
| CYSTM1 | 0.853646 | 0.679 | 0.795412 | down |
| BHLHA15 | 1.244649 | 0.990033 | 0.795431 | down |
| SCAMP4 | 1.299733 | 1.034611 | 0.796018 | down |
| NDUFAB1 | 1.261326 | 1.004066 | 0.79604 | down |
| OAT | 1.2149 | 0.967654 | 0.796489 | down |
| MGAT1 | 1.261218 | 1.004667 | 0.796585 | down |
| NDFIP2 | 1.215983 | 0.968742 | 0.796674 | down |
| CHST3 | 1.115331 | 0.888911 | 0.796993 | down |
| CLU | 1.320736 | 1.053272 | 0.797489 | down |
| ATRN | 1.151207 | 0.91899 | 0.798284 | down |
| INPP5K | 1.170631 | 0.934729 | 0.798483 | down |
| P4HA2 | 1.375499 | 1.098519 | 0.798633 | down |
| TOR2A | 1.092334 | 0.872487 | 0.798737 | down |
| MSH4 | 0.901649 | 0.720181 | 0.798737 | down |
| NR1H2 | 0.915646 | 0.731805 | 0.799222 | down |
| SNX11 | 1.304675 | 1.04283 | 0.799302 | down |
| RFX7 | 1.337041 | 1.068769 | 0.799354 | down |
| GC | 1.235096 | 0.987406 | 0.799457 | down |
| MYORG | 1.35684 | 1.085113 | 0.799735 | down |
| CDH13 | 1.377407 | 1.101756 | 0.799877 | down |
| SEC11A | 1.261008 | 1.008783 | 0.799981 | down |
| TMEM30A | 1.183417 | 0.946839 | 0.800089 | down |
| CD40 | 1.367326 | 1.094231 | 0.800271 | down |
| HAUS3 | 1.45361 | 1.163348 | 0.800317 | down |
| ICOSLG | 1.316574 | 1.053825 | 0.80043 | down |
| RP2 | 1.27376 | 1.019601 | 0.800466 | down |
| COX11 | 1.155804 | 0.92522 | 0.800499 | down |
| ERBB2 | 1.258583 | 1.007501 | 0.800504 | down |
| SOAT1 | 1.193931 | 0.955778 | 0.80053 | down |
| PCYOX1L | 1.044478 | 0.836246 | 0.800636 | down |
| H6PD | 1.263678 | 1.011799 | 0.800678 | down |
| ZKSCAN8 | 1.219688 | 0.976791 | 0.800853 | down |
| LITAF | 1.23001 | 0.985082 | 0.800873 | down |

| | | | | |
|---|---|---|---|---|
| CCDC90B | 1.310193 | 1.049603 | 0.801106 | down |
| IDS | 1.119169 | 0.896675 | 0.801198 | down |
| TMEM8A | 1.330573 | 1.066131 | 0.801257 | down |
| SPARC | 1.291445 | 1.03482 | 0.801288 | down |
| GHITM | 1.237216 | 0.991539 | 0.801428 | down |
| CD9 | 1.273338 | 1.021646 | 0.802337 | down |
| NDEL1 | 1.195674 | 0.959365 | 0.802363 | down |
| TMEM109 | 1.290854 | 1.035761 | 0.802384 | down |
| CLCN7 | 1.190784 | 0.955727 | 0.802603 | down |
| PHYH | 1.263491 | 1.014281 | 0.802761 | down |
| ERLEC1 | 1.296037 | 1.040585 | 0.802898 | down |
| CLSPN | 1.095208 | 0.879357 | 0.802913 | down |
| FAM162A | 1.236458 | 0.992774 | 0.802918 | down |
| H2BFS | 1.130227 | 0.907629 | 0.803051 | down |
| BCKDHA | 1.242281 | 0.997661 | 0.803088 | down |
| CYB5R1 | 1.312647 | 1.054263 | 0.803158 | down |
| SYTL1 | 1.276733 | 1.025434 | 0.80317 | down |
| 44260 | 1.158326 | 0.930365 | 0.803198 | down |
| CHCHD4 | 1.165845 | 0.936546 | 0.80332 | down |
| PDP1 | 1.204713 | 0.968154 | 0.803639 | down |
| POLR2G | 1.165302 | 0.936667 | 0.803797 | down |
| HMOX1 | 1.230771 | 0.989304 | 0.803808 | down |
| SCCPDH | 1.334528 | 1.072998 | 0.804028 | down |
| DNAJC30 | 1.046128 | 0.84149 | 0.804385 | down |
| GPC1 | 1.356989 | 1.091572 | 0.804407 | down |
| SDHAF2 | 1.314392 | 1.057409 | 0.804485 | down |
| RABAC1 | 1.204625 | 0.969259 | 0.804614 | down |
| RBM3 | 1.199187 | 0.965056 | 0.804758 | down |
| NOS1 | 1.181821 | 0.951501 | 0.805115 | down |
| GNA15 | 1.289578 | 1.038393 | 0.80522 | down |
| OR5T2 | 1.485708 | 1.196769 | 0.805521 | down |
| FAM177A1 | 1.214338 | 0.978537 | 0.80582 | down |
| MASP1 | 1.303391 | 1.050635 | 0.806078 | down |
| TMEM120A | 1.207088 | 0.97303 | 0.806097 | down |
| ZRANB1 | 1.497657 | 1.207679 | 0.806379 | down |
| TEAD1 | 1.165042 | 0.939543 | 0.806445 | down |
| MAN1B1 | 1.346565 | 1.085969 | 0.806474 | down |
| STOML2 | 1.202041 | 0.969858 | 0.806843 | down |
| FAM57A | 1.515884 | 1.22317 | 0.806902 | down |
| HBB | 1.134703 | 0.915649 | 0.80695 | down |
| CNNM4 | 1.276062 | 1.029844 | 0.807048 | down |
| NIPSNAP1 | 1.30233 | 1.051074 | 0.807072 | down |
| PCK2 | 1.232693 | 0.994883 | 0.807081 | down |
| ANO10 | 1.183341 | 0.955612 | 0.807554 | down |
| AGPAT3 | 1.199028 | 0.969062 | 0.808207 | down |
| GNB1 | 1.265097 | 1.022487 | 0.808228 | down |
| ZNF12 | 1.243147 | 1.00493 | 0.808376 | down |
| TOR1AIP2 | 1.227583 | 0.992505 | 0.808504 | down |
| TIMM22 | 1.24222 | 1.004426 | 0.808573 | down |
| HCFC1 | 1.037958 | 0.839423 | 0.808726 | down |
| PLA2G12A | 1.071415 | 0.866517 | 0.808759 | down |
| APOBEC3B | 1.17682 | 0.951779 | 0.808772 | down |
| ABHD14A | 1.224103 | 0.990029 | 0.808779 | down |
| MRC1 | 1.341405 | 1.085099 | 0.808927 | down |
| SLC25A3 | 1.25584 | 1.015913 | 0.808951 | down |
| TMEM70 | 1.16588 | 0.943206 | 0.809007 | down |
| MMP14 | 1.164363 | 0.941987 | 0.809015 | down |
| GOLT1B | 1.296491 | 1.048911 | 0.809039 | down |

| | | | | |
|---|---|---|---|---|
| NEDD4L | 1.462251 | 1.183413 | 0.809309 | down |
| NR2C2AP | 1.210632 | 0.979782 | 0.809315 | down |
| ATRAID | 1.189451 | 0.962675 | 0.809344 | down |
| GPT2 | 1.102971 | 0.892851 | 0.809496 | down |
| EPHB4 | 1.201813 | 0.972893 | 0.809521 | down |
| CDH1 | 1.209962 | 0.979798 | 0.809776 | down |
| CES2 | 1.26838 | 1.027286 | 0.809919 | down |
| PLD3 | 1.176201 | 0.952878 | 0.810132 | down |
| ZNF480 | 1.382116 | 1.119789 | 0.810199 | down |
| CSGALNA( | 1.332078 | 1.079665 | 0.810512 | down |
| TTF1 | 1.150308 | 0.932423 | 0.810585 | down |
| SURF1 | 1.287409 | 1.043763 | 0.810747 | down |
| SLC25A4 | 1.254577 | 1.01776 | 0.811238 | down |
| LRRC8D | 1.264341 | 1.025709 | 0.81126 | down |
| RAB15 | 1.325011 | 1.074948 | 0.811275 | down |
| MGAT4B | 1.317793 | 1.069377 | 0.811491 | down |
| CEMIP2 | 1.14598 | 0.930027 | 0.811556 | down |
| AKAP17A | 1.214363 | 0.985996 | 0.811945 | down |
| GALNT2 | 1.231363 | 0.999918 | 0.812042 | down |
| FHAD1 | 1.458491 | 1.184934 | 0.812438 | down |
| TPP1 | 1.236981 | 1.005044 | 0.812498 | down |
| TOMM40L | 1.327692 | 1.078747 | 0.812498 | down |
| SLC35B2 | 1.199981 | 0.975041 | 0.812547 | down |
| ZNRD1 | 1.18424 | 0.962496 | 0.812754 | down |
| TLR3 | 1.360424 | 1.105848 | 0.81287 | down |
| RAP2B | 1.208032 | 0.98198 | 0.812876 | down |
| NUDT2 | 1.172084 | 0.952849 | 0.812953 | down |
| TMEM9B | 1.356697 | 1.10349 | 0.813366 | down |
| KCTD10 | 1.00441 | 0.817026 | 0.813438 | down |
| EI24 | 1.136527 | 0.924513 | 0.813454 | down |
| ITFG1 | 1.210652 | 0.984892 | 0.813522 | down |
| TST | 1.332379 | 1.084391 | 0.813876 | down |
| SLC26A6 | 1.263699 | 1.028498 | 0.81388 | down |
| CLDN1 | 1.028834 | 0.83769 | 0.814213 | down |
| THBD | 1.235635 | 1.006273 | 0.814377 | down |
| RRP15 | 1.218999 | 0.993177 | 0.814748 | down |
| SDF4 | 1.322779 | 1.078082 | 0.815013 | down |
| JAGN1 | 1.303807 | 1.06281 | 0.815159 | down |
| CCNYL1 | 1.210362 | 0.986755 | 0.815256 | down |
| SS18L1 | 1.407589 | 1.147904 | 0.815511 | down |
| MFN1 | 1.245721 | 1.016838 | 0.816265 | down |
| SLC27A4 | 1.263969 | 1.031984 | 0.816464 | down |
| EMB | 1.140541 | 0.931313 | 0.816554 | down |
| GRM4 | 1.070479 | 0.874251 | 0.816691 | down |
| TMEM106( | 1.455037 | 1.188334 | 0.816704 | down |
| OPA3 | 1.223403 | 0.999207 | 0.816744 | down |
| TEX264 | 1.07089 | 0.874645 | 0.816746 | down |
| SLC32A1 | 0.975721 | 0.797014 | 0.816846 | down |
| UBTF | 0.966919 | 0.789828 | 0.81685 | down |
| SURF4 | 1.130334 | 0.923475 | 0.816993 | down |
| MFGE8 | 1.228376 | 1.003852 | 0.817219 | down |
| GAR1 | 1.325901 | 1.083659 | 0.8173 | down |
| RTN4 | 1.199035 | 0.980017 | 0.817338 | down |
| STARD3NL | 1.153988 | 0.943361 | 0.817479 | down |
| SLC12A6 | 1.156665 | 0.945647 | 0.817563 | down |
| RAP1B | 1.24972 | 1.021836 | 0.817652 | down |
| MELTF | 1.347389 | 1.101706 | 0.81766 | down |
| DNAJC19 | 1.200076 | 0.981334 | 0.817726 | down |

| | | | | |
|---|---|---|---|---|
| HSD17B10 | 1.196694 | 0.978969 | 0.818062 | down |
| AGPAT5 | 1.259485 | 1.030884 | 0.818496 | down |
| SLC25A11 | 1.261264 | 1.032479 | 0.818607 | down |
| PDIA5 | 1.20579 | 0.987071 | 0.81861 | down |
| HNRNPH2 | 1.2512 | 1.024302 | 0.818656 | down |
| TMEM87A | 1.24423 | 1.01885 | 0.81886 | down |
| AP2S1 | 1.070962 | 0.877112 | 0.818995 | down |
| HNRNPA1 | 1.320595 | 1.081693 | 0.819095 | down |
| MED31 | 1.092868 | 0.895448 | 0.819357 | down |
| CDH23 | 1.434339 | 1.175299 | 0.819401 | down |
| STOM | 1.226988 | 1.00541 | 0.819413 | down |
| SIDT2 | 1.221174 | 1.000728 | 0.819481 | down |
| ENDOD1 | 1.257161 | 1.030301 | 0.819545 | down |
| CPT2 | 1.201397 | 0.984885 | 0.819784 | down |
| NDUFB7 | 1.357646 | 1.112996 | 0.819798 | down |
| ITIH2 | 1.171579 | 0.960846 | 0.820129 | down |
| CANX | 1.211737 | 0.994228 | 0.820498 | down |
| ING4 | 1.263255 | 1.036505 | 0.820503 | down |
| RGPD3 | 1.126449 | 0.924284 | 0.820529 | down |
| TMEM201 | 1.39512 | 1.145114 | 0.820799 | down |
| BCL2L13 | 1.218041 | 0.999965 | 0.820962 | down |
| PGRMC2 | 1.331578 | 1.093449 | 0.821168 | down |
| S100A10 | 1.34816 | 1.107151 | 0.821231 | down |
| FAHD1 | 1.275984 | 1.048479 | 0.821702 | down |
| RRAGC | 1.164341 | 0.956956 | 0.821887 | down |
| ARF6 | 1.20607 | 0.991311 | 0.821936 | down |
| SLC16A3 | 1.182898 | 0.972487 | 0.822122 | down |
| TSPAN1 | 1.186084 | 0.97583 | 0.822732 | down |
| CTSA | 1.215229 | 0.999851 | 0.822767 | down |
| ATP5PF | 1.154499 | 0.950121 | 0.822973 | down |
| ACSL5 | 1.294645 | 1.066028 | 0.823414 | down |
| HINT3 | 1.172501 | 0.965574 | 0.823517 | down |
| ZADH2 | 1.243281 | 1.024441 | 0.823982 | down |
| TMEM126/ | 1.180557 | 0.97285 | 0.82406 | down |
| ABCB8 | 1.266485 | 1.043755 | 0.824135 | down |
| MSRB2 | 1.23556 | 1.018365 | 0.824213 | down |
| HMGCL | 1.199536 | 0.98878 | 0.824301 | down |
| POFUT2 | 1.211678 | 0.998846 | 0.824349 | down |
| GRN | 1.231156 | 1.01496 | 0.824396 | down |
| USF1 | 1.097316 | 0.90473 | 0.824493 | down |
| LRP8 | 1.275488 | 1.051674 | 0.824527 | down |
| TAP1 | 1.207133 | 0.995369 | 0.824573 | down |
| SYNGR2 | 1.016497 | 0.838367 | 0.82476 | down |
| SEZ6L2 | 1.147428 | 0.946387 | 0.82479 | down |
| NSDHL | 1.24456 | 1.026517 | 0.824803 | down |
| SHISA4 | 0.956462 | 0.788901 | 0.824812 | down |
| TEAD4 | 1.073637 | 0.885549 | 0.824812 | down |
| AGTRAP | 1.286583 | 1.061262 | 0.824868 | down |
| SEC61A1 | 1.164815 | 0.960917 | 0.824953 | down |
| PWWP2B | 1.215614 | 1.003259 | 0.82531 | down |
| SLC1A3 | 1.355657 | 1.118931 | 0.825379 | down |
| HSD17B7 | 1.183879 | 0.97739 | 0.825583 | down |
| GTF2H4 | 1.117489 | 0.922633 | 0.82563 | down |
| PTGS2 | 1.295878 | 1.070191 | 0.825843 | down |
| ARL1 | 1.20022 | 0.991444 | 0.826052 | down |
| POP4 | 1.106987 | 0.914531 | 0.826145 | down |
| RAC2 | 1.183587 | 0.977917 | 0.826232 | down |
| EPHX1 | 1.169703 | 0.966487 | 0.826267 | down |

| | | | | |
|---|---|---|---|---|
| MYH8 | 1.231801 | 1.017929 | 0.826375 | down |
| TM9SF2 | 1.301597 | 1.075706 | 0.826451 | down |
| ACAD9 | 1.229323 | 1.015981 | 0.826456 | down |
| KRT19 | 1.278276 | 1.05708 | 0.826957 | down |
| SPRYD7 | 1.178331 | 0.974634 | 0.827131 | down |
| SNRPF | 1.10062 | 0.910658 | 0.827405 | down |
| PREX2 | 1.638685 | 1.355902 | 0.827433 | down |
| PCYOX1 | 1.250897 | 1.035049 | 0.827445 | down |
| TMEM214 | 1.192855 | 0.98738 | 0.827745 | down |
| CRELD1 | 1.236323 | 1.023387 | 0.827767 | down |
| SARAF | 1.187305 | 0.982813 | 0.827767 | down |
| ECI1 | 1.31655 | 1.089996 | 0.827919 | down |
| SLC44A1 | 1.305015 | 1.080514 | 0.82797 | down |
| PIGT | 1.262797 | 1.045686 | 0.828071 | down |
| HABP2 | 1.141356 | 0.945371 | 0.828288 | down |
| KCNIP3 | 1.19524 | 0.990139 | 0.828402 | down |
| KRT10 | 1.25688 | 1.041217 | 0.828414 | down |
| KRT6B | 1.255024 | 1.03975 | 0.828471 | down |
| ERLIN2 | 1.171453 | 0.970678 | 0.82861 | down |
| MAN2A1 | 1.197544 | 0.992537 | 0.828811 | down |
| AK4 | 1.296241 | 1.074396 | 0.828855 | down |
| TMED4 | 1.161265 | 0.962529 | 0.828862 | down |
| MTG1 | 1.289133 | 1.068833 | 0.829109 | down |
| CDK18 | 1.212822 | 1.005798 | 0.829304 | down |
| MAP1LC3E | 1.142356 | 0.947365 | 0.829308 | down |
| CYP2S1 | 1.14292 | 0.947925 | 0.829389 | down |
| RMND1 | 1.236709 | 1.025984 | 0.829608 | down |
| STEAP3 | 1.310556 | 1.087353 | 0.829688 | down |
| ITPRIPL2 | 1.146217 | 0.951053 | 0.829732 | down |
| POGLUT1 | 1.151177 | 0.95547 | 0.829994 | down |
| MICA | 1.149721 | 0.954404 | 0.830118 | down |
| HSD17B12 | 1.220609 | 1.013334 | 0.830187 | down |
| PRPF4B | 1.244393 | 1.033121 | 0.830221 | down |
| NIPSNAP2 | 1.212743 | 1.007038 | 0.83038 | down |
| SELENOM | 1.263724 | 1.049771 | 0.830696 | down |
| UQCC2 | 1.398623 | 1.162035 | 0.830843 | down |
| ATP6V1F | 1.104016 | 0.917451 | 0.831013 | down |
| DHRS3 | 1.280624 | 1.064235 | 0.831028 | down |
| MYO1D | 1.254393 | 1.042526 | 0.8311 | down |
| SERAC1 | 0.806794 | 0.670681 | 0.831291 | down |
| PPP1R26 | 1.083627 | 0.900865 | 0.831342 | down |
| ALG2 | 1.156352 | 0.96147 | 0.831468 | down |
| SYNJ2 | 1.073659 | 0.892776 | 0.831526 | down |
| RAB32 | 1.170581 | 0.973371 | 0.831529 | down |
| ATP5MF | 1.124567 | 0.935167 | 0.83158 | down |
| CDCA8 | 1.116398 | 0.92847 | 0.831665 | down |
| TYRP1 | 1.551168 | 1.290624 | 0.832034 | down |
| ALCAM | 1.191283 | 0.9912 | 0.832044 | down |
| LEMD2 | 1.268945 | 1.055841 | 0.832062 | down |
| DNAJC16 | 1.256357 | 1.045871 | 0.832463 | down |
| REEP5 | 1.260811 | 1.049593 | 0.832474 | down |
| FAM241A | 1.046789 | 0.871522 | 0.832567 | down |
| GNA13 | 1.212395 | 1.009705 | 0.832818 | down |
| DDX27 | 1.22512 | 1.020346 | 0.832854 | down |
| DHRS1 | 1.233376 | 1.027406 | 0.833003 | down |
| TOR3A | 1.204149 | 1.003254 | 0.833164 | down |

Table S6. related to Figure 5

| Gene Syml | shControl | shRNPS1 a | shRNPS1/shControl | |
|---|---|---|---|---|
| ZNF282 | 4.88398 | 1.132858 | 0.231954 | down |
| CNTN5 | 6.21535 | 1.716336 | 0.276145 | down |
| KNDC1 | 2.963975 | 0.982697 | 0.331547 | down |
| CX3CR1 | 3.35626 | 1.188787 | 0.3542 | down |
| APOLD1 | 2.493969 | 0.925015 | 0.370901 | down |
| CABP2 | 3.796707 | 1.455187 | 0.383276 | down |
| SPHKAP | 3.045546 | 1.1691 | 0.383872 | down |
| SHF | 4.425749 | 1.715974 | 0.387725 | down |
| CIB4 | 4.344682 | 1.740205 | 0.400537 | down |
| ZNF469 | 3.346232 | 1.360437 | 0.406558 | down |
| PLG | 2.088781 | 0.863822 | 0.413553 | down |
| NEB | 2.114149 | 0.910682 | 0.430756 | down |
| MTTP | 3.639659 | 1.574932 | 0.432714 | down |
| ZNF750 | 2.499557 | 1.102568 | 0.441105 | down |
| SDC4 | 2.139825 | 1.020201 | 0.476769 | down |
| FBP2 | 2.20345 | 1.056652 | 0.479544 | down |
| SEMA6D | 1.781326 | 0.863012 | 0.484477 | down |
| KRT4 | 2.528086 | 1.246612 | 0.493105 | down |
| CFAP300 | 2.436094 | 1.21277 | 0.497834 | down |
| DNAH11 | 1.668502 | 0.832965 | 0.499229 | down |
| GPR162 | 2.208801 | 1.116676 | 0.505557 | down |
| NOS2 | 2.503813 | 1.296972 | 0.517999 | down |
| ERVMER34 | 1.504224 | 0.785949 | 0.522495 | down |
| DENND1A | 1.698215 | 0.890078 | 0.524126 | down |
| LIMK1 | 1.983731 | 1.042424 | 0.525487 | down |
| MFAP3 | 1.630508 | 0.857419 | 0.52586 | down |
| SRPX2 | 1.891184 | 1.001543 | 0.529585 | down |
| SPZ1 | 3.548902 | 1.899667 | 0.535283 | down |
| LAPTM4B | 2.159351 | 1.158948 | 0.536711 | down |
| STK36 | 2.395521 | 1.302889 | 0.543885 | down |
| HPX | 1.801603 | 1.000395 | 0.555281 | down |
| SPPL2B | 1.585254 | 0.88869 | 0.560598 | down |
| ART4 | 2.107053 | 1.18292 | 0.56141 | down |
| FUT3 | 1.733304 | 0.976792 | 0.563544 | down |
| HYI | 1.865319 | 1.058618 | 0.567527 | down |
| TTC23L | 1.787451 | 1.024125 | 0.572952 | down |
| MANEAL | 2.470626 | 1.421815 | 0.575488 | down |
| TMEM19 | 1.762263 | 1.016916 | 0.577051 | down |
| VWA5A | 2.05077 | 1.197361 | 0.583859 | down |
| MAPK8IP1 | 1.81266 | 1.060532 | 0.58507 | down |
| CLN3 | 1.602291 | 0.944085 | 0.589209 | down |
| GRIN2D | 1.899048 | 1.119889 | 0.589711 | down |
| TNFRSF12/ | 1.48769 | 0.880683 | 0.591981 | down |
| FAM102B | 1.783326 | 1.060505 | 0.594678 | down |
| TMEM147 | 1.658798 | 0.987851 | 0.595522 | down |
| PCDHB7 | 1.969438 | 1.179431 | 0.598867 | down |
| LRMDA | 1.386466 | 0.837342 | 0.603939 | down |
| PLEC | 1.606593 | 0.978273 | 0.608912 | down |
| PRXL2A | 2.082753 | 1.270715 | 0.610113 | down |
| ETV4 | 1.513889 | 0.927588 | 0.612719 | down |
| HMGCR | 1.84293 | 1.129315 | 0.612782 | down |
| KRT16 | 1.78061 | 1.094798 | 0.614844 | down |
| L1CAM | 1.745017 | 1.078325 | 0.617945 | down |
| FDFT1 | 1.584216 | 0.984228 | 0.621272 | down |
| FAM57A | 1.515884 | 0.944169 | 0.62285 | down |
| MCM9 | 1.492443 | 0.930862 | 0.623717 | down |

| | | | | |
|---|---|---|---|---|
| HM13 | 1.394813 | 0.870717 | 0.624253 | down |
| TPTE2 | 1.532367 | 0.959568 | 0.6262 | down |
| ITIH6 | 1.707737 | 1.07161 | 0.627503 | down |
| KRT13 | 2.016554 | 1.266487 | 0.628045 | down |
| CLSTN3 | 1.847771 | 1.162424 | 0.629095 | down |
| F10 | 1.729791 | 1.099053 | 0.635367 | down |
| HIST1H3A | 1.319292 | 0.839 | 0.635948 | down |
| SPRR3 | 1.677631 | 1.069733 | 0.637645 | down |
| TMEM97 | 1.632101 | 1.042827 | 0.638948 | down |
| NOLC1 | 2.231812 | 1.429786 | 0.640639 | down |
| TTC34 | 1.846226 | 1.187661 | 0.643292 | down |
| BORCS8 | 1.613252 | 1.040105 | 0.644726 | down |
| FGFBP1 | 1.426456 | 0.922095 | 0.646424 | down |
| H3F3A | 1.423909 | 0.920857 | 0.646711 | down |
| COL11A1 | 2.018794 | 1.305921 | 0.646882 | down |
| TSPO | 1.491048 | 0.96545 | 0.647497 | down |
| FCRL5 | 1.495462 | 0.968899 | 0.647892 | down |
| KRT23 | 1.498222 | 0.971462 | 0.64841 | down |
| INS | 1.903939 | 1.235313 | 0.64882 | down |
| ARL8B | 1.487098 | 0.965109 | 0.648989 | down |
| ARL6IP4 | 1.564623 | 1.01667 | 0.649786 | down |
| ETFRF1 | 1.371673 | 0.893042 | 0.651061 | down |
| ZBED8 | 1.640617 | 1.068286 | 0.651149 | down |
| PYURF | 1.527474 | 0.999385 | 0.654273 | down |
| FBXO41 | 1.857088 | 1.218792 | 0.656292 | down |
| B2M | 1.475936 | 0.970499 | 0.657548 | down |
| MSMO1 | 1.361826 | 0.8961 | 0.658014 | down |
| OLR1 | 1.645124 | 1.085557 | 0.659864 | down |
| ABHD2 | 1.740889 | 1.150322 | 0.660767 | down |
| TERB2 | 1.460913 | 0.966345 | 0.661467 | down |
| ISOC2 | 1.508829 | 0.998247 | 0.661604 | down |
| RP1 | 1.512054 | 1.00244 | 0.662966 | down |
| PRSS21 | 2.085428 | 1.383472 | 0.6634 | down |
| TMEM30A | 1.183417 | 0.785775 | 0.663989 | down |
| NID1 | 1.636056 | 1.08716 | 0.664501 | down |
| NDUFB2 | 1.808065 | 1.202097 | 0.664853 | down |
| HLA-B | 1.449723 | 0.965223 | 0.665798 | down |
| MPDU1 | 1.431166 | 0.954458 | 0.66691 | down |
| LAMTOR3 | 1.620503 | 1.081984 | 0.667684 | down |
| LCN2 | 1.601045 | 1.075788 | 0.671929 | down |
| CAPS2 | 1.510107 | 1.01671 | 0.67327 | down |
| TMEM106( | 1.455037 | 0.979839 | 0.673412 | down |
| SC5D | 1.358717 | 0.917838 | 0.675518 | down |
| ARL8A | 1.597061 | 1.081208 | 0.676998 | down |
| RAB10 | 1.609323 | 1.090558 | 0.67765 | down |
| SQLE | 1.480438 | 1.004668 | 0.678629 | down |
| KRT14 | 1.529099 | 1.041412 | 0.681063 | down |
| ACVR1 | 1.592964 | 1.086844 | 0.682278 | down |
| CLCN6 | 1.75135 | 1.195661 | 0.682708 | down |
| SERINC3 | 1.398802 | 0.955717 | 0.683239 | down |
| ZNF30 | 1.426455 | 0.975783 | 0.684061 | down |
| PCNX1 | 1.547901 | 1.060267 | 0.684971 | down |
| AURKB | 1.675943 | 1.150451 | 0.68645 | down |
| SNX7 | 1.462381 | 1.005762 | 0.687756 | down |
| RNF168 | 1.462457 | 1.006274 | 0.688071 | down |
| HERPUD2 | 1.475771 | 1.015688 | 0.688242 | down |
| SCD | 1.365224 | 0.943579 | 0.691154 | down |
| OSTC | 1.302053 | 0.900066 | 0.691266 | down |

| | | | | |
|---|---|---|---|---|
| BEND4 | 1.885441 | 1.304257 | 0.691752 | down |
| TRPM4 | 1.660408 | 1.15024 | 0.692745 | down |
| KRT17 | 1.810094 | 1.254332 | 0.692965 | down |
| SIRT1 | 1.646062 | 1.141546 | 0.693502 | down |
| SCD5 | 1.597024 | 1.107814 | 0.693674 | down |
| MRPL54 | 1.349362 | 0.936473 | 0.694012 | down |
| RNF167 | 1.496483 | 1.040828 | 0.695516 | down |
| PLEKHA7 | 1.495196 | 1.045162 | 0.699013 | down |
| TSPAN31 | 1.399601 | 0.979183 | 0.699616 | down |
| PLGRKT | 1.549742 | 1.085109 | 0.700187 | down |
| SOST | 1.428509 | 1.00079 | 0.700584 | down |
| SOD2 | 1.358794 | 0.954425 | 0.702406 | down |
| TMX4 | 1.21167 | 0.851148 | 0.702458 | down |
| TBRG1 | 1.243343 | 0.873719 | 0.702718 | down |
| ICAM1 | 1.295737 | 0.910984 | 0.703063 | down |
| TMBIM1 | 1.639688 | 1.153461 | 0.703464 | down |
| RNF13 | 1.44844 | 1.019325 | 0.70374 | down |
| APOC3 | 1.32461 | 0.932649 | 0.704093 | down |
| HLA-E | 1.415372 | 0.997363 | 0.704665 | down |
| CLDN7 | 1.313999 | 0.926055 | 0.704761 | down |
| TPBG | 1.43003 | 1.009521 | 0.705944 | down |
| TMEM59 | 1.375178 | 0.970865 | 0.705992 | down |
| FAM3A | 1.44029 | 1.018665 | 0.707264 | down |
| SOX6 | 1.460098 | 1.033782 | 0.708022 | down |
| TMEM40 | 1.2344 | 0.874668 | 0.708577 | down |
| DHCR7 | 1.432278 | 1.015478 | 0.708995 | down |
| LGALS3BP | 1.421921 | 1.008898 | 0.709532 | down |
| LRRTM2 | 1.395444 | 0.99026 | 0.709638 | down |
| APOE | 1.352521 | 0.960475 | 0.710137 | down |
| C3 | 1.368629 | 0.972597 | 0.710636 | down |
| RAP1A | 1.408478 | 1.002002 | 0.711407 | down |
| ZNF174 | 1.286413 | 0.915902 | 0.711982 | down |
| SPINT2 | 1.45438 | 1.039153 | 0.714499 | down |
| ABHD13 | 1.729631 | 1.235902 | 0.714547 | down |
| LOXL2 | 1.45515 | 1.041696 | 0.715869 | down |
| RAP2A | 1.298298 | 0.929445 | 0.715895 | down |
| ADGRG1 | 1.446019 | 1.037683 | 0.717614 | down |
| CCDC88A | 1.665456 | 1.195433 | 0.717781 | down |
| TK2 | 1.537591 | 1.104731 | 0.718481 | down |
| FAM89B | 1.571409 | 1.129456 | 0.718754 | down |
| APP | 1.355308 | 0.974371 | 0.71893 | down |
| KRT15 | 1.50907 | 1.08625 | 0.719814 | down |
| PTTG1IP | 1.35775 | 0.9797 | 0.721561 | down |
| HLA-C | 1.35337 | 0.976665 | 0.721654 | down |
| BTN2A1 | 1.72909 | 1.247915 | 0.721717 | down |
| KAT14 | 1.28672 | 0.928754 | 0.7218 | down |
| CCDC183 | 1.708283 | 1.23317 | 0.721877 | down |
| SLC25A1 | 1.488707 | 1.07517 | 0.722218 | down |
| PDGFC | 1.393037 | 1.006709 | 0.722672 | down |
| OR5T2 | 1.485708 | 1.075314 | 0.723772 | down |
| ATP5MD | 1.414635 | 1.02389 | 0.723784 | down |
| GPR87 | 1.366448 | 0.989085 | 0.723836 | down |
| ABAT | 1.44478 | 1.046515 | 0.724342 | down |
| CD58 | 1.513695 | 1.098472 | 0.725689 | down |
| NDFIP1 | 1.488068 | 1.08 | 0.725773 | down |
| MOS | 1.292261 | 0.938697 | 0.726399 | down |
| CHAD | 1.201033 | 0.873161 | 0.727008 | down |
| ORMDL2 | 1.482381 | 1.079177 | 0.728002 | down |

| | | | | |
|---|---|---|---|---|
| CLCN3 | 1.518827 | 1.105795 | 0.728058 | down |
| BCKDHB | 1.365472 | 0.995787 | 0.729262 | down |
| PRNP | 1.320534 | 0.963266 | 0.729452 | down |
| FAM178B | 1.330306 | 0.970408 | 0.729463 | down |
| ZRANB1 | 1.497657 | 1.093629 | 0.730227 | down |
| TSPAN6 | 1.204558 | 0.880463 | 0.730943 | down |
| KRT6A | 1.871433 | 1.368615 | 0.73132 | down |
| JPH2 | 1.53547 | 1.123512 | 0.731706 | down |
| TMEM201 | 1.39512 | 1.021405 | 0.732127 | down |
| ATP5MF | 1.124567 | 0.823344 | 0.732144 | down |
| TRIM29 | 1.479648 | 1.084113 | 0.732683 | down |
| S100A14 | 1.412408 | 1.035303 | 0.733006 | down |
| RBP3 | 1.470356 | 1.078215 | 0.733302 | down |
| SDHAF2 | 1.314392 | 0.964749 | 0.733989 | down |
| TADA2A | 1.306239 | 0.958811 | 0.734024 | down |
| CD63 | 1.224143 | 0.898614 | 0.734076 | down |
| KRT1 | 1.781503 | 1.307779 | 0.734088 | down |
| SLC25A20 | 1.299282 | 0.954843 | 0.7349 | down |
| LAMP2 | 1.272548 | 0.93549 | 0.735132 | down |
| ITPRIPL1 | 1.501941 | 1.104551 | 0.735416 | down |
| BPHL | 1.367308 | 1.006339 | 0.736 | down |
| SLC35F6 | 1.212065 | 0.892371 | 0.73624 | down |
| HSPG2 | 1.420944 | 1.047128 | 0.736924 | down |
| RRAS | 1.506182 | 1.11097 | 0.737607 | down |
| PIGH | 1.446456 | 1.067051 | 0.7377 | down |
| AP3M2 | 1.157152 | 0.853943 | 0.73797 | down |
| ALDH1L2 | 1.352489 | 0.998507 | 0.738274 | down |
| PBXIP1 | 1.322213 | 0.976483 | 0.738521 | down |
| AURKAIP1 | 1.192909 | 0.88283 | 0.740064 | down |
| CD40 | 1.367326 | 1.013002 | 0.740864 | down |
| HSDL1 | 1.402398 | 1.040194 | 0.741726 | down |
| RP1L1 | 1.307461 | 0.969865 | 0.741793 | down |
| PRRG1 | 1.420285 | 1.054174 | 0.742227 | down |
| LDAH | 1.361625 | 1.011124 | 0.742586 | down |
| SLC1A4 | 1.129657 | 0.839539 | 0.74318 | down |
| CD9 | 1.273338 | 0.948086 | 0.744568 | down |
| RBM3 | 1.199187 | 0.893261 | 0.744889 | down |
| TMEM35B | 1.456385 | 1.087347 | 0.746607 | down |
| ACOT4 | 1.393891 | 1.041475 | 0.747171 | down |
| ACTR3B | 1.410532 | 1.054222 | 0.747394 | down |
| EPC1 | 1.038732 | 0.776468 | 0.747515 | down |
| PLAU | 1.401515 | 1.049143 | 0.748578 | down |
| NDUFA3 | 1.445563 | 1.082339 | 0.748732 | down |
| TMEM222 | 1.395507 | 1.045171 | 0.748954 | down |
| SLC2A1 | 1.378769 | 1.03291 | 0.749154 | down |
| DHRS13 | 1.326918 | 0.994416 | 0.749418 | down |
| FAAP24 | 1.585654 | 1.188412 | 0.749478 | down |
| MMP1 | 1.392683 | 1.045988 | 0.751059 | down |
| SLC16A1 | 1.21929 | 0.916074 | 0.751318 | down |
| PSAP | 1.346283 | 1.011751 | 0.751514 | down |
| KRT5 | 1.635132 | 1.229598 | 0.751987 | down |
| TGOLN2 | 1.276082 | 0.959811 | 0.752155 | down |
| DHCR24 | 1.341213 | 1.009292 | 0.752522 | down |
| TMEM109 | 1.290854 | 0.971532 | 0.752628 | down |
| C4BPA | 1.082792 | 0.814961 | 0.752647 | down |
| CBARP | 1.123178 | 0.846067 | 0.75328 | down |
| SDCBP | 1.224521 | 0.923015 | 0.753776 | down |
| GPC1 | 1.356989 | 1.022965 | 0.753849 | down |

| | | | | |
|---|---|---|---|---|
| TLR3 | 1.360424 | 1.02615 | 0.754286 | down |
| SYVN1 | 1.339107 | 1.01071 | 0.754765 | down |
| PI4K2A | 1.289674 | 0.973774 | 0.755054 | down |
| TRIM22 | 1.468839 | 1.109359 | 0.755262 | down |
| F8 | 1.595564 | 1.205892 | 0.755778 | down |
| ASPG | 1.159468 | 0.876873 | 0.756272 | down |
| REEP2 | 1.438393 | 1.087821 | 0.756275 | down |
| MANSC1 | 1.263806 | 0.957207 | 0.7574 | down |
| SLC25A4 | 1.254577 | 0.951453 | 0.758386 | down |
| MME | 1.334346 | 1.012759 | 0.758993 | down |
| NOS1 | 1.181821 | 0.898197 | 0.760011 | down |
| SLC7A6 | 1.263782 | 0.960691 | 0.760171 | down |
| DPAGT1 | 1.283273 | 0.97604 | 0.760586 | down |
| GSTA4 | 0.983403 | 0.74872 | 0.761357 | down |
| JAGN1 | 1.303807 | 0.992793 | 0.761457 | down |
| UQCC2 | 1.398623 | 1.065077 | 0.761519 | down |
| PREX2 | 1.638685 | 1.248937 | 0.762158 | down |
| GHITM | 1.237216 | 0.943785 | 0.762829 | down |
| LAMB2 | 1.391394 | 1.061609 | 0.762982 | down |
| JUN | 1.330773 | 1.015822 | 0.763332 | down |
| PGS1 | 1.402972 | 1.07117 | 0.763501 | down |
| ITM2B | 1.278514 | 0.976303 | 0.763624 | down |
| CRP | 1.503114 | 1.148004 | 0.76375 | down |
| LACTB | 1.366662 | 1.044503 | 0.764273 | down |
| TMEM115 | 1.127811 | 0.862056 | 0.764362 | down |
| PSEN1 | 1.276362 | 0.975825 | 0.764536 | down |
| CDH13 | 1.377407 | 1.05332 | 0.764713 | down |
| SQOR | 1.285816 | 0.984177 | 0.76541 | down |
| SELENOI | 1.347702 | 1.031865 | 0.765647 | down |
| METTL11B | 1.3192 | 1.010318 | 0.765856 | down |
| MED21 | 1.393552 | 1.067508 | 0.766034 | down |
| SCPEP1 | 1.32466 | 1.015416 | 0.766548 | down |
| TPRG1L | 1.415043 | 1.084786 | 0.76661 | down |
| SRD5A3 | 1.371604 | 1.052509 | 0.767357 | down |
| IL10RB | 1.33468 | 1.024369 | 0.767501 | down |
| CTHRC1 | 1.691845 | 1.298966 | 0.767781 | down |
| PMFBP1 | 1.412918 | 1.084904 | 0.767846 | down |
| CTSB | 1.33052 | 1.021962 | 0.768092 | down |
| TACSTD2 | 1.326442 | 1.018986 | 0.76821 | down |
| RAB5B | 1.401106 | 1.076909 | 0.768613 | down |
| MTCH2 | 1.279115 | 0.983837 | 0.769154 | down |
| MGST3 | 1.37794 | 1.060053 | 0.769302 | down |
| SNX11 | 1.304675 | 1.004303 | 0.769773 | down |
| HAUS3 | 1.45361 | 1.118962 | 0.769781 | down |
| MYL4 | 1.339589 | 1.031463 | 0.769985 | down |
| FGFR2 | 1.361032 | 1.048352 | 0.770263 | down |
| OGDHL | 1.393787 | 1.073666 | 0.770323 | down |
| S100P | 1.960827 | 1.510772 | 0.770477 | down |
| ZNF480 | 1.382116 | 1.064951 | 0.770522 | down |
| MASP1 | 1.303391 | 1.004645 | 0.770793 | down |
| MTFR1L | 1.403434 | 1.08189 | 0.770888 | down |
| STC1 | 1.257661 | 0.969785 | 0.771102 | down |
| MFGE8 | 1.228376 | 0.948355 | 0.77204 | down |
| GRPR | 1.084583 | 0.837867 | 0.772524 | down |
| PPM1J | 1.048614 | 0.810554 | 0.772976 | down |
| ASAH1 | 1.300481 | 1.006023 | 0.773577 | down |
| KDELR2 | 1.204403 | 0.931951 | 0.773787 | down |
| BCO2 | 1.714159 | 1.326601 | 0.773908 | down |

| | | | | |
|---|---|---|---|---|
| APOA1 | 1.161706 | 0.899145 | 0.773986 | down |
| ZNF771 | 1.246151 | 0.965096 | 0.774462 | down |
| FBLN1 | 1.267888 | 0.982086 | 0.774584 | down |
| EPB41L1 | 1.559086 | 1.20796 | 0.774787 | down |
| SMIM13 | 1.469633 | 1.139037 | 0.775049 | down |
| RAB4B | 1.281485 | 0.993541 | 0.775305 | down |
| GUK1 | 1.323848 | 1.026939 | 0.775723 | down |
| CYB5R1 | 1.312647 | 1.018645 | 0.776023 | down |
| SLC15A4 | 1.304752 | 1.013149 | 0.776507 | down |
| TMEM120/ | 1.207088 | 0.937625 | 0.776766 | down |
| PKP1 | 1.217714 | 0.946027 | 0.776887 | down |
| TM9SF2 | 1.301597 | 1.011417 | 0.777058 | down |
| MAN1B1 | 1.346565 | 1.046448 | 0.777124 | down |
| PGRMC2 | 1.331578 | 1.035505 | 0.777652 | down |
| FAM241B | 1.31065 | 1.019443 | 0.777814 | down |
| ALDH5A1 | 1.301637 | 1.012544 | 0.7779 | down |
| LZTS3 | 1.233393 | 0.959891 | 0.778252 | down |
| FMC1 | 1.20261 | 0.936346 | 0.778595 | down |
| CD14 | 1.295093 | 1.008673 | 0.778842 | down |
| PTGS2 | 1.295878 | 1.009801 | 0.779241 | down |
| SYNPO | 1.200939 | 0.935861 | 0.779274 | down |
| RAB6A | 1.326793 | 1.034091 | 0.779392 | down |
| MPZL2 | 1.217094 | 0.949076 | 0.779789 | down |
| RAB3D | 1.321786 | 1.031236 | 0.780184 | down |
| STX11 | 1.448906 | 1.130443 | 0.780205 | down |
| KDM1B | 1.358845 | 1.060218 | 0.780235 | down |
| AGTRAP | 1.286583 | 1.003853 | 0.780247 | down |
| ACSL1 | 1.323423 | 1.032636 | 0.780277 | down |
| THBS2 | 1.501865 | 1.172744 | 0.780858 | down |
| CHCHD4 | 1.165845 | 0.910443 | 0.78093 | down |
| SMPDL3B | 1.247819 | 0.974482 | 0.780948 | down |
| PLXNB3 | 1.546252 | 1.207581 | 0.780973 | down |
| SCARB2 | 1.272205 | 0.993877 | 0.781224 | down |
| CEP250 | 1.169454 | 0.913713 | 0.781316 | down |
| CD151 | 1.256648 | 0.981915 | 0.781376 | down |
| EGFR | 1.295634 | 1.012958 | 0.781824 | down |
| LITAF | 1.23001 | 0.962785 | 0.782746 | down |
| KRT10 | 1.25688 | 0.984061 | 0.782939 | down |
| TFRC | 1.376251 | 1.078104 | 0.783363 | down |
| RETREG2 | 1.433473 | 1.123666 | 0.783877 | down |
| LSS | 1.316713 | 1.032325 | 0.784017 | down |
| TUBGCP6 | 1.197161 | 0.938667 | 0.784077 | down |
| SSR2 | 1.231094 | 0.965476 | 0.784243 | down |
| ACOX3 | 1.432677 | 1.123625 | 0.784283 | down |
| COX6B2 | 1.089789 | 0.854833 | 0.784402 | down |
| RCN1 | 1.189423 | 0.934204 | 0.785426 | down |
| TMEM9B | 1.356697 | 1.065685 | 0.7855 | down |
| CAV2 | 1.188732 | 0.93376 | 0.785509 | down |
| CYB5B | 1.215907 | 0.955107 | 0.78551 | down |
| RGPD3 | 1.126449 | 0.884857 | 0.785528 | down |
| SLC16A3 | 1.182898 | 0.929512 | 0.785792 | down |
| FMR1 | 1.280313 | 1.006067 | 0.785798 | down |
| LY75 | 1.09769 | 0.862629 | 0.785859 | down |
| ENDOG | 1.467187 | 1.153053 | 0.785894 | down |
| GDPD3 | 1.259467 | 0.989917 | 0.785981 | down |
| SHB | 1.73747 | 1.365696 | 0.786026 | down |
| CYSTM1 | 0.853646 | 0.671072 | 0.786125 | down |
| TRIM13 | 1.256827 | 0.988382 | 0.78641 | down |

| | | | | |
|---|---|---|---|---|
| MRPL33 | 1.347555 | 1.060086 | 0.786673 | down |
| JTB | 1.135325 | 0.89322 | 0.786752 | down |
| CERS2 | 1.221824 | 0.961417 | 0.78687 | down |
| ABHD16A | 1.327987 | 1.045013 | 0.786915 | down |
| RAP1B | 1.24972 | 0.983457 | 0.786942 | down |
| ARL6IP5 | 1.328911 | 1.046053 | 0.78715 | down |
| IDH2 | 1.381553 | 1.087512 | 0.787166 | down |
| SCCPDH | 1.334528 | 1.050757 | 0.787362 | down |
| MAS1L | 1.339317 | 1.054644 | 0.787449 | down |
| CDK18 | 1.212822 | 0.955685 | 0.787985 | down |
| CCDC90B | 1.310193 | 1.032691 | 0.788198 | down |
| MMAB | 1.290538 | 1.017221 | 0.788214 | down |
| GNB2 | 1.364392 | 1.07557 | 0.788315 | down |
| CCNL1 | 1.414252 | 1.114898 | 0.788331 | down |
| SOAT1 | 1.193931 | 0.941397 | 0.788486 | down |
| KRT8 | 1.491698 | 1.176464 | 0.788674 | down |
| ITGAV | 1.32675 | 1.046542 | 0.788802 | down |
| CYP51A1 | 1.234348 | 0.973671 | 0.788814 | down |
| IGFBP6 | 1.394156 | 1.100004 | 0.789011 | down |
| SS18L1 | 1.407589 | 1.111071 | 0.789343 | down |
| ZNF205 | 1.301418 | 1.027348 | 0.789406 | down |
| ZNF841 | 1.591838 | 1.256658 | 0.789438 | down |
| RAB32 | 1.170581 | 0.924146 | 0.789477 | down |
| TNNI3K | 1.243471 | 0.982025 | 0.789745 | down |
| HNRNPC | 1.369188 | 1.081639 | 0.789986 | down |
| INPP5D | 1.058999 | 0.83672 | 0.790105 | down |
| GAR1 | 1.325901 | 1.048085 | 0.79047 | down |
| AREG | 1.307615 | 1.033825 | 0.790619 | down |
| HINT2 | 1.335393 | 1.05582 | 0.790644 | down |
| NIPSNAP3 | 1.397479 | 1.105037 | 0.790736 | down |
| GALNS | 1.266099 | 1.002532 | 0.791828 | down |
| HLA-F | 1.139041 | 0.901974 | 0.791871 | down |
| 44257 | 1.267452 | 1.003805 | 0.791987 | down |
| SLX1A | 1.408237 | 1.116368 | 0.792742 | down |
| RRP15 | 1.218999 | 0.966472 | 0.79284 | down |
| S100A10 | 1.34816 | 1.068925 | 0.792877 | down |
| DHRS3 | 1.280624 | 1.015564 | 0.793023 | down |
| GGT6 | 1.289555 | 1.022725 | 0.793084 | down |
| SLC35A2 | 1.19402 | 0.947226 | 0.793309 | down |
| PTDSS1 | 1.281396 | 1.016985 | 0.793654 | down |
| ADGRF4 | 1.22978 | 0.976232 | 0.793826 | down |
| CLIC3 | 1.383549 | 1.098338 | 0.793856 | down |
| FHAD1 | 1.458491 | 1.158024 | 0.793988 | down |
| CSGALNA( | 1.332078 | 1.057837 | 0.794125 | down |
| PDIA5 | 1.20579 | 0.957656 | 0.794215 | down |
| PPTC7 | 1.3413 | 1.065384 | 0.794292 | down |
| GALNT14 | 1.30521 | 1.036995 | 0.794504 | down |
| AK4 | 1.296241 | 1.030499 | 0.79499 | down |
| WFS1 | 1.269956 | 1.010165 | 0.795433 | down |
| TMEM106I | 1.231252 | 0.979425 | 0.795471 | down |
| MED9 | 1.26407 | 1.005688 | 0.795595 | down |
| NSDHL | 1.24456 | 0.990248 | 0.795661 | down |
| GRIA1 | 1.379641 | 1.097943 | 0.795818 | down |
| GJB3 | 1.528261 | 1.216255 | 0.795842 | down |
| MLEC | 1.331823 | 1.060063 | 0.795948 | down |
| CMAS | 1.303072 | 1.037432 | 0.796143 | down |
| TSPAN1 | 1.186084 | 0.944366 | 0.796205 | down |
| LMBRD1 | 1.331434 | 1.060696 | 0.796657 | down |

| | | | | |
|---|---|---|---|---|
| ATP11A | 1.240242 | 0.988074 | 0.796679 | down |
| GNA15 | 1.289578 | 1.027652 | 0.796891 | down |
| SLC44A1 | 1.305015 | 1.039974 | 0.796906 | down |
| MCTP2 | 1.328804 | 1.058955 | 0.796923 | down |
| ACSL5 | 1.294645 | 1.03195 | 0.797091 | down |
| CPT1A | 1.222272 | 0.97447 | 0.797261 | down |
| SDF4 | 1.322779 | 1.054624 | 0.797279 | down |
| DNPEP | 1.27367 | 1.016113 | 0.797783 | down |
| GNG12 | 1.165114 | 0.92972 | 0.797965 | down |
| TRIM36 | 1.387697 | 1.107377 | 0.797996 | down |
| STX8 | 1.237182 | 0.987508 | 0.798191 | down |
| DDRGK1 | 1.314114 | 1.049478 | 0.79862 | down |
| SLC44A2 | 1.266073 | 1.011173 | 0.798669 | down |
| CTSL | 1.215863 | 0.971676 | 0.799166 | down |
| PLCD3 | 1.342358 | 1.073619 | 0.799801 | down |
| SEC11A | 1.261008 | 1.008571 | 0.799813 | down |
| ARSB | 1.254476 | 1.003484 | 0.799922 | down |
| MFSD10 | 1.253561 | 1.002927 | 0.800062 | down |
| CDKN2A | 1.383531 | 1.106967 | 0.800103 | down |
| RPUSD4 | 1.206923 | 0.965856 | 0.800263 | down |
| PHYH | 1.263491 | 1.011189 | 0.800314 | down |
| BCL2L1 | 1.218985 | 0.975572 | 0.800314 | down |
| DNAJC19 | 1.200076 | 0.960552 | 0.800409 | down |
| OAT | 1.2149 | 0.972552 | 0.80052 | down |
| NOX5 | 1.506192 | 1.205856 | 0.800599 | down |
| TOR2A | 1.092334 | 0.874892 | 0.800939 | down |
| PCK2 | 1.232693 | 0.987506 | 0.801096 | down |
| NCOA4 | 1.268176 | 1.016173 | 0.801287 | down |
| TUBGCP4 | 1.44373 | 1.156896 | 0.801324 | down |
| EPHB4 | 1.201813 | 0.963145 | 0.80141 | down |
| ZDHHC7 | 1.240062 | 0.994493 | 0.80197 | down |
| IL6ST | 1.473973 | 1.182191 | 0.802044 | down |
| SCYL3 | 1.199351 | 0.962002 | 0.802102 | down |
| SPARC | 1.291445 | 1.035996 | 0.802199 | down |
| SCAMP4 | 1.299733 | 1.042827 | 0.802339 | down |
| MELTF | 1.347389 | 1.081957 | 0.803003 | down |
| TAX1BP1 | 1.255242 | 1.008172 | 0.80317 | down |
| CXCL16 | 1.547461 | 1.243603 | 0.803641 | down |
| ZDHHC13 | 1.195459 | 0.960724 | 0.803644 | down |
| SLC30A7 | 1.209361 | 0.972236 | 0.803925 | down |
| RAB22A | 1.150588 | 0.925206 | 0.804116 | down |
| TMEM70 | 1.16588 | 0.937996 | 0.804539 | down |
| GINM1 | 1.2619 | 1.015957 | 0.805101 | down |
| CYBC1 | 1.368118 | 1.101936 | 0.805439 | down |
| TMTC2 | 1.340476 | 1.079839 | 0.805564 | down |
| EDNRA | 1.519494 | 1.224208 | 0.805668 | down |
| MRPL51 | 1.169801 | 0.942553 | 0.805738 | down |
| PPP1CC | 1.041333 | 0.839259 | 0.805947 | down |
| FIS1 | 1.291833 | 1.041691 | 0.806367 | down |
| MRC1 | 1.341405 | 1.081688 | 0.806384 | down |
| ARFRP1 | 1.197957 | 0.966076 | 0.806436 | down |
| RFX7 | 1.337041 | 1.078343 | 0.806514 | down |
| SYTL1 | 1.276733 | 1.030507 | 0.807143 | down |
| AGPAT5 | 1.259485 | 1.016845 | 0.80735 | down |
| SIGMAR1 | 1.430115 | 1.154928 | 0.807577 | down |
| SLC1A3 | 1.355657 | 1.094988 | 0.807717 | down |
| KRT6B | 1.255024 | 1.013787 | 0.807783 | down |
| SIDT2 | 1.221174 | 0.986444 | 0.807784 | down |

| | | | | |
|---|---|---|---|---|
| RP2 | 1.27376 | 1.028985 | 0.807833 | down |
| TUBB8 | 1.687245 | 1.36347 | 0.808104 | down |
| DPF2 | 1.19589 | 0.966821 | 0.808453 | down |
| PNPLA6 | 1.283455 | 1.037667 | 0.808495 | down |
| OMA1 | 1.077286 | 0.871217 | 0.808715 | down |
| GBA | 1.382011 | 1.117918 | 0.808907 | down |
| CTSE | 1.25329 | 1.013857 | 0.808957 | down |
| ANKRA2 | 1.644609 | 1.330533 | 0.809027 | down |
| CANX | 1.211737 | 0.980567 | 0.809224 | down |
| ITIH2 | 1.171579 | 0.948279 | 0.809402 | down |
| AGPAT3 | 1.199028 | 0.970619 | 0.809505 | down |
| VKORC1L1 | 1.181388 | 0.956768 | 0.809867 | down |
| NDUFAB1 | 1.261326 | 1.021587 | 0.809931 | down |
| ARF6 | 1.20607 | 0.977048 | 0.810109 | down |
| SH3BP5 | 0.970169 | 0.786137 | 0.810309 | down |
| RAB15 | 1.325011 | 1.073731 | 0.810357 | down |
| LRRC49 | 1.380676 | 1.118879 | 0.810385 | down |
| NEURL4 | 1.118265 | 0.90684 | 0.810935 | down |
| COX7B | 1.165418 | 0.945196 | 0.811036 | down |
| ITM2C | 1.310083 | 1.062552 | 0.811057 | down |
| CLK4 | 1.503194 | 1.219266 | 0.811117 | down |
| ATP2B4 | 1.272687 | 1.032412 | 0.811206 | down |
| 44260 | 1.158326 | 0.93966 | 0.811223 | down |
| SLC25A3 | 1.25584 | 1.019004 | 0.811412 | down |
| TIMM22 | 1.24222 | 1.008045 | 0.811487 | down |
| ZFP28 | 1.319206 | 1.070702 | 0.811626 | down |
| P4HA2 | 1.375499 | 1.116495 | 0.811702 | down |
| NDUFB7 | 1.357646 | 1.102004 | 0.811702 | down |
| ERLIN2 | 1.171453 | 0.950967 | 0.811784 | down |
| ACP5 | 1.200585 | 0.974657 | 0.811818 | down |
| HBB | 1.134703 | 0.921456 | 0.812067 | down |
| TRIQK | 1.053799 | 0.855911 | 0.812215 | down |
| MGAT1 | 1.261218 | 1.024579 | 0.812372 | down |
| ABHD17A | 1.262745 | 1.026409 | 0.812839 | down |
| CRELD1 | 1.236323 | 1.005632 | 0.813406 | down |
| ERLEC1 | 1.296037 | 1.054475 | 0.813615 | down |
| HMOX1 | 1.230771 | 1.001683 | 0.813866 | down |
| GALNT10 | 1.216754 | 0.990407 | 0.813975 | down |
| LNPK | 1.249095 | 1.016933 | 0.814136 | down |
| TOR1AIP2 | 1.227583 | 0.999464 | 0.814172 | down |
| CWC15 | 1.312441 | 1.068715 | 0.814296 | down |
| FAM177A1 | 1.214338 | 0.988916 | 0.814366 | down |
| TST | 1.332379 | 1.08507 | 0.814385 | down |
| DHRS1 | 1.233376 | 1.004848 | 0.814713 | down |
| DEGS1 | 1.326689 | 1.080932 | 0.814759 | down |
| GRN | 1.231156 | 1.003249 | 0.814884 | down |
| RALB | 1.284608 | 1.047077 | 0.815094 | down |
| FUCA1 | 1.273578 | 1.03834 | 0.815294 | down |
| TMCC3 | 1.288632 | 1.050641 | 0.815315 | down |
| POC1B-GA | 1.358206 | 1.1082 | 0.815929 | down |
| GCNT1 | 1.267635 | 1.034373 | 0.815986 | down |
| DPY19L1 | 1.208672 | 0.986456 | 0.816149 | down |
| ATL1 | 1.180058 | 0.963326 | 0.816338 | down |
| GPT2 | 1.102971 | 0.900417 | 0.816357 | down |
| MUC1 | 1.360089 | 1.110349 | 0.816379 | down |
| RAC2 | 1.183587 | 0.96627 | 0.816391 | down |
| ARMCX6 | 1.250317 | 1.020851 | 0.816474 | down |
| LCA5 | 1.350994 | 1.10366 | 0.816924 | down |

| | | | | |
|---|---|---|---|---|
| CD59 | 1.267346 | 1.035449 | 0.817022 | down |
| ATP6AP1 | 1.226558 | 1.002161 | 0.817051 | down |
| RBM25 | 1.206261 | 0.98565 | 0.817112 | down |
| PRRG4 | 1.197091 | 0.978181 | 0.817131 | down |
| TIMM50 | 1.260635 | 1.030137 | 0.817157 | down |
| ACSF3 | 1.23972 | 1.013334 | 0.817389 | down |
| HSD17B7 | 1.183879 | 0.968368 | 0.817962 | down |
| CBR4 | 1.326936 | 1.085486 | 0.818039 | down |
| DECR1 | 1.20223 | 0.983806 | 0.818318 | down |
| RETREG3 | 1.340907 | 1.097403 | 0.818403 | down |
| LRP8 | 1.275488 | 1.043986 | 0.8185 | down |
| ICOSLG | 1.316574 | 1.077638 | 0.818516 | down |
| PPIG | 1.263853 | 1.034854 | 0.818809 | down |
| TMEM11 | 1.311011 | 1.073885 | 0.819128 | down |
| NEDD4L | 1.462251 | 1.197927 | 0.819235 | down |
| GCSH | 1.288419 | 1.055536 | 0.819249 | down |
| SLC23A2 | 1.13972 | 0.934144 | 0.819626 | down |
| SLC33A1 | 1.123993 | 0.921699 | 0.820022 | down |
| PLXNB1 | 1.245929 | 1.022491 | 0.820665 | down |
| PAFAH2 | 1.217966 | 0.999984 | 0.821028 | down |
| PINLYP | 1.478233 | 1.214295 | 0.821451 | down |
| TMEM214 | 1.192855 | 0.979876 | 0.821455 | down |
| MYO1D | 1.254393 | 1.030528 | 0.821535 | down |
| MICU1 | 1.247667 | 1.025094 | 0.821608 | down |
| PRPF4B | 1.244393 | 1.02287 | 0.821983 | down |
| TMEM138 | 1.201909 | 0.98811 | 0.822117 | down |
| STARD3NL | 1.153988 | 0.948952 | 0.822324 | down |
| TFB1M | 1.292526 | 1.062935 | 0.82237 | down |
| SLC27A4 | 1.263969 | 1.040162 | 0.822933 | down |
| P4HTM | 1.356685 | 1.116617 | 0.823048 | down |
| HSD17B12 | 1.220609 | 1.004621 | 0.823049 | down |
| KRT19 | 1.278276 | 1.052093 | 0.823056 | down |
| SPRR1B | 1.596444 | 1.314999 | 0.823705 | down |
| SURF1 | 1.287409 | 1.060709 | 0.82391 | down |
| CKMT1A | 1.264013 | 1.041479 | 0.823947 | down |
| CST3 | 1.253114 | 1.032915 | 0.824278 | down |
| EMC3 | 1.15627 | 0.953186 | 0.824363 | down |
| FAM162A | 1.236458 | 1.01943 | 0.824476 | down |
| RTN4 | 1.199035 | 0.988624 | 0.824516 | down |
| BCL2L13 | 1.218041 | 1.004338 | 0.824552 | down |
| MRPL32 | 1.420191 | 1.171083 | 0.824595 | down |
| LEMD2 | 1.268945 | 1.046368 | 0.824597 | down |
| GC | 1.235096 | 1.01855 | 0.824673 | down |
| TAP2 | 1.230135 | 1.01449 | 0.824697 | down |
| TMEM186 | 1.26498 | 1.043692 | 0.825066 | down |
| CDH1 | 1.209962 | 0.998956 | 0.82561 | down |
| HSD17B10 | 1.196694 | 0.988283 | 0.825845 | down |
| ALCAM | 1.191283 | 0.983971 | 0.825976 | down |
| REEP5 | 1.260811 | 1.041492 | 0.826049 | down |
| SRPRB | 1.146298 | 0.94751 | 0.826583 | down |
| DNAJB2 | 1.281981 | 1.059941 | 0.826799 | down |
| RALA | 1.231202 | 1.018023 | 0.826853 | down |
| EBP | 1.169169 | 0.966794 | 0.826906 | down |
| NDUFS4 | 1.141861 | 0.944433 | 0.8271 | down |
| C7orf50 | 1.194547 | 0.988268 | 0.827316 | down |
| MANEA | 1.133392 | 0.937898 | 0.827515 | down |
| NDUFB1 | 1.31065 | 1.084803 | 0.827683 | down |
| RRAGC | 1.164341 | 0.963788 | 0.827754 | down |

| | | | | |
|---|---|---|---|---|
| MSRB2 | 1.23556 | 1.022788 | 0.827793 | down |
| C4A | 1.166317 | 0.965591 | 0.827897 | down |
| ANO10 | 1.183341 | 0.979695 | 0.827906 | down |
| APOBEC3E | 1.17682 | 0.974585 | 0.828151 | down |
| HMGCL | 1.199536 | 0.993423 | 0.828172 | down |
| C6orf203 | 1.333223 | 1.104172 | 0.828198 | down |
| MYORG | 1.35684 | 1.123967 | 0.828371 | down |
| ARL10 | 1.213383 | 1.005369 | 0.828567 | down |
| EMC10 | 1.209746 | 1.002452 | 0.828646 | down |
| LRRC8D | 1.264341 | 1.048051 | 0.828931 | down |
| EPHX1 | 1.169703 | 0.969617 | 0.828943 | down |
| DGLUCY | 1.129921 | 0.936698 | 0.828994 | down |
| DMWD | 1.312725 | 1.088252 | 0.829002 | down |
| FBF1 | 1.190163 | 0.986901 | 0.829215 | down |
| KRT76 | 1.779164 | 1.47532 | 0.829221 | down |
| SCARB1 | 1.122716 | 0.931033 | 0.829268 | down |
| GPLD1 | 1.156089 | 0.958757 | 0.829311 | down |
| METTL9 | 1.212842 | 1.005973 | 0.829435 | down |
| PPP1R26 | 1.083627 | 0.898896 | 0.829525 | down |
| RHOF | 1.242751 | 1.031022 | 0.829629 | down |
| SLC25A36 | 1.165203 | 0.966704 | 0.829644 | down |
| RAP2B | 1.208032 | 1.002265 | 0.829668 | down |
| TMBIM6 | 1.207766 | 1.002237 | 0.829827 | down |
| TIMP1 | 1.313899 | 1.090384 | 0.829884 | down |
| TMED7-TI | 1.257562 | 1.043655 | 0.829904 | down |
| ZADH2 | 1.243281 | 1.031923 | 0.829999 | down |
| PROM2 | 1.272239 | 1.056129 | 0.830134 | down |
| SEC61A1 | 1.164815 | 0.96716 | 0.830313 | down |
| ECI1 | 1.31655 | 1.093303 | 0.830431 | down |
| TAP1 | 1.207133 | 1.002832 | 0.830755 | down |
| PTPN18 | 0.833689 | 0.692685 | 0.830868 | down |
| PCDH11X | 1.101646 | 0.915472 | 0.831004 | down |
| TMED4 | 1.161265 | 0.965207 | 0.831169 | down |
| SLC18A3 | 1.254059 | 1.042414 | 0.831232 | down |
| NDEL1 | 1.195674 | 0.993942 | 0.831281 | down |
| CMTM4 | 1.099942 | 0.91447 | 0.831381 | down |
| C1RL | 1.329122 | 1.105653 | 0.831867 | down |
| HIGD2A | 1.287872 | 1.071584 | 0.832058 | down |
| GDPD1 | 1.232012 | 1.025482 | 0.832364 | down |
| ADIPOR2 | 1.341249 | 1.116498 | 0.832432 | down |
| CASC4 | 1.222964 | 1.0182 | 0.832568 | down |
| LEPROT | 1.187767 | 0.988909 | 0.832578 | down |
| ABCB8 | 1.266485 | 1.055395 | 0.833326 | down |

Table S7. related to Figure 5

**iTRAQ shNAT10_vs_shControl**

| Ingenuity C | -log(p-valu | p-value | Ratio |
|---|---|---|---|
| Cholestero | 12.9 | 1.26E-13 | 0.769 |
| Cholestero | 12.9 | 1.26E-13 | 0.769 |
| Cholestero | 12.9 | 1.26E-13 | 0.769 |
| Superpathv | 9.53 | 2.95E-10 | 0.379 |
| Glucocortic | 7.88 | 1.32E-08 | 0.0757 |
| LXR/RXR / | 7.65 | 2.24E-08 | 0.146 |
| Acute Phas | 6.2 | 6.31E-07 | 0.108 |
| IL-8 Signali | 5.31 | 4.9E-06 | 0.0948 |
| Phagosom | 5.3 | 5.01E-06 | 0.107 |
| Tumor Mic | 5.2 | 6.31E-06 | 0.101 |
| Sirtuin Sigr | 5.16 | 6.92E-06 | 0.0822 |
| Glioma Inv | 5.03 | 9.33E-06 | 0.151 |
| Zymosterol | 4.97 | 1.07E-05 | 0.667 |
| Mitochondr | 4.91 | 1.23E-05 | 0.1 |
| Colorectal | 4.75 | 1.78E-05 | 0.0818 |
| Ferroptosis | 4.66 | 2.19E-05 | 0.111 |
| Oncostatin | 4.49 | 3.24E-05 | 0.186 |
| Role of Tis | 4.41 | 3.89E-05 | 0.112 |
| Endothelin- | 4.26 | 5.5E-05 | 0.089 |
| Virus Entry | 4.24 | 5.75E-05 | 0.115 |
| Neuroinflar | 4.16 | 6.92E-05 | 0.073 |
| Oxidative F | 4.04 | 9.12E-05 | 0.11 |
| Estrogen-D | 3.97 | 0.000107 | 0.127 |
| Role of MA | 3.92 | 0.00012 | 0.107 |
| Antigen Pre | 3.89 | 0.000129 | 0.179 |
| PTEN Sign | 3.83 | 0.000148 | 0.0933 |
| LPS/IL-1 M | 3.72 | 0.000191 | 0.0754 |
| Agrin Intera | 3.68 | 0.000209 | 0.129 |
| FXR/RXR / | 3.44 | 0.000363 | 0.0952 |
| Macropino | 3.4 | 0.000398 | 0.118 |
| Estrogen R | 3.35 | 0.000447 | 0.0619 |
| ErbB2-ErbI | 3.2 | 0.000631 | 0.123 |
| UVC-Induc | 3.15 | 0.000708 | 0.137 |
| Epoxysqua | 3.06 | 0.000871 | 1 |
| PI3K/AKT | 3.06 | 0.000871 | 0.0754 |
| Inhibition o | 3.04 | 0.000912 | 0.154 |
| Ephrin Rec | 3.01 | 0.000977 | 0.0746 |
| PDGF Sigr | 3.01 | 0.000977 | 0.105 |
| Sertoli Cell | 2.91 | 0.00123 | 0.0728 |
| ERK5 Sign | 2.9 | 0.001259 | 0.111 |
| IL-4 Signali | 2.8 | 0.001585 | 0.0978 |
| Caveolar-m | 2.79 | 0.001622 | 0.107 |
| Mitochondr | 2.78 | 0.00166 | 0.222 |
| Regulation | 2.75 | 0.001778 | 0.0729 |
| Endometria | 2.73 | 0.001862 | 0.117 |
| NRF2-med | 2.72 | 0.001905 | 0.0675 |
| Antiprolifer | 2.71 | 0.00195 | 0.104 |
| HER-2 Sigi | 2.67 | 0.002138 | 0.0714 |
| CCR3 Sigr | 2.65 | 0.002239 | 0.0815 |
| Thyroid Ca | 2.64 | 0.002291 | 0.101 |
| Bladder Ca | 2.63 | 0.002344 | 0.0862 |
| Axonal Gui | 2.62 | 0.002399 | 0.0534 |
| UVA-Induc | 2.61 | 0.002455 | 0.0918 |
| Fc Epsilon | 2.61 | 0.002455 | 0.0855 |
| Synaptoge | 2.61 | 0.002455 | 0.0609 |
| 4-aminobu | 2.59 | 0.00257 | 0.667 |

**iTRAQ shRNPS1_vs_shControl**

| Ingenuity C | -log(p-valu | p-value | Ratio |
|---|---|---|---|
| Cholestero | 15.5 | 3.16E-16 | 0.846 |
| Cholestero | 15.5 | 3.16E-16 | 0.846 |
| Cholestero | 15.5 | 3.16E-16 | 0.846 |
| Superpathv | 11.4 | 3.98E-12 | 0.414 |
| Glucocortic | 10.3 | 5.01E-11 | 0.0792 |
| Acute Phas | 8.31 | 4.9E-09 | 0.119 |
| IL-8 Signali | 7.93 | 1.17E-08 | 0.109 |
| Tumor Mic | 7.86 | 1.38E-08 | 0.117 |
| Glioma Inv | 7.23 | 5.89E-08 | 0.178 |
| Oncostatin | 6.84 | 1.45E-07 | 0.233 |
| Virus Entry | 6.17 | 6.76E-07 | 0.135 |
| Macropino | 6.13 | 7.41E-07 | 0.158 |
| LXR/RXR / | 6.01 | 9.77E-07 | 0.122 |
| Estrogen-D | 5.95 | 1.12E-06 | 0.152 |
| Role of Tis | 5.59 | 2.57E-06 | 0.121 |
| Colorectal | 5.46 | 3.47E-06 | 0.0818 |
| Phagosom | 5.25 | 5.62E-06 | 0.101 |
| Melanoma | 5.23 | 5.89E-06 | 0.18 |
| Ferroptosis | 5.17 | 6.76E-06 | 0.111 |
| UVC-Induc | 5.15 | 7.08E-06 | 0.176 |
| Antigen Pre | 5.15 | 7.08E-06 | 0.205 |
| Zymosterol | 5.15 | 7.08E-06 | 0.667 |
| Estrogen R | 4.9 | 1.26E-05 | 0.0668 |
| Sirtuin Sigr | 4.89 | 1.29E-05 | 0.0753 |
| Mitochondr | 4.88 | 1.32E-05 | 0.0941 |
| Neuroinflar | 4.86 | 1.38E-05 | 0.073 |
| Endothelin- | 4.83 | 1.48E-05 | 0.089 |
| Agrin Intera | 4.81 | 1.55E-05 | 0.143 |
| Ephrin Rec | 4.54 | 2.88E-05 | 0.0846 |
| Antiprolifer | 4.44 | 3.63E-05 | 0.13 |
| Thyroid Ca | 4.34 | 4.57E-05 | 0.127 |
| PTEN Sign | 4.31 | 4.9E-05 | 0.0933 |
| ErbB2-ErbI | 4.27 | 5.37E-05 | 0.138 |
| UVA-Induc | 4.23 | 5.89E-05 | 0.112 |
| Bladder Ca | 4.21 | 6.17E-05 | 0.103 |
| Cholecysto | 4.1 | 7.94E-05 | 0.101 |
| PDGF Sigr | 4.02 | 9.55E-05 | 0.116 |
| ERK5 Sign | 3.92 | 0.00012 | 0.125 |
| CNTF Sign | 3.9 | 0.000126 | 0.14 |
| PPAR Sign | 3.88 | 0.000132 | 0.103 |
| Cancer Dru | 3.85 | 0.000141 | 0.138 |
| CXCR4 Sig | 3.8 | 0.000158 | 0.0838 |
| Caveolar-m | 3.78 | 0.000166 | 0.12 |
| Synaptic Lo | 3.76 | 0.000174 | 0.093 |
| G Beta Gai | 3.76 | 0.000174 | 0.093 |
| Endometria | 3.74 | 0.000182 | 0.133 |
| Actin Nucle | 3.74 | 0.000182 | 0.108 |
| ErbB Signa | 3.7 | 0.0002 | 0.106 |
| IL-2 Signali | 3.69 | 0.000204 | 0.131 |
| HGF Signa | 3.67 | 0.000214 | 0.0909 |
| NRF2-med | 3.66 | 0.000219 | 0.0717 |
| Axonal Gui | 3.63 | 0.000234 | 0.0553 |
| IL-3 Signali | 3.61 | 0.000245 | 0.114 |
| HER-2 Sigi | 3.6 | 0.000251 | 0.0765 |
| Thrombopo | 3.59 | 0.000257 | 0.127 |
| STAT3 Pat | 3.58 | 0.000263 | 0.0889 |

| | | | |
|---|---|---|---|
| Neuroprote | 2.58 | 0.00263 | 0.0847 |
| Cholecysto | 2.55 | 0.002818 | 0.084 |
| Apelin End | 2.55 | 0.002818 | 0.0791 |
| Melanoma | 2.47 | 0.003388 | 0.12 |
| Coagulatio | 2.46 | 0.003467 | 0.143 |
| Apoptosis S | 2.43 | 0.003715 | 0.0865 |
| α-Adrenerg | 2.38 | 0.004169 | 0.0849 |
| SPINK1 Ge | 2.38 | 0.004169 | 0.101 |
| PPAR Sign | 2.35 | 0.004467 | 0.0841 |
| Stearate Bi | 2.34 | 0.004571 | 0.113 |
| IL-6 Signali | 2.33 | 0.004677 | 0.0781 |
| G Beta Gar | 2.3 | 0.005012 | 0.0775 |
| Branched-c | 2.3 | 0.005012 | 0.5 |
| Ceramide S | 2.29 | 0.005129 | 0.0889 |
| Synaptic Lo | 2.28 | 0.005248 | 0.067 |
| Ephrin B Si | 2.27 | 0.00537 | 0.0972 |
| fMLP Signa | 2.26 | 0.005495 | 0.0763 |
| Gα12/13 S | 2.21 | 0.006166 | 0.0752 |
| CNTF Sign | 2.18 | 0.006607 | 0.105 |
| ErbB Signa | 2.18 | 0.006607 | 0.0851 |
| Cancer Dru | 2.15 | 0.007079 | 0.103 |
| Neuregulin | 2.1 | 0.007943 | 0.0769 |
| Gαi Signali | 2.1 | 0.007943 | 0.0725 |
| Citrulline-N | 2.09 | 0.008128 | 0.4 |
| Glutamate | 2.09 | 0.008128 | 0.4 |
| Iron homeo | 2.08 | 0.008318 | 0.0719 |
| Systemic L | 2.07 | 0.008511 | 0.0578 |
| IL-3 Signali | 2.06 | 0.00871 | 0.0886 |
| Superpathw | 2.05 | 0.008913 | 0.2 |
| IL-2 Signali | 2.04 | 0.00912 | 0.0984 |
| MIF Regula | 2.03 | 0.009333 | 0.114 |
| G Protein S | 2.03 | 0.009333 | 0.114 |
| Estrogen B | 2.03 | 0.009333 | 0.114 |
| Renal Cell | 2.03 | 0.009333 | 0.0875 |
| Chemokine | 2.03 | 0.009333 | 0.0875 |

| | | | |
|---|---|---|---|
| Renal Cell | 3.57 | 0.000269 | 0.113 |
| Chemokine | 3.57 | 0.000269 | 0.113 |
| Neuregulin | 3.53 | 0.000295 | 0.094 |
| Natural Kill | 3.53 | 0.000295 | 0.0754 |
| PI3K/AKT S | 3.53 | 0.000295 | 0.0754 |
| VEGF Sign | 3.51 | 0.000309 | 0.101 |
| PAK Signa | 3.5 | 0.000316 | 0.0932 |
| JAK/Stat S | 3.49 | 0.000324 | 0.11 |
| Apelin End | 3.46 | 0.000347 | 0.0863 |
| PEDF Sign | 3.41 | 0.000389 | 0.107 |
| LPS-stimul | 3.37 | 0.000427 | 0.106 |
| Sertoli Cell | 3.37 | 0.000427 | 0.0728 |
| ErbB4 Sign | 3.36 | 0.000437 | 0.118 |
| PI3K Signa | 3.35 | 0.000447 | 0.0839 |
| HIF1α Sign | 3.32 | 0.000479 | 0.0721 |
| SPINK1 Ge | 3.32 | 0.000479 | 0.116 |
| Glioma Sig | 3.32 | 0.000479 | 0.0887 |
| Stearate Bi | 3.32 | 0.000479 | 0.132 |
| Thrombin S | 3.28 | 0.000525 | 0.0714 |
| GM-CSF S | 3.28 | 0.000525 | 0.114 |
| α-Adrenerg | 3.28 | 0.000525 | 0.0943 |
| FXR/RXR A | 3.26 | 0.00055 | 0.0873 |
| Regulation | 3.22 | 0.000603 | 0.101 |
| IL-6 Signali | 3.2 | 0.000631 | 0.0859 |
| Ceramide S | 3.19 | 0.000646 | 0.1 |
| Oxidative P | 3.18 | 0.000661 | 0.0917 |
| P2Y Purige | 3.17 | 0.000676 | 0.0853 |
| Germ Cell- | 3.17 | 0.000676 | 0.076 |
| Glioblastom | 3.17 | 0.000676 | 0.076 |
| PPARα/RX | 3.16 | 0.000692 | 0.0725 |
| Epoxysqua | 3.15 | 0.000708 | 1 |
| Synaptoge | 3.13 | 0.000741 | 0.0609 |
| fMLP Signa | 3.12 | 0.000759 | 0.084 |
| IL-4 Signali | 3.12 | 0.000759 | 0.0978 |
| Role of MA | 3.09 | 0.000813 | 0.0893 |
| Gα12/13 S | 3.06 | 0.000871 | 0.0827 |
| Non-Small | 3.05 | 0.000891 | 0.0957 |
| GDNF Fam | 3.04 | 0.000912 | 0.105 |
| Neurotroph | 3.04 | 0.000912 | 0.105 |
| Hepatic Fib | 3.04 | 0.000912 | 0.0549 |
| CCR3 Sign | 3.01 | 0.000977 | 0.0815 |
| IL-15 Signa | 3 | 0.001 | 0.104 |
| TGF-β Sign | 2.98 | 0.001047 | 0.0938 |
| NF-κB Acti | 2.96 | 0.001096 | 0.103 |
| Ovarian Ca | 2.96 | 0.001096 | 0.0759 |
| Epithelial A | 2.96 | 0.001096 | 0.0759 |
| MSP-RON | 2.94 | 0.001148 | 0.0855 |
| Rac Signal | 2.93 | 0.001175 | 0.0797 |
| FLT3 Signa | 2.89 | 0.001288 | 0.1 |
| Role of MA | 2.85 | 0.001413 | 0.0988 |
| Xenobiotic | 2.8 | 0.001585 | 0.0769 |
| Cardiac Hy | 2.8 | 0.001585 | 0.062 |
| GNRH Sign | 2.79 | 0.001622 | 0.0691 |
| Endocanna | 2.78 | 0.00166 | 0.0813 |
| mTOR Sign | 2.77 | 0.001698 | 0.066 |
| IGF-1 Sign | 2.74 | 0.00182 | 0.0865 |
| Apoptosis S | 2.74 | 0.00182 | 0.0865 |
| ERK/MAPK | 2.73 | 0.001862 | 0.0654 |
| FcγRIIB Sig | 2.72 | 0.001905 | 0.0941 |

| | | | |
|---|---|---|---|
| Regulation | 2.71 | 0.00195 | 0.0677 |
| Reelin Sigr | 2.7 | 0.001995 | 0.0794 |
| Prolactin S | 2.69 | 0.002042 | 0.093 |
| 4-aminobut | 2.68 | 0.002089 | 0.667 |
| 14-3-3-mec | 2.67 | 0.002138 | 0.0787 |
| Synaptic Lo | 2.67 | 0.002138 | 0.067 |
| BMP signa | 2.66 | 0.002188 | 0.092 |
| Chronic My | 2.65 | 0.002239 | 0.0841 |
| Telomerase | 2.65 | 0.002239 | 0.0841 |
| Role of JAK | 2.63 | 0.002344 | 0.101 |
| Paxillin Sig | 2.63 | 0.002344 | 0.0833 |
| Actin Cytos | 2.61 | 0.002455 | 0.0612 |
| p70S6K Sig | 2.55 | 0.002818 | 0.0758 |
| Compleme | 2.55 | 0.002818 | 0.135 |
| Acute Myel | 2.53 | 0.002951 | 0.0879 |
| CDK5 Sign | 2.52 | 0.00302 | 0.0804 |
| Gαi Signali | 2.41 | 0.00389 | 0.0725 |
| MSP-RON | 2.41 | 0.00389 | 0.0725 |
| Fc Epsilon | 2.39 | 0.004074 | 0.0769 |
| FAK Signal | 2.39 | 0.004074 | 0.0769 |
| Insulin Rec | 2.37 | 0.004266 | 0.0714 |
| Neuroprote | 2.37 | 0.004266 | 0.0763 |
| Angiopoieti | 2.36 | 0.004365 | 0.0909 |
| IL-17 Signa | 2.35 | 0.004467 | 0.0642 |
| Integrin Sig | 2.32 | 0.004786 | 0.061 |
| Renin-Angi | 2.32 | 0.004786 | 0.075 |
| Xenobiotic | 2.32 | 0.004786 | 0.0554 |
| HMGB1 Sig | 2.28 | 0.005248 | 0.0659 |
| Production | 2.28 | 0.005248 | 0.0628 |
| B Cell Rec | 2.23 | 0.005888 | 0.0619 |
| Mouse Em | 2.18 | 0.006607 | 0.0769 |
| Citrulline-N | 2.17 | 0.006761 | 0.4 |
| Glutamate | 2.17 | 0.006761 | 0.4 |
| Superpathw | 2.17 | 0.006761 | 0.2 |
| Gap Juncti | 2.16 | 0.006918 | 0.0606 |
| VEGF Fam | 2.15 | 0.007079 | 0.0833 |
| Systemic L | 2.12 | 0.007586 | 0.0542 |
| Relaxin Sig | 2.1 | 0.007943 | 0.0654 |
| Erythropoie | 2.1 | 0.007943 | 0.0621 |
| Glutamate | 2.09 | 0.008128 | 0.0909 |
| Molecular I | 2.07 | 0.008511 | 0.0472 |
